# Human Cytomegalovirus modifies placental small extracellular vesicle secretion and composition towards a proviral phenotype to enhance infection of fetal recipient cells

**DOI:** 10.1101/2021.11.18.468660

**Authors:** Mathilde Bergamelli, Hélène Martin, Yann Aubert, Jean-Michel Mansuy, Marlène Marcellin, Odile Burlet-Schiltz, Ilse Hurbain, Graça Raposo, Jacques Izopet, Thierry Fournier, Alexandra Benchoua, Mélinda Bénard, Marion Groussolles, Géraldine Cartron, Yann Tanguy le Gac, Nathalie Moinard, Gisela D’Angelo, Cécile E. Malnou

## Abstract

Although placental small extracellular vesicles (sEVs) are extensively studied in the context of pregnancy, little is known about their role during human cytomegalovirus (hCMV) congenital infection, especially at the beginning of pregnancy. In this study, we examined the consequences of hCMV infection on sEVs production, composition and function using an immortalized human cytotrophoblast cell line derived from first trimester placenta. By combining complementary approaches of biochemistry, electron microscopy and quantitative proteomic analysis, we showed that hCMV infection increases the yield of sEVs produced by cytotrophoblasts and modifies their protein content towards a proviral phenotype. We further demonstrate that sEVs secreted by hCMV-infected cytotrophoblasts potentiate infection in naive recipient cells of fetal origin, including human neural stem cells. Importantly, these functional consequences are also observed with sEVs prepared from either an *ex vivo* model of infected histocultures from early placenta or from the amniotic fluid of patients naturally infected by hCMV at the beginning of pregnancy. Based on these findings, we propose that placental sEVs could be key actors favoring viral dissemination to the fetal brain during hCMV congenital infection.

**Significance Statement:** Human cytomegalovirus (hCMV) infection is a major issue during pregnancy, affecting 1% of births in western countries. Despite extensive research, the pathophysiology of this congenital infection remains unclear. Recently, increasing evidence point to the key role of placental small extracellular vesicles (sEVs) in materno-fetal communication during pregnancy. Here, we examined the impact of hCMV infection on the protein composition and function of placental sEVs. We observe that hCMV infection leads to major changes in placental sEV protein content. Functional studies show the ability of sEVs produced by placental infected cells to facilitate further infection of naive recipient fetal cells, notably human neural stem cells. Our study demonstrates that placental sEVs are key players of hCMV pathophysiology during congenital infection.

## Introduction

Human cytomegalovirus (hCMV) belongs to the *Herpesviridae* family and its prevalence is of 50 to 90 % in global human population. Most of the hCMV infections occurring among immunocompetent adults induce an asymptomatic acute replication phase followed by a lifelong persistent latent state. However, hCMV primo-infection, reinfection and/or reactivation may severely compromise the health of immunocompromised people, and is a major issue during pregnancy (1, 2). Congenital infection by hCMV affects 1% of live births in western countries, making hCMV the most frequently transmitted virus *in utero* (2, 3), causing placental and fetal impairments of variable severity. The most severe consequences are observed when transmission occurs during peri-conceptional period or first trimester (4). Infection of the placenta itself allows the virus to actively replicate and enable its further access to the fetus (3, 5, 6). The infected placenta can develop a pathology that may lead to miscarriage, premature delivery, *intra* uterine growth retardation or even fetal death (2, 4). On the other side, the infection of the fetus causes various visceral and central nervous system damage, congenital hCMV infection being the most common cause of brain malformations and deafness of infectious origin (2, 7, 8). Despite the extensive research conducted so far, the pathophysiology of hCMV infection remains unclear, especially concerning potential factors which may explain the wide variety of clinical manifestations and their severity (2).

In the course of hCMV congenital infection, the placenta is a central key organ, i.e. the target of viral replication allowing further vertical transmission towards the fetus. Amongst the numerous placental functions, a recently described and extensively studied mode of communication between both maternal and fetal sides consists in the production of placental extracellular vesicles (EVs) (9, 10). EVs are membranous vesicles secreted by cells in both physiological and pathological situations, which main subtypes can be distinguished depending on their biogenesis and size into small EVs (sEVs) and large EVs (lEVs). They are specifically composed of various molecules such as proteins, lipids and coding and non-coding RNAs (11, 12). Once released into the extracellular space, EVs can be internalized by other cells, in their immediate environment or within long distances, wherein they exert regulatory roles (13). For example, placental EVs can be uptaken by Natural Killer cells (14, 15) or by primary placental fibroblasts (16). Although the understanding of the biological relevance of placental EVs *in vivo* remains limited, recent findings highlight their roles in cell-cell communication underlying the feto-placenta-maternal dialogue during pregnancy (17, 18). Interestingly, placental EV content is altered upon gestational diseases such as preeclampsia, preterm birth or gestational diabetes mellitus, and recent literature points towards a putative role of dysregulated placental EVs during pathological pregnancies (19–24).

Although placental sEVs are extensively studied in the context of pregnancy diseases, little is currently known about their role during hCMV congenital infection, especially at the very beginning of pregnancy where the most severe sequelae take their origin. Our recent data described a dysregulation of the surface expression of placental sEV markers upon hCMV infection in an *ex vivo* model of first trimester placental histocultures, suggesting a putative role for viral dissemination (25). In the present study, we used immortalized cytotrophoblasts derived from first trimester placenta, named HIPEC (26), to comprehensively examine the consequences of hCMV infection on sEV production, content and function in recipient cells. We show that hCMV increases the sEV production by HIPECs and alters their protein content towards a proviral phenotype. Finally, we observe that sEVs secreted by hCMV-infected HIPECs potentiate infection in naive recipient cells, including human neural stem cells (NSCs). Importantly, this enhancement of hCMV infection is also observed with sEVs from *ex vivo* early placental histocultures and with sEVs purified from amniotic fluid of patient infected by hCMV during first trimester of pregnancy. We propose that placental sEVs could be key players favoring viral dissemination towards fetal brain during hCMV congenital infection.

## Results

### HIPEC infection by hCMV leads to an increase of sEV secretion without modifying their general features

To study the consequences of hCMV infection on placental sEV secretion, composition and function in early pregnancy, HIPEC conditioned media were collected between 48-72 h post-infection before sEV preparation, times at which 50-80 % of cells were infected (Supplementary Figure S1B). To exclude any viral contamination of sEVs, an infectivity test was systematically carried out for each preparation. No infection was detected in cells incubated with sEVs alone (Supplementary Figure S1C). Moreover, no structure evoking viral particles was observed by transmission electron microscopy (TEM) on sEVs from infected HIPECs (Supplementary Figure S2, and data not shown).

Despite a decrease of cell number upon hCMV infection compared to non-infected (NI) HIPECs (Supplementary Figure S1D), the quantification of sEVs isolated per cell showed a significant higher yield upon infection, with an increase of around 40 % (Figure 1A). No difference in either the mean size or the mode size of the sEVs was observed upon infection when analyzed by nanoparticle tracking analysis (NTA; Figure 1B) and sEVs from both NI or infected HIPECs exhibited the same typical structure and shape as evidenced by TEM (Figure 1C and Supplementary Figure S2).

**Figure 1:**
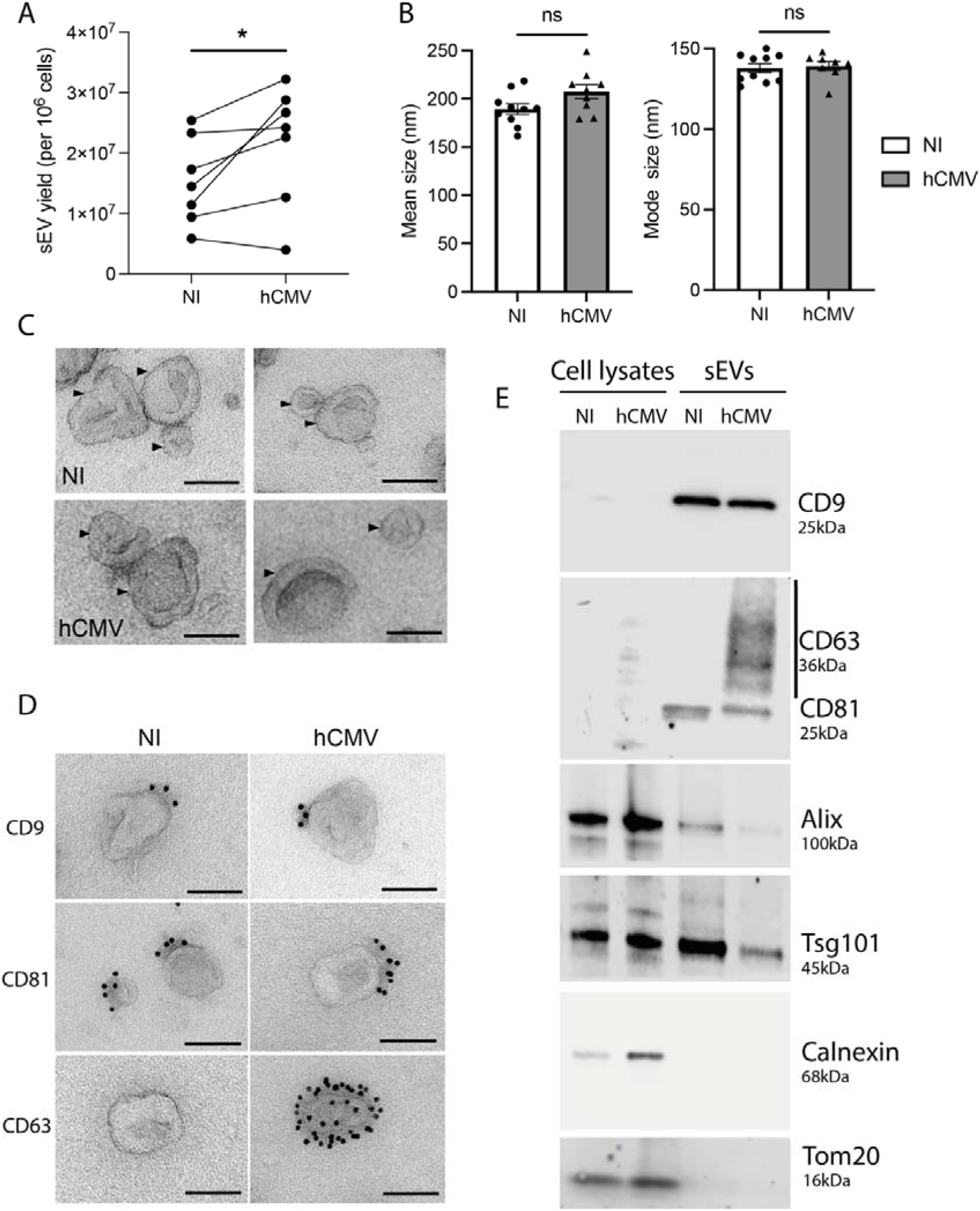
Characterization of sEVs from NI and hCMV infected HIPECs. A) Yield of sEV recovered from non-infected (NI) or infected (hCMV) HIPECs. *, *p* = 0.0464 by paired *t*-test for 7 independent experiments. B) Comparison of mean size (left histogram) and mode size (right histogram) between sEVs from non-infected (NI) or infected (hCMV) HIPECs. Histograms show the mean ± SEM of three independent experiments. ns: non-significant by Mann Whitney test. C) Electron microscopy images of sEVs (indicated by an arrow) prepared from non-infected (NI) or infected (hCMV) HIPECs. Magnification = 26000 X. Scale bar = 100 nm. Images are representative of at least three independent experiments. D) TEM observation of sEV - isolated from non-infected (NI) or infected (hCMV) HIPECs - which were immunogold-labelled for CD9, CD81 or CD63, and revealed with Protein A-gold particle of 10 nm diameter. Scale bar = 100 nm. Magnification = 26000 X. In CD63 IEM, only one example of positive vesicle, representing around 1-5 % of sEVs isolated upon infection, is shown, the other being negative (see Supplementary Figure 3 for wide field image). E) Western-blot realized on either whole cell lysates (left wells) or purified sEVs (right wells), from non-infected (NI) or infected (hCMV) HIPECs. Proteins of interest and their corresponding molecular weight are indicated on the right of the Figure, with a smear for CD63 due to the non-reducing conditions of the western blot, which preserve its rich glycosylated pattern.

We next assessed the impact of hCMV infection on the expression of sEV canonical markers. By combining western-blotting, immunolabeling electron microscopy (IEM) and multiplex bead-based flow cytometry, we observed that sEV preparations expressed specific vesicular markers including CD9, CD81, Alix and Tsg101, but no endoplasmic reticulum or mitochondrial markers (Figure 1D, E and supplementary Figure S3), attesting the purity of the sEV preparations (27). No drastic differences were observed in their expression levels upon infection as compared to NI cells, except for CD63 protein which was not detected in whole cell lysates, but enriched in sEVs from hCMV-infected HIPECs (Figure 1E). By IEM, at the level of individual vesicles, we noticed that this increase was correlated to the presence of a small proportion of sEVs highly positive for CD63 (between 1-5 %), while the others remained negative (Figure 1D and supplementary Figure S4). By multiplex bead-based flow cytometry assay, no significant increase in CD63 expression was detected between sEVs from NI or infected HIPECs upon infection, certainly due to the low proportion of positive vesicles (supplementary Figure S3). Hence, hCMV infection increased HIPEC-sEV production but did not globally impact on their global features, except for CD63 that is detected in a subpopulation of vesicles upon infection.

### sEVs secreted by infected HIPECs harbor a proviral protein cargo

To extend the molecular characterization of sEVs secreted by hCMV-infected HIPECs, a comprehensive proteomic approach that allows for a deeper analysis of sEV composition was carried out. The analysis by mass spectrometry-based quantitative proteomics of equivalent amounts of sEVs from NI or hCMV-infected HIPECs led to the identification of 3079 proteins across all samples (3048 human and 31 viral; Supplementary Table S1). Among the 3048 human proteins identified, the gProfiler2 R package was able to interrogate 2936 proteins, for which the term “extracellular exosome” (Gene ontology GO:0070062) appeared as the most significantly enriched (false discovery rate (FDR)□ = □1.087859e-^259^, R package gProfiler2), with 962 of them (32.8 %) associated with the “extracellular exosome” GO term. Conversely, these 962 proteins constituted 44.1 % of the proteins which define the GO term, and the 94 of the top 100 most frequently identified exosomal proteins, as defined by the Exocarta database (http://exocarta.org/exosome_markers_new), were detected in the sEV preparations.

Proteomics data indicated that 37 proteins, including 31 viral proteins, were significantly over-represented and 15 human proteins under-represented in sEVs from hCMV-infected HIPECs (Figure 2A). Interestingly, the Thy-1 cellular protein, known to play an important role to facilitate hCMV entry into cells *via* macropinocytosis (28, 29), was significantly enriched in sEVs upon infection (Supplementary Figure S5A). Moreover, by using Ingenuity Pathway Analysis (IPA) tool, several biological functions were found to be significantly over- or under-represented in sEVs upon infection (Figure 2B). The most modulated pathway identified was “autophagy”, with several actors showing altered expression in sEVs from infected HIPECs, leading to an overall pattern of autophagy activation (Figure 2C). Since the EV content primarily reflects the composition of the cells they are derived from, this may also reflect the activation of the autophagy pathway induced during early hCMV infection of host cells (30–32). On the other hand, two pathways related to mitochondrial functions (*i.e*., “consumption of oxygen” and “ATP synthesis”) were found to be lower in sEVs from hCMV-infected HIPECs than in NI HIPECs (Figure 2B). This may also be the consequence of hCMV-induced mitochondrial dysfunctions (33–36), likely *via* interference with the antiviral Viperin protein, which leads to decreased cellular ATP levels (37).

**Figure 2:**
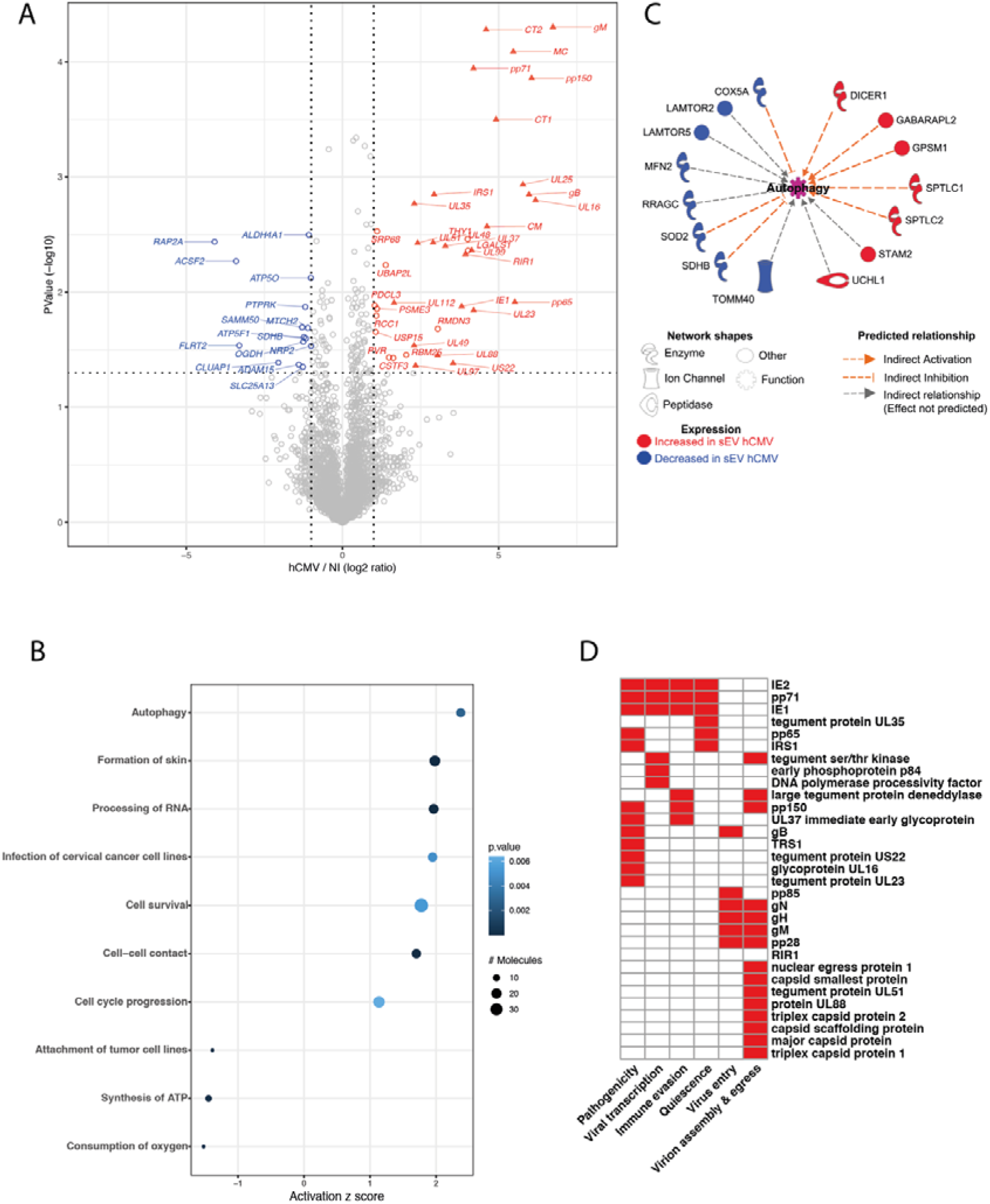
Proteomic analysis of sEV composition upon infection of HIPECs by hCMV. A) Volcano-plot representing differences in normalized mean protein abundance in sEVs hCMV *versus* sEVs NI. Human and viral proteins exhibiting significant differences between the two conditions are represented by circles and triangles, respectively (Student T-test *p*-value <=0.05 and log2 ratio >=1 or <=-1). Red: over-represented proteins; Blue: under-represented proteins. B) Dot plot representation of the top diseases and biological functions associated with human proteins exhibiting an absolute normalized mean abundance log2 ratio greater than 1 or lower than −1, in sEVs hCMV *versus* sEVs NI. Top diseases and biological functions associated with changes in the protein content of sEVs upon hCMV-infection were identified using QIAGEN Ingenuity Pathway Analysis (IPA). Size of the circles depends on the number of the proteins identified in the corresponding pathway; level of blue intensity depends on the *p*-value. C) Human proteins associated with the predicted increased activation state of autophagy pathway as determined by IPA. D) Heatmap representation of the biological functions associated with hCMV viral proteins expressed in sEVs hCMV.

Importantly, the proteomic data also revealed that 31 proteins were of viral origin (Figure 2D and Supplementary Table S2). They are involved in different aspects of hCMV infection, from viral entry to egress, quiescence, as well as pathogenicity and immune evasion, and are mainly immediate early or late proteins (Supplementary Figure S5B). Interestingly, although sEV preparations were devoid of viral particles as assessed by infectivity assays and TEM (supplementary Figures S1C and S2), some of the viral proteins in sEV from hCMV-infected HIPECs are structural proteins (Figure 2D and Supplementary Table S2), i.e. the envelope proteins gB, gH and gM, as it has already been described in other studies (38–40). Most of the other proteins identified were capsid and tegument proteins that are delivered to host cells upon infection, or proteins immediately expressed after virus entry such as IE1 and IE2, which play a role in early transcriptions (41–43). We also identified pp65, which participates to the transactivation of viral major immediate early genes (44, 45), as well as pp71, which stimulates viral immediate early transcription and inhibits the host innate response by targeting STING (46, 47). Finally, IRS1 and TRS1, which inhibit the establishment of an antiviral state in infected cells, in particular by antagonizing the autophagy pathway induced upon hCMV infection (30, 31), were also detected. Altogether, analysis of the proteomic data indicates that sEVs secreted from infected HIPECs carry a protein cargo with proviral properties. The incorporation of these viral proteins into sEVs could enhance the viral spread by providing the recipient cells with elements that may facilitate the early steps of a further hCMV infection.

### sEVs are efficiently uptaken by recipient MCR5 cells

We next examined whether MRC5 cells could uptake sEVs. PKH67-stained sEVs were incubated with MRC5 cells for different times and the internalization of labeled-sEVs was evaluated both by confocal fluorescence (Figures 3A and supplementary S6) and flow cytometry (Figure 3B, C). As early as 2 h post-incubation, some MRC5 cells already showed numerous cytoplasmic puncta for both sEVs produced by NI and infected HIPECs (Supplementary Figure S6). At 16 h, many cells showed bright green puncta in their cytoplasm (Figure 3A), with a percentage of positive cells evaluated to 13% by flow cytometry, which was not altered up to 24h (Figure 3B-C). Moreover, these puncta were visible in the same confocal plan as actin (revealed by phalloidin staining), corroborating the intracytoplasmic localization of PKH67 fluorescence, and excluding therefore a possible binding of sEVs to the cell surface (Figure 3A, Supplementary Figures S6B and C and Supplementary Video S1). However, in comparison to the fluorescence data (Figure 3A), this percentage may be somewhat underestimated, as subtraction of cell autofluorescence background may mask a significant amount of uptaken sEVs, which fluoresce poorly, given the small size of the vesicles and thus, the small number of PKH67 molecules incorporated. Altogether, these data indicate that the sEVs secreted by HIPECs are largely uptaken by recipient cells, and also suggest that the internalized sEVs may thereafter exert a biological function.

**Figure 3:**
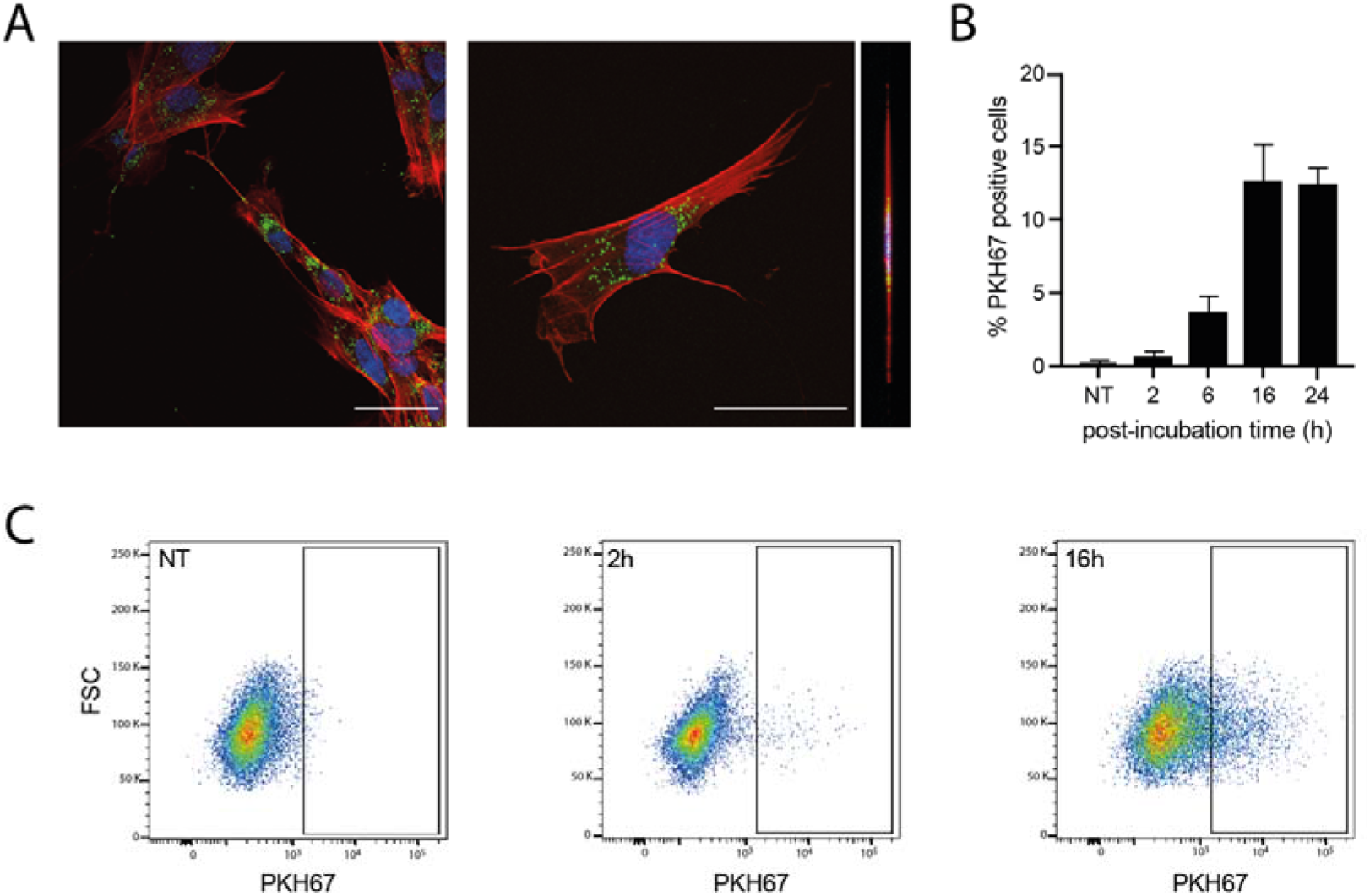
Internalization of sEVs isolated from non-infected HIPECs in fetal MRC5 cells. A) Confocal images of fluorescence microscopy carried out on MRC5 cells after 16h incubation with PKH67-labelled sEVs. Blue: DAPI; Red: Phalloidin; Green: PKH67. Scale bar: 100 μm. Magnification = 63 X. The right image corresponds to the orthogonal projection of the cell z-axis. B) Histogram representing the percentage of PKH67 positive cells along time, upon incubation of MRC5 cells with sEVs. Bars represent the mean ± SEM of three independent experiments. C) Monitoring of PKH67-labeled sEVs internalization by MRC5 cells by flow cytometry. Dot plots represent MRC5 cell fluorescence upon incubation with PKH67-stained sEVs (200 sEVs/cell) for cells that have not been incubated with sEVs (NT, non-treated), or upon 2 h or 16 h of incubation. X-axis: PKH-67 fluorescence intensity; Y-axis: FSC. Gate indicates cells positive for PKH67.

### sEVs from hCMV-infected HIPECs potentialize further infection of recipient MRC5 cells

Based on the proteomic data suggesting a proviral activity of sEVs, we assessed their ability to modulate hCMV infection. As sEV content was composed of proteins prone to act on early steps of infection - entry and immediate early transcriptions - we reasoned that any putative action of sEVs should take place immediately after delivery of their cargo into recipient cells. In this regard, different amounts of sEVs from NI or hCMV-infected HIPECs were incubated with MRC5 cells either concomitantly or 2 h before hCMV infection (Figure 4A). 24 h later, cells were subjected to an anti-IE1/2 immunofluorescence (Figure 4B). When added alone, sEVs from infected HIPECs did not lead to any detectable expression of IE1/2 in MRC5 cells (Figure 4Ba), confirming that they were not contaminated by residual infectious viral particle. This also indicates that the detection of IE proteins upon infection was due to viral gene expression and not to the presence of IE proteins carried by sEV from hCMV-infected HIPECs, even if IE proteins 1 et 2 were detected in proteomic analysis. As shown in Figure 4C, when added simultaneously with hCMV, sEVs did not influence the level of MRC5 cell infection, whatever their origin and quantity, as compared to non-treated cells. In contrast, the addition of increasing doses of sEV from infected HIPECs 2 h before infection led to a potentiation of the infection, when compared to cells treated with sEV from NI HIPECs. This increase was significant for the two highest doses of sEVs applied, with a stimulation of infection of around 17 % and 30 % when 50 and 200 sEVs were added per cell, respectively (Figure 4D).

**Figure 4:**
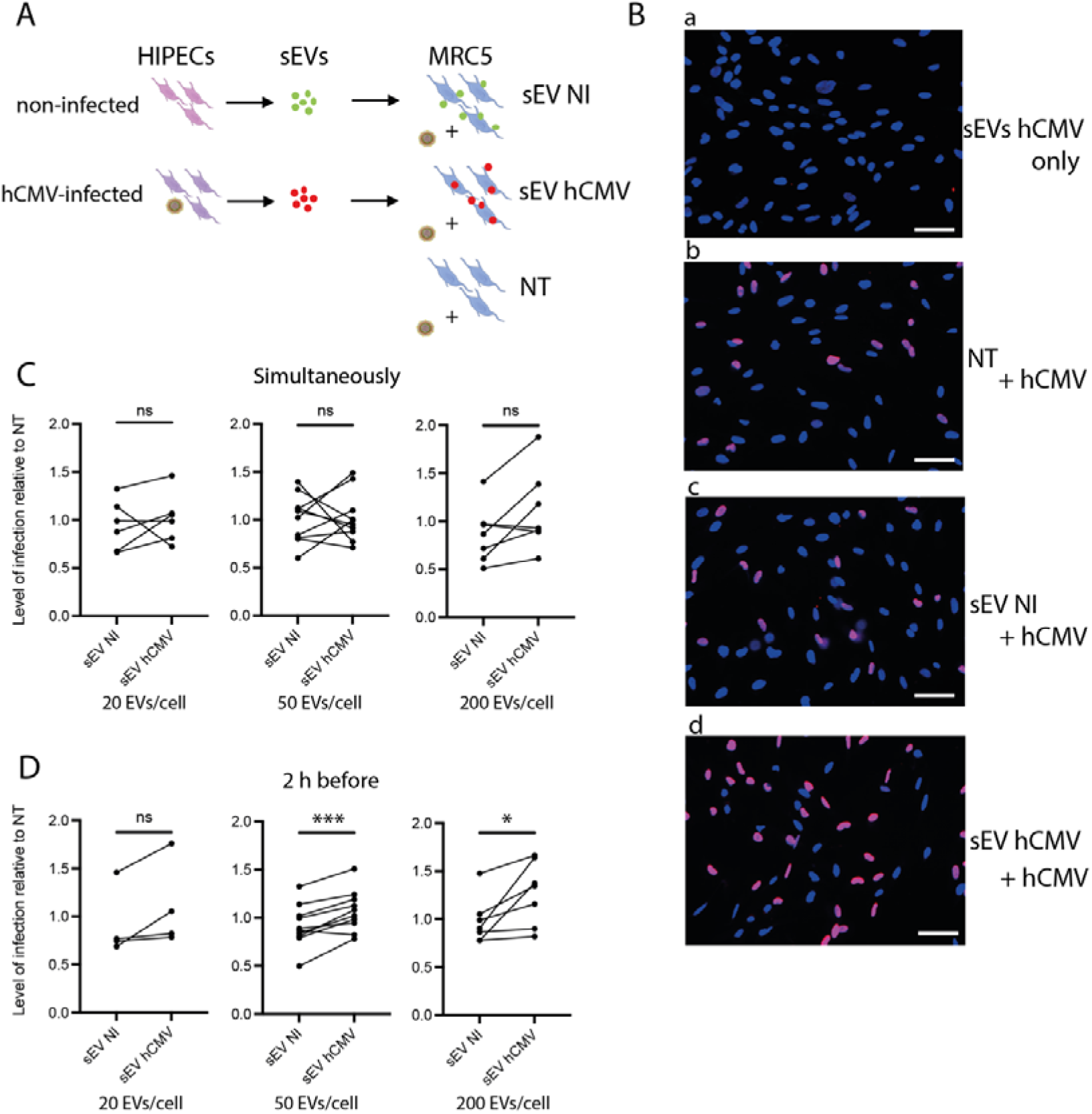
Effect of HIPEC sEVs on MRC5 cell permissiveness for hCMV. A) Experimental procedure (NT=non-treated). B) Immunofluorescence performed on MRC5 cells against IE viral antigen (blue: DAPI; red: IE1/2). a- Non-infected control MRC5 cells upon 24 h incubation with sEVs isolated from hCMV-infected HIPECs. b,c,d- MRC5 cells were either non-treated (b- NT) or incubated during 2 h with sEVs prepared from non-infected (c- sEV NI) or infected HIPECs (d- sEV hCMV) with 50 sEV per cell, then infected during 24 h with hCMV at a MOI of 0.5 before proceeding to immunofluorescence. Scale bar: 200 μm. C-D) MRC5 cells were incubated with sEVs prepared from non-infected (sEV NI) or hCMV-infected (sEV hCMV) HIPECs and infected by hCMV at a MOI of 0.5, concomitantly (C) or 2 h after sEV incubation (D). Three increasing doses of sEVs were used in these experiments (20, 50 or 200 sEV per MRC5 cell, from left to right). 24 h later, expression of IE antigen was assessed by immunofluorescence. Quantification of the percentage of IE positive cells was carried out and normalized by the percentage of cells infected by hCMV without any sEV (NT). Each dot is an independent experiment and corresponds to the mean of the counting of 10 fields, with around 70 cells counted, *i.e*., around 700 cells per dot. n = 4 to 10 independent experiments. Since sEVs used in infection assays were prepared each time in parallel between non-infected or hCMV-infected HIPECs from a given batch, statistical analysis was done by pairing the results between sEV NI and sEV hCMV for each independent experiment. ns, non-significant; *, *p* < 0.05; *** *p* < 0.001 by paired *t*-test.

### sEVs from hCMV-infected HIPECs, placental tissues and amniotic fluids enhance infection of human neural stem cells

To examine the potential proviral role of placental sEVs on hCMV transmission towards the fetal brain, similar experiments using human neural stem cells (NSCs) with different sources of sEVs were performed. NSCs, which are permissive for hCMV infection, constitute a particularly relevant model for studying the consequences of hCMV infection on fetal neural progenitors (7, 48). As observed with MRC5 cells, sEVs from hCMV-infected HIPECs promote a significant enhancement of hCMV infection of NCSs as compared to sEVs from NI HIPECs, with a significant mean increase of 42 % (Figure 5A). To get closer to physiological conditions, an *ex vivo* model of first trimester placental histocultures, that we have previously developed (49, 50), was next infected by hCMV and used for sEV isolation. Recent data obtained from this model showed a modification of sEV surface markers upon hCMV infection and suggested a proviral role for placental sEVs (25). Again, sEVs produced by infected placental histocultures significantly potentiated hCMV infection of NSCs in comparison to sEVs secreted by NI histocultures, with a mean of 48 % of increase (Figure 5B).

**Figure 5:**
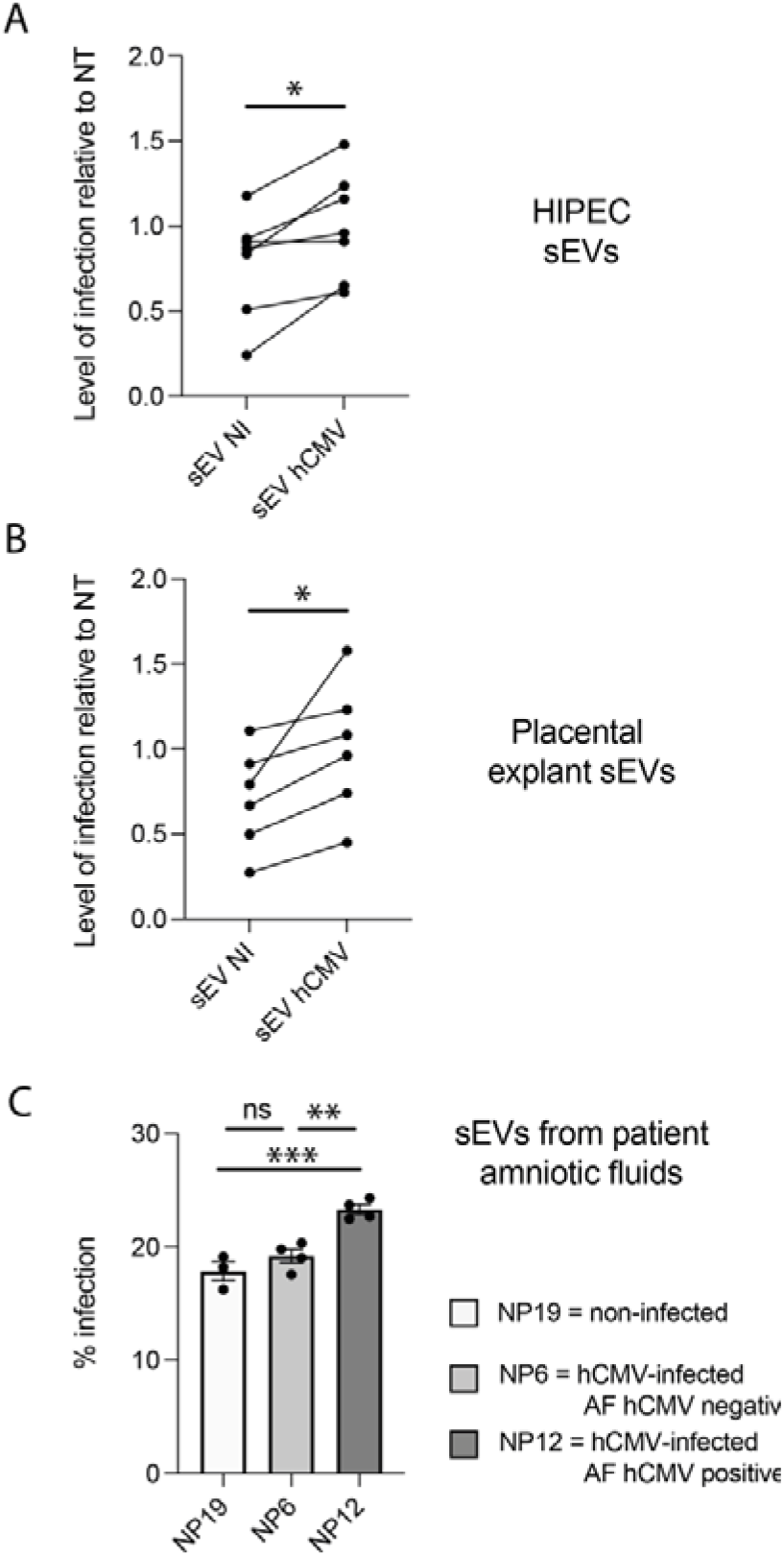
Effect of sEVs from different sources on neural stem cells permissiveness for hCMV. NSCs were incubated with sEVs (200 sEVs per cell) prepared from HIPECs (A), *ex vivo* first trimester placental histoculture (B), or amniotic fluids from naturally infected patients (C), then infected by hCMV at a MOI of 3. 24 h upon infection, expression of IE antigen was assessed by immunofluorescence. Each dot is an independent experiment and corresponds to the mean of the counting of 10 fields, with around 70 cells counted, *i.e*. around 700 cells per dot. n = 3 to 7 independent experiments. For (A) and (B), HIPECs or placental explants were either non-infected (sEV NI) or hCMV-infected (sEV hCMV). Quantification of the percentage of IE positive cells was carried out and normalized by the percentage of infection of cells infected by hCMV without any sEV (NT). Since sEVs used in functional assays were prepared each time in parallel between non-infected or hCMV-infected cytotrophoblasts or placental explants from a given batch, statistical analysis was done by pairing the results between sEV NI and sEV hCMV for each independent experiment. ns, non-significant; *, p < 0.05 by paired *t*-test. For (C), NSCs were incubated with sEVs prepared from amniotic fluid (AF) from patient NP19 (non infected), NP6 (hCMV infected, AF negative for hCMV) or NP12 (hCMV infected, AF positive for hCMV). Histograms represents the mean ± SEM of the percentage of infected cells in each condition. The result of the ordinary one-way ANOVA test performed is *p* = 0.0006 (***). Statistical comparison between the conditions were done by Tukey’s multiple comparisons test upon one-way ANOVA test. ns, non-significant; ** *p* < 0.005; *** *p* < 0.001.

Finally, we assessed the impact of sEVs prepared from amniotic fluid (AF), that contain an important proportion of placental sEVs (51), obtained from naturally infected pregnant women (Figure 5C). AF was collected from 3 women, whose clinical data are summarized in Supplementary Table S3. The first one (NP19) had no history of infection or other maternal disease during pregnancy. The second one (NP6) presented a first trimester hCMV seroconversion with her AF negative for hCMV by qPCR and no fetal defects reported by imaging. The third one (NP12) also seroconverted during first trimester of pregnancy but her AF was positive for hCMV by qPCR and the fetus presented major brain lesions. A control experiment verifying that no infectious particle was present in sEVs preparations was systematically performed in parallel (data not shown). As shown in Figure 5C, level of infection of NSCs incubated with sEVs prepared from the non-infected patient NP19 or from NP6 (infected patient but with AF negative for hCMV) remained similar. Remarkably, when sEVs were prepared from NP12, a case of hCMV positive pregnancy with severe impairment of fetal development, a significant increase of infection of around 20 % was observed in comparison to incubation with sEVs from NP6 or NP19, indicating that NP12 sEVs favored the enhancement of hCMV infection in recipient NSCs. Hence, our data establish the existence of a tight correlation between the clinical severity of hCMV-complicated pregnancy and the proviral effect of sEVs prepared from patient AF.

## Discussion

Although the role of placental sEVs during normal and pathological pregnancy is extensively studied (18, 20, 23, 24), the impact and consequences of hCMV infection on placental sEVs are far to be deciphered, in particular at the beginning of pregnancy. We recently described that infection of first trimester *ex vivo* placental histocultures modified sEV surface markers, suggesting a potential proviral role of the sEVs (25). However, the difficulty of purifying high sEV amounts from this model hampered the possibility to conduct a study combining exhaustive analysis of sEV composition and biological function. In this work, we used a combination of a cytotrophoblast cell line deriving from first trimester placenta (26), *ex vivo* placental histocultures and patient amniotic fluids to isolate sEVs and examine the impact of hCMV infection on their composition and function.

Despite a large number of publications describing a role of EVs on pregnancy regulation, the nature of examined EVs is often not precisely stated, and confusion remains about the subtypes of EVs studied. Here, we focused our study on sEVs, considering their importance in communication between mother and fetus and the fact that they are often dysregulated during pregnancy pathologies (18, 21, 23). Using the gold-standard method based on differential ultracentrifugation followed by density gradient ultracentrifugation, sEVs devoid of viral particles were systematically isolated in a rigorous manner (52). Their characterization by a combination of complementary approaches revealed that they harbor exosome-like features in terms of size, structure and presence of canonical EV markers (27), but their endosomal origin cannot formally be stated, since we did not examine the mechanisms underlying the biogenesis of sEVs in the present study. Instead, we demonstrated that sEVs from hCMV infected cells, histocultures and prepared from amniotic fluids of infected pregnant women present an altered protein content that facilitate the infection of naive recipient fetal cells.

We found that hCMV infection of HIPECs increased the yield of sEV production. However, this increase was counterbalanced by a reduced cell growth upon infection. Hence, the global quantity of sEVs harvested from hCMV- or mock-infected HIPECs remained similar between both conditions. As hCMV mainly replicates in discrete foci in placenta *in vivo*, it is unlikely that hCMV infection would impact on the global quantity of sEVs secreted by the placenta, but we cannot exclude the possibility that local increase of sEV secretion induced by infection may modify placental micro-environment.

Interestingly, most of the viral proteins identified in our proteomic study have not been described in the other proteomic study done of sEVs secreted from hCMV-infected MRC5 cells (39). Conversely, some viral proteins identified by Turner *et al*, such as IR11, IRL12, US14, were not found in our study. One plausible explanation is that, at relatively early time points within the replication cycle (48 h to 72 h upon infection in our present study) the content of sEVs secreted - highly enriched in viral envelop, capsid, tegument and immediate early proteins - may prime the neighboring cells for viral dissemination and spread. In contrast, at latter time point (5 days post infection in Turner *et al* study), the sEV protein cargo could more serve immunoevasion functions, by expressing for example the viral Fc-gamma receptor homologue IR11/gp34 (39). Hence, it seems that composition of sEV secreted by hCMV-infected cells evolves with time, depending on their state of infection and the step of hCMV replication cycle.

In addition to the viral proteins carried by HIPEC sEVs, sEV composition for proteins of cellular origin was also altered upon hCMV infection. By using IPA to analyze biological pathway, the term “autophagy” was the first over-represented in sEVs from infected HIPECs, likely reflecting the autophagy activation induced by hCMV in host cells at the very early times of infection (30–32). Importantly, TRS1 and IRS1, described to antagonize autophagy at latter time points of the viral cycle (30, 31), were also found in sEVs, which may consequently inhibit the induction of autophagy in recipient cells upon sEV uptake. Thy-1, which plays an important role for favoring hCMV entry into cells by macropinocytosis (28, 29), was also highly over-represented in sEVs from infected cells. Hence, all the elements brought by the proteomic analysis of sEVs from infected HIPECs indicate that they were prone to facilitate viral infection of recipient cells.

By examining the function of sEVs from NI or hCMV-infected HIPECs, we observed that sEVs hCMV enhanced further infection of MRC5 cells. Such proviral properties of sEVs produced by infected cells have already been described for other *Herpesviridae* (53, 54) and very recently for hCMV (40) - although not in a placental context - and confirm a general role of sEVs in modulating viral transmission for many viral families (55, 56). Since placental sEVs are found both in maternal and fetal sides at high quantities as soon as the early beginning of pregnancy (57, 58), we hypothesized that they may facilitate hCMV dissemination not only in the placental tissues but also towards the fetus, notably in the fetal brain, by enhancing hCMV infectivity in fetal neural cells. Indeed, we observed that sEVs prepared from HIPECs or first trimester placental explants enhanced hCMV infection of human NSCs. Finally, our observations were confirmed using clinical samples. Strikingly, a significant enhancement of infection was observed with sEVs isolated from the amniotic fluid of an infected woman (with amniotic fluid positive for hCMV) whose fetus showed severe neurodevelopmental impairments, in contrast to sEVs isolated from the amniotic fluid of a non-infected pregnant woman, or an infected woman with amniotic fluid negative for hCMV whose fetus showed no detectable developmental defects. Hence, placental sEVs seem to be key players of hCMV dissemination towards the fetus during congenital infection and may be one of the missing links explaining the variety and severity of sequelae observed during viral congenital infection.

## Materials and Methods

### Human ethic approval

The use of NSCs from human embryonic stem cells was approved by the French authorities (Agence de la Biomédecine, authorization number SASB0920178S).

The biological resource center Germethèque (BB-0033-00081; declaration: DC-2014-2202; authorization: AC-2015-2350) obtained the written consent from each patient (CPP.2.15.27) for the use of human samples and their associated data. For first trimester placenta explants, the steering committee of Germethèque gave its approval for the realization of this study on Feb 5^th^, 2019. The hosting request made to Germethèque bears the number 20190201 and its contract is referenced under the number 19 155C. For amniotic fluid, approval was obtained on July 12^th^, 2019, the hosting request bears the number 20190606 and the contract is referenced under the number DIR-20190823022.

### Cell lines

HIPECs were obtained from Dr T. Fournier (Inserm, Paris; Transfer agreement n°170448). The expression of the cytokeratin 7 specific cytotrophoblastic marker was verified (data not shown). They were cultured in DMEM / F12 medium (Gibco) at 50/50 ratio (v/v), in the presence of 10% fetal bovine serum (FBS, Sigma-Aldrich), 100 U/ml penicillin - 100 μg/ml streptomycin (Gibco) and 100 μg/ml normocin (Invivogen).

MRC5 cells (RD-Biotech) were cultured in Dulbecco’s Modified Eagle Medium (DMEM with Glutamax, Gibco) with the same supplementation as HIPECs.

NSCs were obtained from Dr A. Benchoua (I-Stem, Evry, France), they were produced from ES human cells (SA001, I-STEM, UMR861 France) (48) and maintained in growth medium as described (7). Stem character of NSCs was systematically assessed by immunofluorescence against Nestin and SOX2 proteins (data not shown).

Cell cultures were checked for the absence of mycoplasma (Plasmotest, Invivogen, Toulouse, France).

### Virus production, titration and infection

The endotheliotropic VHL/E strain of hCMV - a gift from Dr C. Sinzger, University of Ulm, Germany - was used in this study (59). Viral stocks were obtained upon amplification of the virus on MRC5 cells, concentrated by ultracentrifugation, and titrated by indirect immunofluorescence as described (7). In some experiments, virus titration was also performed by qPCR from cell culture supernatants (60).

### sEV preparation

Culture medium was previously depleted from EVs to obtain “Exofree” medium: DMEM supplemented with 20 % FBS was ultracentrifuged at 100,000 g for 16 hours at 4 °C and filtered at 0.22 μm. Exofree medium was then obtained by a 1:1 dilution with F12 to reach 10 % FBS.

4 million HIPECs were seeded in 150 cm^2^ flask, with 6 flasks per condition. 24 h later, cells were infected or not by hCMV at multiplicity of infection (MOI) of 10 (Supplementary Figure S1A). Culture medium was replaced by Exofree medium 24 h post-infection, after having previously ensured that this did not affect the cell growth (Supplementary Figure S1E). Culture supernatants were collected at 48 h and 72 h post-infection. Medium were pooled for each condition (non-infected or infected) and submitted to sEV preparation protocol. Cell number at the end of the experiment was determined by counting upon trypsinization (Supplementary Figure 1D) and cell viability evaluated by trypan blue.

Procedures for sEV preparation were realized as described previously (25), according to ISEV guidelines (27). All relevant data were submitted to the EV-TRACK knowledgebase (EV-TRACK ID: EV210154) and obtained an EV-METRIC score of 100 % for HIPEC and placental explant EVs (61).

### Nanoparticle tracking analysis (NTA)

sEV preparations were tracked using a NanoSight LM10 (Malvern Panalytical) equipped with a 405 nm laser. Videos were recorded three times for each sample at constant temperature (22 °C) during 60 s and analyzed with NTA Software 2.0 (Malvern instruments Ltd). Data were analyzed with Excel and GraphPad Prism (v8) softwares.

### Transmission electron microscopy and immunolabeling electron microscopy

Procedures were performed essentially as described (25, 62, 63). Immunodetection was carried out with the following primary antibodies: mouse anti-human CD63 (Abcam ab23792), mouse anti-human CD9 or mouse anti-human CD81 (both from Dr E. Rubinstein, Université Paris-Sud, Institut André Lwoff, Villejuif, France). Secondary incubation was performed with a rabbit anti mouse Fc fragment (Dako Agilent Z0412), then grids were incubated with Protein A-Gold 10 nm (Cell Microscopy Center, Department of Cell Biology, Utrecht University). All samples were observed with a Tecnai Spirit electron microscope (FEI, Eindhoven, The Netherlands), and digital acquisitions were made with a numeric 4k CCD camera (Quemesa, Olympus, Münster, Germany). Images were analysed with iTEM software (EMSIS) and statistical studies were done with GraphPad Prism software (v8).

### Western blot

Western blots were realized as previously described (25), by using the following primary antibodies: mouse anti-CD81 (200 ng/ml, Santa-Cruz), mouse anti-CD63 (500 ng/ml, BD Pharmingen), mouse anti-CD9 (100 ng/ml, Millipore), rabbit anti-Tsg101 (1 μg/ml, Abcam), rabbit anti-Alix (1 μg/ml, Abcam), mouse anti-Thy1 (0.5 μg/ml, Biolegend), rabbit anti-Tom20 (1/500, Sigma) or goat anti-Calnexin (2 μg/ml, Abcam). After incubation with the secondary antibody, membranes were visualized using the Odyssey Infrared Imaging System (LI-COR Biosciences).

### Quantitative proteomic analysis

#### Sample preparation

Protein samples in Laemmli buffer (3 biological replicates of sEVs preparation from non-infected and hCMV-infected cytotrophoblasts cells) were submitted to reduction and alkylation (30 mM DTT and 90 mM iodoacetamide, respectively). Protein samples were digested with trypsin on S-trap Micro devices (Protifi) according to manufacturer’s protocol, with the following modifications: precipitation was performed using 545 μl S-Trap buffer and 1 μg Trypsin was added per sample for digestion.

#### NanoLC-MS/MS analysis

Peptides were analyzed by nanoLC-MS/MS using an UltiMate 3000 RSLCnano system coupled to a Q-Exactive-Plus mass spectrometer (Thermo Fisher Scientific, Bremen, Germany). Five μL of each sample were loaded on a C-18 precolumn (300 μm ID x 5 mm, Dionex) in a solvent made of 5 % acetonitrile and 0.05 % TFA and at a flow rate of 20 μL/min. After 5 min of desalting, the precolumn was switched online with the analytical C-18 column (75 μm ID x 15 cm, Reprosil C18) equilibrated in 95 % solvent A (5 % acetonitrile, 0.2 % formic acid) and 5 % solvent B (80 % acetonitrile, 0.2 % formic acid). Peptides were eluted using a 5 to 50 % gradient of solvent B over 105 min at a flow rate of 300 nL/min. The Q-Exactive-Plus was operated in a data-dependent acquisition mode with the XCalibur software. Survey scan MS were acquired in the Orbitrap on the 350-1500 m/z range with the resolution set to a value of 70000. The 10 most intense ions per survey scan were selected for HCD fragmentation. Dynamic exclusion was employed within 30 s to prevent repetitive selection of the same peptide. At least 3 injections were performed for each sample.

#### Bioinformatics data analysis of mass spectrometry raw files

Raw MS files were processed with the Mascot software for database search and with Proline (64) for label-free quantitative analysis. Data were searched against *Human herpesvirus* 5 and Human entries of the UniProtKB protein database (Human betaherpesvirus 5 clone VHL-E-BAC19 and release Uniprot Swiss-Prot February 2018). Carbamidomethylation of cysteines was set as a fixed modification, whereas oxidation of methionine was set as variable modification. Specificity of trypsin/P digestion was set for cleavage after K or R, and two missed trypsin cleavage sites were allowed. The mass tolerance was set to 10 ppm for the precursor and to 20 mmu in tandem MS mode. Minimum peptide length was set to 7 amino acids, and identification results were further validated in Proline by the target decoy approach using a reverse database at both a PSM and protein false-discovery rate of 1%. After mean of replicate injections, the abundance values were log2 transformed and missing values were replaced by random numbers drawn from a normal distribution with a width of 0.3 and down shift of 1.8 using the Perseus toolbox (version 1.6.7.0). For statistical analysis, a Student *t*-test (two-tailed *t*-test, equal variances) was then performed on log2 transformed values to analyse differences in protein abundances in all biologic group comparisons. Significance level was set at *p*□ = 0.05, and log2 ratios were considered relevant if higher than 1 or lower than −1. The mass spectrometry proteomics data have been deposited to the ProteomeXchange Consortium via the PRIDE (65) partner repository with the dataset identifier PXD029146.

### Functional proteomic data analysis

Volcano plot was established for proteins whose mean abundance exhibited a log2 ratio higher than 1 or lower than −1 and when Student’s *t*-test *p*-values were <= 0.05 between the infected and the non-infected conditions. The list of human proteins exhibiting a normalized mean protein abundance log2 ratio > 1 or < −1 between sEVs from non-infected or hCMV-infected samples was used as an input for analysis with QIAGEN Ingenuity Pathway Analysis (IPA) (66). Results from IPA biological functions and diseases analysis were filtered to retrieve annotations having an absolute activation z-score > 1 and defined by less than 150 molecules. The resulting annotations were manually curated to remove redundant annotations sharing identical genes, keeping annotations defined by the greater number of molecules.

### Flow cytometry analysis

After incubation of cells with PKH67-stained sEVs, cells were washed twice with PBS and trypsinized, before proceeding to flow cytometry analysis. PKH67 positive cells were analyzed on a Macsquant VYB Flow Cytometer (Miltenyi Biotec), by using FCS and FITC fluorescence parameters, and by subtracting cell autofluorescence background. Data were analyzed with FlowJo (BD) and GraphPad Prism (v8) software.

### Immunofluorescence

Cells were fixed using 4 % PFA (Electron microscopy Sciences) at room temperature for 20 min. Permeabilization was performed with PBS 0.3 % Triton-X100 (Thermofisher scientific) for 10 min, followed by 1 h incubation in blocking buffer (PBS with 5 % FBS). Incubation with primary antibodies diluted in blocking buffer was carried out overnight at 4 °C, against hCMV immediate early protein 1 and 2 (1 μg/ml; Abcam IE1/IE2 CH160 ab53495), nestin (4 μg/ml; Abcam 10C2 ab 22035), or SOX2 (1/500 of stock; Cell Signaling D6D9 #3579). Secondary antibody incubation (Goat anti mouse or rabbit - Alexa-fluor 488 or 594 (2 μg/ml; Thermo Fischer Scientific)) was performed at room temperature for 1 h. For actin staining, Alexa-fluor 568 phalloidin (5 μg/ml; Thermo Fischer Scientific A12380) was incubated on cells overnight at 4°C. DAPI staining (1 μg/ml; Sigma) was performed for 10 min at room temperature. ProLong Gold without DAPI (Thermo Fischer Scientific) was used for coverslip mounting.

Widefield acquisitions were realized using Apotome microscope (Zeiss) and confocal acquisitions were made on SP8-STED microscope (Leica). Image processing was performed using ImageJ. GraphPad Prism (v8) software was used to perform data statistical analysis.

### Placental histoculture

Placental histocultures were carried out as described (25, 49) on first trimester placentas (4 placentas; mean = 13.11 ± 0.49 (SEM) weeks of amenorrhea, *i.e*., 11.11 ± 0.49 weeks of pregnancy; age of the women: mean = 23 ± 1.5 (SEM) year-old). Briefly, trophoblastic villi were dissected in small explants and infected or not by hCMV overnight before deposition on gelatin sponges (Gelfoam, Pfizer) in Exofree medium. Conditioned medium was collected and renewed every 3 to 4 days. At 14 days of culture, collected medium was pooled for each condition and used to perform sEV preparation. Placental explants were weighed for normalization of resuspension volume.

### Amniotic fluid collection

Remaining amniotic fluid (AF) collected during classic patient care was saved for the present study. It was subjected to a 10 min centrifugation at 3000 rpm in order to remove the cells and large debris before its storage at −80 °C. The sEV preparation was carried out according to the procedure described above.

## Acknowledgments

We greatly thank C. Chouquet, B. Rauwel, M. Mouysset, L. Battut, C. Mengelle, S. Lhomme, F. Chauvrier, P. Verdy, N. Kopf, as well as the whole ViNeDys team, for their technical assistance and their numerous advice and discussions which allowed the progress of this work. We thank also AL. Iscache, V. Duplan-Eche and F. L’Faqihi-Olive, from the cytometry facility of Infinity, as well as S. Allart and D. Daviaud, from the imaging facility of Infinity, and finally the GeT technical service of Infinity. We thank the medical and paramedical staff of the gynecology unit at Paule de Viguier Hospital, the patients who agreed to participate in the study, as well as L. Bujan and M. Aubry, from the Germethèque. We finally warmly thank E. Haanappel from the Institute of Pharmacology and Structural Biology, who granting us access to the Nanosight device, and Audrey Esclatine and Daniel Gonzalez-Dunia for critical reading of the manuscript and insightful comments.

This project has received financial support from the French Biomedicine Agency, institutional grants from Inserm, CNRS and Toulouse 3 University. This project is part of the doctorate thesis of Mathilde Bergamelli, who was funded by the Ministry of Education and Research (MESR). The TEM experiments were performed on PICT-IBiSA, Institut Curie, Paris, member of the France-BioImaging national research infrastructure, and were supported by the French National Research Agency through the “Investments for the Future” program (France-BioImaging, ANR-11-INSB-04), supported by the CelTisPhyBio Labex (N° ANR-11-LB0038) part of the IDEX PSL (N°ANR-10-IDEX-0001-02 PSL). The proteomic part of this project was supported in part by the Région Occitanie, European funds (Fonds Européens de Développement Régional, FEDER), Toulouse Métropole, and by the French Ministry of Research with the Investissement d’Avenir Infrastructures Nationales en Biologie et Santé program (ProFI, Proteomics French Infrastructure project, ANR-10-INBS-08).

---

**Supplementary Figure S1:**
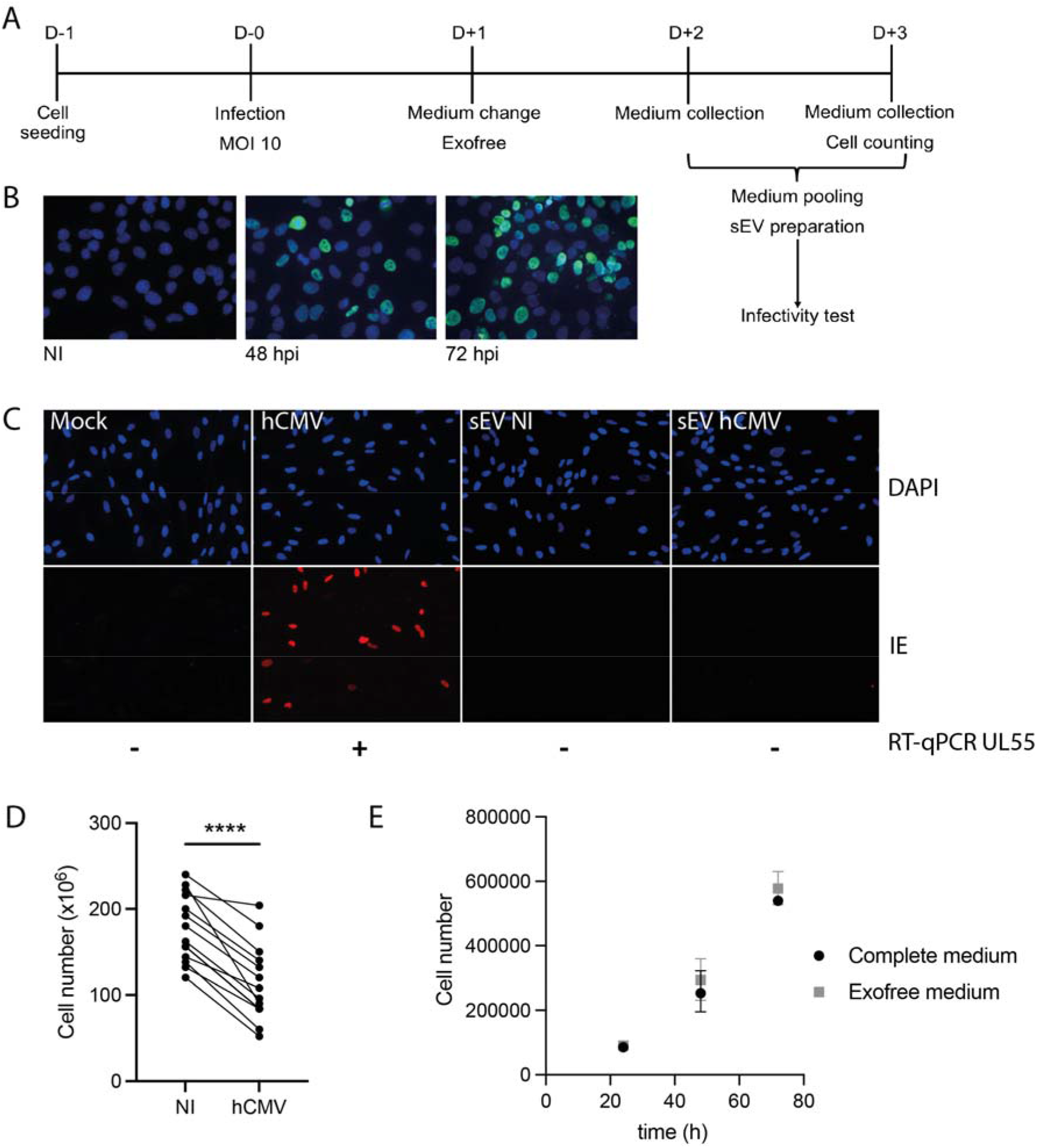
Model system. A) Timeline representing the HIPEC seeding, infection, medium renewal and collection before sEV preparation. D: days; MOI: Multiplicity of infection. B) HIPECs are permissive to hCMV infection. Immunofluorescence against Immediate Early 1/2 (IE) antigens was done on HIPECs either non-infected (NI) or at 48 h or 72 h post-infection (hpi). Green: IE; Blue: DAPI; Magnification: 20 x. C) Immunofluorescence realized against hCMV IE1/2 antigens in nontreated MRC5 cells (Mock), or upon incubation during 24 h with either hCMV at MOI 3 (hCMV), sEVs isolated from non-infected (sEV NI) or from hCMV-infected HIPECs (sEV hCMV). Magnification = 20 x. Blue (upper panel): DAPI; red (lower panel): IE. Below images are indicated the results of RT-qPCR realized against hCMV UL55 mRNAs on RNA extracted from MRC5 cells at 48 h post-incubation with hCMV or sEV preparations (+: amplification; -: no amplification). Data are representative for at least three independent experiments. D) Total cell number counted at the time of sEV preparation, at 72 hpi, for non-infected (NI) or hCMV infected HIPECs. Results are representative of 15 independent experiments. *p<0,0001* by paired *t*-test. E) Comparison of cell growth between complete and Exofree culture medium, by counting of HIPEC number in 12-well plates in which 10^5^ cells were initially seeded. Symbols represent mean ± SEM for three independent experiments. A 2-way ANOVA statistical test indicated no significant difference between complete and Exofree medium culture conditions (*p=0.4414*).

**Supplementary Figure S2:**
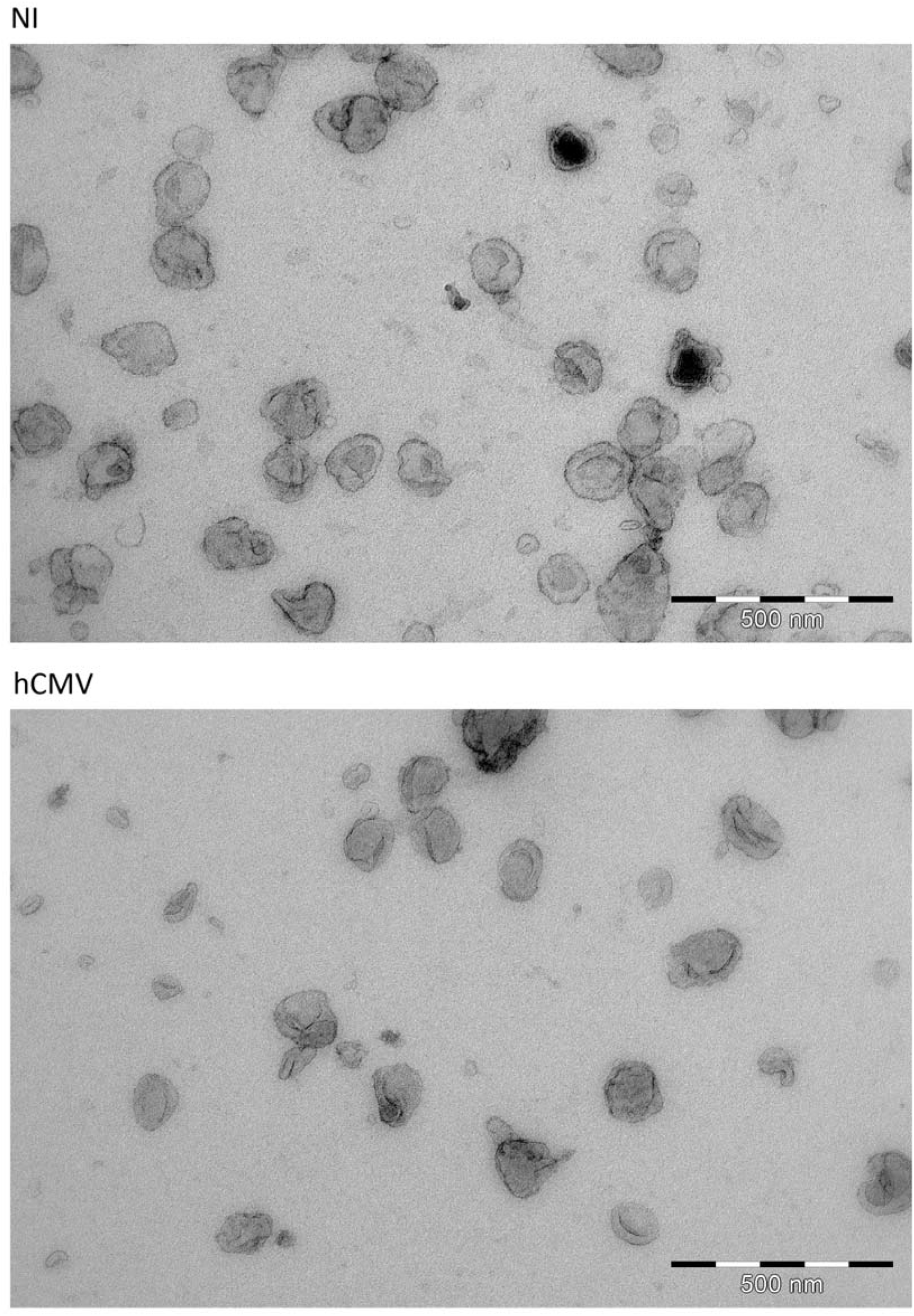
Wide-field transmission electron microscopy. TEM micrographs showing sEV from HIPEC either non-infected (NI; upper panel) or infected
(hCMV; lower panel). Scale bar: 500 nm.

**Supplementary Figure S3:**
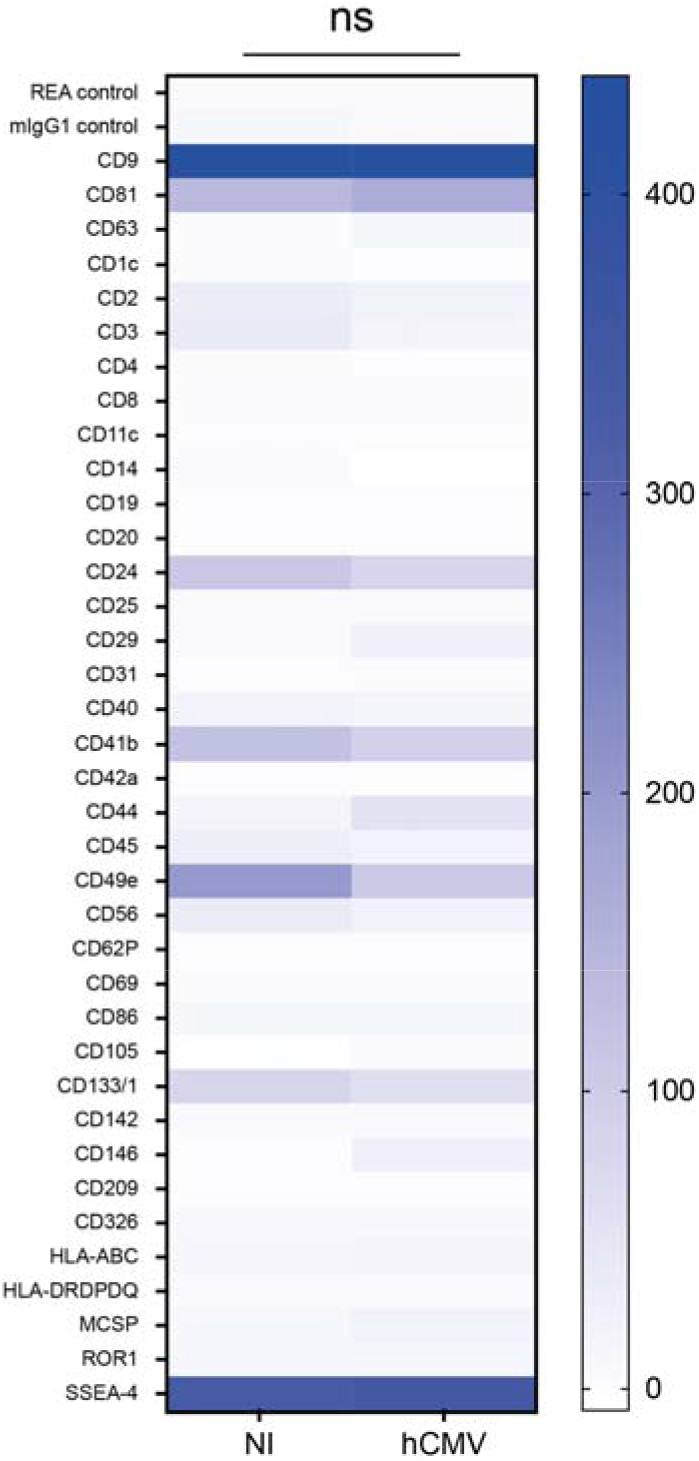
Multiplex bead-based flow cytometry comparison of sEV prepared from non-infected or hCMV-infected HIPECs. Bead-based multiplex analysis using the MACSPlex Exosome Kit, human (Miltenyi Biotec) were realized on sEV preparations by flow cytometry according to the manufacturer’s instructions. This allowed the quantification of 39 different EV markers, distinguishable by flow cytometry by a specific PE and FITC labeling. The MACSQuant Analyzer 10 flow cytometer (Miltenyi Biotec) was used for analysis. The tool MACSQuantify was used to analyze data (v2.11.1746.19438). GraphPad Prism (v8) software was used to perform statistical analysis of the data. The heat-map represents the mean of 3 independent experiments, for different sEV markers indicated on the left column, for sEV prepared from non-infected of hCMV-infected HIPECs. Blue intensity is proportional to the level of expression calculated in Median Fluorescence Intensity, indicated on the right of the heatmap. ns, non-significant by two-way ANOVA.

**Supplementary Figure S4:**
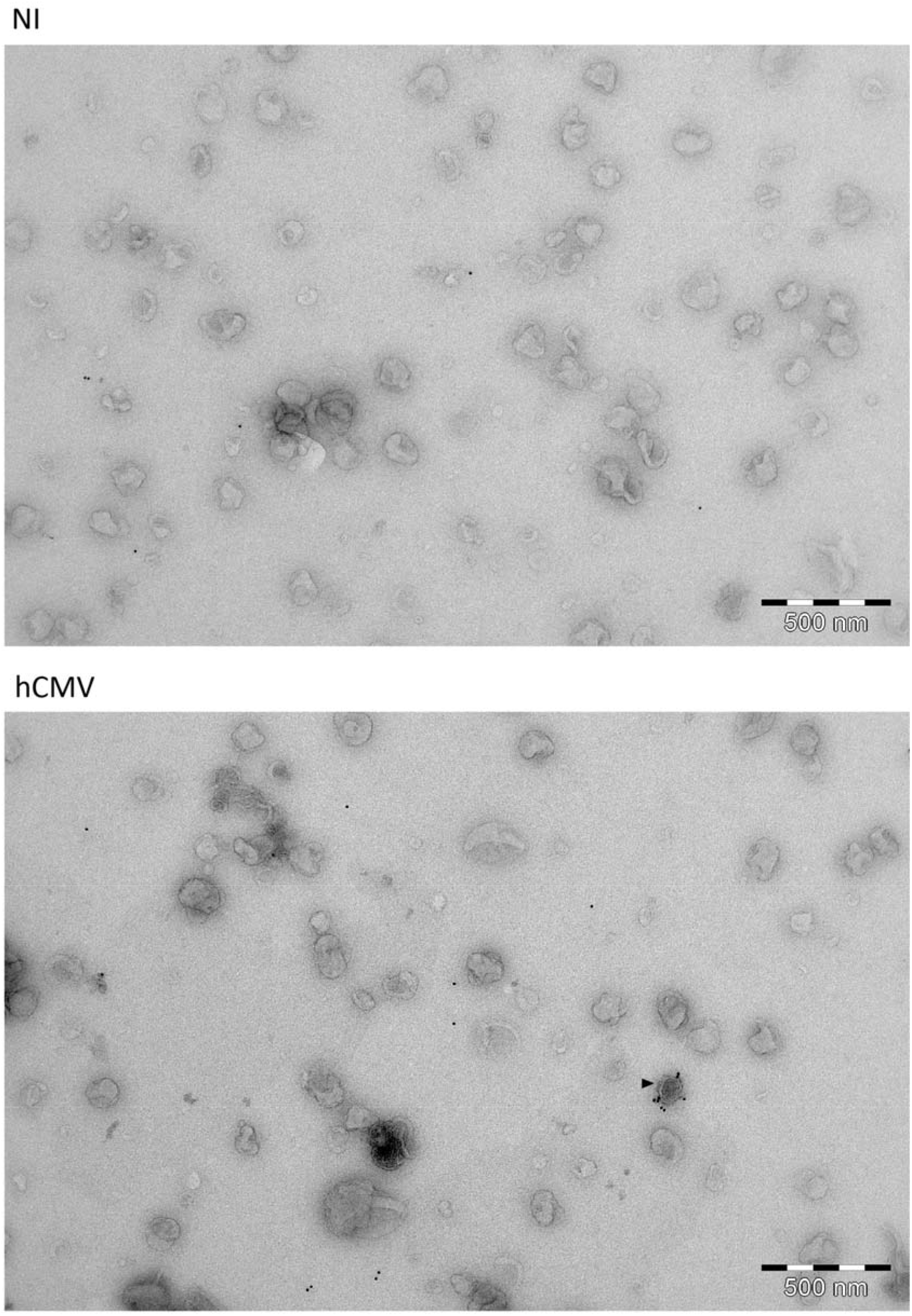
Wide field immuno-electron microscopy anti CD63. TEM micrographs showing sEV from HIPECs either non-infected (NI; upper panel) or infected (hCMV; lower panel), after immunogold labelling against CD63. Arrow in the lower panel indicates a CD63 positive sEV. Scale bar: 500 nm.

**Supplementary Figure S5:**
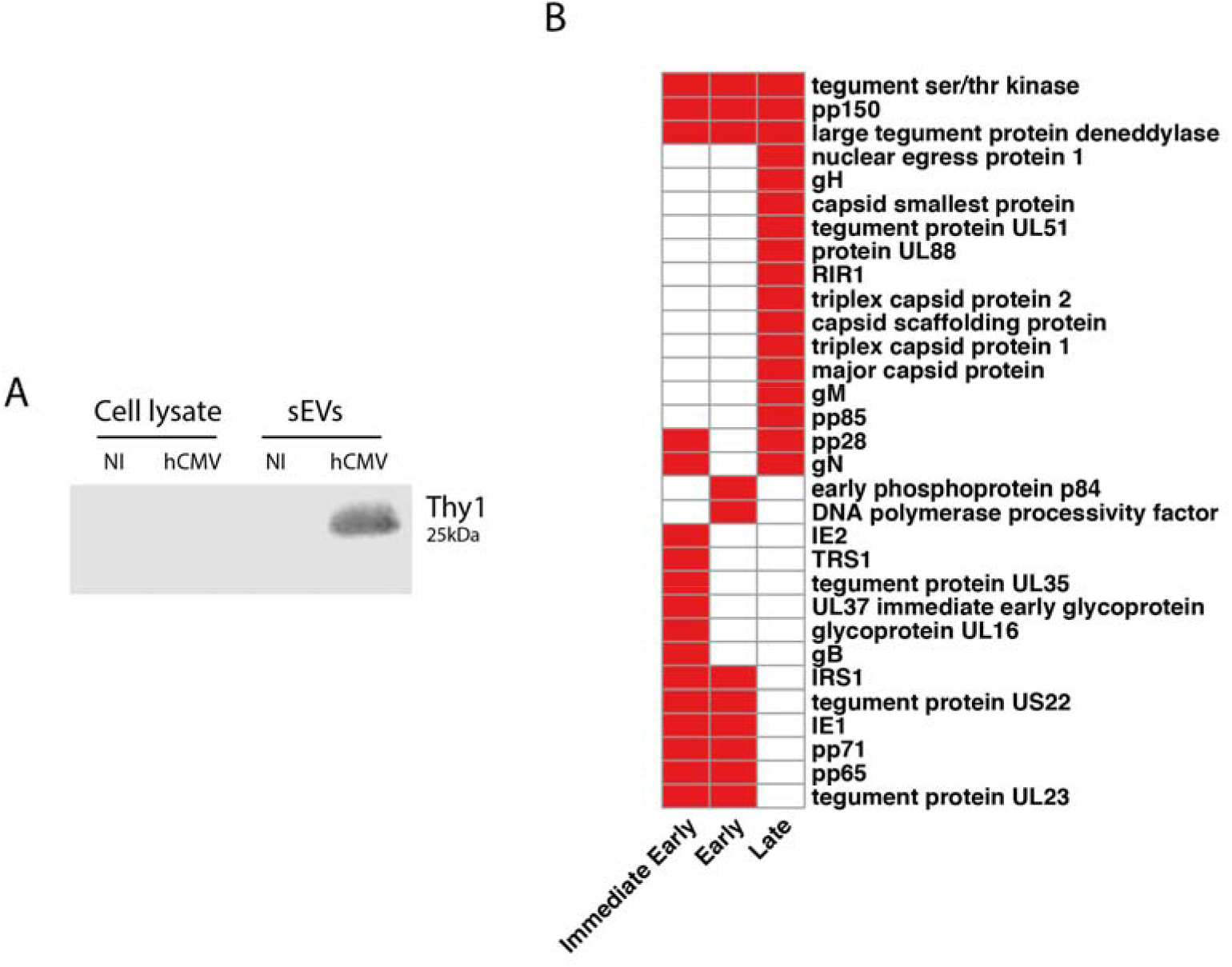
Complement to proteomic analysis. A) Western-blot showing the enrichment of the cellular Thy-1 protein in sEVs from infected HIPECs (NI: non-infected). This result is representative of three independent experiments. B) Heat-map representing the different viral protein found in sEV isolated from infected HIPECs, classified depending on their expression timeline during hCMV infection (Immediate early, early or late).

**Supplementary Figure S6:**
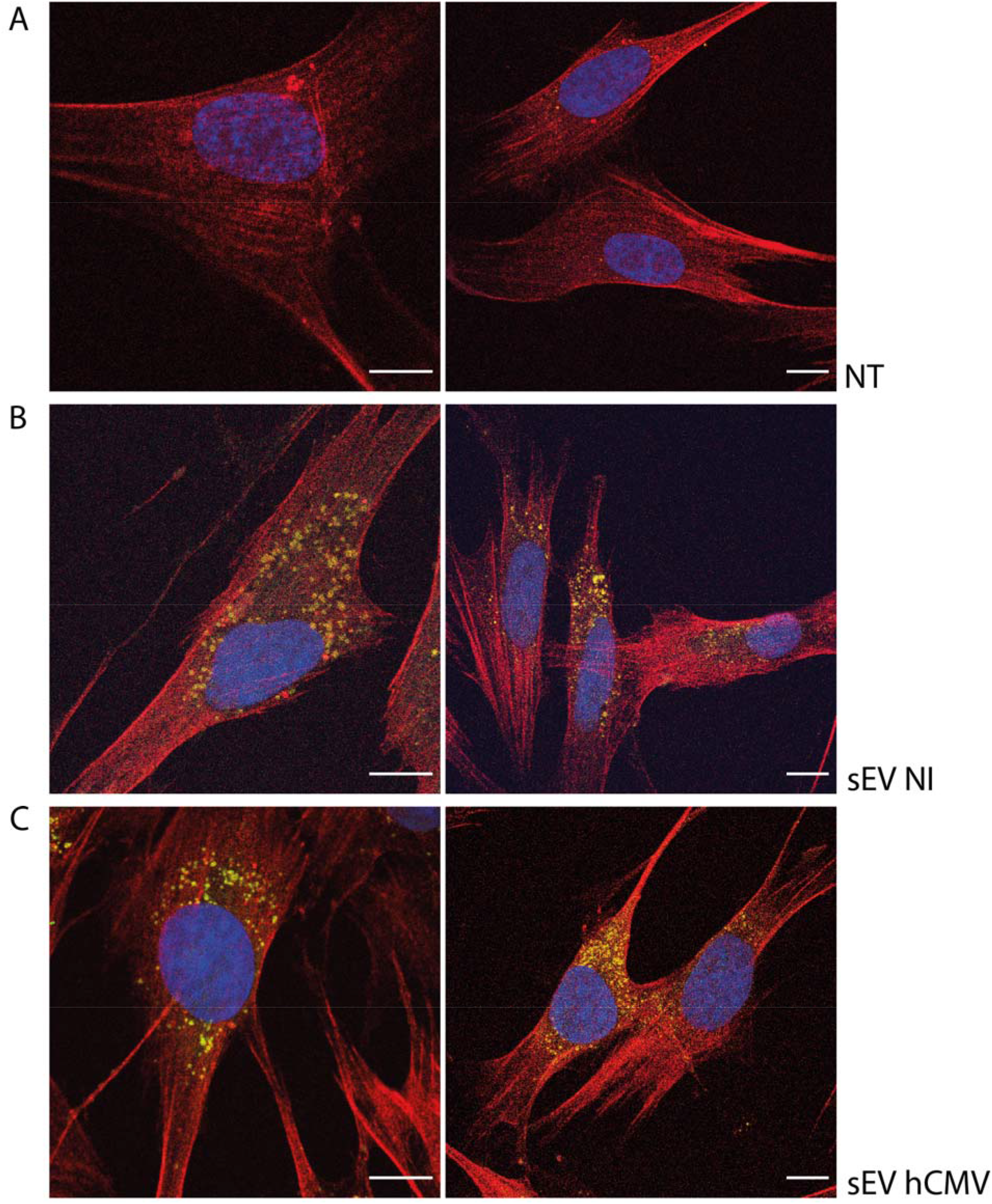
sEV internalization by MRC5 at 2 h post-incubation. MCR5 cells, observed by confocal fluorescence microcopy, upon 2h of incubation with PKH67 stained sEVs (200 sEVs/cell) prepared from non-infected (panels B), hCMV-infected (panels C) or without sEV incubation (panels A). Green: PKH67 staining of sEV; red: actin staining by phalloidin; blue: DAPI. Scale bar: 10 μm.

**Supplementary Video S1: MCR5 cells after 16 h incubation with PKH67 stained sEVs from NI HIPECs (50 sEVs/cells).** Observation by confocal fluorescence microscopy. Blue: DAPI; Red: Phalloidin; Green: PKH67. Magnification = 63 X.

**Supplementary Table S1: Excel file.**

The mass spectrometry proteomics data have been deposited to the ProteomeXchange Consortium via the PRIDE partner repository with the dataset identifier PXD029146.

**Supplementary Table S2:**
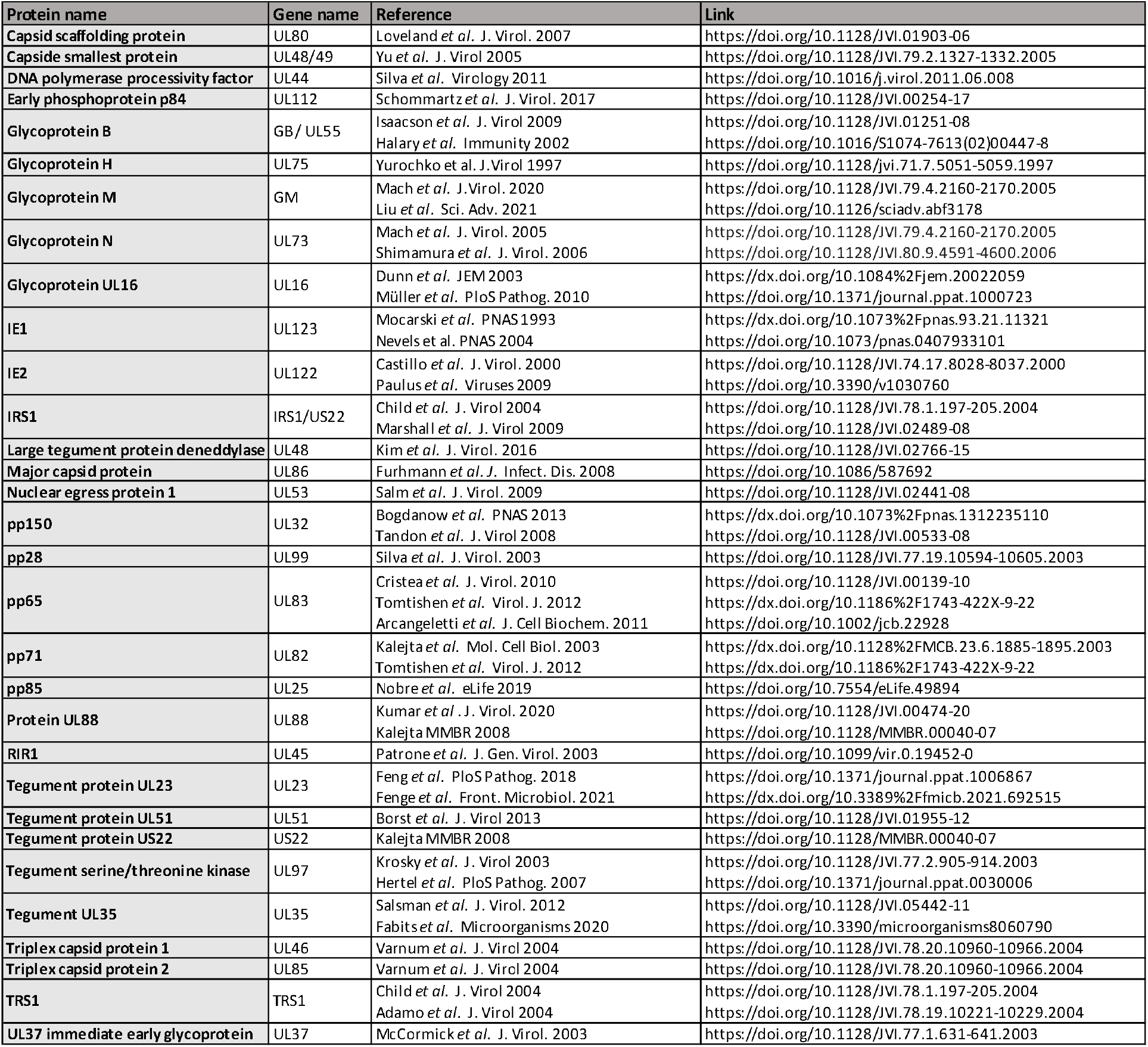
Bibliography references for establishment of Figure 2D and supplementary Figure S5B.

**Supplementary Table S3.**
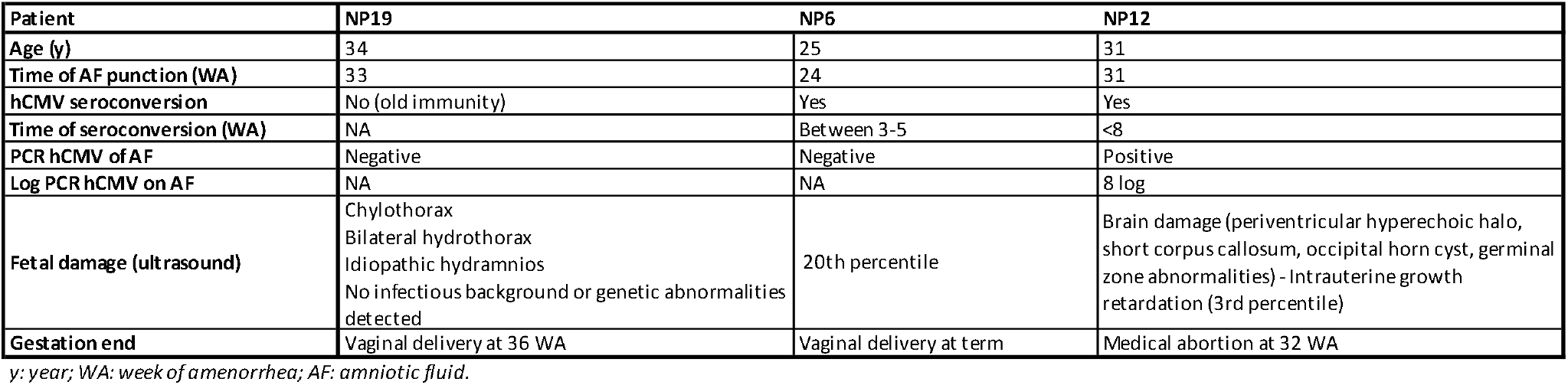

## Notes

**Competing Interest Statement:** The authors declare that the research was conducted in the absence of any commercial or financial relationships that could be construed as a potential conflict of interest.

### Competing Interest Statement

The authors have declared no competing interest.

