## Supplementary material for "Human Cytomegalovirus modifies placental small extracellular vesicle secretion and composition towards a proviral phenotype to enhance infection of fetal recipient cells": Supp Table 1 (from another version of this preprint: version 1, 2021-11-19)

| Accession | Gene_name | Description | Protein_set_score | Coverage | MW | #peptides | Normalized mean protein abundance NI44 | Normalized mean protein abundance NI43 | Normalized mean protein abundance NI37 | Normalized mean protein abundance hCMV44 | Normalized mean protein abundance hCMV43 | Normalized mean protein abundance hCMV37 | Ratio hCMV/NI | T Test | Test | Log 2 ratio hCMV/NI | "-log 10 "T Test |
|---|---|---|---|---|---|---|---|---|---|---|---|---|---|---|---|---|---|
| QBK84361.1 | GM | envelope glycoprotein M [Human betaherpesvirus 5] | 234,229 | 15,01 | 42819 | 5 | 18,63 | 17,76 | 18,93 | 25,30 | 24,99 | 25,24 | **106,637** | 4,97E-05 | 1 | 6,73656336 | 4,30322182 |
| QBK84369.1 | UL16 | tegument protein UL16 [Human betaherpesvirus 5] | 172,205 | 30,43 | 38294 | 7 | 17,70 | 20,28 | 18,54 | 25,54 | 24,57 | 24,95 | **72,503** | 1,59E-03 | 1 | 6,17996724 | 2,79927018 |
| QBK84306.1 | pp150 | tegument protein pp150 [Human betaherpesvirus 5] | 1386,365 | 46,52 | 112773 | 35 | 22,13 | 21,18 | 21,51 | 28,29 | 27,42 | 27,27 | **66,410** | 1,38E-04 | 1 | 6,05333392 | 3,85983375 |
| QBK84312.1 | GB | envelope glycoprotein B [Human betaherpesvirus 5] | 634,712 | 28 | 101855 | 17 | 21,09 | 20,44 | 18,60 | 26,30 | 25,94 | 25,80 | **62,683** | 1,42E-03 | 1 | 5,96999931 | 2,84717455 |
| QBK84326.1 | UL25 | tegument protein UL25 [Human betaherpesvirus 5] | 368,964 | 27,13 | 73525 | 13 | 18,85 | 19,94 | 20,82 | 25,71 | 24,92 | 26,31 | **54,821** | 1,16E-03 | 1 | 5,77666601 | 2,93522096 |
| QBK84340.1 | pp65 | tegument protein pp65 [Human betaherpesvirus 5] | 1980,658 | 72,01 | 62896 | 38 | 26,46 | 22,67 | 23,21 | 29,52 | 28,90 | 30,47 | **45,781** | 1,22E-02 | 1 | 5,51666705 | 1,91465718 |
| QBK84302.1 | MC | major capsid protein [Human betaherpesvirus 5] | 1932,96 | 60,73 | 153866 | 55 | 22,02 | 22,66 | 22,97 | 28,36 | 27,74 | 27,95 | **44,221** | 8,13E-05 | 1 | 5,46666718 | 4,0898573 |
| QBK84386.1 | CT1 | capsid triplex subunit 1 [Human betaherpesvirus 5] | 152,172 | 25,17 | 33027 | 5 | 18,39 | 17,79 | 18,55 | 23,72 | 23,26 | 22,50 | **30,201** | 3,15E-04 | 1 | 4,9165287 | 3,50133475 |
| QBK84322.1 | CM | capsid maturation protease [Human betaherpesvirus 5] | 58,143 | 4,94 | 73886 | 3 | 18,64 | 19,02 | 17,94 | 24,02 | 23,50 | 21,95 | **24,689** | 2,69E-03 | 1 | 4,62580681 | 2,56961975 |
| QBK84381.1 | CT2 | capsid triplex subunit 2 [Human betaherpesvirus 5] | 270,931 | 40,85 | 34595 | 10 | 20,44 | 20,32 | 21,03 | 25,39 | 24,97 | 25,22 | **24,195** | 5,23E-05 | 1 | 4,5966657 | 4,28121836 |
| QBK84387.1 | UL23 | tegument protein UL23 [Human betaherpesvirus 5] | 287,31 | 42,25 | 32957 | 10 | 19,70 | 22,64 | 19,62 | 25,21 | 24,49 | 24,86 | **18,351** | 1,44E-02 | 1 | 4,19780032 | 1,84223379 |
| QBK84341.1 | pp71 | tegument protein pp71 [Human betaherpesvirus 5] | 628,156 | 35,06 | 62008 | 15 | 22,67 | 22,71 | 22,33 | 27,26 | 26,58 | 26,45 | **18,294** | 1,14E-04 | 1 | 4,19333394 | 3,94491169 |
| QBK84419.1 | | myristylated tegument protein [Human betaherpesvirus 5] | 38,345 | 6,32 | 20965 | 2 | 19,30 | 17,66 | 18,02 | 23,17 | 21,48 | 22,74 | **17,574** | 4,34E-03 | 1 | 4,13540395 | 2,36283669 |
| LEG1_HUMA | LGALS1 | sp\|P09382\|LEG1_HUMAN Galectin-1 OS=Homo sapiens GN=LGALS1 PE=1 SV=2 | 86,1 | 33,33 | 14716 | 3 | 18,86 | 19,70 | 18,50 | 24,16 | 22,80 | 22,15 | **16,192** | 4,36E-03 | 1 | 4,0171922 | 2,3607592 |
| THY1_HUMA | THY1 | sp\|P04216\|THY1_HUMAN Thy-1 membrane glycoprotein OS=Homo sapiens GN=THY1 P | 43,572 | 18,01 | 17935 | 2 | 19,29 | 20,56 | 18,33 | 23,48 | 23,35 | 23,37 | **16,058** | 3,45E-03 | 1 | 4,00524076 | 2,46221995 |
| QBK84313.1 | | ribonucleotide reductase subunit 1 [Human betaherpesvirus 5] | 163,71 | 13,58 | 101668 | 12 | 20,66 | 21,25 | 19,18 | 24,85 | 24,29 | 23,76 | **15,307** | 4,69E-03 | 1 | 3,93610128 | 2,32861634 |
| QBK84335.1 | IE1 | regulatory protein IE1 [Human betaherpesvirus 5] | 305,463 | 24,69 | 54876 | 9 | 20,99 | 21,00 | 23,29 | 24,94 | 25,28 | 26,50 | **14,058** | 1,33E-02 | 1 | 3,81333351 | 1,8767252 |
| QBK84338.1 | US22 | tegument protein US22 [Human betaherpesvirus 5] | 406,862 | 29,69 | 65033 | 12 | 21,62 | 18,93 | 22,03 | 23,67 | 23,74 | 25,79 | **11,629** | 4,15E-02 | 1 | 3,53962135 | 1,38229519 |
| QBK84307.1 | UL37 | tegument protein UL37 [Human betaherpesvirus 5] | 111,727 | 6,71 | 110018 | 5 | 18,26 | 17,65 | 17,09 | 21,71 | 20,96 | 20,20 | **9,795** | 3,97E-03 | 1 | 3,29202843 | 2,40164734 |
| RMD3_HUM | RMDN3 | sp\|Q96TC7\|RMD3_HUMAN Regulator of microtubule dynamics protein 3 OS=Homo sap | 9,069 | 1,49 | 52118 | 1 | 18,03 | 19,72 | 19,79 | 22,79 | 21,05 | 22,85 | **8,284** | 2,08E-02 | 1 | 3,05026245 | 1,6820355 |
| QBK84351.1 | UL88 | tegument protein UL88 [Human betaherpesvirus 5] | 22,375 | 5,13 | 47689 | 2 | 18,44 | 17,83 | 20,92 | 22,55 | 22,00 | 21,76 | **8,241** | 3,55E-02 | 1 | 3,04288228 | 1,45006442 |
| QBK84316.1 | IRS1 | tegument protein IRS1 [Human betaherpesvirus 5] | 343,215 | 27,15 | 91114 | 14 | 18,64 | 18,21 | 18,72 | 21,37 | 20,92 | 22,08 | **7,640** | 1,42E-03 | 1 | 2,93353844 | 2,84804343 |
| QBK84300.1 | | large tegument protein [Human betaherpesvirus 5] | 594,245 | 10,93 | 253300 | 16 | 21,28 | 22,00 | 20,97 | 25,05 | 23,97 | 23,93 | **7,464** | 3,70E-03 | 1 | 2,89999962 | 2,43145923 |
| QBK84365.1 | UL51 | tegument protein UL51 [Human betaherpesvirus 5] | 38,828 | 5,82 | 39911 | 2 | 19,12 | 18,42 | 19,55 | 21,72 | 21,00 | 21,58 | **5,294** | 3,74E-03 | 1 | 2,40433884 | 2,42703974 |
| QBK84323.1 | | tegument serinethreonine protein kinase [Human betaherpesvirus 5] | 21,016 | 4,67 | 78246 | 2 | 18,23 | 18,93 | 18,11 | 21,71 | 19,26 | 21,32 | **5,060** | 4,35E-02 | 1 | 2,33913294 | 1,36133746 |
| QBK84327.1 | UL35 | tegument protein UL35 [Human betaherpesvirus 5] | 30,497 | 2,5 | 72630 | 2 | 18,16 | 18,34 | 18,14 | 20,84 | 19,91 | 20,78 | **4,917** | 1,70E-03 | 1 | 2,29780388 | 2,76861861 |
| QBK84448.1 | | small capsid protein [Human betaherpesvirus 5] | 28,467 | 17,33 | 8481 | 1 | 17,60 | 18,03 | 18,55 | 21,54 | 20,13 | 19,39 | **4,895** | 2,91E-02 | 1 | 2,29145241 | 1,53660276 |
| RBM25_HUM | RBM25 | sp\|P49756\|RBM25_HUMAN RNA-binding protein 25 OS=Homo sapiens GN=RBM25 PE= | 108,628 | 7,12 | 100185 | 3 | 19,52 | 21,60 | 20,01 | 22,36 | 22,15 | 22,74 | **4,112** | 3,51E-02 | 1 | 2,03999964 | 1,45492467 |
| QBK84321.1 | UL112 | protein UL112 [Human betaherpesvirus 5] | 185,796 | 13,01 | 70314 | 6 | 22,44 | 22,50 | 22,98 | 23,99 | 23,91 | 24,97 | **3,138** | 1,24E-02 | 1 | 1,64999962 | 1,90770786 |
| CSTF3_HUM | CSTF3 | sp\|Q12996\|CSTF3_HUMAN Cleavage stimulation factor subunit 3 OS=Homo sapiens GN | 51,987 | 1,81 | 82922 | 1 | 18,61 | 18,49 | 18,42 | 19,29 | 20,01 | 21,10 | **3,084** | 3,73E-02 | 1 | 1,62465223 | 1,42866984 |
| PVR_HUMAN | PVR | sp\|P15151\|PVR_HUMAN Poliovirus receptor OS=Homo sapiens GN=PVR PE=1 SV=2 | 14,853 | 3,12 | 45303 | 1 | 19,09 | 19,09 | 19,33 | 21,57 | 20,44 | 19,96 | **2,808** | 3,70E-02 | 1 | 1,48936145 | 1,43195119 |
| UBP2L_HUM | UBAP2L | sp\|Q14157\|UBP2L_HUMAN Ubiquitin-associated protein 2-like OS=Homo sapiens GN=U | 75,671 | 3,5 | 114535 | 3 | 20,98 | 20,42 | 20,32 | 22,23 | 21,95 | 21,69 | **2,609** | 5,83E-03 | 1 | 1,38333384 | 2,23451099 |
| SRP68_HUM | SRP68 | sp\|Q9UHB9\|SRP68_HUMAN Signal recognition particle subunit SRP68 OS=Homo sapiens | 110,109 | 6,38 | 70730 | 3 | 22,27 | 21,82 | 22,07 | 23,24 | 23,30 | 22,94 | **2,153** | 2,96E-03 | 1 | 1,10666656 | 2,52923769 |
| PSME3_HUM | PSME3 | sp\|P61289\|PSME3_HUMAN Proteasome activator complex subunit 3 OS=Homo sapiens | 127,72 | 20,87 | 29506 | 5 | 22,88 | 22,21 | 23,05 | 23,69 | 23,88 | 23,88 | **2,149** | 1,39E-02 | 1 | 1,10333379 | 1,85554107 |
| RCC1_HUMA | RCC1 | sp\|P18754\|RCC1_HUMAN Regulator of chromosome condensation OS=Homo sapiens G | 93,937 | 7,84 | 44969 | 3 | 21,94 | 22,45 | 21,79 | 22,78 | 23,35 | 23,30 | **2,119** | 1,60E-02 | 1 | 1,0833327 | 1,79472761 |
| UBP15_HUM | USP15 | sp\|Q9Y4E8\|UBP15_HUMAN Ubiquitin carboxyl-terminal hydrolase 15 OS=Homo sapiens | 124,631 | 7,44 | 112419 | 6 | 21,70 | 21,11 | 20,77 | 22,04 | 22,33 | 22,40 | **2,090** | 2,22E-02 | 1 | 1,06333288 | 1,65322933 |
| PDCL3_HUM | PDCL3 | sp\|Q9H2J4\|PDCL3_HUMAN Phosducin-like protein 3 OS=Homo sapiens GN=PDCL3 PE=1 | 68,682 | 10,46 | 27614 | 2 | 20,09 | 20,64 | 20,76 | 21,36 | 21,78 | 21,45 | **2,047** | 1,31E-02 | 1 | 1,0333341 | 1,88419671 |
| NRP2_HUMA | NRP2 | sp\|O60462\|NRP2_HUMAN Neuropilin-2 OS=Homo sapiens GN=NRP2 PE=1 SV=2 | 214,344 | 8,7 | 104859 | 6 | 24,59 | 24,40 | 24,09 | 23,80 | 23,38 | 22,88 | **0,498** | 2,94E-02 | 1 | -1,0066675 | 1,53192398 |
| ATPO_HUMA | ATP5O | sp\|P48047\|ATPO_HUMAN ATP synthase subunit O, mitochondrial OS=Homo sapiens GN | 109,129 | 18,31 | 23277 | 4 | 24,62 | 24,76 | 25,16 | 23,66 | 23,77 | 24,07 | **0,495** | 7,53E-03 | 1 | -1,0133336 | 2,12334619 |
| AL4A1_HUM | ALDH4A1 | sp\|P30038\|AL4A1_HUMAN Delta-1-pyrroline-5-carboxylate dehydrogenase, mitochond | 136,565 | 11,55 | 61719 | 4 | 22,57 | 22,79 | 22,23 | 21,35 | 21,50 | 21,50 | **0,473** | 3,17E-03 | 1 | -1,0799999 | 2,49858519 |
| MTCH2_HUM | MTCH2 | sp\|Q9Y6C9\|MTCH2_HUMAN Mitochondrial carrier homolog 2 OS=Homo sapiens GN=M | 204,411 | 17,49 | 33331 | 5 | 24,01 | 24,66 | 24,27 | 23,19 | 23,61 | 22,81 | **0,463** | 2,05E-02 | 1 | -1,11 | 1,68862157 |
| SDHB_HUMA | SDHB | sp\|P21912\|SDHB_HUMAN Succinate dehydrogenase [ubiquinone] iron-sulfur subunit, n | 58,378 | 10,71 | 31630 | 3 | 22,41 | 22,79 | 22,89 | 21,23 | 22,13 | 21,21 | **0,443** | 2,53E-02 | 1 | -1,1733341 | 1,59754874 |
| PTPRK_HUM | PTPRK | sp\|Q15262\|PTPRK_HUMAN Receptor-type tyrosine-protein phosphatase kappa OS=Ho | 40,462 | 1,67 | 162102 | 2 | 20,90 | 20,40 | 20,17 | 19,63 | 19,26 | 19,00 | **0,437** | 1,35E-02 | 1 | -1,1935031 | 1,87124089 |
| AT5F1_HUM | ATP5F1 | sp\|P24539\|AT5F1_HUMAN ATP synthase F(0) complex subunit B1, mitochondrial OS=Ho | 32,425 | 7,81 | 28909 | 2 | 22,94 | 23,38 | 23,90 | 21,81 | 22,14 | 22,56 | **0,424** | 2,47E-02 | 1 | -1,236667 | 1,60811108 |
| ODO1_HUM | OGDH | sp\|Q02218\|ODO1_HUMAN 2-oxoglutarate dehydrogenase, mitochondrial OS=Homo sa | 195,992 | 10,07 | 115935 | 9 | 22,67 | 23,06 | 23,78 | 21,58 | 22,00 | 22,16 | **0,419** | 2,70E-02 | 1 | -1,2566668 | 1,56919295 |
| CMC2_HUM | SLC25A13 | sp\|Q9UJS0\|CMC2_HUMAN Calcium-binding mitochondrial carrier protein Aralar2 OS=H | 81,702 | 5,19 | 74176 | 3 | 21,96 | 22,17 | 23,27 | 21,15 | 20,93 | 21,51 | **0,415** | 4,47E-02 | 1 | -1,2699998 | 1,34947493 |
| SAM50_HUM | SAMM50 | sp\|Q9Y512\|SAM50_HUMAN Sorting and assembly machinery component 50 homolog C | 136,656 | 11,94 | 51976 | 5 | 23,16 | 23,59 | 23,97 | 22,00 | 22,79 | 22,05 | **0,408** | 2,02E-02 | 1 | -1,2933331 | 1,69403537 |
| ADA15_HUM | ADAM15 | sp\|Q13444\|ADA15_HUMAN Disintegrin and metalloproteinase domain-containing prote | 24,013 | 2,09 | 92959 | 1 | 18,27 | 19,21 | 19,48 | 17,77 | 18,00 | 16,99 | **0,379** | 4,27E-02 | 1 | -1,4006589 | 1,36952267 |
| CLUA1_HUM | CLUAP1 | sp\|Q96AJ1\|CLUA1_HUMAN Clusterin-associated protein 1 OS=Homo sapiens GN=CLUA | 56,291 | 3,63 | 48125 | 1 | 19,33 | 19,95 | 21,56 | 18,39 | 18,44 | 17,85 | **0,241** | 4,10E-02 | 1 | -2,0535723 | 1,38717894 |
| FLRT2_HUM | FLRT2 | sp\|O43155\|FLRT2_HUMAN Leucine-rich repeat transmembrane protein FLRT2 OS=Hom | 25,028 | 3,03 | 74049 | 2 | 24,18 | 24,52 | 22,91 | 18,84 | 21,32 | 21,53 | **0,101** | 2,90E-02 | 1 | -3,3054365 | 1,5368732 |
| ACSF2_HUM | ACSF2 | sp\|Q96CM8\|ACSF2_HUMAN Acyl-CoA synthetase family member 2, mitochondrial OS=H | 37,637 | 5,53 | 68125 | 2 | 22,16 | 21,48 | 22,84 | 18,04 | 19,67 | 18,55 | **0,094** | 5,38E-03 | 1 | -3,4066664 | 2,26888702 |
| RAP2A_HUM | RAP2A | sp\|P10114\|RAP2A_HUMAN Ras-related protein Rap-2a OS=Homo sapiens GN=RAP2A PE | 406,51 | 43,17 | 20615 | 7 | 23,51 | 22,30 | 23,53 | 19,84 | 18,02 | 19,19 | **0,059** | 3,65E-03 | 1 | -4,0953045 | 2,43799416 |
| GAK_HUMAN | GAK | sp\|O14976\|GAK_HUMAN Cyclin-G-associated kinase OS=Homo sapiens GN=GAK PE=1 S | 10,083 | 0,69 | 143191 | 1 | 17,31 | 19,18 | 21,40 | 21,74 | 21,39 | 25,42 | **11,739** | 1,12E-01 | 0 | 3,55326589 | 0,9507263 |
| K2C4_HUMA | KRT4 | sp\|P19013\|K2C4_HUMAN Keratin, type II cytoskeletal 4 OS=Homo sapiens GN=KRT4 PE | 767,145 | 43,63 | 57285 | 24 | 17,80 | 21,50 | 19,27 | 20,02 | 21,24 | 27,68 | **10,974** | 2,56E-01 | 0 | 3,4560159 | 0,59211461 |
| DCD_HUMAN | DCD | sp\|P81605\|DCD_HUMAN Dermcidin OS=Homo sapiens GN=DCD PE=1 SV=2 | 75,119 | 20 | 11284 | 2 | 18,55 | 18,08 | 22,95 | 23,05 | 22,35 | 23,45 | **8,523** | 1,23E-01 | 0 | 3,09135183 | 0,91077525 |
| ZG16B_HUM | ZG16B | sp\|Q96DA0\|ZG16B_HUMAN Zymogen granule protein 16 homolog B OS=Homo sapiens | 24,763 | 4,33 | 22739 | 1 | 18,58 | 17,20 | 18,07 | 21,29 | 18,06 | 22,61 | **6,509** | 1,28E-01 | 0 | 2,70236206 | 0,89289859 |
| TOP2B_HUM | TOP2B | sp\|Q02880\|TOP2B_HUMAN DNA topoisomerase 2-beta OS=Homo sapiens GN=TOP2B P | 127,448 | 3,44 | 183267 | 5 | 18,05 | 16,79 | 21,68 | 20,37 | 20,69 | 22,84 | **5,502** | 2,12E-01 | 0 | 2,4600811 | 0,67341958 |
| PRC2A_HUM | PRRC2A | sp\|P48634\|PRC2A_HUMAN Protein PRRC2A OS=Homo sapiens GN=PRRC2A PE=1 SV=3 | 9,256 | 0,32 | 228863 | 1 | 18,74 | 17,65 | 19,30 | 19,11 | 17,66 | 25,88 | **4,999** | 4,19E-01 | 0 | 2,32161204 | 0,37795587 |
| SMRC2_HUM | SMARCC2 | sp\|Q8TAQ2\|SMRC2_HUMAN SWI/SNF complex subunit SMARCC2 OS=Homo sapiens GN | 173,827 | 6,84 | 132879 | 7 | 17,40 | 19,55 | 17,41 | 21,09 | 19,68 | 20,41 | **4,831** | 5,07E-02 | 0 | 2,27240372 | 1,29512489 |
| EI24_HUMAN | EI24 | sp\|O14681\|EI24_HUMAN Etoposide-induced protein 2.4 homolog OS=Homo sapiens GN | 11,45 | 2,35 | 38965 | 1 | 24,00 | 20,54 | 18,68 | 23,92 | 22,64 | 23,30 | **4,634** | 2,39E-01 | 0 | 2,21234512 | 0,62118945 |
| K1C13_HUM | KRT13 | sp\|P13646\|K1C13_HUMAN Keratin, type I cytoskeletal 13 OS=Homo sapiens GN=KRT13 | 653,829 | 33,62 | 49588 | 18 | 20,85 | 20,94 | 20,97 | 19,50 | 22,27 | 27,57 | **4,574** | 4,07E-01 | 0 | 2,19333331 | 0,39072638 |
| SPRY7_HUM | SPRYD7 | sp\|Q5W111\|SPRY7_HUMAN SPRY domain-containing protein 7 OS=Homo sapiens GN=S | 13,719 | 5,1 | 21666 | 1 | 18,36 | 17,19 | 19,22 | 21,50 | 21,60 | 18,14 | **4,458** | 1,68E-01 | 0 | 2,15630277 | 0,77502293 |
| SYF1_HUMA | XAB2 | sp\|Q9HCS7\|SYF1_HUMAN Pre-mRNA-splicing factor SYF1 OS=Homo sapiens GN=XAB2 P | 163,918 | 8,89 | 100010 | 6 | 18,97 | 16,64 | 21,65 | 20,57 | 20,13 | 23,02 | **4,446** | 2,75E-01 | 0 | 2,15266355 | 0,5607288 |
| NOG1_HUM | GTPBP4 | sp\|Q9BZE4\|NOG1_HUMAN Nucleolar GTP-binding protein 1 OS=Homo sapiens GN=GTP | 193,768 | 14,04 | 73964 | 9 | 20,67 | 18,59 | 24,39 | 21,71 | 22,44 | 25,88 | **4,371** | 3,74E-01 | 0 | 2,12804222 | 0,4269039 |
| PRPC_HUMA | PRH1 | sp\|P02810\|PRPC_HUMAN Salivary acidic proline-rich phosphoprotein 1/2 OS=Homo sap | 64,262 | 19,28 | 17016 | 2 | 20,78 | 19,28 | 17,96 | 24,74 | 19,76 | 19,76 | **4,223** | 3,24E-01 | 0 | 2,07837041 | 0,48976307 |
| KBTB3_HUM | KBTBD3 | sp\|Q8NAB2\|KBTB3_HUMAN Kelch repeat and BTB domain-containing protein 3 OS=Hom | 21,019 | 1,15 | 69395 | 1 | 24,26 | 24,52 | 18,22 | 24,19 | 24,78 | 24,08 | **4,047** | 3,85E-01 | 0 | 2,01700592 | 0,41454466 |
| ICLN_HUMA | CLNS1A | sp\|P54105\|ICLN_HUMAN Methylosome subunit pICln OS=Homo sapiens GN=CLNS1A PE | 43,765 | 5,49 | 26215 | 1 | 17,91 | 20,16 | 21,96 | 23,15 | 20,18 | 22,65 | **3,958** | 2,53E-01 | 0 | 1,98476537 | 0,59619331 |
| PLPP_HUMA | PDXP | sp\|Q96GD0\|PLPP_HUMAN Pyridoxal phosphate phosphatase OS=Homo sapiens GN=PD | 20,319 | 3,38 | 31698 | 1 | 17,97 | 17,46 | 20,04 | 21,57 | 21,75 | 18,09 | **3,947** | 2,38E-01 | 0 | 1,98078156 | 0,62324431 |
| RALB_HUMA | RALB | sp\|P11234\|RALB_HUMAN Ras-related protein Ral-B OS=Homo sapiens GN=RALB PE=1 S | 540,388 | 48,06 | 23409 | 13 | 18,43 | 18,78 | 23,51 | 21,92 | 21,24 | 23,44 | **3,897** | 3,28E-01 | 0 | 1,96236928 | 0,48388465 |
| PTMS_HUMA | PTMS | sp\|P20962\|PTMS_HUMAN Parathymosin OS=Homo sapiens GN=PTMS PE=1 SV=2 | 24,364 | 10,78 | 11530 | 1 | 18,97 | 24,37 | 22,89 | 24,26 | 24,21 | 23,64 | **3,888** | 2,94E-01 | 0 | 1,95906639 | 0,53203982 |
| K2C78_HUM | KRT78 | sp\|Q8N1N4\|K2C78_HUMAN Keratin, type II cytoskeletal 78 OS=Homo sapiens GN=KRT7 | 188,524 | 15,19 | 56866 | 7 | 22,54 | 20,72 | 20,10 | 24,65 | 22,17 | 22,28 | **3,767** | 1,54E-01 | 0 | 1,91333326 | 0,8114227 |
| 1433S_HUM | SFN | sp\|P31947\|1433S_HUMAN 14-3-3 protein sigma OS=Homo sapiens GN=SFN PE=1 SV=1 | 104,615 | 10,48 | 27774 | 4 | 22,90 | 19,38 | 26,79 | 25,04 | 22,99 | 26,77 | **3,758** | 4,71E-01 | 0 | 1,910127 | 0,32684724 |
| TAOK3_HUM | TAOK3 | sp\|Q9H2K8\|TAOK3_HUMAN Serine/threonine-protein kinase TAO3 OS=Homo sapiens G | 76,881 | 5,01 | 105406 | 5 | 21,73 | 21,24 | 20,37 | 21,97 | 21,23 | 25,81 | **3,706** | 2,69E-01 | 0 | 1,88999939 | 0,57012897 |
| TMED1_HUM | TMED1 | sp\|Q13445\|TMED1_HUMAN Transmembrane emp24 domain-containing protein 1 OS=H | 52,872 | 9,69 | 25206 | 2 | 17,49 | 20,92 | 23,08 | 21,78 | 22,08 | 23,26 | **3,669** | 3,29E-01 | 0 | 1,8753891 | 0,4830444 |
| OSGEP_HUM | OSGEP | sp\|Q9NPF4\|OSGEP_HUMAN Probable tRNA N6-adenosine threonylcarbamoyltransferas | 26,188 | 4,78 | 36427 | 1 | 18,35 | 19,84 | 16,86 | 20,20 | 20,08 | 20,34 | **3,621** | 9,81E-02 | 0 | 1,85645485 | 1,00840514 |
| RER1_HUMA | RER1 | sp\|O15258\|RER1_HUMAN Protein RER1 OS=Homo sapiens GN=RER1 PE=1 SV=1 | 75,641 | 9,18 | 22958 | 1 | 18,31 | 17,95 | 22,26 | 20,62 | 21,15 | 22,31 | **3,609** | 2,75E-01 | 0 | 1,85174815 | 0,5598892 |
| CATZ_HUMA | CTSZ | sp\|Q9UBR2\|CATZ_HUMAN Cathepsin Z OS=Homo sapiens GN=CTSZ PE=1 SV=1 | 13,265 | 3,3 | 33868 | 1 | 18,29 | 18,39 | 22,62 | 21,53 | 21,27 | 22,04 | **3,596** | 2,70E-01 | 0 | 1,84645398 | 0,56809507 |
| AMY1_HUM | AMY1A | sp\|P04745\|AMY1_HUMAN Alpha-amylase 1 OS=Homo sapiens GN=AMY1A PE=1 SV=2 | 537,526 | 43,05 | 57768 | 15 | 22,69 | 19,49 | 22,75 | 26,82 | 21,10 | 22,51 | **3,564** | 4,17E-01 | 0 | 1,83333333 | 0,37937714 |
| LAT1_HUMA | SLC7A5 | sp\|Q01650\|LAT1_HUMAN Large neutral amino acids transporter small subunit 1 OS=Ho | 187,587 | 13,41 | 55010 | 4 | 20,57 | 21,22 | 18,82 | 22,96 | 22,06 | 21,01 | **3,502** | 1,19E-01 | 0 | 1,80807177 | 0,92566108 |
| PDS5B_HUM | PDS5B | sp\|Q9NTI5\|PDS5B_HUMAN Sister chromatid cohesion protein PDS5 homolog B OS=Hom | 289,573 | 10,92 | 164667 | 10 | 20,82 | 21,82 | 24,20 | 23,49 | 23,94 | 24,81 | **3,482** | 1,69E-01 | 0 | 1,79999987 | 0,77140616 |
| HMGB3_HU | HMGB3 | sp\|O15347\|HMGB3_HUMAN High mobility group protein B3 OS=Homo sapiens GN=HM | 14,102 | 4 | 22980 | 1 | 17,43 | 17,72 | 21,78 | 20,85 | 19,51 | 21,92 | **3,442** | 3,19E-01 | 0 | 1,78310966 | 0,49638056 |
| AT12A_HUM | ATP12A | sp\|P54707\|AT12A_HUMAN Potassium-transporting ATPase alpha chain 2 OS=Homo sapi | 206,78 | 3,56 | 115511 | 6 | 19,20 | 20,64 | 25,81 | 23,32 | 21,50 | 26,17 | **3,436** | 5,03E-01 | 0 | 1,78068097 | 0,29814214 |
| COMD2_HU | COMMD2 | sp\|Q86X83\|COMD2_HUMAN COMM domain-containing protein 2 OS=Homo sapiens GN | 14,207 | 3,52 | 22745 | 1 | 22,94 | 20,91 | 17,87 | 22,62 | 22,96 | 21,42 | **3,390** | 3,18E-01 | 0 | 1,76147143 | 0,49707669 |
| BASP1_HUM | BASP1 | sp\|P80723\|BASP1_HUMAN Brain acid soluble protein 1 OS=Homo sapiens GN=BASP1 PE | 184,537 | 27,75 | 22693 | 5 | 24,52 | 20,52 | 24,18 | 26,13 | 24,65 | 23,71 | **3,379** | 2,96E-01 | 0 | 1,75666555 | 0,52924063 |
| RAB24_HUM | RAB24 | sp\|Q969Q5\|RAB24_HUMAN Ras-related protein Rab-24 OS=Homo sapiens GN=RAB24 P | 138,09 | 18,23 | 23124 | 3 | 17,63 | 21,75 | 23,14 | 22,02 | 23,13 | 22,59 | **3,340** | 3,60E-01 | 0 | 1,73974164 | 0,44351123 |
| TGM3_HUM | TGM3 | sp\|Q08188\|TGM3_HUMAN Protein-glutamine gamma-glutamyltransferase E OS=Homo | 75,503 | 4,47 | 76632 | 3 | 21,08 | 17,16 | 20,74 | 22,76 | 20,35 | 21,04 | **3,302** | 2,99E-01 | 0 | 1,72324498 | 0,52478527 |
| MYO6_HUM | MYO6 | sp\|Q9UM54\|MYO6_HUMAN Unconventional myosin-VI OS=Homo sapiens GN=MYO6 P | 78,392 | 3,94 | 149691 | 5 | 18,34 | 20,82 | 20,56 | 21,18 | 22,18 | 21,50 | **3,278** | 1,11E-01 | 0 | 1,71295166 | 0,95453158 |
| XPO4_HUMA | XPO4 | sp\|Q9C0E2\|XPO4_HUMAN Exportin-4 OS=Homo sapiens GN=XPO4 PE=1 SV=2 | 26,513 | 0,61 | 130139 | 1 | 18,45 | 21,27 | 19,55 | 21,04 | 21,44 | 21,89 | **3,248** | 1,18E-01 | 0 | 1,69974264 | 0,92843162 |
| DPP9_HUMA | DPP9 | sp\|Q86TI2\|DPP9_HUMAN Dipeptidyl peptidase 9 OS=Homo sapiens GN=DPP9 PE=1 SV= | 32,535 | 1,27 | 98263 | 1 | 21,44 | 21,32 | 17,57 | 21,51 | 21,27 | 22,59 | **3,206** | 2,76E-01 | 0 | 1,68092155 | 0,55855101 |
| B3AT_HUMA | SLC4A1 | sp\|P02730\|B3AT_HUMAN Band 3 anion transport protein OS=Homo sapiens GN=SLC4A | 17,975 | 0,99 | 101792 | 1 | 24,80 | 17,45 | 22,58 | 24,69 | 22,80 | 22,31 | **3,155** | 5,10E-01 | 0 | 1,65776507 | 0,29234258 |
| PLPHP_HUM | PLPBP | sp\|O94903\|PLPHP_HUMAN Pyridoxal phosphate homeostasis protein OS=Homo sapiens | 48,386 | 4,73 | 30344 | 1 | 20,66 | 21,36 | 17,03 | 21,20 | 21,44 | 21,38 | **3,152** | 2,85E-01 | 0 | 1,65631294 | 0,54491654 |
| CBSL_HUMA | CBSL | sp\|P0DN79\|CBSL_HUMAN Cystathionine beta-synthase-like protein OS=Homo sapiens G | 16,925 | 2,36 | 60587 | 1 | 17,65 | 18,43 | 20,04 | 17,41 | 22,50 | 21,16 | **3,143** | 3,81E-01 | 0 | 1,65224457 | 0,41951955 |
| INT11_HUM | INTS11 | sp\|Q5TA45\|INT11_HUMAN Integrator complex subunit 11 OS=Homo sapiens GN=INTS1 | 31,653 | 1,67 | 67663 | 1 | 17,23 | 18,62 | 21,20 | 19,90 | 20,33 | 21,74 | **3,119** | 2,72E-01 | 0 | 1,64096006 | 0,56515661 |
| ERG7_HUMA | LSS | sp\|P48449\|ERG7_HUMAN Lanosterol synthase OS=Homo sapiens GN=LSS PE=1 SV=1 | 55,891 | 3,42 | 83309 | 2 | 19,98 | 18,43 | 22,92 | 21,42 | 21,49 | 23,27 | **3,068** | 3,27E-01 | 0 | 1,61732419 | 0,48537191 |
| TSN3_HUMA | TSPAN3 | sp\|O60637\|TSN3_HUMAN Tetraspanin-3 OS=Homo sapiens GN=TSPAN3 PE=2 SV=1 | 16,128 | 2,77 | 28018 | 1 | 23,77 | 17,86 | 23,81 | 24,05 | 22,87 | 23,36 | **3,062** | 4,66E-01 | 0 | 1,61449496 | 0,3313429 |
| NUP54_HUM | NUP54 | sp\|Q7Z3B4\|NUP54_HUMAN Nucleoporin p54 OS=Homo sapiens GN=NUP54 PE=1 SV=2 | 40,31 | 4,54 | 55435 | 2 | 18,24 | 18,50 | 21,59 | 21,56 | 19,21 | 22,38 | **3,055** | 3,24E-01 | 0 | 1,61137962 | 0,48882993 |
| SEC62_HUM | SEC62 | sp\|Q99442\|SEC62_HUMAN Translocation protein SEC62 OS=Homo sapiens GN=SEC62 P | 68,447 | 8,02 | 45862 | 4 | 23,49 | 18,76 | 22,05 | 23,67 | 22,72 | 22,72 | **3,038** | 3,26E-01 | 0 | 1,60333316 | 0,4861222 |
| SCAM1_HUM | SCAMP1 | sp\|O15126\|SCAM1_HUMAN Secretory carrier-associated membrane protein 1 OS=Hom | 20,919 | 3,85 | 37920 | 1 | 18,27 | 18,16 | 18,99 | 20,95 | 20,70 | 18,53 | **2,998** | 1,22E-01 | 0 | 1,58413378 | 0,91196969 |
| THUM3_HUM | THUMPD3 | sp\|Q9BV44\|THUM3_HUMAN THUMP domain-containing protein 3 OS=Homo sapiens G | 10,887 | 2,17 | 57003 | 1 | 17,80 | 21,15 | 17,65 | 21,57 | 21,30 | 18,38 | **2,921** | 3,70E-01 | 0 | 1,54668808 | 0,43200329 |
| DCUP_HUMA | UROD | sp\|P06132\|DCUP_HUMAN Uroporphyrinogen decarboxylase OS=Homo sapiens GN=UR | 58,799 | 5,72 | 40787 | 2 | 21,59 | 20,91 | 19,02 | 22,23 | 22,21 | 21,69 | **2,898** | 1,23E-01 | 0 | 1,53527069 | 0,90994904 |
| KIF11_HUMA | KIF11 | sp\|P52732\|KIF11_HUMAN Kinesin-like protein KIF11 OS=Homo sapiens GN=KIF11 PE=1 | 38,441 | 0,95 | 119159 | 1 | 18,37 | 17,73 | 21,15 | 20,05 | 20,01 | 21,77 | **2,880** | 2,72E-01 | 0 | 1,52616819 | 0,56540093 |
| PP4R1_HUM | PPP4R1 | sp\|Q8TF05\|PP4R1_HUMAN Serine/threonine-protein phosphatase 4 regulatory subunit | 65,954 | 3,89 | 107004 | 3 | 19,84 | 19,16 | 22,18 | 21,54 | 21,43 | 22,78 | **2,875** | 2,07E-01 | 0 | 1,52333387 | 0,68472462 |
| AAMP_HUM | AAMP | sp\|Q13685\|AAMP_HUMAN Angio-associated migratory cell protein OS=Homo sapiens G | 59,131 | 3 | 46751 | 1 | 16,40 | 19,25 | 20,44 | 20,14 | 19,55 | 20,95 | **2,859** | 2,97E-01 | 0 | 1,51535988 | 0,52738026 |
| SPTC1_HUM | SPTLC1 | sp\|O15269\|SPTC1_HUMAN Serine palmitoyltransferase 1 OS=Homo sapiens GN=SPTLC1 | 89,65 | 5,71 | 52744 | 2 | 20,73 | 19,57 | 19,74 | 20,96 | 21,06 | 22,56 | **2,855** | 7,46E-02 | 0 | 1,513333 | 1,1272787 |
| FBLL1_HUM | FBLL1 | sp\|A6NHQ2\|FBLL1_HUMAN rRNA/tRNA 2'-O-methyltransferase fibrillarin-like protein 1 | 39,845 | 6,59 | 34803 | 2 | 18,18 | 21,93 | 23,91 | 22,00 | 22,30 | 24,24 | **2,838** | 4,55E-01 | 0 | 1,50509262 | 0,34233211 |
| ARL15_HUM | ARL15 | sp\|Q9NXU5\|ARL15_HUMAN ADP-ribosylation factor-like protein 15 OS=Homo sapiens G | 38,106 | 10,29 | 22876 | 2 | 22,64 | 23,14 | 17,78 | 23,37 | 23,02 | 21,68 | **2,835** | 4,47E-01 | 0 | 1,50352097 | 0,34954531 |
| LZIC_HUMAN | LZIC | sp\|Q8WZA0\|LZIC_HUMAN Protein LZIC OS=Homo sapiens GN=LZIC PE=1 SV=1 | 69,887 | 18,95 | 21495 | 2 | 21,21 | 21,43 | 23,66 | 23,95 | 22,93 | 23,90 | **2,815** | 1,54E-01 | 0 | 1,49333382 | 0,81301435 |
| PRP4_HUMA | PRPF4 | sp\|O43172\|PRP4_HUMAN U4/U6 small nuclear ribonucleoprotein Prp4 OS=Homo sapie | 83,186 | 10,34 | 58449 | 4 | 18,45 | 21,09 | 22,78 | 21,69 | 21,98 | 23,11 | **2,803** | 3,27E-01 | 0 | 1,48710378 | 0,48545656 |
| NU205_HUM | NUP205 | sp\|Q92621\|NU205_HUMAN Nuclear pore complex protein Nup205 OS=Homo sapiens G | 292,034 | 5,96 | 227922 | 11 | 19,87 | 21,75 | 24,31 | 22,35 | 22,97 | 25,07 | **2,802** | 3,86E-01 | 0 | 1,48666636 | 0,41396421 |
| BLVRB_HUM | BLVRB | sp\|P30043\|BLVRB_HUMAN Flavin reductase (NADPH) OS=Homo sapiens GN=BLVRB PE= | 31,95 | 7,77 | 22119 | 2 | 24,64 | 24,43 | 18,11 | 24,63 | 24,64 | 22,34 | **2,781** | 5,52E-01 | 0 | 1,47551155 | 0,25822957 |
| FILA2_HUMA | FLG2 | sp\|Q5D862\|FILA2_HUMAN Filaggrin-2 OS=Homo sapiens GN=FLG2 PE=1 SV=1 | 319,692 | 7,49 | 248073 | 8 | 22,87 | 20,77 | 22,84 | 24,81 | 22,91 | 23,16 | **2,764** | 1,84E-01 | 0 | 1,4666659 | 0,73410915 |
| AMGO2_HU | AMIGO2 | sp\|Q86SJ2\|AMGO2_HUMAN Amphoterin-induced protein 2 OS=Homo sapiens GN=AMI | 19,501 | 1,53 | 57934 | 1 | 20,53 | 18,85 | 19,49 | 20,93 | 20,82 | 21,42 | **2,702** | 5,19E-02 | 0 | 1,43391101 | 1,28509102 |
| UBR5_HUMA | UBR5 | sp\|O95071\|UBR5_HUMAN E3 ubiquitin-protein ligase UBR5 OS=Homo sapiens GN=UBR | 42,168 | 0,82 | 309352 | 2 | 14,55 | 17,64 | 18,94 | 19,73 | 16,49 | 19,16 | **2,669** | 4,37E-01 | 0 | 1,41637389 | 0,35962162 |
| ZN169_HUM | ZNF169 | sp\|Q14929\|ZN169_HUMAN Zinc finger protein 169 OS=Homo sapiens GN=ZNF169 PE=1 | 15,473 | 2,65 | 68488 | 1 | 18,33 | 20,66 | 22,18 | 21,42 | 21,50 | 22,47 | **2,652** | 2,95E-01 | 0 | 1,40706698 | 0,5301811 |
| GLMN_HUM | GLMN | sp\|Q92990\|GLMN_HUMAN Glomulin OS=Homo sapiens GN=GLMN PE=1 SV=2 | 38,028 | 1,85 | 68208 | 1 | 19,15 | 17,25 | 21,01 | 20,46 | 20,86 | 20,31 | **2,652** | 2,69E-01 | 0 | 1,40686417 | 0,56975088 |
| NCLN_HUMA | NCLN | sp\|Q969V3\|NCLN_HUMAN Nicalin OS=Homo sapiens GN=NCLN PE=1 SV=2 | 98,977 | 9,06 | 62974 | 5 | 20,82 | 20,29 | 22,52 | 21,99 | 22,15 | 23,69 | **2,639** | 1,80E-01 | 0 | 1,39999962 | 0,74378157 |
| NALDL_HUM | NAALADL1 | sp\|Q9UQQ1\|NALDL_HUMAN N-acetylated-alpha-linked acidic dipeptidase-like protein C | 48,762 | 1,49 | 80558 | 1 | 18,73 | 23,67 | 24,44 | 23,17 | 23,85 | 24,01 | **2,635** | 4,83E-01 | 0 | 1,39759318 | 0,31631558 |
| NICA_HUMA | NCSTN | sp\|Q92542\|NICA_HUMAN Nicastrin OS=Homo sapiens GN=NCSTN PE=1 SV=2 | 106,403 | 7,33 | 78411 | 2 | 23,47 | 23,64 | 17,36 | 23,03 | 23,22 | 22,41 | **2,633** | 5,39E-01 | 0 | 1,39653524 | 0,26863858 |
| BMP15_HUM | BMP15 | sp\|O95972\|BMP15_HUMAN Bone morphogenetic protein 15 OS=Homo sapiens GN=BM | 18,119 | 1,79 | 45055 | 1 | 24,10 | 23,55 | 19,42 | 23,26 | 22,80 | 25,19 | **2,628** | 4,45E-01 | 0 | 1,39417521 | 0,35115352 |
| PELO_HUMA | PELO | sp\|Q9BRX2\|PELO_HUMAN Protein pelota homolog OS=Homo sapiens GN=PELO PE=1 S | 20,838 | 7,53 | 43359 | 2 | 20,91 | 17,60 | 18,41 | 21,00 | 20,02 | 20,02 | **2,591** | 2,60E-01 | 0 | 1,37333361 | 0,58441524 |
| K1C19_HUM | KRT19 | sp\|P08727\|K1C19_HUMAN Keratin, type I cytoskeletal 19 OS=Homo sapiens GN=KRT19 | 285,575 | 13,75 | 44106 | 9 | 20,93 | 20,33 | 19,47 | 19,03 | 20,83 | 24,98 | **2,585** | 4,92E-01 | 0 | 1,3700002 | 0,30832005 |
| CCNB1_HUM | CCNB1 | sp\|P14635\|CCNB1_HUMAN G2/mitotic-specific cyclin-B1 OS=Homo sapiens GN=CCNB1 | 23,009 | 3 | 48337 | 1 | 18,74 | 21,44 | 21,26 | 21,90 | 21,67 | 21,98 | **2,583** | 1,93E-01 | 0 | 1,36927795 | 0,71450031 |
| TOP2A_HUM | TOP2A | sp\|P11388\|TOP2A_HUMAN DNA topoisomerase 2-alpha OS=Homo sapiens GN=TOP2A | 385,917 | 10,39 | 174385 | 13 | 20,49 | 22,32 | 24,06 | 22,76 | 23,05 | 25,16 | **2,579** | 3,45E-01 | 0 | 1,36666679 | 0,46183843 |
| CTBL1_HUM | CTNNBL1 | sp\|Q8WYA6\|CTBL1_HUMAN Beta-catenin-like protein 1 OS=Homo sapiens GN=CTNNBL | 51,294 | 5,51 | 65173 | 3 | 20,33 | 20,18 | 21,26 | 21,51 | 21,47 | 22,88 | **2,573** | 7,62E-02 | 0 | 1,36333275 | 1,11791697 |
| SC61G_HUM | SEC61G | sp\|P60059\|SC61G_HUMAN Protein transport protein Sec61 subunit gamma OS=Homo s | 51,65 | 17,65 | 7741 | 1 | 18,22 | 20,01 | 19,13 | 20,63 | 21,14 | 19,65 | **2,553** | 1,16E-01 | 0 | 1,35225487 | 0,93530206 |
| OSTF1_HUM | OSTF1 | sp\|Q92882\|OSTF1_HUMAN Osteoclast-stimulating factor 1 OS=Homo sapiens GN=OSTF | 169,699 | 26,64 | 23787 | 4 | 24,16 | 24,36 | 22,24 | 24,26 | 27,66 | 22,86 | **2,532** | 4,43E-01 | 0 | 1,34000015 | 0,3531785 |
| SMRD2_HUM | SMARCD2 | sp\|Q92925\|SMRD2_HUMAN SWI/SNF-related matrix-associated actin-dependent regula | 23,497 | 2,64 | 58921 | 1 | 18,49 | 19,10 | 20,50 | 20,47 | 20,77 | 20,87 | **2,529** | 9,20E-02 | 0 | 1,33845329 | 1,03609882 |
| HMCS1_HUM | HMGCS1 | sp\|Q01581\|HMCS1_HUMAN Hydroxymethylglutaryl-CoA synthase, cytoplasmic OS=Hom | 158,96 | 12,5 | 57294 | 6 | 23,06 | 22,92 | 23,30 | 24,47 | 23,54 | 25,28 | **2,526** | 6,03E-02 | 0 | 1,33666738 | 1,21990884 |
| ANS1B_HUM | ANKS1B | sp\|Q7Z6G8\|ANS1B_HUMAN Ankyrin repeat and sterile alpha motif domain-containing p | 11,39 | 0,56 | 138066 | 1 | 17,56 | 17,39 | 23,86 | 20,38 | 18,17 | 24,25 | **2,515** | 6,56E-01 | 0 | 1,33075905 | 0,18282089 |
| DDX21_HUM | DDX21 | sp\|Q9NR30\|DDX21_HUMAN Nucleolar RNA helicase 2 OS=Homo sapiens GN=DDX21 PE | 188,75 | 9,58 | 87344 | 7 | 23,21 | 22,61 | 25,17 | 23,89 | 24,23 | 26,84 | **2,502** | 3,36E-01 | 0 | 1,3233331 | 0,47391684 |
| ICAM1_HUM | ICAM1 | sp\|P05362\|ICAM1_HUMAN Intercellular adhesion molecule 1 OS=Homo sapiens GN=IC | 41,272 | 2,63 | 57825 | 1 | 18,55 | 19,27 | 17,69 | 20,28 | 19,56 | 19,62 | **2,492** | 6,20E-02 | 0 | 1,31712596 | 1,20792948 |
| FACE1_HUM | ZMPSTE24 | sp\|O75844\|FACE1_HUMAN CAAX prenyl protease 1 homolog OS=Homo sapiens GN=ZM | 152,904 | 13,68 | 54813 | 4 | 18,42 | 20,28 | 22,43 | 19,77 | 21,80 | 23,50 | **2,485** | 4,53E-01 | 0 | 1,31333288 | 0,34362638 |
| MBB1A_HUM | MYBBP1A | sp\|Q9BQG0\|MBB1A_HUMAN Myb-binding protein 1A OS=Homo sapiens GN=MYBBP1A | 96,764 | 1,96 | 148855 | 2 | 20,82 | 21,45 | 24,05 | 22,66 | 22,82 | 24,76 | **2,474** | 3,36E-01 | 0 | 1,30666669 | 0,47323187 |
| RCN3_HUMA | RCN3 | sp\|Q96D15\|RCN3_HUMAN Reticulocalbin-3 OS=Homo sapiens GN=RCN3 PE=1 SV=1 | 53,378 | 9,15 | 37493 | 2 | 17,88 | 21,21 | 23,32 | 21,75 | 21,75 | 22,83 | **2,472** | 4,66E-01 | 0 | 1,30540975 | 0,33139135 |
| SYMPK_HUM | SYMPK | sp\|Q92797\|SYMPK_HUMAN Symplekin OS=Homo sapiens GN=SYMPK PE=1 SV=2 | 20,449 | 1,49 | 141148 | 2 | 17,40 | 19,02 | 19,66 | 19,20 | 19,62 | 21,17 | **2,470** | 2,21E-01 | 0 | 1,30475426 | 0,65555983 |
| S35E1_HUM | SLC35E1 | sp\|Q96K37\|S35E1_HUMAN Solute carrier family 35 member E1 OS=Homo sapiens GN=S | 46,653 | 2,93 | 44773 | 1 | 16,51 | 18,58 | 19,82 | 20,29 | 18,37 | 20,17 | **2,470** | 3,19E-01 | 0 | 1,3047231 | 0,4966301 |
| WFS1_HUM | WFS1 | sp\|O76024\|WFS1_HUMAN Wolframin OS=Homo sapiens GN=WFS1 PE=1 SV=2 | 12,048 | 0,9 | 100292 | 1 | 18,34 | 18,34 | 22,12 | 19,25 | 21,25 | 22,15 | **2,436** | 4,47E-01 | 0 | 1,28422483 | 0,34999027 |
| EVA1C_HUM | EVA1C | sp\|P58658\|EVA1C_HUMAN Protein eva-1 homolog C OS=Homo sapiens GN=EVA1C PE= | 20,881 | 2,72 | 49483 | 1 | 22,14 | 17,92 | 20,40 | 22,35 | 21,73 | 20,21 | **2,425** | 4,07E-01 | 0 | 1,2779452 | 0,3903617 |
| NFIA_HUMA | NFIA | sp\|Q12857\|NFIA_HUMAN Nuclear factor 1 A-type OS=Homo sapiens GN=NFIA PE=1 SV= | 23,206 | 2,16 | 55944 | 1 | 21,78 | 21,34 | 18,98 | 21,96 | 21,29 | 22,68 | **2,423** | 2,53E-01 | 0 | 1,27676773 | 0,59637237 |
| I5P1_HUMAN | INPP5A | sp\|Q14642\|I5P1_HUMAN Type I inositol 1,4,5-trisphosphate 5-phosphatase OS=Homo s | 77,91 | 12,62 | 47820 | 4 | 21,71 | 22,11 | 19,21 | 22,34 | 23,23 | 21,29 | **2,423** | 2,97E-01 | 0 | 1,27666728 | 0,5266279 |
| WDR3_HUM | WDR3 | sp\|Q9UNX4\|WDR3_HUMAN WD repeat-containing protein 3 OS=Homo sapiens GN=WD | 26,356 | 1,06 | 106099 | 1 | 18,89 | 18,70 | 18,61 | 18,78 | 19,97 | 21,26 | **2,412** | 1,53E-01 | 0 | 1,27048747 | 0,81642512 |
| CATA_HUMA | CAT | sp\|P04040\|CATA_HUMAN Catalase OS=Homo sapiens GN=CAT PE=1 SV=3 | 102,909 | 6,26 | 59756 | 3 | 18,70 | 18,66 | 20,34 | 21,58 | 18,85 | 21,08 | **2,412** | 2,75E-01 | 0 | 1,27019819 | 0,56080358 |
| PGRC2_HUM | PGRMC2 | sp\|O15173\|PGRC2_HUMAN Membrane-associated progesterone receptor component | 74,237 | 13,45 | 23818 | 3 | 21,60 | 22,60 | 23,93 | 23,89 | 24,11 | 23,94 | **2,412** | 1,34E-01 | 0 | 1,26999982 | 0,87160824 |
| MMP3_HUM | MMP3 | sp\|P08254\|MMP3_HUMAN Stromelysin-1 OS=Homo sapiens GN=MMP3 PE=1 SV=2 | 12,663 | 1,47 | 53977 | 1 | 22,94 | 22,95 | 18,75 | 23,47 | 23,63 | 21,34 | **2,404** | 4,68E-01 | 0 | 1,26550674 | 0,32968308 |
| AGRG1_HUM | ADGRG1 | sp\|Q9Y653\|AGRG1_HUMAN Adhesion G-protein coupled receptor G1 OS=Homo sapien | 29,991 | 3,17 | 77738 | 2 | 23,08 | 21,77 | 20,89 | 23,78 | 23,23 | 22,52 | **2,400** | 1,60E-01 | 0 | 1,26333364 | 0,79573958 |
| PTSS2_HUM | PTDSS2 | sp\|Q9BVG9\|PTSS2_HUMAN Phosphatidylserine synthase 2 OS=Homo sapiens GN=PTDS | 12,275 | 1,64 | 56253 | 1 | 19,22 | 21,20 | 19,08 | 21,12 | 21,15 | 21,02 | **2,397** | 1,39E-01 | 0 | 1,26103973 | 0,85617911 |
| VMP1_HUM | VMP1 | sp\|Q96GC9\|VMP1_HUMAN Vacuole membrane protein 1 OS=Homo sapiens GN=VMP1 | 13,901 | 1,72 | 46238 | 1 | 18,45 | 21,10 | 21,69 | 21,76 | 21,75 | 21,51 | **2,396** | 2,76E-01 | 0 | 1,26052793 | 0,558968 |
| HCFC1_HUM | HCFC1 | sp\|P51610\|HCFC1_HUMAN Host cell factor 1 OS=Homo sapiens GN=HCFC1 PE=1 SV=2 | 73,327 | 2,06 | 208732 | 4 | 19,22 | 20,83 | 21,55 | 21,30 | 21,46 | 22,62 | **2,395** | 1,92E-01 | 0 | 1,26000023 | 0,71576984 |
| APOM_HUM | APOM | sp\|O95445\|APOM_HUMAN Apolipoprotein M OS=Homo sapiens GN=APOM PE=1 SV=2 | 29,864 | 3,72 | 21253 | 1 | 19,74 | 23,48 | 25,72 | 23,98 | 23,42 | 25,32 | **2,393** | 5,30E-01 | 0 | 1,25909424 | 0,27593147 |
| RNPS1_HUM | RNPS1 | sp\|Q15287\|RNPS1_HUMAN RNA-binding protein with serine-rich domain 1 OS=Homo s | 42,106 | 4,92 | 34208 | 1 | 18,31 | 19,08 | 20,27 | 20,12 | 20,41 | 20,88 | **2,380** | 1,11E-01 | 0 | 1,25076103 | 0,95494791 |
| CORO6_HUM | CORO6 | sp\|Q6QEF8\|CORO6_HUMAN Coronin-6 OS=Homo sapiens GN=CORO6 PE=1 SV=2 | 73,695 | 4,66 | 52762 | 3 | 23,43 | 24,26 | 19,21 | 23,60 | 23,83 | 23,21 | **2,373** | 4,73E-01 | 0 | 1,24650955 | 0,32546134 |
| DICER_HUM | DICER1 | sp\|Q9UPY3\|DICER_HUMAN Endoribonuclease Dicer OS=Homo sapiens GN=DICER1 PE= | 23,497 | 1,04 | 218682 | 1 | 16,70 | 17,98 | 18,45 | 19,20 | 18,79 | 18,87 | **2,368** | 8,20E-02 | 0 | 1,24374135 | 1,08633441 |
| WDR5_HUM | WDR5 | sp\|P61964\|WDR5_HUMAN WD repeat-containing protein 5 OS=Homo sapiens GN=WDR | 75,781 | 19,16 | 36588 | 4 | 19,84 | 21,83 | 22,62 | 22,25 | 22,54 | 23,21 | **2,357** | 2,30E-01 | 0 | 1,23666636 | 0,6378134 |
| CHSP1_HUM | CARHSP1 | sp\|Q9Y2V2\|CHSP1_HUMAN Calcium-regulated heat-stable protein 1 OS=Homo sapiens | 27,912 | 10,88 | 15892 | 1 | 18,81 | 18,63 | 23,06 | 19,64 | 21,47 | 23,10 | **2,356** | 5,21E-01 | 0 | 1,2363205 | 0,28316845 |
| DSG1_HUMA | DSG1 | sp\|Q02413\|DSG1_HUMAN Desmoglein-1 OS=Homo sapiens GN=DSG1 PE=1 SV=2 | 129,325 | 7,91 | 113748 | 5 | 22,37 | 21,00 | 19,88 | 24,16 | 22,49 | 20,31 | **2,355** | 4,04E-01 | 0 | 1,23566373 | 0,39327474 |
| ZNT1_HUMA | SLC30A1 | sp\|Q9Y6M5\|ZNT1_HUMAN Zinc transporter 1 OS=Homo sapiens GN=SLC30A1 PE=1 SV= | 38,394 | 2,37 | 55300 | 1 | 21,87 | 21,87 | 18,04 | 21,38 | 22,32 | 21,77 | **2,347** | 3,99E-01 | 0 | 1,23075231 | 0,39857111 |
| KIFC1_HUM | KIFC1 | sp\|Q9BW19\|KIFC1_HUMAN Kinesin-like protein KIFC1 OS=Homo sapiens GN=KIFC1 PE= | 59,319 | 2,08 | 73748 | 1 | 17,65 | 20,04 | 20,72 | 20,30 | 20,10 | 21,68 | **2,337** | 3,11E-01 | 0 | 1,22478867 | 0,50753206 |
| GPSM1_HUM | GPSM1 | sp\|Q86YR5\|GPSM1_HUMAN G-protein-signaling modulator 1 OS=Homo sapiens GN=GP | 26,616 | 1,48 | 74510 | 1 | 20,65 | 17,39 | 20,78 | 20,83 | 20,54 | 21,12 | **2,333** | 3,37E-01 | 0 | 1,22237396 | 0,47281878 |
| AGFG2_HUM | AGFG2 | sp\|O95081\|AGFG2_HUMAN Arf-GAP domain and FG repeat-containing protein 2 OS=H | 47,459 | 2,49 | 48963 | 1 | 20,97 | 21,19 | 17,10 | 21,11 | 20,92 | 20,88 | **2,325** | 4,12E-01 | 0 | 1,217254 | 0,38503191 |
| SPTC2_HUM | SPTLC2 | sp\|O15270\|SPTC2_HUMAN Serine palmitoyltransferase 2 OS=Homo sapiens GN=SPTLC | 42,053 | 5,52 | 62924 | 2 | 20,04 | 19,16 | 18,05 | 20,08 | 20,54 | 20,27 | **2,319** | 1,09E-01 | 0 | 1,21362432 | 0,961115 |
| GRTP1_HUM | GRTP1 | sp\|Q5TC63\|GRTP1_HUMAN Growth hormone-regulated TBC protein 1 OS=Homo sapien | 22,64 | 2,08 | 38554 | 1 | 21,59 | 21,35 | 19,45 | 21,26 | 22,29 | 22,41 | **2,284** | 1,97E-01 | 0 | 1,19163895 | 0,70587336 |
| CYTA_HUMA | CSTA | sp\|P01040\|CYTA_HUMAN Cystatin-A OS=Homo sapiens GN=CSTA PE=1 SV=1 | 29,037 | 12,24 | 11006 | 1 | 20,60 | 17,95 | 20,93 | 21,93 | 20,08 | 21,03 | **2,276** | 3,35E-01 | 0 | 1,18638229 | 0,47482414 |
| PTCD1_HUM | PTCD1 | sp\|O75127\|PTCD1_HUMAN Pentatricopeptide repeat-containing protein 1, mitochondr | 10,625 | 2,29 | 78856 | 1 | 24,87 | 21,14 | 25,77 | 24,88 | 25,12 | 25,33 | **2,271** | 4,52E-01 | 0 | 1,18333308 | 0,3443848 |
| S29A1_HUM | SLC29A1 | sp\|Q99808\|S29A1_HUMAN Equilibrative nucleoside transporter 1 OS=Homo sapiens GN | 54,637 | 6,8 | 50219 | 3 | 21,85 | 21,01 | 22,51 | 25,06 | 21,35 | 22,50 | **2,266** | 3,74E-01 | 0 | 1,17999967 | 0,42756005 |
| AAAS_HUMA | AAAS | sp\|Q9NRG9\|AAAS_HUMAN Aladin OS=Homo sapiens GN=AAAS PE=1 SV=1 | 34,878 | 2,75 | 59574 | 1 | 16,84 | 20,22 | 21,31 | 18,42 | 21,79 | 21,67 | **2,250** | 5,38E-01 | 0 | 1,17019018 | 0,2690504 |
| FOCAD_HUM | FOCAD | sp\|Q5VW36\|FOCAD_HUMAN Focadhesin OS=Homo sapiens GN=FOCAD PE=1 SV=1 | 36,763 | 0,72 | 200072 | 1 | 18,80 | 17,95 | 18,42 | 19,17 | 19,10 | 20,38 | **2,241** | 7,32E-02 | 0 | 1,16443634 | 1,13549939 |
| UBE2A_HUM | UBE2A | sp\|P49459\|UBE2A_HUMAN Ubiquitin-conjugating enzyme E2 A OS=Homo sapiens GN=U | 45,391 | 17,76 | 17315 | 2 | 20,77 | 20,54 | 19,79 | 21,82 | 22,07 | 20,65 | **2,212** | 9,57E-02 | 0 | 1,14560699 | 1,01902409 |
| QBK84349.1 | | DNA polymerase processivity subunit [Human betaherpesvirus 5] | 92,416 | 8,08 | 46263 | 2 | 18,44 | 18,95 | 19,07 | 19,25 | 18,97 | 21,68 | **2,212** | 2,64E-01 | 0 | 1,1454366 | 0,57856027 |
| SPB12_HUM | SERPINB12 | sp\|Q96P63\|SPB12_HUMAN Serpin B12 OS=Homo sapiens GN=SERPINB12 PE=1 SV=1 | 105,347 | 10,62 | 46276 | 3 | 22,41 | 22,50 | 19,26 | 22,74 | 22,55 | 22,31 | **2,211** | 3,47E-01 | 0 | 1,14459991 | 0,46026644 |
| MBOA5_HUM | LPCAT3 | sp\|Q6P1A2\|MBOA5_HUMAN Lysophospholipid acyltransferase 5 OS=Homo sapiens GN | 57,275 | 2,67 | 56035 | 1 | 18,42 | 18,16 | 21,90 | 20,21 | 20,20 | 21,50 | **2,203** | 4,24E-01 | 0 | 1,1394062 | 0,37308452 |
| PRP31_HUM | PRPF31 | sp\|Q8WWY3\|PRP31_HUMAN U4/U6 small nuclear ribonucleoprotein Prp31 OS=Homo s | 64,124 | 8,02 | 55456 | 3 | 18,48 | 21,65 | 22,71 | 21,46 | 21,45 | 23,32 | **2,189** | 4,69E-01 | 0 | 1,12997182 | 0,32868052 |
| NP1L4_HUM | NAP1L4 | sp\|Q99733\|NP1L4_HUMAN Nucleosome assembly protein 1-like 4 OS=Homo sapiens G | 145,082 | 13,07 | 42823 | 4 | 20,40 | 20,82 | 21,36 | 21,30 | 22,48 | 22,18 | **2,184** | 6,65E-02 | 0 | 1,12666639 | 1,17693549 |
| CO1A2_HUM | COL1A2 | sp\|P08123\|CO1A2_HUMAN Collagen alpha-2(I) chain OS=Homo sapiens GN=COL1A2 PE | 74,354 | 1,61 | 129314 | 2 | 21,70 | 17,49 | 23,25 | 21,82 | 21,52 | 22,44 | **2,163** | 5,58E-01 | 0 | 1,11279678 | 0,25368186 |
| FILA_HUMAN | FLG | sp\|P20930\|FILA_HUMAN Filaggrin OS=Homo sapiens GN=FLG PE=1 SV=3 | 47,264 | 0,59 | 435170 | 4 | 21,06 | 17,58 | 21,05 | 22,30 | 19,98 | 20,72 | **2,149** | 4,58E-01 | 0 | 1,10333316 | 0,33894426 |
| DNJC9_HUM | DNAJC9 | sp\|Q8WXX5\|DNJC9_HUMAN DnaJ homolog subfamily C member 9 OS=Homo sapiens G | 58,85 | 7,31 | 29910 | 2 | 20,51 | 20,90 | 22,11 | 22,60 | 21,64 | 22,58 | **2,144** | 1,29E-01 | 0 | 1,09999975 | 0,88934094 |
| K1C16_HUM | KRT16 | sp\|P08779\|K1C16_HUMAN Keratin, type I cytoskeletal 16 OS=Homo sapiens GN=KRT16 | 1386,594 | 59,2 | 51268 | 31 | 25,91 | 24,58 | 21,69 | 27,53 | 25,09 | 22,83 | **2,129** | 5,86E-01 | 0 | 1,09000015 | 0,2322017 |
| K2C1B_HUM | KRT77 | sp\|Q7Z794\|K2C1B_HUMAN Keratin, type II cytoskeletal 1b OS=Homo sapiens GN=KRT7 | 286,412 | 13,19 | 61688 | 8 | 20,68 | 19,70 | 19,84 | 24,53 | 19,13 | 19,83 | **2,129** | 5,61E-01 | 0 | 1,08999952 | 0,25080321 |
| CHTOP_HUM | CHTOP | sp\|Q9Y3Y2\|CHTOP_HUMAN Chromatin target of PRMT1 protein OS=Homo sapiens GN= | 56,368 | 5,24 | 26397 | 1 | 18,44 | 19,60 | 17,83 | 19,59 | 19,49 | 20,06 | **2,128** | 1,17E-01 | 0 | 1,08931796 | 0,93045022 |
| NAA25_HUM | NAA25 | sp\|Q14CX7\|NAA25_HUMAN N-alpha-acetyltransferase 25, NatB auxiliary subunit OS=H | 43,273 | 1,95 | 112292 | 2 | 20,57 | 21,93 | 20,14 | 21,47 | 22,31 | 22,12 | **2,124** | 1,43E-01 | 0 | 1,08666674 | 0,84604435 |
| SRP54_HUM | SRP54 | sp\|P61011\|SRP54_HUMAN Signal recognition particle 54 kDa protein OS=Homo sapiens | 71,222 | 7,34 | 55705 | 3 | 19,33 | 22,59 | 22,74 | 22,37 | 22,22 | 23,33 | **2,122** | 4,04E-01 | 0 | 1,08568255 | 0,39361554 |
| CSTF1_HUM | CSTF1 | sp\|Q05048\|CSTF1_HUMAN Cleavage stimulation factor subunit 1 OS=Homo sapiens GN | 53 | 3,71 | 48358 | 1 | 19,55 | 16,60 | 19,24 | 18,46 | 19,60 | 20,57 | **2,116** | 3,87E-01 | 0 | 1,08100382 | 0,41178672 |
| TIM14_HUM | DNAJC19 | sp\|Q96DA6\|TIM14_HUMAN Mitochondrial import inner membrane translocase subunit | 62,178 | 16,38 | 12499 | 1 | 19,18 | 16,28 | 19,18 | 20,57 | 19,05 | 18,25 | **2,109** | 4,14E-01 | 0 | 1,07634036 | 0,38257384 |
| PDXD1_HUM | PDXDC1 | sp\|Q6P996\|PDXD1_HUMAN Pyridoxal-dependent decarboxylase domain-containing pro | 67,356 | 3,3 | 86707 | 2 | 20,69 | 21,50 | 19,87 | 20,76 | 22,04 | 22,49 | **2,107** | 1,99E-01 | 0 | 1,07551893 | 0,70142544 |
| CDSN_HUMA | CDSN | sp\|Q15517\|CDSN_HUMAN Corneodesmosin OS=Homo sapiens GN=CDSN PE=1 SV=3 | 24,781 | 3,4 | 51522 | 1 | 19,88 | 19,30 | 18,53 | 21,76 | 19,87 | 19,30 | **2,105** | 2,70E-01 | 0 | 1,07358615 | 0,56837878 |
| MYLK2_HUM | MYLK2 | sp\|Q9H1R3\|MYLK2_HUMAN Myosin light chain kinase 2, skeletal/cardiac muscle OS=Ho | 12,078 | 1,34 | 64685 | 1 | 22,72 | 22,22 | 18,69 | 22,33 | 22,06 | 22,46 | **2,105** | 4,47E-01 | 0 | 1,07354609 | 0,34985987 |
| MRT4_HUM | MRTO4 | sp\|Q9UKD2\|MRT4_HUMAN mRNA turnover protein 4 homolog OS=Homo sapiens GN= | 33,4 | 8,79 | 27560 | 2 | 17,49 | 18,20 | 20,68 | 17,56 | 19,39 | 22,62 | **2,096** | 5,79E-01 | 0 | 1,06752968 | 0,23769355 |
| CNDD3_HUM | NCAPD3 | sp\|P42695\|CNDD3_HUMAN Condensin-2 complex subunit D3 OS=Homo sapiens GN=N | 85,956 | 1,67 | 168891 | 2 | 19,15 | 17,58 | 19,38 | 20,45 | 18,65 | 20,20 | **2,088** | 2,55E-01 | 0 | 1,06178093 | 0,59415343 |
| DESP_HUMA | DSP | sp\|P15924\|DESP_HUMAN Desmoplakin OS=Homo sapiens GN=DSP PE=1 SV=3 | 273,152 | 7 | 331774 | 15 | 24,65 | 23,34 | 23,39 | 26,01 | 24,31 | 24,24 | **2,085** | 2,15E-01 | 0 | 1,0600001 | 0,66756627 |
| LAMP1_HUM | LAMP1 | sp\|P11279\|LAMP1_HUMAN Lysosome-associated membrane glycoprotein 1 OS=Homo | 41,995 | 4,56 | 44882 | 2 | 20,60 | 21,42 | 23,60 | 23,44 | 22,39 | 22,97 | **2,085** | 3,25E-01 | 0 | 1,05999947 | 0,48823603 |
| SRPRA_HUM | SRPRA | sp\|P08240\|SRPRA_HUMAN Signal recognition particle receptor subunit alpha OS=Homo | 230,863 | 15,05 | 69811 | 7 | 20,65 | 22,40 | 22,52 | 21,55 | 23,05 | 24,15 | **2,085** | 3,34E-01 | 0 | 1,05999947 | 0,47613673 |
| RANB3_HUM | RANBP3 | sp\|Q9H6Z4\|RANB3_HUMAN Ran-binding protein 3 OS=Homo sapiens GN=RANBP3 PE=1 | 35,341 | 3,7 | 60210 | 2 | 20,56 | 21,12 | 18,11 | 20,56 | 21,24 | 21,16 | **2,081** | 3,28E-01 | 0 | 1,05718549 | 0,48432798 |
| CLDN1_HUM | CLDND1 | sp\|Q9NY35\|CLDN1_HUMAN Claudin domain-containing protein 1 OS=Homo sapiens GN | 112,093 | 18,18 | 28603 | 4 | 25,28 | 25,21 | 23,74 | 25,06 | 25,29 | 27,05 | **2,080** | 2,59E-01 | 0 | 1,05666669 | 0,58624557 |
| 5NTD_HUMA | NT5E | sp\|P21589\|5NTD_HUMAN 5'-nucleotidase OS=Homo sapiens GN=NT5E PE=1 SV=1 | 146,853 | 10,45 | 63368 | 5 | 23,84 | 23,58 | 22,99 | 25,00 | 24,78 | 23,80 | **2,080** | 7,71E-02 | 0 | 1,05666669 | 1,11312668 |
| ACTT3_HUM | ACTRT3 | sp\|Q9BYD9\|ACTT3_HUMAN Actin-related protein T3 OS=Homo sapiens GN=ACTRT3 PE | 32,26 | 1,88 | 41008 | 1 | 24,69 | 24,28 | 18,34 | 24,70 | 23,78 | 21,99 | **2,075** | 6,57E-01 | 0 | 1,05339622 | 0,18232662 |
| FETUA_HUM | AHSG | sp\|P02765\|FETUA_HUMAN Alpha-2-HS-glycoprotein OS=Homo sapiens GN=AHSG PE=1 | 168,922 | 5,45 | 39325 | 3 | 26,66 | 22,54 | 29,14 | 27,39 | 25,97 | 28,13 | **2,071** | 6,32E-01 | 0 | 1,04999924 | 0,19949551 |
| COMT_HUM | COMT | sp\|P21964\|COMT_HUMAN Catechol O-methyltransferase OS=Homo sapiens GN=COMT | 74,856 | 9,23 | 30037 | 2 | 22,38 | 22,48 | 20,05 | 22,60 | 23,13 | 22,31 | **2,059** | 2,77E-01 | 0 | 1,04180463 | 0,55799009 |
| STAM2_HUM | STAM2 | sp\|O75886\|STAM2_HUMAN Signal transducing adapter molecule 2 OS=Homo sapiens G | 106,202 | 6,48 | 58164 | 4 | 21,51 | 21,58 | 18,56 | 21,77 | 21,41 | 21,58 | **2,052** | 3,59E-01 | 0 | 1,03709221 | 0,44524659 |
| UCHL1_HUM | UCHL1 | sp\|P09936\|UCHL1_HUMAN Ubiquitin carboxyl-terminal hydrolase isozyme L1 OS=Hom | 129,603 | 37,22 | 24824 | 5 | 24,88 | 24,98 | 20,98 | 25,29 | 25,02 | 23,62 | **2,042** | 5,07E-01 | 0 | 1,03000132 | 0,29499441 |
| RRP44_HUM | DIS3 | sp\|Q9Y2L1\|RRP44_HUMAN Exosome complex exonuclease RRP44 OS=Homo sapiens GN | 288,86 | 13,36 | 109003 | 10 | 21,87 | 21,89 | 23,90 | 23,14 | 22,91 | 24,70 | **2,042** | 3,05E-01 | 0 | 1,03000005 | 0,51502474 |
| BNIP2_HUM | BNIP2 | sp\|Q12982\|BNIP2_HUMAN BCL2/adenovirus E1B 19 kDa protein-interacting protein 2 O | 34,122 | 11,46 | 36018 | 2 | 17,78 | 21,35 | 21,00 | 20,58 | 21,49 | 21,15 | **2,042** | 4,27E-01 | 0 | 1,02999942 | 0,36935125 |
| DDX5_HUMA | DDX5 | sp\|P17844\|DDX5_HUMAN Probable ATP-dependent RNA helicase DDX5 OS=Homo sapie | 1119,965 | 44,46 | 69148 | 33 | 24,78 | 24,86 | 27,70 | 26,13 | 26,17 | 28,13 | **2,042** | 4,27E-01 | 0 | 1,02999878 | 0,36990475 |
| IDI1_HUMAN | IDI1 | sp\|Q13907\|IDI1_HUMAN Isopentenyl-diphosphate Delta-isomerase 1 OS=Homo sapiens | 62,449 | 14,98 | 26319 | 3 | 20,60 | 19,43 | 21,97 | 21,38 | 20,74 | 22,95 | **2,033** | 3,57E-01 | 0 | 1,02333323 | 0,44682425 |
| SNUT2_HUM | USP39 | sp\|Q53GS9\|SNUT2_HUMAN U4/U6.U5 tri-snRNP-associated protein 2 OS=Homo sapien | 48,632 | 6,73 | 65381 | 3 | 16,90 | 18,06 | 20,21 | 18,21 | 18,81 | 21,22 | **2,033** | 4,87E-01 | 0 | 1,02333323 | 0,31287152 |
| GBRL2_HUM | GABARAPL2 | sp\|P60520\|GBRL2_HUMAN Gamma-aminobutyric acid receptor-associated protein-like | 77,316 | 35,04 | 13667 | 4 | 22,24 | 21,80 | 21,42 | 23,53 | 22,98 | 22,01 | **2,028** | 1,13E-01 | 0 | 1,02000046 | 0,94778408 |
| PFD4_HUMA | PFDN4 | sp\|Q9NQP4\|PFD4_HUMAN Prefoldin subunit 4 OS=Homo sapiens GN=PFDN4 PE=1 SV=1 | 109,58 | 19,4 | 15314 | 2 | 20,88 | 21,63 | 17,89 | 21,31 | 20,93 | 21,22 | **2,026** | 4,25E-01 | 0 | 1,0186526 | 0,37203245 |
| K2C6A_HUM | KRT6A | sp\|P02538\|K2C6A_HUMAN Keratin, type II cytoskeletal 6A OS=Homo sapiens GN=KRT6 | 1149,335 | 54,08 | 60045 | 36 | 23,78 | 22,68 | 20,37 | 25,31 | 22,70 | 21,87 | **2,023** | 5,20E-01 | 0 | 1,01666641 | 0,28391568 |
| CPSF2_HUM | CPSF2 | sp\|Q9P2I0\|CPSF2_HUMAN Cleavage and polyadenylation specificity factor subunit 2 OS | 121,478 | 3,58 | 88487 | 2 | 19,95 | 20,23 | 20,02 | 20,68 | 20,63 | 21,94 | **2,023** | 8,04E-02 | 0 | 1,01666641 | 1,09448154 |
| CASPE_HUM | CASP14 | sp\|P31944\|CASPE_HUMAN Caspase-14 OS=Homo sapiens GN=CASP14 PE=1 SV=2 | 37,281 | 10,74 | 27680 | 2 | 20,72 | 18,91 | 19,05 | 22,67 | 20,24 | 18,79 | **2,010** | 4,72E-01 | 0 | 1,00742213 | 0,32564127 |
| MSS4_HUM | RABIF | sp\|P47224\|MSS4_HUMAN Guanine nucleotide exchange factor MSS4 OS=Homo sapiens | 22,813 | 8,94 | 13839 | 1 | 22,15 | 22,65 | 18,69 | 22,46 | 22,33 | 21,72 | **2,007** | 4,71E-01 | 0 | 1,00518354 | 0,32688204 |
| SMU1_HUM | SMU1 | sp\|Q2TAY7\|SMU1_HUMAN WD40 repeat-containing protein SMU1 OS=Homo sapiens G | 129,839 | 10,92 | 57544 | 5 | 21,77 | 21,96 | 23,57 | 22,97 | 23,18 | 24,16 | **2,005** | 2,13E-01 | 0 | 1,00333341 | 0,6709274 |

| | | | | | | | | | | | | | | | | |
|---|---|---|---|---|---|---|---|---|---|---|---|---|---|---|---|---|
| DFFA_HUMA | DFFA | sp\|O00273\|DFFA_HUMAN DNA fragmentation factor subunit alpha OS=Homo sapiens G | 10,738 | 2,72 | 36522 | 1 | 19,32 | 20,89 | 19,82 | 21,43 | 21,28 | 20,32 | **2,000** | 1,59E-01 | **0** | 1,00000064 | 0,79788521 |
| DCPS_HUMA | DCPS | sp\|Q96C86\|DCPS_HUMAN m7GpppX diphosphatase OS=Homo sapiens GN=DCPS PE=1 S | 34,283 | 5,93 | 38609 | 2 | 19,90 | 21,67 | 21,26 | 22,05 | 21,94 | 21,84 | **2,000** | 1,37E-01 | **0** | 1 | 0,86393831 |
| UBE2H_HUM | UBE2H | sp\|P62256\|UBE2H_HUMAN Ubiquitin-conjugating enzyme E2 H OS=Homo sapiens GN=U | 19,156 | 8,2 | 20655 | 1 | 18,20 | 21,36 | 20,40 | 22,12 | 22,30 | 18,53 | **1,994** | 5,55E-01 | **0** | 0,99540583 | 0,25597527 |
| QBK84320.1 | | envelope glycoprotein H [Human betaherpesvirus 5] | 91,385 | 4,85 | 84227 | 3 | 20,01 | 19,22 | 18,83 | 21,93 | 21,10 | 18,02 | **1,993** | 4,67E-01 | **0** | 0,99479993 | 0,33082008 |
| K22E_HUMA | KRT2 | sp\|P35908\|K22E_HUMAN Keratin, type II cytoskeletal 2 epidermal OS=Homo sapiens GN | 3518,817 | 84,19 | 65433 | 70 | 30,24 | 29,12 | 28,80 | 32,47 | 30,13 | 28,53 | **1,986** | 4,64E-01 | **0** | 0,99000041 | 0,33334085 |
| STMN2_HUN | STMN2 | sp\|Q93045\|STMN2_HUMAN Stathmin-2 OS=Homo sapiens GN=STMN2 PE=1 SV=3 | 33,237 | 5,59 | 20828 | 1 | 21,48 | 19,53 | 21,30 | 22,68 | 21,43 | 21,16 | **1,983** | 2,74E-01 | **0** | 0,98741913 | 0,56246971 |
| K2C6B_HUM | KRT6B | sp\|P04259\|K2C6B_HUMAN Keratin, type II cytoskeletal 6B OS=Homo sapiens GN=KRT6E | 1055,775 | 49,47 | 60067 | 30 | 18,79 | 19,69 | 18,33 | 22,78 | 17,82 | 19,17 | **1,980** | 5,56E-01 | **0** | 0,9852651 | 0,25515026 |
| PRI1_HUMAI | PRIM1 | sp\|P49642\|PRI1_HUMAN DNA primase small subunit OS=Homo sapiens GN=PRIM1 PE=1 | 13,469 | 2,14 | 49902 | 1 | 21,19 | 21,50 | 18,50 | 21,27 | 21,15 | 21,72 | **1,978** | 3,67E-01 | **0** | 0,98392677 | 0,43503912 |
| PSB7_HUMA | PSMB7 | sp\|Q99436\|PSB7_HUMAN Proteasome subunit beta type-7 OS=Homo sapiens GN=PSMI | 274,663 | 31,41 | 29965 | 7 | 25,00 | 24,76 | 23,42 | 25,46 | 25,72 | 24,95 | **1,977** | 1,43E-01 | **0** | 0,98333295 | 0,84366831 |
| ODPA_HUM/ | PDHA1 | sp\|P08559\|ODPA_HUMAN Pyruvate dehydrogenase E1 component subunit alpha, soma | 73,798 | 7,18 | 43296 | 3 | 23,99 | 19,59 | 24,33 | 23,71 | 23,70 | 23,44 | **1,973** | 5,56E-01 | **0** | 0,98073324 | 0,25465335 |
| BPTF_HUMA | BPTF | sp\|Q12830\|BPTF_HUMAN Nucleosome-remodeling factor subunit BPTF OS=Homo sapie | 18,701 | 0,23 | 338262 | 1 | 20,69 | 17,75 | 18,86 | 20,31 | 20,15 | 19,78 | **1,973** | 3,24E-01 | **0** | 0,98005931 | 0,48960924 |
| EMC4_HUM/ | EMC4 | sp\|Q5J8M3\|EMC4_HUMAN ER membrane protein complex subunit 4 OS=Homo sapiens | 22,55 | 6,56 | 20087 | 1 | 18,05 | 17,94 | 19,95 | 19,54 | 19,08 | 20,26 | **1,972** | 2,55E-01 | **0** | 0,97985713 | 0,59417098 |
| PANX2_HUM | PANX2 | sp\|Q96RD6\|PANX2_HUMAN Pannexin-2 OS=Homo sapiens GN=PANX2 PE=2 SV=2 | 18,593 | 1,18 | 74447 | 1 | 24,65 | 23,90 | 24,38 | 25,76 | 25,52 | 24,58 | **1,968** | 8,14E-02 | **0** | 0,9766674 | 1,08915622 |
| ERI3_HUMAI | ERI3 | sp\|O43414\|ERI3_HUMAN ERI1 exoribonuclease 3 OS=Homo sapiens GN=ERI3 PE=1 SV=: | 16,813 | 2,67 | 37238 | 1 | 18,21 | 22,48 | 22,01 | 22,09 | 21,90 | 21,63 | **1,964** | 5,13E-01 | **0** | 0,97402954 | 0,28972134 |
| MEP50_HUN | WDR77 | sp\|Q9BQA1\|MEP50_HUMAN Methylosome protein 50 OS=Homo sapiens GN=WDR77 Pl | 41,539 | 5,56 | 36724 | 2 | 21,37 | 21,45 | 20,71 | 22,22 | 22,65 | 21,57 | **1,959** | 6,85E-02 | **0** | 0,96999931 | 1,16410909 |
| TRM6_HUM/ | TRMT6 | sp\|Q9UJA5\|TRM6_HUMAN tRNA (adenine(58)-N(1))-methyltransferase non-catalytic sul | 10,747 | 2,01 | 55799 | 1 | 21,19 | 21,09 | 18,96 | 21,80 | 21,54 | 20,79 | **1,951** | 2,89E-01 | **0** | 0,96437772 | 0,53977766 |
| PHF5A_HUM | PHF5A | sp\|Q7RTV0\|PHF5A_HUMAN PHD finger-like domain-containing protein 5A OS=Homo sa | 160,488 | 54,55 | 12405 | 5 | 23,10 | 23,89 | 22,40 | 24,39 | 23,92 | 23,97 | **1,950** | 1,02E-01 | **0** | 0,96333313 | 0,99180066 |
| NEP1_HUMA | EMG1 | sp\|Q92979\|NEP1_HUMAN Ribosomal RNA small subunit methyltransferase NEP1 OS=Hc | 102,818 | 19,67 | 26720 | 4 | 21,74 | 19,56 | 23,37 | 21,41 | 22,02 | 24,12 | **1,945** | 5,24E-01 | **0** | 0,96000036 | 0,28096738 |
| NELFB_HUM | NELFB | sp\|Q8WX92\|NELFB_HUMAN Negative elongation factor B OS=Homo sapiens GN=NELFB | 21,586 | 2,24 | 65697 | 1 | 18,57 | 20,06 | 20,70 | 20,57 | 20,36 | 21,27 | **1,941** | 2,37E-01 | **0** | 0,9570605 | 0,62529767 |
| NOP56_HUN | NOP56 | sp\|O00567\|NOP56_HUMAN Nucleolar protein 56 OS=Homo sapiens GN=NOP56 PE=1 SV | 233,64 | 18,86 | 66050 | 9 | 22,34 | 22,07 | 24,58 | 23,41 | 23,07 | 25,38 | **1,941** | 4,23E-01 | **0** | 0,95666631 | 0,37366387 |
| RAB4B_HUM | RAB4B | sp\|P61018\|RAB4B_HUMAN Ras-related protein Rab-4B OS=Homo sapiens GN=RAB4B PE | 114,153 | 16,43 | 23587 | 3 | 17,74 | 22,07 | 20,53 | 21,51 | 21,54 | 20,16 | **1,939** | 5,17E-01 | **0** | 0,95517921 | 0,28663712 |
| COPZ1_HUM | COPZ1 | sp\|P61923\|COPZ1_HUMAN Coatomer subunit zeta-1 OS=Homo sapiens GN=COPZ1 PE=: | 75,578 | 14,12 | 20198 | 2 | 20,95 | 22,58 | 22,45 | 22,17 | 23,16 | 23,51 | **1,936** | 2,22E-01 | **0** | 0,9533329 | 0,65430872 |
| ACADV_HUN | ACADVL | sp\|P49748\|ACADV_HUMAN Very long-chain specific acyl-CoA dehydrogenase, mitochor | 10,577 | 1,22 | 70390 | 1 | 17,81 | 20,31 | 22,33 | 19,22 | 21,55 | 22,54 | **1,933** | 5,92E-01 | **0** | 0,95065371 | 0,22746204 |
| ACTBL_HUM | ACTBL2 | sp\|Q562R1\|ACTBL_HUMAN Beta-actin-like protein 2 OS=Homo sapiens GN=ACTBL2 PE= | 866,731 | 33,78 | 42003 | 22 | 24,14 | 22,75 | 24,19 | 24,95 | 24,91 | 24,07 | **1,932** | 1,61E-01 | **0** | 0,95000013 | 0,79449652 |
| TUT7_HUMA | ZCCHC6 | sp\|Q5VYS8\|TUT7_HUMAN Terminal uridylyltransferase 7 OS=Homo sapiens GN=ZCCHC | 16,176 | 0,47 | 171229 | 1 | 23,89 | 23,28 | 21,55 | 24,08 | 23,41 | 24,05 | **1,919** | 2,70E-01 | **0** | 0,9399999 | 0,56931116 |
| SMYD5_HUN | SMYD5 | sp\|Q6GMV2\|SMYD5_HUMAN SET and MYND domain-containing protein 5 OS=Homo sa | 10,053 | 3,35 | 47341 | 1 | 16,85 | 18,20 | 18,55 | 19,79 | 17,79 | 18,84 | **1,916** | 2,93E-01 | **0** | 0,93814341 | 0,53295964 |
| PURA2_HUM | ADSS | sp\|P30520\|PURA2_HUMAN Adenylosuccinate synthetase isozyme 2 OS=Homo sapiens ( | 270,401 | 24,78 | 50097 | 9 | 23,24 | 23,22 | 25,08 | 24,68 | 24,38 | 25,29 | **1,914** | 2,36E-01 | **0** | 0,93666712 | 0,62711896 |
| K2C80_HUM | KRT80 | sp\|Q6KB66\|K2C80_HUMAN Keratin, type II cytoskeletal 80 OS=Homo sapiens GN=KRT8( | 132,707 | 9,51 | 50525 | 4 | 20,21 | 19,23 | 19,50 | 23,29 | 19,73 | 18,72 | **1,912** | 5,45E-01 | **0** | 0,93482208 | 0,2632768 |
| COIA1_HUM | COL18A1 | sp\|P39060\|COIA1_HUMAN Collagen alpha-1(XVIII) chain OS=Homo sapiens GN=COL18A | 60,243 | 1,14 | 178188 | 2 | 24,05 | 22,92 | 21,76 | 24,11 | 23,75 | 23,67 | **1,910** | 2,39E-01 | **0** | 0,93333371 | 0,6219433 |
| HDGF_HUM/ | HDGF | sp\|P51858\|HDGF_HUMAN Hepatoma-derived growth factor OS=Homo sapiens GN=HD( | 174,685 | 22,08 | 26788 | 6 | 23,47 | 24,09 | 22,98 | 24,44 | 24,62 | 24,27 | **1,905** | 5,07E-02 | **0** | 0,93000094 | 1,29471104 |
| NOP2_HUM/ | NOP2 | sp\|P46087\|NOP2_HUMAN Probable 28S rRNA (cytosine(4447)-C(5))-methyltransferase ( | 46,854 | 3,82 | 89302 | 3 | 20,05 | 20,31 | 22,42 | 21,39 | 20,92 | 23,26 | **1,905** | 4,20E-01 | **0** | 0,93000031 | 0,3765234 |
| API5_HUMAI | API5 | sp\|Q9BZZ5\|API5_HUMAN Apoptosis inhibitor 5 OS=Homo sapiens GN=API5 PE=1 SV=3 | 255,843 | 16,6 | 59005 | 6 | 21,62 | 21,81 | 22,63 | 22,16 | 22,96 | 23,72 | **1,901** | 1,65E-01 | **0** | 0,92666626 | 0,78168629 |
| SHCBP_HUM | SHCBP1 | sp\|Q8NEM2\|SHCBP_HUMAN SHC SH2 domain-binding protein 1 OS=Homo sapiens GN= | 21,042 | 4,02 | 75690 | 3 | 20,82 | 19,77 | 23,51 | 21,21 | 20,66 | 25,00 | **1,896** | 6,28E-01 | **0** | 0,92333285 | 0,20218496 |
| PURB_HUMA | PURB | sp\|Q96QR8\|PURB_HUMAN Transcriptional activator protein Pur-beta OS=Homo sapiens | 117,659 | 23,08 | 33241 | 4 | 23,39 | 22,97 | 24,17 | 23,91 | 23,95 | 25,42 | **1,888** | 2,06E-01 | **0** | 0,9166673 | 0,68509486 |
| NPM_HUMA | NPM1 | sp\|P06748\|NPM_HUMAN Nucleophosmin OS=Homo sapiens GN=NPM1 PE=1 SV=2 | 502,86 | 52,04 | 32575 | 9 | 26,08 | 26,51 | 26,21 | 27,12 | 27,15 | 27,28 | **1,888** | 2,56E-03 | **0** | 0,9166673 | 2,59221875 |
| IPYR2_HUM/ | PPA2 | sp\|Q9H2U2\|IPYR2_HUMAN Inorganic pyrophosphatase 2, mitochondrial OS=Homo sapi | 96,731 | 9,88 | 37920 | 3 | 22,59 | 25,27 | 24,34 | 24,30 | 25,71 | 24,94 | **1,888** | 3,59E-01 | **0** | 0,91666603 | 0,44508876 |
| UB2R2_HUM | UBE2R2 | sp\|Q712K3\|UB2R2_HUMAN Ubiquitin-conjugating enzyme E2 R2 OS=Homo sapiens GN: | 14,23 | 2,94 | 27166 | 1 | 21,24 | 18,73 | 20,04 | 21,50 | 21,11 | 20,15 | **1,888** | 3,30E-01 | **0** | 0,91662598 | 0,48116493 |
| PSPC1_HUM | PSPC1 | sp\|Q8WXF1\|PSPC1_HUMAN Paraspeckle component 1 OS=Homo sapiens GN=PSPC1 PE | 210,599 | 17,02 | 58744 | 7 | 22,91 | 23,52 | 24,64 | 24,21 | 24,40 | 25,19 | **1,879** | 1,97E-01 | **0** | 0,90999985 | 0,70530218 |
| GRB10_HUM | GRB10 | sp\|Q13322\|GRB10_HUMAN Growth factor receptor-bound protein 10 OS=Homo sapier | 96,08 | 5,89 | 67231 | 2 | 18,75 | 18,09 | 22,06 | 20,37 | 20,23 | 21,02 | **1,873** | 5,09E-01 | **0** | 0,9057045 | 0,2930048 |
| WBP2_HUM/ | WBP2 | sp\|Q969T9\|WBP2_HUMAN WW domain-binding protein 2 OS=Homo sapiens GN=WBP2 | 51,2 | 9,2 | 28087 | 2 | 23,98 | 23,72 | 17,79 | 22,89 | 23,58 | 21,73 | **1,872** | 6,88E-01 | **0** | 0,90480868 | 0,16247727 |
| NASP_HUMA | NASP | sp\|P49321\|NASP_HUMAN Nuclear autoantigenic sperm protein OS=Homo sapiens GN=I | 102,62 | 5,71 | 85238 | 3 | 22,30 | 22,05 | 22,81 | 22,99 | 22,96 | 23,91 | **1,866** | 7,89E-02 | **0** | 0,90000025 | 1,102962 |
| HAP28_HUM | PDAP1 | sp\|Q13442\|HAP28_HUMAN 28 kDa heat- and acid-stable phosphoprotein OS=Homo sap | 117,591 | 22,1 | 20630 | 3 | 22,81 | 22,88 | 18,61 | 23,06 | 22,99 | 20,94 | **1,863** | 5,99E-01 | **0** | 0,89768092 | 0,22249059 |
| CDIPT_HUM/ | CDIPT | sp\|O14735\|CDIPT_HUMAN CDP-diacylglycerol--inositol 3-phosphatidyltransferase OS=H | 96,375 | 10,33 | 23539 | 2 | 22,30 | 22,34 | 22,37 | 23,06 | 23,37 | 23,27 | **1,862** | 6,63E-04 | **0** | 0,89666684 | 3,17868622 |
| FXL15_HUM/ | FBXL15 | sp\|Q9H469\|FXL15_HUMAN F-box/LRR-repeat protein 15 OS=Homo sapiens GN=FBXL15 | 14,866 | 4 | 32998 | 1 | 16,57 | 13,55 | 17,54 | 17,18 | 16,30 | 16,87 | **1,862** | 5,06E-01 | **0** | 0,89666653 | 0,29586138 |
| CADM1_HUN | CADM1 | sp\|Q9BY67\|CADM1_HUMAN Cell adhesion molecule 1 OS=Homo sapiens GN=CADM1 PI | 57,56 | 15,38 | 48509 | 4 | 21,05 | 21,59 | 20,89 | 22,52 | 21,96 | 21,73 | **1,857** | 4,75E-02 | **0** | 0,89333344 | 1,32326803 |
| SLMAP_HUN | SLMAP | sp\|Q14BN4\|SLMAP_HUMAN Sarcolemmal membrane-associated protein OS=Homo sap | 91,458 | 5,68 | 95198 | 4 | 19,52 | 21,39 | 23,67 | 21,48 | 22,29 | 23,48 | **1,854** | 5,41E-01 | **0** | 0,89040502 | 0,26692921 |
| LMAN1_HUN | LMAN1 | sp\|P49257\|LMAN1_HUMAN Protein ERGIC-53 OS=Homo sapiens GN=LMAN1 PE=1 SV=2 | 71,848 | 7,06 | 57549 | 4 | 22,83 | 21,92 | 23,05 | 23,27 | 23,33 | 23,87 | **1,853** | 8,73E-02 | **0** | 0,89000066 | 1,0587659 |
| KDM1A_HUN | KDM1A | sp\|O60341\|KDM1A_HUMAN Lysine-specific histone demethylase 1A OS=Homo sapiens | 29,112 | 2,46 | 92903 | 2 | 20,41 | 20,90 | 19,62 | 21,27 | 21,25 | 21,08 | **1,852** | 7,78E-02 | **0** | 0,88913091 | 1,10928791 |
| QBK84433.1 | | envelope glycoprotein N [Human betaherpesvirus 5] | 17,463 | 6,57 | 14801 | 2 | 18,37 | 18,01 | 20,51 | 18,84 | 17,89 | 22,82 | **1,850** | 6,29E-01 | **0** | 0,88744354 | 0,20133297 |
| MOGS_HUM | MOGS | sp\|Q13724\|MOGS_HUMAN Mannosyl-oligosaccharide glucosidase OS=Homo sapiens G | 25,681 | 1,79 | 91918 | 1 | 20,20 | 17,92 | 20,79 | 20,58 | 19,60 | 21,39 | **1,849** | 4,32E-01 | **0** | 0,88694382 | 0,36488534 |
| SPCS3_HUM | SPCS3 | sp\|P61009\|SPCS3_HUMAN Signal peptidase complex subunit 3 OS=Homo sapiens GN=SI | 83,244 | 21,67 | 20313 | 4 | 24,43 | 23,59 | 24,34 | 24,88 | 24,69 | 25,45 | **1,849** | 6,48E-02 | **0** | 0,88666662 | 1,18827999 |
| UTP4_HUMA | UTP4 | sp\|Q969X6\|UTP4_HUMAN U3 small nucleolar RNA-associated protein 4 homolog OS=Hc | 30,771 | 1,75 | 76890 | 1 | 17,57 | 15,97 | 18,99 | 18,67 | 17,77 | 18,74 | **1,845** | 3,94E-01 | **0** | 0,88333352 | 0,40407291 |
| C1D_HUMAN | C1D | sp\|Q13901\|C1D_HUMAN Nuclear nucleic acid-binding protein C1D OS=Homo sapiens G | 18,021 | 7,09 | 16019 | 1 | 17,66 | 17,31 | 18,58 | 18,81 | 18,19 | 19,18 | **1,836** | 1,39E-01 | **0** | 0,87666702 | 0,85544545 |
| CNTP4_HUM | CNTNAP4 | sp\|Q9C0A0\|CNTP4_HUMAN Contactin-associated protein-like 4 OS=Homo sapiens GN=( | 9,027 | 0,54 | 145274 | 1 | 22,17 | 22,47 | 24,00 | 23,61 | 23,13 | 24,51 | **1,828** | 2,80E-01 | **0** | 0,8700002 | 0,55352474 |
| RINT1_HUM | RINT1 | sp\|Q6NUQ1\|RINT1_HUMAN RAD50-interacting protein 1 OS=Homo sapiens GN=RINT1 F | 14,356 | 1,52 | 90632 | 1 | 17,34 | 19,99 | 20,88 | 19,14 | 20,16 | 21,51 | **1,821** | 5,32E-01 | **0** | 0,86447016 | 0,27426509 |
| NOMO1_HUI | NOMO1 | sp\|Q15155\|NOMO1_HUMAN Nodal modulator 1 OS=Homo sapiens GN=NOMO1 PE=1 S | 155,875 | 7,45 | 134324 | 7 | 22,38 | 21,57 | 24,47 | 22,85 | 23,22 | 24,94 | **1,819** | 4,68E-01 | **0** | 0,86333402 | 0,32986394 |
| MSH6_HUM/ | MSH6 | sp\|P52701\|MSH6_HUMAN DNA mismatch repair protein Msh6 OS=Homo sapiens GN=M | 84,579 | 2,21 | 152786 | 2 | 17,13 | 18,85 | 21,27 | 18,22 | 19,95 | 21,66 | **1,815** | 6,10E-01 | **0** | 0,85999997 | 0,21439054 |
| RPAB1_HUM | POLR2E | sp\|P19388\|RPAB1_HUMAN DNA-directed RNA polymerases I, II, and III subunit RPABC1 ( | 22,522 | 4,76 | 24551 | 1 | 20,64 | 18,13 | 21,48 | 20,94 | 20,32 | 21,54 | **1,801** | 4,70E-01 | **0** | 0,84913953 | 0,32783991 |
| RBM39_HUN | RBM39 | sp\|Q14498\|RBM39_HUMAN RNA-binding protein 39 OS=Homo sapiens GN=RBM39 PE= | 61,43 | 7,92 | 59380 | 3 | 18,59 | 21,20 | 22,59 | 22,88 | 20,67 | 21,37 | **1,800** | 5,62E-01 | **0** | 0,84769885 | 0,25024037 |
| ABHDA_HUN | ABHD10 | sp\|Q9NUJ1\|ABHDA_HUMAN Mycophenolic acid acyl-glucuronide esterase, mitochondri | 44,769 | 5,23 | 33933 | 1 | 17,97 | 18,05 | 18,72 | 18,84 | 18,99 | 19,46 | **1,799** | 4,86E-02 | **0** | 0,84707769 | 1,31335022 |
| CDC27_HUM | CDC27 | sp\|P30260\|CDC27_HUMAN Cell division cycle protein 27 homolog OS=Homo sapiens GN | 14,403 | 3,52 | 91867 | 1 | 17,12 | 19,85 | 20,28 | 19,22 | 19,62 | 20,95 | **1,798** | 4,91E-01 | **0** | 0,84666634 | 0,30852188 |
| YTHD3_HUM | YTHDF3 | sp\|Q7Z739\|YTHD3_HUMAN YTH domain-containing family protein 3 OS=Homo sapiens ( | 30,85 | 3,25 | 63861 | 2 | 16,96 | 18,29 | 22,35 | 20,00 | 18,25 | 21,89 | **1,798** | 6,84E-01 | **0** | 0,84662755 | 0,16509303 |
| ANM5_HUM | PRMT5 | sp\|O14744\|ANM5_HUMAN Protein arginine N-methyltransferase 5 OS=Homo sapiens G | 356,569 | 24,96 | 72684 | 12 | 24,31 | 24,76 | 24,64 | 25,09 | 25,42 | 25,73 | **1,794** | 2,10E-02 | **0** | 0,84333356 | 1,67724159 |
| UFSP2_HUM | UFSP2 | sp\|Q9NUQ7\|UFSP2_HUMAN Ufm1-specific protease 2 OS=Homo sapiens GN=UFSP2 PE: | 30,19 | 4,26 | 53261 | 1 | 17,62 | 17,48 | 21,39 | 18,43 | 18,90 | 21,69 | **1,791** | 6,34E-01 | **0** | 0,8411293 | 0,19784115 |
| OSBP1_HUM | OSBP | sp\|P22059\|OSBP1_HUMAN Oxysterol-binding protein 1 OS=Homo sapiens GN=OSBP PE | 165,351 | 11,77 | 89421 | 6 | 21,22 | 21,78 | 22,27 | 22,43 | 22,18 | 23,18 | **1,790** | 1,21E-01 | **0** | 0,84000015 | 0,91892643 |
| ACLY_HUMA | ACLY | sp\|P53396\|ACLY_HUMAN ATP-citrate synthase OS=Homo sapiens GN=ACLY PE=1 SV=3 | 1843,594 | 41,96 | 120839 | 41 | 29,21 | 28,89 | 28,65 | 30,99 | 29,06 | 29,20 | **1,782** | 2,64E-01 | **0** | 0,83333397 | 0,5781436 |
| TOP1_HUMA | TOP1 | sp\|P11387\|TOP1_HUMAN DNA topoisomerase 1 OS=Homo sapiens GN=TOP1 PE=1 SV=: | 39,218 | 4,31 | 90726 | 3 | 21,92 | 21,18 | 22,56 | 22,32 | 22,82 | 23,01 | **1,778** | 1,38E-01 | **0** | 0,82999992 | 0,86006959 |
| CYBP_HUMA | CACYBP | sp\|Q9HB71\|CYBP_HUMAN Calcyclin-binding protein OS=Homo sapiens GN=CACYBP PE= | 183,958 | 25,44 | 26210 | 7 | 24,30 | 24,62 | 22,76 | 24,98 | 25,14 | 24,05 | **1,778** | 2,81E-01 | **0** | 0,82999929 | 0,55067156 |
| SELB_HUMA | EEFSEC | sp\|P57772\|SELB_HUMAN Selenocysteine-specific elongation factor OS=Homo sapiens G | 23,256 | 4,19 | 65305 | 2 | 17,53 | 19,10 | 21,91 | 20,41 | 18,70 | 21,90 | **1,773** | 6,29E-01 | **0** | 0,82610512 | 0,20147282 |
| TRM61_HUN | TRMT61A | sp\|Q96FX7\|TRM61_HUMAN tRNA (adenine(58)-N(1))-methyltransferase catalytic subun | 37,603 | 19,03 | 31382 | 3 | 20,59 | 20,93 | 21,16 | 22,19 | 21,03 | 21,93 | **1,769** | 1,01E-01 | **0** | 0,82333374 | 0,99388456 |
| CERS2_HUM | CERS2 | sp\|Q96G23\|CERS2_HUMAN Ceramide synthase 2 OS=Homo sapiens GN=CERS2 PE=1 SV | 78,878 | 4,74 | 44876 | 2 | 22,64 | 22,69 | 22,93 | 23,63 | 23,73 | 23,37 | **1,769** | 4,15E-03 | **0** | 0,8233331 | 2,38219415 |
| CTBP1_HUM | CTBP1 | sp\|Q13363\|CTBP1_HUMAN C-terminal-binding protein 1 OS=Homo sapiens GN=CTBP1 I | 158,804 | 19,09 | 47535 | 7 | 22,62 | 21,83 | 23,99 | 23,09 | 23,16 | 24,66 | **1,769** | 3,68E-01 | **0** | 0,8233331 | 0,43382236 |
| DKC1_HUMA | DKC1 | sp\|O60832\|DKC1_HUMAN H/ACA ribonucleoprotein complex subunit 4 OS=Homo sapie | 58,967 | 9,14 | 57674 | 3 | 18,57 | 19,94 | 22,78 | 19,18 | 21,18 | 23,39 | **1,764** | 6,62E-01 | **0** | 0,81906446 | 0,1793792 |
| HEAT3_HUM | HEATR3 | sp\|Q7Z4Q2\|HEAT3_HUMAN HEAT repeat-containing protein 3 OS=Homo sapiens GN=HI | 38,778 | 1,62 | 74583 | 1 | 16,39 | 19,97 | 20,92 | 18,25 | 20,12 | 21,36 | **1,761** | 6,46E-01 | **0** | 0,81666756 | 0,18949986 |
| IPO11_HUM/ | IPO11 | sp\|Q9UI26\|IPO11_HUMAN Importin-11 OS=Homo sapiens GN=IPO11 PE=1 SV=1 | 222,548 | 14,36 | 112535 | 9 | 21,98 | 21,92 | 22,97 | 22,79 | 22,88 | 23,65 | **1,761** | 1,35E-01 | **0** | 0,81666692 | 0,87106565 |
| DSC1_HUMA | DSC1 | sp\|Q08554\|DSC1_HUMAN Desmocollin-1 OS=Homo sapiens GN=DSC1 PE=1 SV=2 | 71,494 | 4,36 | 99987 | 3 | 22,97 | 21,68 | 21,72 | 24,32 | 23,03 | 21,47 | **1,761** | 4,28E-01 | **0** | 0,81666692 | 0,36874415 |
| ZN326_HUM | ZNF326 | sp\|Q5BKZ1\|ZN326_HUMAN DBIRD complex subunit ZNF326 OS=Homo sapiens GN=ZNF: | 103,278 | 7,56 | 65654 | 4 | 20,46 | 21,11 | 22,29 | 21,69 | 21,57 | 23,05 | **1,761** | 3,17E-01 | **0** | 0,81666629 | 0,49829377 |
| RBMX_HUM/ | RBMX | sp\|P38159\|RBMX_HUMAN RNA-binding motif protein, X chromosome OS=Homo sapien | 73,652 | 8,7 | 42332 | 3 | 22,42 | 22,53 | 22,31 | 23,43 | 23,45 | 22,82 | **1,757** | 1,98E-02 | **0** | 0,81333351 | 1,70408234 |
| NH2L1_HUM | SNU13 | sp\|P55769\|NH2L1_HUMAN NHP2-like protein 1 OS=Homo sapiens GN=SNU13 PE=1 SV= | 143,231 | 37,5 | 14174 | 5 | 22,79 | 23,92 | 23,43 | 23,81 | 24,16 | 24,61 | **1,757** | 1,12E-01 | **0** | 0,81333288 | 0,94958711 |
| CUL4A_HUM | CUL4A | sp\|Q13619\|CUL4A_HUMAN Cullin-4A OS=Homo sapiens GN=CUL4A PE=1 SV=3 | 246,769 | 10,41 | 87680 | 7 | 21,33 | 21,03 | 21,13 | 21,61 | 21,79 | 22,52 | **1,753** | 5,01E-02 | **0** | 0,81000074 | 1,30047616 |
| K1C10_HUM | KRT10 | sp\|P13645\|K1C10_HUMAN Keratin, type I cytoskeletal 10 OS=Homo sapiens GN=KRT10 | 3282,328 | 60,03 | 59511 | 56 | 31,07 | 29,88 | 30,24 | 32,73 | 30,71 | 30,18 | **1,753** | 3,96E-01 | **0** | 0,8100001 | 0,40217786 |
| NSDHL_HUM | NSDHL | sp\|Q15738\|NSDHL_HUMAN Sterol-4-alpha-carboxylate 3-dehydrogenase, decarboxylat | 44,047 | 11,26 | 41900 | 3 | 21,58 | 20,87 | 23,45 | 22,02 | 22,39 | 23,91 | **1,749** | 4,49E-01 | **0** | 0,80666606 | 0,34760338 |
| CSN7A_HUM | COPS7A | sp\|Q9UBW8\|CSN7A_HUMAN COP9 signalosome complex subunit 7a OS=Homo sapiens | 108,334 | 10,91 | 30277 | 2 | 19,14 | 19,22 | 20,03 | 19,61 | 20,06 | 21,13 | **1,745** | 2,06E-01 | **0** | 0,80333328 | 0,68571785 |
| TM9S4_HUN | TM9SF4 | sp\|Q92544\|TM9S4_HUMAN Transmembrane 9 superfamily member 4 OS=Homo sapien | 79,224 | 10,44 | 74519 | 5 | 21,55 | 21,31 | 23,49 | 21,80 | 23,13 | 23,83 | **1,745** | 4,28E-01 | **0** | 0,80333328 | 0,36866731 |
| SERB_HUMA | PSPH | sp\|P78330\|SERB_HUMAN Phosphoserine phosphatase OS=Homo sapiens GN=PSPH PE= | 30,453 | 4 | 25008 | 1 | 21,82 | 21,13 | 21,23 | 22,51 | 22,37 | 21,70 | **1,741** | 7,24E-02 | **0** | 0,80000114 | 1,14048914 |
| S61A2_HUM | SEC61A2 | sp\|Q9H9S3\|S61A2_HUMAN Protein transport protein Sec61 subunit alpha isoform 2 OS | 261,554 | 17,23 | 52248 | 9 | 18,70 | 18,99 | 17,55 | 19,35 | 18,85 | 19,44 | **1,741** | 1,68E-01 | **0** | 0,80000051 | 0,77353286 |
| UBFD1_HUM | UBFD1 | sp\|O14562\|UBFD1_HUMAN Ubiquitin domain-containing protein UBFD1 OS=Homo sapi | 16,691 | 3,56 | 33382 | 1 | 21,72 | 20,92 | 18,82 | 21,18 | 21,40 | 21,27 | **1,738** | 4,10E-01 | **0** | 0,79709689 | 0,38703015 |
| ACSL1_HUM | ACSL1 | sp\|P33121\|ACSL1_HUMAN Long-chain-fatty-acid--CoA ligase 1 OS=Homo sapiens GN=A | 92,81 | 7,45 | 77943 | 4 | 23,22 | 23,45 | 20,13 | 22,81 | 23,68 | 22,70 | **1,737** | 5,14E-01 | **0** | 0,7966671 | 0,28888805 |
| RTCA_HUMA | RTCA | sp\|O00442\|RTCA_HUMAN RNA 3'-terminal phosphate cyclase OS=Homo sapiens GN=RT | 65,159 | 8,47 | 39337 | 3 | 21,92 | 21,23 | 23,06 | 22,27 | 22,52 | 23,80 | **1,733** | 3,29E-01 | **0** | 0,79333369 | 0,48336651 |
| K2C5_HUMA | KRT5 | sp\|P13647\|K2C5_HUMAN Keratin, type II cytoskeletal 5 OS=Homo sapiens GN=KRT5 PE= | 1237,15 | 51,86 | 62378 | 43 | 28,08 | 26,94 | 26,24 | 29,29 | 27,67 | 26,68 | **1,733** | 4,42E-01 | **0** | 0,79333369 | 0,35455626 |
| SYLC_HUMA | LARS | sp\|Q9P2J5\|SYLC_HUMAN Leucine--tRNA ligase, cytoplasmic OS=Homo sapiens GN=LARS | 554,102 | 18,11 | 134466 | 18 | 24,78 | 25,35 | 26,12 | 25,89 | 26,05 | 26,69 | **1,733** | 1,59E-01 | **0** | 0,79333242 | 0,79907779 |
| ARFG1_HUM | ARFGAP1 | sp\|Q8N6T3\|ARFG1_HUMAN ADP-ribosylation factor GTPase-activating protein 1 OS=Ho | 13,466 | 3,69 | 44668 | 1 | 19,41 | 17,15 | 21,00 | 19,42 | 19,23 | 21,28 | **1,729** | 5,75E-01 | **0** | 0,79031245 | 0,24057823 |
| K2C1_HUMA | KRT1 | sp\|P04264\|K2C1_HUMAN Keratin, type II cytoskeletal 1 OS=Homo sapiens GN=KRT1 PE= | 3105,999 | 67,55 | 66039 | 58 | 31,81 | 30,58 | 30,02 | 33,13 | 31,44 | 30,20 | **1,725** | 4,76E-01 | **0** | 0,78666751 | 0,32278408 |
| PSB4_HUMA | PSMB4 | sp\|P28070\|PSB4_HUMAN Proteasome subunit beta type-4 OS=Homo sapiens GN=PSME | 212,665 | 21,59 | 29204 | 4 | 23,28 | 23,52 | 23,30 | 23,56 | 24,20 | 24,69 | **1,721** | 8,02E-02 | **0** | 0,78333346 | 1,09594665 |
| MET16_HUN | METTL16 | sp\|Q86W50\|MET16_HUMAN U6 small nuclear RNA (adenine-(43)-N(6))-methyltransfera | 41,478 | 3,2 | 63621 | 1 | 15,80 | 16,27 | 18,59 | 16,45 | 17,60 | 18,96 | **1,721** | 5,25E-01 | **0** | 0,78333314 | 0,27962801 |
| TWF2_HUM/ | TWF2 | sp\|Q6IBS0\|TWF2_HUMAN Twinfilin-2 OS=Homo sapiens GN=TWF2 PE=1 SV=2 | 165,852 | 20,34 | 39548 | 5 | 22,93 | 22,78 | 20,66 | 23,24 | 23,02 | 22,44 | **1,713** | 3,71E-01 | **0** | 0,77666664 | 0,43103589 |
| PTMA_HUM/ | PTMA | sp\|P06454\|PTMA_HUMAN Prothymosin alpha OS=Homo sapiens GN=PTMA PE=1 SV=2 | 133,134 | 21,62 | 12203 | 2 | 21,94 | 21,77 | 22,09 | 22,53 | 22,69 | 22,91 | **1,713** | 5,69E-03 | **0** | 0,77666664 | 2,2450419 |
| SPSY_HUMAI | SMS | sp\|P52788\|SPSY_HUMAN Spermine synthase OS=Homo sapiens GN=SMS PE=1 SV=2 | 182,306 | 26,5 | 41268 | 6 | 23,84 | 23,93 | 20,75 | 24,01 | 24,21 | 22,63 | **1,713** | 5,39E-01 | **0** | 0,77666601 | 0,26846944 |
| HORN_HUM/ | HRNR | sp\|Q86YZ3\|HORN_HUMAN Hornerin OS=Homo sapiens GN=HRNR PE=1 SV=2 | 436,201 | 18,11 | 282390 | 12 | 24,87 | 22,40 | 23,20 | 25,89 | 24,07 | 22,84 | **1,713** | 5,35E-01 | **0** | 0,77666601 | 0,27142457 |
| XRN2_HUMA | XRN2 | sp\|Q9H0D6\|XRN2_HUMAN 5'-3' exoribonuclease 2 OS=Homo sapiens GN=XRN2 PE=1 S\ | 46,906 | 3,89 | 108582 | 3 | 21,59 | 22,04 | 24,33 | 22,86 | 22,76 | 24,66 | **1,709** | 5,02E-01 | **0** | 0,77333323 | 0,29932218 |
| LNP_HUMAN | LNPK | sp\|Q9C0E8\|LNP_HUMAN Endoplasmic reticulum junction formation protein lunapark O | 43,719 | 3,04 | 47740 | 1 | 19,24 | 19,07 | 21,44 | 20,10 | 20,28 | 21,69 | **1,709** | 4,45E-01 | **0** | 0,7731603 | 0,35147957 |
| SC61B_HUM | SEC61B | sp\|P60468\|SC61B_HUMAN Protein transport protein Sec61 subunit beta OS=Homo sapi | 89,699 | 37,5 | 9974 | 3 | 24,11 | 24,30 | 23,36 | 25,04 | 25,20 | 23,84 | **1,705** | 2,10E-01 | **0** | 0,77000046 | 0,67757644 |
| CCD47_HUM | CCDC47 | sp\|Q96A33\|CCD47_HUMAN Coiled-coil domain-containing protein 47 OS=Homo sapiens | 82,517 | 7,25 | 55874 | 3 | 24,13 | 21,79 | 28,29 | 24,90 | 23,94 | 27,68 | **1,705** | 7,45E-01 | **0** | 0,76999982 | 0,12798715 |
| TMF1_HUM/ | TMF1 | sp\|P82094\|TMF1_HUMAN TATA element modulatory factor OS=Homo sapiens GN=TMF | 17,275 | 0,64 | 122842 | 1 | 21,82 | 22,00 | 23,28 | 22,76 | 23,08 | 23,57 | **1,705** | 2,10E-01 | **0** | 0,76999982 | 0,67728852 |
| GOT1B_HUN | GOLT1B | sp\|Q9Y3E0\|GOT1B_HUMAN Vesicle transport protein GOT1B OS=Homo sapiens GN=GC | 121,263 | 10,87 | 15426 | 2 | 21,82 | 21,80 | 22,15 | 22,16 | 22,35 | 23,56 | **1,701** | 1,66E-01 | **0** | 0,76666705 | 0,78056541 |
| LSM8_HUM/ | LSM8 | sp\|O95777\|LSM8_HUMAN U6 snRNA-associated Sm-like protein LSm8 OS=Homo sapier | 74,928 | 36,46 | 10403 | 3 | 21,97 | 22,30 | 22,02 | 22,77 | 22,94 | 22,87 | **1,697** | 2,58E-03 | **0** | 0,76333427 | 2,58819251 |
| TBB4B_HUM | TUBB4B | sp\|P68371\|TBB4B_HUMAN Tubulin beta-4B chain OS=Homo sapiens GN=TUBB4B PE=1 ! | 2709,341 | 84,27 | 49831 | 55 | 25,49 | 25,72 | 26,75 | 26,49 | 26,62 | 27,14 | **1,697** | 1,54E-01 | **0** | 0,76333364 | 0,8113802 |
| NUCL_HUM/ | NCL | sp\|P19338\|NUCL_HUMAN Nucleolin OS=Homo sapiens GN=NCL PE=1 SV=3 | 479,811 | 16,34 | 76614 | 12 | 25,16 | 25,27 | 25,56 | 25,94 | 25,95 | 26,38 | **1,693** | 1,55E-02 | **0** | 0,76000023 | 1,8092723 |
| LRC59_HUM | LRRC59 | sp\|Q96AG4\|LRC59_HUMAN Leucine-rich repeat-containing protein 59 OS=Homo sapien | 143,453 | 16,94 | 34930 | 5 | 24,26 | 24,84 | 25,33 | 25,16 | 25,52 | 26,03 | **1,693** | 1,30E-01 | **0** | 0,76000023 | 0,8872483 |
| CBX3_HUMA | CBX3 | sp\|Q13185\|CBX3_HUMAN Chromobox protein homolog 3 OS=Homo sapiens GN=CBX3 I | 188,855 | 37,16 | 20811 | 5 | 22,36 | 23,36 | 23,67 | 23,50 | 23,95 | 24,22 | **1,693** | 1,65E-01 | **0** | 0,75999959 | 0,78319777 |
| SEC63_HUM | SEC63 | sp\|Q9UGP8\|SEC63_HUMAN Translocation protein SEC63 homolog OS=Homo sapiens GI | 34,079 | 3,03 | 87997 | 2 | 20,39 | 20,95 | 22,28 | 21,42 | 21,97 | 22,51 | **1,693** | 3,03E-01 | **0** | 0,75999959 | 0,5192194 |
| COG5_HUM/ | COG5 | sp\|Q9UP83\|COG5_HUMAN Conserved oligomeric Golgi complex subunit 5 OS=Homo sa | 26,028 | 3,34 | 92743 | 1 | 19,74 | 17,36 | 19,49 | 20,12 | 19,69 | 19,06 | **1,693** | 4,04E-01 | **0** | 0,75927798 | 0,39345548 |
| MTREX_HUN | MTREX | sp\|P42285\|MTREX_HUMAN Exosome RNA helicase MTR4 OS=Homo sapiens GN=MTRE) | 282,253 | 11,32 | 117805 | 9 | 22,40 | 22,48 | 22,98 | 22,80 | 23,23 | 24,10 | **1,690** | 1,48E-01 | **0** | 0,75666682 | 0,82873667 |
| MP2K4_HUN | MAP2K4 | sp\|P45985\|MP2K4_HUMAN Dual specificity mitogen-activated protein kinase kinase 4 O | 51,66 | 10,03 | 44288 | 3 | 20,55 | 21,61 | 20,69 | 21,38 | 21,90 | 21,84 | **1,690** | 1,11E-01 | **0** | 0,75666618 | 0,95514083 |
| DEK_HUMAN | DEK | sp\|P35659\|DEK_HUMAN Protein DEK OS=Homo sapiens GN=DEK PE=1 SV=1 | 103,675 | 8,27 | 42674 | 4 | 22,81 | 24,29 | 24,25 | 24,10 | 24,64 | 24,88 | **1,690** | 2,33E-01 | **0** | 0,75666618 | 0,63299155 |
| H90B4_HUM | HSP90AB4P | sp\|Q58FF6\|H90B4_HUMAN Putative heat shock protein HSP 90-beta 4 OS=Homo sapien | 326,675 | 11,88 | 58264 | 7 | 21,95 | 22,27 | 22,23 | 22,42 | 23,22 | 23,08 | **1,690** | 4,69E-02 | **0** | 0,75666618 | 1,32919943 |
| FUBP3_HUM | FUBP3 | sp\|Q96I24\|FUBP3_HUMAN Far upstream element-binding protein 3 OS=Homo sapiens ( | 46,419 | 3,85 | 61640 | 2 | 20,86 | 21,91 | 22,34 | 22,43 | 21,99 | 22,95 | **1,686** | 2,21E-01 | **0** | 0,75333341 | 0,65590305 |
| ACPH_HUM/ | APEH | sp\|P13798\|ACPH_HUMAN Acylamino-acid-releasing enzyme OS=Homo sapiens GN=APE | 69,422 | 5,19 | 81225 | 4 | 23,15 | 22,78 | 22,00 | 23,53 | 23,90 | 22,76 | **1,686** | 1,89E-01 | **0** | 0,75333341 | 0,72243079 |
| AP1S1_HUM | AP1S1 | sp\|P61966\|AP1S1_HUMAN AP-1 complex subunit sigma-1A OS=Homo sapiens GN=AP1S | 117,541 | 14,56 | 18733 | 2 | 22,03 | 22,20 | 18,93 | 22,32 | 22,17 | 20,92 | **1,682** | 5,50E-01 | **0** | 0,75030454 | 0,25933887 |
| BASI_HUMAI | BSG | sp\|P35613\|BASI_HUMAN Basigin OS=Homo sapiens GN=BSG PE=1 SV=2 | 11,253 | 2,34 | 42200 | 1 | 20,59 | 22,85 | 19,79 | 22,87 | 22,05 | 20,56 | **1,682** | 5,46E-01 | **0** | 0,75028165 | 0,26279224 |
| ERLN2_HUM | ERLIN2 | sp\|O94905\|ERLN2_HUMAN Erlin-2 OS=Homo sapiens GN=ERLIN2 PE=1 SV=1 | 206,149 | 18,29 | 37840 | 6 | 22,57 | 22,63 | 23,01 | 23,66 | 23,47 | 23,33 | **1,682** | 1,11E-02 | **0** | 0,75 | 1,9564546 |
| ROAA_HUM/ | HNRNPAB | sp\|Q99729\|ROAA_HUMAN Heterogeneous nuclear ribonucleoprotein A/B OS=Homo sa | 393,809 | 24,4 | 36225 | 10 | 25,61 | 26,00 | 26,61 | 26,52 | 26,70 | 27,25 | **1,682** | 1,09E-01 | **0** | 0,75 | 0,96347542 |
| UBP7_HUMA | USP7 | sp\|Q93009\|UBP7_HUMAN Ubiquitin carboxyl-terminal hydrolase 7 OS=Homo sapiens G | 300,066 | 16,79 | 128302 | 12 | 22,78 | 23,45 | 24,17 | 23,71 | 24,00 | 24,94 | **1,682** | 2,42E-01 | **0** | 0,74999936 | 0,61614544 |
| ISOC1_HUM/ | ISOC1 | sp\|Q96CN7\|ISOC1_HUMAN Isochorismatase domain-containing protein 1 OS=Homo sap | 55,775 | 6,38 | 32237 | 1 | 17,62 | 17,21 | 18,27 | 18,21 | 18,61 | 18,52 | **1,678** | 8,74E-02 | **0** | 0,74666659 | 1,05832582 |
| SFPQ_HUMA | SFPQ | sp\|P23246\|SFPQ_HUMAN Splicing factor, proline- and glutamine-rich OS=Homo sapiens | 590,73 | 32,53 | 76149 | 18 | 25,71 | 25,77 | 27,36 | 26,69 | 26,49 | 27,90 | **1,678** | 3,44E-01 | **0** | 0,74666659 | 0,46289629 |
| LA_HUMAN | SSB | sp\|P05455\|LA_HUMAN Lupus La protein OS=Homo sapiens GN=SSB PE=1 SV=2 | 555,939 | 36,27 | 46837 | 14 | 24,77 | 25,06 | 25,89 | 25,57 | 25,57 | 26,81 | **1,674** | 2,35E-01 | **0** | 0,74333318 | 0,62860774 |
| ROCK2_HUN | ROCK2 | sp\|O75116\|ROCK2_HUMAN Rho-associated protein kinase 2 OS=Homo sapiens GN=RO( | 55,154 | 3,24 | 160900 | 3 | 20,47 | 21,62 | 20,45 | 21,74 | 21,47 | 21,55 | **1,670** | 1,34E-01 | **0** | 0,73999914 | 0,87213415 |
| UB2E1_HUM | UBE2E1 | sp\|P51965\|UB2E1_HUMAN Ubiquitin-conjugating enzyme E2 E1 OS=Homo sapiens GN= | 11,975 | 5,7 | 21404 | 1 | 18,89 | 18,05 | 20,32 | 18,19 | 20,53 | 20,76 | **1,669** | 5,22E-01 | **0** | 0,73888588 | 0,28193986 |
| TCRG1_HUM | TCERG1 | sp\|O14776\|TCRG1_HUMAN Transcription elongation regulator 1 OS=Homo sapiens GN= | 21,652 | 2,64 | 123901 | 3 | 21,65 | 22,11 | 22,03 | 23,22 | 22,91 | 21,87 | **1,666** | 1,64E-01 | **0** | 0,73666636 | 0,7864528 |
| KCMF1_HUN | KCMF1 | sp\|Q9P0J7\|KCMF1_HUMAN E3 ubiquitin-protein ligase KCMF1 OS=Homo sapiens GN=K | 52,841 | 3,67 | 41945 | 1 | 17,60 | 18,28 | 19,45 | 19,24 | 18,63 | 19,65 | **1,661** | 3,00E-01 | **0** | 0,7316316 | 0,5230654 |
| DJC17_HUM. | DNAJC17 | sp\|Q9NVM6\|DJC17_HUMAN DnaJ homolog subfamily C member 17 OS=Homo sapiens ( | 54,936 | 6,91 | 34687 | 1 | 17,68 | 17,29 | 20,65 | 17,93 | 19,28 | 20,60 | **1,659** | 6,07E-01 | **0** | 0,73000018 | 0,21649846 |
| UBP48_HUM | USP48 | sp\|Q86UV5\|UBP48_HUMAN Ubiquitin carboxyl-terminal hydrolase 48 OS=Homo sapien | 51,059 | 1,64 | 119032 | 1 | 20,64 | 18,43 | 20,75 | 20,34 | 21,14 | 20,52 | **1,655** | 4,12E-01 | **0** | 0,72691282 | 0,38550834 |
| PUF60_HUM | PUF60 | sp\|Q9UHX1\|PUF60_HUMAN Poly(U)-binding-splicing factor PUF60 OS=Homo sapiens GI | 193,804 | 14,13 | 59875 | 5 | 21,60 | 21,58 | 22,69 | 22,53 | 22,40 | 23,12 | **1,655** | 1,65E-01 | **0** | 0,72666677 | 0,78223178 |
| SCYL1_HUM. | SCYL1 | sp\|Q96KG9\|SCYL1_HUMAN N-terminal kinase-like protein OS=Homo sapiens GN=SCYL1 | 10,381 | 1,86 | 89631 | 1 | 20,46 | 21,31 | 20,48 | 21,71 | 21,55 | 21,17 | **1,655** | 8,74E-02 | **0** | 0,72666677 | 1,05845343 |
| K1C14_HUM | KRT14 | sp\|P02533\|K1C14_HUMAN Keratin, type I cytoskeletal 14 OS=Homo sapiens GN=KRT14 | 1291,608 | 64,41 | 51561 | 34 | 26,02 | 25,01 | 24,04 | 27,39 | 25,57 | 24,29 | **1,655** | 5,33E-01 | **0** | 0,72666613 | 0,27347457 |
| LEMD2_HUN | LEMD2 | sp\|Q8NC56\|LEMD2_HUMAN LEM domain-containing protein 2 OS=Homo sapiens GN=L | 46,215 | 4,77 | 56975 | 2 | 21,62 | 23,65 | 22,30 | 24,33 | 23,20 | 22,19 | **1,643** | 4,51E-01 | **0** | 0,71666718 | 0,34577965 |
| CDK5_HUMA | CDK5 | sp\|Q00535\|CDK5_HUMAN Cyclin-dependent-like kinase 5 OS=Homo sapiens GN=CDK5 I | 22,403 | 10,62 | 33304 | 3 | 19,98 | 20,59 | 19,13 | 19,22 | 20,89 | 21,73 | **1,640** | 4,49E-01 | **0** | 0,71408335 | 0,3481426 |
| NOLC1_HUN | NOLC1 | sp\|Q14978\|NOLC1_HUMAN Nucleolar and coiled-body phosphoprotein 1 OS=Homo sap | 92,788 | 3,58 | 73603 | 2 | 21,98 | 21,89 | 22,04 | 22,70 | 22,58 | 22,77 | **1,640** | 5,39E-04 | **0** | 0,71333377 | 3,26874477 |
| ACATN_HUN | SLC33A1 | sp\|O00400\|ACATN_HUMAN Acetyl-coenzyme A transporter 1 OS=Homo sapiens GN=SL | 37 | 5,65 | 60909 | 2 | 16,83 | 17,90 | 19,25 | 17,54 | 18,44 | 20,14 | **1,640** | 5,29E-01 | **0** | 0,71333377 | 0,27686917 |
| SPRR3_HUM. | SPRR3 | sp\|Q9UBC9\|SPRR3_HUMAN Small proline-rich protein 3 OS=Homo sapiens GN=SPRR3 P | 29,393 | 9,47 | 18154 | 1 | 19,08 | 19,22 | 18,63 | 19,55 | 18,56 | 20,95 | **1,637** | 3,77E-01 | **0** | 0,71086693 | 0,42354761 |
| CO9_HUMAN | C9 | sp\|P02748\|CO9_HUMAN Complement component C9 OS=Homo sapiens GN=C9 PE=1 S\ | 31,306 | 1,61 | 63173 | 1 | 23,68 | 18,52 | 23,79 | 23,86 | 21,03 | 23,23 | **1,636** | 7,33E-01 | **0** | 0,70999972 | 0,13504428 |
| FKBP4_HUM | FKBP4 | sp\|Q02790\|FKBP4_HUMAN Peptidyl-prolyl cis-trans isomerase FKBP4 OS=Homo sapiens | 248,228 | 16,12 | 51805 | 10 | 25,65 | 25,36 | 26,12 | 26,30 | 26,10 | 26,86 | **1,636** | 8,90E-02 | **0** | 0,70999972 | 1,0508484 |
| FKB11_HUM. | FKBP11 | sp\|Q9NYL4\|FKB11_HUMAN Peptidyl-prolyl cis-trans isomerase FKBP11 OS=Homo sapier | 73,741 | 14,43 | 22180 | 2 | 22,31 | 22,95 | 22,41 | 23,26 | 23,16 | 23,38 | **1,636** | 2,72E-02 | **0** | 0,70999972 | 1,56500288 |
| TPRKB_HUM | TPRKB | sp\|Q9Y3C4\|TPRKB_HUMAN EKC/KEOPS complex subunit TPRKB OS=Homo sapiens GN=' | 12,28 | 4 | 19661 | 1 | 20,30 | 20,69 | 20,20 | 20,96 | 20,97 | 21,39 | **1,636** | 2,61E-02 | **0** | 0,70999908 | 1,58298911 |
| TBCA_HUMA | TBCA | sp\|O75347\|TBCA_HUMAN Tubulin-specific chaperone A OS=Homo sapiens GN=TBCA PE | 57,104 | 15,74 | 12855 | 2 | 24,25 | 23,85 | 20,02 | 24,40 | 24,02 | 21,82 | **1,634** | 6,76E-01 | **0** | 0,70801862 | 0,17032283 |
| PRP19_HUM. | PRPF19 | sp\|Q9UMS4\|PRP19_HUMAN Pre-mRNA-processing factor 19 OS=Homo sapiens GN=PRF | 422,313 | 52,98 | 55181 | 14 | 24,39 | 24,63 | 26,29 | 25,05 | 25,55 | 26,83 | **1,632** | 4,26E-01 | **0** | 0,70666631 | 0,37037375 |
| S23IP_HUM/ | SEC23IP | sp\|Q9Y6Y8\|S23IP_HUMAN SEC23-interacting protein OS=Homo sapiens GN=SEC23IP PE | 188,958 | 10,8 | 111076 | 7 | 22,76 | 23,72 | 23,85 | 23,91 | 24,20 | 24,33 | **1,628** | 1,27E-01 | **0** | 0,70333354 | 0,89754594 |
| CDC73_HUM | CDC73 | sp\|Q6P1J9\|CDC73_HUMAN Parafibromin OS=Homo sapiens GN=CDC73 PE=1 SV=1 | 23,122 | 3,77 | 60577 | 1 | 17,18 | 16,68 | 18,57 | 17,63 | 17,62 | 19,28 | **1,625** | 4,26E-01 | **0** | 0,70000013 | 0,37099041 |
| RUXGL_HUN | SNRPGP15 | sp\|A8MWD9\|RUXGL_HUMAN Putative small nuclear ribonucleoprotein G-like protein 1! | 62,294 | 26,32 | 8544 | 2 | 23,52 | 21,75 | 23,12 | 24,03 | 23,62 | 22,84 | **1,625** | 3,35E-01 | **0** | 0,70000013 | 0,47468075 |
| LOXL2_HUM | LOXL2 | sp\|Q9Y4K0\|LOXL2_HUMAN Lysyl oxidase homolog 2 OS=Homo sapiens GN=LOXL2 PE=1 | 169,169 | 6,07 | 86725 | 5 | 22,34 | 20,53 | 24,46 | 22,75 | 22,22 | 24,46 | **1,625** | 6,24E-01 | **0** | 0,69999949 | 0,2045547 |
| DPOD1_HUN | POLD1 | sp\|P28340\|DPOD1_HUMAN DNA polymerase delta catalytic subunit OS=Homo sapiens ( | 227,683 | 8,13 | 123631 | 7 | 21,97 | 22,34 | 22,63 | 22,77 | 23,01 | 23,25 | **1,621** | 4,19E-02 | **0** | 0,69666735 | 1,37813175 |
| HNRPF_HUM | HNRNPF | sp\|P52597\|HNRPF_HUMAN Heterogeneous nuclear ribonucleoprotein F OS=Homo sapi | 494,011 | 36,63 | 45672 | 12 | 25,37 | 24,90 | 26,35 | 26,00 | 26,15 | 26,56 | **1,621** | 2,03E-01 | **0** | 0,69666608 | 0,69150154 |
| OSTC_HUMA | OSTC | sp\|Q9NRP0\|OSTC_HUMAN Oligosaccharyltransferase complex subunit OSTC OS=Homo | 60,725 | 8,05 | 16829 | 1 | 22,53 | 22,19 | 23,71 | 23,14 | 23,31 | 24,07 | **1,621** | 2,68E-01 | **0** | 0,69666608 | 0,57167535 |
| POSTN_HUN | POSTN | sp\|Q15063\|POSTN_HUMAN Periostin OS=Homo sapiens GN=POSTN PE=1 SV=2 | 81,685 | 6,46 | 93314 | 5 | 20,08 | 17,57 | 24,46 | 18,55 | 21,47 | 24,18 | **1,618** | 8,02E-01 | **0** | 0,69438299 | 0,09596714 |
| ARFRP_HUM | ARFRP1 | sp\|Q13795\|ARFRP_HUMAN ADP-ribosylation factor-related protein 1 OS=Homo sapiens | 44,094 | 4,98 | 22614 | 1 | 18,70 | 18,36 | 19,83 | 19,43 | 19,22 | 20,33 | **1,618** | 2,82E-01 | **0** | 0,6942927 | 0,54925398 |
| MYPT1_HUN | PPP1R12A | sp\|O14974\|MYPT1_HUMAN Protein phosphatase 1 regulatory subunit 12A OS=Homo sa | 203,205 | 6,31 | 115281 | 6 | 22,13 | 20,92 | 23,56 | 22,65 | 22,35 | 23,69 | **1,617** | 4,67E-01 | **0** | 0,69333394 | 0,33032333 |
| CTR9_HUMA | CTR9 | sp\|Q6PD62\|CTR9_HUMAN RNA polymerase-associated protein CTR9 homolog OS=Hom | 49,565 | 1,71 | 133502 | 2 | 19,77 | 18,57 | 21,66 | 20,78 | 19,74 | 21,56 | **1,617** | 5,42E-01 | **0** | 0,69333331 | 0,26568629 |
| NAA10_HUN | NAA10 | sp\|P41227\|NAA10_HUMAN N-alpha-acetyltransferase 10 OS=Homo sapiens GN=NAA10 | 112,857 | 29,36 | 26459 | 5 | 22,44 | 22,04 | 23,72 | 22,88 | 22,84 | 24,56 | **1,617** | 4,13E-01 | **0** | 0,69333267 | 0,38366977 |
| PSMD8_HUN | PSMD8 | sp\|P48556\|PSMD8_HUMAN 26S proteasome non-ATPase regulatory subunit 8 OS=Hom | 327,976 | 31,43 | 39612 | 10 | 24,78 | 24,79 | 26,02 | 25,40 | 25,55 | 26,72 | **1,617** | 3,02E-01 | **0** | 0,69333204 | 0,51945839 |
| CLPT1_HUM. | CLPTM1 | sp\|O96005\|CLPT1_HUMAN Cleft lip and palate transmembrane protein 1 OS=Homo sap | 199,916 | 9,87 | 76097 | 6 | 22,76 | 22,55 | 23,17 | 23,30 | 23,40 | 23,85 | **1,613** | 5,00E-02 | **0** | 0,6899999 | 1,30109041 |
| AMPN_HUM | ANPEP | sp\|P15144\|AMPN_HUMAN Aminopeptidase N OS=Homo sapiens GN=ANPEP PE=1 SV=4 | 40,413 | 2,38 | 109540 | 2 | 24,24 | 23,42 | 22,69 | 25,96 | 22,84 | 23,62 | **1,613** | 5,43E-01 | **0** | 0,6899999 | 0,26528086 |
| VAPB_HUMA | VAPB | sp\|O95292\|VAPB_HUMAN Vesicle-associated membrane protein-associated protein B/( | 84,261 | 18,93 | 27228 | 4 | 22,83 | 23,27 | 22,82 | 23,89 | 23,95 | 23,15 | **1,613** | 8,08E-02 | **0** | 0,6899999 | 1,09244733 |
| COPE_HUMA | COPE | sp\|O14579\|COPE_HUMAN Coatomer subunit epsilon OS=Homo sapiens GN=COPE PE=1 | 72,772 | 11,36 | 34482 | 3 | 22,13 | 22,21 | 22,75 | 22,22 | 23,39 | 23,55 | **1,613** | 2,10E-01 | **0** | 0,6899999 | 0,67822456 |
| SC22B_HUM. | SEC22B | sp\|O75396\|SC22B_HUMAN Vesicle-trafficking protein SEC22b OS=Homo sapiens GN=SE | 207,276 | 28,37 | 24593 | 5 | 23,09 | 23,24 | 24,36 | 24,22 | 23,98 | 24,56 | **1,613** | 1,88E-01 | **0** | 0,68999926 | 0,72691676 |
| IL1AP_HUMA | IL1RAP | sp\|Q9NPH3\|IL1AP_HUMAN Interleukin-1 receptor accessory protein OS=Homo sapiens | 28,803 | 1,93 | 65418 | 1 | 20,11 | 18,46 | 19,03 | 20,44 | 19,92 | 19,29 | **1,610** | 3,07E-01 | **0** | 0,68726285 | 0,5127889 |
| NAA15_HUN | NAA15 | sp\|Q9BXJ9\|NAA15_HUMAN N-alpha-acetyltransferase 15, NatA auxiliary subunit OS=Ho | 503,122 | 26,67 | 101272 | 18 | 23,49 | 23,42 | 25,21 | 23,94 | 24,39 | 25,85 | **1,610** | 4,50E-01 | **0** | 0,68666712 | 0,34651052 |
| HNRL2_HUM | HNRNPUL2 | sp\|Q1KMD3\|HNRL2_HUMAN Heterogeneous nuclear ribonucleoprotein U-like protein 2 | 300,647 | 18,21 | 85105 | 11 | 24,31 | 24,18 | 25,85 | 24,98 | 25,13 | 26,29 | **1,610** | 3,68E-01 | **0** | 0,68666649 | 0,43401244 |
| MBOA7_HUI | MBOAT7 | sp\|Q96N66\|MBOA7_HUMAN Lysophospholipid acyltransferase 7 OS=Homo sapiens GN | 92,078 | 8,26 | 52765 | 2 | 19,59 | 21,00 | 22,02 | 20,84 | 21,44 | 22,39 | **1,610** | 4,58E-01 | **0** | 0,68666649 | 0,339256 |
| H2AY_HUMA | H2AFY | sp\|O75367\|H2AY_HUMAN Core histone macro-H2A.1 OS=Homo sapiens GN=H2AFY PE= | 38,394 | 5,91 | 39617 | 1 | 16,84 | 17,61 | 19,81 | 18,44 | 17,78 | 20,09 | **1,608** | 5,75E-01 | **0** | 0,68537585 | 0,23996269 |
| AN32E_HUM | ANP32E | sp\|Q9BTT0\|AN32E_HUMAN Acidic leucine-rich nuclear phosphoprotein 32 family memb | 198,046 | 18,28 | 30692 | 4 | 23,09 | 22,55 | 25,06 | 23,97 | 24,04 | 24,74 | **1,606** | 4,42E-01 | **0** | 0,68333371 | 0,3546885 |
| PRAF2_HUM | PRAF2 | sp\|O60831\|PRAF2_HUMAN PRA1 family protein 2 OS=Homo sapiens GN=PRAF2 PE=1 SV | 83,603 | 10,67 | 19258 | 2 | 21,24 | 22,14 | 22,65 | 22,00 | 23,03 | 23,05 | **1,606** | 2,73E-01 | **0** | 0,68333371 | 0,56326243 |
| RL31_HUMA | RPL31 | sp\|P62899\|RL31_HUMAN 60S ribosomal protein L31 OS=Homo sapiens GN=RPL31 PE=1 | 52,825 | 30,4 | 14463 | 4 | 24,14 | 22,48 | 23,51 | 24,66 | 23,19 | 24,32 | **1,602** | 3,59E-01 | **0** | 0,68000031 | 0,44484112 |
| PSN1_HUMA | PSEN1 | sp\|P49768\|PSN1_HUMAN Presenilin-1 OS=Homo sapiens GN=PSEN1 PE=1 SV=1 | 109,102 | 5,14 | 52668 | 2 | 22,39 | 22,11 | 19,72 | 23,07 | 22,98 | 20,21 | **1,602** | 6,19E-01 | **0** | 0,67999967 | 0,20812813 |
| RPC3_HUMA | POLR3C | sp\|Q9BUI4\|RPC3_HUMAN DNA-directed RNA polymerase III subunit RPC3 OS=Homo sap | 27,022 | 1,87 | 60612 | 1 | 20,29 | 19,34 | 18,15 | 19,83 | 19,76 | 20,22 | **1,600** | 3,47E-01 | **0** | 0,67821312 | 0,45987569 |
| RBP10_HUM | RANBP10 | sp\|Q6VN20\|RBP10_HUMAN Ran-binding protein 10 OS=Homo sapiens GN=RANBP10 PE | 32,013 | 1,61 | 67257 | 1 | 19,15 | 18,00 | 20,97 | 19,90 | 19,19 | 21,06 | **1,599** | 5,44E-01 | **0** | 0,67687225 | 0,26435455 |
| PSA2_HUMA | PSMA2 | sp\|P25787\|PSA2_HUMAN Proteasome subunit alpha type-2 OS=Homo sapiens GN=PSM | 502,151 | 61,11 | 25899 | 13 | 25,31 | 25,18 | 25,64 | 25,86 | 26,05 | 26,25 | **1,598** | 1,88E-02 | **0** | 0,6766669 | 1,72534108 |
| ZFR_HUMAN | ZFR | sp\|Q96KR1\|ZFR_HUMAN Zinc finger RNA-binding protein OS=Homo sapiens GN=ZFR PE: | 63,969 | 1,49 | 117012 | 1 | 20,03 | 17,72 | 17,65 | 18,01 | 18,80 | 20,60 | **1,595** | 5,72E-01 | **0** | 0,67367744 | 0,24274624 |
| FBRL_HUMA | FBL | sp\|P22087\|FBRL_HUMAN rRNA 2'-O-methyltransferase fibrillarin OS=Homo sapiens GN: | 322,31 | 47,98 | 33784 | 11 | 22,80 | 23,09 | 25,80 | 23,55 | 23,77 | 26,39 | **1,595** | 6,37E-01 | **0** | 0,67333349 | 0,19581637 |
| NCBP1_HUM | NCBP1 | sp\|Q09161\|NCBP1_HUMAN Nuclear cap-binding protein subunit 1 OS=Homo sapiens GI | 326,782 | 22,78 | 91839 | 12 | 22,69 | 22,69 | 24,41 | 23,02 | 23,79 | 25,00 | **1,595** | 4,54E-01 | **0** | 0,67333349 | 0,34289683 |
| AN32A_HUN | ANP32A | sp\|P39687\|AN32A_HUMAN Acidic leucine-rich nuclear phosphoprotein 32 family memb | 70,358 | 14,06 | 28585 | 4 | 22,76 | 23,27 | 23,21 | 23,98 | 23,68 | 23,60 | **1,595** | 2,73E-02 | **0** | 0,67333349 | 1,56319095 |
| SARNP_HUM | SARNP | sp\|P82979\|SARNP_HUMAN SAP domain-containing ribonucleoprotein OS=Homo sapien | 36,313 | 9,05 | 23671 | 1 | 17,61 | 18,89 | 18,41 | 17,78 | 19,39 | 19,76 | **1,592** | 4,01E-01 | **0** | 0,67055257 | 0,39727025 |
| DHR11_HUM | DHRS11 | sp\|Q6UWP2\|DHR11_HUMAN Dehydrogenase/reductase SDR family member 11 OS=Hoi | 51,973 | 5 | 28308 | 1 | 21,58 | 21,90 | 19,77 | 21,82 | 21,87 | 21,56 | **1,589** | 3,76E-01 | **0** | 0,66779582 | 0,42474146 |
| LC7L3_HUM/ | LUC7L3 | sp\|O95232\|LC7L3_HUMAN Luc7-like protein 3 OS=Homo sapiens GN=LUC7L3 PE=1 SV=: | 181,891 | 7,41 | 51466 | 2 | 17,60 | 17,76 | 21,19 | 17,52 | 19,43 | 21,60 | **1,587** | 7,09E-01 | **0** | 0,66666667 | 0,14952011 |
| GUAA_HUM/ | GMPS | sp\|P49915\|GUAA_HUMAN GMP synthase [glutamine-hydrolyzing] OS=Homo sapiens GI | 291,958 | 23,67 | 76715 | 12 | 21,97 | 21,75 | 24,42 | 22,68 | 22,48 | 24,98 | **1,587** | 6,00E-01 | **0** | 0,66666667 | 0,22174758 |
| ORML1_HUN | ORMDL1 | sp\|Q9P0S3\|ORML1_HUMAN ORM1-like protein 1 OS=Homo sapiens GN=ORMDL1 PE=1 | 26,478 | 7,19 | 17371 | 1 | 20,46 | 21,22 | 18,02 | 21,06 | 21,14 | 19,49 | **1,583** | 5,81E-01 | **0** | 0,66307322 | 0,23607166 |
| FBF1_HUMA | FBF1 | sp\|Q8TES7\|FBF1_HUMAN Fas-binding factor 1 OS=Homo sapiens GN=FBF1 PE=1 SV=2 | 9,622 | 0,88 | 125446 | 1 | 18,89 | 19,41 | 21,28 | 20,29 | 19,81 | 21,46 | **1,580** | 4,93E-01 | **0** | 0,65999985 | 0,30720258 |
| HMOX2_HUI | HMOX2 | sp\|P30519\|HMOX2_HUMAN Heme oxygenase 2 OS=Homo sapiens GN=HMOX2 PE=1 SV | 58,851 | 8,86 | 36033 | 2 | 22,37 | 23,32 | 22,91 | 23,25 | 23,76 | 23,57 | **1,580** | 1,02E-01 | **0** | 0,65999985 | 0,98941885 |
| BLMH_HUM/ | BLMH | sp\|Q13867\|BLMH_HUMAN Bleomycin hydrolase OS=Homo sapiens GN=BLMH PE=1 SV= | 158,795 | 13,19 | 52562 | 7 | 24,50 | 24,32 | 24,59 | 25,44 | 24,89 | 25,06 | **1,580** | 2,18E-02 | **0** | 0,65999985 | 1,66132284 |
| LAMB1_HUN | LAMB1 | sp\|P07942\|LAMB1_HUMAN Laminin subunit beta-1 OS=Homo sapiens GN=LAMB1 PE=1 | 183,361 | 6,05 | 198038 | 10 | 24,28 | 21,09 | 24,83 | 24,25 | 22,79 | 25,14 | **1,580** | 6,51E-01 | **0** | 0,65999985 | 0,18639722 |
| RS6_HUMAN | RPS6 | sp\|P62753\|RS6_HUMAN 40S ribosomal protein S6 OS=Homo sapiens GN=RPS6 PE=1 SV= | 483,117 | 29,72 | 28681 | 11 | 25,68 | 26,12 | 26,37 | 26,49 | 26,32 | 27,34 | **1,580** | 1,53E-01 | **0** | 0,65999921 | 0,81597224 |
| EOGT_HUM/ | EOGT | sp\|Q5NDL2\|EOGT_HUMAN EGF domain-specific O-linked N-acetylglucosamine transfera | 28,56 | 2,09 | 62011 | 1 | 15,23 | 14,07 | 19,63 | 17,01 | 15,91 | 17,98 | **1,576** | 7,33E-01 | **0** | 0,65666707 | 0,13481037 |
| SPS1_HUMA | SEPHS1 | sp\|P49903\|SPS1_HUMAN Selenide, water dikinase 1 OS=Homo sapiens GN=SEPHS1 PE= | 65,947 | 17,6 | 42911 | 4 | 22,59 | 22,46 | 21,13 | 22,51 | 23,47 | 22,17 | **1,576** | 3,41E-01 | **0** | 0,65666707 | 0,46776184 |
| GSTM3_HUN | GSTM3 | sp\|P21266\|GSTM3_HUMAN Glutathione S-transferase Mu 3 OS=Homo sapiens GN=GST | 113,502 | 16,44 | 26560 | 5 | 24,16 | 24,81 | 23,97 | 24,89 | 25,06 | 24,96 | **1,576** | 6,43E-02 | **0** | 0,65666644 | 1,19156038 |
| RBBP5_HUM | RBBP5 | sp\|Q15291\|RBBP5_HUMAN Retinoblastoma-binding protein 5 OS=Homo sapiens GN=RE | 23,033 | 1,67 | 59153 | 1 | 18,54 | 19,62 | 20,47 | 19,66 | 19,76 | 21,17 | **1,574** | 4,28E-01 | **0** | 0,65475273 | 0,3690113 |
| HDAC2_HUN | HDAC2 | sp\|Q92769\|HDAC2_HUMAN Histone deacetylase 2 OS=Homo sapiens GN=HDAC2 PE=1 ! | 327,581 | 29,51 | 55364 | 13 | 22,76 | 22,06 | 23,98 | 22,64 | 23,46 | 24,66 | **1,573** | 4,66E-01 | **0** | 0,65333303 | 0,33164429 |
| SPCS2_HUM. | SPCS2 | sp\|Q15005\|SPCS2_HUMAN Signal peptidase complex subunit 2 OS=Homo sapiens GN=S | 75,247 | 19,47 | 25003 | 4 | 23,42 | 22,02 | 24,59 | 23,98 | 23,03 | 24,98 | **1,573** | 5,22E-01 | **0** | 0,65333303 | 0,28234387 |
| PRS8_HUMA | PSMC5 | sp\|P62195\|PRS8_HUMAN 26S proteasome regulatory subunit 8 OS=Homo sapiens GN=F | 861,752 | 59,36 | 45626 | 23 | 25,37 | 25,58 | 27,13 | 26,12 | 26,25 | 27,66 | **1,569** | 4,31E-01 | **0** | 0,65000025 | 0,36581843 |
| ABD12_HUM | ABHD12 | sp\|Q8N2K0\|ABD12_HUMAN Monoacylglycerol lipase ABHD12 OS=Homo sapiens GN=AE | 60,42 | 6,78 | 45097 | 2 | 19,84 | 19,55 | 21,51 | 20,43 | 20,46 | 21,96 | **1,569** | 4,58E-01 | **0** | 0,64999962 | 0,33894886 |
| RAB31_HUM | RAB31 | sp\|Q13636\|RAB31_HUMAN Ras-related protein Rab-31 OS=Homo sapiens GN=RAB31 PI | 217,168 | 30,41 | 21569 | 5 | 23,76 | 22,99 | 21,10 | 24,05 | 23,63 | 22,12 | **1,569** | 5,45E-01 | **0** | 0,64999962 | 0,26366765 |
| U520_HUMA | SNRNP200 | sp\|O75643\|U520_HUMAN U5 small nuclear ribonucleoprotein 200 kDa helicase OS=Hor | 448,571 | 8,05 | 244508 | 12 | 22,79 | 23,46 | 23,77 | 23,37 | 24,14 | 24,46 | **1,569** | 2,08E-01 | **0** | 0,64999962 | 0,68093716 |
| OSBP2_HUM | OSBP2 | sp\|Q969R2\|OSBP2_HUMAN Oxysterol-binding protein 2 OS=Homo sapiens GN=OSBP2 F | 40,384 | 2,62 | 101266 | 2 | 19,13 | 20,92 | 21,89 | 20,89 | 21,19 | 21,80 | **1,566** | 4,90E-01 | **0** | 0,64677747 | 0,3099032 |
| SMC1A_HUN | SMC1A | sp\|Q14683\|SMC1A_HUMAN Structural maintenance of chromosomes protein 1A OS=Hc | 303,489 | 14,27 | 143233 | 17 | 22,91 | 24,42 | 25,07 | 24,44 | 24,52 | 25,38 | **1,566** | 4,12E-01 | **0** | 0,64666684 | 0,38496605 |
| SYVC_HUMA | VARS | sp\|P26640\|SYVC_HUMAN Valine--tRNA ligase OS=Homo sapiens GN=VARS PE=1 SV=4 | 600,936 | 15,9 | 140476 | 15 | 24,32 | 24,92 | 25,66 | 25,10 | 25,47 | 26,27 | **1,566** | 2,81E-01 | **0** | 0,64666684 | 0,55162756 |
| IDE_HUMAN | IDE | sp\|P14735\|IDE_HUMAN Insulin-degrading enzyme OS=Homo sapiens GN=IDE PE=1 SV=4 | 171,857 | 13,05 | 117968 | 12 | 24,70 | 24,79 | 22,74 | 25,01 | 25,12 | 24,04 | **1,566** | 4,38E-01 | **0** | 0,64666684 | 0,35836758 |
| MSTO1_HUN | MSTO1 | sp\|Q9BUK6\|MSTO1_HUMAN Protein misato homolog 1 OS=Homo sapiens GN=MSTO1 F | 37,756 | 2,11 | 61835 | 1 | 20,86 | 21,23 | 21,11 | 22,53 | 21,31 | 21,29 | **1,562** | 2,04E-01 | **0** | 0,64333344 | 0,69030166 |
| HDAC1_HUN | HDAC1 | sp\|Q13547\|HDAC1_HUMAN Histone deacetylase 1 OS=Homo sapiens GN=HDAC1 PE=1 ! | 289,917 | 21,58 | 55103 | 9 | 20,92 | 21,79 | 23,26 | 21,98 | 22,26 | 23,66 | **1,562** | 4,95E-01 | **0** | 0,6433328 | 0,30529459 |
| UB2L3_HUM | UBE2L3 | sp\|P68036\|UB2L3_HUMAN Ubiquitin-conjugating enzyme E2 L3 OS=Homo sapiens GN=I | 281,436 | 69,48 | 17862 | 9 | 26,01 | 26,09 | 23,96 | 26,12 | 26,57 | 25,29 | **1,558** | 4,64E-01 | **0** | 0,64000066 | 0,33340656 |
| PSB5_HUMA | PSMB5 | sp\|P28074\|PSB5_HUMAN Proteasome subunit beta type-5 OS=Homo sapiens GN=PSME | 473,016 | 41,83 | 28480 | 10 | 25,66 | 25,78 | 25,16 | 26,07 | 26,28 | 26,16 | **1,555** | 3,31E-02 | **0** | 0,63666662 | 1,48012024 |
| SSRG_HUMA | SSR3 | sp\|Q9UNL2\|SSRG_HUMAN Translocon-associated protein subunit gamma OS=Homo sa | 63,116 | 7,57 | 21080 | 1 | 21,69 | 22,45 | 23,11 | 22,51 | 22,96 | 23,69 | **1,555** | 3,00E-01 | **0** | 0,63666598 | 0,52280001 |
| RFC4_HUMA | RFC4 | sp\|P35249\|RFC4_HUMAN Replication factor C subunit 4 OS=Homo sapiens GN=RFC4 PE: | 12,029 | 2,48 | 39682 | 1 | 21,13 | 19,59 | 22,01 | 20,96 | 21,34 | 22,34 | **1,553** | 4,80E-01 | **0** | 0,63509687 | 0,31844467 |
| HNRPU_HUN | HNRNPU | sp\|Q00839\|HNRPU_HUMAN Heterogeneous nuclear ribonucleoprotein U OS=Homo sap | 1317,794 | 42,42 | 90584 | 38 | 26,30 | 27,00 | 29,60 | 27,42 | 27,69 | 29,69 | **1,551** | 6,35E-01 | **0** | 0,63333384 | 0,19752072 |
| DHX29_HUM | DHX29 | sp\|Q7Z478\|DHX29_HUMAN ATP-dependent RNA helicase DHX29 OS=Homo sapiens GN | 190,422 | 10,59 | 155236 | 10 | 23,01 | 22,34 | 24,44 | 23,25 | 23,33 | 25,11 | **1,551** | 5,06E-01 | **0** | 0,63333321 | 0,2961147 |
| CTBP2_HUM | CTBP2 | sp\|P56545\|CTBP2_HUMAN C-terminal-binding protein 2 OS=Homo sapiens GN=CTBP2 F | 153,461 | 24,04 | 48945 | 7 | 21,06 | 21,44 | 24,15 | 21,90 | 22,25 | 24,40 | **1,551** | 6,39E-01 | **0** | 0,63333321 | 0,19483889 |
| SF01_HUMA | SF1 | sp\|Q15637\|SF01_HUMAN Splicing factor 1 OS=Homo sapiens GN=SF1 PE=1 SV=4 | 157,413 | 8,92 | 68330 | 4 | 21,64 | 20,79 | 22,86 | 22,53 | 21,57 | 23,09 | **1,551** | 4,44E-01 | **0** | 0,63333321 | 0,35239742 |
| RFC2_HUMA | RFC2 | sp\|P35250\|RFC2_HUMAN Replication factor C subunit 2 OS=Homo sapiens GN=RFC2 PE: | 66 | 8,47 | 39157 | 2 | 20,88 | 20,60 | 22,69 | 21,40 | 21,57 | 23,10 | **1,551** | 4,97E-01 | **0** | 0,63333321 | 0,30341676 |

| | | | | | | | | | | | | | | | | |
|---|---|---|---|---|---|---|---|---|---|---|---|---|---|---|---|---|
| FUS_HUMAN | FUS | sp\|P35637\|FUS_HUMAN RNA-binding protein FUS OS=Homo sapiens GN=FUS PE=1 SV= | 119,959 | 5,7 | 53426 | 3 | 22,33 | 22,55 | 22,69 | 23,05 | 23,32 | 23,09 | 1,548 | 9,39E-03 | 0 | 0,6299998 | 2,02741994 |
| TALDO_HUM | TALDO1 | sp\|P37837\|TALDO_HUMAN Transaldolase OS=Homo sapiens GN=TALDO1 PE=1 SV=2 | 210,018 | 19,88 | 37540 | 8 | 25,36 | 25,59 | 24,15 | 25,73 | 25,89 | 25,37 | 1,548 | 2,53E-01 | 0 | 0,6299998 | 0,59659006 |
| DSRAD_HUM | ADAR | sp\|P55265\|DSRAD_HUMAN Double-stranded RNA-specific adenosine deaminase OS=Ho | 118,826 | 5,3 | 136066 | 5 | 21,27 | 21,37 | 24,08 | 22,48 | 22,16 | 23,97 | 1,548 | 5,90E-01 | 0 | 0,62999916 | 0,22934901 |
| OR6N2_HUM | OR6N2 | sp\|Q8NGY6\|OR6N2_HUMAN Olfactory receptor 6N2 OS=Homo sapiens GN=OR6N2 PE= | 11,493 | 2,21 | 35692 | 1 | 21,82 | 21,84 | 17,76 | 21,69 | 21,28 | 20,33 | 1,544 | 6,80E-01 | 0 | 0,62701925 | 0,16718665 |
| HMGB1_HUM | HMGB1 | sp\|P09429\|HMGB1_HUMAN High mobility group protein B1 OS=Homo sapiens GN=HM | 317,366 | 37,67 | 24894 | 9 | 25,14 | 25,33 | 25,63 | 25,94 | 25,98 | 26,06 | 1,544 | 1,30E-02 | 0 | 0,62666702 | 1,88600894 |
| 6PGD_HUMA | PGD | sp\|P52209\|6PGD_HUMAN 6-phosphogluconate dehydrogenase, decarboxylating OS=Ho | 384,829 | 24,22 | 53140 | 11 | 26,10 | 26,27 | 24,13 | 26,32 | 26,50 | 25,56 | 1,544 | 4,47E-01 | 0 | 0,62666639 | 0,34925869 |
| UBP19_HUM | USP19 | sp\|O94966\|UBP19_HUMAN Ubiquitin carboxyl-terminal hydrolase 19 OS=Homo sapiens | 99,103 | 3,26 | 145651 | 3 | 19,96 | 20,47 | 21,55 | 20,39 | 21,15 | 22,31 | 1,540 | 4,41E-01 | 0 | 0,62333361 | 0,35588186 |
| SRPRB_HUM | SRPRB | sp\|Q9Y5M8\|SRPRB_HUMAN Signal recognition particle receptor subunit beta OS=Homo | 117,11 | 15,87 | 29702 | 3 | 22,74 | 22,93 | 23,18 | 23,27 | 23,71 | 23,74 | 1,540 | 3,47E-02 | 0 | 0,62333298 | 1,45943814 |
| PARP1_HUM | PARP1 | sp\|P09874\|PARP1_HUMAN Poly [ADP-ribose] polymerase 1 OS=Homo sapiens GN=PARI | 81,296 | 5,92 | 113084 | 4 | 22,05 | 22,35 | 22,22 | 22,81 | 22,64 | 23,03 | 1,537 | 1,21E-02 | 0 | 0,6200002 | 1,91609887 |
| KS6A3_HUM | RPS6KA3 | sp\|P51812\|KS6A3_HUMAN Ribosomal protein S6 kinase alpha-3 OS=Homo sapiens GN= | 104,868 | 5,54 | 83736 | 4 | 21,90 | 22,94 | 22,79 | 23,14 | 22,74 | 23,61 | 1,537 | 2,06E-01 | 0 | 0,61999957 | 0,68714324 |
| UBE4B_HUM | UBE4B | sp\|O95155\|UBE4B_HUMAN Ubiquitin conjugation factor E4 B OS=Homo sapiens GN=UE | 56,622 | 2,53 | 146185 | 2 | 18,45 | 19,23 | 20,23 | 18,88 | 19,64 | 21,25 | 1,537 | 5,15E-01 | 0 | 0,61999957 | 0,28857656 |
| EI2BA_HUM | EIF2B1 | sp\|Q14232\|EI2BA_HUMAN Translation initiation factor eIF-2B subunit alpha OS=Homo s | 144,09 | 20,98 | 33712 | 5 | 23,15 | 23,00 | 21,41 | 23,33 | 23,52 | 22,56 | 1,533 | 3,83E-01 | 0 | 0,61666679 | 0,41723171 |
| CSN3_HUMA | COPS3 | sp\|Q9UNS2\|CSN3_HUMAN COP9 signalosome complex subunit 3 OS=Homo sapiens GN | 243,865 | 21,99 | 47873 | 7 | 23,64 | 23,44 | 24,33 | 24,17 | 24,36 | 24,73 | 1,533 | 1,23E-01 | 0 | 0,61666679 | 0,91168931 |
| EXOC7_HUM | EXOC7 | sp\|Q9UPT5\|EXOC7_HUMAN Exocyst complex component 7 OS=Homo sapiens GN=EXO | 113,559 | 5,44 | 83382 | 3 | 21,11 | 21,18 | 20,67 | 21,56 | 21,94 | 21,31 | 1,533 | 6,41E-02 | 0 | 0,61666616 | 1,19316207 |
| NUDC1_HUM | NUDCD1 | sp\|Q96RS6\|NUDC1_HUMAN NudC domain-containing protein 1 OS=Homo sapiens GN= | 36,847 | 2,74 | 66756 | 1 | 17,41 | 20,92 | 20,19 | 19,17 | 20,00 | 21,20 | 1,532 | 6,41E-01 | 0 | 0,61527952 | 0,19339141 |
| PYGB_HUMA | PYGB | sp\|P11216\|PYGB_HUMAN Glycogen phosphorylase, brain form OS=Homo sapiens GN=F | 646,443 | 26,22 | 96696 | 15 | 23,94 | 24,06 | 23,11 | 24,50 | 24,59 | 23,86 | 1,530 | 1,79E-01 | 0 | 0,61333338 | 0,74723853 |
| SET_HUMAN | SET | sp\|Q01105\|SET_HUMAN Protein SET OS=Homo sapiens GN=SET PE=1 SV=3 | 277,836 | 28,97 | 33489 | 6 | 25,88 | 26,16 | 25,63 | 26,40 | 26,72 | 26,39 | 1,530 | 3,08E-02 | 0 | 0,61333338 | 1,51164629 |
| STAT1_HUM | STAT1 | sp\|P42224\|STAT1_HUMAN Signal transducer and activator of transcription 1-alpha/beta | 163,43 | 9,73 | 87335 | 6 | 22,91 | 21,83 | 22,51 | 22,35 | 22,73 | 24,01 | 1,530 | 3,59E-01 | 0 | 0,61333338 | 0,44448606 |
| RBX1_HUMA | RBX1 | sp\|P62877\|RBX1_HUMAN E3 ubiquitin-protein ligase RBX1 OS=Homo sapiens GN=RBX1 | 150,31 | 30,56 | 12274 | 3 | 20,49 | 22,04 | 18,95 | 20,45 | 22,45 | 20,41 | 1,527 | 6,14E-01 | 0 | 0,61075719 | 0,21185645 |
| PYRG1_HUM | CTPS1 | sp\|P17812\|PYRG1_HUMAN CTP synthase 1 OS=Homo sapiens GN=CTPS1 PE=1 SV=2 | 281,323 | 19,97 | 66690 | 9 | 24,67 | 23,96 | 24,96 | 24,72 | 25,02 | 25,68 | 1,526 | 2,12E-01 | 0 | 0,61000061 | 0,67447011 |
| TM9S2_HUM | TM9SF2 | sp\|Q99805\|TM9S2_HUMAN Transmembrane 9 superfamily member 2 OS=Homo sapien | 181,639 | 8,75 | 75776 | 4 | 21,17 | 22,20 | 24,08 | 22,38 | 22,69 | 24,21 | 1,526 | 5,83E-01 | 0 | 0,60999934 | 0,23439806 |
| FUBP2_HUM | KHSRP | sp\|Q92945\|FUBP2_HUMAN Far upstream element-binding protein 2 OS=Homo sapiens | 286,337 | 16,17 | 73115 | 9 | 23,62 | 24,27 | 25,05 | 24,63 | 24,89 | 25,25 | 1,526 | 2,47E-01 | 0 | 0,60999934 | 0,60660425 |
| SSA27_HUM | SSSCA1 | sp\|O60232\|SSA27_HUMAN Sjoegren syndrome/scleroderma autoantigen 1 OS=Homo s | 14,487 | 4,02 | 21474 | 1 | 21,31 | 21,18 | 18,73 | 20,94 | 21,50 | 20,60 | 1,524 | 5,28E-01 | 0 | 0,60771116 | 0,27743673 |
| QBK84305.1 | | regulatory protein IE2 [Human betaherpesvirus 5] | 70,097 | 5,86 | 62947 | 3 | 19,82 | 18,63 | 18,49 | 19,26 | 18,67 | 20,83 | 1,523 | 4,75E-01 | 0 | 0,60671043 | 0,322891 |
| CC126_HUM | CCDC126 | sp\|Q96EE4\|CC126_HUMAN Coiled-coil domain-containing protein 126 OS=Homo sapien | 25,278 | 8,57 | 15668 | 1 | 17,88 | 17,27 | 19,03 | 18,17 | 17,94 | 19,88 | 1,521 | 4,92E-01 | 0 | 0,60549037 | 0,30829212 |
| RU17_HUMA | SNRNP70 | sp\|P08621\|RU17_HUMAN U1 small nuclear ribonucleoprotein 70 kDa OS=Homo sapiens | 134,36 | 13,73 | 51557 | 6 | 22,55 | 23,00 | 23,00 | 23,59 | 22,92 | 23,85 | 1,519 | 1,28E-01 | 0 | 0,60333379 | 0,8928466 |
| PSA7_HUMA | PSMA7 | sp\|O14818\|PSA7_HUMAN Proteasome subunit alpha type-7 OS=Homo sapiens GN=PSN | 650,314 | 58,47 | 27887 | 16 | 26,68 | 26,93 | 26,43 | 27,25 | 27,36 | 27,24 | 1,519 | 1,56E-02 | 0 | 0,60333316 | 1,80648038 |
| DPP4_HUMA | DPP4 | sp\|P27487\|DPP4_HUMAN Dipeptidyl peptidase 4 OS=Homo sapiens GN=DPP4 PE=1 SV= | 16,257 | 0,91 | 88279 | 1 | 19,39 | 19,87 | 19,73 | 20,58 | 20,40 | 19,82 | 1,517 | 8,99E-02 | 0 | 0,60086632 | 1,04629596 |
| TIGAR_HUM | TIGAR | sp\|Q9NQ88\|TIGAR_HUMAN Fructose-2,6-bisphosphatase TIGAR OS=Homo sapiens GN= | 54,2 | 6,3 | 30063 | 1 | 20,93 | 20,69 | 19,57 | 20,96 | 21,04 | 20,99 | 1,516 | 2,26E-01 | 0 | 0,60008812 | 0,64566231 |
| RPN1_HUMA | RPN1 | sp\|P04843\|RPN1_HUMAN Dolichyl-diphosphooligosaccharide--protein glycosyltransfera | 960,83 | 44,48 | 68569 | 21 | 26,60 | 26,25 | 28,20 | 27,11 | 27,15 | 28,59 | 1,516 | 4,81E-01 | 0 | 0,59999975 | 0,31796481 |
| TCEA1_HUM | TCEA1 | sp\|P23193\|TCEA1_HUMAN Transcription elongation factor A protein 1 OS=Homo sapier | 95,523 | 10,63 | 33970 | 3 | 22,98 | 22,62 | 24,67 | 23,48 | 23,52 | 25,07 | 1,516 | 5,05E-01 | 0 | 0,59999975 | 0,29654353 |
| ROA1_HUMA | HNRNPA1 | sp\|P09651\|ROA1_HUMAN Heterogeneous nuclear ribonucleoprotein A1 OS=Homo sapi | 1236,174 | 60,48 | 38747 | 29 | 27,56 | 27,54 | 28,52 | 28,16 | 28,29 | 28,97 | 1,516 | 2,17E-01 | 0 | 0,59999975 | 0,66415319 |
| CHRD1_HUM | CHORDC1 | sp\|Q9UHD1\|CHRD1_HUMAN Cysteine and histidine-rich domain-containing protein 1 O | 46,628 | 6,63 | 37490 | 2 | 21,31 | 21,85 | 21,96 | 21,67 | 22,15 | 23,09 | 1,512 | 2,67E-01 | 0 | 0,59666697 | 0,57368492 |
| ZA2G_HUMA | AZGP1 | sp\|P25311\|ZA2G_HUMAN Zinc-alpha-2-glycoprotein OS=Homo sapiens GN=AZGP1 PE=1 | 42,483 | 7,72 | 34259 | 2 | 21,49 | 20,87 | 22,27 | 23,66 | 21,11 | 21,65 | 1,512 | 5,33E-01 | 0 | 0,59666634 | 0,27342021 |
| SGMR1_HUM | SIGMAR1 | sp\|Q99720\|SGMR1_HUMAN Sigma non-opioid intracellular receptor 1 OS=Homo sapien | 59,107 | 28,7 | 25128 | 3 | 22,70 | 22,32 | 23,03 | 23,19 | 23,51 | 23,13 | 1,509 | 6,62E-02 | 0 | 0,59333293 | 1,17893592 |
| RABL6_HUM | RABL6 | sp\|Q3YEC7\|RABL6_HUMAN Rab-like protein 6 OS=Homo sapiens GN=RABL6 PE=1 SV=2 | 25,747 | 1,51 | 79549 | 1 | 18,64 | 19,32 | 20,15 | 19,66 | 19,13 | 21,09 | 1,506 | 4,64E-01 | 0 | 0,59053294 | 0,33348711 |
| K1C9_HUMA | KRT9 | sp\|P35527\|K1C9_HUMAN Keratin, type I cytoskeletal 9 OS=Homo sapiens GN=KRT9 PE= | 2768,14 | 81,7 | 62064 | 52 | 30,64 | 29,77 | 28,93 | 31,87 | 30,59 | 28,65 | 1,505 | 6,07E-01 | 0 | 0,59000015 | 0,2168931 |
| TPR_HUMAN | TPR | sp\|P12270\|TPR_HUMAN Nucleoprotein TPR OS=Homo sapiens GN=TPR PE=1 SV=3 | 23,399 | 0,42 | 267293 | 1 | 19,10 | 19,13 | 21,06 | 19,77 | 20,15 | 21,14 | 1,505 | 4,84E-01 | 0 | 0,59000015 | 0,31491499 |
| HM13_HUM | HM13 | sp\|Q8TCT9\|HM13_HUMAN Minor histocompatibility antigen H13 OS=Homo sapiens GN | 148,189 | 14,06 | 41488 | 4 | 24,24 | 23,34 | 24,76 | 24,46 | 24,43 | 25,22 | 1,505 | 2,94E-01 | 0 | 0,58999952 | 0,53186957 |
| SSRP1_HUM | SSRP1 | sp\|Q08945\|SSRP1_HUMAN FACT complex subunit SSRP1 OS=Homo sapiens GN=SSRP1 F | 372,733 | 28,63 | 81075 | 14 | 23,86 | 24,29 | 25,71 | 24,48 | 24,96 | 26,19 | 1,505 | 4,79E-01 | 0 | 0,58999952 | 0,31983241 |
| TNPO2_HUM | TNPO2 | sp\|O14787\|TNPO2_HUMAN Transportin-2 OS=Homo sapiens GN=TNPO2 PE=1 SV=3 | 223,297 | 16,28 | 101388 | 10 | 21,35 | 22,75 | 23,25 | 22,49 | 23,12 | 23,50 | 1,502 | 4,11E-01 | 0 | 0,58666674 | 0,38570142 |
| HMGB2_HUM | HMGB2 | sp\|P26583\|HMGB2_HUMAN High mobility group protein B2 OS=Homo sapiens GN=HM | 178,469 | 35,41 | 24034 | 8 | 23,12 | 23,60 | 23,71 | 24,17 | 24,02 | 24,00 | 1,502 | 3,60E-02 | 0 | 0,58666674 | 1,44333635 |
| PESC_HUMA | PES1 | sp\|O00541\|PESC_HUMAN Pescadillo homolog OS=Homo sapiens GN=PES1 PE=1 SV=1 | 23,928 | 1,53 | 68003 | 1 | 19,84 | 19,70 | 20,05 | 20,52 | 20,05 | 20,78 | 1,502 | 6,82E-02 | 0 | 0,58666674 | 1,16596292 |
| LKHA4_HUM | LTA4H | sp\|P09960\|LKHA4_HUMAN Leukotriene A-4 hydrolase OS=Homo sapiens GN=LTA4H PE | 162,469 | 7,69 | 69285 | 3 | 23,06 | 22,72 | 21,28 | 23,26 | 23,01 | 22,55 | 1,502 | 3,72E-01 | 0 | 0,58666674 | 0,42965255 |
| HYPK_HUMA | HYPK | sp\|Q9NX55\|HYPK_HUMAN Huntingtin-interacting protein K OS=Homo sapiens GN=HYPI | 14,356 | 11,63 | 14665 | 1 | 18,93 | 19,21 | 18,65 | 19,14 | 19,28 | 20,13 | 1,501 | 1,68E-01 | 0 | 0,58615494 | 0,77425941 |
| DIAP3_HUM | DIAPH3 | sp\|Q9NSV4\|DIAP3_HUMAN Protein diaphanous homolog 3 OS=Homo sapiens GN=DIAPI | 18,701 | 0,59 | 136926 | 1 | 20,69 | 17,51 | 18,99 | 20,31 | 20,15 | 18,48 | 1,498 | 6,20E-01 | 0 | 0,58350182 | 0,20727156 |
| S27A4_HUM | SLC27A4 | sp\|Q6P1M0\|S27A4_HUMAN Long-chain fatty acid transport protein 4 OS=Homo sapiens | 51,778 | 5,75 | 72064 | 3 | 20,33 | 21,13 | 21,99 | 20,99 | 21,84 | 22,37 | 1,498 | 4,04E-01 | 0 | 0,58333397 | 0,39378929 |
| HNRPM_HUM | HNRNPM | sp\|P52272\|HNRPM_HUMAN Heterogeneous nuclear ribonucleoprotein M OS=Homo sa | 761,378 | 39,32 | 77516 | 23 | 24,68 | 24,84 | 27,32 | 25,49 | 25,58 | 27,52 | 1,498 | 6,18E-01 | 0 | 0,58333333 | 0,20892252 |
| HNRPD_HUM | HNRNPD | sp\|Q14103\|HNRPD_HUMAN Heterogeneous nuclear ribonucleoprotein D0 OS=Homo sa | 346,635 | 31,83 | 38434 | 12 | 25,10 | 25,53 | 25,99 | 25,99 | 26,09 | 26,29 | 1,498 | 9,83E-02 | 0 | 0,58333333 | 1,00745116 |
| OST48_HUM | DDOST | sp\|P39656\|OST48_HUMAN Dolichyl-diphosphooligosaccharide--protein glycosyltransfe | 216,155 | 23,46 | 50801 | 6 | 25,67 | 25,29 | 26,35 | 26,11 | 26,13 | 26,82 | 1,498 | 2,07E-01 | 0 | 0,5833327 | 0,68352751 |
| TMED4_HUM | TMED4 | sp\|Q7Z7H5\|TMED4_HUMAN Transmembrane emp24 domain-containing protein 4 OS=I | 232,33 | 37,44 | 25943 | 6 | 24,89 | 24,63 | 23,75 | 25,10 | 25,08 | 24,83 | 1,495 | 1,78E-01 | 0 | 0,58000056 | 0,7488224 |
| UBP14_HUM | USP14 | sp\|P54578\|UBP14_HUMAN Ubiquitin carboxyl-terminal hydrolase 14 OS=Homo sapiens | 310,609 | 34,01 | 56069 | 11 | 24,75 | 25,01 | 24,63 | 24,99 | 25,68 | 25,46 | 1,495 | 6,70E-02 | 0 | 0,57999992 | 1,17366619 |
| SRPK1_HUM | SRPK1 | sp\|Q96SB4\|SRPK1_HUMAN SRSF protein kinase 1 OS=Homo sapiens GN=SRPK1 PE=1 SV | 104,697 | 10,38 | 74325 | 6 | 23,02 | 23,03 | 23,76 | 23,61 | 23,65 | 24,29 | 1,495 | 1,53E-01 | 0 | 0,57999992 | 0,81483102 |
| GORS2_HUM | GORASP2 | sp\|Q9H8Y8\|GORS2_HUMAN Golgi reassembly-stacking protein 2 OS=Homo sapiens GN= | 66,071 | 4,2 | 47145 | 2 | 22,03 | 22,70 | 22,59 | 22,62 | 23,03 | 23,41 | 1,495 | 1,33E-01 | 0 | 0,57999992 | 0,87578801 |
| RIF1_HUMAN | RIF1 | sp\|Q5UIP0\|RIF1_HUMAN Telomere-associated protein RIF1 OS=Homo sapiens GN=RIF1 | 52,578 | 0,69 | 274466 | 1 | 17,88 | 18,10 | 19,20 | 19,35 | 17,83 | 19,72 | 1,492 | 4,61E-01 | 0 | 0,57694689 | 0,33670567 |
| MY18A_HUM | MYO18A | sp\|Q92614\|MY18A_HUMAN Unconventional myosin-XVIIIa OS=Homo sapiens GN=MYO | 283,04 | 5,11 | 233115 | 8 | 25,58 | 24,98 | 24,25 | 26,02 | 26,00 | 24,52 | 1,491 | 4,11E-01 | 0 | 0,57666715 | 0,38663037 |
| EMC1_HUM | EMC1 | sp\|Q8N766\|EMC1_HUMAN ER membrane protein complex subunit 1 OS=Homo sapiens | 273,314 | 9,97 | 111759 | 6 | 20,89 | 20,97 | 23,94 | 21,67 | 21,82 | 24,04 | 1,491 | 6,72E-01 | 0 | 0,57666715 | 0,17290108 |
| TMCO1_HUM | TMCO1 | sp\|Q9UM00\|TMCO1_HUMAN Calcium load-activated calcium channel OS=Homo sapien | 87,93 | 15,43 | 21175 | 2 | 22,16 | 22,52 | 23,25 | 22,85 | 23,38 | 23,43 | 1,491 | 1,95E-01 | 0 | 0,57666651 | 0,71096201 |
| NUDC_HUM | NUDC | sp\|Q9Y266\|NUDC_HUMAN Nuclear migration protein nudC OS=Homo sapiens GN=NUD | 187,713 | 17,82 | 38243 | 6 | 25,07 | 24,95 | 23,99 | 25,44 | 25,26 | 25,03 | 1,488 | 1,88E-01 | 0 | 0,57333374 | 0,72540399 |
| ERGI1_HUM | ERGIC1 | sp\|Q969X5\|ERGI1_HUMAN Endoplasmic reticulum-Golgi intermediate compartment pro | 159,384 | 24,48 | 32592 | 7 | 24,76 | 24,08 | 25,19 | 25,25 | 25,04 | 25,46 | 1,488 | 1,72E-01 | 0 | 0,5733331 | 0,76446524 |
| ERG28_HUM | ERG28 | sp\|Q9UKR5\|ERG28_HUMAN Probable ergosterol biosynthetic protein 28 OS=Homo sapi | 61,116 | 20 | 15864 | 2 | 21,25 | 21,38 | 19,09 | 21,52 | 21,70 | 20,22 | 1,487 | 5,49E-01 | 0 | 0,57241313 | 0,26030188 |
| FANCI_HUM | FANCI | sp\|Q9NVI1\|FANCI_HUMAN Fanconi anemia group I protein OS=Homo sapiens GN=FAN( | 74,218 | 1,2 | 149324 | 2 | 18,19 | 17,56 | 21,37 | 18,78 | 18,33 | 21,72 | 1,485 | 7,38E-01 | 0 | 0,56999969 | 0,13211603 |
| PSB1_HUMA | PSMB1 | sp\|P20618\|PSB1_HUMAN Proteasome subunit beta type-1 OS=Homo sapiens GN=PSME | 458,909 | 47,72 | 26489 | 9 | 25,57 | 26,02 | 24,95 | 26,21 | 26,20 | 25,84 | 1,485 | 1,62E-01 | 0 | 0,56999969 | 0,78965302 |
| STT3A_HUM | STT3A | sp\|P46977\|STT3A_HUMAN Dolichyl-diphosphooligosaccharide--protein glycosyltransfer | 262,599 | 13,9 | 80530 | 10 | 25,17 | 24,03 | 25,87 | 25,26 | 25,32 | 26,20 | 1,485 | 4,07E-01 | 0 | 0,56999969 | 0,39000523 |
| PGM2_HUM | PGM2 | sp\|Q96G03\|PGM2_HUMAN Phosphoglucomutase-2 OS=Homo sapiens GN=PGM2 PE=1 | 66,851 | 7,84 | 68283 | 4 | 24,02 | 24,34 | 22,08 | 24,65 | 24,58 | 22,92 | 1,485 | 5,63E-01 | 0 | 0,56999969 | 0,24965107 |
| TRFE_HUMA | TF | sp\|P02787\|TRFE_HUMAN Serotransferrin OS=Homo sapiens GN=TF PE=1 SV=3 | 23,424 | 1 | 77064 | 1 | 21,47 | 20,28 | 23,74 | 22,61 | 21,35 | 23,24 | 1,484 | 6,48E-01 | 0 | 0,56914775 | 0,18815375 |
| DCAKD_HUM | DCAKD | sp\|Q8WVC6\|DCAKD_HUMAN Dephospho-CoA kinase domain-containing protein OS=Ho | 28,241 | 5,63 | 26550 | 1 | 17,99 | 18,58 | 19,85 | 20,29 | 18,23 | 19,61 | 1,482 | 5,25E-01 | 0 | 0,56786346 | 0,27959949 |
| SEC13_HUM | SEC13 | sp\|P55735\|SEC13_HUMAN Protein SEC13 homolog OS=Homo sapiens GN=SEC13 PE=1 S | 149,194 | 19,88 | 35541 | 3 | 22,23 | 23,15 | 23,85 | 22,79 | 23,83 | 24,31 | 1,481 | 4,32E-01 | 0 | 0,56666692 | 0,36460211 |
| SNRPA_HUM | SNRPA | sp\|P09012\|SNRPA_HUMAN U1 small nuclear ribonucleoprotein A OS=Homo sapiens GN | 59,463 | 17,73 | 31280 | 4 | 23,86 | 22,89 | 24,04 | 24,16 | 24,11 | 24,22 | 1,481 | 1,89E-01 | 0 | 0,56666629 | 0,72317837 |
| GELS_HUMA | GSN | sp\|P06396\|GELS_HUMAN Gelsolin OS=Homo sapiens GN=GSN PE=1 SV=1 | 687,497 | 30,43 | 85698 | 20 | 28,02 | 27,50 | 27,54 | 28,04 | 27,47 | 29,25 | 1,481 | 3,62E-01 | 0 | 0,56666629 | 0,44168743 |
| SMCA5_HUM | SMARCA5 | sp\|O60264\|SMCA5_HUMAN SWI/SNF-related matrix-associated actin-dependent regula | 154,169 | 8,84 | 121905 | 8 | 21,88 | 19,62 | 24,22 | 20,97 | 21,94 | 24,50 | 1,479 | 7,56E-01 | 0 | 0,56437747 | 0,12151181 |
| PSB2_HUMA | PSMB2 | sp\|P49721\|PSB2_HUMAN Proteasome subunit beta type-2 OS=Homo sapiens GN=PSME | 263,668 | 54,73 | 22836 | 9 | 24,84 | 25,32 | 25,29 | 25,43 | 25,94 | 25,77 | 1,478 | 5,94E-02 | 0 | 0,56333351 | 1,22616498 |
| UCHL5_HUM | UCHL5 | sp\|Q9Y5K5\|UCHL5_HUMAN Ubiquitin carboxyl-terminal hydrolase isozyme L5 OS=Hom | 182,093 | 24,01 | 37607 | 7 | 23,20 | 24,11 | 24,05 | 23,85 | 24,12 | 25,08 | 1,478 | 3,01E-01 | 0 | 0,56333351 | 0,52101956 |
| PPID_HUMA | PPID | sp\|Q08752\|PPID_HUMAN Peptidyl-prolyl cis-trans isomerase D OS=Homo sapiens GN=P | 225,309 | 21,62 | 40764 | 6 | 24,69 | 24,99 | 23,93 | 25,22 | 25,11 | 24,97 | 1,478 | 1,57E-01 | 0 | 0,56333288 | 0,80485468 |
| HNRH1_HUM | HNRNPH1 | sp\|P31943\|HNRH1_HUMAN Heterogeneous nuclear ribonucleoprotein H OS=Homo sap | 628,947 | 46,77 | 49229 | 14 | 24,48 | 24,68 | 26,62 | 25,40 | 25,35 | 26,72 | 1,478 | 5,28E-01 | 0 | 0,56333288 | 0,27715543 |
| ZSWM8_HUM | ZSWIM8 | sp\|A7E2V4\|ZSWM8_HUMAN Zinc finger SWIM domain-containing protein 8 OS=Homo s | 64,544 | 1,58 | 197297 | 2 | 18,07 | 16,28 | 19,45 | 18,31 | 17,39 | 19,79 | 1,478 | 6,51E-01 | 0 | 0,56333288 | 0,18649945 |
| FDFT_HUMA | FDFT1 | sp\|P37268\|FDFT_HUMAN Squalene synthase OS=Homo sapiens GN=FDFT1 PE=1 SV=1 | 31,287 | 3,12 | 48115 | 1 | 17,25 | 18,36 | 21,38 | 18,62 | 18,67 | 21,38 | 1,475 | 7,34E-01 | 0 | 0,56032753 | 0,13450462 |
| FRYL_HUMA | FRYL | sp\|O94915\|FRYL_HUMAN Protein furry homolog-like OS=Homo sapiens GN=FRYL PE=1 | 124,257 | 1,92 | 339598 | 3 | 21,05 | 21,65 | 19,79 | 21,61 | 21,77 | 20,79 | 1,474 | 4,22E-01 | 0 | 0,56000074 | 0,37475293 |
| CHD4_HUMA | CHD4 | sp\|Q14839\|CHD4_HUMAN Chromodomain-helicase-DNA-binding protein 4 OS=Homo sa | 320,639 | 10,36 | 218005 | 15 | 25,43 | 23,30 | 26,81 | 26,10 | 24,73 | 26,39 | 1,474 | 6,50E-01 | 0 | 0,5600001 | 0,18736984 |
| DHCR7_HUM | DHCR7 | sp\|Q9UBM7\|DHCR7_HUMAN 7-dehydrocholesterol reductase OS=Homo sapiens GN=D | 38,359 | 2,32 | 54489 | 1 | 22,78 | 22,59 | 23,03 | 23,11 | 23,31 | 23,66 | 1,474 | 5,24E-02 | 0 | 0,55999947 | 1,28047039 |
| HACD3_HUM | HACD3 | sp\|Q9P035\|HACD3_HUMAN Very-long-chain (3R)-3-hydroxyacyl-CoA dehydratase 3 OS= | 67,143 | 7,18 | 43160 | 3 | 22,46 | 22,31 | 23,08 | 22,77 | 23,25 | 23,50 | 1,471 | 1,55E-01 | 0 | 0,55666733 | 0,80858316 |
| PUR4_HUMA | PFAS | sp\|O15067\|PUR4_HUMAN Phosphoribosylformylglycinamidine synthase OS=Homo sapi | 239,628 | 4,56 | 144734 | 4 | 23,04 | 22,99 | 22,31 | 23,62 | 23,72 | 22,67 | 1,471 | 2,45E-01 | 0 | 0,55666669 | 0,61039827 |
| CYB5_HUMA | CYB5A | sp\|P00167\|CYB5_HUMAN Cytochrome b5 OS=Homo sapiens GN=CYB5A PE=1 SV=2 | 192,883 | 41,04 | 15330 | 4 | 23,90 | 24,18 | 24,92 | 24,66 | 24,87 | 25,14 | 1,471 | 1,71E-01 | 0 | 0,55666669 | 0,76601259 |
| TX264_HUM | TEX264 | sp\|Q9Y6I9\|TX264_HUMAN Testis-expressed protein 264 OS=Homo sapiens GN=TEX264 | 13,85 | 3,51 | 34189 | 1 | 21,06 | 20,74 | 20,71 | 21,89 | 21,12 | 21,16 | 1,467 | 1,14E-01 | 0 | 0,55333392 | 0,94407331 |
| NC2A_HUMA | DRAP1 | sp\|Q14919\|NC2A_HUMAN Dr1-associated corepressor OS=Homo sapiens GN=DRAP1 PE | 45,562 | 5,37 | 22350 | 1 | 17,69 | 19,54 | 19,72 | 18,96 | 19,80 | 19,85 | 1,467 | 4,79E-01 | 0 | 0,55333265 | 0,31933567 |
| CYTN_HUMA | CST1 | sp\|P01037\|CYTN_HUMAN Cystatin-SN OS=Homo sapiens GN=CST1 PE=1 SV=3 | 169,003 | 32,62 | 16388 | 3 | 19,32 | 21,95 | 19,49 | 24,27 | 18,95 | 19,20 | 1,464 | 7,90E-01 | 0 | 0,55003993 | 0,10242009 |
| CKAP4_HUM | CKAP4 | sp\|Q07065\|CKAP4_HUMAN Cytoskeleton-associated protein 4 OS=Homo sapiens GN=C | 514,309 | 24,25 | 66022 | 13 | 25,89 | 25,54 | 27,10 | 26,45 | 26,30 | 27,43 | 1,464 | 4,04E-01 | 0 | 0,54999987 | 0,39309557 |
| A4_HUMAN | APP | sp\|P05067\|A4_HUMAN Amyloid-beta A4 protein OS=Homo sapiens GN=APP PE=1 SV=3 | 39,781 | 1,69 | 86943 | 1 | 18,84 | 17,79 | 19,97 | 19,27 | 19,51 | 19,46 | 1,462 | 4,36E-01 | 0 | 0,54833857 | 0,36059135 |
| SRSF6_HUM | SRSF6 | sp\|Q13247\|SRSF6_HUMAN Serine/arginine-rich splicing factor 6 OS=Homo sapiens GN= | 154,897 | 16,57 | 39587 | 6 | 22,96 | 23,06 | 23,94 | 23,70 | 23,39 | 24,51 | 1,461 | 2,97E-01 | 0 | 0,5466671 | 0,52690042 |
| CLU_HUMAN | CLUH | sp\|O75153\|CLU_HUMAN Clustered mitochondria protein homolog OS=Homo sapiens G | 202,716 | 9,24 | 146670 | 11 | 23,67 | 23,54 | 23,64 | 23,52 | 24,40 | 24,57 | 1,461 | 1,71E-01 | 0 | 0,54666646 | 0,76787592 |
| CSN2_HUMA | COPS2 | sp\|P61201\|CSN2_HUMAN COP9 signalosome complex subunit 2 OS=Homo sapiens GN= | 290,636 | 38,15 | 51597 | 10 | 23,73 | 23,85 | 22,94 | 24,05 | 24,30 | 23,81 | 1,461 | 1,61E-01 | 0 | 0,54666583 | 0,79234737 |
| EIF3J_HUMA | EIF3J | sp\|O75822\|EIF3J_HUMAN Eukaryotic translation initiation factor 3 subunit J OS=Homo s | 80,098 | 12,02 | 29062 | 3 | 23,31 | 22,74 | 20,81 | 23,57 | 23,37 | 21,55 | 1,457 | 6,13E-01 | 0 | 0,54333369 | 0,21239521 |
| PSMD1_HUM | PSMD1 | sp\|Q99460\|PSMD1_HUMAN 26S proteasome non-ATPase regulatory subunit 1 OS=Hom | 1451,483 | 49,32 | 105836 | 35 | 26,43 | 26,77 | 27,80 | 26,97 | 27,32 | 28,34 | 1,457 | 4,03E-01 | 0 | 0,54333305 | 0,39443073 |
| SUMO1_HUM | SUMO1 | sp\|P63165\|SUMO1_HUMAN Small ubiquitin-related modifier 1 OS=Homo sapiens GN=S | 20,975 | 15,84 | 11557 | 1 | 20,92 | 21,28 | 20,38 | 21,73 | 21,31 | 21,17 | 1,457 | 1,56E-01 | 0 | 0,54333305 | 0,80817159 |
| AT2B4_HUM | ATP2B4 | sp\|P23634\|AT2B4_HUMAN Plasma membrane calcium-transporting ATPase 4 OS=Homo | 1067,433 | 20,55 | 137920 | 25 | 24,53 | 23,77 | 23,71 | 24,93 | 24,73 | 23,98 | 1,457 | 2,37E-01 | 0 | 0,54333305 | 0,62438441 |
| TBB4A_HUM | TUBB4A | sp\|P04350\|TBB4A_HUMAN Tubulin beta-4A chain OS=Homo sapiens GN=TUBB4A PE=1 | 1951,231 | 72,07 | 49586 | 41 | 18,23 | 18,62 | 21,14 | 19,13 | 19,51 | 20,98 | 1,457 | 6,39E-01 | 0 | 0,54325867 | 0,19420222 |
| CBPM_HUM | CPM | sp\|P14384\|CBPM_HUMAN Carboxypeptidase M OS=Homo sapiens GN=CPM PE=1 SV=2 | 11,778 | 2,93 | 50514 | 1 | 19,17 | 19,85 | 17,95 | 20,47 | 19,76 | 18,36 | 1,457 | 5,50E-01 | 0 | 0,5426623 | 0,25971524 |
| PROS_HUMA | PROS1 | sp\|P07225\|PROS_HUMAN Vitamin K-dependent protein S OS=Homo sapiens GN=PROS1 | 10,999 | 1,18 | 75123 | 1 | 18,86 | 20,55 | 22,57 | 20,76 | 20,70 | 22,14 | 1,454 | 6,69E-01 | 0 | 0,54027303 | 0,17486292 |
| SGT1_HUMA | SUGT1 | sp\|Q9Y2Z0\|SGT1_HUMAN Protein SGT1 homolog OS=Homo sapiens GN=SUGT1 PE=1 SV | 28,784 | 1,55 | 72758 | 1 | 22,42 | 22,14 | 22,03 | 22,85 | 22,77 | 22,59 | 1,454 | 1,79E-02 | 0 | 0,54000028 | 1,74786903 |
| XPP3_HUMA | XPNPEP3 | sp\|Q9NQH7\|XPP3_HUMAN Probable Xaa-Pro aminopeptidase 3 OS=Homo sapiens GN=. | 28,541 | 1,38 | 57034 | 1 | 26,68 | 26,91 | 19,63 | 26,87 | 27,17 | 20,80 | 1,454 | 8,73E-01 | 0 | 0,54000028 | 0,05908898 |
| 2ABA_HUMA | PPP2R2A | sp\|P63151\|2ABA_HUMAN Serine/threonine-protein phosphatase 2A 55 kDa regulatory | 298,843 | 30,65 | 51692 | 10 | 24,87 | 24,98 | 25,44 | 25,31 | 25,53 | 26,07 | 1,454 | 1,31E-01 | 0 | 0,53999964 | 0,88127648 |
| MTA2_HUM | MTA2 | sp\|O94776\|MTA2_HUMAN Metastasis-associated protein MTA2 OS=Homo sapiens GN= | 259,574 | 17,22 | 75023 | 10 | 23,45 | 23,31 | 24,33 | 23,77 | 23,97 | 24,97 | 1,454 | 3,32E-01 | 0 | 0,53999964 | 0,47895466 |
| KIF4A_HUM | KIF4A | sp\|O95239\|KIF4A_HUMAN Chromosome-associated kinesin KIF4A OS=Homo sapiens GI | 617,837 | 19,48 | 139881 | 20 | 20,89 | 23,23 | 25,91 | 24,04 | 23,56 | 24,04 | 1,451 | 7,32E-01 | 0 | 0,53666751 | 0,13569948 |
| RAE1L_HUM | RAE1 | sp\|P78406\|RAE1L_HUMAN mRNA export factor OS=Homo sapiens GN=RAE1 PE=1 SV=1 | 67,01 | 12,5 | 40968 | 4 | 21,18 | 20,93 | 20,96 | 21,57 | 21,14 | 21,97 | 1,451 | 1,01E-01 | 0 | 0,53666623 | 0,99776746 |
| TBB3_HUMA | TUBB3 | sp\|Q13509\|TBB3_HUMAN Tubulin beta-3 chain OS=Homo sapiens GN=TUBB3 PE=1 SV= | 1290,774 | 48,89 | 50433 | 27 | 20,46 | 21,67 | 20,68 | 20,97 | 22,58 | 20,86 | 1,447 | 4,70E-01 | 0 | 0,53333346 | 0,32794859 |
| KAP3_HUMA | PRKAR2B | sp\|P31323\|KAP3_HUMAN cAMP-dependent protein kinase type II-beta regulatory subu | 177,388 | 20,1 | 46302 | 5 | 16,78 | 17,17 | 21,00 | 16,19 | 19,59 | 20,77 | 1,446 | 7,96E-01 | 0 | 0,53189977 | 0,09921944 |
| FAF2_HUMA | FAF2 | sp\|Q96CS3\|FAF2_HUMAN FAS-associated factor 2 OS=Homo sapiens GN=FAF2 PE=1 SV= | 231,286 | 16,85 | 52623 | 5 | 20,82 | 22,64 | 24,18 | 22,01 | 23,20 | 24,02 | 1,444 | 6,64E-01 | 0 | 0,53000069 | 0,17766227 |
| S61A1_HUM | SEC61A1 | sp\|P61619\|S61A1_HUMAN Protein transport protein Sec61 subunit alpha isoform 1 OS= | 532,609 | 30,67 | 52265 | 14 | 24,88 | 24,71 | 25,14 | 25,17 | 25,31 | 25,84 | 1,444 | 9,11E-02 | 0 | 0,53000069 | 1,04027822 |
| SSRA_HUMA | SSR1 | sp\|P43307\|SSRA_HUMAN Translocon-associated protein subunit alpha OS=Homo sapier | 128,141 | 11,89 | 32235 | 3 | 25,73 | 25,10 | 26,16 | 26,13 | 25,93 | 26,52 | 1,444 | 2,08E-01 | 0 | 0,53000005 | 0,6822111 |
| TMED2_HUM | TMED2 | sp\|Q15363\|TMED2_HUMAN Transmembrane emp24 domain-containing protein 2 OS=I | 294,443 | 30,85 | 22761 | 7 | 25,42 | 24,74 | 25,69 | 25,52 | 25,63 | 26,29 | 1,444 | 2,26E-01 | 0 | 0,53000005 | 0,64513956 |
| CBX1_HUMA | CBX1 | sp\|P83916\|CBX1_HUMAN Chromobox protein homolog 1 OS=Homo sapiens GN=CBX1 F | 128,491 | 15,68 | 21418 | 3 | 21,79 | 22,28 | 21,65 | 22,48 | 22,34 | 22,49 | 1,444 | 5,47E-02 | 0 | 0,52999942 | 1,26229706 |
| STRN_HUMA | STRN | sp\|O43815\|STRN_HUMAN Striatin OS=Homo sapiens GN=STRN PE=1 SV=4 | 11,909 | 1,41 | 86132 | 1 | 18,46 | 19,21 | 18,61 | 18,12 | 19,85 | 19,90 | 1,444 | 4,46E-01 | 0 | 0,52983983 | 0,35035526 |
| DPM1_HUM | DPM1 | sp\|O60762\|DPM1_HUMAN Dolichol-phosphate mannosyltransferase subunit 1 OS=Hon | 121,568 | 24,62 | 29634 | 5 | 23,51 | 24,01 | 23,31 | 24,02 | 24,36 | 24,03 | 1,441 | 8,97E-02 | 0 | 0,52666728 | 1,04731925 |
| RL22_HUMA | RPL22 | sp\|P35268\|RL22_HUMAN 60S ribosomal protein L22 OS=Homo sapiens GN=RPL22 PE=1 | 155,664 | 49,22 | 14787 | 6 | 25,60 | 25,39 | 24,50 | 25,91 | 26,21 | 24,95 | 1,441 | 3,58E-01 | 0 | 0,52666664 | 0,44560196 |
| DDX17_HUM | DDX17 | sp\|Q92841\|DDX17_HUMAN Probable ATP-dependent RNA helicase DDX17 OS=Homo sa | 1216,464 | 40,47 | 80272 | 33 | 26,01 | 26,32 | 27,13 | 26,63 | 26,67 | 27,74 | 1,441 | 3,46E-01 | 0 | 0,52666664 | 0,46082634 |
| TMX4_HUM | TMX4 | sp\|Q9H1E5\|TMX4_HUMAN Thioredoxin-related transmembrane protein 4 OS=Homo sa | 47,169 | 3,72 | 38952 | 1 | 20,39 | 19,85 | 21,44 | 20,89 | 20,81 | 21,56 | 1,441 | 3,72E-01 | 0 | 0,52666601 | 0,42990161 |
| PTPRJ_HUM | PTPRJ | sp\|Q12913\|PTPRJ_HUMAN Receptor-type tyrosine-protein phosphatase eta OS=Homo | 10,925 | 0,9 | 145941 | 1 | 22,29 | 21,97 | 18,91 | 22,11 | 21,80 | 20,83 | 1,439 | 6,71E-01 | 0 | 0,52474022 | 0,17348617 |
| PPIC_HUMA | PPIC | sp\|P45877\|PPIC_HUMAN Peptidyl-prolyl cis-trans isomerase C OS=Homo sapiens GN=PF | 13,881 | 3,77 | 22763 | 1 | 18,37 | 20,04 | 21,63 | 21,41 | 18,66 | 21,54 | 1,438 | 7,13E-01 | 0 | 0,52446302 | 0,14666907 |
| HNRLL_HUM | HNRNPLL | sp\|Q8WVV9\|HNRLL_HUMAN Heterogeneous nuclear ribonucleoprotein L-like OS=Hom | 26,309 | 2,77 | 60083 | 1 | 18,26 | 19,16 | 20,10 | 19,33 | 19,94 | 19,82 | 1,438 | 4,05E-01 | 0 | 0,52362378 | 0,392208 |
| CUL5_HUMA | CUL5 | sp\|Q93034\|CUL5_HUMAN Cullin-5 OS=Homo sapiens GN=CUL5 PE=1 SV=4 | 131,879 | 8,33 | 90955 | 4 | 21,86 | 20,81 | 21,78 | 21,86 | 22,04 | 22,12 | 1,437 | 2,05E-01 | 0 | 0,52333387 | 0,68812622 |
| SFR15_HUM | SCAF4 | sp\|O95104\|SFR15_HUMAN Splicing factor, arginine/serine-rich 15 OS=Homo sapiens GN | 104,688 | 2,18 | 125869 | 2 | 21,21 | 20,83 | 22,80 | 22,42 | 21,14 | 22,85 | 1,437 | 5,45E-01 | 0 | 0,52333387 | 0,26356935 |
| GSLG1_HUM | GLG1 | sp\|Q92896\|GSLG1_HUMAN Golgi apparatus protein 1 OS=Homo sapiens GN=GLG1 PE= | 460,454 | 15,18 | 134552 | 16 | 24,47 | 23,92 | 25,79 | 24,86 | 24,50 | 26,39 | 1,437 | 5,50E-01 | 0 | 0,52333323 | 0,25982092 |
| HNRPL_HUM | HNRNPL | sp\|P14866\|HNRPL_HUMAN Heterogeneous nuclear ribonucleoprotein L OS=Homo sapi | 477,12 | 41,43 | 64133 | 15 | 24,44 | 24,91 | 26,87 | 25,31 | 25,45 | 27,03 | 1,437 | 6,02E-01 | 0 | 0,52333323 | 0,22022868 |
| MCA3_HUM | EEF1E1 | sp\|O43324\|MCA3_HUMAN Eukaryotic translation elongation factor 1 epsilon-1 OS=Hon | 75,047 | 14,37 | 19811 | 2 | 21,99 | 22,07 | 23,00 | 22,16 | 23,00 | 23,47 | 1,437 | 3,56E-01 | 0 | 0,52333323 | 0,44861885 |
| GMPR2_HUM | GMPR2 | sp\|Q9P2T1\|GMPR2_HUMAN GMP reductase 2 OS=Homo sapiens GN=GMPR2 PE=1 SV= | 229,493 | 26,15 | 37874 | 6 | 23,72 | 23,27 | 22,56 | 23,81 | 23,99 | 23,32 | 1,437 | 2,53E-01 | 0 | 0,52333323 | 0,59631249 |
| SCMC2_HUM | SLC25A25 | sp\|Q6KCM7\|SCMC2_HUMAN Calcium-binding mitochondrial carrier protein SCaMC-2 O | 14,189 | 1,71 | 52663 | 1 | 18,11 | 19,13 | 20,22 | 18,49 | 18,16 | 22,38 | 1,437 | 7,43E-01 | 0 | 0,52255885 | 0,12914876 |
| DDX47_HUM | DDX47 | sp\|Q9H0S4\|DDX47_HUMAN Probable ATP-dependent RNA helicase DDX47 OS=Homo sa | 99,305 | 11,43 | 50647 | 4 | 23,29 | 23,25 | 23,59 | 23,73 | 23,87 | 24,09 | 1,434 | 2,56E-02 | 0 | 0,51999982 | 1,59105774 |
| SPEE_HUMA | SRM | sp\|P19623\|SPEE_HUMAN Spermidine synthase OS=Homo sapiens GN=SRM PE=1 SV=1 | 303,655 | 38,08 | 33825 | 8 | 25,56 | 25,77 | 25,36 | 25,74 | 26,13 | 26,38 | 1,434 | 7,80E-02 | 0 | 0,51999919 | 1,10818085 |
| SMD2_HUM | SNRPD2 | sp\|P62316\|SMD2_HUMAN Small nuclear ribonucleoprotein Sm D2 OS=Homo sapiens GI | 203,082 | 61,86 | 13527 | 8 | 25,23 | 25,47 | 25,35 | 26,09 | 25,81 | 25,70 | 1,431 | 1,88E-02 | 0 | 0,51666705 | 1,72695338 |
| UBP8_HUMA | USP8 | sp\|P40818\|UBP8_HUMAN Ubiquitin carboxyl-terminal hydrolase 8 OS=Homo sapiens GI | 156,881 | 9,93 | 127523 | 8 | 23,72 | 22,69 | 23,83 | 23,54 | 23,74 | 24,51 | 1,431 | 3,32E-01 | 0 | 0,51666705 | 0,47918188 |
| RFOX1_HUM | RBFOX1 | sp\|Q9NWB1\|RFOX1_HUMAN RNA binding protein fox-1 homolog 1 OS=Homo sapiens G | 24,654 | 2,77 | 42784 | 1 | 20,08 | 18,95 | 20,39 | 20,52 | 20,24 | 20,21 | 1,431 | 3,14E-01 | 0 | 0,51666641 | 0,50367133 |
| AL9A1_HUM | ALDH9A1 | sp\|P49189\|AL9A1_HUMAN 4-trimethylaminobutyraldehyde dehydrogenase OS=Homo | 92,414 | 8,7 | 53802 | 4 | 23,40 | 23,16 | 20,95 | 23,27 | 23,49 | 22,30 | 1,431 | 5,81E-01 | 0 | 0,51666641 | 0,23591839 |
| HP1B3_HUM | HP1BP3 | sp\|Q5SSJ5\|HP1B3_HUMAN Heterochromatin protein 1-binding protein 3 OS=Homo sapi | 153,876 | 9,4 | 61207 | 4 | 20,34 | 20,68 | 23,93 | 21,42 | 21,12 | 23,96 | 1,431 | 7,41E-01 | 0 | 0,51666641 | 0,1303779 |
| UBP12_HUM | USP12 | sp\|O75317\|UBP12_HUMAN Ubiquitin carboxyl-terminal hydrolase 12 OS=Homo sapiens | 24,584 | 4,86 | 42858 | 1 | 16,84 | 19,76 | 17,08 | 17,77 | 18,44 | 19,01 | 1,427 | 6,35E-01 | 0 | 0,51333618 | 0,19690089 |
| NHLC2_HUM | NHLRC2 | sp\|Q8NBF2\|NHLC2_HUMAN NHL repeat-containing protein 2 OS=Homo sapiens GN=NH | 59,319 | 7,3 | 79444 | 4 | 22,97 | 22,56 | 22,86 | 23,56 | 22,94 | 23,43 | 1,427 | 8,47E-02 | 0 | 0,51333364 | 1,07217434 |
| TECR_HUMA | TECR | sp\|Q9NZ01\|TECR_HUMAN Very-long-chain enoyl-CoA reductase OS=Homo sapiens GN= | 85,506 | 9,09 | 36034 | 4 | 21,99 | 21,87 | 22,79 | 22,34 | 22,43 | 23,42 | 1,427 | 3,18E-01 | 0 | 0,513333 | 0,49724236 |
| MAT2B_HUM | MAT2B | sp\|Q9NZL9\|MAT2B_HUMAN Methionine adenosyltransferase 2 subunit beta OS=Homo | 114,191 | 13,47 | 37552 | 3 | 21,84 | 21,70 | 22,59 | 22,23 | 22,10 | 23,34 | 1,427 | 3,46E-01 | 0 | 0,513333 | 0,4611597 |
| VPS52_HUM | VPS52 | sp\|Q8N1B4\|VPS52_HUMAN Vacuolar protein sorting-associated protein 52 homolog OS | 22,101 | 3,6 | 82221 | 2 | 18,66 | 18,30 | 24,39 | 20,59 | 17,82 | 24,47 | 1,424 | 8,62E-01 | 0 | 0,5103817 | 0,06438342 |
| BTF3_HUMA | BTF3 | sp\|P20290\|BTF3_HUMAN Transcription factor BTF3 OS=Homo sapiens GN=BTF3 PE=1 S | 241,753 | 33,5 | 22168 | 3 | 23,74 | 24,32 | 23,09 | 24,43 | 24,78 | 23,47 | 1,424 | 3,89E-01 | 0 | 0,51000023 | 0,40962604 |
| DHPR_HUMA | QDPR | sp\|P09417\|DHPR_HUMAN Dihydropteridine reductase OS=Homo sapiens GN=QDPR PE= | 189,683 | 18,44 | 25790 | 3 | 22,82 | 23,04 | 22,58 | 23,23 | 23,34 | 23,40 | 1,424 | 2,29E-02 | 0 | 0,50999959 | 1,64103211 |
| NELFA_HUM | NELFA | sp\|Q9H3P2\|NELFA_HUMAN Negative elongation factor A OS=Homo sapiens GN=NELFA | 22,344 | 4,92 | 57277 | 1 | 17,35 | 17,12 | 19,88 | 17,60 | 18,15 | 20,13 | 1,423 | 6,86E-01 | 0 | 0,50901922 | 0,16348207 |
| HNRL1_HUM | HNRNPUL1 | sp\|Q9BUJ2\|HNRL1_HUMAN Heterogeneous nuclear ribonucleoprotein U-like protein 1 | 80,349 | 3,86 | 95739 | 3 | 21,16 | 21,31 | 21,38 | 21,76 | 22,22 | 21,39 | 1,421 | 1,11E-01 | 0 | 0,50666682 | 0,95360456 |
| DDX23_HUM | DDX23 | sp\|Q9BUQ8\|DDX23_HUMAN Probable ATP-dependent RNA helicase DDX23 OS=Homo s | 53,631 | 3,29 | 95583 | 2 | 16,85 | 18,19 | 19,56 | 18,05 | 17,83 | 20,24 | 1,421 | 6,68E-01 | 0 | 0,50666618 | 0,17507071 |
| RU2B_HUMA | SNRPB2 | sp\|P08579\|RU2B_HUMAN U2 small nuclear ribonucleoprotein B'' OS=Homo sapiens GN | 178,124 | 30,22 | 25486 | 5 | 24,18 | 24,11 | 23,12 | 24,33 | 24,25 | 24,35 | 1,421 | 2,14E-01 | 0 | 0,50666618 | 0,66883715 |
| LSM2_HUMA | LSM2 | sp\|Q9Y333\|LSM2_HUMAN U6 snRNA-associated Sm-like protein LSm2 OS=Homo sapier | 21,509 | 8,42 | 10835 | 1 | 20,71 | 20,69 | 18,44 | 20,73 | 21,08 | 19,54 | 1,420 | 5,99E-01 | 0 | 0,50575193 | 0,2228807 |
| ERLEC_HUM | ERLEC1 | sp\|Q96DZ1\|ERLEC_HUMAN Endoplasmic reticulum lectin 1 OS=Homo sapiens GN=ERLE | 24,086 | 2,07 | 54858 | 1 | 18,32 | 17,08 | 20,04 | 20,21 | 17,18 | 19,56 | 1,420 | 7,09E-01 | 0 | 0,50538699 | 0,14942722 |
| STK36_HUM | STK36 | sp\|Q9NRP7\|STK36_HUMAN Serine/threonine-protein kinase 36 OS=Homo sapiens GN= | 39,11 | 1,6 | 143995 | 2 | 16,49 | 18,33 | 18,83 | 18,31 | 18,69 | 18,16 | 1,419 | 5,26E-01 | 0 | 0,50520388 | 0,27898592 |
| PIGS_HUMA | PIGS | sp\|Q96S52\|PIGS_HUMAN GPI transamidase component PIG-S OS=Homo sapiens GN=PI | 61,878 | 3,78 | 61656 | 1 | 18,98 | 18,24 | 20,05 | 19,24 | 19,13 | 20,41 | 1,417 | 4,92E-01 | 0 | 0,50333341 | 0,30806895 |
| LC7L2_HUM | LUC7L2 | sp\|Q9Y383\|LC7L2_HUMAN Putative RNA-binding protein Luc7-like 2 OS=Homo sapiens | 147,322 | 13,27 | 46514 | 4 | 21,69 | 21,74 | 21,07 | 22,18 | 21,30 | 22,53 | 1,417 | 3,01E-01 | 0 | 0,50333341 | 0,52072845 |
| NOP58_HUM | NOP58 | sp\|Q9Y2X3\|NOP58_HUMAN Nucleolar protein 58 OS=Homo sapiens GN=NOP58 PE=1 S | 634,531 | 29,49 | 59578 | 12 | 22,95 | 23,09 | 25,30 | 23,41 | 23,65 | 25,79 | 1,417 | 6,63E-01 | 0 | 0,50333341 | 0,1781853 |
| TMEDA_HUM | TMED10 | sp\|P49755\|TMEDA_HUMAN Transmembrane emp24 domain-containing protein 10 OS= | 368,658 | 33,33 | 24976 | 9 | 26,51 | 26,27 | 27,24 | 26,92 | 26,92 | 27,69 | 1,417 | 2,65E-01 | 0 | 0,50333341 | 0,57698466 |
| COPB2_HUM | COPB2 | sp\|P35606\|COPB2_HUMAN Coatomer subunit beta' OS=Homo sapiens GN=COPB2 PE=1 | 855,511 | 40,4 | 102487 | 28 | 26,07 | 26,23 | 26,87 | 26,38 | 26,90 | 27,40 | 1,417 | 2,59E-01 | 0 | 0,50333277 | 0,58712928 |
| RTRAF_HUM | RTRAF | sp\|Q9Y224\|RTRAF_HUMAN RNA transcription, translation and transport factor protein | 223,792 | 33,61 | 28068 | 6 | 22,85 | 23,28 | 24,34 | 23,16 | 23,75 | 25,07 | 1,417 | 5,22E-01 | 0 | 0,50333277 | 0,28260632 |
| AP4A_HUMA | NUDT2 | sp\|P50583\|AP4A_HUMAN Bis(5'-nucleosyl)-tetraphosphatase [asymmetrical] OS=Homo | 17,422 | 6,8 | 16829 | 1 | 20,24 | 18,67 | 21,77 | 19,31 | 20,76 | 22,11 | 1,415 | 6,99E-01 | 0 | 0,50072479 | 0,15526348 |
| S12A5_HUM | SLC12A5 | sp\|Q9H2X9\|S12A5_HUMAN Solute carrier family 12 member 5 OS=Homo sapiens GN=SI | 30,519 | 1,23 | 126184 | 1 | 20,84 | 21,19 | 19,36 | 20,83 | 20,89 | 21,17 | 1,414 | 4,30E-01 | 0 | 0,50003052 | 0,36621855 |
| TIF1B_HUMA | TRIM28 | sp\|Q13263\|TIF1B_HUMAN Transcription intermediary factor 1-beta OS=Homo sapiens | 710,161 | 26,35 | 88550 | 23 | 25,05 | 25,15 | 27,21 | 25,36 | 25,96 | 27,59 | 1,414 | 6,33E-01 | 0 | 0,50000064 | 0,19849776 |
| HABP2_HUM | HABP2 | sp\|Q14520\|HABP2_HUMAN Hyaluronan-binding protein 2 OS=Homo sapiens GN=HABP | 88,525 | 6,25 | 62672 | 4 | 23,41 | 22,71 | 25,21 | 23,83 | 23,61 | 25,39 | 1,414 | 6,20E-01 | 0 | 0,50000064 | 0,2075941 |
| DPY30_HUM | DPY30 | sp\|Q9C005\|DPY30_HUMAN Protein dpy-30 homolog OS=Homo sapiens GN=DPY30 PE=1 | 96,579 | 36,36 | 11250 | 2 | 19,65 | 20,04 | 21,09 | 20,62 | 20,45 | 21,21 | 1,414 | 3,63E-01 | 0 | 0,5 | 0,43975482 |
| GPD1L_HUM | GPD1L | sp\|Q8N335\|GPD1L_HUMAN Glycerol-3-phosphate dehydrogenase 1-like protein OS=Ho | 94,216 | 12,25 | 38419 | 4 | 21,92 | 22,50 | 22,36 | 22,78 | 22,98 | 22,52 | 1,414 | 8,52E-02 | 0 | 0,5 | 1,06969173 |
| CNO11_HUM | CNOT11 | sp\|Q9UKZ1\|CNO11_HUMAN CCR4-NOT transcription complex subunit 11 OS=Homo sap | 55,522 | 3,53 | 55215 | 1 | 17,03 | 18,15 | 19,67 | 18,58 | 17,86 | 19,90 | 1,411 | 6,35E-01 | 0 | 0,49717776 | 0,19689693 |
| PBDC1_HUM | PBDC1 | sp\|Q9BVG4\|PBDC1_HUMAN Protein PBDC1 OS=Homo sapiens GN=PBDC1 PE=1 SV=1 | 87,381 | 11,59 | 26057 | 2 | 21,13 | 21,75 | 19,78 | 21,65 | 21,69 | 20,81 | 1,411 | 4,86E-01 | 0 | 0,49666659 | 0,31297544 |
| PGM1_HUM | PGM1 | sp\|P36871\|PGM1_HUMAN Phosphoglucomutase-1 OS=Homo sapiens GN=PGM1 PE=1 S | 327,691 | 24,38 | 61449 | 9 | 24,20 | 24,07 | 22,98 | 24,60 | 24,47 | 23,67 | 1,411 | 3,63E-01 | 0 | 0,49666659 | 0,44040424 |
| PSMD5_HUM | PSMD5 | sp\|Q16401\|PSMD5_HUMAN 26S proteasome non-ATPase regulatory subunit 5 OS=Hom | 61,744 | 6,15 | 56196 | 3 | 23,45 | 23,06 | 22,67 | 23,70 | 23,92 | 23,05 | 1,411 | 2,23E-01 | 0 | 0,49666659 | 0,65129063 |
| MMGT1_HU | MMGT1 | sp\|Q8N4V1\|MMGT1_HUMAN Membrane magnesium transporter 1 OS=Homo sapiens | 45,828 | 16,03 | 14686 | 1 | 19,27 | 19,28 | 16,36 | 19,03 | 19,46 | 17,91 | 1,411 | 6,68E-01 | 0 | 0,49666595 | 0,1750036 |
| LYRIC_HUM | MTDH | sp\|Q86UE4\|LYRIC_HUMAN Protein LYRIC OS=Homo sapiens GN=MTDH PE=1 SV=2 | 57,697 | 2,75 | 63837 | 1 | 19,81 | 18,32 | 18,72 | 18,86 | 20,20 | 19,27 | 1,410 | 4,53E-01 | 0 | 0,4960537 | 0,34422133 |
| PUR8_HUMA | ADSL | sp\|P30566\|PUR8_HUMAN Adenylosuccinate lyase OS=Homo sapiens GN=ADSL PE=1 SV= | 296,785 | 21,49 | 54889 | 8 | 24,57 | 24,47 | 23,12 | 24,78 | 24,83 | 24,03 | 1,408 | 4,08E-01 | 0 | 0,49333382 | 0,38917915 |
| MTA1_HUM | MTA1 | sp\|Q13330\|MTA1_HUMAN Metastasis-associated protein MTA1 OS=Homo sapiens GN= | 110,187 | 7,83 | 80786 | 5 | 20,30 | 20,73 | 22,16 | 20,65 | 21,71 | 22,31 | 1,408 | 5,43E-01 | 0 | 0,49333318 | 0,26530981 |
| TCPH_HUMA | CCT7 | sp\|Q99832\|TCPH_HUMAN T-complex protein 1 subunit eta OS=Homo sapiens GN=CCT7 | 1506,016 | 57,09 | 59367 | 36 | 28,12 | 28,25 | 28,16 | 28,64 | 28,78 | 28,59 | 1,408 | 1,99E-03 | 0 | 0,49333318 | 2,70222637 |
| KCY_HUMAN | CMPK1 | sp\|P30085\|KCY_HUMAN UMP-CMP kinase OS=Homo sapiens GN=CMPK1 PE=1 SV=3 | 321,935 | 53,57 | 22222 | 9 | 24,77 | 25,23 | 24,96 | 25,48 | 25,32 | 25,64 | 1,408 | 3,84E-02 | 0 | 0,49333318 | 1,41541386 |
| P66B_HUMA | GATAD2B | sp\|Q8WXI9\|P66B_HUMAN Transcriptional repressor p66-beta OS=Homo sapiens GN=G | 47,031 | 3,88 | 65261 | 2 | 19,34 | 18,53 | 20,57 | 19,81 | 18,73 | 21,37 | 1,407 | 6,37E-01 | 0 | 0,49310239 | 0,19552801 |
| SBSN_HUMA | SBSN | sp\|Q6UWP8\|SBSN_HUMAN Suprabasin OS=Homo sapiens GN=SBSN PE=1 SV=2 | 79,112 | 15,25 | 60541 | 2 | 20,85 | 19,35 | 20,38 | 22,20 | 19,98 | 19,87 | 1,404 | 6,07E-01 | 0 | 0,49000041 | 0,2169109 |
| GARS_HUMA | GARS | sp\|P41250\|GARS_HUMAN Glycine--tRNA ligase OS=Homo sapiens GN=GARS PE=1 SV=3 | 663,804 | 34,91 | 83166 | 22 | 27,04 | 27,31 | 26,89 | 27,38 | 27,75 | 27,58 | 1,404 | 3,96E-02 | 0 | 0,48999977 | 1,40205897 |
| NAT10_HUM | NAT10 | sp\|Q9H0A0\|NAT10_HUMAN RNA cytidine acetyltransferase OS=Homo sapiens GN=NAT | 273,331 | 13,76 | 115730 | 10 | 21,39 | 18,69 | 24,11 | 20,50 | 20,61 | 24,55 | 1,404 | 8,23E-01 | 0 | 0,48999977 | 0,08447246 |
| TMED9_HUM | TMED9 | sp\|Q9BVK6\|TMED9_HUMAN Transmembrane emp24 domain-containing protein 9 OS=I | 211,519 | 37,87 | 27277 | 7 | 25,34 | 24,89 | 25,99 | 25,54 | 25,62 | 26,52 | 1,401 | 3,38E-01 | 0 | 0,48666763 | 0,47060451 |
| ATLA1_HUM | ATL1 | sp\|Q8WXF7\|ATLA1_HUMAN Atlastin-1 OS=Homo sapiens GN=ATL1 PE=1 SV=1 | 94,241 | 7,89 | 63544 | 4 | 22,31 | 20,93 | 22,17 | 22,48 | 22,69 | 21,70 | 1,401 | 4,12E-01 | 0 | 0,486667 | 0,3850505 |
| ACTC_HUMA | ACTC1 | sp\|P68032\|ACTC_HUMAN Actin, alpha cardiac muscle 1 OS=Homo sapiens GN=ACTC1 P | 3014,223 | 83,29 | 42019 | 60 | 24,43 | 27,07 | 24,31 | 26,53 | 26,07 | 24,67 | 1,401 | 6,70E-01 | 0 | 0,486667 | 0,17389302 |
| SYYC_HUMA | YARS | sp\|P54577\|SYYC_HUMAN Tyrosine--tRNA ligase, cytoplasmic OS=Homo sapiens GN=YAI | 349,239 | 31,82 | 59143 | 14 | 25,56 | 26,02 | 24,85 | 26,06 | 26,22 | 25,61 | 1,401 | 2,76E-01 | 0 | 0,48666636 | 0,55888591 |
| STMN1_HUM | STMN1 | sp\|P16949\|STMN1_HUMAN Stathmin OS=Homo sapiens GN=STMN1 PE=1 SV=3 | 63,25 | 15,44 | 17303 | 2 | 22,23 | 21,62 | 21,30 | 23,05 | 22,40 | 21,16 | 1,401 | 4,75E-01 | 0 | 0,48666636 | 0,3233547 |
| ERD22_HUM | KDELR2 | sp\|P33947\|ERD22_HUMAN ER lumen protein-retaining receptor 2 OS=Homo sapiens GN | 110,3 | 22,64 | 24422 | 4 | 20,92 | 21,33 | 20,43 | 20,76 | 21,66 | 21,72 | 1,401 | 2,96E-01 | 0 | 0,48666636 | 0,52893334 |
| MYG1_HUM | C12orf10 | sp\|Q9HB07\|MYG1_HUMAN UPF0160 protein MYG1, mitochondrial OS=Homo sapiens G | 120,568 | 12,77 | 42449 | 5 | 23,29 | 23,99 | 23,46 | 24,26 | 24,21 | 23,73 | 1,401 | 1,46E-01 | 0 | 0,48666636 | 0,83570064 |
| PSB6_HUMA | PSMB6 | sp\|P28072\|PSB6_HUMAN Proteasome subunit beta type-6 OS=Homo sapiens GN=PSME | 180,629 | 38,49 | 25358 | 5 | 25,84 | 25,88 | 25,36 | 26,24 | 26,38 | 25,92 | 1,401 | 8,69E-02 | 0 | 0,48666636 | 1,06122123 |
| KC1D_HUMA | CSNK1D | sp\|P48730\|KC1D_HUMAN Casein kinase I isoform delta OS=Homo sapiens GN=CSNK1D | 129,864 | 13,25 | 47330 | 5 | 21,37 | 17,85 | 20,57 | 21,26 | 20,82 | 19,17 | 1,400 | 7,15E-01 | 0 | 0,4853255 | 0,14550442 |
| CP51A_HUM | CYP51A1 | sp\|Q16850\|CP51A_HUMAN Lanosterol 14-alpha demethylase OS=Homo sapiens GN=CY | 19,475 | 2,19 | 56806 | 1 | 19,30 | 20,02 | 20,90 | 20,73 | 19,44 | 21,51 | 1,399 | 5,58E-01 | 0 | 0,48485756 | 0,25323696 |
| PP1R8_HUM | PPP1R8 | sp\|Q12972\|PP1R8_HUMAN Nuclear inhibitor of protein phosphatase 1 OS=Homo sapier | 30,803 | 11,11 | 38479 | 2 | 20,41 | 20,34 | 18,95 | 20,90 | 20,54 | 19,71 | 1,399 | 4,60E-01 | 0 | 0,48411115 | 0,33754586 |
| BUB3_HUMA | BUB3 | sp\|O43684\|BUB3_HUMAN Mitotic checkpoint protein BUB3 OS=Homo sapiens GN=BUB | 454,445 | 49,09 | 37155 | 13 | 25,60 | 25,02 | 26,51 | 25,76 | 26,04 | 26,78 | 1,398 | 4,13E-01 | 0 | 0,48333359 | 0,38387608 |
| SYWC_HUM | WARS | sp\|P23381\|SYWC_HUMAN Tryptophan--tRNA ligase, cytoplasmic OS=Homo sapiens GN= | 344,393 | 23,35 | 53165 | 10 | 25,65 | 25,51 | 24,00 | 25,75 | 26,14 | 24,72 | 1,398 | 5,15E-01 | 0 | 0,48333295 | 0,2883994 |
| HS90B_HUM | HSP90AB1 | sp\|P08238\|HS90B_HUMAN Heat shock protein HSP 90-beta OS=Homo sapiens GN=HSP | 2926,237 | 57,87 | 83264 | 59 | 29,21 | 29,33 | 29,55 | 29,66 | 29,79 | 30,08 | 1,395 | 3,93E-02 | 0 | 0,48000081 | 1,405591 |
| SAHH2_HUM | AHCYL1 | sp\|O43865\|SAHH2_HUMAN S-adenosylhomocysteine hydrolase-like protein 1 OS=Hom | 316,295 | 27,55 | 58951 | 13 | 21,53 | 20,55 | 20,50 | 20,42 | 21,75 | 21,85 | 1,395 | 4,47E-01 | 0 | 0,48000018 | 0,34958427 |
| BAP31_HUM | BCAP31 | sp\|P51572\|BAP31_HUMAN B-cell receptor-associated protein 31 OS=Homo sapiens GN= | 180,527 | 10,98 | 27992 | 7 | 24,21 | 24,01 | 24,54 | 24,64 | 24,77 | 24,79 | 1,395 | 4,11E-02 | 0 | 0,48000018 | 1,3864444 |
| RL8_HUMAN | RPL8 | sp\|P62917\|RL8_HUMAN 60S ribosomal protein L8 OS=Homo sapiens GN=RPL8 PE=1 SV= | 431,971 | 50,97 | 28025 | 17 | 26,69 | 26,51 | 27,67 | 27,32 | 26,61 | 28,38 | 1,395 | 4,87E-01 | 0 | 0,47999954 | 0,31222835 |
| RB3GP_HUM | RAB3GAP1 | sp\|Q15042\|RB3GP_HUMAN Rab3 GTPase-activating protein catalytic subunit OS=Homo | 37,619 | 2,14 | 110524 | 2 | 20,97 | 20,95 | 20,70 | 21,36 | 21,15 | 21,55 | 1,395 | 2,93E-02 | 0 | 0,47999954 | 1,53242201 |
| ZPR1_HUMA | ZPR1 | sp\|O75312\|ZPR1_HUMAN Zinc finger protein ZPR1 OS=Homo sapiens GN=ZPR1 PE=1 SV | 64,062 | 6,32 | 50925 | 2 | 18,88 | 17,34 | 21,82 | 17,76 | 19,57 | 22,14 | 1,392 | 8,07E-01 | 0 | 0,47686513 | 0,09313019 |
| GSH0_HUMA | GCLM | sp\|P48507\|GSH0_HUMAN Glutamate--cysteine ligase regulatory subunit OS=Homo sapi | 96,889 | 16,79 | 30727 | 4 | 22,88 | 22,56 | 23,17 | 23,10 | 23,24 | 23,70 | 1,392 | 1,32E-01 | 0 | 0,4766674 | 0,87823812 |
| FLII_HUMAN | FLII | sp\|Q13045\|FLII_HUMAN Protein flightless-1 homolog OS=Homo sapiens GN=FLII PE=1 S | 72,587 | 3,31 | 144751 | 4 | 22,73 | 22,90 | 23,34 | 23,37 | 23,54 | 23,49 | 1,392 | 6,49E-02 | 0 | 0,4766674 | 1,1880016 |
| UMPS_HUM | UMPS | sp\|P11172\|UMPS_HUMAN Uridine 5'-monophosphate synthase OS=Homo sapiens GN= | 148,552 | 13,12 | 52222 | 5 | 23,33 | 23,33 | 23,84 | 23,68 | 23,77 | 24,48 | 1,392 | 1,93E-01 | 0 | 0,47666677 | 0,71465029 |
| NONO_HUM | NONO | sp\|Q15233\|NONO_HUMAN Non-POU domain-containing octamer-binding protein OS=I | 401,39 | 27,6 | 54232 | 16 | 25,28 | 25,43 | 26,38 | 25,78 | 25,69 | 27,05 | 1,392 | 4,41E-01 | 0 | 0,47666677 | 0,35541169 |
| ERLN1_HUM | ERLIN1 | sp\|O75477\|ERLN1_HUMAN Erlin-1 OS=Homo sapiens GN=ERLIN1 PE=1 SV=1 | 201,138 | 19,36 | 38926 | 6 | 21,76 | 21,85 | 22,30 | 21,98 | 22,39 | 22,97 | 1,392 | 2,25E-01 | 0 | 0,47666613 | 0,64843618 |
| GLRX3_HUM | GLRX3 | sp\|O76003\|GLRX3_HUMAN Glutaredoxin-3 OS=Homo sapiens GN=GLRX3 PE=1 SV=2 | 246,081 | 22,09 | 37432 | 7 | 29,11 | 25,36 | 29,35 | 29,41 | 27,13 | 28,71 | 1,392 | 7,60E-01 | 0 | 0,4766655 | 0,11918846 |
| PPAC_HUMA | ACP1 | sp\|P24666\|PPAC_HUMAN Low molecular weight phosphotyrosine protein phosphatase | 24,922 | 5,7 | 18042 | 1 | 22,81 | 22,82 | 22,59 | 23,24 | 23,32 | 23,08 | 1,388 | 1,01E-02 | 0 | 0,47333336 | 1,9970302 |
| SSRD_HUMA | SSR4 | sp\|P51571\|SSRD_HUMAN Translocon-associated protein subunit delta OS=Homo sapier | 123,112 | 23,12 | 18999 | 3 | 24,69 | 24,40 | 25,57 | 24,92 | 25,24 | 25,92 | 1,388 | 3,61E-01 | 0 | 0,47333336 | 0,44285459 |
| TMED3_HUM | TMED3 | sp\|Q9Y3Q3\|TMED3_HUMAN Transmembrane emp24 domain-containing protein 3 OS=I | 25,485 | 4,61 | 24777 | 1 | 20,79 | 20,17 | 22,10 | 20,93 | 20,96 | 22,59 | 1,388 | 5,81E-01 | 0 | 0,47333272 | 0,23550053 |
| CDC23_HUM | CDC23 | sp\|Q9UJX2\|CDC23_HUMAN Cell division cycle protein 23 homolog OS=Homo sapiens GI | 116,457 | 9,21 | 68834 | 5 | 21,70 | 21,62 | 23,30 | 22,02 | 22,63 | 23,39 | 1,388 | 5,22E-01 | 0 | 0,47333272 | 0,28220785 |
| KITH_HUMA | TK1 | sp\|P04183\|KITH_HUMAN Thymidine kinase, cytosolic OS=Homo sapiens GN=TK1 PE=1 S | 15,697 | 6,84 | 25469 | 1 | 20,29 | 20,20 | 19,45 | 20,70 | 20,32 | 20,34 | 1,387 | 1,82E-01 | 0 | 0,47230657 | 0,74049313 |
| APEX1_HUM | APEX1 | sp\|P27695\|APEX1_HUMAN DNA-(apurinic or apyrimidinic site) lyase OS=Homo sapiens | 206,397 | 22,96 | 35555 | 5 | 23,56 | 24,42 | 23,68 | 24,00 | 24,79 | 24,28 | 1,385 | 2,56E-01 | 0 | 0,47000058 | 0,59220748 |
| TBA1C_HUM | TUBA1C | sp\|Q9BQE3\|TBA1C_HUMAN Tubulin alpha-1C chain OS=Homo sapiens GN=TUBA1C PE= | 2041,443 | 83,96 | 49895 | 38 | 22,23 | 22,36 | 24,37 | 22,69 | 23,31 | 24,37 | 1,385 | 6,09E-01 | 0 | 0,46999995 | 0,21520791 |
| SC24D_HUM | SEC24D | sp\|O94855\|SC24D_HUMAN Protein transport protein Sec24D OS=Homo sapiens GN=SE | 20,763 | 3,2 | 113010 | 2 | 22,51 | 22,00 | 21,96 | 22,77 | 22,99 | 22,11 | 1,382 | 2,16E-01 | 0 | 0,46666718 | 0,6647695 |
| PSA4_HUMA | PSMA4 | sp\|P25789\|PSA4_HUMAN Proteasome subunit alpha type-4 OS=Homo sapiens GN=PSM | 600,653 | 70,88 | 29484 | 14 | 25,88 | 26,00 | 26,19 | 26,15 | 26,61 | 26,71 | 1,382 | 7,45E-02 | 0 | 0,46666654 | 1,12764838 |
| TMED7_HUM | TMED7 | sp\|Q9Y3B3\|TMED7_HUMAN Transmembrane emp24 domain-containing protein 7 OS=I | 238,497 | 32,14 | 25172 | 5 | 24,95 | 24,05 | 25,63 | 24,88 | 25,08 | 26,07 | 1,382 | 4,71E-01 | 0 | 0,46666654 | 0,32678479 |
| PSMD3_HUM | PSMD3 | sp\|O43242\|PSMD3_HUMAN 26S proteasome non-ATPase regulatory subunit 3 OS=Hom | 936,24 | 47,38 | 60978 | 29 | 26,97 | 27,10 | 27,90 | 27,39 | 27,59 | 28,39 | 1,382 | 3,31E-01 | 0 | 0,46666654 | 0,48072365 |
| MSH2_HUM | MSH2 | sp\|P43246\|MSH2_HUMAN DNA mismatch repair protein Msh2 OS=Homo sapiens GN=N | 159,518 | 4,39 | 104743 | 3 | 18,40 | 18,87 | 20,84 | 18,45 | 19,21 | 21,84 | 1,379 | 7,34E-01 | 0 | 0,46333313 | 0,13445071 |
| MFGM_HUM | MFGE8 | sp\|Q08431\|MFGM_HUMAN Lactadherin OS=Homo sapiens GN=MFGE8 PE=1 SV=2 | 613,157 | 49,35 | 43123 | 16 | 28,35 | 28,15 | 28,60 | 28,62 | 28,71 | 29,16 | 1,379 | 9,39E-02 | 0 | 0,46333313 | 1,02730696 |
| BZW1_HUM | BZW1 | sp\|Q7L1Q6\|BZW1_HUMAN Basic leucine zipper and W2 domain-containing protein 1 O | 487,189 | 46,78 | 48043 | 14 | 23,05 | 23,37 | 25,57 | 23,64 | 23,90 | 25,84 | 1,379 | 6,83E-01 | 0 | 0,46333313 | 0,16577099 |
| SYFB_HUMA | FARSB | sp\|Q9NSD9\|SYFB_HUMAN Phenylalanine--tRNA ligase beta subunit OS=Homo sapiens G | 73,311 | 9 | 66116 | 5 | 22,29 | 22,78 | 22,91 | 22,43 | 23,42 | 23,52 | 1,379 | 3,07E-01 | 0 | 0,46333313 | 0,513266 |
| MLEC_HUM | MLEC | sp\|Q14165\|MLEC_HUMAN Malectin OS=Homo sapiens GN=MLEC PE=1 SV=1 | 215,295 | 27,4 | 32234 | 6 | 22,44 | 22,24 | 24,78 | 22,63 | 23,18 | 25,04 | 1,379 | 6,94E-01 | 0 | 0,46333313 | 0,15885466 |
| EMC7_HUM | EMC7 | sp\|Q9NPA0\|EMC7_HUMAN ER membrane protein complex subunit 7 OS=Homo sapiens | 43,384 | 9,5 | 26471 | 2 | 22,31 | 21,63 | 22,27 | 22,58 | 22,40 | 22,61 | 1,376 | 1,16E-01 | 0 | 0,46000036 | 0,93573088 |

| | | | | | | | | | | | | | | | | | |
|---|---|---|---|---|---|---|---|---|---|---|---|---|---|---|---|---|---|
| PAIRB_HUM/ | SERBP1 | sp\|Q8NC51\|PAIRB_HUMAN Plasminogen activator inhibitor 1 RNA-binding protein OS=H | 134,789 | 17,16 | 44965 | 6 | 23,58 | 23,95 | 23,63 | 24,27 | 24,27 | 24,00 | **1,376** | 3,50E-02 | **0** | 0,46000036 | 1,45562396 |
| AIMP1_HUM | AIMP1 | sp\|Q12904\|AIMP1_HUMAN Aminoacyl tRNA synthase complex-interacting multifunctio | 73,056 | 9,62 | 34353 | 2 | 21,13 | 21,09 | 22,73 | 21,56 | 22,04 | 22,73 | **1,376** | 5,11E-01 | **0** | 0,46000036 | 0,29175635 |
| CD109_HUM | CD109 | sp\|Q6YHK3\|CD109_HUMAN CD109 antigen OS=Homo sapiens GN=CD109 PE=1 SV=2 | 110,133 | 3,46 | 161689 | 6 | 24,47 | 23,81 | 23,06 | 24,13 | 24,17 | 24,42 | **1,376** | 3,32E-01 | **0** | 0,46000036 | 0,4786308 |
| IF4B_HUMAI | EIF4B | sp\|P23588\|IF4B_HUMAN Eukaryotic translation initiation factor 4B OS=Homo sapiens GI | 73,491 | 8,84 | 69151 | 3 | 22,28 | 21,05 | 21,54 | 22,26 | 22,25 | 21,74 | **1,376** | 3,11E-01 | **0** | 0,45999972 | 0,50777372 |
| EMC2_HUM/ | EMC2 | sp\|Q15006\|EMC2_HUMAN ER membrane protein complex subunit 2 OS=Homo sapiens | 188,337 | 21,89 | 34834 | 5 | 22,76 | 21,89 | 23,95 | 23,18 | 22,74 | 24,06 | **1,376** | 5,53E-01 | **0** | 0,45999972 | 0,25690201 |
| IPYR_HUMAI | PPA1 | sp\|Q15181\|IPYR_HUMAN Inorganic pyrophosphatase OS=Homo sapiens GN=PPA1 PE=1 | 706,964 | 66,78 | 32660 | 16 | 26,60 | 27,19 | 26,46 | 26,92 | 27,63 | 27,08 | **1,376** | 2,12E-01 | **0** | 0,45999972 | 0,67299898 |
| TCPG_HUMA | CCT3 | sp\|P49368\|TCPG_HUMAN T-complex protein 1 subunit gamma OS=Homo sapiens GN=C | 1015,572 | 46,79 | 60534 | 33 | 28,20 | 28,50 | 28,08 | 28,69 | 28,83 | 28,64 | **1,376** | 2,85E-02 | **0** | 0,45999972 | 1,54488305 |
| NCPR_HUM/ | POR | sp\|P16435\|NCPR_HUMAN NADPH--cytochrome P450 reductase OS=Homo sapiens GN=I | 259,019 | 14,48 | 76690 | 8 | 24,32 | 24,68 | 25,52 | 24,98 | 25,12 | 25,80 | **1,376** | 3,51E-01 | **0** | 0,45999972 | 0,45419056 |
| DPYL3_HUM | DPYSL3 | sp\|Q14195\|DPYL3_HUMAN Dihydropyrimidinase-related protein 3 OS=Homo sapiens GI | 286,089 | 16,14 | 61963 | 8 | 22,90 | 23,24 | 19,59 | 23,22 | 23,49 | 20,39 | **1,373** | 7,80E-01 | **0** | 0,45710246 | 0,10794942 |
| ACSL3_HUM | ACSL3 | sp\|O95573\|ACSL3_HUMAN Long-chain-fatty-acid--CoA ligase 3 OS=Homo sapiens GN=A | 217,018 | 10 | 80420 | 6 | 24,12 | 23,49 | 23,25 | 23,93 | 23,71 | 24,59 | **1,372** | 2,85E-01 | **0** | 0,45666631 | 0,54494104 |
| RIFK_HUMAI | RFK | sp\|Q969G6\|RIFK_HUMAN Riboflavin kinase OS=Homo sapiens GN=RFK PE=1 SV=2 | 96,537 | 13,55 | 17623 | 2 | 19,11 | 19,43 | 20,66 | 19,48 | 20,05 | 21,04 | **1,372** | 5,25E-01 | **0** | 0,45666631 | 0,27989417 |
| TM9S3_HUN | TM9SF3 | sp\|Q9HD45\|TM9S3_HUMAN Transmembrane 9 superfamily member 3 OS=Homo sapiei | 255,508 | 18 | 67888 | 9 | 24,69 | 24,25 | 25,82 | 24,98 | 25,10 | 26,04 | **1,369** | 4,75E-01 | **0** | 0,45333354 | 0,32352951 |
| UBR4_HUM/ | UBR4 | sp\|Q5T4S7\|UBR4_HUMAN E3 ubiquitin-protein ligase UBR4 OS=Homo sapiens GN=UBR4 | 844,487 | 8,14 | 573841 | 27 | 23,20 | 23,87 | 24,59 | 23,77 | 24,16 | 25,09 | **1,369** | 4,64E-01 | **0** | 0,4533329 | 0,33335707 |
| CDC20_HUM | CDC20 | sp\|Q12834\|CDC20_HUMAN Cell division cycle protein 20 homolog OS=Homo sapiens GI | 32,749 | 5,61 | 54723 | 2 | 24,35 | 23,86 | 24,97 | 24,75 | 24,80 | 24,99 | **1,369** | 2,41E-01 | **0** | 0,4533329 | 0,61845017 |
| GCP60_HUM | ACBD3 | sp\|Q9H3P7\|GCP60_HUMAN Golgi resident protein GCP60 OS=Homo sapiens GN=ACBD3 | 59,437 | 6,44 | 60593 | 3 | 21,35 | 22,17 | 21,54 | 21,82 | 22,30 | 22,30 | **1,369** | 1,99E-01 | **0** | 0,45333227 | 0,7008348 |
| AT1B3_HUM | ATP1B3 | sp\|P54709\|AT1B3_HUMAN Sodium/potassium-transporting ATPase subunit beta-3 OS=I | 45,944 | 11,83 | 31513 | 3 | 24,35 | 24,79 | 23,41 | 25,03 | 24,96 | 23,91 | **1,366** | 4,55E-01 | **0** | 0,44999949 | 0,34173855 |
| ANXA3_HUN | ANXA3 | sp\|P12429\|ANXA3_HUMAN Annexin A3 OS=Homo sapiens GN=ANXA3 PE=1 SV=3 | 341,988 | 29,72 | 36375 | 10 | 27,25 | 27,07 | 26,60 | 27,57 | 27,65 | 27,05 | **1,366** | 1,71E-01 | **0** | 0,44999949 | 0,76710532 |
| PSA3_HUMA | PSMA3 | sp\|P25788\|PSA3_HUMAN Proteasome subunit alpha type-3 OS=Homo sapiens GN=PSM | 508,655 | 42,35 | 28433 | 11 | 26,33 | 26,26 | 25,93 | 26,39 | 26,77 | 26,71 | **1,366** | 5,78E-02 | **0** | 0,44999949 | 1,23833357 |
| AMERL_HUN | AMMECR1L | sp\|Q6DCA0\|AMERL_HUMAN AMMECR1-like protein OS=Homo sapiens GN=AMMECR1L | 10,203 | 3,23 | 34499 | 1 | 20,98 | 18,66 | 20,19 | 20,79 | 20,34 | 20,04 | **1,364** | 5,66E-01 | **0** | 0,44773801 | 0,24745583 |
| RTCB_HUMA | RTCB | sp\|Q9Y3I0\|RTCB_HUMAN tRNA-splicing ligase RtcB homolog OS=Homo sapiens GN=RTC | 574,691 | 40,99 | 55210 | 17 | 25,69 | 25,87 | 25,99 | 26,04 | 26,20 | 26,65 | **1,363** | 9,19E-02 | **0** | 0,44666672 | 1,03673809 |
| IPO5_HUMAI | IPO5 | sp\|O00410\|IPO5_HUMAN Importin-5 OS=Homo sapiens GN=IPO5 PE=1 SV=4 | 1347,792 | 38,74 | 123630 | 31 | 26,33 | 26,63 | 26,27 | 26,66 | 27,09 | 26,82 | **1,363** | 5,62E-02 | **0** | 0,44666672 | 1,24992145 |
| EF1G_HUMA | EEF1G | sp\|P26641\|EF1G_HUMAN Elongation factor 1-gamma OS=Homo sapiens GN=EEF1G PE= | 1066,491 | 59,73 | 50119 | 23 | 28,81 | 28,86 | 28,74 | 29,05 | 29,35 | 29,35 | **1,363** | 1,35E-02 | **0** | 0,44666672 | 1,8698635 |
| UBQL1_HUN | UBQLN1 | sp\|Q9UMX0\|UBQL1_HUMAN Ubiquilin-1 OS=Homo sapiens GN=UBQLN1 PE=1 SV=2 | 208,525 | 18,34 | 62519 | 7 | 21,53 | 22,64 | 18,35 | 21,85 | 22,40 | 19,61 | **1,362** | 7,87E-01 | **0** | 0,446256 | 0,10421955 |
| USO1_HUM/ | USO1 | sp\|O60763\|USO1_HUMAN General vesicular transport factor p115 OS=Homo sapiens G | 156,441 | 6,76 | 107895 | 5 | 22,93 | 23,30 | 22,42 | 23,45 | 23,43 | 23,10 | **1,360** | 1,88E-01 | **0** | 0,44333394 | 0,72696198 |
| RSSA_HUMA | RPSA | sp\|P08865\|RSSA_HUMAN 40S ribosomal protein SA OS=Homo sapiens GN=RPSA PE=1 S' | 705,596 | 54,92 | 32854 | 19 | 27,56 | 27,73 | 28,50 | 27,93 | 28,11 | 29,08 | **1,360** | 3,89E-01 | **0** | 0,44333394 | 0,40970399 |
| THOC4_HUN | ALYREF | sp\|Q86V81\|THOC4_HUMAN THO complex subunit 4 OS=Homo sapiens GN=ALYREF PE= | 66,684 | 14,79 | 26888 | 3 | 22,31 | 22,86 | 21,48 | 22,97 | 23,02 | 21,99 | **1,360** | 4,44E-01 | **0** | 0,44333331 | 0,35242599 |
| LRC41_HUM | LRRC41 | sp\|Q15345\|LRC41_HUMAN Leucine-rich repeat-containing protein 41 OS=Homo sapien | 27,322 | 2,22 | 88650 | 1 | 17,93 | 18,11 | 19,66 | 18,36 | 18,72 | 19,95 | **1,360** | 5,77E-01 | **0** | 0,44333331 | 0,23916086 |
| TRAP1_HUM | TRAP1 | sp\|Q12931\|TRAP1_HUMAN Heat shock protein 75 kDa, mitochondrial OS=Homo sapien | 103,219 | 3,12 | 80110 | 2 | 26,33 | 26,59 | 27,55 | 26,82 | 27,27 | 27,71 | **1,360** | 3,82E-01 | **0** | 0,44333331 | 0,41847188 |
| CHIP_HUMAI | STUB1 | sp\|Q9UNE7\|CHIP_HUMAN E3 ubiquitin-protein ligase CHIP OS=Homo sapiens GN=STUB | 63,197 | 12,54 | 34856 | 3 | 22,53 | 21,71 | 19,15 | 22,47 | 23,00 | 19,25 | **1,360** | 7,89E-01 | **0** | 0,44333331 | 0,10273647 |
| RAN_HUMAI | RAN | sp\|P62826\|RAN_HUMAN GTP-binding nuclear protein Ran OS=Homo sapiens GN=RAN F | 695,423 | 76,85 | 24423 | 19 | 28,66 | 28,69 | 29,28 | 28,99 | 29,08 | 29,89 | **1,360** | 2,74E-01 | **0** | 0,44333267 | 0,56185188 |
| SYIC_HUMAI | IARS | sp\|P41252\|SYIC_HUMAN Isoleucine--tRNA ligase, cytoplasmic OS=Homo sapiens GN=IAI | 396,627 | 18,23 | 144498 | 21 | 25,54 | 25,69 | 26,40 | 25,82 | 26,33 | 26,81 | **1,360** | 3,19E-01 | **0** | 0,44333267 | 0,49614883 |
| KHDR1_HUN | KHDRBS1 | sp\|Q07666\|KHDR1_HUMAN KH domain-containing, RNA-binding, signal transduction-as | 102,733 | 12,19 | 48227 | 6 | 23,03 | 23,59 | 24,19 | 23,64 | 23,72 | 24,78 | **1,360** | 4,23E-01 | **0** | 0,44333267 | 0,37374016 |
| DYLT3_HUM | DYNLT3 | sp\|P51808\|DYLT3_HUMAN Dynein light chain Tctex-type 3 OS=Homo sapiens GN=DYNL | 36,697 | 20,69 | 13062 | 1 | 20,48 | 20,94 | 19,25 | 19,99 | 20,97 | 21,03 | **1,357** | 5,08E-01 | **0** | 0,44050916 | 0,29388857 |
| MATR3_HUN | MATR3 | sp\|P43243\|MATR3_HUMAN Matrin-3 OS=Homo sapiens GN=MATR3 PE=1 SV=2 | 367,905 | 22,67 | 94623 | 14 | 22,88 | 23,56 | 26,55 | 23,95 | 23,83 | 26,53 | **1,357** | 7,74E-01 | **0** | 0,44000117 | 0,11140356 |
| ADRM1_HUI | ADRM1 | sp\|Q16186\|ADRM1_HUMAN Proteasomal ubiquitin receptor ADRM1 OS=Homo sapiens | 19,381 | 2,21 | 42153 | 1 | 22,14 | 22,14 | 22,28 | 22,23 | 22,78 | 22,87 | **1,357** | 9,88E-02 | **0** | 0,44000053 | 1,00503631 |
| RL28_HUMA | RPL28 | sp\|P46779\|RL28_HUMAN 60S ribosomal protein L28 OS=Homo sapiens GN=RPL28 PE=1 | 209,363 | 40,88 | 15748 | 7 | 26,06 | 25,54 | 26,52 | 26,45 | 26,04 | 26,95 | **1,357** | 3,18E-01 | **0** | 0,44000053 | 0,49692141 |
| DAZP1_HUM | DAZAP1 | sp\|Q96EP5\|DAZP1_HUMAN DAZ-associated protein 1 OS=Homo sapiens GN=DAZAP1 PE | 150,178 | 19,16 | 43383 | 5 | 23,79 | 23,97 | 24,57 | 24,38 | 24,40 | 24,87 | **1,357** | 1,98E-01 | **0** | 0,4399999 | 0,70439077 |
| MAGT1_HUI | MAGT1 | sp\|Q9H0U3\|MAGT1_HUMAN Magnesium transporter protein 1 OS=Homo sapiens GN=I | 19,017 | 2,69 | 38037 | 1 | 22,56 | 22,61 | 22,10 | 22,99 | 23,11 | 22,49 | **1,357** | 1,53E-01 | **0** | 0,4399999 | 0,81548708 |
| FUBP1_HUM | FUBP1 | sp\|Q96AE4\|FUBP1_HUMAN Far upstream element-binding protein 1 OS=Homo sapiens | 172,771 | 8,85 | 67560 | 6 | 23,94 | 23,91 | 23,77 | 24,69 | 24,08 | 24,17 | **1,357** | 8,95E-02 | **0** | 0,4399999 | 1,04822015 |
| TPP2_HUMA | TPP2 | sp\|P29144\|TPP2_HUMAN Tripeptidyl-peptidase 2 OS=Homo sapiens GN=TPP2 PE=1 SV= | 260,336 | 7,53 | 138350 | 8 | 23,99 | 23,93 | 23,74 | 24,21 | 24,31 | 24,46 | **1,357** | 1,37E-02 | **0** | 0,43999926 | 1,86471435 |
| BCCIP_HUM/ | BCCIP | sp\|Q9P287\|BCCIP_HUMAN BRCA2 and CDKN1A-interacting protein OS=Homo sapiens G | 31,362 | 2,55 | 35979 | 1 | 20,96 | 21,04 | 22,20 | 21,06 | 21,72 | 22,74 | **1,357** | 5,25E-01 | **0** | 0,43999926 | 0,28017743 |
| TAP1_HUMA | TAP1 | sp\|Q03518\|TAP1_HUMAN Antigen peptide transporter 1 OS=Homo sapiens GN=TAP1 PI | 40,653 | 1,36 | 87218 | 1 | 18,62 | 19,40 | 19,96 | 19,04 | 20,25 | 20,01 | **1,356** | 4,57E-01 | **0** | 0,4395256 | 0,33968422 |
| HSP74_HUM | HSPA4 | sp\|P34932\|HSP74_HUMAN Heat shock 70 kDa protein 4 OS=Homo sapiens GN=HSPA4 P | 1565,589 | 47,26 | 94331 | 37 | 27,59 | 27,73 | 27,72 | 27,94 | 28,00 | 28,41 | **1,353** | 4,74E-02 | **0** | 0,43666712 | 1,32378141 |
| RANG_HUM/ | RANBP1 | sp\|P43487\|RANG_HUMAN Ran-specific GTPase-activating protein OS=Homo sapiens GN | 241,305 | 38,31 | 23310 | 7 | 26,46 | 26,52 | 26,55 | 26,89 | 27,00 | 26,95 | **1,353** | 4,56E-04 | **0** | 0,43666712 | 3,34143015 |
| EWS_HUMAI | EWSR1 | sp\|Q01844\|EWS_HUMAN RNA-binding protein EWS OS=Homo sapiens GN=EWSR1 PE=1 | 149,573 | 7,32 | 68478 | 3 | 23,15 | 23,19 | 22,29 | 23,58 | 23,70 | 22,66 | **1,353** | 3,78E-01 | **0** | 0,43666649 | 0,42284466 |
| APC7_HUMA | ANAPC7 | sp\|Q9UJX3\|APC7_HUMAN Anaphase-promoting complex subunit 7 OS=Homo sapiens G | 58,834 | 6,01 | 66855 | 2 | 20,42 | 20,79 | 21,23 | 21,25 | 20,27 | 22,23 | **1,353** | 5,15E-01 | **0** | 0,43666649 | 0,28805516 |
| H2A1B_HUM | HIST1H2AB | sp\|P04908\|H2A1B_HUMAN Histone H2A type 1-B/E OS=Homo sapiens GN=HIST1H2AB F | 264,582 | 57,69 | 14135 | 5 | 22,75 | 22,63 | 22,80 | 23,71 | 23,05 | 22,73 | **1,353** | 2,10E-01 | **0** | 0,43666649 | 0,67724207 |
| SYK_HUMAN | KARS | sp\|Q15046\|SYK_HUMAN Lysine--tRNA ligase OS=Homo sapiens GN=KARS PE=1 SV=3 | 288,39 | 20,94 | 68048 | 12 | 25,36 | 25,42 | 25,80 | 25,80 | 25,74 | 26,35 | **1,353** | 1,40E-01 | **0** | 0,43666649 | 0,85237176 |
| IQGA1_HUM | IQGAP1 | sp\|P46940\|IQGA1_HUMAN Ras GTPase-activating-like protein IQGAP1 OS=Homo sapien | 2370,887 | 41,7 | 189252 | 52 | 30,29 | 28,66 | 28,90 | 30,35 | 30,11 | 28,69 | **1,350** | 5,83E-01 | **0** | 0,43333371 | 0,23469086 |
| PSDE_HUMA | PSMD14 | sp\|O00487\|PSDE_HUMAN 26S proteasome non-ATPase regulatory subunit 14 OS=Homo | 641,445 | 53,23 | 34577 | 14 | 24,68 | 24,92 | 26,31 | 25,19 | 25,37 | 26,65 | **1,350** | 5,61E-01 | **0** | 0,43333371 | 0,2507157 |
| AP1G1_HUM | AP1G1 | sp\|O43747\|AP1G1_HUMAN AP-1 complex subunit gamma-1 OS=Homo sapiens GN=AP1 | 164,886 | 13,75 | 91351 | 8 | 24,09 | 23,73 | 23,61 | 24,03 | 24,39 | 24,31 | **1,350** | 7,47E-02 | **0** | 0,43333308 | 1,12676638 |
| RL21_HUMA | RPL21 | sp\|P46778\|RL21_HUMAN 60S ribosomal protein L21 OS=Homo sapiens GN=RPL21 PE=1 | 236,219 | 38,75 | 18565 | 7 | 26,93 | 26,78 | 26,34 | 27,19 | 26,93 | 27,23 | **1,350** | 9,67E-02 | **0** | 0,43333308 | 1,01448787 |
| FBX50_HUM | NCCRP1 | sp\|Q6ZVX7\|FBX50_HUMAN F-box only protein 50 OS=Homo sapiens GN=NCCRP1 PE=1 | 26,303 | 4 | 30847 | 1 | 20,96 | 19,15 | 19,20 | 22,01 | 19,13 | 19,47 | **1,350** | 7,10E-01 | **0** | 0,43333308 | 0,14854542 |
| NU155_HUN | NUP155 | sp\|O75694\|NU155_HUMAN Nuclear pore complex protein Nup155 OS=Homo sapiens G | 251,262 | 9,06 | 155199 | 9 | 22,37 | 22,27 | 23,95 | 22,31 | 23,38 | 24,20 | **1,350** | 6,04E-01 | **0** | 0,43333244 | 0,21868437 |
| ITPA_HUMAI | ITPA | sp\|Q9BY32\|ITPA_HUMAN Inosine triphosphate pyrophosphatase OS=Homo sapiens GN | 114,268 | 27,32 | 21446 | 4 | 24,64 | 24,88 | 21,52 | 24,78 | 24,94 | 22,61 | **1,347** | 7,61E-01 | **0** | 0,43000094 | 0,11889853 |
| TINAL_HUM/ | TINAGL1 | sp\|Q9GZM7\|TINAL_HUMAN Tubulointerstitial nephritis antigen-like OS=Homo sapiens ( | 170,117 | 7,92 | 52387 | 3 | 23,87 | 22,96 | 25,48 | 24,48 | 23,53 | 25,59 | **1,347** | 6,73E-01 | **0** | 0,43000031 | 0,17173583 |
| HS12B_HUM | HSPA12B | sp\|Q96MM6\|HS12B_HUMAN Heat shock 70 kDa protein 12B OS=Homo sapiens GN=HSF | 21,809 | 1,46 | 75688 | 1 | 20,10 | 20,28 | 20,59 | 20,33 | 20,62 | 21,31 | **1,347** | 2,55E-01 | **0** | 0,42999967 | 0,5932859 |
| FA98B_HUM | FAM98B | sp\|Q52LJ0\|FA98B_HUMAN Protein FAM98B OS=Homo sapiens GN=FAM98B PE=1 SV=1 | 53,819 | 5,76 | 37191 | 2 | 18,35 | 18,00 | 22,33 | 18,87 | 18,50 | 22,59 | **1,346** | 8,33E-01 | **0** | 0,42851957 | 0,07929264 |
| EF1B_HUMA | EEF1B2 | sp\|P24534\|EF1B_HUMAN Elongation factor 1-beta OS=Homo sapiens GN=EEF1B2 PE=1 | 272,518 | 44,44 | 24764 | 8 | 26,30 | 26,81 | 25,52 | 26,85 | 27,03 | 26,03 | **1,344** | 4,29E-01 | **0** | 0,42666753 | 0,36769163 |
| ANM1_HUM | PRMT1 | sp\|Q99873\|ANM1_HUMAN Protein arginine N-methyltransferase 1 OS=Homo sapiens G | 718,619 | 59,83 | 41516 | 18 | 26,58 | 26,65 | 26,50 | 26,44 | 27,24 | 27,33 | **1,344** | 2,10E-01 | **0** | 0,4266669 | 0,67731662 |
| FIBP_HUMAI | FIBP | sp\|O43427\|FIBP_HUMAN Acidic fibroblast growth factor intracellular-binding protein O! | 32,973 | 6,04 | 41878 | 2 | 22,09 | 21,38 | 20,96 | 22,22 | 22,33 | 21,16 | **1,344** | 4,40E-01 | **0** | 0,4266669 | 0,35677044 |
| GSTM2_HUN | GSTM2 | sp\|P28161\|GSTM2_HUMAN Glutathione S-transferase Mu 2 OS=Homo sapiens GN=GST | 127,98 | 9,63 | 25745 | 3 | 22,57 | 22,53 | 20,69 | 22,74 | 22,75 | 21,58 | **1,344** | 5,91E-01 | **0** | 0,42666626 | 0,2283426 |
| RL19_HUMA | RPL19 | sp\|P84098\|RL19_HUMAN 60S ribosomal protein L19 OS=Homo sapiens GN=RPL19 PE=1 | 190,645 | 26,02 | 23466 | 6 | 26,10 | 25,74 | 25,37 | 26,35 | 25,92 | 26,22 | **1,344** | 1,58E-01 | **0** | 0,42666626 | 0,80094181 |
| CJ076_HUM/ | C10orf76 | sp\|Q5T2E6\|CJ076_HUMAN UPF0668 protein C10orf76 OS=Homo sapiens GN=C10orf76 | 77,247 | 6,1 | 78710 | 2 | 19,42 | 20,27 | 18,77 | 19,54 | 20,05 | 20,15 | **1,343** | 4,19E-01 | **0** | 0,42570623 | 0,37791238 |
| DTX2_HUMA | DTX2 | sp\|Q86UW9\|DTX2_HUMAN Probable E3 ubiquitin-protein ligase DTX2 OS=Homo sapien | 29,009 | 3,86 | 67246 | 1 | 17,93 | 18,79 | 17,33 | 17,83 | 17,67 | 19,82 | **1,341** | 6,29E-01 | **0** | 0,42353948 | 0,20108679 |
| KATL2_HUM | KATNAL2 | sp\|Q8IYT4\|KATL2_HUMAN Katanin p60 ATPase-containing subunit A-like 2 OS=Homo sa | 37,604 | 2,23 | 61253 | 1 | 24,94 | 25,35 | 24,27 | 25,49 | 25,66 | 24,68 | **1,341** | 3,87E-01 | **0** | 0,42333285 | 0,41229035 |
| TAF5L_HUM | TAF5L | sp\|O75529\|TAF5L_HUMAN TAF5-like RNA polymerase II p300/CBP-associated factor-as | 20,891 | 1,7 | 66155 | 1 | 19,32 | 18,88 | 19,23 | 20,34 | 19,36 | 18,99 | **1,339** | 3,77E-01 | **0** | 0,42165057 | 0,42382507 |
| S38A1_HUM | SLC38A1 | sp\|Q9H2H9\|S38A1_HUMAN Sodium-coupled neutral amino acid transporter 1 OS=Hom | 176,736 | 16,63 | 54048 | 8 | 24,59 | 24,37 | 24,76 | 25,14 | 24,89 | 24,95 | **1,338** | 3,64E-02 | **0** | 0,41999944 | 1,43868488 |
| TRXR1_HUM | TXNRD1 | sp\|Q16881\|TRXR1_HUMAN Thioredoxin reductase 1, cytoplasmic OS=Homo sapiens GN | 210,212 | 12,48 | 70906 | 6 | 23,12 | 24,24 | 23,40 | 23,64 | 24,38 | 24,00 | **1,338** | 3,51E-01 | **0** | 0,41999944 | 0,4541054 |
| DPP3_HUMA | DPP3 | sp\|Q9NY33\|DPP3_HUMAN Dipeptidyl peptidase 3 OS=Homo sapiens GN=DPP3 PE=1 SV= | 280,284 | 13,43 | 82589 | 9 | 25,34 | 25,30 | 23,90 | 25,60 | 25,77 | 24,42 | **1,335** | 5,48E-01 | **0** | 0,4166673 | 0,26114596 |
| HS90A_HUM | HSP90AA1 | sp\|P07900\|HS90A_HUMAN Heat shock protein HSP 90-alpha OS=Homo sapiens GN=HSF | 2872,873 | 54,23 | 84660 | 58 | 29,74 | 29,84 | 29,54 | 30,00 | 30,34 | 30,03 | **1,335** | 4,09E-02 | **0** | 0,41666667 | 1,38871821 |
| ASNA_HUM/ | ASNA1 | sp\|O43681\|ASNA_HUMAN ATPase ASNA1 OS=Homo sapiens GN=ASNA1 PE=1 SV=2 | 194,043 | 23,28 | 38793 | 7 | 23,96 | 24,27 | 25,00 | 24,52 | 24,57 | 25,39 | **1,335** | 3,75E-01 | **0** | 0,41666667 | 0,4258612 |
| AAPK2_HUM | PRKAA2 | sp\|P54646\|AAPK2_HUMAN 5'-AMP-activated protein kinase catalytic subunit alpha-2 O! | 102,226 | 8,88 | 62320 | 3 | 17,42 | 17,83 | 17,77 | 17,93 | 18,66 | 17,68 | **1,334** | 2,65E-01 | **0** | 0,41611989 | 0,57729079 |
| PSMD7_HUN | PSMD7 | sp\|P51665\|PSMD7_HUMAN 26S proteasome non-ATPase regulatory subunit 7 OS=Hom | 644,95 | 52,47 | 37025 | 15 | 25,69 | 26,21 | 27,08 | 26,21 | 26,49 | 27,52 | **1,332** | 5,07E-01 | **0** | 0,41333326 | 0,29467762 |
| KCC2D_HUM | CAMK2D | sp\|Q13557\|KCC2D_HUMAN Calcium/calmodulin-dependent protein kinase type II subu | 211,512 | 15,83 | 56369 | 7 | 22,75 | 22,98 | 23,78 | 23,23 | 23,53 | 23,99 | **1,332** | 3,41E-01 | **0** | 0,41333326 | 0,46771646 |
| CD44_HUMA | CD44 | sp\|P16070\|CD44_HUMAN CD44 antigen OS=Homo sapiens GN=CD44 PE=1 SV=3 | 281,495 | 7,95 | 81538 | 6 | 27,91 | 26,75 | 27,47 | 28,30 | 27,69 | 27,38 | **1,332** | 3,94E-01 | **0** | 0,41333326 | 0,40485577 |
| KLC4_HUMA | KLC4 | sp\|Q9NSK0\|KLC4_HUMAN Kinesin light chain 4 OS=Homo sapiens GN=KLC4 PE=1 SV=3 | 74,413 | 7,11 | 68640 | 4 | 16,83 | 17,19 | 19,32 | 17,09 | 18,05 | 19,44 | **1,332** | 7,10E-01 | **0** | 0,41333326 | 0,1488894 |
| CYC_HUMAN | CYCS | sp\|P99999\|CYC_HUMAN Cytochrome c OS=Homo sapiens GN=CYCS PE=1 SV=2 | 65,575 | 19,05 | 11749 | 3 | 25,54 | 25,52 | 21,92 | 25,66 | 25,75 | 22,81 | **1,332** | 8,02E-01 | **0** | 0,41333262 | 0,0958232 |
| GGCT_HUM/ | GGCT | sp\|O75223\|GGCT_HUMAN Gamma-glutamylcyclotransferase OS=Homo sapiens GN=GG | 46,019 | 15,96 | 21008 | 2 | 20,67 | 20,22 | 19,19 | 20,73 | 20,93 | 19,66 | **1,330** | 5,22E-01 | **0** | 0,41189194 | 0,28197376 |
| GPDA_HUM/ | GPD1 | sp\|P21695\|GPDA_HUMAN Glycerol-3-phosphate dehydrogenase [NAD(+)], cytoplasmic | 30,05 | 3,15 | 37568 | 1 | 20,97 | 19,82 | 18,83 | 19,84 | 20,89 | 20,12 | **1,329** | 5,86E-01 | **0** | 0,41068268 | 0,23223344 |
| DHX15_HUM | DHX15 | sp\|O43143\|DHX15_HUMAN Pre-mRNA-splicing factor ATP-dependent RNA helicase DH) | 852,851 | 44,53 | 90933 | 29 | 26,32 | 26,32 | 27,17 | 26,52 | 26,78 | 27,74 | **1,329** | 4,29E-01 | **0** | 0,41000048 | 0,36715557 |
| SAHH_HUM/ | AHCY | sp\|P23526\|SAHH_HUMAN Adenosylhomocysteinase OS=Homo sapiens GN=AHCY PE=1 | 773,306 | 43,29 | 47716 | 21 | 28,06 | 27,80 | 27,92 | 28,31 | 28,26 | 28,44 | **1,329** | 1,13E-02 | **0** | 0,41000048 | 1,94604537 |
| PLEC_HUMA | PLEC | sp\|Q15149\|PLEC_HUMAN Plectin OS=Homo sapiens GN=PLEC PE=1 SV=3 | 1066,603 | 11,46 | 531791 | 46 | 26,51 | 26,73 | 26,50 | 27,04 | 27,03 | 26,90 | **1,329** | 9,43E-03 | **0** | 0,41000048 | 2,02549155 |
| RPR1B_HUM | RPRD1B | sp\|Q9NQG5\|RPR1B_HUMAN Regulation of nuclear pre-mRNA domain-containing protei | 52,685 | 7,98 | 36900 | 2 | 20,64 | 20,41 | 20,08 | 21,10 | 20,34 | 20,92 | **1,329** | 2,18E-01 | **0** | 0,41000048 | 0,66079873 |
| CAMLG_HUN | CAMLG | sp\|P49069\|CAMLG_HUMAN Calcium signal-modulating cyclophilin ligand OS=Homo sapi | 65,778 | 6,76 | 32953 | 1 | 19,66 | 19,69 | 20,83 | 19,85 | 20,28 | 21,28 | **1,329** | 5,13E-01 | **0** | 0,41000048 | 0,28948859 |
| PSMD6_HUN | PSMD6 | sp\|Q15008\|PSMD6_HUMAN 26S proteasome non-ATPase regulatory subunit 6 OS=Hom | 1066,278 | 65,81 | 45531 | 27 | 26,32 | 26,38 | 27,45 | 26,59 | 26,87 | 27,92 | **1,329** | 4,95E-01 | **0** | 0,41000048 | 0,30556281 |
| PIMT_HUMA | PCMT1 | sp\|P22061\|PIMT_HUMAN Protein-L-isoaspartate(D-aspartate) O-methyltransferase OS= | 80,197 | 15,42 | 24636 | 4 | 22,77 | 23,41 | 22,62 | 23,94 | 23,60 | 22,49 | **1,329** | 4,59E-01 | **0** | 0,40999985 | 0,33860121 |
| THIC_HUMAI | ACAT2 | sp\|Q9BWD1\|THIC_HUMAN Acetyl-CoA acetyltransferase, cytosolic OS=Homo sapiens GI | 23,071 | 2,02 | 41351 | 1 | 21,94 | 22,21 | 22,19 | 22,46 | 22,35 | 22,76 | **1,329** | 5,24E-02 | **0** | 0,40999985 | 1,28029748 |
| SURF4_HUM | SURF4 | sp\|O15260\|SURF4_HUMAN Surfeit locus protein 4 OS=Homo sapiens GN=SURF4 PE=1 S | 217,891 | 19,33 | 30394 | 6 | 24,35 | 24,64 | 25,06 | 24,81 | 24,94 | 25,52 | **1,326** | 2,47E-01 | **0** | 0,40666707 | 0,60735634 |
| COPD_HUM/ | ARCN1 | sp\|P48444\|COPD_HUMAN Coatomer subunit delta OS=Homo sapiens GN=ARCN1 PE=1 S | 381,172 | 32,09 | 57210 | 16 | 25,73 | 25,26 | 24,74 | 25,74 | 25,72 | 25,49 | **1,326** | 2,43E-01 | **0** | 0,40666644 | 0,61493534 |
| EH1L1_HUM | EHBP1L1 | sp\|Q8N3D4\|EH1L1_HUMAN EH domain-binding protein 1-like protein 1 OS=Homo sapie | 43,485 | 0,72 | 161854 | 1 | 22,24 | 21,65 | 18,99 | 22,31 | 21,73 | 20,06 | **1,325** | 7,53E-01 | **0** | 0,40584819 | 0,12307064 |
| CC50A_HUM | TMEM30A | sp\|Q9NV96\|CC50A_HUMAN Cell cycle control protein 50A OS=Homo sapiens GN=TMEN | 100,899 | 12,47 | 40684 | 3 | 24,42 | 23,76 | 20,81 | 24,43 | 24,41 | 21,36 | **1,323** | 8,02E-01 | **0** | 0,40333366 | 0,09568753 |
| PUR9_HUM/ | ATIC | sp\|P31939\|PUR9_HUMAN Bifunctional purine biosynthesis protein PURH OS=Homo sap | 701,694 | 34,46 | 64616 | 16 | 25,98 | 26,14 | 26,34 | 26,37 | 26,42 | 26,88 | **1,323** | 1,05E-01 | **0** | 0,40333366 | 0,98024912 |
| SH3L3_HUM | SH3BGRL3 | sp\|Q9H299\|SH3L3_HUMAN SH3 domain-binding glutamic acid-rich-like protein 3 OS=Hc | 101,171 | 66,67 | 10438 | 3 | 23,13 | 23,11 | 22,31 | 23,36 | 23,32 | 23,08 | **1,323** | 2,28E-01 | **0** | 0,40333366 | 0,64131702 |
| GNA1_HUM/ | GNPNAT1 | sp\|Q96EK6\|GNA1_HUMAN Glucosamine 6-phosphate N-acetyltransferase OS=Homo sa | 138,138 | 19,02 | 20749 | 3 | 22,33 | 23,11 | 22,14 | 22,95 | 23,27 | 22,57 | **1,323** | 3,24E-01 | **0** | 0,40333366 | 0,48903874 |
| BACH_HUM/ | ACOT7 | sp\|O00154\|BACH_HUMAN Cytosolic acyl coenzyme A thioester hydrolase OS=Homo sap | 230,332 | 13,95 | 41796 | 6 | 25,44 | 25,55 | 25,19 | 25,76 | 25,98 | 25,65 | **1,323** | 4,88E-02 | **0** | 0,40333303 | 1,31130447 |
| TERA_HUMA | VCP | sp\|P55072\|TERA_HUMAN Transitional endoplasmic reticulum ATPase OS=Homo sapien: | 2817,634 | 78,91 | 89322 | 62 | 29,30 | 29,46 | 28,87 | 29,73 | 29,80 | 29,31 | **1,323** | 1,59E-01 | **0** | 0,40333303 | 0,79874611 |
| NDKA_HUM/ | NME1 | sp\|P15531\|NDKA_HUMAN Nucleoside diphosphate kinase A OS=Homo sapiens GN=NM | 340,286 | 51,97 | 17149 | 7 | 23,04 | 23,77 | 22,52 | 23,58 | 23,65 | 23,31 | **1,323** | 3,45E-01 | **0** | 0,40333239 | 0,46215065 |
| IPO8_HUMAI | IPO8 | sp\|O15397\|IPO8_HUMAN Importin-8 OS=Homo sapiens GN=IPO8 PE=1 SV=2 | 92,866 | 2,41 | 119938 | 2 | 18,68 | 20,74 | 21,91 | 20,19 | 20,32 | 22,03 | **1,322** | 7,36E-01 | **0** | 0,4023304 | 0,13296827 |
| LRC8D_HUM | LRRC8D | sp\|Q7L1W4\|LRC8D_HUMAN Volume-regulated anion channel subunit LRRC8D OS=Hom | 17,52 | 0,93 | 98201 | 1 | 21,72 | 21,53 | 18,43 | 21,42 | 21,77 | 19,70 | **1,321** | 7,63E-01 | **0** | 0,40182304 | 0,11773049 |
| VAPA_HUMA | VAPA | sp\|Q9P0L0\|VAPA_HUMAN Vesicle-associated membrane protein-associated protein A C | 213,152 | 37,75 | 27893 | 8 | 24,23 | 24,56 | 24,74 | 24,75 | 25,01 | 24,97 | **1,320** | 7,80E-02 | **0** | 0,40000025 | 1,10773 |
| FXR1_HUMA | FXR1 | sp\|P51114\|FXR1_HUMAN Fragile X mental retardation syndrome-related protein 1 OS=I | 333,789 | 16,59 | 69721 | 9 | 23,29 | 22,51 | 24,65 | 23,81 | 23,00 | 24,84 | **1,320** | 6,52E-01 | **0** | 0,39999962 | 0,18595141 |
| METK2_HUN | MAT2A | sp\|P31153\|METK2_HUMAN S-adenosylmethionine synthase isoform type-2 OS=Homo s | 342,454 | 33,16 | 43661 | 11 | 24,28 | 25,02 | 25,60 | 25,00 | 25,23 | 25,87 | **1,320** | 4,36E-01 | **0** | 0,39999962 | 0,3608619 |
| MON2_HUM | MON2 | sp\|Q7Z3U7\|MON2_HUMAN Protein MON2 homolog OS=Homo sapiens GN=MON2 PE=1 | 74,127 | 3,38 | 190487 | 4 | 21,45 | 20,76 | 21,52 | 21,71 | 21,33 | 21,89 | **1,320** | 2,44E-01 | **0** | 0,39999898 | 0,61188424 |
| NMT1_HUM | NMT1 | sp\|P30419\|NMT1_HUMAN Glycylpeptide N-tetradecanoyltransferase 1 OS=Homo sapien | 210,511 | 16,53 | 56806 | 5 | 22,38 | 22,30 | 22,89 | 22,10 | 22,91 | 23,75 | **1,316** | 4,81E-01 | **0** | 0,39666748 | 0,31796358 |
| LAR4B_HUM | LARP4B | sp\|Q92615\|LAR4B_HUMAN La-related protein 4B OS=Homo sapiens GN=LARP4B PE=1 S | 68,004 | 3,52 | 80552 | 2 | 20,98 | 20,83 | 23,41 | 21,61 | 21,49 | 23,31 | **1,316** | 7,18E-01 | **0** | 0,39666684 | 0,14406209 |
| SRP72_HUM | SRP72 | sp\|O76094\|SRP72_HUMAN Signal recognition particle subunit SRP72 OS=Homo sapiens | 462,147 | 23,25 | 74606 | 12 | 24,65 | 24,71 | 25,33 | 24,90 | 25,16 | 25,82 | **1,316** | 3,20E-01 | **0** | 0,39666684 | 0,49496354 |
| NDKB_HUM/ | NME2 | sp\|P22392\|NDKB_HUMAN Nucleoside diphosphate kinase B OS=Homo sapiens GN=NMI | 522,454 | 66,45 | 17298 | 12 | 27,67 | 27,73 | 26,18 | 27,96 | 27,88 | 26,93 | **1,316** | 5,48E-01 | **0** | 0,39666621 | 0,26116795 |
| TRPV4_HUM | TRPV4 | sp\|Q9HBA0\|TRPV4_HUMAN Transient receptor potential cation channel subfamily V me | 11,74 | 1,49 | 98281 | 1 | 21,08 | 21,15 | 18,45 | 21,12 | 21,04 | 19,70 | **1,314** | 7,14E-01 | **0** | 0,39451218 | 0,14648886 |
| PRS4_HUMA | PSMC1 | sp\|P62191\|PRS4_HUMAN 26S proteasome regulatory subunit 4 OS=Homo sapiens GN=I | 646,715 | 42,95 | 49185 | 15 | 25,81 | 25,96 | 26,64 | 26,25 | 26,25 | 27,09 | **1,313** | 3,58E-01 | **0** | 0,39333407 | 0,44620864 |
| CSN5_HUMA | COPS5 | sp\|Q92905\|CSN5_HUMAN COP9 signalosome complex subunit 5 OS=Homo sapiens GN= | 126,311 | 17,96 | 37579 | 5 | 23,62 | 23,23 | 23,92 | 23,50 | 24,09 | 24,36 | **1,313** | 2,90E-01 | **0** | 0,39333344 | 0,53709777 |
| SYTC_HUMA | TARS | sp\|P26639\|SYTC_HUMAN Threonine--tRNA ligase, cytoplasmic OS=Homo sapiens GN=T/ | 408,136 | 21,44 | 83435 | 15 | 26,18 | 26,21 | 25,75 | 26,53 | 26,58 | 26,21 | **1,313** | 1,05E-01 | **0** | 0,39333344 | 0,97817074 |
| TBCB_HUMA | TBCB | sp\|Q99426\|TBCB_HUMAN Tubulin-folding cofactor B OS=Homo sapiens GN=TBCB PE=1 | 22,199 | 6,97 | 27326 | 2 | 22,78 | 23,55 | 22,28 | 23,48 | 24,12 | 22,19 | **1,313** | 5,93E-01 | **0** | 0,39333344 | 0,22730896 |
| IF2B_HUMAI | EIF2S2 | sp\|P20042\|IF2B_HUMAN Eukaryotic translation initiation factor 2 subunit 2 OS=Homo sa | 223,082 | 23,12 | 38388 | 9 | 24,85 | 24,83 | 25,32 | 25,16 | 25,25 | 25,77 | **1,313** | 1,89E-01 | **0** | 0,39333344 | 0,72422838 |
| MCM2_HUM | MCM2 | sp\|P49736\|MCM2_HUMAN DNA replication licensing factor MCM2 OS=Homo sapiens G | 690,169 | 27,32 | 101896 | 21 | 24,70 | 24,71 | 26,75 | 25,07 | 25,41 | 26,86 | **1,313** | 6,76E-01 | **0** | 0,39333344 | 0,1698017 |
| ANXA6_HUN | ANXA6 | sp\|P08133\|ANXA6_HUMAN Annexin A6 OS=Homo sapiens GN=ANXA6 PE=1 SV=3 | 1925,292 | 71,17 | 75873 | 52 | 31,06 | 31,03 | 28,88 | 31,48 | 31,53 | 29,14 | **1,313** | 7,32E-01 | **0** | 0,39333344 | 0,13576499 |
| IF4G2_HUM/ | EIF4G2 | sp\|P78344\|IF4G2_HUMAN Eukaryotic translation initiation factor 4 gamma 2 OS=Homo | 312,144 | 16,65 | 102362 | 15 | 25,42 | 25,14 | 25,78 | 26,25 | 25,21 | 26,06 | **1,313** | 3,47E-01 | **0** | 0,3933328 | 0,45952409 |
| TCPA_HUMA | TCP1 | sp\|P17987\|TCPA_HUMAN T-complex protein 1 subunit alpha OS=Homo sapiens GN=TCI | 1252,451 | 52,16 | 60344 | 27 | 27,89 | 28,00 | 28,11 | 28,21 | 28,41 | 28,56 | **1,313** | 3,03E-02 | **0** | 0,3933328 | 1,51892557 |
| CRNN_HUM/ | CRNN | sp\|Q9UBG3\|CRNN_HUMAN Cornulin OS=Homo sapiens GN=CRNN PE=1 SV=1 | 53,469 | 6,67 | 53533 | 1 | 19,23 | 18,43 | 17,79 | 18,94 | 17,42 | 20,26 | **1,311** | 6,93E-01 | **0** | 0,3909537 | 0,15944457 |
| SYDC_HUMA | DARS | sp\|P14868\|SYDC_HUMAN Aspartate--tRNA ligase, cytoplasmic OS=Homo sapiens GN=D/ | 630,353 | 42,32 | 57136 | 19 | 25,48 | 25,73 | 26,96 | 25,69 | 26,03 | 27,62 | **1,310** | 6,31E-01 | **0** | 0,3900013 | 0,20019892 |
| MTAP_HUM/ | MTAP | sp\|Q13126\|MTAP_HUMAN S-methyl-5'-thioadenosine phosphorylase OS=Homo sapien | 197,751 | 32,16 | 31236 | 6 | 24,48 | 25,00 | 23,73 | 24,67 | 25,17 | 24,54 | **1,310** | 4,01E-01 | **0** | 0,39000066 | 0,39663225 |
| UBE4A_HUM | UBE4A | sp\|Q14139\|UBE4A_HUMAN Ubiquitin conjugation factor E4 A OS=Homo sapiens GN=UE | 33,973 | 2,35 | 122561 | 2 | 20,69 | 21,13 | 20,64 | 21,03 | 21,47 | 21,13 | **1,310** | 1,30E-01 | **0** | 0,39000003 | 0,88714483 |
| RBBP4_HUM | RBBP4 | sp\|Q09028\|RBBP4_HUMAN Histone-binding protein RBBP4 OS=Homo sapiens GN=RBBF | 348,445 | 49,88 | 47656 | 12 | 23,82 | 24,72 | 25,10 | 24,49 | 24,84 | 25,48 | **1,310** | 4,60E-01 | **0** | 0,39000003 | 0,33727565 |
| CND2_HUM/ | NCAPH | sp\|Q15003\|CND2_HUMAN Condensin complex subunit 2 OS=Homo sapiens GN=NCAPH | 22,1 | 2,16 | 82563 | 1 | 17,80 | 18,40 | 17,92 | 18,01 | 18,39 | 18,89 | **1,310** | 2,82E-01 | **0** | 0,39000003 | 0,54984548 |
| PRS10_HUM | PSMC6 | sp\|P62333\|PRS10_HUMAN 26S proteasome regulatory subunit 10B OS=Homo sapiens G | 729,916 | 59,38 | 44173 | 20 | 26,09 | 26,46 | 27,50 | 26,59 | 26,83 | 27,80 | **1,310** | 5,25E-01 | **0** | 0,39000003 | 0,27954708 |
| XPO1_HUM/ | XPO1 | sp\|O14980\|XPO1_HUMAN Exportin-1 OS=Homo sapiens GN=XPO1 PE=1 SV=1 | 833,486 | 27,54 | 123386 | 23 | 25,79 | 25,87 | 25,75 | 25,85 | 26,39 | 26,34 | **1,310** | 9,08E-02 | **0** | 0,38999939 | 1,04176686 |
| TKT_HUMAN | TKT | sp\|P29401\|TKT_HUMAN Transketolase OS=Homo sapiens GN=TKT PE=1 SV=3 | 350,898 | 20,55 | 67878 | 11 | 26,86 | 26,61 | 26,27 | 27,15 | 27,01 | 26,75 | **1,310** | 1,33E-01 | **0** | 0,38999939 | 0,87597055 |
| MRCKB_HUN | CDC42BPB | sp\|Q9Y5S2\|MRCKB_HUMAN Serine/threonine-protein kinase MRCK beta OS=Homo sap | 209,598 | 7,66 | 194315 | 10 | 24,13 | 24,00 | 23,25 | 24,10 | 23,70 | 24,74 | **1,307** | 3,98E-01 | **0** | 0,38666725 | 0,40060134 |
| ERGI3_HUM/ | ERGIC3 | sp\|Q9Y282\|ERGI3_HUMAN Endoplasmic reticulum-Golgi intermediate compartment pro | 60,881 | 12,53 | 43222 | 3 | 22,33 | 21,09 | 23,13 | 22,35 | 22,17 | 23,19 | **1,307** | 5,96E-01 | **0** | 0,38666725 | 0,22504236 |
| DHX9_HUM/ | DHX9 | sp\|Q08211\|DHX9_HUMAN ATP-dependent RNA helicase A OS=Homo sapiens GN=DHX9 | 425,357 | 11,18 | 140958 | 13 | 23,21 | 23,35 | 25,17 | 23,56 | 23,73 | 25,60 | **1,307** | 6,92E-01 | **0** | 0,38666662 | 0,15968432 |
| VKOR1_HUN | VKORC1 | sp\|Q9BQB6\|VKOR1_HUMAN Vitamin K epoxide reductase complex subunit 1 OS=Homo | 54,305 | 7,98 | 18235 | 1 | 21,69 | 21,96 | 20,94 | 21,60 | 21,97 | 22,18 | **1,307** | 3,30E-01 | **0** | 0,38666662 | 0,48135285 |
| RL6_HUMAN | RPL6 | sp\|Q02878\|RL6_HUMAN 60S ribosomal protein L6 OS=Homo sapiens GN=RPL6 PE=1 SV= | 556,951 | 32,29 | 32728 | 15 | 27,25 | 27,45 | 27,18 | 27,62 | 27,23 | 28,19 | **1,307** | 2,54E-01 | **0** | 0,38666662 | 0,59579782 |
| DCTN5_HUN | DCTN5 | sp\|Q9BTE1\|DCTN5_HUMAN Dynactin subunit 5 OS=Homo sapiens GN=DCTN5 PE=1 SV= | 78,738 | 23,08 | 20127 | 4 | 22,48 | 22,91 | 20,09 | 22,21 | 22,72 | 21,71 | **1,307** | 6,97E-01 | **0** | 0,38666598 | 0,15664151 |
| IF4G1_HUM/ | EIF4G1 | sp\|Q04637\|IF4G1_HUMAN Eukaryotic translation initiation factor 4 gamma 1 OS=Homo | 473,346 | 12,01 | 175491 | 17 | 25,03 | 24,86 | 25,39 | 25,22 | 25,24 | 25,98 | **1,307** | 2,60E-01 | **0** | 0,38666598 | 0,58510976 |
| SPTN1_HUM | SPTAN1 | sp\|Q13813\|SPTN1_HUMAN Spectrin alpha chain, non-erythrocytic 1 OS=Homo sapiens | 1511,053 | 26,42 | 284539 | 49 | 26,98 | 27,24 | 25,24 | 27,16 | 27,50 | 25,95 | **1,304** | 6,51E-01 | **0** | 0,38333384 | 0,18663786 |
| CNN2_HUM/ | CNN2 | sp\|Q99439\|CNN2_HUMAN Calponin-2 OS=Homo sapiens GN=CNN2 PE=1 SV=4 | 56,082 | 10,36 | 33697 | 3 | 20,24 | 20,25 | 21,08 | 21,04 | 20,33 | 21,35 | **1,304** | 4,03E-01 | **0** | 0,38333384 | 0,3942609 |
| LRRK1_HUM | LRRK1 | sp\|Q38SD2\|LRRK1_HUMAN Leucine-rich repeat serine/threonine-protein kinase 1 OS=H | 14,116 | 0,35 | 225393 | 1 | 23,57 | 23,75 | 23,17 | 23,71 | 24,18 | 23,75 | **1,304** | 1,68E-01 | **0** | 0,38333321 | 0,77446372 |
| RL3_HUMAN | RPL3 | sp\|P39023\|RL3_HUMAN 60S ribosomal protein L3 OS=Homo sapiens GN=RPL3 PE=1 SV= | 801,332 | 43,92 | 46109 | 23 | 27,50 | 27,53 | 28,03 | 27,75 | 27,62 | 28,84 | **1,304** | 4,16E-01 | **0** | 0,38333321 | 0,38052507 |
| CSN4_HUMA | COPS4 | sp\|Q9BT78\|CSN4_HUMAN COP9 signalosome complex subunit 4 OS=Homo sapiens GN= | 491,82 | 43,1 | 46269 | 16 | 25,13 | 25,16 | 25,32 | 25,63 | 25,59 | 25,53 | **1,301** | 4,45E-03 | **0** | 0,38000043 | 2,35162208 |
| RNBP6_HUM | RANBP6 | sp\|O60518\|RNBP6_HUMAN Ran-binding protein 6 OS=Homo sapiens GN=RANBP6 PE=1 | 66,67 | 3,62 | 124714 | 3 | 21,32 | 21,50 | 22,73 | 21,49 | 22,03 | 23,17 | **1,301** | 5,98E-01 | **0** | 0,38000043 | 0,22330195 |
| BAG6_HUM/ | BAG6 | sp\|P46379\|BAG6_HUMAN Large proline-rich protein BAG6 OS=Homo sapiens GN=BAG6 | 296,98 | 13,87 | 119409 | 10 | 24,03 | 24,05 | 25,56 | 24,29 | 24,64 | 25,85 | **1,301** | 6,13E-01 | **0** | 0,38000043 | 0,21284662 |
| LTN1_HUMA | LTN1 | sp\|O94822\|LTN1_HUMAN E3 ubiquitin-protein ligase listerin OS=Homo sapiens GN=LTN | 11,291 | 0,91 | 200552 | 1 | 16,53 | 17,83 | 18,88 | 17,16 | 18,18 | 19,04 | **1,301** | 6,85E-01 | **0** | 0,38000043 | 0,16439264 |
| PKP4_HUMA | PKP4 | sp\|Q99569\|PKP4_HUMAN Plakophilin-4 OS=Homo sapiens GN=PKP4 PE=1 SV=2 | 48,474 | 1,51 | 131868 | 2 | 21,89 | 22,09 | 19,60 | 21,72 | 21,98 | 21,02 | **1,301** | 6,77E-01 | **0** | 0,3799998 | 0,16909335 |
| OSB10_HUM | OSBPL10 | sp\|Q9BXB5\|OSB10_HUMAN Oxysterol-binding protein-related protein 10 OS=Homo sap | 11,722 | 1,31 | 83970 | 1 | 21,00 | 20,66 | 21,58 | 21,22 | 21,48 | 21,68 | **1,301** | 2,74E-01 | **0** | 0,3799998 | 0,56271438 |
| EXOS1_HUM | EXOSC1 | sp\|Q9Y3B2\|EXOS1_HUMAN Exosome complex component CSL4 OS=Homo sapiens GN= | 49,398 | 13,85 | 21452 | 2 | 20,79 | 20,40 | 21,84 | 21,39 | 21,10 | 21,68 | **1,301** | 4,56E-01 | **0** | 0,3799998 | 0,34056782 |
| GLOD4_HUN | GLOD4 | sp\|Q9HC38\|GLOD4_HUMAN Glyoxalase domain-containing protein 4 OS=Homo sapiens | 71,897 | 13,1 | 34793 | 3 | 22,92 | 23,32 | 21,50 | 23,55 | 23,80 | 21,53 | **1,301** | 6,97E-01 | **0** | 0,3799998 | 0,15705258 |
| MTMR9_HUI | MTMR9 | sp\|Q96QG7\|MTMR9_HUMAN Myotubularin-related protein 9 OS=Homo sapiens GN=M | 31,86 | 1,46 | 63462 | 1 | 18,97 | 19,47 | 18,37 | 19,14 | 19,14 | 19,67 | **1,301** | 3,55E-01 | **0** | 0,3799998 | 0,44957553 |
| TFP11_HUM | TFIP11 | sp\|Q9UBB9\|TFP11_HUMAN Tuftelin-interacting protein 11 OS=Homo sapiens GN=TFIP1 | 28,466 | 1,31 | 96820 | 1 | 18,63 | 16,20 | 18,87 | 18,23 | 17,31 | 19,30 | **1,301** | 7,31E-01 | **0** | 0,37945938 | 0,13609547 |
| HS2ST_HUM | HS2ST1 | sp\|Q7LGA3\|HS2ST_HUMAN Heparan sulfate 2-O-sulfotransferase 1 OS=Homo sapiens G | 55,7 | 4,78 | 41881 | 1 | 17,78 | 18,77 | 20,54 | 18,53 | 19,54 | 20,16 | **1,300** | 7,07E-01 | **0** | 0,37851079 | 0,15068971 |
| IFT88_HUM/ | IFT88 | sp\|Q13099\|IFT88_HUMAN Intraflagellar transport protein 88 homolog OS=Homo sapien | 45,45 | 3 | 94270 | 2 | 18,48 | 18,01 | 24,28 | 21,16 | 18,20 | 22,54 | **1,299** | 8,82E-01 | **0** | 0,37790298 | 0,05457417 |
| DC1L1_HUM | DYNC1LI1 | sp\|Q9Y6G9\|DC1L1_HUMAN Cytoplasmic dynein 1 light intermediate chain 1 OS=Homo s | 195,539 | 23,9 | 56579 | 9 | 24,74 | 25,15 | 25,22 | 25,23 | 25,31 | 25,70 | **1,298** | 1,45E-01 | **0** | 0,37666702 | 0,83807557 |
| GLO2_HUM/ | HAGH | sp\|Q16775\|GLO2_HUMAN Hydroxyacylglutathione hydrolase, mitochondrial OS=Homo | 94,328 | 18,18 | 33806 | 4 | 23,43 | 23,72 | 20,17 | 23,72 | 23,29 | 21,44 | **1,298** | 7,92E-01 | **0** | 0,37666702 | 0,10130096 |
| ASNS_HUMA | ASNS | sp\|P08243\|ASNS_HUMAN Asparagine synthetase [glutamine-hydrolyzing] OS=Homo saj | 167,324 | 14,62 | 64370 | 7 | 24,50 | 24,70 | 24,12 | 24,94 | 25,11 | 24,40 | **1,298** | 2,40E-01 | **0** | 0,37666639 | 0,61916952 |
| TSN_HUMAN | TSN | sp\|Q15631\|TSN_HUMAN Translin OS=Homo sapiens GN=TSN PE=1 SV=1 | 345,745 | 47,81 | 26183 | 8 | 24,41 | 24,45 | 24,76 | 24,41 | 24,96 | 25,38 | **1,298** | 2,80E-01 | **0** | 0,37666575 | 0,55262538 |
| TES_HUMAN | TES | sp\|Q9UGI8\|TES_HUMAN Testin OS=Homo sapiens GN=TES PE=1 SV=1 | 80,564 | 10,21 | 47996 | 3 | 22,94 | 21,66 | 23,21 | 23,15 | 22,76 | 23,02 | **1,295** | 4,90E-01 | **0** | 0,37333361 | 0,30988645 |
| STRAP_HUM | STRAP | sp\|Q9Y3F4\|STRAP_HUMAN Serine-threonine kinase receptor-associated protein OS=Ho | 741,867 | 57,14 | 38438 | 16 | 26,82 | 26,80 | 26,52 | 26,86 | 27,14 | 27,26 | **1,295** | 7,13E-02 | **0** | 0,37333361 | 1,14719032 |
| NUP93_HUN | NUP93 | sp\|Q8N1F7\|NUP93_HUMAN Nuclear pore complex protein Nup93 OS=Homo sapiens GI | 211,972 | 11,36 | 93488 | 7 | 22,49 | 23,04 | 24,00 | 22,68 | 23,37 | 24,60 | **1,295** | 6,29E-01 | **0** | 0,37333361 | 0,20152543 |
| RS21_HUMA | RPS21 | sp\|P63220\|RS21_HUMAN 40S ribosomal protein S21 OS=Homo sapiens GN=RPS21 PE=1 | 331,128 | 63,86 | 9111 | 7 | 25,70 | 25,87 | 22,40 | 25,74 | 25,79 | 23,56 | **1,295** | 7,95E-01 | **0** | 0,37333298 | 0,09937211 |
| PSD12_HUM | PSMD12 | sp\|O00232\|PSD12_HUMAN 26S proteasome non-ATPase regulatory subunit 12 OS=Hon | 948,661 | 49,12 | 52904 | 23 | 26,30 | 26,19 | 27,45 | 26,42 | 26,75 | 27,89 | **1,295** | 5,68E-01 | **0** | 0,37333298 | 0,24577436 |
| SRSF1_HUM/ | SRSF1 | sp\|Q07955\|SRSF1_HUMAN Serine/arginine-rich splicing factor 1 OS=Homo sapiens GN= | 239,985 | 31,45 | 27745 | 7 | 24,42 | 24,47 | 25,17 | 24,77 | 24,70 | 25,70 | **1,292** | 4,11E-01 | **0** | 0,37000084 | 0,38656583 |
| H32_HUMAN | HIST2H3A | sp\|Q71DI3\|H32_HUMAN Histone H3.2 OS=Homo sapiens GN=HIST2H3A PE=1 SV=3 | 248,29 | 43,38 | 15388 | 6 | 21,46 | 21,93 | 23,92 | 22,09 | 22,15 | 24,18 | **1,292** | 7,35E-01 | **0** | 0,3700002 | 0,13363178 |
| P3H1_HUMA | P3H1 | sp\|Q32P28\|P3H1_HUMAN Prolyl 3-hydroxylase 1 OS=Homo sapiens GN=P3H1 PE=1 SV= | 80,09 | 4,76 | 83394 | 3 | 21,33 | 21,09 | 22,04 | 22,01 | 21,73 | 21,83 | **1,292** | 2,80E-01 | **0** | 0,36999957 | 0,55211404 |
| ZC3H6_HUM | ZC3H6 | sp\|P61129\|ZC3H6_HUMAN Zinc finger CCCH domain-containing protein 6 OS=Homo sap | 20,47 | 0,59 | 131670 | 1 | 17,38 | 18,82 | 22,86 | 18,53 | 18,65 | 22,98 | **1,291** | 8,75E-01 | **0** | 0,36828105 | 0,05795083 |
| RL32_HUMA | RPL32 | sp\|P62910\|RL32_HUMAN 60S ribosomal protein L32 OS=Homo sapiens GN=RPL32 PE=1 | 156,887 | 34,81 | 15860 | 5 | 25,26 | 25,39 | 23,30 | 25,51 | 25,11 | 24,43 | **1,289** | 6,49E-01 | **0** | 0,36666743 | 0,18789933 |
| ELOB_HUMA | ELOB | sp\|Q15370\|ELOB_HUMAN Elongin-B OS=Homo sapiens GN=ELOB PE=1 SV=1 | 194,259 | 44,92 | 13133 | 6 | 24,50 | 24,38 | 23,18 | 24,51 | 24,78 | 23,87 | **1,289** | 5,04E-01 | **0** | 0,36666743 | 0,29727175 |
| RL15_HUMA | RPL15 | sp\|P61313\|RL15_HUMAN 60S ribosomal protein L15 OS=Homo sapiens GN=RPL15 PE=1 | 383,144 | 50 | 24146 | 13 | 26,86 | 26,88 | 26,67 | 27,18 | 26,70 | 27,63 | **1,289** | 2,56E-01 | **0** | 0,36666679 | 0,59211983 |
| EIF3G_HUM/ | EIF3G | sp\|O75821\|EIF3G_HUMAN Eukaryotic translation initiation factor 3 subunit G OS=Homo | 146,783 | 19,38 | 35611 | 5 | 22,01 | 22,95 | 25,71 | 23,20 | 23,65 | 24,92 | **1,289** | 7,79E-01 | **0** | 0,36666679 | 0,10822741 |
| RL7_HUMAN | RPL7 | sp\|P18124\|RL7_HUMAN 60S ribosomal protein L7 OS=Homo sapiens GN=RPL7 PE=1 SV= | 305,727 | 39,52 | 29226 | 13 | 26,97 | 27,65 | 25,87 | 27,00 | 27,31 | 27,28 | **1,289** | 5,26E-01 | **0** | 0,36666679 | 0,27938345 |
| H90B2_HUM | HSP90AB2P | sp\|Q58FF8\|H90B2_HUMAN Putative heat shock protein HSP 90-beta 2 OS=Homo sapien | 615,222 | 25,2 | 44314 | 13 | 23,82 | 24,27 | 24,27 | 24,05 | 24,37 | 25,04 | **1,289** | 3,26E-01 | **0** | 0,36666679 | 0,48650457 |
| DOCK7_HUN | DOCK7 | sp\|Q96N67\|DOCK7_HUMAN Dedicator of cytokinesis protein 7 OS=Homo sapiens GN=D | 94,913 | 2,01 | 242561 | 3 | 20,78 | 21,01 | 19,25 | 20,91 | 21,16 | 20,07 | **1,289** | 5,99E-01 | **0** | 0,36666616 | 0,22246681 |
| LSM3_HUM/ | LSM3 | sp\|P62310\|LSM3_HUMAN U6 snRNA-associated Sm-like protein LSm3 OS=Homo sapien | 14,422 | 7,84 | 11845 | 1 | 20,72 | 21,25 | 19,15 | 21,02 | 21,03 | 20,17 | **1,289** | 6,25E-01 | **0** | 0,3656985 | 0,20424763 |
| YLPM1_HUN | YLPM1 | sp\|P49750\|YLPM1_HUMAN YLP motif-containing protein 1 OS=Homo sapiens GN=YLPM | 42,725 | 0,92 | 219985 | 1 | 18,50 | 17,12 | 19,70 | 17,31 | 19,01 | 20,09 | **1,288** | 7,57E-01 | **0** | 0,3650322 | 0,12106964 |
| TCPE_HUMA | CCT5 | sp\|P48643\|TCPE_HUMAN T-complex protein 1 subunit epsilon OS=Homo sapiens GN=C | 1312,57 | 63,03 | 59671 | 29 | 28,00 | 28,06 | 27,53 | 28,33 | 28,45 | 27,90 | **1,286** | 1,99E-01 | **0** | 0,36333338 | 0,70038892 |
| VATE1_HUM | ATP6V1E1 | sp\|P36543\|VATE1_HUMAN V-type proton ATPase subunit E 1 OS=Homo sapiens GN=AT | 97,309 | 18,58 | 26145 | 3 | 22,23 | 22,44 | 20,94 | 22,51 | 22,36 | 21,83 | **1,286** | 5,17E-01 | **0** | 0,36333338 | 0,28622927 |
| AT2A2_HUM | ATP2A2 | sp\|P16615\|AT2A2_HUMAN Sarcoplasmic/endoplasmic reticulum calcium ATPase 2 OS=I | 1336,222 | 32,25 | 114757 | 29 | 26,92 | 26,78 | 27,56 | 27,22 | 27,34 | 27,79 | **1,286** | 2,87E-01 | **0** | 0,36333338 | 0,54174859 |
| TMM33_HUI | TMEM33 | sp\|P57088\|TMM33_HUMAN Transmembrane protein 33 OS=Homo sapiens GN=TMEM3 | 91,104 | 12,55 | 27978 | 3 | 24,65 | 24,85 | 23,97 | 24,82 | 25,08 | 24,66 | **1,286** | 2,83E-01 | **0** | 0,36333338 | 0,5483958 |
| THG1_HUM/ | THG1L | sp\|Q9NWX6\|THG1_HUMAN Probable tRNA(His) guanylyltransferase OS=Homo sapiens ( | 19,803 | 3,69 | 34831 | 1 | 21,10 | 21,25 | 21,51 | 21,50 | 21,72 | 21,73 | **1,286** | 6,19E-02 | **0** | 0,36333275 | 1,20802384 |
| K1C17_HUM | KRT17 | sp\|Q04695\|K1C17_HUMAN Keratin, type I cytoskeletal 17 OS=Homo sapiens GN=KRT17 | 421,999 | 25,46 | 48106 | 15 | 24,79 | 24,65 | 23,70 | 25,38 | 24,73 | 24,12 | **1,286** | 5,07E-01 | **0** | 0,36333275 | 0,29470354 |
| UBA3_HUM/ | UBA3 | sp\|Q8TBC4\|UBA3_HUMAN NEDD8-activating enzyme E1 catalytic subunit OS=Homo sap | 94,763 | 15,12 | 51852 | 4 | 23,02 | 23,03 | 22,61 | 23,13 | 23,43 | 23,19 | **1,286** | 9,38E-02 | **0** | 0,36333275 | 1,02791366 |
| SIAS_HUMAI | NANS | sp\|Q9NR45\|SIAS_HUMAN Sialic acid synthase OS=Homo sapiens GN=NANS PE=1 SV=2 | 191,792 | 20,06 | 40308 | 4 | 23,54 | 23,69 | 24,03 | 23,88 | 24,01 | 24,46 | **1,286** | 1,86E-01 | **0** | 0,36333211 | 0,73063594 |
| JUPI1_HUM/ | JPT1 | sp\|Q9UK76\|JUPI1_HUMAN Jupiter microtubule associated homolog 1 OS=Homo sapien: | 51,247 | 15,58 | 16015 | 1 | 21,11 | 21,49 | 17,87 | 21,56 | 21,82 | 18,17 | **1,285** | 8,37E-01 | **0** | 0,36121305 | 0,07731312 |
| HAT1_HUM/ | HAT1 | sp\|O14929\|HAT1_HUMAN Histone acetyltransferase type B catalytic subunit OS=Homo | 205,45 | 15,27 | 49513 | 5 | 22,21 | 22,71 | 23,27 | 22,78 | 23,04 | 23,45 | **1,283** | 3,77E-01 | **0** | 0,36000125 | 0,42313032 |
| PSA1_HUMA | PSMA1 | sp\|P25786\|PSA1_HUMAN Proteasome subunit alpha type-1 OS=Homo sapiens GN=PSM | 465,13 | 75,67 | 29556 | 19 | 26,91 | 27,16 | 26,23 | 27,37 | 27,26 | 26,75 | **1,283** | 3,46E-01 | **0** | 0,36000061 | 0,46115589 |
| PSMD2_HUN | PSMD2 | sp\|Q13200\|PSMD2_HUMAN 26S proteasome non-ATPase regulatory subunit 2 OS=Hom | 1699,414 | 59,69 | 100200 | 42 | 26,99 | 27,30 | 28,06 | 27,39 | 27,59 | 28,45 | **1,283** | 4,73E-01 | **0** | 0,36000061 | 0,32528965 |
| UBA1_HUM/ | UBA1 | sp\|P22314\|UBA1_HUMAN Ubiquitin-like modifier-activating enzyme 1 OS=Homo sapien | 1058,536 | 31,38 | 117849 | 26 | 26,94 | 27,01 | 27,02 | 27,11 | 27,44 | 27,50 | **1,283** | 4,38E-02 | **0** | 0,35999997 | 1,3585466 |
| UCHL3_HUM | UCHL3 | sp\|P15374\|UCHL3_HUMAN Ubiquitin carboxyl-terminal hydrolase isozyme L3 OS=Homc | 238,991 | 36,96 | 26183 | 6 | 26,09 | 24,88 | 23,91 | 24,76 | 26,73 | 24,47 | **1,283** | 7,24E-01 | **0** | 0,35999997 | 0,14034342 |
| STA5A_HUM | STAT5A | sp\|P42229\|STA5A_HUMAN Signal transducer and activator of transcription 5A OS=Hom | 22,203 | 1,51 | 90647 | 1 | 19,29 | 19,26 | 20,22 | 19,61 | 19,73 | 20,51 | **1,282** | 4,45E-01 | **0** | 0,35871951 | 0,35193987 |
| VINEX_HUM | SORBS3 | sp\|O60504\|VINEX_HUMAN Vinexin OS=Homo sapiens GN=SORBS3 PE=1 SV=2 | 60,09 | 3,13 | 75341 | 2 | 21,92 | 21,57 | 19,21 | 22,00 | 21,42 | 20,36 | **1,282** | 7,32E-01 | **0** | 0,35854594 | 0,13553381 |
| TCPQ_HUM/ | CCT8 | sp\|P50990\|TCPQ_HUMAN T-complex protein 1 subunit theta OS=Homo sapiens GN=CC | 1225,1 | 51,64 | 59621 | 29 | 28,39 | 28,38 | 28,18 | 28,66 | 28,76 | 28,60 | **1,280** | 1,26E-02 | **0** | 0,3566672 | 1,9006971 |
| KCTD5_HUM | KCTD5 | sp\|Q9NXV2\|KCTD5_HUMAN BTB/POZ domain-containing protein KCTD5 OS=Homo sapi | 72,519 | 8,97 | 26093 | 1 | 17,99 | 19,41 | 20,06 | 18,95 | 18,92 | 20,66 | **1,280** | 6,93E-01 | **0** | 0,3566672 | 0,15945866 |
| PPM1G_HUN | PPM1G | sp\|O15355\|PPM1G_HUMAN Protein phosphatase 1G OS=Homo sapiens GN=PPM1G PE | 75,754 | 7,69 | 59272 | 4 | 23,06 | 23,56 | 22,53 | 23,59 | 23,40 | 23,23 | **1,280** | 3,21E-01 | **0** | 0,35666656 | 0,49371905 |
| AP3S1_HUM | AP3S1 | sp\|Q92572\|AP3S1_HUMAN AP-3 complex subunit sigma-1 OS=Homo sapiens GN=AP3S1 | 46,45 | 13,47 | 21732 | 2 | 19,83 | 20,53 | 22,28 | 21,20 | 20,87 | 21,64 | **1,280** | 6,64E-01 | **0** | 0,35666656 | 0,17777615 |
| IF5_HUMAN | EIF5 | sp\|P55010\|IF5_HUMAN Eukaryotic translation initiation factor 5 OS=Homo sapiens GN= | 282,582 | 22,74 | 49223 | 8 | 25,11 | 25,08 | 25,27 | 25,28 | 25,45 | 25,80 | **1,280** | 9,54E-02 | **0** | 0,35666656 | 1,02062495 |
| ACACA_HUN | ACACA | sp\|Q13085\|ACACA_HUMAN Acetyl-CoA carboxylase 1 OS=Homo sapiens GN=ACACA PE | 298,413 | 5,75 | 265554 | 11 | 22,00 | 22,74 | 23,78 | 21,69 | 23,51 | 24,39 | **1,280** | 7,26E-01 | **0** | 0,35666656 | 0,13916382 |
| TCPD_HUMA | CCT4 | sp\|P50991\|TCPD_HUMAN T-complex protein 1 subunit delta OS=Homo sapiens GN=CCT | 1662,535 | 68,09 | 57924 | 34 | 28,28 | 28,39 | 28,17 | 28,61 | 28,70 | 28,59 | **1,278** | 7,98E-03 | **0** | 0,35333379 | 2,09776145 |
| PRS7_HUMA | PSMC2 | sp\|P35998\|PRS7_HUMAN 26S proteasome regulatory subunit 7 OS=Homo sapiens GN=I | 956,554 | 71,82 | 48634 | 27 | 26,51 | 26,38 | 27,47 | 26,93 | 26,68 | 27,81 | **1,278** | 5,07E-01 | **0** | 0,35333379 | 0,29503183 |
| COPG1_HUN | COPG1 | sp\|Q9Y678\|COPG1_HUMAN Coatomer subunit gamma-1 OS=Homo sapiens GN=COPG1 | 734,49 | 40,27 | 97718 | 24 | 25,46 | 25,98 | 26,52 | 25,77 | 26,23 | 27,02 | **1,278** | 4,99E-01 | **0** | 0,35333379 | 0,3015528 |
| RDH11_HUM | RDH11 | sp\|Q8TC12\|RDH11_HUMAN Retinol dehydrogenase 11 OS=Homo sapiens GN=RDH11 PE | 198,877 | 13,21 | 35386 | 3 | 24,30 | 24,03 | 24,09 | 24,29 | 24,68 | 24,51 | **1,278** | 6,44E-02 | **0** | 0,35333379 | 1,19113207 |
| CYB5B_HUM | CYB5B | sp\|O43169\|CYB5B_HUMAN Cytochrome b5 type B OS=Homo sapiens GN=CYB5B PE=1 S | 50,084 | 23,29 | 16332 | 1 | 22,89 | 24,08 | 24,34 | 23,51 | 24,42 | 24,44 | **1,278** | 5,50E-01 | **0** | 0,35333379 | 0,25985063 |
| RL24_HUMA | RPL24 | sp\|P83731\|RL24_HUMAN 60S ribosomal protein L24 OS=Homo sapiens GN=RPL24 PE=1 | 292,207 | 31,85 | 17779 | 7 | 26,37 | 26,53 | 26,22 | 26,70 | 26,67 | 26,81 | **1,278** | 2,35E-02 | **0** | 0,35333316 | 1,62935111 |
| 2AAB_HUMA | PPP2R1B | sp\|P30154\|2AAB_HUMAN Serine/threonine-protein phosphatase 2A 65 kDa regulatory | 644,733 | 29,12 | 66214 | 17 | 24,24 | 24,01 | 24,10 | 24,21 | 24,35 | 24,85 | **1,278** | 1,61E-01 | **0** | 0,35333316 | 0,79427298 |
| TBB2A_HUM | TUBB2A | sp\|Q13885\|TBB2A_HUMAN Tubulin beta-2A chain OS=Homo sapiens GN=TUBB2A PE=1 | 2503,288 | 75,06 | 49907 | 53 | 22,96 | 23,19 | 24,44 | 23,39 | 23,64 | 24,62 | **1,278** | 5,84E-01 | **0** | 0,35333316 | 0,23381026 |
| TNR12_HUM | TNFRSF12A | sp\|Q9NP84\|TNR12_HUMAN Tumor necrosis factor receptor superfamily member 12A C | 33,137 | 7,75 | 13911 | 1 | 22,67 | 22,94 | 19,88 | 22,42 | 22,49 | 21,64 | **1,276** | 7,46E-01 | **0** | 0,35181046 | 0,12721362 |
| LDHA_HUM/ | LDHA | sp\|P00338\|LDHA_HUMAN L-lactate dehydrogenase A chain OS=Homo sapiens GN=LDH/ | 466,691 | 32,23 | 36689 | 10 | 27,41 | 27,31 | 26,30 | 27,45 | 27,57 | 27,05 | **1,275** | 4,18E-01 | **0** | 0,35000038 | 0,37901958 |
| CSN1_HUMA | GPS1 | sp\|Q13098\|CSN1_HUMAN COP9 signalosome complex subunit 1 OS=Homo sapiens GN= | 117,591 | 13,44 | 55537 | 6 | 23,84 | 23,62 | 23,38 | 23,57 | 23,95 | 24,37 | **1,275** | 2,59E-01 | **0** | 0,35000038 | 0,58612467 |
| NID2_HUMA | NID2 | sp\|Q14112\|NID2_HUMAN Nidogen-2 OS=Homo sapiens GN=NID2 PE=1 SV=3 | 46,591 | 1,6 | 151254 | 2 | 23,95 | 21,10 | 22,73 | 24,08 | 22,18 | 22,57 | **1,275** | 7,46E-01 | **0** | 0,34999975 | 0,12722528 |
| SF3B4_HUM | SF3B4 | sp\|Q15427\|SF3B4_HUMAN Splicing factor 3B subunit 4 OS=Homo sapiens GN=SF3B4 PE | 201,919 | 12,74 | 44386 | 3 | 21,18 | 22,43 | 23,39 | 22,15 | 22,49 | 23,41 | **1,275** | 6,62E-01 | **0** | 0,34999975 | 0,17924657 |
| CALX_HUMA | CANX | sp\|P27824\|CALX_HUMAN Calnexin OS=Homo sapiens GN=CANX PE=1 SV=2 | 772,391 | 29,56 | 67568 | 16 | 26,92 | 26,88 | 27,94 | 27,43 | 27,55 | 27,81 | **1,275** | 3,91E-01 | **0** | 0,34999975 | 0,40741472 |
| PSA6_HUMA | PSMA6 | sp\|P60900\|PSA6_HUMAN Proteasome subunit alpha type-6 OS=Homo sapiens GN=PSM | 457,483 | 56,5 | 27399 | 14 | 26,29 | 26,62 | 26,26 | 26,68 | 26,91 | 26,63 | **1,275** | 7,19E-02 | **0** | 0,34999911 | 1,14311214 |
| TCPZ_HUMA | CCT6A | sp\|P40227\|TCPZ_HUMAN T-complex protein 1 subunit zeta OS=Homo sapiens GN=CCTE | 1176,469 | 48,4 | 58024 | 28 | 28,07 | 28,04 | 28,14 | 28,40 | 28,45 | 28,44 | **1,272** | 4,83E-04 | **0** | 0,34666697 | 3,31629869 |
| PSA5_HUMA | PSMA5 | sp\|P28066\|PSA5_HUMAN Proteasome subunit alpha type-5 OS=Homo sapiens GN=PSM | 391,784 | 56,43 | 26411 | 11 | 25,96 | 26,17 | 26,64 | 26,31 | 26,58 | 26,92 | **1,272** | 2,65E-01 | **0** | 0,34666697 | 0,57722811 |
| TTL12_HUM/ | TTLL12 | sp\|Q14166\|TTL12_HUMAN Tubulin--tyrosine ligase-like protein 12 OS=Homo sapiens GI | 222,127 | 19,72 | 74404 | 9 | 24,41 | 24,57 | 23,03 | 24,48 | 24,62 | 23,95 | **1,272** | 5,49E-01 | **0** | 0,34666697 | 0,26077834 |
| PGRC1_HUM | PGRMC1 | sp\|O00264\|PGRC1_HUMAN Membrane-associated progesterone receptor component : | 168,053 | 26,15 | 21671 | 5 | 24,06 | 24,21 | 25,28 | 24,48 | 24,77 | 25,34 | **1,272** | 4,93E-01 | **0** | 0,34666697 | 0,30735371 |
| PGBM_HUM | HSPG2 | sp\|P98160\|PGBM_HUMAN Basement membrane-specific heparan sulfate proteoglycan | 1428,607 | 12,39 | 468830 | 38 | 28,21 | 26,41 | 27,19 | 28,38 | 27,44 | 27,03 | **1,272** | 6,26E-01 | **0** | 0,34666697 | 0,20376553 |
| HPCA_HUM/ | HPCA | sp\|P84074\|HPCA_HUMAN Neuron-specific calcium-binding protein hippocalcin OS=Hon | 50,978 | 12,44 | 22427 | 2 | 18,53 | 19,16 | 20,75 | 20,36 | 17,81 | 21,31 | **1,271** | 7,93E-01 | **0** | 0,34650548 | 0,10051162 |

| | | | | | | | | | | | | | | | | |
|---|---|---|---|---|---|---|---|---|---|---|---|---|---|---|---|---|
| CLCC1_HUM | CLCC1 | sp\|Q96S66\|CLCC1_HUMAN Chloride channel CLIC-like protein 1 OS=Homo sapiens GN= | 29,497 | 3,09 | 62023 | 1 | 17,74 | 19,07 | 20,16 | 18,57 | 19,30 | 20,12 | **1,270** | 6,99E-01 | **0** | 0,34516462 | 0,15546794 |
| HPRT_HUMA | HPRT1 | sp\|P00492\|HPRT_HUMAN Hypoxanthine-guanine phosphoribosyltransferase OS=Homo | 91,088 | 18,35 | 24579 | 5 | 23,85 | 23,56 | 22,59 | 23,89 | 24,04 | 23,10 | **1,269** | 5,14E-01 | **0** | 0,34333356 | 0,28924226 |
| IF4G3_HUM | EIF4G3 | sp\|O43432\|IF4G3_HUMAN Eukaryotic translation initiation factor 4 gamma 3 OS=Homo | 163,539 | 3,85 | 176652 | 6 | 22,40 | 22,06 | 22,85 | 22,65 | 22,73 | 22,96 | **1,269** | 2,37E-01 | **0** | 0,34333293 | 0,62565159 |
| SRSF7_HUM | SRSF7 | sp\|Q16629\|SRSF7_HUMAN Serine/arginine-rich splicing factor 7 OS=Homo sapiens GN= | 108,285 | 19,75 | 27367 | 4 | 22,79 | 22,36 | 23,52 | 22,32 | 22,54 | 24,84 | **1,269** | 7,15E-01 | **0** | 0,34333293 | 0,14599476 |
| NACAM_HU | NACA | sp\|E9PAV3\|NACAM_HUMAN Nascent polypeptide-associated complex subunit alpha, m | 310,776 | 3,37 | 205422 | 5 | 24,65 | 24,40 | 24,84 | 24,81 | 24,88 | 25,23 | **1,269** | 1,32E-01 | **0** | 0,34333293 | 0,87864794 |
| FRMD6_HUN | FRMD6 | sp\|Q96NE9\|FRMD6_HUMAN FERM domain-containing protein 6 OS=Homo sapiens GN= | 26,6 | 3,38 | 72044 | 1 | 18,88 | 18,17 | 17,85 | 18,48 | 17,61 | 19,83 | **1,266** | 6,59E-01 | **0** | 0,34021187 | 0,18134795 |
| LAMC1_HUN | LAMC1 | sp\|P11047\|LAMC1_HUMAN Laminin subunit gamma-1 OS=Homo sapiens GN=LAMC1 PE | 216,248 | 5,22 | 177603 | 7 | 23,51 | 20,63 | 24,21 | 23,12 | 21,77 | 24,48 | **1,266** | 8,13E-01 | **0** | 0,34000079 | 0,08988632 |
| LAMA5_HUN | LAMA5 | sp\|O15230\|LAMA5_HUMAN Laminin subunit alpha-5 OS=Homo sapiens GN=LAMA5 PE= | 90,466 | 1,35 | 399737 | 4 | 22,89 | 21,52 | 23,37 | 22,18 | 21,84 | 24,78 | **1,266** | 7,69E-01 | **0** | 0,34000015 | 0,11411634 |
| PYR1_HUMA | CAD | sp\|P27708\|PYR1_HUMAN CAD protein OS=Homo sapiens GN=CAD PE=1 SV=3 | 455,366 | 10,74 | 242984 | 21 | 24,55 | 24,95 | 25,81 | 24,94 | 25,16 | 26,23 | **1,266** | 5,66E-01 | **0** | 0,34000015 | 0,2468207 |
| VAV2_HUMA | VAV2 | sp\|P52735\|VAV2_HUMAN Guanine nucleotide exchange factor VAV2 OS=Homo sapiens | 11,272 | 1,59 | 101289 | 1 | 17,53 | 17,51 | 19,74 | 18,01 | 18,42 | 19,37 | **1,266** | 7,07E-01 | **0** | 0,34000015 | 0,15046411 |
| EIF3D_HUM | EIF3D | sp\|O15371\|EIF3D_HUMAN Eukaryotic translation initiation factor 3 subunit D OS=Homo | 367,148 | 25 | 63973 | 12 | 25,70 | 25,09 | 25,68 | 25,82 | 25,52 | 26,15 | **1,266** | 2,77E-01 | **0** | 0,33999952 | 0,55741702 |
| MED30_HUN | MED30 | sp\|Q96HR3\|MED30_HUMAN Mediator of RNA polymerase II transcription subunit 30 OS | 59,292 | 11,8 | 20277 | 2 | 20,24 | 18,63 | 21,67 | 20,80 | 18,87 | 21,88 | **1,264** | 7,99E-01 | **0** | 0,33777618 | 0,09721203 |
| PSD13_HUM | PSMD13 | sp\|Q9UNM6\|PSD13_HUMAN 26S proteasome non-ATPase regulatory subunit 13 OS=Ho | 617,152 | 43,09 | 42945 | 17 | 26,81 | 26,92 | 27,15 | 26,92 | 27,29 | 27,68 | **1,263** | 2,35E-01 | **0** | 0,33666738 | 0,62843971 |
| ANXA7_HUN | ANXA7 | sp\|P20073\|ANXA7_HUMAN Annexin A7 OS=Homo sapiens GN=ANXA7 PE=1 SV=3 | 447,772 | 22,54 | 52739 | 11 | 26,26 | 26,14 | 25,73 | 26,81 | 26,70 | 25,63 | **1,263** | 4,57E-01 | **0** | 0,33666674 | 0,34028393 |
| PDS5A_HUM | PDS5A | sp\|Q29RF7\|PDS5A_HUMAN Sister chromatid cohesion protein PDS5 homolog A OS=Hon | 232,067 | 11,74 | 150830 | 10 | 21,80 | 21,69 | 23,97 | 21,63 | 22,30 | 24,54 | **1,263** | 7,84E-01 | **0** | 0,33666674 | 0,1054165 |
| TMX2_HUM | TMX2 | sp\|Q9Y320\|TMX2_HUMAN Thioredoxin-related transmembrane protein 2 OS=Homo sa | 58,812 | 5,07 | 34038 | 1 | 18,62 | 19,22 | 20,13 | 19,07 | 19,14 | 20,77 | **1,263** | 6,59E-01 | **0** | 0,33666674 | 0,18099975 |
| LGUL_HUMA | GLO1 | sp\|Q04760\|LGUL_HUMAN Lactoylglutathione lyase OS=Homo sapiens GN=GLO1 PE=1 S | 36,609 | 7,61 | 20778 | 2 | 23,83 | 22,98 | 21,76 | 23,38 | 23,64 | 22,56 | **1,263** | 6,48E-01 | **0** | 0,33666611 | 0,18842268 |
| TBA1A_HUM | TUBA1A | sp\|Q71U36\|TBA1A_HUMAN Tubulin alpha-1A chain OS=Homo sapiens GN=TUBA1A PE= | 2338,417 | 93,57 | 50136 | 44 | 22,89 | 23,21 | 26,80 | 22,92 | 23,98 | 27,00 | **1,260** | 8,58E-01 | **0** | 0,33333397 | 0,06637712 |
| SC11A_HUM | SEC11A | sp\|P67812\|SC11A_HUMAN Signal peptidase complex catalytic subunit SEC11A OS=Hom | 133,256 | 34,64 | 20625 | 6 | 24,68 | 23,92 | 24,99 | 24,66 | 24,64 | 25,29 | **1,260** | 4,33E-01 | **0** | 0,33333333 | 0,36343411 |
| HIC2_HUMA | HIC2 | sp\|Q96JB3\|HIC2_HUMAN Hypermethylated in cancer 2 protein OS=Homo sapiens GN=H | 32,339 | 1,63 | 66156 | 1 | 33,11 | 32,81 | 31,76 | 33,18 | 33,13 | 32,37 | **1,260** | 5,30E-01 | **0** | 0,3333327 | 0,27534098 |
| RBBP7_HUM | RBBP7 | sp\|Q16576\|RBBP7_HUMAN Histone-binding protein RBBP7 OS=Homo sapiens GN=RBBF | 368,554 | 56,24 | 47820 | 12 | 24,50 | 24,99 | 25,38 | 24,75 | 25,26 | 25,85 | **1,257** | 4,63E-01 | **0** | 0,33000056 | 0,3342549 |
| TBB5_HUMA | TUBB | sp\|P07437\|TBB5_HUMAN Tubulin beta chain OS=Homo sapiens GN=TUBB PE=1 SV=2 | 2734,568 | 84,46 | 49671 | 55 | 27,55 | 27,73 | 30,02 | 27,96 | 28,32 | 30,01 | **1,257** | 7,62E-01 | **0** | 0,32999992 | 0,11832131 |
| HNRPK_HUN | HNRNPK | sp\|P61978\|HNRPK_HUMAN Heterogeneous nuclear ribonucleoprotein K OS=Homo sapi | 947,892 | 50,54 | 50976 | 22 | 27,44 | 27,59 | 28,92 | 27,75 | 27,94 | 29,25 | **1,257** | 6,46E-01 | **0** | 0,32999992 | 0,18960823 |
| CSN6_HUMA | COPS6 | sp\|Q7L5N1\|CSN6_HUMAN COP9 signalosome complex subunit 6 OS=Homo sapiens GN= | 153,344 | 19,88 | 36163 | 5 | 24,12 | 23,94 | 23,81 | 24,19 | 24,43 | 24,24 | **1,257** | 4,65E-02 | **0** | 0,32999992 | 1,33281733 |
| PRP6_HUMA | PRPF6 | sp\|O94906\|PRP6_HUMAN Pre-mRNA-processing factor 6 OS=Homo sapiens GN=PRPF6 | 72,192 | 6,38 | 106925 | 6 | 22,14 | 22,31 | 24,41 | 22,39 | 22,74 | 24,72 | **1,257** | 7,65E-01 | **0** | 0,32999992 | 0,11661118 |
| HNRPC_HUN | HNRNPC | sp\|P07910\|HNRPC_HUMAN Heterogeneous nuclear ribonucleoproteins C1/C2 OS=Hom | 306,102 | 19,61 | 33670 | 7 | 25,79 | 25,74 | 26,85 | 25,93 | 26,34 | 27,09 | **1,254** | 5,46E-01 | **0** | 0,32666651 | 0,26249326 |
| COPA_HUM | COPA | sp\|P53621\|COPA_HUMAN Coatomer subunit alpha OS=Homo sapiens GN=COPA PE=1 S | 1194,47 | 38,64 | 138346 | 38 | 26,94 | 27,10 | 27,38 | 27,15 | 27,38 | 27,87 | **1,254** | 2,59E-01 | **0** | 0,32666651 | 0,58752756 |
| CLCB_HUMA | CLTB | sp\|P09497\|CLCB_HUMAN Clathrin light chain B OS=Homo sapiens GN=CLTB PE=1 SV=1 | 172,651 | 24,45 | 25190 | 8 | 25,22 | 24,77 | 25,35 | 25,82 | 25,19 | 25,31 | **1,254** | 2,79E-01 | **0** | 0,32666651 | 0,55426175 |
| UNC5B_HUN | UNC5B | sp\|Q8IZJ1\|UNC5B_HUMAN Netrin receptor UNC5B OS=Homo sapiens GN=UNC5B PE=1 | 51,586 | 2,54 | 103638 | 2 | 20,29 | 20,69 | 18,28 | 20,53 | 19,87 | 19,84 | **1,254** | 6,96E-01 | **0** | 0,32654762 | 0,15737558 |
| ROA2_HUM | HNRNPA2B1 | sp\|P22626\|ROA2_HUMAN Heterogeneous nuclear ribonucleoproteins A2/B1 OS=Homo | 412,887 | 43,34 | 37430 | 16 | 25,67 | 25,64 | 26,64 | 26,08 | 26,31 | 26,53 | **1,251** | 4,12E-01 | **0** | 0,32333374 | 0,3853517 |
| EIF3M_HUM | EIF3M | sp\|Q7L2H7\|EIF3M_HUMAN Eukaryotic translation initiation factor 3 subunit M OS=Hom | 357,319 | 33,69 | 42503 | 9 | 24,26 | 24,80 | 29,73 | 24,22 | 25,10 | 30,44 | **1,251** | 9,07E-01 | **0** | 0,32333374 | 0,04223087 |
| SUMO2_HU | SUMO2 | sp\|P61956\|SUMO2_HUMAN Small ubiquitin-related modifier 2 OS=Homo sapiens GN=S | 109,669 | 27,37 | 10871 | 3 | 24,72 | 25,13 | 24,19 | 25,51 | 25,18 | 24,32 | **1,251** | 5,10E-01 | **0** | 0,32333374 | 0,29282063 |
| SPT6H_HUM | SUPT6H | sp\|Q7KZ85\|SPT6H_HUMAN Transcription elongation factor SPT6 OS=Homo sapiens GN= | 88,327 | 1,74 | 199073 | 3 | 20,74 | 20,28 | 22,56 | 20,16 | 21,53 | 22,86 | **1,251** | 7,72E-01 | **0** | 0,32333374 | 0,11212009 |
| SYAC_HUMA | AARS | sp\|P49588\|SYAC_HUMAN Alanine--tRNA ligase, cytoplasmic OS=Homo sapiens GN=AAR | 1154,429 | 34,92 | 106810 | 26 | 26,97 | 27,33 | 27,07 | 27,37 | 27,50 | 27,47 | **1,251** | 4,74E-02 | **0** | 0,32333374 | 1,32463003 |
| PGP_HUMAN | PGP | sp\|A6NDG6\|PGP_HUMAN Glycerol-3-phosphate phosphatase OS=Homo sapiens GN=PG | 101,782 | 12,46 | 34006 | 2 | 20,28 | 20,94 | 19,55 | 20,67 | 20,92 | 20,15 | **1,251** | 5,22E-01 | **0** | 0,3233331 | 0,28251096 |
| FPPS_HUMA | FDPS | sp\|P14324\|FPPS_HUMAN Farnesyl pyrophosphate synthase OS=Homo sapiens GN=FDP | 58,723 | 3,1 | 48275 | 2 | 22,83 | 23,10 | 23,10 | 23,37 | 23,38 | 23,25 | **1,251** | 3,11E-02 | **0** | 0,3233331 | 1,50700444 |
| STIP1_HUM | STIP1 | sp\|P31948\|STIP1_HUMAN Stress-induced-phosphoprotein 1 OS=Homo sapiens GN=STII | 1085,249 | 42,17 | 62639 | 34 | 28,52 | 28,45 | 27,36 | 28,65 | 28,61 | 28,04 | **1,251** | 4,88E-01 | **0** | 0,3233331 | 0,31132861 |
| NUDT5_HUN | NUDT5 | sp\|Q9UKK9\|NUDT5_HUMAN ADP-sugar pyrophosphatase OS=Homo sapiens GN=NUDT | 28,234 | 9,13 | 24328 | 2 | 22,74 | 22,73 | 21,61 | 22,82 | 23,01 | 22,22 | **1,251** | 5,07E-01 | **0** | 0,3233331 | 0,29499734 |
| FKBP2_HUM | FKBP2 | sp\|P26885\|FKBP2_HUMAN Peptidyl-prolyl cis-trans isomerase FKBP2 OS=Homo sapiens | 39,481 | 8,45 | 15649 | 1 | 22,12 | 22,02 | 21,95 | 22,73 | 22,57 | 21,76 | **1,251** | 3,48E-01 | **0** | 0,32333247 | 0,4586293 |
| TBB8_HUMA | TUBB8 | sp\|Q3ZCM7\|TBB8_HUMAN Tubulin beta-8 chain OS=Homo sapiens GN=TUBB8 PE=1 SV= | 887,961 | 25,23 | 49776 | 20 | 26,42 | 26,98 | 28,30 | 26,77 | 27,54 | 28,35 | **1,248** | 6,80E-01 | **0** | 0,32000097 | 0,16763136 |
| CKAP5_HUM | CKAP5 | sp\|Q14008\|CKAP5_HUMAN Cytoskeleton-associated protein 5 OS=Homo sapiens GN=C | 407,452 | 8,76 | 225495 | 13 | 25,00 | 25,05 | 23,71 | 25,21 | 25,32 | 24,19 | **1,248** | 6,03E-01 | **0** | 0,32000033 | 0,21983531 |
| RPAB3_HUM | POLR2H | sp\|P52434\|RPAB3_HUMAN DNA-directed RNA polymerases I, II, and III subunit RPABC3 | 131,617 | 21,33 | 17143 | 3 | 20,49 | 20,76 | 23,02 | 20,86 | 21,03 | 23,34 | **1,248** | 7,92E-01 | **0** | 0,32000033 | 0,10150573 |
| TXNL1_HUM | TXNL1 | sp\|O43396\|TXNL1_HUMAN Thioredoxin-like protein 1 OS=Homo sapiens GN=TXNL1 PE | 484,228 | 51,9 | 32251 | 13 | 25,81 | 25,88 | 25,48 | 25,99 | 26,16 | 25,98 | **1,248** | 7,89E-02 | **0** | 0,32000033 | 1,10265199 |
| PSB3_HUMA | PSMB3 | sp\|P49720\|PSB3_HUMAN Proteasome subunit beta type-3 OS=Homo sapiens GN=PSME | 440,602 | 48,29 | 22949 | 9 | 25,64 | 25,49 | 25,34 | 25,56 | 26,02 | 25,85 | **1,248** | 1,16E-01 | **0** | 0,32000033 | 0,93639985 |
| SYCC_HUMA | CARS | sp\|P49589\|SYCC_HUMAN Cysteine--tRNA ligase, cytoplasmic OS=Homo sapiens GN=CAF | 242,351 | 12,03 | 85473 | 8 | 23,96 | 24,74 | 24,08 | 24,35 | 25,08 | 24,31 | **1,248** | 4,10E-01 | **0** | 0,32000033 | 0,38676403 |
| NUDC2_HUN | NUDCD2 | sp\|Q8WVJ2\|NUDC2_HUMAN NudC domain-containing protein 2 OS=Homo sapiens GN= | 166,594 | 29,3 | 17676 | 3 | 22,42 | 21,51 | 23,17 | 22,72 | 22,47 | 22,87 | **1,248** | 5,52E-01 | **0** | 0,31999969 | 0,25776726 |
| HSPB1_HUM | HSPB1 | sp\|P04792\|HSPB1_HUMAN Heat shock protein beta-1 OS=Homo sapiens GN=HSPB1 PE= | 285,886 | 25,37 | 22783 | 10 | 26,86 | 26,96 | 26,45 | 27,40 | 27,14 | 26,69 | **1,248** | 2,85E-01 | **0** | 0,31999969 | 0,54502763 |
| FHOD1_HUN | FHOD1 | sp\|Q9Y613\|FHOD1_HUMAN FH1/FH2 domain-containing protein 1 OS=Homo sapiens G | 14,919 | 2,49 | 126551 | 1 | 18,29 | 19,13 | 18,67 | 18,76 | 19,32 | 18,97 | **1,248** | 3,36E-01 | **0** | 0,31974983 | 0,4736965 |
| KAT7_HUMA | KAT7 | sp\|O95251\|KAT7_HUMAN Histone acetyltransferase KAT7 OS=Homo sapiens GN=KAT7 | 13,081 | 3,93 | 70642 | 1 | 19,34 | 18,07 | 19,66 | 20,09 | 18,08 | 19,86 | **1,248** | 7,10E-01 | **0** | 0,31921514 | 0,14855373 |
| UD3A2_HUN | UGT3A2 | sp\|Q3SY77\|UD3A2_HUMAN UDP-glucuronosyltransferase 3A2 OS=Homo sapiens GN=U | 15,973 | 2,1 | 59547 | 1 | 21,30 | 19,14 | 18,18 | 21,53 | 20,75 | 17,29 | **1,246** | 8,52E-01 | **0** | 0,31764857 | 0,06954365 |
| ALG9_HUMA | ALG9 | sp\|Q9H6U8\|ALG9_HUMAN Alpha-1,2-mannosyltransferase ALG9 OS=Homo sapiens GN | 58,766 | 2,62 | 69863 | 1 | 19,79 | 19,94 | 19,70 | 20,07 | 20,15 | 20,16 | **1,246** | 1,36E-02 | **0** | 0,31698291 | 1,8667805 |
| RSU1_HUMA | RSU1 | sp\|Q15404\|RSU1_HUMAN Ras suppressor protein 1 OS=Homo sapiens GN=RSU1 PE=1 S | 302,866 | 52,71 | 31540 | 11 | 26,87 | 26,34 | 25,67 | 27,02 | 26,69 | 26,12 | **1,245** | 5,07E-01 | **0** | 0,31666692 | 0,29464882 |
| XPP1_HUMA | XPNPEP1 | sp\|Q9NQW7\|XPP1_HUMAN Xaa-Pro aminopeptidase 1 OS=Homo sapiens GN=XPNPEP1 | 221,304 | 16,69 | 69918 | 8 | 23,81 | 24,46 | 23,77 | 24,13 | 24,58 | 24,28 | **1,245** | 2,90E-01 | **0** | 0,31666692 | 0,53775439 |
| CUL2_HUMA | CUL2 | sp\|Q13617\|CUL2_HUMAN Cullin-2 OS=Homo sapiens GN=CUL2 PE=1 SV=2 | 132,909 | 11,41 | 86983 | 6 | 22,30 | 23,29 | 22,43 | 22,33 | 22,96 | 23,68 | **1,245** | 5,60E-01 | **0** | 0,31666629 | 0,25192653 |
| PPP5_HUMA | PPP5C | sp\|P53041\|PPP5_HUMAN Serine/threonine-protein phosphatase 5 OS=Homo sapiens G | 135,238 | 8,22 | 56879 | 4 | 22,44 | 22,25 | 22,52 | 22,44 | 23,13 | 22,59 | **1,245** | 2,31E-01 | **0** | 0,31666629 | 0,63665162 |
| RL7A_HUMA | RPL7A | sp\|P62424\|RL7A_HUMAN 60S ribosomal protein L7a OS=Homo sapiens GN=RPL7A PE=1 | 847,491 | 54,89 | 29996 | 21 | 27,70 | 27,88 | 27,84 | 27,99 | 27,74 | 28,64 | **1,245** | 3,12E-01 | **0** | 0,31666629 | 0,50626822 |
| BIEA_HUMA | BLVRA | sp\|P53004\|BIEA_HUMAN Biliverdin reductase A OS=Homo sapiens GN=BLVRA PE=1 SV= | 118,918 | 16,22 | 33428 | 3 | 22,74 | 23,04 | 22,09 | 22,80 | 22,97 | 23,05 | **1,245** | 3,36E-01 | **0** | 0,31666565 | 0,47356473 |
| QBK84317.1 | | tegument protein TRS1 [Human betaherpesvirus 5] | 395,699 | 30,93 | 84110 | 15 | 18,44 | 17,18 | 19,87 | 18,54 | 18,35 | 19,54 | **1,243** | 7,33E-01 | **0** | 0,31404813 | 0,13490851 |
| UBA5_HUM | UBA5 | sp\|Q9GZZ9\|UBA5_HUMAN Ubiquitin-like modifier-activating enzyme 5 OS=Homo sapier | 23,321 | 8,42 | 44863 | 2 | 19,75 | 21,06 | 18,71 | 19,90 | 20,28 | 20,28 | **1,243** | 6,74E-01 | **0** | 0,31333415 | 0,17134949 |
| PFD3_HUMA | VBP1 | sp\|P61758\|PFD3_HUMAN Prefoldin subunit 3 OS=Homo sapiens GN=VBP1 PE=1 SV=3 | 138,165 | 32,99 | 22658 | 4 | 23,58 | 23,89 | 22,66 | 23,81 | 23,67 | 23,59 | **1,243** | 4,50E-01 | **0** | 0,31333351 | 0,34650918 |
| TBA4A_HUM | TUBA4A | sp\|P68366\|TBA4A_HUMAN Tubulin alpha-4A chain OS=Homo sapiens GN=TUBA4A PE=1 | 1823,774 | 71,65 | 49924 | 35 | 26,95 | 26,98 | 29,07 | 27,32 | 27,51 | 29,11 | **1,243** | 7,46E-01 | **0** | 0,31333351 | 0,12727809 |
| TBC13_HUM | TBC1D13 | sp\|Q9NVG8\|TBC13_HUMAN TBC1 domain family member 13 OS=Homo sapiens GN=TB | 78,527 | 5,5 | 46554 | 2 | 20,23 | 21,52 | 20,03 | 21,32 | 21,09 | 20,31 | **1,243** | 6,04E-01 | **0** | 0,31333288 | 0,21864673 |
| HNRDL_HUN | HNRNPDL | sp\|O14979\|HNRDL_HUMAN Heterogeneous nuclear ribonucleoprotein D-like OS=Homo | 206,84 | 24,29 | 46438 | 8 | 23,61 | 23,91 | 24,01 | 24,01 | 24,23 | 24,23 | **1,243** | 9,01E-02 | **0** | 0,31333288 | 1,04547555 |
| EPN4_HUMA | CLINT1 | sp\|Q14677\|EPN4_HUMAN Clathrin interactor 1 OS=Homo sapiens GN=CLINT1 PE=1 SV= | 143,603 | 9,92 | 68259 | 5 | 21,86 | 22,41 | 22,10 | 22,54 | 22,64 | 22,13 | **1,243** | 2,33E-01 | **0** | 0,31333288 | 0,63349685 |
| UBP24_HUM | USP24 | sp\|Q9UPU5\|UBP24_HUMAN Ubiquitin carboxyl-terminal hydrolase 24 OS=Homo sapien | 14,45 | 0,69 | 294365 | 1 | 17,45 | 17,90 | 18,40 | 17,22 | 18,12 | 19,34 | **1,240** | 6,69E-01 | **0** | 0,31017749 | 0,17476631 |
| EI2BG_HUM | EIF2B3 | sp\|Q9NR50\|EI2BG_HUMAN Translation initiation factor eIF-2B subunit gamma OS=Hom | 61,879 | 8,63 | 50240 | 3 | 21,31 | 21,70 | 22,25 | 21,52 | 21,86 | 22,81 | **1,240** | 5,48E-01 | **0** | 0,3100001 | 0,26147548 |
| MANF_HUM | MANF | sp\|P55145\|MANF_HUMAN Mesencephalic astrocyte-derived neurotrophic factor OS=H | 36,781 | 19,78 | 20700 | 3 | 23,30 | 22,96 | 23,36 | 23,65 | 23,51 | 23,39 | **1,240** | 1,00E-01 | **0** | 0,3100001 | 0,99974279 |
| PRP8_HUMA | PRPF8 | sp\|Q6P2Q9\|PRP8_HUMAN Pre-mRNA-processing-splicing factor 8 OS=Homo sapiens GN | 711,632 | 16,32 | 273600 | 30 | 25,10 | 25,12 | 26,08 | 25,27 | 25,66 | 26,30 | **1,240** | 5,21E-01 | **0** | 0,30999947 | 0,28308692 |
| DNJC3_HUM | DNAJC3 | sp\|Q13217\|DNJC3_HUMAN DnaJ homolog subfamily C member 3 OS=Homo sapiens GN | 22,011 | 1,98 | 57580 | 1 | 20,05 | 19,84 | 21,97 | 20,17 | 19,87 | 22,74 | **1,238** | 8,00E-01 | **0** | 0,30819829 | 0,09715399 |
| PA2G4_HUM | PA2G4 | sp\|Q9UQ80\|PA2G4_HUMAN Proliferation-associated protein 2G4 OS=Homo sapiens GN | 752,939 | 55,33 | 43787 | 19 | 26,89 | 26,98 | 27,43 | 26,98 | 27,19 | 28,05 | **1,237** | 4,51E-01 | **0** | 0,30666669 | 0,34588432 |
| NSF1C_HUM | NSFL1C | sp\|Q9UNZ2\|NSF1C_HUMAN NSFL1 cofactor p47 OS=Homo sapiens GN=NSFL1C PE=1 SV | 268,52 | 31,62 | 40573 | 10 | 25,83 | 25,83 | 23,85 | 26,01 | 26,04 | 24,38 | **1,237** | 7,39E-01 | **0** | 0,30666669 | 0,13143302 |
| GNAZ_HUM | GNAZ | sp\|P19086\|GNAZ_HUMAN Guanine nucleotide-binding protein G(z) subunit alpha OS=H | 94,168 | 6,2 | 40924 | 2 | 20,83 | 20,65 | 19,21 | 20,78 | 20,57 | 20,26 | **1,237** | 5,97E-01 | **0** | 0,30659167 | 0,22413601 |
| ROA3_HUM | HNRNPA3 | sp\|P51991\|ROA3_HUMAN Heterogeneous nuclear ribonucleoprotein A3 OS=Homo sapi | 537,086 | 42,33 | 39595 | 17 | 25,14 | 25,58 | 27,19 | 25,69 | 26,09 | 27,04 | **1,234** | 7,03E-01 | **0** | 0,30333392 | 0,15297967 |
| DAD1_HUM | DAD1 | sp\|P61803\|DAD1_HUMAN Dolichyl-diphosphooligosaccharide--protein glycosyltransfer | 48,574 | 19,47 | 12497 | 2 | 22,14 | 21,01 | 22,06 | 21,90 | 21,63 | 22,59 | **1,234** | 5,48E-01 | **0** | 0,30333328 | 0,26119848 |
| TPM3_HUM | TPM3 | sp\|P06753\|TPM3_HUMAN Tropomyosin alpha-3 chain OS=Homo sapiens GN=TPM3 PE= | 710,189 | 42,46 | 32950 | 19 | 26,16 | 26,41 | 24,45 | 26,57 | 26,50 | 24,86 | **1,234** | 7,34E-01 | **0** | 0,30333328 | 0,1344546 |
| SND1_HUM | SND1 | sp\|Q7KZF4\|SND1_HUMAN Staphylococcal nuclease domain-containing protein 1 OS=Ho | 846,066 | 27,58 | 101997 | 21 | 26,86 | 26,84 | 27,17 | 27,07 | 27,02 | 27,69 | **1,234** | 2,76E-01 | **0** | 0,30333328 | 0,55944594 |
| PDLI4_HUM | PDLIM4 | sp\|P50479\|PDLI4_HUMAN PDZ and LIM domain protein 4 OS=Homo sapiens GN=PDLIM | 13,654 | 3,03 | 35398 | 1 | 21,32 | 20,90 | 20,01 | 21,41 | 21,22 | 20,51 | **1,234** | 5,57E-01 | **0** | 0,30333328 | 0,25446611 |
| PSD11_HUM | PSMD11 | sp\|O00231\|PSD11_HUMAN 26S proteasome non-ATPase regulatory subunit 11 OS=Hon | 869,007 | 64,93 | 47464 | 26 | 26,91 | 27,14 | 27,68 | 27,18 | 27,41 | 28,05 | **1,234** | 4,30E-01 | **0** | 0,30333328 | 0,36618726 |
| FAS_HUMAN | FASN | sp\|P49327\|FAS_HUMAN Fatty acid synthase OS=Homo sapiens GN=FASN PE=1 SV=3 | 487,408 | 6,25 | 273427 | 16 | 26,95 | 26,98 | 26,24 | 27,26 | 27,16 | 26,66 | **1,234** | 3,76E-01 | **0** | 0,30333328 | 0,42479322 |
| SPTB2_HUM | SPTBN1 | sp\|Q01082\|SPTB2_HUMAN Spectrin beta chain, non-erythrocytic 1 OS=Homo sapiens G | 1252,173 | 21,79 | 274609 | 38 | 26,29 | 26,84 | 25,27 | 26,71 | 26,92 | 25,68 | **1,234** | 6,39E-01 | **0** | 0,30333265 | 0,19449117 |
| PLCB_HUMA | AGPAT2 | sp\|O15120\|PLCB_HUMAN 1-acyl-sn-glycerol-3-phosphate acyltransferase beta OS=Hor | 16,025 | 3,24 | 30914 | 1 | 21,90 | 21,66 | 22,07 | 21,96 | 22,37 | 22,20 | **1,231** | 1,49E-01 | **0** | 0,30000051 | 0,82667826 |
| TMM43_HUI | TMEM43 | sp\|Q9BTV4\|TMM43_HUMAN Transmembrane protein 43 OS=Homo sapiens GN=TMEM | 248,729 | 24,75 | 44876 | 8 | 24,38 | 23,83 | 25,14 | 24,62 | 24,39 | 25,24 | **1,231** | 5,47E-01 | **0** | 0,30000051 | 0,26184585 |
| RL35A_HUM | RPL35A | sp\|P18077\|RL35A_HUMAN 60S ribosomal protein L35a OS=Homo sapiens GN=RPL35A P | 106,453 | 35,45 | 12538 | 8 | 26,65 | 26,72 | 26,80 | 26,89 | 26,89 | 27,29 | **1,231** | 9,91E-02 | **0** | 0,30000051 | 1,00389359 |
| CBPD_HUMA | CPD | sp\|O75976\|CBPD_HUMAN Carboxypeptidase D OS=Homo sapiens GN=CPD PE=1 SV=2 | 15,181 | 0,94 | 152931 | 1 | 20,40 | 20,36 | 20,21 | 20,93 | 20,16 | 20,78 | **1,231** | 2,84E-01 | **0** | 0,30000051 | 0,54668447 |
| CDC37_HUM | CDC37 | sp\|Q16543\|CDC37_HUMAN Hsp90 co-chaperone Cdc37 OS=Homo sapiens GN=CDC37 F | 71,108 | 7,94 | 44468 | 3 | 23,28 | 23,47 | 23,13 | 23,73 | 23,72 | 23,33 | **1,231** | 1,42E-01 | **0** | 0,29999987 | 0,84755525 |
| RS20_HUMA | RPS20 | sp\|P60866\|RS20_HUMAN 40S ribosomal protein S20 OS=Homo sapiens GN=RPS20 PE=1 | 108,814 | 22,69 | 13373 | 3 | 26,16 | 26,43 | 26,10 | 26,54 | 26,56 | 26,49 | **1,231** | 4,43E-02 | **0** | 0,29999987 | 1,35351161 |
| ERF3A_HUM | GSPT1 | sp\|P15170\|ERF3A_HUMAN Eukaryotic peptide chain release factor GTP-binding subunit | 507,964 | 47,29 | 55756 | 19 | 23,93 | 24,11 | 24,35 | 23,76 | 25,01 | 24,52 | **1,231** | 4,78E-01 | **0** | 0,29999987 | 0,32082732 |
| ECT2_HUMA | ECT2 | sp\|Q9H8V3\|ECT2_HUMAN Protein ECT2 OS=Homo sapiens GN=ECT2 PE=1 SV=4 | 16,147 | 1,53 | 103505 | 1 | 18,06 | 17,33 | 18,92 | 17,81 | 18,06 | 19,33 | **1,229** | 6,74E-01 | **0** | 0,29782104 | 0,17133006 |
| ABCG2_HUN | ABCG2 | sp\|Q9UNQ0\|ABCG2_HUMAN ATP-binding cassette sub-family G member 2 OS=Homo sa | 102,978 | 5,65 | 72314 | 4 | 25,25 | 25,46 | 20,48 | 25,02 | 25,40 | 21,66 | **1,228** | 8,90E-01 | **0** | 0,2966671 | 0,05059763 |
| PHAX_HUM | PHAX | sp\|Q9H814\|PHAX_HUMAN Phosphorylated adapter RNA export protein OS=Homo sapie | 38,853 | 2,79 | 44403 | 1 | 20,80 | 20,85 | 20,94 | 21,00 | 21,29 | 21,19 | **1,228** | 3,48E-02 | **0** | 0,2966671 | 1,45895067 |
| GBG12_HUN | GNG12 | sp\|Q9UBI6\|GBG12_HUMAN Guanine nucleotide-binding protein G(I)/G(S)/G(O) subunit | 55,522 | 44,44 | 8006 | 3 | 23,51 | 22,49 | 22,40 | 23,67 | 23,01 | 22,61 | **1,228** | 5,63E-01 | **0** | 0,2966671 | 0,24927537 |
| COPB_HUM | COPB1 | sp\|P53618\|COPB_HUMAN Coatomer subunit beta OS=Homo sapiens GN=COPB1 PE=1 S | 620,997 | 27,49 | 107142 | 21 | 25,86 | 26,18 | 26,57 | 26,01 | 26,54 | 26,95 | **1,228** | 4,33E-01 | **0** | 0,2966671 | 0,3633007 |
| SC31A_HUM | SEC31A | sp\|O94979\|SC31A_HUMAN Protein transport protein Sec31A OS=Homo sapiens GN=SE | 344,899 | 10,82 | 133015 | 12 | 23,76 | 24,16 | 24,35 | 24,24 | 24,40 | 24,52 | **1,228** | 1,97E-01 | **0** | 0,29666646 | 0,70565533 |
| HS105_HUM | HSPH1 | sp\|Q92598\|HS105_HUMAN Heat shock protein 105 kDa OS=Homo sapiens GN=HSPH1 P | 555,086 | 21,56 | 96865 | 14 | 25,00 | 24,98 | 24,98 | 25,01 | 25,38 | 25,46 | **1,228** | 9,93E-02 | **0** | 0,29666646 | 1,00296508 |
| SAC1_HUMA | SACM1L | sp\|Q9NTJ5\|SAC1_HUMAN Phosphatidylinositide phosphatase SAC1 OS=Homo sapiens G | 230,967 | 17,38 | 66967 | 10 | 24,89 | 24,25 | 25,27 | 24,79 | 25,05 | 25,46 | **1,228** | 4,51E-01 | **0** | 0,29666646 | 0,34551699 |
| DHX32_HUM | DHX32 | sp\|Q7L7V1\|DHX32_HUMAN Putative pre-mRNA-splicing factor ATP-dependent RNA heli | 46,24 | 1,62 | 84419 | 1 | 17,49 | 18,19 | 20,11 | 18,64 | 17,88 | 20,16 | **1,226** | 7,89E-01 | **0** | 0,29436556 | 0,10281846 |
| IF2G_HUMA | EIF2S3 | sp\|P41091\|IF2G_HUMAN Eukaryotic translation initiation factor 2 subunit 3 OS=Homo s | 574,412 | 52,54 | 51109 | 20 | 26,89 | 27,14 | 26,66 | 27,10 | 27,36 | 27,11 | **1,225** | 1,46E-01 | **0** | 0,29333433 | 0,83687443 |
| TBB1_HUMA | TUBB1 | sp\|Q9H4B7\|TBB1_HUMAN Tubulin beta-1 chain OS=Homo sapiens GN=TUBB1 PE=1 SV= | 506,523 | 21,95 | 50327 | 12 | 26,22 | 23,05 | 24,69 | 25,57 | 25,32 | 23,95 | **1,225** | 7,93E-01 | **0** | 0,29333369 | 0,10084093 |
| DC1I2_HUM | DYNC1I2 | sp\|Q13409\|DC1I2_HUMAN Cytoplasmic dynein 1 intermediate chain 2 OS=Homo sapien | 375,105 | 31,03 | 71457 | 10 | 24,99 | 25,17 | 24,92 | 25,28 | 25,56 | 25,12 | **1,225** | 1,20E-01 | **0** | 0,29333369 | 0,92228134 |
| SYSC_HUMA | SARS | sp\|P49591\|SYSC_HUMAN Serine--tRNA ligase, cytoplasmic OS=Homo sapiens GN=SARS | 422,754 | 26,46 | 58777 | 11 | 25,37 | 25,47 | 25,55 | 25,63 | 25,76 | 25,88 | **1,225** | 3,01E-02 | **0** | 0,29333305 | 1,52206553 |
| PCH2_HUMA | TRIP13 | sp\|Q15645\|PCH2_HUMAN Pachytene checkpoint protein 2 homolog OS=Homo sapiens | 64,622 | 12,27 | 48551 | 4 | 23,90 | 22,55 | 24,09 | 23,72 | 23,31 | 24,39 | **1,225** | 6,39E-01 | **0** | 0,29333305 | 0,19483136 |
| AK1A1_HUM | AKR1A1 | sp\|P14550\|AK1A1_HUMAN Alcohol dehydrogenase [NADP(+)] OS=Homo sapiens GN=AI | 296,716 | 32 | 36573 | 12 | 26,64 | 26,62 | 25,62 | 26,75 | 26,71 | 26,30 | **1,225** | 4,68E-01 | **0** | 0,29333242 | 0,32982055 |
| PAR4_HUMA | F2RL3 | sp\|Q96RI0\|PAR4_HUMAN Proteinase-activated receptor 4 OS=Homo sapiens GN=F2RL3 | 27,988 | 2,6 | 41133 | 1 | 21,66 | 20,61 | 18,93 | 21,29 | 20,86 | 19,93 | **1,224** | 7,59E-01 | **0** | 0,29211171 | 0,1196971 |
| LAP2A_HUM | TMPO | sp\|P42166\|LAP2A_HUMAN Lamina-associated polypeptide 2, isoform alpha OS=Homo s | 66,774 | 4,18 | 75492 | 2 | 22,07 | 22,37 | 20,71 | 22,61 | 22,55 | 20,86 | **1,223** | 7,25E-01 | **0** | 0,29000028 | 0,13971416 |
| PRDX6_HUM | PRDX6 | sp\|P30041\|PRDX6_HUMAN Peroxiredoxin-6 OS=Homo sapiens GN=PRDX6 PE=1 SV=3 | 256,781 | 39,73 | 25035 | 9 | 26,99 | 27,13 | 25,66 | 27,14 | 27,16 | 26,35 | **1,223** | 6,19E-01 | **0** | 0,29000028 | 0,20822911 |
| MCM7_HUM | MCM7 | sp\|P33993\|MCM7_HUMAN DNA replication licensing factor MCM7 OS=Homo sapiens G | 634,057 | 39,92 | 81308 | 21 | 25,01 | 24,94 | 27,19 | 25,28 | 25,50 | 27,23 | **1,223** | 7,78E-01 | **0** | 0,28999964 | 0,10895036 |
| ABHGA_HUN | ABHD16A | sp\|O95870\|ABHGA_HUMAN Protein ABHD16A OS=Homo sapiens GN=ABHD16A PE=1 S\ | 14,974 | 1,61 | 63243 | 1 | 20,61 | 21,00 | 21,28 | 21,11 | 21,27 | 21,38 | **1,223** | 2,38E-01 | **0** | 0,28999964 | 0,62252128 |
| CSN8_HUMA | COPS8 | sp\|Q99627\|CSN8_HUMAN COP9 signalosome complex subunit 8 OS=Homo sapiens GN= | 210,427 | 46,89 | 23226 | 4 | 22,19 | 22,66 | 22,76 | 22,60 | 22,67 | 23,21 | **1,223** | 3,28E-01 | **0** | 0,28999964 | 0,48346977 |
| H33_HUMAN | H3F3A | sp\|P84243\|H33_HUMAN Histone H3.3 OS=Homo sapiens GN=H3F3A PE=1 SV=2 | 166,921 | 43,38 | 15328 | 5 | 21,78 | 22,89 | 23,29 | 22,55 | 22,74 | 23,54 | **1,223** | 6,22E-01 | **0** | 0,28999964 | 0,20602562 |
| GEMI4_HUM | GEMIN4 | sp\|P57678\|GEMI4_HUMAN Gem-associated protein 4 OS=Homo sapiens GN=GEMIN4 P | 41,882 | 1,23 | 120037 | 1 | 18,26 | 18,67 | 21,58 | 20,01 | 18,31 | 21,06 | **1,223** | 8,37E-01 | **0** | 0,28994815 | 0,07752946 |
| CBX5_HUMA | CBX5 | sp\|P45973\|CBX5_HUMAN Chromobox protein homolog 5 OS=Homo sapiens GN=CBX5 F | 164,75 | 25,65 | 22225 | 4 | 22,82 | 22,83 | 23,01 | 23,20 | 22,99 | 23,33 | **1,220** | 7,00E-02 | **0** | 0,28666687 | 1,15501745 |
| S38A2_HUM | SLC38A2 | sp\|Q96QD8\|S38A2_HUMAN Sodium-coupled neutral amino acid transporter 2 OS=Hom | 205,875 | 9,29 | 56026 | 3 | 23,30 | 23,97 | 24,82 | 24,23 | 24,32 | 24,40 | **1,220** | 5,52E-01 | **0** | 0,28666687 | 0,25771762 |
| GDIA_HUMA | GDI1 | sp\|P31150\|GDIA_HUMAN Rab GDP dissociation inhibitor alpha OS=Homo sapiens GN=G | 1054,747 | 58,17 | 50583 | 23 | 25,72 | 25,89 | 24,09 | 25,49 | 26,05 | 25,02 | **1,220** | 6,80E-01 | **0** | 0,28666687 | 0,1672661 |
| TCPB_HUMA | CCT2 | sp\|P78371\|TCPB_HUMAN T-complex protein 1 subunit beta OS=Homo sapiens GN=CCT | 1405,346 | 64,49 | 57488 | 29 | 28,25 | 28,41 | 28,18 | 28,53 | 28,60 | 28,57 | **1,220** | 1,57E-02 | **0** | 0,28666687 | 1,80539693 |
| SYEP_HUMA | EPRS | sp\|P07814\|SYEP_HUMAN Bifunctional glutamate/proline--tRNA ligase OS=Homo sapien | 700,309 | 20,3 | 170591 | 26 | 26,21 | 26,58 | 27,00 | 26,50 | 26,66 | 27,49 | **1,220** | 4,95E-01 | **0** | 0,28666687 | 0,30527457 |
| CAPG_HUM | CAPG | sp\|P40121\|CAPG_HUMAN Macrophage-capping protein OS=Homo sapiens GN=CAPG PI | 231,14 | 13,79 | 38499 | 4 | 25,49 | 25,59 | 24,71 | 25,71 | 25,75 | 25,19 | **1,220** | 4,36E-01 | **0** | 0,28666687 | 0,36052662 |
| RL17_HUMA | RPL17 | sp\|P18621\|RL17_HUMAN 60S ribosomal protein L17 OS=Homo sapiens GN=RPL17 PE=1 | 318,604 | 41,85 | 21397 | 7 | 25,65 | 26,31 | 26,93 | 26,08 | 26,23 | 27,44 | **1,220** | 6,40E-01 | **0** | 0,28666687 | 0,19385067 |
| PA1B2_HUM | PAFAH1B2 | sp\|P68402\|PA1B2_HUMAN Platelet-activating factor acetylhydrolase IB subunit beta OS | 266,223 | 48,47 | 25569 | 7 | 24,52 | 24,59 | 25,00 | 24,54 | 24,98 | 25,45 | **1,220** | 3,97E-01 | **0** | 0,28666687 | 0,40140715 |
| NALCN_HUN | NALCN | sp\|Q8IZF0\|NALCN_HUMAN Sodium leak channel non-selective protein OS=Homo sapier | 10,755 | 0,63 | 200331 | 1 | 21,79 | 20,85 | 22,89 | 21,81 | 21,69 | 22,89 | **1,220** | 7,04E-01 | **0** | 0,28666623 | 0,15241949 |
| DHR13_HUM | DHRS13 | sp\|Q6UX07\|DHR13_HUMAN Dehydrogenase/reductase SDR family member 13 OS=Hom | 21,845 | 3,45 | 40849 | 1 | 20,64 | 20,37 | 20,61 | 21,15 | 20,47 | 20,86 | **1,220** | 2,53E-01 | **0** | 0,28666623 | 0,59722799 |
| ILEU_HUMA | SERPINB1 | sp\|P30740\|ILEU_HUMAN Leukocyte elastase inhibitor OS=Homo sapiens GN=SERPINB1 | 35,375 | 2,64 | 42742 | 1 | 21,58 | 21,62 | 21,95 | 21,97 | 21,93 | 22,11 | **1,220** | 9,09E-02 | **0** | 0,28666623 | 1,04127305 |
| FA98A_HUM | FAM98A | sp\|Q8NCA5\|FA98A_HUMAN Protein FAM98A OS=Homo sapiens GN=FAM98A PE=1 SV= | 130,624 | 10,79 | 55401 | 5 | 21,29 | 20,95 | 21,10 | 21,26 | 21,14 | 21,80 | **1,220** | 2,73E-01 | **0** | 0,2866656 | 0,56440083 |
| CSTF2_HUM | CSTF2 | sp\|P33240\|CSTF2_HUMAN Cleavage stimulation factor subunit 2 OS=Homo sapiens GN= | 61,58 | 12,65 | 60959 | 4 | 17,02 | 20,13 | 21,20 | 16,96 | 19,99 | 22,26 | **1,219** | 8,92E-01 | **0** | 0,28540357 | 0,04942749 |
| DRG1_HUM | DRG1 | sp\|Q9Y295\|DRG1_HUMAN Developmentally-regulated GTP-binding protein 1 OS=Homo | 74,771 | 13,08 | 40542 | 3 | 22,60 | 22,96 | 22,89 | 22,81 | 23,31 | 23,18 | **1,217** | 2,02E-01 | **0** | 0,28333346 | 0,69409551 |
| SYTC2_HUM | TARSL2 | sp\|A2RTX5\|SYTC2_HUMAN Probable threonine--tRNA ligase 2, cytoplasmic OS=Homo sa | 46,079 | 2,99 | 92646 | 3 | 18,75 | 18,02 | 18,13 | 18,58 | 18,70 | 18,47 | **1,217** | 2,97E-01 | **0** | 0,28333346 | 0,52657635 |
| ERGI2_HUM | ERGIC2 | sp\|Q96RQ1\|ERGI2_HUMAN Endoplasmic reticulum-Golgi intermediate compartment pr | 11,118 | 2,39 | 42549 | 1 | 20,85 | 19,34 | 21,30 | 20,84 | 19,81 | 21,69 | **1,217** | 7,42E-01 | **0** | 0,28333346 | 0,12936939 |
| FKBP3_HUM | FKBP3 | sp\|Q00688\|FKBP3_HUMAN Peptidyl-prolyl cis-trans isomerase FKBP3 OS=Homo sapiens | 89,817 | 21,43 | 25177 | 4 | 24,64 | 25,26 | 22,56 | 24,63 | 25,00 | 23,68 | **1,217** | 7,70E-01 | **0** | 0,28333346 | 0,11341441 |
| G3P_HUMAN | GAPDH | sp\|P04406\|G3P_HUMAN Glyceraldehyde-3-phosphate dehydrogenase OS=Homo sapier | 1097,103 | 59,7 | 36053 | 23 | 30,90 | 30,85 | 30,50 | 31,23 | 31,13 | 30,74 | **1,217** | 2,21E-01 | **0** | 0,28333282 | 0,65631151 |
| STXB2_HUM | STXBP2 | sp\|Q15833\|STXB2_HUMAN Syntaxin-binding protein 2 OS=Homo sapiens GN=STXBP2 P | 178,813 | 12,82 | 66453 | 7 | 23,79 | 22,57 | 19,69 | 23,75 | 22,92 | 20,23 | **1,217** | 8,69E-01 | **0** | 0,28333282 | 0,06087857 |
| TRMB_HUM | METTL1 | sp\|Q9UBP6\|TRMB_HUMAN tRNA (guanine-N(7)-)-methyltransferase OS=Homo sapiens | 20,947 | 5,43 | 31471 | 1 | 18,25 | 18,27 | 18,95 | 17,59 | 19,07 | 19,66 | **1,217** | 6,89E-01 | **0** | 0,28333282 | 0,16201357 |
| C42S2_HUM | CDC42SE2 | sp\|Q9NRR3\|C42S2_HUMAN CDC42 small effector protein 2 OS=Homo sapiens GN=CDC | 27,153 | 44,05 | 9223 | 1 | 22,33 | 22,41 | 20,40 | 22,33 | 22,40 | 21,26 | **1,216** | 7,27E-01 | **0** | 0,28225072 | 0,13866164 |
| G6PI_HUMA | GPI | sp\|P06744\|G6PI_HUMAN Glucose-6-phosphate isomerase OS=Homo sapiens GN=GPI PI | 456,522 | 31,54 | 63147 | 12 | 25,92 | 26,14 | 23,88 | 25,75 | 26,16 | 24,87 | **1,214** | 7,48E-01 | **0** | 0,28000069 | 0,1260074 |
| RAGP1_HUM | RANGAP1 | sp\|P46060\|RAGP1_HUMAN Ran GTPase-activating protein 1 OS=Homo sapiens GN=RAN | 160,427 | 11,93 | 63542 | 5 | 23,88 | 24,16 | 22,97 | 24,02 | 24,26 | 23,57 | **1,214** | 5,34E-01 | **0** | 0,28000069 | 0,27224662 |
| RL38_HUMA | RPL38 | sp\|P63173\|RL38_HUMAN 60S ribosomal protein L38 OS=Homo sapiens GN=RPL38 PE=1 | 112,876 | 35,71 | 8218 | 2 | 24,03 | 24,50 | 23,81 | 24,44 | 24,54 | 24,20 | **1,214** | 2,85E-01 | **0** | 0,28000069 | 0,54490762 |
| PTBP1_HUM | PTBP1 | sp\|P26599\|PTBP1_HUMAN Polypyrimidine tract-binding protein 1 OS=Homo sapiens GN | 550,409 | 35,59 | 57221 | 11 | 24,43 | 24,73 | 25,63 | 24,66 | 25,01 | 25,96 | **1,214** | 6,25E-01 | **0** | 0,28000005 | 0,20397953 |
| MAGD2_HUI | MAGED2 | sp\|Q9UNF1\|MAGD2_HUMAN Melanoma-associated antigen D2 OS=Homo sapiens GN=I | 62,907 | 8,75 | 64954 | 5 | 23,26 | 23,24 | 22,15 | 23,43 | 23,32 | 22,74 | **1,214** | 5,46E-01 | **0** | 0,28000005 | 0,26308781 |
| PTBP2_HUM | PTBP2 | sp\|Q9UKA9\|PTBP2_HUMAN Polypyrimidine tract-binding protein 2 OS=Homo sapiens G | 23,88 | 1,51 | 57491 | 1 | 23,43 | 23,58 | 23,34 | 23,86 | 23,92 | 23,41 | **1,214** | 1,86E-01 | **0** | 0,28000005 | 0,7308533 |
| PP2AA_HUM | PPP2CA | sp\|P67775\|PP2AA_HUMAN Serine/threonine-protein phosphatase 2A catalytic subunit | 497,019 | 60,84 | 35594 | 11 | 26,25 | 26,57 | 26,15 | 26,42 | 26,91 | 26,48 | **1,214** | 2,33E-01 | **0** | 0,28000005 | 0,63189896 |
| AP1M1_HUN | AP1M1 | sp\|Q9BXS5\|AP1M1_HUMAN AP-1 complex subunit mu-1 OS=Homo sapiens GN=AP1M1 | 77,296 | 16,08 | 48587 | 5 | 23,69 | 23,43 | 22,45 | 23,74 | 23,78 | 22,89 | **1,214** | 5,88E-01 | **0** | 0,27999942 | 0,23052309 |
| ELP3_HUMA | ELP3 | sp\|Q9H9T3\|ELP3_HUMAN Elongator complex protein 3 OS=Homo sapiens GN=ELP3 PE= | 11,3 | 1,83 | 62259 | 1 | 19,98 | 19,79 | 21,03 | 20,18 | 20,40 | 21,05 | **1,212** | 5,84E-01 | **0** | 0,27688344 | 0,23341086 |
| SAR1B_HUM | SAR1B | sp\|Q9Y6B6\|SAR1B_HUMAN GTP-binding protein SAR1b OS=Homo sapiens GN=SAR1B P | 247,286 | 56,06 | 22410 | 9 | 22,80 | 23,42 | 24,63 | 23,18 | 23,63 | 24,87 | **1,211** | 7,27E-01 | **0** | 0,27666728 | 0,13867935 |
| HSP7E_HUM | HSPA14 | sp\|Q0VDF9\|HSP7E_HUMAN Heat shock 70 kDa protein 14 OS=Homo sapiens GN=HSPA1 | 48,511 | 8,64 | 54794 | 4 | 22,14 | 21,88 | 23,30 | 22,23 | 22,16 | 23,76 | **1,211** | 7,05E-01 | **0** | 0,27666728 | 0,15172466 |
| SC23A_HUM | SEC23A | sp\|Q15436\|SC23A_HUMAN Protein transport protein Sec23A OS=Homo sapiens GN=SE | 676,961 | 28,5 | 86161 | 17 | 26,22 | 26,07 | 24,87 | 26,35 | 26,22 | 25,42 | **1,211** | 6,21E-01 | **0** | 0,27666664 | 0,20707299 |
| AIMP2_HUM | AIMP2 | sp\|Q13155\|AIMP2_HUMAN Aminoacyl tRNA synthase complex-interacting multifunctio | 128,88 | 10,94 | 35349 | 3 | 21,79 | 21,77 | 24,18 | 21,62 | 22,49 | 24,46 | **1,211** | 8,23E-01 | **0** | 0,27666601 | 0,08448347 |
| H2B1B_HUM | HIST1H2BB | sp\|P33778\|H2B1B_HUMAN Histone H2B type 1-B OS=Homo sapiens GN=HIST1H2BB PE= | 465,055 | 35,71 | 13950 | 8 | 23,22 | 22,94 | 24,37 | 23,80 | 23,57 | 23,99 | **1,211** | 5,75E-01 | **0** | 0,27666601 | 0,24015611 |
| MCM3_HUM | MCM3 | sp\|P25205\|MCM3_HUMAN DNA replication licensing factor MCM3 OS=Homo sapiens G | 518,951 | 20,79 | 90981 | 15 | 24,51 | 24,96 | 26,36 | 25,23 | 25,21 | 26,22 | **1,211** | 6,92E-01 | **0** | 0,27666601 | 0,15995205 |
| DNM1L_HU | DNM1L | sp\|O00429\|DNM1L_HUMAN Dynamin-1-like protein OS=Homo sapiens GN=DNM1L PE= | 991,987 | 46,06 | 81877 | 22 | 25,27 | 25,62 | 26,40 | 25,40 | 26,06 | 26,66 | **1,211** | 6,05E-01 | **0** | 0,27666601 | 0,21805936 |
| PLAP_HUMA | PLAA | sp\|Q9Y263\|PLAP_HUMAN Phospholipase A-2-activating protein OS=Homo sapiens GN=I | 239,753 | 14,09 | 87157 | 9 | 22,36 | 22,62 | 24,08 | 22,63 | 22,89 | 24,37 | **1,211** | 7,35E-01 | **0** | 0,27666601 | 0,13381974 |
| PP14B_HUM | PPP1R14B | sp\|Q96C90\|PP14B_HUMAN Protein phosphatase 1 regulatory subunit 14B OS=Homo sa | 88,925 | 25,85 | 15911 | 2 | 21,92 | 21,65 | 18,47 | 21,69 | 21,46 | 19,71 | **1,210** | 8,40E-01 | **0** | 0,27460607 | 0,07583677 |
| INT2_HUMA | INTS2 | sp\|Q9H0H0\|INT2_HUMAN Integrator complex subunit 2 OS=Homo sapiens GN=INTS2 PI | 40,269 | 1 | 134323 | 1 | 18,05 | 17,75 | 20,43 | 18,09 | 18,30 | 20,66 | **1,209** | 8,28E-01 | **0** | 0,27353795 | 0,08179093 |
| RS7_HUMAN | RPS7 | sp\|P62081\|RS7_HUMAN 40S ribosomal protein S7 OS=Homo sapiens GN=RPS7 PE=1 SV= | 489,985 | 61,86 | 22127 | 14 | 26,92 | 26,97 | 27,48 | 27,03 | 27,18 | 27,98 | **1,209** | 4,72E-01 | **0** | 0,27333387 | 0,32567275 |
| 5NTC_HUMA | NT5C2 | sp\|P49902\|5NTC_HUMAN Cytosolic purine 5'-nucleotidase OS=Homo sapiens GN=NT5C | 473,839 | 33,16 | 64970 | 14 | 25,10 | 25,14 | 25,07 | 25,25 | 25,60 | 25,28 | **1,209** | 7,42E-02 | **0** | 0,27333387 | 1,12934377 |
| ZFR2_HUMA | ZFR2 | sp\|Q9UPR6\|ZFR2_HUMAN Zinc finger RNA-binding protein 2 OS=Homo sapiens GN=ZFR | 40,006 | 1,28 | 101330 | 1 | 21,90 | 21,90 | 22,49 | 22,43 | 22,07 | 22,61 | **1,209** | 3,40E-01 | **0** | 0,27333387 | 0,46810198 |
| TTC7B_HUM | TTC7B | sp\|Q86TV6\|TTC7B_HUMAN Tetratricopeptide repeat protein 7B OS=Homo sapiens GN= | 77,741 | 6,41 | 94179 | 5 | 22,39 | 22,82 | 20,71 | 22,75 | 23,23 | 20,76 | **1,209** | 7,97E-01 | **0** | 0,27333387 | 0,09867228 |
| IDHC_HUMA | IDH1 | sp\|O75874\|IDHC_HUMAN Isocitrate dehydrogenase [NADP] cytoplasmic OS=Homo sapi | 315,865 | 31,64 | 46659 | 11 | 25,63 | 25,49 | 25,58 | 25,68 | 25,90 | 25,94 | **1,209** | 3,93E-02 | **0** | 0,27333387 | 1,40559632 |
| EPHA1_HUM | EPHA1 | sp\|P21709\|EPHA1_HUMAN Ephrin type-A receptor 1 OS=Homo sapiens GN=EPHA1 PE= | 48,789 | 1,02 | 108127 | 1 | 23,63 | 24,02 | 23,77 | 23,94 | 24,45 | 23,85 | **1,209** | 2,80E-01 | **0** | 0,27333387 | 0,55307205 |
| G6PD_HUM | G6PD | sp\|P11413\|G6PD_HUMAN Glucose-6-phosphate 1-dehydrogenase OS=Homo sapiens G | 192,556 | 13,79 | 59257 | 7 | 24,85 | 25,05 | 24,10 | 24,94 | 25,17 | 24,71 | **1,209** | 4,39E-01 | **0** | 0,27333323 | 0,3577756 |
| PRS6B_HUM | PSMC4 | sp\|P43686\|PRS6B_HUMAN 26S proteasome regulatory subunit 6B OS=Homo sapiens GN | 764,762 | 67,7 | 47366 | 21 | 26,08 | 26,51 | 27,35 | 26,38 | 26,54 | 27,84 | **1,209** | 6,69E-01 | **0** | 0,27333323 | 0,17438229 |
| CCS_HUMAN | CCS | sp\|O14618\|CCS_HUMAN Copper chaperone for superoxide dismutase OS=Homo sapier | 22,053 | 6,2 | 29041 | 2 | 21,83 | 22,60 | 21,05 | 22,23 | 22,34 | 21,73 | **1,209** | 6,03E-01 | **0** | 0,27333323 | 0,21946491 |
| SMC3_HUM | SMC3 | sp\|Q9UQE7\|SMC3_HUMAN Structural maintenance of chromosomes protein 3 OS=Hon | 730,105 | 24,65 | 141542 | 24 | 24,77 | 24,80 | 26,08 | 25,13 | 25,13 | 26,21 | **1,209** | 6,52E-01 | **0** | 0,2733326 | 0,18559131 |
| PRI2_HUMA | PRIM2 | sp\|P49643\|PRI2_HUMAN DNA primase large subunit OS=Homo sapiens GN=PRIM2 PE=1 | 49,334 | 2,16 | 58806 | 1 | 18,88 | 19,23 | 21,49 | 19,47 | 19,62 | 21,32 | **1,206** | 8,02E-01 | **0** | 0,27043978 | 0,09573055 |
| PTGIS_HUM | PTGIS | sp\|Q16647\|PTGIS_HUMAN Prostacyclin synthase OS=Homo sapiens GN=PTGIS PE=1 SV= | 55,861 | 7,2 | 57104 | 3 | 24,14 | 24,22 | 24,57 | 24,40 | 24,50 | 24,84 | **1,206** | 2,23E-01 | **0** | 0,27000046 | 0,65102715 |
| RUXE_HUM | SNRPE | sp\|P62304\|RUXE_HUMAN Small nuclear ribonucleoprotein E OS=Homo sapiens GN=SNI | 126,082 | 39,13 | 10804 | 2 | 23,88 | 23,88 | 24,04 | 23,92 | 24,54 | 24,15 | **1,206** | 2,26E-01 | **0** | 0,27000046 | 0,64662758 |
| MPU1_HUM | MPDU1 | sp\|O75352\|MPU1_HUMAN Mannose-P-dolichol utilization defect 1 protein OS=Homo s | 45,218 | 4,05 | 26638 | 1 | 22,01 | 21,96 | 22,43 | 22,10 | 22,62 | 22,49 | **1,206** | 2,79E-01 | **0** | 0,27000046 | 0,55394386 |
| VATC1_HUM | ATP6V1C1 | sp\|P21283\|VATC1_HUMAN V-type proton ATPase subunit C 1 OS=Homo sapiens GN=AT | 115,721 | 14,14 | 43942 | 5 | 22,65 | 22,69 | 22,73 | 23,21 | 23,35 | 22,32 | **1,206** | 4,51E-01 | **0** | 0,26999982 | 0,34610596 |
| U2AF2_HUM | U2AF2 | sp\|P26368\|U2AF2_HUMAN Splicing factor U2AF 65 kDa subunit OS=Homo sapiens GN=I | 396,625 | 29,68 | 53501 | 10 | 25,25 | 25,27 | 25,34 | 25,28 | 25,65 | 25,74 | **1,206** | 1,33E-01 | **0** | 0,26999982 | 0,87684416 |
| RL18A_HUM | RPL18A | sp\|Q02543\|RL18A_HUMAN 60S ribosomal protein L18a OS=Homo sapiens GN=RPL18A F | 337,854 | 35,23 | 20762 | 11 | 26,09 | 26,13 | 27,14 | 26,32 | 26,21 | 27,64 | **1,206** | 6,62E-01 | **0** | 0,26999982 | 0,17887848 |
| ADK_HUMAN | ADK | sp\|P55263\|ADK_HUMAN Adenosine kinase OS=Homo sapiens GN=ADK PE=1 SV=2 | 463,273 | 35,64 | 40545 | 10 | 23,44 | 23,81 | 24,82 | 23,98 | 24,26 | 24,64 | **1,206** | 5,85E-01 | **0** | 0,26999982 | 0,23320439 |
| EIF3K_HUM | EIF3K | sp\|Q9UBQ5\|EIF3K_HUMAN Eukaryotic translation initiation factor 3 subunit K OS=Homo | 172,499 | 36,7 | 25060 | 6 | 23,93 | 24,01 | 24,77 | 24,23 | 24,31 | 24,98 | **1,206** | 4,93E-01 | **0** | 0,26999919 | 0,30738005 |
| GFRP_HUMA | GCHFR | sp\|P30047\|GFRP_HUMAN GTP cyclohydrolase 1 feedback regulatory protein OS=Homo | 33,294 | 21,43 | 9698 | 1 | 21,66 | 22,10 | 17,88 | 21,40 | 21,96 | 19,09 | **1,204** | 8,75E-01 | **0** | 0,26759338 | 0,05779524 |
| COR1A_HUN | CORO1A | sp\|P31146\|COR1A_HUMAN Coronin-1A OS=Homo sapiens GN=CORO1A PE=1 SV=4 | 17,365 | 1,95 | 51026 | 1 | 18,33 | 18,30 | 20,65 | 18,29 | 19,40 | 20,39 | **1,203** | 8,00E-01 | **0** | 0,26712926 | 0,09687699 |
| KAPCB_HUM | PRKACB | sp\|P22694\|KAPCB_HUMAN cAMP-dependent protein kinase catalytic subunit beta OS=H | 154,03 | 28,49 | 40623 | 12 | 18,29 | 19,67 | 18,42 | 19,69 | 19,56 | 17,93 | **1,203** | 7,30E-01 | **0** | 0,26670011 | 0,13697063 |
| SYRC_HUMA | RARS | sp\|P54136\|SYRC_HUMAN Arginine--tRNA ligase, cytoplasmic OS=Homo sapiens GN=RAF | 457,806 | 26,06 | 75379 | 15 | 25,30 | 25,64 | 26,60 | 25,70 | 25,98 | 26,66 | **1,203** | 6,10E-01 | **0** | 0,26666705 | 0,21475786 |
| ANXA5_HUN | ANXA5 | sp\|P08758\|ANXA5_HUMAN Annexin A5 OS=Homo sapiens GN=ANXA5 PE=1 SV=2 | 1138,267 | 65,62 | 35937 | 25 | 29,75 | 29,67 | 29,30 | 30,06 | 29,86 | 29,60 | **1,203** | 2,38E-01 | **0** | 0,26666705 | 0,62411724 |
| PON2_HUM | PON2 | sp\|Q15165\|PON2_HUMAN Serum paraoxonase/arylesterase 2 OS=Homo sapiens GN=P | 69,582 | 9,32 | 39397 | 2 | 21,05 | 20,83 | 22,41 | 21,34 | 21,48 | 22,27 | **1,203** | 6,66E-01 | **0** | 0,26666705 | 0,17673238 |
| CP20A_HUM | CYP20A1 | sp\|Q6UW02\|CP20A_HUMAN Cytochrome P450 20A1 OS=Homo sapiens GN=CYP20A1 PI | 78,929 | 13,42 | 52432 | 5 | 22,26 | 22,97 | 24,65 | 22,94 | 23,69 | 24,05 | **1,203** | 7,50E-01 | **0** | 0,26666705 | 0,12506988 |
| LMNB1_HUN | LMNB1 | sp\|P20700\|LMNB1_HUMAN Lamin-B1 OS=Homo sapiens GN=LMNB1 PE=1 SV=2 | 557,596 | 31,57 | 66408 | 19 | 24,71 | 24,58 | 26,52 | 25,05 | 24,94 | 26,62 | **1,203** | 7,64E-01 | **0** | 0,26666705 | 0,11709015 |
| IPO9_HUMA | IPO9 | sp\|Q96P70\|IPO9_HUMAN Importin-9 OS=Homo sapiens GN=IPO9 PE=1 SV=3 | 593,358 | 24,69 | 115963 | 15 | 24,10 | 24,32 | 24,01 | 24,01 | 24,51 | 24,71 | **1,203** | 3,06E-01 | **0** | 0,26666641 | 0,51370678 |
| RL13_HUMA | RPL13 | sp\|P26373\|RL13_HUMAN 60S ribosomal protein L13 OS=Homo sapiens GN=RPL13 PE=1 | 125,024 | 23,22 | 24261 | 6 | 26,57 | 26,59 | 26,25 | 26,71 | 26,31 | 27,19 | **1,203** | 3,91E-01 | **0** | 0,26666641 | 0,4083344 |
| RS11_HUMA | RPS11 | sp\|P62280\|RS11_HUMAN 40S ribosomal protein S11 OS=Homo sapiens GN=RPS11 PE=1 | 406,971 | 62,66 | 18431 | 16 | 27,70 | 27,70 | 28,28 | 27,95 | 27,84 | 28,69 | **1,203** | 4,64E-01 | **0** | 0,26666641 | 0,33366121 |
| RS5_HUMAN | RPS5 | sp\|P46782\|RS5_HUMAN 40S ribosomal protein S5 OS=Homo sapiens GN=RPS5 PE=1 SV= | 567,969 | 61,76 | 22876 | 14 | 26,18 | 26,32 | 26,83 | 26,59 | 26,49 | 27,05 | **1,203** | 3,67E-01 | **0** | 0,26666641 | 0,43576514 |
| FAAH1_HUM | FAAH | sp\|O00519\|FAAH1_HUMAN Fatty-acid amide hydrolase 1 OS=Homo sapiens GN=FAAH F | 58,893 | 10,71 | 63066 | 5 | 23,53 | 23,44 | 23,45 | 23,88 | 24,10 | 23,24 | **1,203** | 3,62E-01 | **0** | 0,26666578 | 0,44102487 |
| DYN1_HUM | DNM1 | sp\|Q05193\|DYN1_HUMAN Dynamin-1 OS=Homo sapiens GN=DNM1 PE=1 SV=2 | 72,316 | 6,13 | 97408 | 6 | 22,54 | 22,61 | 19,90 | 22,04 | 22,27 | 21,53 | **1,201** | 7,88E-01 | **0** | 0,26425934 | 0,10346328 |
| ARFG2_HUM | ARFGAP2 | sp\|Q8N6H7\|ARFG2_HUMAN ADP-ribosylation factor GTPase-activating protein 2 OS=Ho | 20,586 | 1,92 | 56720 | 1 | 20,90 | 20,89 | 19,34 | 21,12 | 20,86 | 19,94 | **1,201** | 6,96E-01 | **0** | 0,26402473 | 0,15743867 |
| PSME2_HUN | PSME2 | sp\|Q9UL46\|PSME2_HUMAN Proteasome activator complex subunit 2 OS=Homo sapiens | 121,097 | 15,48 | 27402 | 3 | 23,80 | 23,54 | 23,07 | 23,59 | 24,07 | 23,54 | **1,200** | 3,88E-01 | **0** | 0,26333364 | 0,41076613 |
| TNPO1_HUN | TNPO1 | sp\|Q92973\|TNPO1_HUMAN Transportin-1 OS=Homo sapiens GN=TNPO1 PE=1 SV=2 | 602,379 | 27,95 | 102355 | 18 | 25,72 | 25,63 | 26,04 | 25,74 | 25,97 | 26,47 | **1,200** | 3,50E-01 | **0** | 0,263333 | 0,45644944 |
| STK39_HUM | STK39 | sp\|Q9UEW8\|STK39_HUMAN STE20/SPS1-related proline-alanine-rich protein kinase OS | 215,222 | 18,53 | 59474 | 6 | 19,24 | 20,09 | 22,03 | 19,03 | 20,96 | 22,16 | **1,200** | 8,41E-01 | **0** | 0,263333 | 0,07523211 |
| SKP1_HUMA | SKP1 | sp\|P63208\|SKP1_HUMAN S-phase kinase-associated protein 1 OS=Homo sapiens GN=SK | 237,485 | 35,58 | 18658 | 7 | 24,35 | 24,10 | 23,97 | 24,60 | 24,73 | 23,88 | **1,200** | 4,11E-01 | **0** | 0,263333 | 0,38655004 |
| EXOC6_HUM | EXOC6 | sp\|Q8TAG9\|EXOC6_HUMAN Exocyst complex component 6 OS=Homo sapiens GN=EXO | 9,692 | 1,24 | 93722 | 1 | 18,07 | 18,62 | 20,02 | 18,21 | 19,33 | 19,95 | **1,200** | 7,51E-01 | **0** | 0,26277479 | 0,12456363 |
| PDCD5_HUM | PDCD5 | sp\|O14737\|PDCD5_HUMAN Programmed cell death protein 5 OS=Homo sapiens GN=PD | 68,106 | 19,2 | 14285 | 2 | 22,55 | 23,07 | 17,63 | 22,64 | 22,67 | 18,72 | **1,198** | 9,10E-01 | **0** | 0,26118215 | 0,04090632 |
| TM9S1_HUN | TM9SF1 | sp\|O15321\|TM9S1_HUMAN Transmembrane 9 superfamily member 1 OS=Homo sapien | 51,45 | 3,63 | 68861 | 2 | 22,05 | 22,01 | 22,30 | 22,45 | 21,98 | 22,71 | **1,197** | 3,25E-01 | **0** | 0,26000023 | 0,48768376 |
| CFA20_HUM | CFAP20 | sp\|Q9Y6A4\|CFA20_HUMAN Cilia- and flagella-associated protein 20 OS=Homo sapiens G | 35,631 | 11,92 | 22774 | 2 | 21,10 | 20,91 | 20,40 | 21,25 | 21,61 | 20,33 | **1,197** | 5,82E-01 | **0** | 0,26000023 | 0,23512013 |
| SPT5H_HUM | SUPT5H | sp\|O00267\|SPT5H_HUMAN Transcription elongation factor SPT5 OS=Homo sapiens GN= | 204,912 | 7,73 | 121000 | 6 | 22,00 | 22,51 | 24,10 | 22,51 | 22,52 | 24,36 | **1,197** | 7,83E-01 | **0** | 0,26000023 | 0,10631983 |
| DDB1_HUM | DDB1 | sp\|Q16531\|DDB1_HUMAN DNA damage-binding protein 1 OS=Homo sapiens GN=DDB1 | 713,93 | 31,67 | 126968 | 28 | 26,46 | 26,37 | 26,49 | 26,57 | 26,70 | 26,83 | **1,197** | 3,54E-02 | **0** | 0,26000023 | 1,45049761 |
| P53_HUMAN | TP53 | sp\|P04637\|P53_HUMAN Cellular tumor antigen p53 OS=Homo sapiens GN=TP53 PE=1 S | 74,591 | 4,83 | 43653 | 1 | 21,07 | 21,84 | 23,49 | 21,43 | 21,98 | 23,77 | **1,197** | 8,08E-01 | **0** | 0,26000023 | 0,09232362 |
| UBE2K_HUM | UBE2K | sp\|P61086\|UBE2K_HUMAN Ubiquitin-conjugating enzyme E2 K OS=Homo sapiens GN=U | 439,813 | 72 | 22407 | 11 | 26,48 | 26,25 | 24,84 | 26,35 | 25,75 | 26,25 | **1,197** | 6,58E-01 | **0** | 0,26000023 | 0,18155507 |
| SRS10_HUM | SRSF10 | sp\|O75494\|SRS10_HUMAN Serine/arginine-rich splicing factor 10 OS=Homo sapiens GN | 39,444 | 6,11 | 31301 | 1 | 21,18 | 21,23 | 22,45 | 21,40 | 21,69 | 22,55 | **1,197** | 6,55E-01 | **0** | 0,25999959 | 0,18351576 |
| AFAD_HUM | AFDN | sp\|P55196\|AFAD_HUMAN Afadin OS=Homo sapiens GN=AFDN PE=1 SV=3 | 112,374 | 3,34 | 206804 | 5 | 23,70 | 23,59 | 20,58 | 23,40 | 23,26 | 21,99 | **1,197** | 8,27E-01 | **0** | 0,25999959 | 0,08236267 |
| RS24_HUMA | RPS24 | sp\|P62847\|RS24_HUMAN 40S ribosomal protein S24 OS=Homo sapiens GN=RPS24 PE=1 | 246,153 | 35,34 | 15423 | 5 | 24,96 | 25,24 | 25,17 | 25,30 | 25,14 | 25,71 | **1,197** | 2,42E-01 | **0** | 0,25999959 | 0,61644815 |
| DTD1_HUM | DTD1 | sp\|Q8TEA8\|DTD1_HUMAN D-aminoacyl-tRNA deacylase 1 OS=Homo sapiens GN=DTD1 | 93,944 | 14,83 | 23424 | 2 | 20,05 | 19,50 | 20,14 | 19,27 | 19,84 | 21,35 | **1,195** | 7,14E-01 | **0** | 0,25666746 | 0,14635732 |
| DCTN4_HUN | DCTN4 | sp\|Q9UJW0\|DCTN4_HUMAN Dynactin subunit 4 OS=Homo sapiens GN=DCTN4 PE=1 SV | 71,31 | 12,39 | 52337 | 4 | 22,28 | 23,06 | 20,76 | 22,37 | 22,69 | 21,81 | **1,195** | 7,40E-01 | **0** | 0,25666682 | 0,1305424 |
| RL12_HUMA | RPL12 | sp\|P30050\|RL12_HUMAN 60S ribosomal protein L12 OS=Homo sapiens GN=RPL12 PE=1 | 356,555 | 54,55 | 17819 | 6 | 26,39 | 26,44 | 26,87 | 26,61 | 26,67 | 27,19 | **1,195** | 3,43E-01 | **0** | 0,25666682 | 0,4642871 |
| RAB6A_HUM | RAB6A | sp\|P20340\|RAB6A_HUMAN Ras-related protein Rab-6A OS=Homo sapiens GN=RAB6A PI | 390,142 | 55,29 | 23593 | 14 | 25,33 | 25,32 | 24,59 | 25,50 | 25,49 | 25,02 | **1,195** | 4,29E-01 | **0** | 0,25666682 | 0,36788995 |
| GALE_HUMA | GALE | sp\|Q14376\|GALE_HUMAN UDP-glucose 4-epimerase OS=Homo sapiens GN=GALE PE=1 | 194,857 | 18,39 | 38282 | 4 | 23,23 | 23,52 | 22,81 | 23,25 | 23,85 | 23,23 | **1,195** | 4,26E-01 | **0** | 0,25666682 | 0,37109539 |
| SRSF2_HUM | SRSF2 | sp\|Q01130\|SRSF2_HUMAN Serine/arginine-rich splicing factor 2 OS=Homo sapiens GN= | 24,43 | 6,79 | 25476 | 2 | 19,40 | 22,77 | 21,31 | 19,54 | 22,32 | 22,39 | **1,195** | 8,59E-01 | **0** | 0,25666682 | 0,06607379 |
| PRP4B_HUM | PRPF4B | sp\|Q13523\|PRP4B_HUMAN Serine/threonine-protein kinase PRP4 homolog OS=Homo s | 21,125 | 1,39 | 116987 | 1 | 19,17 | 19,59 | 19,72 | 19,67 | 19,63 | 19,95 | **1,194** | 2,59E-01 | **0** | 0,25534058 | 0,58625739 |
| FAT2_HUMA | FAT2 | sp\|Q9NYQ8\|FAT2_HUMAN Protocadherin Fat 2 OS=Homo sapiens GN=FAT2 PE=1 SV=2 | 41,928 | 0,28 | 479317 | 1 | 21,66 | 18,83 | 19,04 | 21,68 | 20,51 | 18,11 | **1,193** | 8,64E-01 | **0** | 0,25459735 | 0,06363873 |
| PSA_HUMAN | NPEPPS | sp\|P55786\|PSA_HUMAN Puromycin-sensitive aminopeptidase OS=Homo sapiens GN=NI | 538,562 | 26,22 | 103276 | 22 | 26,51 | 26,71 | 25,18 | 26,58 | 26,83 | 25,75 | **1,192** | 6,85E-01 | **0** | 0,25333341 | 0,16423878 |
| H15_HUMAN | HIST1H1B | sp\|P16401\|H15_HUMAN Histone H1.5 OS=Homo sapiens GN=HIST1H1B PE=1 SV=3 | 21,847 | 4,87 | 22580 | 1 | 22,33 | 22,30 | 22,40 | 22,94 | 22,46 | 22,39 | **1,192** | 2,22E-01 | **0** | 0,25333341 | 0,65346734 |
| AN32B_HUM | ANP32B | sp\|Q92688\|AN32B_HUMAN Acidic leucine-rich nuclear phosphoprotein 32 family memb | 93,608 | 19,12 | 28788 | 5 | 24,66 | 24,91 | 23,47 | 25,16 | 25,16 | 23,48 | **1,192** | 7,41E-01 | **0** | 0,25333341 | 0,13022485 |
| SPD2A_HUM | SH3PXD2A | sp\|Q5TCZ1\|SPD2A_HUMAN SH3 and PX domain-containing protein 2A OS=Homo sapien | 23,713 | 1,94 | 125289 | 1 | 17,10 | 18,96 | 19,77 | 18,85 | 18,78 | 18,96 | **1,191** | 7,66E-01 | **0** | 0,25203896 | 0,11561038 |
| MGAT3_HUN | MGAT3 | sp\|Q09327\|MGAT3_HUMAN Beta-1,4-mannosyl-glycoprotein 4-beta-N-acetylglucosami | 18,265 | 2,25 | 61313 | 1 | 21,44 | 21,58 | 20,37 | 21,63 | 21,35 | 21,16 | **1,189** | 5,71E-01 | **0** | 0,25000636 | 0,24323549 |
| RS8_HUMAN | RPS8 | sp\|P62241\|RS8_HUMAN 40S ribosomal protein S8 OS=Homo sapiens GN=RPS8 PE=1 SV= | 481,049 | 52,88 | 24205 | 12 | 26,96 | 27,32 | 27,57 | 27,34 | 27,22 | 28,04 | **1,189** | 4,67E-01 | **0** | 0,25000064 | 0,33111501 |
| MTMR1_HU | MTMR1 | sp\|Q13613\|MTMR1_HUMAN Myotubularin-related protein 1 OS=Homo sapiens GN=MT | 14,414 | 1,05 | 74678 | 1 | 20,96 | 20,76 | 21,07 | 21,06 | 21,15 | 21,33 | **1,189** | 1,07E-01 | **0** | 0,25 | 0,9715305 |
| CNDP2_HUN | CNDP2 | sp\|Q96KP4\|CNDP2_HUMAN Cytosolic non-specific dipeptidase OS=Homo sapiens GN=C | 64,797 | 6,53 | 52878 | 2 | 22,01 | 21,92 | 21,05 | 22,04 | 22,39 | 21,30 | **1,189** | 6,03E-01 | **0** | 0,25 | 0,21948353 |

| | | | | | | | | | | | | | | | | | |
|---|---|---|---|---|---|---|---|---|---|---|---|---|---|---|---|---|---|
| ALDR_HUMA | AKR1B1 | sp\|P15121\|ALDR_HUMAN Aldose reductase OS=Homo sapiens GN=AKR1B1 PE=1 SV=3 | 267,276 | 31,65 | 35853 | 9 | 26,44 | 26,46 | 24,95 | 26,49 | 26,52 | 25,59 | 1,189 | 6,92E-01 | 0 | 0,25 | 0,1602052 |
| MYO5A_HUM | MYO5A | sp\|Q9Y4I1\|MYO5A_HUMAN Unconventional myosin-Va OS=Homo sapiens GN=MYO5A | 379,135 | 7,12 | 215405 | 10 | 23,21 | 23,86 | 21,87 | 23,35 | 23,79 | 22,55 | 1,189 | 7,35E-01 | 0 | 0,25 | 0,13362614 |
| E41L5_HUM | EPB41L5 | sp\|Q9HCM4\|E41L5_HUMAN Band 4.1-like protein 5 OS=Homo sapiens GN=EPB41L5 PE= | 32,938 | 1,09 | 81856 | 1 | 20,70 | 20,68 | 17,65 | 20,52 | 20,39 | 18,87 | 1,188 | 8,38E-01 | 0 | 0,24884605 | 0,07652676 |
| NAMPT_HUM | NAMPT | sp\|P43490\|NAMPT_HUMAN Nicotinamide phosphoribosyltransferase OS=Homo sapien | 24,294 | 2,44 | 55521 | 1 | 21,12 | 21,16 | 20,07 | 20,95 | 21,47 | 20,67 | 1,187 | 5,94E-01 | 0 | 0,24765714 | 0,22646282 |
| RUVB1_HUM | RUVBL1 | sp\|Q9Y265\|RUVB1_HUMAN RuvB-like 1 OS=Homo sapiens GN=RUVBL1 PE=1 SV=1 | 934,672 | 62,94 | 50228 | 21 | 26,63 | 26,71 | 27,81 | 26,79 | 27,08 | 28,02 | 1,186 | 6,67E-01 | 0 | 0,24666786 | 0,17597383 |
| RL27A_HUM | RPL27A | sp\|P46776\|RL27A_HUMAN 60S ribosomal protein L27a OS=Homo sapiens GN=RPL27A P | 223,022 | 31,76 | 16561 | 7 | 26,25 | 26,67 | 26,48 | 26,64 | 26,55 | 26,95 | 1,186 | 2,24E-01 | 0 | 0,24666659 | 0,65013096 |
| GNPI1_HUM | GNPDA1 | sp\|P46926\|GNPI1_HUMAN Glucosamine-6-phosphate isomerase 1 OS=Homo sapiens G | 164,617 | 30,1 | 32669 | 6 | 21,27 | 22,71 | 21,61 | 21,83 | 22,47 | 22,03 | 1,186 | 6,30E-01 | 0 | 0,24666659 | 0,20052682 |
| MK01_HUM | MAPK1 | sp\|P28482\|MK01_HUMAN Mitogen-activated protein kinase 1 OS=Homo sapiens GN=M | 285,983 | 39,17 | 41390 | 12 | 24,87 | 24,60 | 23,55 | 24,91 | 24,63 | 24,22 | 1,186 | 6,13E-01 | 0 | 0,24666595 | 0,212873 |
| LASP1_HUM | LASP1 | sp\|Q14847\|LASP1_HUMAN LIM and SH3 domain protein 1 OS=Homo sapiens GN=LASP1 | 270,28 | 39,85 | 29717 | 11 | 26,10 | 25,25 | 24,66 | 26,40 | 25,89 | 24,46 | 1,186 | 7,48E-01 | 0 | 0,24666595 | 0,12631843 |
| ACTY_HUMA | ACTR1B | sp\|P42025\|ACTY_HUMAN Beta-centractin OS=Homo sapiens GN=ACTR1B PE=1 SV=1 | 332,694 | 28,46 | 42293 | 10 | 21,95 | 21,86 | 23,49 | 22,11 | 22,22 | 23,71 | 1,186 | 7,55E-01 | 0 | 0,24666595 | 0,12188545 |
| ROA0_HUM | HNRNPA0 | sp\|Q13151\|ROA0_HUMAN Heterogeneous nuclear ribonucleoprotein A0 OS=Homo sap | 35,78 | 2,62 | 30841 | 2 | 24,65 | 24,42 | 24,99 | 24,84 | 24,93 | 25,02 | 1,184 | 2,33E-01 | 0 | 0,24333382 | 0,63177543 |
| MCMBP_HU | MCMBP | sp\|Q9BTE3\|MCMBP_HUMAN Mini-chromosome maintenance complex-binding protein | 72,347 | 6,23 | 72980 | 3 | 21,08 | 21,85 | 22,51 | 21,44 | 21,94 | 22,79 | 1,184 | 6,92E-01 | 0 | 0,24333382 | 0,15994453 |
| KCRB_HUMA | CKB | sp\|P12277\|KCRB_HUMAN Creatine kinase B-type OS=Homo sapiens GN=CKB PE=1 SV=1 | 962,112 | 47,77 | 42644 | 19 | 27,79 | 27,98 | 26,16 | 28,04 | 28,21 | 26,41 | 1,184 | 7,80E-01 | 0 | 0,24333318 | 0,10795172 |
| RS19_HUMA | RPS19 | sp\|P39019\|RS19_HUMAN 40S ribosomal protein S19 OS=Homo sapiens GN=RPS19 PE=1 | 187,612 | 42,76 | 16060 | 7 | 26,55 | 26,75 | 25,55 | 26,83 | 26,84 | 25,90 | 1,181 | 6,46E-01 | 0 | 0,24000041 | 0,18946452 |
| MTPN_HUM | MTPN | sp\|P58546\|MTPN_HUMAN Myotrophin OS=Homo sapiens GN=MTPN PE=1 SV=2 | 31,306 | 8,47 | 12895 | 1 | 21,21 | 21,18 | 22,14 | 21,52 | 21,59 | 22,14 | 1,181 | 5,53E-01 | 0 | 0,24000041 | 0,25721854 |
| ESTD_HUMA | ESD | sp\|P10768\|ESTD_HUMAN S-formylglutathione hydrolase OS=Homo sapiens GN=ESD PE | 28,871 | 8,51 | 31463 | 3 | 24,09 | 24,38 | 21,31 | 24,55 | 24,17 | 21,78 | 1,181 | 8,63E-01 | 0 | 0,24000041 | 0,06384888 |
| NF1_HUMAN | NF1 | sp\|P21359\|NF1_HUMAN Neurofibromin OS=Homo sapiens GN=NF1 PE=1 SV=2 | 112,443 | 2,15 | 319372 | 4 | 20,67 | 20,83 | 21,63 | 20,96 | 21,09 | 21,80 | 1,181 | 5,77E-01 | 0 | 0,23999977 | 0,23911877 |
| DDX46_HUM | DDX46 | sp\|Q7L014\|DDX46_HUMAN Probable ATP-dependent RNA helicase DDX46 OS=Homo sa | 177,712 | 7,86 | 117362 | 8 | 22,68 | 22,83 | 24,85 | 22,98 | 23,18 | 24,92 | 1,181 | 8,10E-01 | 0 | 0,23999977 | 0,09176401 |
| KISHA_HUM | TMEM167A | sp\|Q8TBQ9\|KISHA_HUMAN Protein kish-A OS=Homo sapiens GN=TMEM167A PE=1 SV= | 15,929 | 12,5 | 8060 | 1 | 20,46 | 21,12 | 19,63 | 21,00 | 21,48 | 19,45 | 1,181 | 7,65E-01 | 0 | 0,23952929 | 0,11628562 |
| RIC8A_HUM | RIC8A | sp\|Q9NPQ8\|RIC8A_HUMAN Synembryn-A OS=Homo sapiens GN=RIC8A PE=1 SV=3 | 54,935 | 4,33 | 59710 | 2 | 20,46 | 21,48 | 20,88 | 20,08 | 20,84 | 22,61 | 1,178 | 7,84E-01 | 0 | 0,23666763 | 0,10590711 |
| EZRI_HUMAN | EZR | sp\|P15311\|EZRI_HUMAN Ezrin OS=Homo sapiens GN=EZR PE=1 SV=4 | 965,397 | 40,61 | 69413 | 28 | 26,80 | 26,50 | 24,41 | 26,71 | 26,37 | 25,34 | 1,178 | 7,96E-01 | 0 | 0,236667 | 0,09901369 |
| IMPA2_HUM | IMPA2 | sp\|O14732\|IMPA2_HUMAN Inositol monophosphatase 2 OS=Homo sapiens GN=IMPA2 | 47,131 | 7,99 | 31321 | 1 | 18,59 | 19,24 | 18,73 | 19,17 | 18,95 | 19,15 | 1,178 | 3,22E-01 | 0 | 0,236667 | 0,4921052 |
| PRPS1_HUM | PRPS1 | sp\|P60891\|PRPS1_HUMAN Ribose-phosphate pyrophosphokinase 1 OS=Homo sapiens | 380,709 | 52,83 | 34834 | 13 | 24,86 | 24,91 | 25,13 | 25,07 | 25,17 | 25,37 | 1,178 | 1,22E-01 | 0 | 0,236667 | 0,9127152 |
| TTC7A_HUM | TTC7A | sp\|Q9ULT0\|TTC7A_HUMAN Tetratricopeptide repeat protein 7A OS=Homo sapiens GN= | 62,824 | 2,68 | 96185 | 2 | 18,76 | 21,65 | 19,66 | 20,44 | 20,96 | 19,38 | 1,178 | 8,20E-01 | 0 | 0,23666636 | 0,08637606 |
| EIF3H_HUM | EIF3H | sp\|O15372\|EIF3H_HUMAN Eukaryotic translation initiation factor 3 subunit H OS=Homo | 334,306 | 46,59 | 39930 | 13 | 24,92 | 24,83 | 25,80 | 25,22 | 25,07 | 25,97 | 1,178 | 6,00E-01 | 0 | 0,23666636 | 0,22182698 |
| RIR1_HUMA | RRM1 | sp\|P23921\|RIR1_HUMAN Ribonucleoside-diphosphate reductase large subunit OS=Hom | 909,962 | 37,63 | 90070 | 23 | 26,19 | 26,06 | 27,68 | 26,17 | 26,42 | 28,05 | 1,178 | 7,78E-01 | 0 | 0,23666636 | 0,10885835 |
| IDH3G_HUM | IDH3G | sp\|P51553\|IDH3G_HUMAN Isocitrate dehydrogenase [NAD] subunit gamma, mitochond | 33,185 | 11,7 | 42794 | 2 | 19,96 | 20,10 | 21,63 | 19,54 | 21,19 | 21,66 | 1,178 | 7,92E-01 | 0 | 0,23601151 | 0,10147277 |
| TMX1_HUM | TMX1 | sp\|Q9H3N1\|TMX1_HUMAN Thioredoxin-related transmembrane protein 1 OS=Homo sa | 67 | 7,5 | 31791 | 2 | 23,92 | 24,46 | 24,89 | 24,36 | 24,75 | 24,86 | 1,176 | 5,05E-01 | 0 | 0,23333422 | 0,29667902 |
| GNA14_HUM | GNA14 | sp\|O95837\|GNA14_HUMAN Guanine nucleotide-binding protein subunit alpha-14 OS=H | 337,96 | 29,86 | 41571 | 12 | 24,22 | 21,89 | 23,65 | 24,24 | 23,10 | 23,12 | 1,176 | 7,84E-01 | 0 | 0,23333422 | 0,10568102 |
| PRS6A_HUM | PSMC3 | sp\|P17980\|PRS6A_HUMAN 26S proteasome regulatory subunit 6A OS=Homo sapiens GN | 793,125 | 54,9 | 49204 | 23 | 26,49 | 26,69 | 27,46 | 26,78 | 26,86 | 27,70 | 1,176 | 6,06E-01 | 0 | 0,23333422 | 0,21770318 |
| SNX4_HUMA | SNX4 | sp\|O95219\|SNX4_HUMAN Sorting nexin-4 OS=Homo sapiens GN=SNX4 PE=1 SV=1 | 51,058 | 4,89 | 51909 | 2 | 21,42 | 21,56 | 21,67 | 21,45 | 21,85 | 22,05 | 1,176 | 2,88E-01 | 0 | 0,23333359 | 0,5404002 |
| NP1L1_HUM | NAP1L1 | sp\|P55209\|NP1L1_HUMAN Nucleosome assembly protein 1-like 1 OS=Homo sapiens GN | 548,897 | 43,73 | 45374 | 13 | 25,89 | 26,45 | 26,86 | 26,12 | 26,74 | 27,04 | 1,176 | 5,82E-01 | 0 | 0,23333359 | 0,23489418 |
| GSTM5_HUM | GSTM5 | sp\|P46439\|GSTM5_HUMAN Glutathione S-transferase Mu 5 OS=Homo sapiens GN=GST | 71,647 | 8,72 | 25675 | 2 | 23,66 | 23,97 | 23,45 | 23,94 | 24,18 | 23,66 | 1,176 | 3,35E-01 | 0 | 0,23333359 | 0,47503501 |
| NUCB1_HUM | NUCB1 | sp\|Q02818\|NUCB1_HUMAN Nucleobindin-1 OS=Homo sapiens GN=NUCB1 PE=1 SV=4 | 77,232 | 14,53 | 53879 | 6 | 23,87 | 23,50 | 22,55 | 24,15 | 23,74 | 22,73 | 1,176 | 7,06E-01 | 0 | 0,23333295 | 0,15090576 |
| EFGM_HUM | GFM1 | sp\|Q96RP9\|EFGM_HUMAN Elongation factor G, mitochondrial OS=Homo sapiens GN=G | 24,706 | 1,6 | 83471 | 1 | 17,61 | 20,10 | 21,36 | 19,28 | 19,70 | 20,78 | 1,174 | 8,55E-01 | 0 | 0,23113378 | 0,06780586 |
| PARK7_HUM | PARK7 | sp\|Q99497\|PARK7_HUMAN Protein/nucleic acid deglycase DJ-1 OS=Homo sapiens GN= | 342,579 | 45,5 | 19891 | 8 | 25,98 | 26,16 | 25,41 | 26,15 | 26,24 | 25,85 | 1,173 | 4,18E-01 | 0 | 0,23000018 | 0,37882732 |
| ILKAP_HUM | ILKAP | sp\|Q9H0C8\|ILKAP_HUMAN Integrin-linked kinase-associated serine/threonine phospha | 42,804 | 5,1 | 42907 | 2 | 20,32 | 20,66 | 21,83 | 20,51 | 20,46 | 22,53 | 1,173 | 7,93E-01 | 0 | 0,23000018 | 0,10058796 |
| RS25_HUMA | RPS25 | sp\|P62851\|RS25_HUMAN 40S ribosomal protein S25 OS=Homo sapiens GN=RPS25 PE=1 | 182,648 | 25,6 | 13742 | 5 | 26,59 | 26,83 | 26,47 | 27,00 | 26,90 | 26,68 | 1,173 | 1,80E-01 | 0 | 0,23000018 | 0,74389668 |
| APT_HUMAN | APRT | sp\|P07741\|APT_HUMAN Adenine phosphoribosyltransferase OS=Homo sapiens GN=AP | 73,71 | 11,11 | 19608 | 1 | 20,07 | 20,46 | 20,57 | 19,93 | 20,48 | 21,38 | 1,173 | 6,35E-01 | 0 | 0,23000018 | 0,19689794 |
| RL37A_HUM | RPL37A | sp\|P61513\|RL37A_HUMAN 60S ribosomal protein L37a OS=Homo sapiens GN=RPL37A P | 183,374 | 51,09 | 10275 | 4 | 23,86 | 23,65 | 24,73 | 24,03 | 24,14 | 24,76 | 1,173 | 5,97E-01 | 0 | 0,23000018 | 0,22394323 |
| SAE1_HUMA | SAE1 | sp\|Q9UBE0\|SAE1_HUMAN SUMO-activating enzyme subunit 1 OS=Homo sapiens GN=S | 86,456 | 13,29 | 38450 | 3 | 19,67 | 17,88 | 20,79 | 19,76 | 19,17 | 20,10 | 1,172 | 8,09E-01 | 0 | 0,22909419 | 0,09181499 |
| PLCB1_HUM | PLCB1 | sp\|Q9NQ66\|PLCB1_HUMAN 1-phosphatidylinositol 4,5-bisphosphate phosphodiesteras | 9,528 | 1,23 | 138567 | 1 | 17,71 | 20,10 | 19,51 | 18,61 | 19,70 | 19,70 | 1,172 | 7,91E-01 | 0 | 0,22898229 | 0,10206703 |
| SF3B3_HUM | SF3B3 | sp\|Q15393\|SF3B3_HUMAN Splicing factor 3B subunit 3 OS=Homo sapiens GN=SF3B3 PE | 1089,804 | 32,79 | 135577 | 29 | 26,35 | 26,00 | 27,55 | 26,41 | 26,58 | 27,59 | 1,170 | 7,23E-01 | 0 | 0,22666677 | 0,14068109 |
| SE1L1_HUM | SEL1L | sp\|Q9UBV2\|SE1L1_HUMAN Protein sel-1 homolog 1 OS=Homo sapiens GN=SEL1L PE=1 | 44,572 | 2,52 | 88755 | 1 | 21,06 | 20,52 | 21,44 | 20,93 | 21,17 | 21,60 | 1,170 | 5,31E-01 | 0 | 0,22666677 | 0,2746915 |
| G3BP1_HUM | G3BP1 | sp\|Q13283\|G3BP1_HUMAN Ras GTPase-activating protein-binding protein 1 OS=Homo | 344,708 | 28,54 | 52164 | 9 | 24,76 | 24,85 | 24,52 | 24,71 | 25,11 | 24,99 | 1,170 | 2,15E-01 | 0 | 0,22666613 | 0,66703854 |
| RS4X_HUMA | RPS4X | sp\|P62701\|RS4X_HUMAN 40S ribosomal protein S4, X isoform OS=Homo sapiens GN=R | 883,706 | 65,4 | 29598 | 24 | 28,44 | 28,39 | 29,10 | 28,64 | 28,48 | 29,49 | 1,170 | 5,90E-01 | 0 | 0,22666613 | 0,22878611 |
| CSK2B_HUM | CSNK2B | sp\|P67870\|CSK2B_HUMAN Casein kinase II subunit beta OS=Homo sapiens GN=CSNK2B | 260,07 | 49,3 | 24942 | 8 | 23,66 | 24,68 | 25,98 | 24,23 | 24,80 | 25,97 | 1,170 | 8,02E-01 | 0 | 0,22666613 | 0,0960205 |
| WASC5_HUM | WASHC5 | sp\|Q12768\|WASC5_HUMAN WASH complex subunit 5 OS=Homo sapiens GN=WASHC5 | 29,769 | 1,04 | 134286 | 1 | 19,20 | 18,46 | 18,99 | 19,44 | 19,04 | 18,85 | 1,170 | 4,65E-01 | 0 | 0,22619883 | 0,33275595 |
| SP16H_HUM | SUPT16H | sp\|Q9Y5B9\|SP16H_HUMAN FACT complex subunit SPT16 OS=Homo sapiens GN=SUPT16 | 411,962 | 16,81 | 119914 | 14 | 23,32 | 24,01 | 25,33 | 23,53 | 24,10 | 25,70 | 1,167 | 8,12E-01 | 0 | 0,22333399 | 0,09065895 |
| ERH_HUMAN | ERH | sp\|P84090\|ERH_HUMAN Enhancer of rudimentary homolog OS=Homo sapiens GN=ERH | 115,715 | 41,35 | 12259 | 3 | 23,50 | 24,46 | 22,41 | 24,45 | 23,85 | 22,74 | 1,167 | 7,88E-01 | 0 | 0,22333399 | 0,10364331 |
| GALT2_HUM | GALNT2 | sp\|Q10471\|GALT2_HUMAN Polypeptide N-acetylgalactosaminyltransferase 2 OS=Homo | 32,472 | 8,06 | 64733 | 2 | 21,13 | 20,81 | 21,04 | 20,69 | 21,54 | 21,42 | 1,167 | 4,73E-01 | 0 | 0,22333399 | 0,32518047 |
| SERA_HUMA | PHGDH | sp\|O43175\|SERA_HUMAN D-3-phosphoglycerate dehydrogenase OS=Homo sapiens GN | 475,049 | 28,52 | 56650 | 12 | 26,87 | 27,34 | 27,58 | 27,33 | 27,60 | 27,53 | 1,167 | 3,75E-01 | 0 | 0,22333336 | 0,42652347 |
| IF6_HUMAN | EIF6 | sp\|P56537\|IF6_HUMAN Eukaryotic translation initiation factor 6 OS=Homo sapiens GN= | 287,488 | 36,73 | 26599 | 4 | 24,50 | 24,65 | 24,86 | 24,47 | 24,92 | 25,29 | 1,167 | 4,37E-01 | 0 | 0,22333336 | 0,35926793 |
| WDR82_HUM | WDR82 | sp\|Q6UXN9\|WDR82_HUMAN WD repeat-containing protein 82 OS=Homo sapiens GN=V | 131,343 | 13,42 | 35079 | 3 | 21,34 | 20,78 | 21,95 | 20,63 | 21,53 | 22,58 | 1,167 | 7,51E-01 | 0 | 0,22333272 | 0,12435453 |
| OTUB1_HUM | OTUB1 | sp\|Q96FW1\|OTUB1_HUMAN Ubiquitin thioesterase OTUB1 OS=Homo sapiens GN=OTU | 468,683 | 46,86 | 31284 | 12 | 25,87 | 26,17 | 25,41 | 25,89 | 26,39 | 25,84 | 1,167 | 4,73E-01 | 0 | 0,22333272 | 0,32504627 |
| SMG1_HUM | SMG1 | sp\|Q96Q15\|SMG1_HUMAN Serine/threonine-protein kinase SMG1 OS=Homo sapiens G | 38,347 | 0,36 | 410501 | 1 | 17,02 | 17,01 | 20,09 | 16,98 | 17,73 | 20,08 | 1,167 | 8,80E-01 | 0 | 0,22333272 | 0,05559275 |
| RL9_HUMAN | RPL9 | sp\|P32969\|RL9_HUMAN 60S ribosomal protein L9 OS=Homo sapiens GN=RPL9 PE=1 SV= | 248,905 | 41,15 | 21863 | 8 | 25,05 | 25,11 | 26,70 | 25,22 | 25,15 | 27,16 | 1,167 | 8,06E-01 | 0 | 0,22333272 | 0,09358754 |
| WBP11_HUM | WBP11 | sp\|Q9Y2W2\|WBP11_HUMAN WW domain-binding protein 11 OS=Homo sapiens GN=W | 20,572 | 1,56 | 69998 | 1 | 19,32 | 19,06 | 19,71 | 19,97 | 19,10 | 19,68 | 1,165 | 5,27E-01 | 0 | 0,22000058 | 0,27815729 |
| GTR14_HUM | SLC2A14 | sp\|Q8TDB8\|GTR14_HUMAN Solute carrier family 2, facilitated glucose transporter mem | 36,459 | 3,08 | 56320 | 2 | 26,39 | 26,37 | 24,50 | 26,21 | 26,28 | 25,43 | 1,165 | 7,64E-01 | 0 | 0,21999995 | 0,11712525 |
| HOOK3_HUM | HOOK3 | sp\|Q86VS8\|HOOK3_HUMAN Protein Hook homolog 3 OS=Homo sapiens GN=HOOK3 PE | 87,497 | 6,13 | 83126 | 3 | 21,34 | 20,99 | 19,67 | 20,52 | 20,97 | 21,17 | 1,165 | 7,06E-01 | 0 | 0,21999995 | 0,1509535 |
| RBM8A_HUM | RBM8A | sp\|Q9Y5S9\|RBM8A_HUMAN RNA-binding protein 8A OS=Homo sapiens GN=RBM8A PE= | 318,905 | 50,57 | 19889 | 7 | 24,94 | 24,75 | 25,27 | 24,92 | 25,09 | 25,61 | 1,165 | 4,41E-01 | 0 | 0,21999995 | 0,35598387 |
| RL35_HUMA | RPL35 | sp\|P42766\|RL35_HUMAN 60S ribosomal protein L35 OS=Homo sapiens GN=RPL35 PE=1 | 54,221 | 26,02 | 14551 | 3 | 25,70 | 26,23 | 24,47 | 25,76 | 25,57 | 25,73 | 1,165 | 6,96E-01 | 0 | 0,21999995 | 0,1570792 |
| OSBL8_HUM | OSBPL8 | sp\|Q9BZF1\|OSBL8_HUMAN Oxysterol-binding protein-related protein 8 OS=Homo sapie | 79,785 | 3,37 | 101196 | 3 | 21,19 | 21,70 | 22,04 | 22,08 | 21,88 | 21,63 | 1,165 | 4,75E-01 | 0 | 0,21999868 | 0,32349132 |
| SEC20_HUM | BNIP1 | sp\|Q12981\|SEC20_HUMAN Vesicle transport protein SEC20 OS=Homo sapiens GN=BNIP | 14,853 | 4,39 | 26132 | 1 | 18,22 | 20,82 | 21,06 | 18,27 | 21,13 | 21,36 | 1,164 | 8,79E-01 | 0 | 0,2191569 | 0,05625264 |
| RS28_HUMA | RPS28 | sp\|P62857\|RS28_HUMAN 40S ribosomal protein S28 OS=Homo sapiens GN=RPS28 PE=1 | 91,004 | 30,43 | 7841 | 2 | 25,07 | 25,33 | 22,29 | 25,40 | 25,59 | 22,35 | 1,162 | 8,87E-01 | 0 | 0,21666654 | 0,05207462 |
| GLU2B_HUM | PRKCSH | sp\|P14314\|GLU2B_HUMAN Glucosidase 2 subunit beta OS=Homo sapiens GN=PRKCSH | 131,452 | 8,71 | 59425 | 3 | 22,32 | 22,27 | 23,27 | 22,94 | 22,96 | 22,61 | 1,162 | 5,64E-01 | 0 | 0,21666654 | 0,24905026 |
| FKBP5_HUM | FKBP5 | sp\|Q13451\|FKBP5_HUMAN Peptidyl-prolyl cis-trans isomerase FKBP5 OS=Homo sapiens | 11,112 | 2,19 | 51212 | 1 | 19,36 | 18,47 | 20,62 | 19,26 | 19,13 | 20,71 | 1,162 | 8,01E-01 | 0 | 0,21665827 | 0,09652652 |
| CCYL1_HUM | CCNYL1 | sp\|Q8N7R7\|CCYL1_HUMAN Cyclin-Y-like protein 1 OS=Homo sapiens GN=CCNYL1 PE=1 | 71,203 | 8,64 | 40705 | 3 | 22,12 | 22,33 | 18,50 | 22,05 | 22,20 | 19,35 | 1,162 | 8,96E-01 | 0 | 0,21605428 | 0,04775991 |
| PUR6_HUMA | PAICS | sp\|P22234\|PUR6_HUMAN Multifunctional protein ADE2 OS=Homo sapiens GN=PAICS P | 393,786 | 28 | 47079 | 11 | 26,22 | 26,06 | 25,72 | 26,20 | 26,37 | 26,07 | 1,159 | 2,80E-01 | 0 | 0,2133344 | 0,5520686 |
| TXD17_HUM | TXNDC17 | sp\|Q9BRA2\|TXD17_HUMAN Thioredoxin domain-containing protein 17 OS=Homo sapie | 16,539 | 7,32 | 13941 | 1 | 23,90 | 24,14 | 21,78 | 24,01 | 24,30 | 22,15 | 1,159 | 8,43E-01 | 0 | 0,21333313 | 0,07431246 |
| FNTA_HUMA | FNTA | sp\|P49354\|FNTA_HUMAN Protein farnesyltransferase/geranylgeranyltransferase type-1 | 186,352 | 21,9 | 44409 | 7 | 22,82 | 22,93 | 23,11 | 22,80 | 23,08 | 23,62 | 1,159 | 4,50E-01 | 0 | 0,21333313 | 0,3468026 |
| OLA1_HUMA | OLA1 | sp\|Q9NTK5\|OLA1_HUMAN Obg-like ATPase 1 OS=Homo sapiens GN=OLA1 PE=1 SV=2 | 407,639 | 39,65 | 44744 | 14 | 26,28 | 25,56 | 26,11 | 26,17 | 25,99 | 26,43 | 1,159 | 4,45E-01 | 0 | 0,21333313 | 0,35166277 |
| NC2B_HUMA | DR1 | sp\|Q01658\|NC2B_HUMAN Protein Dr1 OS=Homo sapiens GN=DR1 PE=1 SV=1 | 22,203 | 13,64 | 19444 | 1 | 17,82 | 18,65 | 19,25 | 17,95 | 19,35 | 19,05 | 1,157 | 7,42E-01 | 0 | 0,21000036 | 0,12983831 |
| RL10A_HUM | RPL10A | sp\|P62906\|RL10A_HUMAN 60S ribosomal protein L10a OS=Homo sapiens GN=RPL10A P | 400,327 | 59,45 | 24831 | 13 | 26,59 | 26,82 | 26,76 | 26,97 | 26,74 | 27,09 | 1,157 | 1,65E-01 | 0 | 0,20999972 | 0,78338451 |
| KPYM_HUM | PKM | sp\|P14618\|KPYM_HUMAN Pyruvate kinase PKM OS=Homo sapiens GN=PKM PE=1 SV=4 | 1878,815 | 56,69 | 57937 | 39 | 29,26 | 29,35 | 29,82 | 29,40 | 29,52 | 30,14 | 1,157 | 5,06E-01 | 0 | 0,20999972 | 0,2960514 |
| PNPH_HUM | PNP | sp\|P00491\|PNPH_HUMAN Purine nucleoside phosphorylase OS=Homo sapiens GN=PNP | 153,516 | 13,49 | 32118 | 4 | 26,02 | 26,34 | 23,72 | 26,02 | 26,48 | 24,21 | 1,157 | 8,55E-01 | 0 | 0,20999972 | 0,06804841 |
| KAT3_HUMA | KYAT3 | sp\|Q6YP21\|KAT3_HUMAN Kynurenine--oxoglutarate transaminase 3 OS=Homo sapiens | 41,809 | 6,17 | 51400 | 2 | 21,80 | 21,76 | 18,96 | 21,74 | 21,96 | 19,45 | 1,156 | 8,74E-01 | 0 | 0,20877584 | 0,05841434 |
| S14L1_HUM | SEC14L1 | sp\|Q92503\|S14L1_HUMAN SEC14-like protein 1 OS=Homo sapiens GN=SEC14L1 PE=1 S | 13,485 | 2,52 | 81250 | 1 | 17,51 | 17,86 | 17,44 | 17,17 | 17,24 | 19,03 | 1,155 | 7,54E-01 | 0 | 0,20850182 | 0,12268731 |
| CIB1_HUMA | CIB1 | sp\|Q99828\|CIB1_HUMAN Calcium and integrin-binding protein 1 OS=Homo sapiens GN= | 45,088 | 6,81 | 21703 | 1 | 20,73 | 21,07 | 18,36 | 20,29 | 20,39 | 20,10 | 1,155 | 8,20E-01 | 0 | 0,20788892 | 0,08596261 |
| INT4_HUMA | INTS4 | sp\|Q96HW7\|INT4_HUMAN Integrator complex subunit 4 OS=Homo sapiens GN=INTS4 P | 44,44 | 1,87 | 108171 | 1 | 17,86 | 19,14 | 17,99 | 19,27 | 16,86 | 19,48 | 1,155 | 8,35E-01 | 0 | 0,20787621 | 0,07848424 |
| RL13A_HUM | RPL13A | sp\|P40429\|RL13A_HUMAN 60S ribosomal protein L13a OS=Homo sapiens GN=RPL13A P | 106,07 | 24,14 | 23577 | 5 | 24,55 | 24,81 | 24,39 | 24,82 | 24,43 | 25,12 | 1,154 | 4,27E-01 | 0 | 0,20666758 | 0,36907586 |
| CRKL_HUMA | CRKL | sp\|P46109\|CRKL_HUMAN Crk-like protein OS=Homo sapiens GN=CRKL PE=1 SV=1 | 187,835 | 24,75 | 33777 | 5 | 23,96 | 24,22 | 23,23 | 24,10 | 24,09 | 23,84 | 1,154 | 5,39E-01 | 0 | 0,20666758 | 0,26813477 |
| ELOC_HUMA | ELOC | sp\|Q15369\|ELOC_HUMAN Elongin-C OS=Homo sapiens GN=ELOC PE=1 SV=1 | 247,742 | 66,07 | 12473 | 6 | 24,69 | 24,97 | 23,48 | 24,74 | 24,91 | 24,11 | 1,154 | 7,10E-01 | 0 | 0,20666695 | 0,14858456 |
| NB5R3_HUM | CYB5R3 | sp\|P00387\|NB5R3_HUMAN NADH-cytochrome b5 reductase 3 OS=Homo sapiens GN=C | 295,182 | 23,26 | 34235 | 9 | 25,43 | 25,38 | 26,28 | 25,65 | 25,72 | 26,34 | 1,154 | 6,02E-01 | 0 | 0,20666631 | 0,22063454 |
| GOLI4_HUM | GOLIM4 | sp\|O00461\|GOLI4_HUMAN Golgi integral membrane protein 4 OS=Homo sapiens GN=G | 26,129 | 1,58 | 81880 | 1 | 22,02 | 22,57 | 22,91 | 22,38 | 22,49 | 23,25 | 1,154 | 6,13E-01 | 0 | 0,20666631 | 0,21280977 |
| DUT_HUMAN | DUT | sp\|P33316\|DUT_HUMAN Deoxyuridine 5'-triphosphate nucleotidohydrolase, mitochon | 109,792 | 20,63 | 26563 | 5 | 23,83 | 24,49 | 24,60 | 24,39 | 24,49 | 24,66 | 1,154 | 4,60E-01 | 0 | 0,20666631 | 0,33732987 |
| UFC1_HUMA | UFC1 | sp\|Q9Y3C8\|UFC1_HUMAN Ubiquitin-fold modifier-conjugating enzyme 1 OS=Homo sap | 36,747 | 22,75 | 19458 | 2 | 20,74 | 22,13 | 18,99 | 21,10 | 20,94 | 20,43 | 1,152 | 8,37E-01 | 0 | 0,20467504 | 0,0773307 |
| TPM4_HUM | TPM4 | sp\|P67936\|TPM4_HUMAN Tropomyosin alpha-4 chain OS=Homo sapiens GN=TPM4 PE= | 1363,185 | 83,06 | 28522 | 36 | 28,21 | 28,23 | 26,98 | 28,67 | 28,33 | 27,03 | 1,151 | 7,70E-01 | 0 | 0,20333417 | 0,11376222 |
| DRG2_HUM | DRG2 | sp\|P55039\|DRG2_HUMAN Developmentally-regulated GTP-binding protein 2 OS=Homo | 90,656 | 17,86 | 40746 | 3 | 19,50 | 21,27 | 22,08 | 19,90 | 21,45 | 22,11 | 1,151 | 8,49E-01 | 0 | 0,20333354 | 0,07084348 |
| PPCE_HUMA | PREP | sp\|P48147\|PPCE_HUMAN Prolyl endopeptidase OS=Homo sapiens GN=PREP PE=1 SV=2 | 258,349 | 13,24 | 80700 | 8 | 24,18 | 24,37 | 23,21 | 24,04 | 24,49 | 23,84 | 1,151 | 6,44E-01 | 0 | 0,20333354 | 0,19117274 |
| IMA1_HUMA | KPNA2 | sp\|P52292\|IMA1_HUMAN Importin subunit alpha-1 OS=Homo sapiens GN=KPNA2 PE=1 | 661,91 | 44,05 | 57862 | 14 | 26,14 | 26,15 | 26,90 | 26,41 | 26,57 | 26,82 | 1,151 | 5,06E-01 | 0 | 0,20333354 | 0,2959978 |
| U5S1_HUMA | EFTUD2 | sp\|Q15029\|U5S1_HUMAN 116 kDa U5 small nuclear ribonucleoprotein component OS= | 1104,514 | 45,27 | 109436 | 31 | 25,96 | 26,34 | 27,11 | 26,29 | 26,40 | 27,33 | 1,151 | 6,89E-01 | 0 | 0,20333354 | 0,16170753 |
| AK1BA_HUM | AKR1B10 | sp\|O60218\|AK1BA_HUMAN Aldo-keto reductase family 1 member B10 OS=Homo sapier | 61,674 | 10,44 | 36020 | 4 | 23,44 | 23,60 | 19,49 | 23,69 | 23,71 | 19,73 | 1,150 | 9,20E-01 | 0 | 0,20196851 | 0,03624967 |
| IF2A_HUMA | EIF2S1 | sp\|P05198\|IF2A_HUMAN Eukaryotic translation initiation factor 2 subunit 1 OS=Homo s | 423,818 | 52,06 | 36112 | 15 | 26,46 | 26,60 | 26,22 | 26,58 | 26,79 | 26,51 | 1,149 | 2,24E-01 | 0 | 0,20000076 | 0,64925135 |
| HINT1_HUM | HINT1 | sp\|P49773\|HINT1_HUMAN Histidine triad nucleotide-binding protein 1 OS=Homo sapier | 151,527 | 30,16 | 13802 | 4 | 25,43 | 25,75 | 23,92 | 25,59 | 25,62 | 24,49 | 1,149 | 7,82E-01 | 0 | 0,20000013 | 0,10680404 |
| RS13_HUMA | RPS13 | sp\|P62277\|RS13_HUMAN 40S ribosomal protein S13 OS=Homo sapiens GN=RPS13 PE=1 | 294,898 | 47,68 | 17222 | 9 | 27,26 | 27,38 | 27,16 | 27,45 | 27,57 | 27,38 | 1,149 | 7,68E-02 | 0 | 0,20000013 | 1,11445507 |
| MIA2_HUMA | MIA2 | sp\|Q96PC5\|MIA2_HUMAN Melanoma inhibitory activity protein 2 OS=Homo sapiens GN | 13,279 | 0,5 | 159836 | 1 | 24,18 | 21,78 | 23,47 | 24,36 | 22,56 | 23,11 | 1,149 | 8,33E-01 | 0 | 0,20000013 | 0,07934277 |
| IF5A1_HUM | EIF5A | sp\|P63241\|IF5A1_HUMAN Eukaryotic translation initiation factor 5A-1 OS=Homo sapien | 938,82 | 88,31 | 16832 | 22 | 28,17 | 28,12 | 27,96 | 28,32 | 28,20 | 28,33 | 1,149 | 5,78E-02 | 0 | 0,20000013 | 1,23801943 |
| SEH1_HUMA | SEH1L | sp\|Q96EE3\|SEH1_HUMAN Nucleoporin SEH1 OS=Homo sapiens GN=SEH1L PE=1 SV=3 | 14,394 | 3,06 | 39649 | 1 | 20,07 | 20,02 | 20,97 | 20,33 | 20,21 | 21,12 | 1,149 | 6,59E-01 | 0 | 0,20000013 | 0,18105182 |
| RUVB2_HUM | RUVBL2 | sp\|Q9Y230\|RUVB2_HUMAN RuvB-like 2 OS=Homo sapiens GN=RUVBL2 PE=1 SV=3 | 1148,543 | 51,19 | 51157 | 24 | 26,88 | 27,12 | 27,76 | 27,04 | 27,27 | 28,05 | 1,149 | 6,46E-01 | 0 | 0,20000013 | 0,18996161 |
| PSMG2_HUM | PSMG2 | sp\|Q969U7\|PSMG2_HUMAN Proteasome assembly chaperone 2 OS=Homo sapiens GN= | 22,944 | 4,55 | 29396 | 1 | 19,97 | 20,34 | 20,40 | 20,17 | 20,40 | 20,74 | 1,149 | 4,01E-01 | 0 | 0,20000013 | 0,39634242 |
| UBE2N_HUM | UBE2N | sp\|P61088\|UBE2N_HUMAN Ubiquitin-conjugating enzyme E2 N OS=Homo sapiens GN= | 276,043 | 79,61 | 17138 | 10 | 26,74 | 27,08 | 25,36 | 26,75 | 27,02 | 26,01 | 1,149 | 7,58E-01 | 0 | 0,20000013 | 0,12027986 |
| SDC2_HUMA | SDC2 | sp\|P34741\|SDC2_HUMAN Syndecan-2 OS=Homo sapiens GN=SDC2 PE=1 SV=2 | 109,059 | 5,97 | 22160 | 3 | 24,73 | 22,04 | 24,37 | 24,38 | 23,84 | 23,52 | 1,149 | 8,31E-01 | 0 | 0,19999949 | 0,08024725 |
| YES_HUMAN | YES1 | sp\|P07947\|YES_HUMAN Tyrosine-protein kinase Yes OS=Homo sapiens GN=YES1 PE=1 | 466,952 | 37,2 | 60801 | 16 | 25,01 | 24,45 | 22,94 | 25,00 | 25,01 | 22,99 | 1,149 | 8,37E-01 | 0 | 0,19999949 | 0,07711781 |
| HEAT1_HUM | HEATR1 | sp\|Q9H583\|HEAT1_HUMAN HEAT repeat-containing protein 1 OS=Homo sapiens GN=H | 79,397 | 2,52 | 242370 | 3 | 17,17 | 17,80 | 20,35 | 17,97 | 17,48 | 20,47 | 1,146 | 8,90E-01 | 0 | 0,19699605 | 0,05041347 |
| ANXA8_HUM | ANXA8 | sp\|P13928\|ANXA8_HUMAN Annexin A8 OS=Homo sapiens GN=ANXA8 PE=1 SV=3 | 427,769 | 29,97 | 36881 | 12 | 27,07 | 27,13 | 27,23 | 27,10 | 27,52 | 27,40 | 1,146 | 2,14E-01 | 0 | 0,19666735 | 0,66910856 |
| EF2_HUMAN | EEF2 | sp\|P13639\|EF2_HUMAN Elongation factor 2 OS=Homo sapiens GN=EEF2 PE=1 SV=4 | 3474,534 | 70,05 | 95338 | 81 | 30,30 | 30,46 | 30,81 | 30,36 | 30,57 | 31,23 | 1,146 | 5,51E-01 | 0 | 0,19666735 | 0,25901663 |
| CRK_HUMAN | CRK | sp\|P46108\|CRK_HUMAN Adapter molecule crk OS=Homo sapiens GN=CRK PE=1 SV=2 | 191,301 | 38,49 | 33831 | 8 | 23,96 | 24,21 | 21,12 | 23,77 | 24,09 | 22,02 | 1,146 | 8,76E-01 | 0 | 0,19666735 | 0,05755971 |
| NUP37_HUM | NUP37 | sp\|Q8NFH4\|NUP37_HUMAN Nucleoporin Nup37 OS=Homo sapiens GN=NUP37 PE=1 SV | 15,519 | 4,6 | 36708 | 1 | 20,10 | 19,94 | 20,86 | 20,27 | 20,02 | 21,20 | 1,146 | 6,90E-01 | 0 | 0,19666672 | 0,16146196 |
| ODPB_HUM | PDHB | sp\|P11177\|ODPB_HUMAN Pyruvate dehydrogenase E1 component subunit beta, mitocl | 171,612 | 20,33 | 39233 | 6 | 23,19 | 23,64 | 25,40 | 24,10 | 23,96 | 24,76 | 1,146 | 7,98E-01 | 0 | 0,19666672 | 0,09815125 |
| ADHX_HUM | ADH5 | sp\|P11766\|ADHX_HUMAN Alcohol dehydrogenase class-3 OS=Homo sapiens GN=ADH5 | 164,717 | 14,71 | 39724 | 4 | 24,98 | 25,09 | 24,57 | 25,06 | 25,23 | 24,94 | 1,146 | 3,34E-01 | 0 | 0,19666672 | 0,47618265 |
| PCD15_HUM | PCDH15 | sp\|Q96QU1\|PCD15_HUMAN Protocadherin-15 OS=Homo sapiens GN=PCDH15 PE=1 SV= | 10,39 | 0,36 | 216069 | 1 | 22,50 | 22,43 | 19,37 | 22,52 | 22,16 | 20,21 | 1,145 | 8,84E-01 | 0 | 0,19576136 | 0,05363166 |
| CAN1_HUM | CAPN1 | sp\|P07384\|CAN1_HUMAN Calpain-1 catalytic subunit OS=Homo sapiens GN=CAPN1 PE= | 346,526 | 22,41 | 81890 | 14 | 25,24 | 25,57 | 24,02 | 25,27 | 25,60 | 24,54 | 1,143 | 7,50E-01 | 0 | 0,19333394 | 0,12503566 |
| RPTOR_HUM | RPTOR | sp\|Q8N122\|RPTOR_HUMAN Regulatory-associated protein of mTOR OS=Homo sapiens | 89,555 | 3,82 | 149038 | 3 | 20,76 | 20,29 | 22,17 | 20,95 | 20,26 | 22,59 | 1,143 | 8,39E-01 | 0 | 0,19333331 | 0,07618429 |
| GTF2I_HUM | GTF2I | sp\|P78347\|GTF2I_HUMAN General transcription factor II-I OS=Homo sapiens GN=GTF2I | 194,8 | 14,43 | 112416 | 9 | 21,62 | 22,42 | 23,58 | 21,99 | 22,68 | 23,53 | 1,143 | 8,02E-01 | 0 | 0,19333331 | 0,095685 |
| AAPK1_HUM | PRKAA1 | sp\|Q13131\|AAPK1_HUMAN 5'-AMP-activated protein kinase catalytic subunit alpha-1 O | 132,876 | 10,91 | 64009 | 5 | 21,90 | 21,44 | 22,99 | 22,34 | 21,91 | 22,66 | 1,143 | 7,23E-01 | 0 | 0,19333331 | 0,14082756 |
| ENOA_HUM | ENO1 | sp\|P06733\|ENOA_HUMAN Alpha-enolase OS=Homo sapiens GN=ENO1 PE=1 SV=2 | 1405,938 | 54,15 | 47169 | 26 | 30,34 | 30,35 | 29,27 | 30,51 | 30,52 | 29,51 | 1,143 | 7,14E-01 | 0 | 0,19333331 | 0,14654089 |
| VATB2_HUM | ATP6V1B2 | sp\|P21281\|VATB2_HUMAN V-type proton ATPase subunit B, brain isoform OS=Homo sa | 317 | 25,64 | 56501 | 8 | 24,01 | 24,42 | 24,29 | 24,40 | 24,65 | 24,25 | 1,143 | 3,14E-01 | 0 | 0,19333267 | 0,50295338 |
| ARF1_HUMA | ARF1 | sp\|P84077\|ARF1_HUMAN ADP-ribosylation factor 1 OS=Homo sapiens GN=ARF1 PE=1 S | 999,084 | 86,19 | 20697 | 22 | 22,51 | 22,95 | 19,67 | 22,60 | 22,65 | 20,46 | 1,143 | 8,85E-01 | 0 | 0,19333267 | 0,05299726 |
| RS10_HUMA | RPS10 | sp\|P46783\|RS10_HUMAN 40S ribosomal protein S10 OS=Homo sapiens GN=RPS10 PE=1 | 190,681 | 33,33 | 18898 | 6 | 25,39 | 25,41 | 25,95 | 25,47 | 25,63 | 26,23 | 1,143 | 5,48E-01 | 0 | 0,19333267 | 0,26098446 |
| RS27A_HUM | RPS27A | sp\|P62979\|RS27A_HUMAN Ubiquitin-40S ribosomal protein S27a OS=Homo sapiens GN | 1002,377 | 68,59 | 17965 | 20 | 23,95 | 24,27 | 22,78 | 24,07 | 24,21 | 23,30 | 1,143 | 7,36E-01 | 0 | 0,19333204 | 0,13334944 |
| UBE2Z_HUM | UBE2Z | sp\|Q9H832\|UBE2Z_HUMAN Ubiquitin-conjugating enzyme E2 Z OS=Homo sapiens GN=U | 112,464 | 16,38 | 38210 | 6 | 24,48 | 24,06 | 23,29 | 24,63 | 24,44 | 23,33 | 1,141 | 7,40E-01 | 0 | 0,1899999 | 0,1306409 |
| SF3B6_HUM | SF3B6 | sp\|Q9Y3B4\|SF3B6_HUMAN Splicing factor 3B subunit 6 OS=Homo sapiens GN=SF3B6 PE | 63,228 | 9,6 | 14585 | 1 | 21,76 | 21,76 | 22,42 | 21,79 | 22,09 | 22,63 | 1,141 | 5,95E-01 | 0 | 0,1899999 | 0,22514906 |
| H2AV_HUM | H2AFV | sp\|Q71UI9\|H2AV_HUMAN Histone H2A.V OS=Homo sapiens GN=H2AFV PE=1 SV=3 | 179,848 | 53,91 | 13509 | 5 | 23,94 | 24,41 | 23,25 | 24,04 | 24,17 | 23,96 | 1,141 | 6,09E-01 | 0 | 0,1899999 | 0,21573807 |
| RL11_HUMA | RPL11 | sp\|P62913\|RL11_HUMAN 60S ribosomal protein L11 OS=Homo sapiens GN=RPL11 PE=1 | 375,721 | 53,37 | 20252 | 9 | 26,10 | 26,09 | 27,61 | 26,28 | 25,95 | 28,14 | 1,141 | 8,34E-01 | 0 | 0,1899999 | 0,07895715 |
| ACL6A_HUM | ACTL6A | sp\|O96019\|ACL6A_HUMAN Actin-like protein 6A OS=Homo sapiens GN=ACTL6A PE=1 S | 66,225 | 9,79 | 47461 | 3 | 21,85 | 21,86 | 22,35 | 21,76 | 22,16 | 22,71 | 1,141 | 5,86E-01 | 0 | 0,18999926 | 0,2322521 |
| PAPOA_HUM | PAPOLA | sp\|P51003\|PAPOA_HUMAN Poly(A) polymerase alpha OS=Homo sapiens GN=PAPOLA P | 10,353 | 1,07 | 82843 | 1 | 20,45 | 20,28 | 18,58 | 20,50 | 20,81 | 18,57 | 1,139 | 8,49E-01 | 0 | 0,18720245 | 0,07116906 |
| H2B1C_HUM | HIST1H2BC | sp\|P62807\|H2B1C_HUMAN Histone H2B type 1-C/E/F/G/I OS=Homo sapiens GN=HIST1 | 465,828 | 35,71 | 13906 | 8 | 25,80 | 26,05 | 26,90 | 26,39 | 26,26 | 26,66 | 1,138 | 6,25E-01 | 0 | 0,18666712 | 0,20407649 |
| VIGLN_HUM | HDLBP | sp\|Q00341\|VIGLN_HUMAN Vigilin OS=Homo sapiens GN=HDLBP PE=1 SV=2 | 329,537 | 13,17 | 141456 | 14 | 25,72 | 24,99 | 25,82 | 25,82 | 25,73 | 25,54 | 1,138 | 5,34E-01 | 0 | 0,18666712 | 0,27283612 |
| EXOC5_HUM | EXOC5 | sp\|O00471\|EXOC5_HUMAN Exocyst complex component 5 OS=Homo sapiens GN=EXO | 77,981 | 5,79 | 81853 | 3 | 22,59 | 22,15 | 20,95 | 22,29 | 22,67 | 21,29 | 1,138 | 7,85E-01 | 0 | 0,18666712 | 0,10511355 |
| EXOC8_HUM | EXOC8 | sp\|Q8IYI6\|EXOC8_HUMAN Exocyst complex component 8 OS=Homo sapiens GN=EXOC | 23,2 | 1,52 | 81799 | 1 | 19,18 | 19,51 | 18,92 | 20,24 | 19,09 | 18,84 | 1,138 | 7,08E-01 | 0 | 0,18666649 | 0,15006915 |
| PLST_HUMA | PLS3 | sp\|P13797\|PLST_HUMAN Plastin-3 OS=Homo sapiens GN=PLS3 PE=1 SV=4 | 632,839 | 46,35 | 70811 | 21 | 25,61 | 25,41 | 23,36 | 25,20 | 25,66 | 24,08 | 1,138 | 8,39E-01 | 0 | 0,18666649 | 0,07648788 |
| RSMB_HUM | SNRPB | sp\|P14678\|RSMB_HUMAN Small nuclear ribonucleoprotein-associated proteins B and B | 223,09 | 24,17 | 24610 | 8 | 25,72 | 25,77 | 25,76 | 25,86 | 25,99 | 25,95 | 1,136 | 1,14E-02 | 0 | 0,18333371 | 1,94294395 |
| 2AAA_HUM | PPP2R1A | sp\|P30153\|2AAA_HUMAN Serine/threonine-protein phosphatase 2A 65 kDa regulatory | 1419,674 | 57,39 | 65309 | 32 | 27,23 | 27,40 | 27,60 | 27,40 | 27,59 | 27,79 | 1,136 | 3,03E-01 | 0 | 0,18333371 | 0,51835571 |
| NDRG1_HUM | NDRG1 | sp\|Q92597\|NDRG1_HUMAN Protein NDRG1 OS=Homo sapiens GN=NDRG1 PE=1 SV=1 | 133,106 | 12,18 | 42835 | 3 | 21,59 | 21,82 | 22,72 | 21,78 | 21,96 | 22,94 | 1,136 | 7,32E-01 | 0 | 0,18333371 | 0,13559322 |
| PIPNA_HUM | PITPNA | sp\|Q00169\|PIPNA_HUMAN Phosphatidylinositol transfer protein alpha isoform OS=Hom | 107,113 | 22,96 | 31806 | 5 | 22,99 | 22,84 | 21,71 | 22,82 | 23,19 | 22,08 | 1,136 | 7,42E-01 | 0 | 0,18333371 | 0,12966331 |
| RL5_HUMAN | RPL5 | sp\|P46777\|RL5_HUMAN 60S ribosomal protein L5 OS=Homo sapiens GN=RPL5 PE=1 SV= | 521,698 | 49,83 | 34363 | 15 | 26,79 | 27,03 | 27,22 | 27,13 | 27,01 | 27,45 | 1,136 | 3,68E-01 | 0 | 0,18333308 | 0,43397742 |
| MX1_HUMA | MX1 | sp\|P20591\|MX1_HUMAN Interferon-induced GTP-binding protein Mx1 OS=Homo sapie | 52,777 | 3,93 | 75520 | 2 | 20,37 | 20,94 | 23,17 | 20,50 | 21,17 | 23,36 | 1,136 | 8,87E-01 | 0 | 0,18333308 | 0,0519076 |
| MDHC_HUM | MDH1 | sp\|P40925\|MDHC_HUMAN Malate dehydrogenase, cytoplasmic OS=Homo sapiens GN= | 422,58 | 35,03 | 36426 | 8 | 26,42 | 26,62 | 25,25 | 26,58 | 26,76 | 25,50 | 1,136 | 7,68E-01 | 0 | 0,18333308 | 0,11460921 |
| AIFM2_HUM | AIFM2 | sp\|Q9BRQ8\|AIFM2_HUMAN Apoptosis-inducing factor 2 OS=Homo sapiens GN=AIFM2 | 120,25 | 12,6 | 40527 | 4 | 25,31 | 25,59 | 27,54 | 26,15 | 25,58 | 27,26 | 1,136 | 8,41E-01 | 0 | 0,18333308 | 0,07512876 |
| AMPL_HUM | LAP3 | sp\|P28838\|AMPL_HUMAN Cytosol aminopeptidase OS=Homo sapiens GN=LAP3 PE=1 S | 367,848 | 24,66 | 56166 | 10 | 25,32 | 25,36 | 24,85 | 25,53 | 25,47 | 25,08 | 1,136 | 4,44E-01 | 0 | 0,18333308 | 0,35254839 |
| RPN2_HUMA | RPN2 | sp\|P04844\|RPN2_HUMAN Dolichyl-diphosphooligosaccharide--protein glycosyltransfera | 188,883 | 18,54 | 69284 | 6 | 24,85 | 23,85 | 26,41 | 24,41 | 24,94 | 26,31 | 1,136 | 8,54E-01 | 0 | 0,18333308 | 0,06844356 |
| SYMC_HUM | MARS | sp\|P56192\|SYMC_HUMAN Methionine--tRNA ligase, cytoplasmic OS=Homo sapiens GN= | 186,655 | 8,78 | 101116 | 6 | 23,45 | 23,81 | 24,73 | 23,53 | 23,96 | 25,05 | 1,136 | 7,72E-01 | 0 | 0,18333308 | 0,11233765 |
| MCM5_HUM | MCM5 | sp\|P33992\|MCM5_HUMAN DNA replication licensing factor MCM5 OS=Homo sapiens G | 433,287 | 24,93 | 82286 | 14 | 24,61 | 24,85 | 26,21 | 24,90 | 25,01 | 26,31 | 1,136 | 7,99E-01 | 0 | 0,18333308 | 0,09753729 |
| RS23_HUMA | RPS23 | sp\|P62266\|RS23_HUMAN 40S ribosomal protein S23 OS=Homo sapiens GN=RPS23 PE=1 | 305,922 | 49,65 | 15808 | 8 | 25,21 | 25,72 | 26,10 | 25,64 | 25,38 | 26,56 | 1,136 | 6,99E-01 | 0 | 0,18333308 | 0,15549654 |
| ABCF1_HUM | ABCF1 | sp\|Q8NE71\|ABCF1_HUMAN ATP-binding cassette sub-family F member 1 OS=Homo sap | 296,717 | 21,07 | 95926 | 10 | 24,46 | 24,51 | 25,13 | 24,40 | 24,65 | 25,59 | 1,133 | 6,91E-01 | 0 | 0,18000031 | 0,1603135 |
| COR1B_HUM | CORO1B | sp\|Q9BR76\|COR1B_HUMAN Coronin-1B OS=Homo sapiens GN=CORO1B PE=1 SV=1 | 303,945 | 26,18 | 54235 | 8 | 26,22 | 26,01 | 24,89 | 26,16 | 26,36 | 25,14 | 1,133 | 7,64E-01 | 0 | 0,18000031 | 0,11701594 |
| P5CR3_HUM | PYCR3 | sp\|Q53H96\|P5CR3_HUMAN Pyrroline-5-carboxylate reductase 3 OS=Homo sapiens GN= | 129,263 | 8,76 | 28649 | 3 | 24,06 | 24,10 | 23,82 | 24,14 | 24,34 | 24,04 | 1,133 | 2,21E-01 | 0 | 0,18000031 | 0,6559825 |
| CAND1_HUM | CAND1 | sp\|Q86VP6\|CAND1_HUMAN Cullin-associated NEDD8-dissociated protein 1 OS=Homo s | 1288,802 | 40,98 | 136376 | 37 | 27,31 | 27,24 | 26,66 | 27,14 | 27,50 | 27,11 | 1,133 | 4,97E-01 | 0 | 0,18000031 | 0,30379874 |
| NTF2_HUMA | NUTF2 | sp\|P61970\|NTF2_HUMAN Nuclear transport factor 2 OS=Homo sapiens GN=NUTF2 PE= | 138,158 | 55,12 | 14478 | 5 | 25,27 | 25,90 | 21,90 | 25,40 | 25,76 | 22,45 | 1,133 | 9,17E-01 | 0 | 0,18000031 | 0,0375625 |
| KCAB2_HUM | KCNAB2 | sp\|Q13303\|KCAB2_HUMAN Voltage-gated potassium channel subunit beta-2 OS=Homo | 129,769 | 13,9 | 41000 | 3 | 22,59 | 23,23 | 20,17 | 22,56 | 22,88 | 21,09 | 1,132 | 8,76E-01 | 0 | 0,17941348 | 0,05731821 |
| RBGPR_HUM | RAB3GAP2 | sp\|Q9H2M9\|RBGPR_HUMAN Rab3 GTPase-activating protein non-catalytic subunit OS= | 20,403 | 1,65 | 155985 | 1 | 18,47 | 19,04 | 18,07 | 18,09 | 18,94 | 19,08 | 1,131 | 6,93E-01 | 0 | 0,17752647 | 0,15904405 |
| PP6R3_HUM | PPP6R3 | sp\|Q5H9R7\|PP6R3_HUMAN Serine/threonine-protein phosphatase 6 regulatory subuni | 179,861 | 7,79 | 97669 | 7 | 23,32 | 23,42 | 20,76 | 22,34 | 23,67 | 22,02 | 1,130 | 8,69E-01 | 0 | 0,1766669 | 0,06088655 |
| RS17_HUMA | RPS17 | sp\|P08708\|RS17_HUMAN 40S ribosomal protein S17 OS=Homo sapiens GN=RPS17 PE=1 | 418,508 | 59,26 | 15550 | 10 | 25,23 | 25,51 | 25,95 | 25,61 | 25,24 | 26,37 | 1,130 | 6,76E-01 | 0 | 0,1766669 | 0,16978251 |
| UBC12_HUM | UBE2M | sp\|P61081\|UBC12_HUMAN NEDD8-conjugating enzyme Ubc12 OS=Homo sapiens GN=U | 307,386 | 42,08 | 20900 | 8 | 26,41 | 26,44 | 25,66 | 26,50 | 26,67 | 25,87 | 1,130 | 6,43E-01 | 0 | 0,1766669 | 0,19203284 |
| AKCL2_HUM | AKR1E2 | sp\|Q96JD6\|AKCL2_HUMAN 1,5-anhydro-D-fructose reductase OS=Homo sapiens GN=A | 11,541 | 2,5 | 36589 | 1 | 24,29 | 24,18 | 23,66 | 24,37 | 24,25 | 24,04 | 1,130 | 4,61E-01 | 0 | 0,1766669 | 0,33624151 |
| TTYH3_HUM | TTYH3 | sp\|Q9C0H2\|TTYH3_HUMAN Protein tweety homolog 3 OS=Homo sapiens GN=TTYH3 PE | 141,869 | 3,06 | 57545 | 2 | 23,26 | 22,46 | 23,10 | 23,44 | 23,39 | 22,52 | 1,130 | 6,71E-01 | 0 | 0,1766669 | 0,17335285 |
| UGPA_HUM | UGP2 | sp\|Q16851\|UGPA_HUMAN UTP--glucose-1-phosphate uridylyltransferase OS=Homo sap | 121,718 | 14,96 | 56940 | 5 | 23,76 | 23,87 | 23,16 | 23,69 | 24,08 | 23,55 | 1,130 | 5,51E-01 | 0 | 0,17666626 | 0,25884266 |
| PPME1_HUM | PPME1 | sp\|Q9Y570\|PPME1_HUMAN Protein phosphatase methylesterase 1 OS=Homo sapiens G | 230,558 | 25,91 | 42315 | 9 | 25,26 | 25,25 | 23,97 | 25,14 | 25,39 | 24,48 | 1,130 | 7,45E-01 | 0 | 0,17666626 | 0,12777582 |
| EFMT1_HUM | EEF1AKMT1 | sp\|Q8WVE0\|EFMT1_HUMAN EEF1A lysine methyltransferase 1 OS=Homo sapiens GN=E | 13,306 | 5,61 | 24506 | 1 | 21,08 | 20,91 | 18,85 | 19,35 | 21,04 | 20,97 | 1,128 | 8,57E-01 | 0 | 0,17427826 | 0,06723113 |
| TPIS_HUMAN | TPI1 | sp\|P60174\|TPIS_HUMAN Triosephosphate isomerase OS=Homo sapiens GN=TPI1 PE=1 | 890,147 | 68,18 | 30791 | 19 | 28,82 | 28,98 | 28,53 | 29,00 | 29,09 | 28,76 | 1,128 | 3,51E-01 | 0 | 0,17333349 | 0,45427853 |
| RUXF_HUMA | SNRPF | sp\|P62306\|RUXF_HUMAN Small nuclear ribonucleoprotein F OS=Homo sapiens GN=SN | 159,397 | 58,14 | 9725 | 6 | 23,20 | 23,67 | 21,39 | 23,00 | 23,85 | 21,93 | 1,128 | 8,55E-01 | 0 | 0,17333349 | 0,06801747 |
| DDX1_HUMA | DDX1 | sp\|Q92499\|DDX1_HUMAN ATP-dependent RNA helicase DDX1 OS=Homo sapiens GN=D | 714,649 | 34,73 | 82432 | 22 | 26,00 | 26,36 | 26,56 | 26,27 | 26,37 | 26,80 | 1,128 | 4,94E-01 | 0 | 0,17333349 | 0,30587966 |
| ELMO2_HUM | ELMO2 | sp\|Q96JJ3\|ELMO2_HUMAN Engulfment and cell motility protein 2 OS=Homo sapiens GN | 76,993 | 7,36 | 82615 | 4 | 21,78 | 22,60 | 21,51 | 22,24 | 22,56 | 21,61 | 1,128 | 7,08E-01 | 0 | 0,17333285 | 0,15010265 |
| SF3B2_HUM | SF3B2 | sp\|Q13435\|SF3B2_HUMAN Splicing factor 3B subunit 2 OS=Homo sapiens GN=SF3B2 PE | 182,376 | 12,96 | 100228 | 8 | 24,62 | 24,35 | 24,46 | 24,71 | 24,51 | 24,73 | 1,128 | 1,75E-01 | 0 | 0,17333285 | 0,75712662 |
| TSP4_HUMA | THBS4 | sp\|P35443\|TSP4_HUMAN Thrombospondin-4 OS=Homo sapiens GN=THBS4 PE=1 SV=2 | 127,326 | 6,35 | 105869 | 4 | 22,06 | 19,33 | 22,52 | 22,38 | 20,17 | 21,88 | 1,128 | 8,92E-01 | 0 | 0,17333285 | 0,04959059 |
| SPRC_HUMA | SPARC | sp\|P09486\|SPRC_HUMAN SPARC OS=Homo sapiens GN=SPARC PE=1 SV=1 | 44,756 | 7,92 | 34632 | 3 | 23,86 | 23,58 | 22,46 | 24,06 | 23,89 | 22,47 | 1,128 | 8,06E-01 | 0 | 0,17333285 | 0,09360252 |
| COTL1_HUM | COTL1 | sp\|Q14019\|COTL1_HUMAN Coactosin-like protein OS=Homo sapiens GN=COTL1 PE=1 S | 128,772 | 33,8 | 15945 | 6 | 25,26 | 24,79 | 23,62 | 25,30 | 25,22 | 23,67 | 1,128 | 8,22E-01 | 0 | 0,17333221 | 0,08527434 |
| ELOV1_HUM | ELOVL1 | sp\|Q9BW60\|ELOV1_HUMAN Elongation of very long chain fatty acids protein 1 OS=Hom | 49,719 | 5,02 | 32663 | 1 | 19,54 | 19,66 | 18,42 | 19,84 | 18,14 | 20,15 | 1,126 | 8,28E-01 | 0 | 0,17146683 | 0,08178662 |
| SNX2_HUMA | SNX2 | sp\|O60749\|SNX2_HUMAN Sorting nexin-2 OS=Homo sapiens GN=SNX2 PE=1 SV=2 | 482,16 | 29,48 | 58471 | 14 | 24,88 | 24,89 | 24,84 | 24,79 | 25,11 | 25,22 | 1,125 | 2,61E-01 | 0 | 0,17000071 | 0,58390727 |
| PAK2_HUMA | PAK2 | sp\|Q13177\|PAK2_HUMAN Serine/threonine-protein kinase PAK 2 OS=Homo sapiens GN | 186,258 | 16,41 | 58043 | 7 | 24,23 | 24,35 | 21,21 | 23,89 | 23,49 | 22,92 | 1,125 | 8,81E-01 | 0 | 0,17000008 | 0,0550611 |
| PSMD4_HUM | PSMD4 | sp\|P55036\|PSMD4_HUMAN 26S proteasome non-ATPase regulatory subunit 4 OS=Hom | 328,646 | 24,4 | 40737 | 7 | 26,47 | 26,29 | 25,54 | 26,47 | 26,37 | 25,97 | 1,125 | 6,27E-01 | 0 | 0,16999944 | 0,20296171 |
| RFA3_HUMA | RPA3 | sp\|P35244\|RFA3_HUMAN Replication protein A 14 kDa subunit OS=Homo sapiens GN=R | 23,778 | 8,26 | 13569 | 1 | 21,20 | 21,62 | 20,64 | 21,38 | 21,59 | 21,00 | 1,125 | 6,36E-01 | 0 | 0,16999944 | 0,19665315 |
| PGK1_HUMA | PGK1 | sp\|P00558\|PGK1_HUMAN Phosphoglycerate kinase 1 OS=Homo sapiens GN=PGK1 PE=1 | 1177,505 | 73,14 | 44615 | 30 | 28,61 | 28,61 | 27,76 | 28,66 | 28,74 | 28,09 | 1,125 | 6,52E-01 | 0 | 0,16999944 | 0,18566426 |
| CUL1_HUMA | CUL1 | sp\|Q13616\|CUL1_HUMAN Cullin-1 OS=Homo sapiens GN=CUL1 PE=1 SV=2 | 215,362 | 11,6 | 89679 | 7 | 24,34 | 24,09 | 23,01 | 24,14 | 24,24 | 23,57 | 1,125 | 7,30E-01 | 0 | 0,16999944 | 0,13695914 |
| RS9_HUMAN | RPS9 | sp\|P46781\|RS9_HUMAN 40S ribosomal protein S9 OS=Homo sapiens GN=RPS9 PE=1 SV | 322,661 | 45,88 | 22591 | 15 | 26,92 | 27,65 | 27,06 | 27,10 | 27,19 | 27,84 | 1,122 | 6,33E-01 | 0 | 0,1666673 | 0,19849706 |
| ANXA4_HUM | ANXA4 | sp\|P09525\|ANXA4_HUMAN Annexin A4 OS=Homo sapiens GN=ANXA4 PE=1 SV=4 | 762,341 | 47,96 | 35883 | 17 | 27,91 | 27,73 | 26,07 | 28,15 | 28,00 | 26,06 | 1,122 | 8,61E-01 | 0 | 0,16666667 | 0,06504555 |
| PEPL1_HUM | NPEPL1 | sp\|Q8NDH3\|PEPL1_HUMAN Probable aminopeptidase NPEPL1 OS=Homo sapiens GN=N | 56,112 | 4,59 | 55861 | 2 | 22,72 | 21,95 | 21,51 | 22,42 | 22,20 | 22,06 | 1,122 | 6,75E-01 | 0 | 0,16666667 | 0,17085835 |
| IF4A1_HUM | EIF4A1 | sp\|P60842\|IF4A1_HUMAN Eukaryotic initiation factor 4A-I OS=Homo sapiens GN=EIF4A | 2652,046 | 83,99 | 46154 | 51 | 28,76 | 28,70 | 29,56 | 28,71 | 28,99 | 29,82 | 1,122 | 7,20E-01 | 0 | 0,16666603 | 0,14254775 |
| ATLA3_HUM | ATL3 | sp\|Q6DD88\|ATLA3_HUMAN Atlastin-3 OS=Homo sapiens GN=ATL3 PE=1 SV=1 | 503,139 | 27,36 | 60542 | 14 | 26,04 | 26,14 | 26,57 | 26,05 | 26,55 | 26,65 | 1,122 | 5,36E-01 | 0 | 0,16666603 | 0,27050926 |
| AT131_HUM | ATP13A1 | sp\|Q9HD20\|AT131_HUMAN Manganese-transporting ATPase 13A1 OS=Homo sapiens G | 12,143 | 0,91 | 132955 | 1 | 19,70 | 17,97 | 20,79 | 19,86 | 18,73 | 20,37 | 1,122 | 8,70E-01 | 0 | 0,16599909 | 0,06040263 |
| DDX42_HUM | DDX42 | sp\|Q86XP3\|DDX42_HUMAN ATP-dependent RNA helicase DDX42 OS=Homo sapiens GN | 67,666 | 3,3 | 102975 | 2 | 18,31 | 18,64 | 18,95 | 18,42 | 18,61 | 19,36 | 1,120 | 6,57E-01 | 0 | 0,16333389 | 0,18227806 |
| ANX11_HUM | ANXA11 | sp\|P50995\|ANX11_HUMAN Annexin A11 OS=Homo sapiens GN=ANXA11 PE=1 SV=1 | 557,799 | 26,53 | 54390 | 16 | 28,02 | 27,96 | 26,31 | 28,42 | 28,10 | 26,26 | 1,120 | 8,61E-01 | 0 | 0,16333389 | 0,06493704 |
| NID1_HUMA | NID1 | sp\|P14543\|NID1_HUMAN Nidogen-1 OS=Homo sapiens GN=NID1 PE=1 SV=3 | 161,554 | 4,41 | 136377 | 5 | 25,15 | 22,28 | 24,31 | 24,58 | 22,88 | 24,77 | 1,120 | 8,83E-01 | 0 | 0,16333326 | 0,05399581 |
| HNRPR_HUM | HNRNPR | sp\|O43390\|HNRPR_HUMAN Heterogeneous nuclear ribonucleoprotein R OS=Homo sapiens | 348,257 | 23,7 | 70943 | 8 | 22,99 | 23,62 | 25,41 | 23,49 | 23,61 | 25,41 | 1,120 | 8,72E-01 | 0 | 0,16333326 | 0,05926769 |
| DDAH2_HUM | DDAH2 | sp\|O95865\|DDAH2_HUMAN N(G),N(G)-dimethylarginine dimethylaminohydrolase 2 OS= | 59,326 | 13,33 | 29644 | 3 | 23,02 | 22,80 | 22,63 | 23,30 | 22,72 | 22,92 | 1,120 | 4,68E-01 | 0 | 0,16333326 | 0,32929355 |
| UBE2O_HUM | UBE2O | sp\|Q9C0C9\|UBE2O_HUMAN (E3-independent) E2 ubiquitin-conjugating enzyme OS=Ho | 48,882 | 2,4 | 141293 | 2 | 20,19 | 20,21 | 21,20 | 20,39 | 20,12 | 21,58 | 1,120 | 7,85E-01 | 0 | 0,16333326 | 0,10534772 |
| VTDB_HUMA | GC | sp\|P02774\|VTDB_HUMAN Vitamin D-binding protein OS=Homo sapiens GN=GC PE=1 S | 191,211 | 12,03 | 52964 | 4 | 22,42 | 21,00 | 24,99 | 22,88 | 21,91 | 24,01 | 1,119 | 9,07E-01 | 0 | 0,16333263 | 0,04230247 |
| PRDX1_HUM | PRDX1 | sp\|Q06830\|PRDX1_HUMAN Peroxiredoxin-1 OS=Homo sapiens GN=PRDX1 PE=1 SV=1 | 522,717 | 46,73 | 22110 | 13 | 29,67 | 30,17 | 28,89 | 29,83 | 30,10 | 29,28 | 1,117 | 7,37E-01 | 0 | 0,16000048 | 0,13273015 |
| CNOT1_HUM | CNOT1 | sp\|A5YKK6\|CNOT1_HUMAN CCR4-NOT transcription complex subunit 1 OS=Homo sapiens | 137,71 | 3,16 | 266939 | 3 | 22,14 | 22,12 | 23,67 | 21,68 | 22,95 | 23,78 | 1,117 | 8,15E-01 | 0 | 0,16000048 | 0,07051929 |
| GMDS_HUM | GMDS | sp\|O60547\|GMDS_HUMAN GDP-mannose 4,6 dehydratase OS=Homo sapiens GN=GMD | 45,034 | 6,18 | 41950 | 2 | 20,79 | 20,80 | 20,71 | 21,51 | 21,03 | 20,24 | 1,117 | 6,89E-01 | 0 | 0,16000048 | 0,1619327 |
| EPHA2_HUM | EPHA2 | sp\|P29317\|EPHA2_HUMAN Ephrin type-A receptor 2 OS=Homo sapiens GN=EPHA2 PE= | 957,477 | 27,15 | 108266 | 23 | 27,67 | 28,10 | 27,22 | 28,17 | 28,17 | 27,13 | 1,117 | 7,29E-01 | 0 | 0,15999985 | 0,13752577 |
| LMNA_HUM | LMNA | sp\|P02545\|LMNA_HUMAN Prelamin-A/C OS=Homo sapiens GN=LMNA PE=1 SV=1 | 1255,868 | 49,4 | 74139 | 33 | 27,42 | 27,69 | 27,93 | 27,73 | 27,78 | 28,01 | 1,117 | 4,02E-01 | 0 | 0,15999985 | 0,39617905 |
| SMD3_HUM | SNRPD3 | sp\|P62318\|SMD3_HUMAN Small nuclear ribonucleoprotein Sm D3 OS=Homo sapiens GN | 133,776 | 31,75 | 13916 | 3 | 24,29 | 25,26 | 25,16 | 24,63 | 26,05 | 25,65 | 1,117 | 3,44E-01 | 0 | 0,15999985 | 0,46317902 |
| AR13B_HUM | ARL13B | sp\|Q3SXY8\|AR13B_HUMAN ADP-ribosylation factor-like protein 13B OS=Homo sapiens | 39,575 | 2,8 | 48643 | 1 | 20,42 | 20,54 | 22,42 | 21,05 | 21,10 | 21,71 | 1,117 | 8,76E-01 | 0 | 0,15999921 | 0,08307147 |
| DNPEP_HUM | DNPEP | sp\|Q9ULA0\|DNPEP_HUMAN Aspartyl aminopeptidase OS=Homo sapiens GN=DNPEP PE= | 33,78 | 5,26 | 52428 | 2 | 22,35 | 22,61 | 22,02 | 22,22 | 22,50 | 22,64 | 1,117 | 5,06E-01 | 0 | 0,15999921 | 0,29568468 |
| DHX16_HUM | DHX16 | sp\|O60231\|DHX16_HUMAN Pre-mRNA-splicing factor ATP-dependent RNA helicase DHX | 30,95 | 0,86 | 119264 | 1 | 18,45 | 19,29 | 22,23 | 18,76 | 19,25 | 22,43 | 1,116 | 9,28E-01 | 0 | 0,15676092 | 0,03257332 |
| TAGL2_HUM | TAGLN2 | sp\|P37802\|TAGL2_HUMAN Transgelin-2 OS=Homo sapiens GN=TAGLN2 PE=1 SV=3 | 738,842 | 64,32 | 22391 | 15 | 27,80 | 27,76 | 27,39 | 27,87 | 27,85 | 27,70 | 1,115 | 3,64E-01 | 0 | 0,15666771 | 0,48264032 |
| FKB10_HUM | FKBP10 | sp\|Q96AY3\|FKB10_HUMAN Peptidyl-prolyl cis-trans isomerase FKBP10 OS=Homo sapiens | 35,169 | 2,06 | 64245 | 1 | 21,26 | 20,78 | 23,05 | 21,34 | 21,78 | 22,44 | 1,115 | 8,47E-01 | 0 | 0,15666771 | 0,07213545 |
| EIF3L_HUM | EIF3L | sp\|Q9Y262\|EIF3L_HUMAN Eukaryotic translation initiation factor 3 subunit L OS=Homo | 808,714 | 37,23 | 66727 | 20 | 26,46 | 26,51 | 26,60 | 26,44 | 26,79 | 26,81 | 1,115 | 2,85E-01 | 0 | 0,15666771 | 0,54568243 |
| HNRH3_HUM | HNRNPH3 | sp\|P31942\|HNRH3_HUMAN Heterogeneous nuclear ribonucleoprotein H3 OS=Homo sa | 408,049 | 33,82 | 36926 | 10 | 24,21 | 24,56 | 26,60 | 24,14 | 24,93 | 25,97 | 1,115 | 8,91E-01 | 0 | 0,15666707 | 0,04988246 |
| SRSF3_HUM | SRSF3 | sp\|P84103\|SRSF3_HUMAN Serine/arginine-rich splicing factor 3 OS=Homo sapiens GN= | 185,79 | 30,49 | 19330 | 5 | 25,30 | 25,53 | 25,13 | 25,57 | 25,41 | 25,35 | 1,115 | 3,61E-01 | 0 | 0,15666707 | 0,44288079 |
| RL34_HUMA | RPL34 | sp\|P49207\|RL34_HUMAN 60S ribosomal protein L34 OS=Homo sapiens GN=RPL34 PE=1 | 63 | 13,68 | 13293 | 3 | 26,40 | 26,06 | 24,36 | 26,54 | 26,30 | 25,57 | 1,115 | 7,13E-01 | 0 | 0,15666707 | 0,14696223 |
| RS3_HUMAN | RPS3 | sp\|P23396\|RS3_HUMAN 40S ribosomal protein S3 OS=Homo sapiens GN=RPS3 PE=1 SV= | 707,588 | 70,37 | 26688 | 21 | 28,37 | 28,48 | 29,20 | 28,24 | 28,66 | 29,62 | 1,115 | 6,42E-01 | 0 | 0,15666644 | 0,11774343 |
| HLAG_HUMA | HLA-G | sp\|P17693\|HLAG_HUMAN HLA class I histocompatibility antigen, alpha chain G OS=Hom | 98,86 | 10,65 | 38224 | 3 | 22,89 | 21,54 | 22,13 | 23,28 | 22,02 | 21,72 | 1,112 | 8,16E-01 | 0 | 0,15333367 | 0,08827509 |
| RAB6B_HUM | RAB6B | sp\|Q9NRW1\|RAB6B_HUMAN Ras-related protein Rab-6B OS=Homo sapiens GN=RAB6B PE | 290,717 | 28,85 | 23462 | 8 | 24,26 | 24,53 | 23,64 | 24,43 | 24,62 | 23,82 | 1,112 | 6,99E-01 | 0 | 0,15333366 | 0,16173623 |
| ABCE1_HUM | ABCE1 | sp\|P61221\|ABCE1_HUMAN ATP-binding cassette sub-family E member 1 OS=Homo sapi | 739,598 | 48,58 | 67314 | 23 | 25,80 | 25,82 | 26,94 | 25,73 | 26,15 | 27,14 | 1,112 | 7,99E-01 | 0 | 0,15333303 | 0,09760349 |
| GFPT1_HUM | GFPT1 | sp\|Q06210\|GFPT1_HUMAN Glutamine--fructose-6-phosphate aminotransferase [isome | 874,267 | 36,48 | 78806 | 25 | 25,25 | 25,62 | 26,03 | 25,22 | 25,60 | 26,54 | 1,112 | 7,52E-01 | 0 | 0,15333303 | 0,12395508 |
| EIF3F_HUM | EIF3F | sp\|O00303\|EIF3F_HUMAN Eukaryotic translation initiation factor 3 subunit F OS=Homo | 523,212 | 54,06 | 37564 | 13 | 25,78 | 25,78 | 26,47 | 25,68 | 25,92 | 26,89 | 1,112 | 8,28E-01 | 0 | 0,15333303 | 0,12926047 |
| CYTS_HUMA | CST4 | sp\|P01036\|CYTS_HUMAN Cystatin-S OS=Homo sapiens GN=CST4 PE=1 SV=3 | 151,678 | 32,62 | 16214 | 3 | 19,19 | 18,72 | 19,53 | 22,59 | 17,56 | 17,75 | 1,111 | 9,32E-01 | 0 | 0,15133031 | 0,03062987 |
| RL27_HUMA | RPL27 | sp\|P61353\|RL27_HUMAN 60S ribosomal protein L27 OS=Homo sapiens GN=RPL27 PE=1 | 201,015 | 42,65 | 15798 | 8 | 26,82 | 26,75 | 26,73 | 26,96 | 26,79 | 27,07 | 1,110 | 9,85E-02 | 0 | 0,15000025 | 1,00656746 |
| PLD2_HUMA | PLD2 | sp\|O14939\|PLD2_HUMAN Phospholipase D2 OS=Homo sapiens GN=PLD2 PE=1 SV=2 | 249,661 | 17,9 | 105987 | 13 | 25,21 | 25,15 | 23,83 | 25,26 | 25,14 | 24,24 | 1,110 | 8,00E-01 | 0 | 0,15000025 | 0,09701283 |
| RS15_HUMA | RPS15 | sp\|P62841\|RS15_HUMAN 40S ribosomal protein S15 OS=Homo sapiens GN=RPS15 PE=1 | 422,145 | 55,86 | 17040 | 9 | 24,86 | 25,31 | 24,76 | 25,10 | 25,06 | 25,22 | 1,110 | 3,05E-01 | 0 | 0,14999962 | 0,35485336 |
| LDHB_HUMA | LDHB | sp\|P07195\|LDHB_HUMAN L-lactate dehydrogenase B chain OS=Homo sapiens GN=LDHE | 1267,111 | 78,14 | 36638 | 26 | 29,33 | 29,72 | 28,53 | 29,42 | 29,64 | 28,97 | 1,110 | 7,28E-01 | 0 | 0,14999962 | 0,13790289 |
| ATX2_HUMA | ATXN2 | sp\|Q99700\|ATX2_HUMAN Ataxin-2 OS=Homo sapiens GN=ATXN2 PE=1 SV=2 | 124,953 | 3,27 | 140283 | 3 | 22,70 | 23,11 | 20,07 | 22,71 | 22,49 | 21,13 | 1,110 | 8,96E-01 | 0 | 0,14999898 | 0,04789954 |

| | | | | | | | | | | | | | | | | | | |
|---|---|---|---|---|---|---|---|---|---|---|---|---|---|---|---|---|---|---|
| CISD2_HUM/ | CISD2 | sp\|Q8N5K1\|CISD2_HUMAN CDGSH iron-sulfur domain-containing protein 2 OS=Homo s | 32,846 | 8,89 | 15278 | 1 | 22,56 | 23,23 | 22,48 | 23,01 | 23,52 | 22,18 | 1,107 | 7,64E-01 | 0 | 0,14666748 | 0,11666727 |
| TEN3_HUMA | TENM3 | sp\|Q9P273\|TEN3_HUMAN Teneurin-3 OS=Homo sapiens GN=TENM3 PE=2 SV=3 | 102,785 | 1,89 | 300950 | 4 | 23,06 | 21,80 | 21,96 | 22,17 | 22,17 | 22,92 | 1,107 | 7,70E-01 | 0 | 0,14666748 | 0,11361862 |
| DX39B_HUM | DDX39B | sp\|Q13838\|DX39B_HUMAN Spliceosome RNA helicase DDX39B OS=Homo sapiens GN=D | 620,588 | 50,7 | 48991 | 20 | 27,60 | 27,38 | 27,92 | 27,24 | 27,60 | 28,50 | 1,107 | 7,36E-01 | 0 | 0,14666684 | 0,13294901 |
| RL23_HUMA | RPL23 | sp\|P62829\|RL23_HUMAN 60S ribosomal protein L23 OS=Homo sapiens GN=RPL23 PE=1 | 423,603 | 52,86 | 14865 | 11 | 26,35 | 26,64 | 27,31 | 26,38 | 26,45 | 27,91 | 1,107 | 8,11E-01 | 0 | 0,14666684 | 0,09100751 |
| PCNA_HUM/ | PCNA | sp\|P12004\|PCNA_HUMAN Proliferating cell nuclear antigen OS=Homo sapiens GN=PCN/ | 493,889 | 61,69 | 28769 | 12 | 26,47 | 26,69 | 26,64 | 26,72 | 26,71 | 26,81 | 1,107 | 1,18E-01 | 0 | 0,14666621 | 0,92900421 |
| SYNC_HUMA | NARS | sp\|O43776\|SYNC_HUMAN Asparagine--tRNA ligase, cytoplasmic OS=Homo sapiens GN= | 322,174 | 20,07 | 62943 | 9 | 24,85 | 24,94 | 25,34 | 24,80 | 25,17 | 25,60 | 1,107 | 6,23E-01 | 0 | 0,14666621 | 0,20541538 |
| AQR_HUMAI | AQR | sp\|O60306\|AQR_HUMAN Intron-binding protein aquarius OS=Homo sapiens GN=AQR P | 13,128 | 0,94 | 171295 | 1 | 19,35 | 19,63 | 21,13 | 19,04 | 19,86 | 21,64 | 1,104 | 8,87E-01 | 0 | 0,14333407 | 0,05214231 |
| AAAT_HUM/ | SLC1A5 | sp\|Q15758\|AAAT_HUMAN Neutral amino acid transporter B(0) OS=Homo sapiens GN=S | 66,55 | 4,25 | 56598 | 2 | 24,94 | 24,82 | 24,64 | 24,56 | 25,37 | 24,90 | 1,104 | 5,98E-01 | 0 | 0,14333344 | 0,22344358 |
| CPNE1_HUM | CPNE1 | sp\|Q99829\|CPNE1_HUMAN Copine-1 OS=Homo sapiens GN=CPNE1 PE=1 SV=1 | 151,351 | 11,55 | 59059 | 6 | 25,90 | 25,73 | 25,35 | 26,06 | 26,18 | 25,17 | 1,104 | 7,09E-01 | 0 | 0,14333344 | 0,14930793 |
| PEPD_HUMA | PEPD | sp\|P12955\|PEPD_HUMAN Xaa-Pro dipeptidase OS=Homo sapiens GN=PEPD PE=1 SV=3 | 44,506 | 2,03 | 54548 | 1 | 21,61 | 21,79 | 21,88 | 21,77 | 22,02 | 21,92 | 1,104 | 2,54E-01 | 0 | 0,14333344 | 0,59575122 |
| RS15A_HUM | RPS15A | sp\|P62244\|RS15A_HUMAN 40S ribosomal protein S15a OS=Homo sapiens GN=RPS15A F | 309,666 | 59,23 | 14840 | 8 | 27,19 | 27,23 | 26,78 | 27,21 | 27,52 | 26,90 | 1,104 | 5,66E-01 | 0 | 0,1433328 | 0,24697564 |
| ERF1_HUMA | ETF1 | sp\|P62495\|ERF1_HUMAN Eukaryotic peptide chain release factor subunit 1 OS=Homo sa | 510,444 | 38,9 | 49031 | 14 | 26,28 | 26,03 | 25,79 | 26,17 | 26,41 | 25,95 | 1,104 | 5,01E-01 | 0 | 0,1433328 | 0,30005746 |
| C1TC_HUMA | MTHFD1 | sp\|P11586\|C1TC_HUMAN C-1-tetrahydrofolate synthase, cytoplasmic OS=Homo sapiens | 513,838 | 23,96 | 101559 | 16 | 25,78 | 26,66 | 26,82 | 26,16 | 26,30 | 27,23 | 1,104 | 7,74E-01 | 0 | 0,1433328 | 0,11136474 |
| VIR_HUMAN | VIRMA | sp\|Q69YN4\|VIR_HUMAN Protein virilizer homolog OS=Homo sapiens GN=VIRMA PE=1 S | 79,732 | 1,55 | 202025 | 2 | 18,34 | 20,11 | 18,62 | 18,63 | 18,74 | 20,12 | 1,102 | 8,56E-01 | 0 | 0,14059957 | 0,06740226 |
| RL30_HUMA | RPL30 | sp\|P62888\|RL30_HUMAN 60S ribosomal protein L30 OS=Homo sapiens GN=RPL30 PE=1 | 252,318 | 44,35 | 12784 | 5 | 25,76 | 25,56 | 26,32 | 25,78 | 25,74 | 26,54 | 1,102 | 7,06E-01 | 0 | 0,14000066 | 0,15109132 |
| PI42A_HUM/ | PIP4K2A | sp\|P48426\|PI42A_HUMAN Phosphatidylinositol 5-phosphate 4-kinase type-2 alpha OS=I | 137,256 | 13,55 | 46225 | 5 | 21,99 | 20,87 | 22,49 | 22,55 | 21,54 | 21,68 | 1,102 | 8,19E-01 | 0 | 0,14000003 | 0,08659503 |
| PPIL1_HUM/ | PPIL1 | sp\|Q9Y3C6\|PPIL1_HUMAN Peptidyl-prolyl cis-trans isomerase-like 1 OS=Homo sapiens ( | 219,316 | 57,83 | 18237 | 7 | 25,85 | 25,78 | 23,92 | 25,86 | 25,71 | 24,40 | 1,102 | 8,67E-01 | 0 | 0,13999939 | 0,06201627 |
| PUR2_HUMA | GART | sp\|P22102\|PUR2_HUMAN Trifunctional purine biosynthetic protein adenosine-3 OS=Ho | 250,532 | 10,89 | 107767 | 9 | 25,44 | 25,56 | 25,42 | 25,55 | 25,63 | 25,66 | 1,102 | 6,26E-02 | 0 | 0,13999939 | 1,20352021 |
| BYST_HUMA | BYSL | sp\|Q13895\|BYST_HUMAN Bystin OS=Homo sapiens GN=BYSL PE=1 SV=3 | 52,843 | 9,84 | 49601 | 2 | 18,76 | 18,99 | 18,91 | 17,69 | 19,87 | 19,52 | 1,102 | 8,47E-01 | 0 | 0,13986397 | 0,07215709 |
| ANXA2_HUN | ANXA2 | sp\|P07355\|ANXA2_HUMAN Annexin A2 OS=Homo sapiens GN=ANXA2 PE=1 SV=2 | 2140,826 | 86,14 | 38604 | 49 | 33,12 | 33,46 | 32,66 | 33,30 | 33,54 | 32,81 | 1,099 | 6,88E-01 | 0 | 0,13666789 | 0,16259432 |
| MCM4_HUM | MCM4 | sp\|P33991\|MCM4_HUMAN DNA replication licensing factor MCM4 OS=Homo sapiens G | 611,112 | 21,44 | 96558 | 17 | 24,92 | 25,15 | 26,89 | 25,27 | 25,08 | 27,02 | 1,099 | 8,84E-01 | 0 | 0,13666725 | 0,05373196 |
| PFD6_HUMA | PFDN6 | sp\|O15212\|PFD6_HUMAN Prefoldin subunit 6 OS=Homo sapiens GN=PFDN6 PE=1 SV=1 | 12,875 | 6,2 | 14583 | 1 | 21,64 | 21,27 | 20,48 | 21,70 | 21,02 | 21,08 | 1,099 | 7,53E-01 | 0 | 0,13666725 | 0,1232488 |
| AP1B1_HUM | AP1B1 | sp\|Q10567\|AP1B1_HUMAN AP-1 complex subunit beta-1 OS=Homo sapiens GN=AP1B1 | 1166,108 | 27,5 | 104637 | 31 | 25,00 | 24,97 | 22,99 | 25,10 | 25,23 | 23,04 | 1,099 | 8,95E-01 | 0 | 0,13666725 | 0,04816953 |
| GLYC_HUMA | SHMT1 | sp\|P34896\|GLYC_HUMAN Serine hydroxymethyltransferase, cytosolic OS=Homo sapien | 65,154 | 5,59 | 53083 | 2 | 23,43 | 23,32 | 23,43 | 23,52 | 23,51 | 23,56 | 1,099 | 2,63E-02 | 0 | 0,13666662 | 1,58039994 |
| GPI8_HUMAI | PIGK | sp\|Q92643\|GPI8_HUMAN GPI-anchor transamidase OS=Homo sapiens GN=PIGK PE=1 SV | 26,413 | 2,78 | 45252 | 1 | 19,78 | 19,40 | 20,91 | 19,98 | 19,34 | 21,18 | 1,099 | 8,56E-01 | 0 | 0,13666662 | 0,06769934 |
| UBP5_HUMA | USP5 | sp\|P45974\|UBP5_HUMAN Ubiquitin carboxyl-terminal hydrolase 5 OS=Homo sapiens GI | 1042,446 | 40,91 | 95786 | 24 | 26,67 | 26,96 | 26,79 | 26,84 | 26,85 | 27,14 | 1,099 | 3,51E-01 | 0 | 0,13666662 | 0,45518193 |
| SYQ_HUMAN | QARS | sp\|P47897\|SYQ_HUMAN Glutamine--tRNA ligase OS=Homo sapiens GN=QARS PE=1 SV= | 290,286 | 12,39 | 87799 | 10 | 24,60 | 24,61 | 25,41 | 24,63 | 24,64 | 25,76 | 1,099 | 7,82E-01 | 0 | 0,13666598 | 0,10696056 |
| AB17B_HUM | ABHD17B | sp\|Q5VST6\|AB17B_HUMAN Protein ABHD17B OS=Homo sapiens GN=ABHD17B PE=1 SV | 67,065 | 14,24 | 32215 | 2 | 19,56 | 20,00 | 20,73 | 20,44 | 19,74 | 20,51 | 1,097 | 7,67E-01 | 0 | 0,13333384 | 0,11517316 |
| CDK6_HUMA | CDK6 | sp\|Q00534\|CDK6_HUMAN Cyclin-dependent kinase 6 OS=Homo sapiens GN=CDK6 PE=1 | 134,173 | 20,55 | 36938 | 5 | 23,63 | 23,74 | 22,85 | 23,66 | 23,61 | 23,35 | 1,097 | 6,76E-01 | 0 | 0,13333384 | 0,17012894 |
| ARP10_HUM | ACTR10 | sp\|Q9NZ32\|ARP10_HUMAN Actin-related protein 10 OS=Homo sapiens GN=ACTR10 PE= | 100,813 | 12,95 | 46307 | 4 | 22,63 | 22,81 | 23,81 | 21,82 | 23,91 | 23,92 | 1,097 | 8,74E-01 | 0 | 0,13333384 | 0,05849251 |
| COR1C_HUN | CORO1C | sp\|Q9ULV4\|COR1C_HUMAN Coronin-1C OS=Homo sapiens GN=CORO1C PE=1 SV=1 | 574,425 | 39,87 | 53249 | 15 | 27,08 | 26,85 | 26,44 | 27,08 | 27,10 | 26,59 | 1,097 | 6,23E-01 | 0 | 0,13333321 | 0,20552297 |
| ENPP1_HUM | ENPP1 | sp\|P22413\|ENPP1_HUMAN Ectonucleotide pyrophosphatase/phosphodiesterase family | 18,003 | 1,73 | 104924 | 1 | 21,34 | 20,98 | 21,34 | 21,23 | 21,41 | 21,42 | 1,097 | 3,79E-01 | 0 | 0,13333321 | 0,42127874 |
| EIF3I_HUMA | EIF3I | sp\|Q13347\|EIF3I_HUMAN Eukaryotic translation initiation factor 3 subunit I OS=Homo s | 662,202 | 68,31 | 36502 | 17 | 26,29 | 26,60 | 26,26 | 26,52 | 26,55 | 26,48 | 1,097 | 2,94E-01 | 0 | 0,13333257 | 0,53127093 |
| RS14_HUMA | RPS14 | sp\|P62263\|RS14_HUMAN 40S ribosomal protein S14 OS=Homo sapiens GN=RPS14 PE=1 | 419,348 | 68,21 | 16273 | 10 | 26,54 | 26,61 | 27,06 | 26,74 | 26,47 | 27,40 | 1,097 | 6,99E-01 | 0 | 0,13333257 | 0,15557251 |
| SORCN_HUN | SRI | sp\|P30626\|SORCN_HUMAN Sorcin OS=Homo sapiens GN=SRI PE=1 SV=1 | 132,986 | 27,27 | 21676 | 5 | 25,85 | 25,63 | 23,37 | 25,97 | 25,57 | 23,71 | 1,097 | 9,06E-01 | 0 | 0,13333257 | 0,04309912 |
| RL14_HUMA | RPL14 | sp\|P50914\|RL14_HUMAN 60S ribosomal protein L14 OS=Homo sapiens GN=RPL14 PE=1 | 164,26 | 15,81 | 23432 | 3 | 25,60 | 25,85 | 26,11 | 25,60 | 25,47 | 26,89 | 1,097 | 7,94E-01 | 0 | 0,13333257 | 0,10044866 |
| PFKAP_HUM | PFKP | sp\|Q01813\|PFKAP_HUMAN ATP-dependent 6-phosphofructokinase, platelet type OS=H | 511,539 | 14,29 | 85596 | 12 | 23,26 | 23,89 | 25,67 | 23,51 | 23,77 | 25,93 | 1,094 | 9,08E-01 | 0 | 0,13000043 | 0,04205281 |
| PPP6_HUMA | PPP6C | sp\|O00743\|PPP6_HUMAN Serine/threonine-protein phosphatase 6 catalytic subunit OS | 182,264 | 34,43 | 35144 | 7 | 23,48 | 23,83 | 23,20 | 23,13 | 23,82 | 23,95 | 1,094 | 6,99E-01 | 0 | 0,1299998 | 0,15540475 |
| CAPR1_HUM | CAPRIN1 | sp\|Q14444\|CAPR1_HUMAN Caprin-1 OS=Homo sapiens GN=CAPRIN1 PE=1 SV=2 | 111,584 | 6,06 | 78366 | 4 | 23,15 | 23,57 | 23,52 | 23,63 | 23,50 | 23,50 | 1,094 | 4,04E-01 | 0 | 0,1299998 | 0,39391485 |
| COG7_HUM/ | COG7 | sp\|P83436\|COG7_HUMAN Conserved oligomeric Golgi complex subunit 7 OS=Homo sap | 28,166 | 2,21 | 86344 | 1 | 17,48 | 17,76 | 19,02 | 17,94 | 17,89 | 18,82 | 1,094 | 8,28E-01 | 0 | 0,1299998 | 0,08179432 |
| ERF3B_HUM | GSPT2 | sp\|Q8IYD1\|ERF3B_HUMAN Eukaryotic peptide chain release factor GTP-binding subunit | 399,08 | 25,32 | 68883 | 14 | 22,29 | 22,25 | 22,38 | 22,48 | 22,85 | 21,98 | 1,094 | 6,37E-01 | 0 | 0,1299998 | 0,19584478 |
| PGTA_HUM/ | RABGGTA | sp\|Q92696\|PGTA_HUMAN Geranylgeranyl transferase type-2 subunit alpha OS=Homo s | 55,574 | 5,29 | 65072 | 3 | 20,99 | 22,12 | 20,24 | 21,05 | 22,43 | 20,26 | 1,094 | 8,84E-01 | 0 | 0,1299998 | 0,05349917 |
| TM50B_HUN | TMEM50B | sp\|P56557\|TM50B_HUMAN Transmembrane protein 50B OS=Homo sapiens GN=TMEM | 26,131 | 6,96 | 17936 | 1 | 20,59 | 20,18 | 17,54 | 20,29 | 20,26 | 18,14 | 1,093 | 9,19E-01 | 0 | 0,12892087 | 0,03663401 |
| MAOX_HUM | ME1 | sp\|P48163\|MAOX_HUMAN NADP-dependent malic enzyme OS=Homo sapiens GN=ME1 | 27,899 | 1,92 | 64150 | 1 | 20,82 | 20,47 | 19,73 | 21,02 | 20,63 | 19,75 | 1,092 | 8,10E-01 | 0 | 0,12666702 | 0,09130851 |
| GMPPB_HUN | GMPPB | sp\|Q9Y5P6\|GMPPB_HUMAN Mannose-1-phosphate guanyltransferase beta OS=Homo s | 81,731 | 9,72 | 39834 | 3 | 23,46 | 23,63 | 23,29 | 23,49 | 23,44 | 23,83 | 1,092 | 4,65E-01 | 0 | 0,12666702 | 0,3325649 |
| PDIA6_HUM/ | PDIA6 | sp\|Q15084\|PDIA6_HUMAN Protein disulfide-isomerase A6 OS=Homo sapiens GN=PDIA6 | 188,443 | 9,55 | 48121 | 4 | 24,57 | 24,13 | 25,97 | 24,85 | 24,82 | 25,38 | 1,092 | 8,39E-01 | 0 | 0,12666702 | 0,07631755 |
| KAD1_HUMA | AK1 | sp\|P00568\|KAD1_HUMAN Adenylate kinase isoenzyme 1 OS=Homo sapiens GN=AK1 PE= | 161,651 | 28,87 | 21635 | 5 | 24,25 | 24,33 | 23,12 | 24,35 | 24,48 | 23,25 | 1,092 | 8,30E-01 | 0 | 0,12666639 | 0,08102686 |
| RPA2_HUMA | POLR1B | sp\|Q9H9Y6\|RPA2_HUMAN DNA-directed RNA polymerase I subunit RPA2 OS=Homo sap | 42,162 | 1,06 | 128229 | 1 | 18,25 | 18,37 | 19,54 | 20,23 | 16,28 | 20,02 | 1,090 | 9,31E-01 | 0 | 0,12484233 | 0,03124861 |
| PEBP1_HUM | PEBP1 | sp\|P30086\|PEBP1_HUMAN Phosphatidylethanolamine-binding protein 1 OS=Homo sapi | 52,793 | 10,16 | 21057 | 2 | 26,12 | 25,74 | 24,14 | 25,73 | 26,12 | 24,52 | 1,089 | 8,81E-01 | 0 | 0,12333361 | 0,05491823 |
| NIT1_HUMAI | NIT1 | sp\|Q86X76\|NIT1_HUMAN Deaminated glutathione amidase OS=Homo sapiens GN=NIT1 | 93,233 | 7,03 | 35896 | 2 | 21,90 | 22,71 | 20,79 | 22,23 | 22,45 | 21,09 | 1,089 | 8,68E-01 | 0 | 0,12333361 | 0,06130427 |
| NPL4_HUMA | NPLOC4 | sp\|Q8TAT6\|NPL4_HUMAN Nuclear protein localization protein 4 homolog OS=Homo sap | 365,445 | 24,84 | 68120 | 13 | 25,46 | 25,60 | 24,55 | 25,42 | 25,64 | 24,92 | 1,089 | 7,69E-01 | 0 | 0,12333361 | 0,11417795 |
| CK068_HUM | C11orf68 | sp\|Q9H3H3\|CK068_HUMAN UPF0696 protein C11orf68 OS=Homo sapiens GN=C11orf68 | 16,259 | 4,38 | 27355 | 1 | 21,12 | 21,19 | 18,88 | 20,95 | 20,89 | 19,72 | 1,089 | 8,93E-01 | 0 | 0,12333298 | 0,04933087 |
| TBB6_HUMA | TUBB6 | sp\|Q9BUF5\|TBB6_HUMAN Tubulin beta-6 chain OS=Homo sapiens GN=TUBB6 PE=1 SV= | 1343,956 | 69,28 | 49857 | 34 | 26,41 | 26,79 | 27,52 | 26,42 | 26,98 | 27,69 | 1,089 | 8,14E-01 | 0 | 0,12333298 | 0,08934523 |
| WDR91_HUN | WDR91 | sp\|A4D1P6\|WDR91_HUMAN WD repeat-containing protein 91 OS=Homo sapiens GN=W | 21,968 | 1,61 | 83344 | 1 | 19,75 | 19,14 | 20,42 | 19,64 | 19,61 | 20,42 | 1,088 | 8,03E-01 | 0 | 0,12122663 | 0,09507276 |
| LRC57_HUM | LRRC57 | sp\|Q8N9N7\|LRC57_HUMAN Leucine-rich repeat-containing protein 57 OS=Homo sapier | 104,22 | 13,39 | 26754 | 3 | 24,46 | 25,15 | 23,70 | 24,61 | 24,91 | 24,15 | 1,087 | 8,12E-01 | 0 | 0,1200002 | 0,09021864 |
| VATA_HUMA | ATP6V1A | sp\|P38606\|VATA_HUMAN V-type proton ATPase catalytic subunit A OS=Homo sapiens ( | 329,793 | 31,44 | 68304 | 12 | 25,39 | 25,11 | 24,64 | 25,44 | 25,42 | 24,64 | 1,087 | 7,44E-01 | 0 | 0,1200002 | 0,1286116 |
| VPS29_HUM | VPS29 | sp\|Q9UBQ0\|VPS29_HUMAN Vacuolar protein sorting-associated protein 29 OS=Homo s | 257,424 | 47,8 | 20506 | 8 | 24,68 | 24,71 | 24,90 | 24,55 | 24,80 | 25,30 | 1,087 | 6,31E-01 | 0 | 0,11999957 | 0,20008693 |
| AL7A1_HUM | ALDH7A1 | sp\|P49419\|AL7A1_HUMAN Alpha-aminoadipic semialdehyde dehydrogenase OS=Homo | 210,139 | 17,25 | 58487 | 8 | 23,97 | 24,04 | 24,44 | 24,11 | 23,99 | 24,70 | 1,084 | 6,81E-01 | 0 | 0,11666679 | 0,16676438 |
| AHSA1_HUM | AHSA1 | sp\|O95433\|AHSA1_HUMAN Activator of 90 kDa heat shock protein ATPase homolog 1 O | 184,812 | 25,44 | 38274 | 8 | 25,57 | 25,48 | 26,09 | 25,82 | 25,74 | 25,93 | 1,084 | 5,87E-01 | 0 | 0,11666679 | 0,23116974 |
| CLC11_HUM | CLEC11A | sp\|Q9Y240\|CLC11_HUMAN C-type lectin domain family 11 member A OS=Homo sapiens | 85,487 | 8,67 | 35695 | 2 | 22,19 | 20,03 | 22,05 | 22,50 | 20,43 | 21,69 | 1,084 | 9,05E-01 | 0 | 0,11666679 | 0,04316418 |
| TBCD_HUMA | TBCD | sp\|Q9BTW9\|TBCD_HUMAN Tubulin-specific chaperone D OS=Homo sapiens GN=TBCD F | 173,589 | 5,54 | 132600 | 6 | 24,26 | 24,20 | 24,30 | 24,16 | 24,45 | 24,50 | 1,084 | 3,48E-01 | 0 | 0,11666679 | 0,45808499 |
| CNOT9_HUN | CNOT9 | sp\|Q92600\|CNOT9_HUMAN CCR4-NOT transcription complex subunit 9 OS=Homo sapie | 23,533 | 6,69 | 33631 | 2 | 22,02 | 22,31 | 22,43 | 21,97 | 22,37 | 22,77 | 1,084 | 6,78E-01 | 0 | 0,11666679 | 0,16873028 |
| DPYL2_HUM | DPYSL2 | sp\|Q16555\|DPYL2_HUMAN Dihydropyrimidinase-related protein 2 OS=Homo sapiens GI | 1069,504 | 65,56 | 62294 | 23 | 27,10 | 27,03 | 26,95 | 27,18 | 27,27 | 26,98 | 1,084 | 2,91E-01 | 0 | 0,11666616 | 0,53576883 |
| IF1AX_HUM/ | EIF1AX | sp\|P47813\|IF1AX_HUMAN Eukaryotic translation initiation factor 1A, X-chromosomal O: | 290,392 | 43,75 | 16460 | 6 | 25,25 | 25,31 | 23,87 | 25,23 | 24,98 | 24,57 | 1,084 | 8,30E-01 | 0 | 0,11666616 | 0,08109419 |
| PIGT_HUMAI | PIGT | sp\|Q969N2\|PIGT_HUMAN GPI transamidase component PIG-T OS=Homo sapiens GN=PI | 40,512 | 2,6 | 65700 | 1 | 19,27 | 19,55 | 20,91 | 19,23 | 19,47 | 21,37 | 1,082 | 9,00E-01 | 0 | 0,11350187 | 0,04590823 |
| GTR1_HUMA | SLC2A1 | sp\|P11166\|GTR1_HUMAN Solute carrier family 2, facilitated glucose transporter membe | 322,222 | 16,67 | 54084 | 7 | 27,19 | 26,80 | 25,05 | 26,92 | 26,90 | 25,56 | 1,082 | 8,94E-01 | 0 | 0,11333338 | 0,04874762 |
| DJB11_HUM | DNAJB11 | sp\|Q9UBS4\|DJB11_HUMAN DnaJ homolog subfamily B member 11 OS=Homo sapiens GI | 89,735 | 4,47 | 40514 | 1 | 20,76 | 20,51 | 21,19 | 21,17 | 20,70 | 20,93 | 1,082 | 6,62E-01 | 0 | 0,11333338 | 0,17912092 |
| PERP_HUMA | PERP | sp\|Q96FX8\|PERP_HUMAN p53 apoptosis effector related to PMP-22 OS=Homo sapiens ( | 37,051 | 8,81 | 21386 | 1 | 22,61 | 21,99 | 23,41 | 22,17 | 23,09 | 23,09 | 1,082 | 8,36E-01 | 0 | 0,11333338 | 0,07783894 |
| ACOC_HUM/ | ACO1 | sp\|P21399\|ACOC_HUMAN Cytoplasmic aconitate hydratase OS=Homo sapiens GN=ACO | 174,415 | 10,57 | 98399 | 7 | 25,01 | 24,36 | 23,84 | 24,88 | 24,61 | 24,06 | 1,082 | 7,99E-01 | 0 | 0,11333275 | 0,09766875 |
| SMD1_HUM/ | SNRPD1 | sp\|P62314\|SMD1_HUMAN Small nuclear ribonucleoprotein Sm D1 OS=Homo sapiens GI | 221,697 | 54,62 | 13282 | 4 | 25,11 | 25,06 | 24,87 | 25,14 | 25,17 | 25,07 | 1,082 | 2,24E-01 | 0 | 0,11333275 | 0,64947124 |
| GPT_HUMAN | DPAGT1 | sp\|Q9H3H5\|GPT_HUMAN UDP-N-acetylglucosamine--dolichyl-phosphate N-acetylgluco: | 28,728 | 2,7 | 46090 | 1 | 19,51 | 18,28 | 20,37 | 18,81 | 18,83 | 20,85 | 1,081 | 9,07E-01 | 0 | 0,11302884 | 0,04237161 |
| K2C3_HUMA | KRT3 | sp\|P12035\|K2C3_HUMAN Keratin, type II cytoskeletal 3 OS=Homo sapiens GN=KRT3 PE= | 365,583 | 12,74 | 64417 | 12 | 19,18 | 19,34 | 17,26 | 20,01 | 18,57 | 17,54 | 1,080 | 9,15E-01 | 0 | 0,11150551 | 0,03859444 |
| SMC4_HUM/ | SMC4 | sp\|Q9NTJ3\|SMC4_HUMAN Structural maintenance of chromosomes protein 4 OS=Hom | 243,157 | 9,01 | 147182 | 8 | 23,09 | 22,38 | 24,44 | 23,34 | 22,61 | 24,29 | 1,079 | 8,94E-01 | 0 | 0,11000061 | 0,04862672 |
| QOR_HUMAI | CRYZ | sp\|Q08257\|QOR_HUMAN Quinone oxidoreductase OS=Homo sapiens GN=CRYZ PE=1 SV | 40,738 | 7,6 | 35207 | 2 | 23,63 | 23,26 | 21,69 | 23,77 | 23,50 | 21,64 | 1,079 | 9,08E-01 | 0 | 0,10999997 | 0,04183671 |
| PTN11_HUM | PTPN11 | sp\|Q06124\|PTN11_HUMAN Tyrosine-protein phosphatase non-receptor type 11 OS=Hc | 373,893 | 28,81 | 68436 | 14 | 25,54 | 25,66 | 23,97 | 25,64 | 25,48 | 24,38 | 1,079 | 8,78E-01 | 0 | 0,10999934 | 0,05643807 |
| CAN2_HUM/ | CAPN2 | sp\|P17655\|CAN2_HUMAN Calpain-2 catalytic subunit OS=Homo sapiens GN=CAPN2 PE= | 448,497 | 21 | 79995 | 14 | 25,88 | 26,52 | 26,92 | 26,05 | 26,64 | 26,96 | 1,079 | 7,99E-01 | 0 | 0,10999934 | 0,09766085 |
| NPA1P_HUM | URB1 | sp\|O60287\|NPA1P_HUMAN Nucleolar pre-ribosomal-associated protein 1 OS=Homo sa | 18,04 | 0,44 | 254389 | 1 | 19,04 | 19,99 | 19,96 | 20,00 | 19,27 | 20,04 | 1,077 | 8,01E-01 | 0 | 0,10765139 | 0,09614057 |
| ZDH18_HUM | ZDHHC18 | sp\|Q9NUE0\|ZDH18_HUMAN Palmitoyltransferase ZDHHC18 OS=Homo sapiens GN=ZDH | 10,625 | 2,84 | 42031 | 1 | 21,18 | 21,07 | 20,08 | 21,10 | 20,64 | 20,91 | 1,077 | 7,89E-01 | 0 | 0,10698255 | 0,10278383 |
| CPNE3_HUM | CPNE3 | sp\|O75131\|CPNE3_HUMAN Copine-3 OS=Homo sapiens GN=CPNE3 PE=1 SV=1 | 528,8 | 36,5 | 60131 | 12 | 26,55 | 25,69 | 25,55 | 26,66 | 26,45 | 25,00 | 1,077 | 8,69E-01 | 0 | 0,1066672 | 0,06081556 |
| THIO_HUMA | TXN | sp\|P10599\|THIO_HUMAN Thioredoxin OS=Homo sapiens GN=TXN PE=1 SV=3 | 26,516 | 8,57 | 11737 | 1 | 26,38 | 26,43 | 23,75 | 26,49 | 26,49 | 23,90 | 1,077 | 9,35E-01 | 0 | 0,10666656 | 0,02900348 |
| TRY1_HUMA | PRSS1 | sp\|P07477\|TRY1_HUMAN Trypsin-1 OS=Homo sapiens GN=PRSS1 PE=1 SV=1 | 41,394 | 8,1 | 26558 | 1 | 21,48 | 26,03 | 30,25 | 21,99 | 26,44 | 29,65 | 1,077 | 9,76E-01 | 0 | 0,10666656 | 0,01043743 |
| MCM6_HUM | MCM6 | sp\|Q14566\|MCM6_HUMAN DNA replication licensing factor MCM6 OS=Homo sapiens G | 696,602 | 33,62 | 92889 | 25 | 26,24 | 26,16 | 27,08 | 26,24 | 26,21 | 27,35 | 1,077 | 8,34E-01 | 0 | 0,10666656 | 0,07887554 |
| RS18_HUMA | RPS18 | sp\|P62269\|RS18_HUMAN 40S ribosomal protein S18 OS=Homo sapiens GN=RPS18 PE=1 | 334,259 | 49,34 | 17719 | 11 | 27,16 | 27,40 | 27,11 | 27,41 | 27,06 | 27,52 | 1,077 | 5,53E-01 | 0 | 0,10666656 | 0,25701094 |
| LGAT1_HUM | LPGAT1 | sp\|Q92604\|LGAT1_HUMAN Acyl-CoA:lysophosphatidylglycerol acyltransferase 1 OS=Ho | 33,758 | 2,7 | 43089 | 1 | 17,40 | 16,59 | 18,83 | 17,81 | 17,07 | 18,26 | 1,077 | 8,93E-01 | 0 | 0,10666656 | 0,04938755 |
| H2AX_HUM/ | H2AFX | sp\|P16104\|H2AX_HUMAN Histone H2AX OS=Homo sapiens GN=H2AFX PE=1 SV=2 | 195,283 | 52,45 | 15145 | 5 | 20,74 | 21,69 | 21,36 | 21,96 | 21,09 | 21,06 | 1,077 | 8,06E-01 | 0 | 0,10666593 | 0,09386421 |
| CHMP7_HUN | CHMP7 | sp\|Q8WUX9\|CHMP7_HUMAN Charged multivesicular body protein 7 OS=Homo sapiens | 112,244 | 8,39 | 50911 | 4 | 19,78 | 19,85 | 21,87 | 20,64 | 20,06 | 21,12 | 1,077 | 8,94E-01 | 0 | 0,10666593 | 0,04872133 |
| HSPB7_HUM | HSPB7 | sp\|Q9UBY9\|HSPB7_HUMAN Heat shock protein beta-7 OS=Homo sapiens GN=HSPB7 PE | 36,036 | 7,65 | 18611 | 2 | 18,89 | 19,31 | 16,65 | 19,69 | 18,83 | 16,64 | 1,074 | 9,37E-01 | 0 | 0,10333379 | 0,02828808 |
| RS2_HUMAN | RPS2 | sp\|P15880\|RS2_HUMAN 40S ribosomal protein S2 OS=Homo sapiens GN=RPS2 PE=1 SV= | 470,263 | 39,59 | 31324 | 12 | 26,31 | 27,31 | 26,74 | 26,68 | 26,63 | 27,36 | 1,074 | 7,96E-01 | 0 | 0,10333379 | 0,09929949 |
| BPNT1_HUM | BPNT1 | sp\|O95861\|BPNT1_HUMAN 3'(2'),5'-bisphosphate nucleotidase 1 OS=Homo sapiens GN | 54,662 | 6,17 | 33392 | 2 | 23,65 | 23,71 | 22,87 | 23,53 | 23,78 | 23,23 | 1,074 | 7,58E-01 | 0 | 0,10333379 | 0,12006743 |
| EFHD2_HUM | EFHD2 | sp\|Q96C19\|EFHD2_HUMAN EF-hand domain-containing protein D2 OS=Homo sapiens G | 136,889 | 25,42 | 26697 | 6 | 24,37 | 24,44 | 23,23 | 24,55 | 24,62 | 23,18 | 1,074 | 8,74E-01 | 0 | 0,10333316 | 0,05851425 |
| ALDOA_HUN | ALDOA | sp\|P04075\|ALDOA_HUMAN Fructose-bisphosphate aldolase A OS=Homo sapiens GN=Al | 1227,164 | 63,74 | 39420 | 27 | 29,38 | 29,53 | 28,40 | 29,48 | 29,43 | 28,71 | 1,074 | 8,23E-01 | 0 | 0,10333316 | 0,08455893 |
| EIF2A_HUM/ | EIF2A | sp\|Q9BY44\|EIF2A_HUMAN Eukaryotic translation initiation factor 2A OS=Homo sapiens | 33,224 | 3,25 | 64990 | 2 | 22,64 | 22,45 | 21,16 | 22,73 | 22,65 | 21,18 | 1,074 | 8,87E-01 | 0 | 0,10333316 | 0,05183687 |
| CAHM2_HUN | CALHM2 | sp\|Q9HA72\|CAHM2_HUMAN Calcium homeostasis modulator protein 2 OS=Homo sapie | 118,85 | 9,6 | 36175 | 3 | 22,36 | 21,86 | 23,65 | 21,90 | 22,80 | 23,48 | 1,074 | 8,90E-01 | 0 | 0,10333252 | 0,05051299 |
| ML12A_HUN | MYL12A | sp\|P19105\|ML12A_HUMAN Myosin regulatory light chain 12A OS=Homo sapiens GN=M | 506,191 | 56,14 | 19794 | 10 | 25,04 | 25,47 | 25,49 | 25,36 | 25,68 | 25,26 | 1,072 | 6,33E-01 | 0 | 0,10000038 | 0,19845865 |
| TIPRL_HUM/ | TIPRL | sp\|O75663\|TIPRL_HUMAN TIP41-like protein OS=Homo sapiens GN=TIPRL PE=1 SV=2 | 126,246 | 19,12 | 31444 | 4 | 22,72 | 23,41 | 22,96 | 22,88 | 23,02 | 23,49 | 1,072 | 7,33E-01 | 0 | 0,10000038 | 0,13467336 |
| PCDGD_HUN | PCDHGB1 | sp\|Q9Y5G3\|PCDGD_HUMAN Protocadherin gamma-B1 OS=Homo sapiens GN=PCDHGB1 | 121,912 | 4,53 | 100360 | 3 | 25,69 | 23,66 | 23,85 | 25,75 | 24,52 | 23,23 | 1,072 | 9,23E-01 | 0 | 0,09999975 | 0,03472633 |
| IF4A2_HUM/ | EIF4A2 | sp\|Q14240\|IF4A2_HUMAN Eukaryotic initiation factor 4A-II OS=Homo sapiens GN=EIF4/ | 852,193 | 41,03 | 46402 | 19 | 26,30 | 26,50 | 26,52 | 26,50 | 26,55 | 26,57 | 1,072 | 2,44E-01 | 0 | 0,09999975 | 0,61265071 |
| AGRIN_HUM | AGRN | sp\|O00468\|AGRIN_HUMAN Agrin OS=Homo sapiens GN=AGRN PE=1 SV=5 | 34,03 | 0,82 | 217232 | 2 | 24,19 | 24,57 | 22,39 | 24,79 | 24,40 | 22,25 | 1,069 | 9,30E-01 | 0 | 0,09666697 | 0,03141233 |
| KLC1_HUMA | KLC1 | sp\|Q07866\|KLC1_HUMAN Kinesin light chain 1 OS=Homo sapiens GN=KLC1 PE=1 SV=2 | 572,768 | 36,3 | 65310 | 17 | 24,53 | 24,82 | 24,80 | 24,65 | 24,78 | 25,01 | 1,069 | 5,30E-01 | 0 | 0,09666697 | 0,2756676 |
| HYCCI_HUM/ | FAM126A | sp\|Q9BYI3\|HYCCI_HUMAN Hyccin OS=Homo sapiens GN=FAM126A PE=1 SV=2 | 111,606 | 9,02 | 57625 | 5 | 24,03 | 23,83 | 22,71 | 23,85 | 23,93 | 23,08 | 1,069 | 8,54E-01 | 0 | 0,09666697 | 0,06862398 |
| KAP0_HUMA | PRKAR1A | sp\|P10644\|KAP0_HUMAN cAMP-dependent protein kinase type I-alpha regulatory subu | 237,659 | 29,4 | 42982 | 9 | 25,11 | 24,32 | 25,38 | 25,03 | 24,93 | 25,14 | 1,069 | 7,80E-01 | 0 | 0,09666697 | 0,10783105 |
| PRAF3_HUM | ARL6IP5 | sp\|O75915\|PRAF3_HUMAN PRA1 family protein 3 OS=Homo sapiens GN=ARL6IP5 PE=1 ! | 110,883 | 14,36 | 21615 | 4 | 25,18 | 25,63 | 25,54 | 24,98 | 25,95 | 25,71 | 1,069 | 7,79E-01 | 0 | 0,09666634 | 0,10830845 |
| SC24C_HUM | SEC24C | sp\|P53992\|SC24C_HUMAN Protein transport protein Sec24C OS=Homo sapiens GN=SEC | 283,342 | 13,62 | 118325 | 11 | 24,79 | 24,92 | 25,07 | 24,80 | 25,07 | 25,20 | 1,069 | 5,36E-01 | 0 | 0,09666634 | 0,27094615 |
| SGTA_HUM/ | SGTA | sp\|O43765\|SGTA_HUMAN Small glutamine-rich tetratricopeptide repeat-containing pro | 149,822 | 11,18 | 34063 | 3 | 23,19 | 24,15 | 23,41 | 23,72 | 23,61 | 23,71 | 1,069 | 7,58E-01 | 0 | 0,09666634 | 0,12055766 |
| NAA40_HUN | NAA40 | sp\|Q86UY6\|NAA40_HUMAN N-alpha-acetyltransferase 40 OS=Homo sapiens GN=NAA40 | 45,086 | 14,77 | 27194 | 3 | 20,23 | 20,60 | 21,79 | 20,34 | 20,89 | 21,68 | 1,069 | 8,82E-01 | 0 | 0,09666634 | 0,05459951 |
| MGN_HUMA | MAGOH | sp\|P61326\|MGN_HUMAN Protein mago nashi homolog OS=Homo sapiens GN=MAGOH | 325,103 | 84,93 | 17164 | 11 | 26,03 | 25,53 | 26,37 | 25,82 | 25,85 | 26,55 | 1,069 | 7,91E-01 | 0 | 0,0966657 | 0,10183904 |
| TCAF1_HUM | TCAF1 | sp\|Q9Y4C2\|TCAF1_HUMAN TRPM8 channel-associated factor 1 OS=Homo sapiens GN=T | 25,63 | 1,09 | 102126 | 1 | 20,36 | 17,21 | 21,63 | 20,35 | 18,32 | 20,82 | 1,068 | 9,53E-01 | 0 | 0,09551748 | 0,0209191 |
| CDK14_HUM | CDK14 | sp\|O94921\|CDK14_HUMAN Cyclin-dependent kinase 14 OS=Homo sapiens GN=CDK14 F | 28,919 | 3,41 | 53057 | 2 | 19,67 | 19,33 | 20,12 | 20,46 | 19,14 | 19,81 | 1,068 | 8,40E-01 | 0 | 0,09539477 | 0,07560341 |
| 5NT3B_HUM | NT5C3B | sp\|Q969T7\|5NT3B_HUMAN 7-methylguanosine phosphate-specific 5'-nucleotidase OS= | 62,487 | 10,67 | 34389 | 3 | 22,85 | 22,57 | 22,35 | 22,77 | 22,77 | 22,51 | 1,067 | 6,09E-01 | 0 | 0,09333356 | 0,21505202 |
| UBC9_HUMA | UBE2I | sp\|P63279\|UBC9_HUMAN SUMO-conjugating enzyme UBC9 OS=Homo sapiens GN=UBE | 82,297 | 22,78 | 18007 | 3 | 23,81 | 24,07 | 23,51 | 23,75 | 23,86 | 24,06 | 1,067 | 6,41E-01 | 0 | 0,09333356 | 0,19293343 |
| SYHC_HUMA | HARS | sp\|P12081\|SYHC_HUMAN Histidine--tRNA ligase, cytoplasmic OS=Homo sapiens GN=HA | 422,753 | 37,33 | 57411 | 15 | 25,40 | 26,03 | 24,28 | 25,33 | 25,87 | 24,79 | 1,067 | 8,84E-01 | 0 | 0,09333356 | 0,05366075 |
| SMCO1_HUN | SMCO1 | sp\|Q147U7\|SMCO1_HUMAN Single-pass membrane and coiled-coil domain-containing | | 11,239 | 3,27 | 24598 | 1 | 25,34 | 23,19 | 23,53 | 24,75 | 23,44 | 24,15 | 1,067 | 9,09E-01 | 0 | 0,09333293 | 0,0414178 |
| DYLT1_HUM | DYNLT1 | sp\|P63172\|DYLT1_HUMAN Dynein light chain Tctex-type 1 OS=Homo sapiens GN=DYNL | 136,328 | 51,33 | 12452 | 3 | 22,98 | 22,59 | 21,61 | 22,74 | 22,81 | 21,91 | 1,067 | 8,61E-01 | 0 | 0,09333293 | 0,06504528 |
| BAG2_HUM/ | BAG2 | sp\|O95816\|BAG2_HUMAN BAG family molecular chaperone regulator 2 OS=Homo sapie | 37,841 | 5,21 | 23772 | 1 | 21,14 | 21,05 | 22,12 | 21,03 | 21,43 | 22,12 | 1,064 | 8,57E-01 | 0 | 0,09000079 | 0,06712955 |
| CCNY_HUMA | CCNY | sp\|Q8ND76\|CCNY_HUMAN Cyclin-Y OS=Homo sapiens GN=CCNY PE=1 SV=2 | 76,15 | 7,62 | 39337 | 2 | 21,65 | 21,67 | 21,55 | 21,68 | 21,92 | 21,54 | 1,064 | 4,85E-01 | 0 | 0,09000079 | 0,31456666 |
| TBK1_HUMA | TBK1 | sp\|Q9UHD2\|TBK1_HUMAN Serine/threonine-protein kinase TBK1 OS=Homo sapiens GN | 43,65 | 7,82 | 83642 | 3 | 21,25 | 21,21 | 19,99 | 20,75 | 21,11 | 20,86 | 1,064 | 8,43E-01 | 0 | 0,09000079 | 0,07398578 |
| ARHG2_HUN | ARHGEF2 | sp\|Q92974\|ARHG2_HUMAN Rho guanine nucleotide exchange factor 2 OS=Homo sapie | 106,214 | 7,91 | 111543 | 5 | 21,93 | 22,85 | 22,82 | 22,20 | 22,90 | 22,77 | 1,064 | 8,20E-01 | 0 | 0,09000015 | 0,08616984 |
| CAP1_HUMA | CAP1 | sp\|Q01518\|CAP1_HUMAN Adenylyl cyclase-associated protein 1 OS=Homo sapiens GN= | 842,331 | 40,42 | 51901 | 23 | 27,48 | 27,52 | 26,63 | 27,44 | 27,58 | 26,88 | 1,064 | 8,15E-01 | 0 | 0,09000015 | 0,08875898 |
| RS27_HUMA | RPS27 | sp\|P42677\|RS27_HUMAN 40S ribosomal protein S27 OS=Homo sapiens GN=RPS27 PE=1 | 198,803 | 40,48 | 9461 | 5 | 24,28 | 24,47 | 23,67 | 24,53 | 24,07 | 24,09 | 1,064 | 7,67E-01 | 0 | 0,09000015 | 0,11502249 |
| PDIA1_HUM/ | P4HB | sp\|P07237\|PDIA1_HUMAN Protein disulfide-isomerase OS=Homo sapiens GN=P4HB PE= | 561,569 | 28,54 | 57116 | 14 | 27,24 | 26,91 | 28,11 | 27,54 | 27,46 | 27,53 | 1,064 | 8,14E-01 | 0 | 0,09000015 | 0,08921781 |
| RS12_HUMA | RPS12 | sp\|P25398\|RS12_HUMAN 40S ribosomal protein S12 OS=Homo sapiens GN=RPS12 PE=1 | 516,211 | 70,45 | 14515 | 10 | 26,87 | 27,31 | 26,53 | 27,06 | 27,12 | 26,80 | 1,064 | 7,33E-01 | 0 | 0,08999952 | 0,13476314 |
| FLNC_HUMA | FLNC | sp\|Q14315\|FLNC_HUMAN Filamin-C OS=Homo sapiens GN=FLNC PE=1 SV=3 | 4217,772 | 46,9 | 290959 | 103 | 29,36 | 29,58 | 28,61 | 29,58 | 29,71 | 28,53 | 1,064 | 8,59E-01 | 0 | 0,08999952 | 0,06601267 |
| ANXA1_HUN | ANXA1 | sp\|P04083\|ANXA1_HUMAN Annexin A1 OS=Homo sapiens GN=ANXA1 PE=1 SV=2 | 1011,666 | 50,87 | 38714 | 21 | 30,39 | 30,71 | 30,22 | 30,51 | 30,69 | 30,38 | 1,062 | 6,36E-01 | 0 | 0,08666738 | 0,19657383 |
| H4_HUMAN | HIST1H4A | sp\|P62805\|H4_HUMAN Histone H4 OS=Homo sapiens GN=HIST1H4A PE=1 SV=2 | 473,416 | 57,28 | 11367 | 13 | 28,54 | 28,51 | 29,14 | 28,79 | 28,81 | 28,85 | 1,062 | 6,96E-01 | 0 | 0,08666674 | 0,15770208 |
| BCDO1_HUN | BCO1 | sp\|Q9HAY6\|BCDO1_HUMAN Beta,beta-carotene 15,15'-dioxygenase OS=Homo sapiens | 57,559 | 1,28 | 62637 | 2 | 22,44 | 24,15 | 22,72 | 23,46 | 24,51 | 21,60 | 1,062 | 9,35E-01 | 0 | 0,08666674 | 0,02907672 |
| ESYT2_HUM | ESYT2 | sp\|A0FGR8\|ESYT2_HUMAN Extended synaptotagmin-2 OS=Homo sapiens GN=ESYT2 PE | 290,791 | 11,07 | 102357 | 8 | 23,74 | 23,83 | 24,64 | 24,14 | 24,34 | 23,99 | 1,062 | 7,89E-01 | 0 | 0,08666674 | 0,10266645 |
| SNX6_HUMA | SNX6 | sp\|Q9UNH7\|SNX6_HUMAN Sorting nexin-6 OS=Homo sapiens GN=SNX6 PE=1 SV=1 | 452,045 | 34,73 | 46649 | 13 | 26,05 | 25,92 | 25,45 | 25,79 | 26,00 | 25,89 | 1,062 | 6,75E-01 | 0 | 0,08666674 | 0,17056761 |
| MUC18_HUN | MCAM | sp\|P43121\|MUC18_HUMAN Cell surface glycoprotein MUC18 OS=Homo sapiens GN=M( | 63,818 | 2,94 | 71607 | 1 | 22,75 | 22,49 | 21,39 | 22,81 | 22,42 | 21,66 | 1,062 | 8,79E-01 | 0 | 0,08666674 | 0,0557739 |
| ARHG1_HUN | ARHGEF1 | sp\|Q92888\|ARHG1_HUMAN Rho guanine nucleotide exchange factor 1 OS=Homo sapie | 75,776 | 6,91 | 102435 | 5 | 23,04 | 23,31 | 22,52 | 22,67 | 23,33 | 23,13 | 1,062 | 7,89E-01 | 0 | 0,08666611 | 0,10282126 |
| JAG1_HUMA | JAG1 | sp\|P78504\|JAG1_HUMAN Protein jagged-1 OS=Homo sapiens GN=JAG1 PE=1 SV=3 | 21,031 | 1,48 | 133799 | 1 | 17,15 | 17,85 | 20,66 | 17,90 | 17,79 | 20,22 | 1,060 | 9,53E-01 | 0 | 0,08372943 | 0,02090919 |
| TEBP_HUMA | PTGES3 | sp\|Q15185\|TEBP_HUMAN Prostaglandin E synthase 3 OS=Homo sapiens GN=PTGES3 PE | 446,951 | 52,5 | 18697 | 8 | 26,16 | 26,68 | 26,90 | 26,17 | 26,71 | 27,11 | 1,059 | 8,23E-01 | 0 | 0,08333333 | 0,08440306 |
| XPO2_HUM/ | CSE1L | sp\|P55060\|XPO2_HUMAN Exportin-2 OS=Homo sapiens GN=CSE1L PE=1 SV=3 | 1313,569 | 48,3 | 110417 | 35 | 27,26 | 27,65 | 28,46 | 27,27 | 27,47 | 28,88 | 1,059 | 8,99E-01 | 0 | 0,08333333 | 0,04614217 |
| PDLI7_HUM/ | PDLIM7 | sp\|Q9NR12\|PDLI7_HUMAN PDZ and LIM domain protein 7 OS=Homo sapiens GN=PDLIM | 198,843 | 15,54 | 49845 | 6 | 24,92 | 24,57 | 24,45 | 24,41 | 24,75 | 25,03 | 1,059 | 7,33E-01 | 0 | 0,08333333 | 0,13471007 |
| PPIH_HUMAI | PPIH | sp\|O43447\|PPIH_HUMAN Peptidyl-prolyl cis-trans isomerase H OS=Homo sapiens GN=P | 134,83 | 28,81 | 19208 | 4 | 23,54 | 23,68 | 22,12 | 23,63 | 23,27 | 22,69 | 1,059 | 8,91E-01 | 0 | 0,0833327 | 0,05033295 |
| FXR2_HUMA | FXR2 | sp\|P51116\|FXR2_HUMAN Fragile X mental retardation syndrome-related protein 2 OS=I | 282,931 | 18,87 | 74223 | 9 | 22,38 | 22,26 | 24,79 | 22,57 | 22,39 | 24,72 | 1,059 | 9,44E-01 | 0 | 0,0833327 | 0,02506337 |
| A16A1_HUM | ALDH16A1 | sp\|Q8IZ83\|A16A1_HUMAN Aldehyde dehydrogenase family 16 member A1 OS=Homo sa | 60,27 | 1,5 | 85127 | 1 | 18,82 | 19,60 | 20,45 | 19,64 | 18,84 | 20,63 | 1,057 | 9,14E-01 | 0 | 0,08064524 | 0,0390672 |
| ARP3_HUMA | ACTR3 | sp\|P61158\|ARP3_HUMAN Actin-related protein 3 OS=Homo sapiens GN=ACTR3 PE=1 SV | 934,429 | 74,4 | 47371 | 23 | 27,82 | 27,57 | 27,29 | 27,67 | 27,75 | 27,50 | 1,057 | 6,62E-01 | 0 | 0,07999992 | 0,17896474 |
| EIF3A_HUM/ | EIF3A | sp\|Q14152\|EIF3A_HUMAN Eukaryotic translation initiation factor 3 subunit A OS=Homo | 886,844 | 23,81 | 166569 | 31 | 26,62 | 26,80 | 27,10 | 26,63 | 26,85 | 27,28 | 1,057 | 7,52E-01 | 0 | 0,07999992 | 0,12355951 |
| MK14_HUM/ | MAPK14 | sp\|Q16539\|MK14_HUMAN Mitogen-activated protein kinase 14 OS=Homo sapiens GN= | 198,848 | 30,83 | 41293 | 8 | 23,65 | 23,13 | 23,10 | 23,46 | 23,26 | 23,40 | 1,057 | 6,93E-01 | 0 | 0,07999992 | 0,15956375 |
| RALA_HUMA | RALA | sp\|P11233\|RALA_HUMAN Ras-related protein Ral-A OS=Homo sapiens GN=RALA PE=1 S | 682,945 | 71,36 | 23567 | 16 | 24,31 | 24,41 | 23,45 | 24,49 | 24,11 | 23,81 | 1,057 | 8,36E-01 | 0 | 0,07999992 | 0,07767311 |
| S4A10_HUM | SLC4A10 | sp\|Q6U841\|S4A10_HUMAN Sodium-driven chloride bicarbonate exchanger OS=Homo s | 50,972 | 0,98 | 125946 | 1 | 21,07 | 21,95 | 17,99 | 21,69 | 20,92 | 18,64 | 1,057 | 9,61E-01 | 0 | 0,07952499 | 0,01749965 |
| STX8_HUMA | STX8 | sp\|Q9UNK0\|STX8_HUMAN Syntaxin-8 OS=Homo sapiens GN=STX8 PE=1 SV=2 | 12,819 | 2,97 | 26907 | 1 | 20,59 | 20,14 | 18,83 | 20,57 | 20,53 | 18,70 | 1,056 | 9,28E-01 | 0 | 0,07793872 | 0,03238813 |
| RACK1_HUM | RACK1 | sp\|P63244\|RACK1_HUMAN Receptor of activated protein C kinase 1 OS=Homo sapiens ( | 990,837 | 81,7 | 35077 | 22 | 28,14 | 28,91 | 28,49 | 28,11 | 28,54 | 29,12 | 1,055 | 8,45E-01 | 0 | 0,07666779 | 0,07314004 |
| SWP70_HUN | SWAP70 | sp\|Q9UH65\|SWP70_HUMAN Switch-associated protein 70 OS=Homo sapiens GN=SWAF | 32,778 | 1,71 | 68998 | 1 | 20,99 | 20,67 | 20,70 | 21,11 | 20,99 | 20,49 | 1,055 | 7,40E-01 | 0 | 0,07666651 | 0,13075105 |
| RL23A_HUM | RPL23A | sp\|P62750\|RL23A_HUMAN 60S ribosomal protein L23a OS=Homo sapiens GN=RPL23A P | 211,368 | 34,62 | 17695 | 7 | 26,03 | 26,34 | 25,25 | 26,33 | 25,90 | 25,62 | 1,055 | 8,52E-01 | 0 | 0,07666651 | 0,06973862 |
| RS3A_HUMA | RPS3A | sp\|P61247\|RS3A_HUMAN 40S ribosomal protein S3a OS=Homo sapiens GN=RPS3A PE=1 | 779,635 | 61,36 | 29945 | 21 | 28,03 | 28,33 | 28,15 | 27,90 | 28,08 | 28,76 | 1,055 | 7,95E-01 | 0 | 0,07666651 | 0,09965911 |
| SNX12_HUM | SNX12 | sp\|Q9UMY4\|SNX12_HUMAN Sorting nexin-12 OS=Homo sapiens GN=SNX12 PE=1 SV=3 | 29,066 | 10,47 | 19730 | 2 | 20,97 | 19,43 | 17,61 | 21,40 | 20,54 | 16,30 | 1,054 | 9,69E-01 | 0 | 0,0762914 | 0,01363224 |
| ATRN_HUM/ | ATRN | sp\|O75882\|ATRN_HUMAN Attractin OS=Homo sapiens GN=ATRN PE=1 SV=2 | 58,084 | 1,82 | 158537 | 2 | 21,81 | 21,07 | 20,07 | 21,58 | 21,69 | 19,90 | 1,052 | 9,29E-01 | 0 | 0,07333374 | 0,03221315 |
| GRWD1_HUI | GRWD1 | sp\|Q9BQ67\|GRWD1_HUMAN Glutamate-rich WD repeat-containing protein 1 OS=Homo | 42,785 | 5,38 | 49419 | 2 | 21,76 | 21,41 | 22,10 | 22,21 | 21,02 | 22,26 | 1,052 | 8,79E-01 | 0 | 0,0733331 | 0,05607716 |
| EIF3B_HUM/ | EIF3B | sp\|P55884\|EIF3B_HUMAN Eukaryotic translation initiation factor 3 subunit B OS=Homo | 520,229 | 24,94 | 92482 | 17 | 25,87 | 26,02 | 26,24 | 25,77 | 26,26 | 26,32 | 1,052 | 7,38E-01 | 0 | 0,0733331 | 0,13181097 |
| SNX1_HUMA | SNX1 | sp\|Q13596\|SNX1_HUMAN Sorting nexin-1 OS=Homo sapiens GN=SNX1 PE=1 SV=3 | 265,445 | 19,92 | 59070 | 9 | 25,32 | 25,27 | 23,65 | 25,06 | 25,01 | 24,39 | 1,052 | 9,07E-01 | 0 | 0,0733331 | 0,04240516 |
| ABCF2_HUM | ABCF2 | sp\|Q9UG63\|ABCF2_HUMAN ATP-binding cassette sub-family F member 2 OS=Homo sap | 209,245 | 13,32 | 71290 | 7 | 24,12 | 24,02 | 24,07 | 24,13 | 24,01 | 24,29 | 1,052 | 4,42E-01 | 0 | 0,0733331 | 0,35428304 |
| CKLF6_HUM | CMTM6 | sp\|Q9NX76\|CKLF6_HUMAN CKLF-like MARVEL transmembrane domain-containing prot | 50,853 | 10,38 | 20419 | 1 | 21,14 | 21,22 | 19,64 | 21,01 | 21,46 | 19,75 | 1,052 | 9,24E-01 | 0 | 0,07321103 | 0,03412186 |
| MYPT2_HUN | PPP1R12B | sp\|O60237\|MYPT2_HUMAN Protein phosphatase 1 regulatory subunit 12B OS=Homo sa | 29,732 | 1,83 | 110404 | 2 | 26,52 | 26,47 | 18,38 | 26,18 | 26,03 | 19,38 | 1,051 | 9,85E-01 | 0 | 0,07220395 | 0,00674522 |
| TRA2B_HUM | TRA2B | sp\|P62995\|TRA2B_HUMAN Transformer-2 protein homolog beta OS=Homo sapiens GN= | 21,235 | 6,6 | 33666 | 2 | 20,89 | 20,10 | 22,73 | 20,49 | 20,58 | 22,86 | 1,050 | 9,52E-01 | 0 | 0,07000033 | 0,02123538 |
| TXND5_HUN | TXNDC5 | sp\|Q8NBS9\|TXND5_HUMAN Thioredoxin domain-containing protein 5 OS=Homo sapien | 113,572 | 13,19 | 47629 | 4 | 23,00 | 23,08 | 25,12 | 23,36 | 23,53 | 24,52 | 1,050 | 9,33E-01 | 0 | 0,07000033 | 0,03011603 |
| RU2A_HUMA | SNRPA1 | sp\|P09661\|RU2A_HUMAN U2 small nuclear ribonucleoprotein A' OS=Homo sapiens GN= | 322,78 | 43,14 | 28416 | 10 | 25,75 | 25,23 | 26,50 | 26,00 | 25,84 | 25,85 | 1,050 | 8,60E-01 | 0 | 0,07000033 | 0,06550794 |
| XRP2_HUMA | RP2 | sp\|O75695\|XRP2_HUMAN Protein XRP2 OS=Homo sapiens GN=RP2 PE=1 SV=4 | 149,316 | 20 | 39641 | 7 | 25,51 | 25,26 | 24,00 | 25,61 | 25,31 | 24,06 | 1,050 | 9,21E-01 | 0 | 0,06999969 | 0,03557146 |
| VP33A_HUM | VPS33A | sp\|Q96AX1\|VP33A_HUMAN Vacuolar protein sorting-associated protein 33A OS=Homo | 71,134 | 7,72 | 67611 | 2 | 19,87 | 20,32 | 18,42 | 19,66 | 19,86 | 19,30 | 1,050 | 9,12E-01 | 0 | 0,06999969 | 0,0399124 |
| ARHL2_HUM | ADPRHL2 | sp\|Q9NX46\|ARHL2_HUMAN Poly(ADP-ribose) glycohydrolase ARH3 OS=Homo sapiens G | 72,713 | 6,34 | 38947 | 2 | 21,95 | 22,33 | 21,14 | 22,15 | 22,17 | 21,31 | 1,050 | 8,84E-01 | 0 | 0,06999969 | 0,05345513 |
| HS74L_HUM | HSPA4L | sp\|O95757\|HS74L_HUMAN Heat shock 70 kDa protein 4L OS=Homo sapiens GN=HSPA4L | 308,143 | 11,8 | 94512 | 7 | 23,68 | 23,23 | 22,62 | 23,61 | 23,58 | 22,54 | 1,047 | 8,93E-01 | 0 | 0,06666692 | 0,04896121 |
| RL10_HUMA | RPL10 | sp\|P27635\|RL10_HUMAN 60S ribosomal protein L10 OS=Homo sapiens GN=RPL10 PE=1 | 514,154 | 46,26 | 24604 | 16 | 26,74 | 27,00 | 27,90 | 26,87 | 26,66 | 28,31 | 1,047 | 9,20E-01 | 0 | 0,06666692 | 0,03603768 |
| PAX6_HUMA | PAX6 | sp\|P26367\|PAX6_HUMAN Paired box protein Pax-6 OS=Homo sapiens GN=PAX6 PE=1 SV | 19,353 | 1,66 | 46683 | 1 | 24,78 | 23,89 | 23,46 | 24,79 | 24,57 | 22,97 | 1,047 | 9,28E-01 | 0 | 0,06666692 | 0,03246573 |
| RHEB_HUMA | RHEB | sp\|Q15382\|RHEB_HUMAN GTP-binding protein Rheb OS=Homo sapiens GN=RHEB PE=1 | 149,203 | 28,8 | 20497 | 5 | 24,87 | 24,57 | 24,21 | 24,81 | 24,97 | 24,07 | 1,047 | 8,53E-01 | 0 | 0,06666629 | 0,06924297 |
| BOLA2_HUM | BOLA2 | sp\|Q9H3K6\|BOLA2_HUMAN BolA-like protein 2 OS=Homo sapiens GN=BOLA2 PE=1 SV= | 69,634 | 29,07 | 10117 | 2 | 22,69 | 23,10 | 21,34 | 22,67 | 22,93 | 21,73 | 1,047 | 9,23E-01 | 0 | 0,06666629 | 0,03498346 |
| GHC1_HUM/ | SLC25A22 | sp\|Q9H936\|GHC1_HUMAN Mitochondrial glutamate carrier 1 OS=Homo sapiens GN=SL( | 11,412 | 6,19 | 34470 | 1 | 18,24 | 18,80 | 19,65 | 19,48 | 18,64 | 18,76 | 1,046 | 9,00E-01 | 0 | 0,06502342 | 0,04558488 |
| SC24B_HUM | SEC24B | sp\|O95487\|SC24B_HUMAN Protein transport protein Sec24B OS=Homo sapiens GN=SE( | 73,713 | 1,74 | 137418 | 2 | 21,07 | 21,05 | 21,01 | 20,84 | 20,99 | 21,49 | 1,045 | 7,64E-01 | 0 | 0,06333351 | 0,11675516 |
| GLGB_HUM/ | GBE1 | sp\|Q04446\|GLGB_HUMAN 1,4-alpha-glucan-branching enzyme OS=Homo sapiens GN=( | 15,603 | 1,99 | 80474 | 1 | 21,16 | 21,00 | 20,04 | 21,35 | 20,45 | 20,59 | 1,045 | 8,94E-01 | 0 | 0,06333351 | 0,04848965 |
| CSDE1_HUM | CSDE1 | sp\|O75534\|CSDE1_HUMAN Cold shock domain-containing protein E1 OS=Homo sapiens | 570,574 | 28,32 | 88885 | 20 | 25,52 | 25,23 | 25,83 | 25,65 | 25,35 | 25,77 | 1,045 | 7,82E-01 | 0 | 0,06333351 | 0,10703191 |
| GBF1_HUMA | GBF1 | sp\|Q92538\|GBF1_HUMAN Golgi-specific brefeldin A-resistance guanine nucleotide exch | 49,274 | 1,24 | 206446 | 2 | 19,10 | 20,56 | 19,60 | 19,78 | 20,19 | 19,48 | 1,045 | 9,00E-01 | 0 | 0,06333351 | 0,04555543 |
| NRAP_HUM/ | NRAP | sp\|Q86VF7\|NRAP_HUMAN Nebulin-related-anchoring protein OS=Homo sapiens GN=NI | 14,957 | 0,4 | 197074 | 1 | 24,27 | 22,41 | 20,88 | 24,44 | 23,89 | 19,42 | 1,045 | 9,75E-01 | 0 | 0,06333351 | 0,0111865 |
| U119B_HUM | UNC119B | sp\|A6NIH7\|U119B_HUMAN Protein unc-119 homolog B OS=Homo sapiens GN=UNC119I | 26,594 | 11,55 | 28137 | 2 | 21,94 | 21,92 | 21,02 | 22,04 | 21,88 | 21,15 | 1,045 | 8,84E-01 | 0 | 0,06333288 | 0,05336555 |
| EF1A2_HUM | EEF1A2 | sp\|Q05639\|EF1A2_HUMAN Elongation factor 1-alpha 2 OS=Homo sapiens GN=EEF1A2 F | 1149,596 | 56,16 | 50470 | 28 | 23,68 | 23,88 | 18,88 | 23,41 | 23,91 | 19,31 | 1,044 | 9,79E-01 | 0 | 0,0618604 | 0,0092959 |
| VATD_HUM/ | ATP6V1D | sp\|Q9Y5K8\|VATD_HUMAN V-type proton ATPase subunit D OS=Homo sapiens GN=ATP( | 24,453 | 4,45 | 28263 | 1 | 20,89 | 21,08 | 20,30 | 21,10 | 21,02 | 20,33 | 1,042 | 8,68E-01 | 0 | 0,06000074 | 0,06143215 |
| NINJ1_HUM/ | NINJ1 | sp\|Q92982\|NINJ1_HUMAN Ninjurin-1 OS=Homo sapiens GN=NINJ1 PE=1 SV=2 | 93,144 | 13,16 | 16345 | 1 | 21,99 | 21,90 | 22,29 | 21,85 | 22,33 | 22,18 | 1,042 | 7,61E-01 | 0 | 0,0600001 | 0,11850639 |
| RAP2B_HUM | RAP2B | sp\|P61225\|RAP2B_HUMAN Ras-related protein Rap-2b OS=Homo sapiens GN=RAP2B PE | 525,548 | 58,47 | 20504 | 10 | 25,16 | 24,91 | 23,22 | 25,16 | 25,06 | 23,25 | 1,042 | 9,48E-01 | 0 | 0,0600001 | 0,02304866 |
| LANC2_HUM | LANCL2 | sp\|Q9NS86\|LANC2_HUMAN LanC-like protein 2 OS=Homo sapiens GN=LANCL2 PE=1 SV: | 71,046 | 5,11 | 50854 | 2 | 21,67 | 21,50 | 21,95 | 21,50 | 21,43 | 22,37 | 1,042 | 8,64E-01 | 0 | 0,0600001 | 0,06328694 |
| IF2P_HUMAI | EIF5B | sp\|O60841\|IF2P_HUMAN Eukaryotic translation initiation factor 5B OS=Homo sapiens G | 341,554 | 12,7 | 138827 | 12 | 24,72 | 24,95 | 24,36 | 24,69 | 24,66 | 24,86 | 1,042 | 7,59E-01 | 0 | 0,0600001 | 0,11976152 |
| FHL3_HUMA | FHL3 | sp\|Q13643\|FHL3_HUMAN Four and a half LIM domains protein 3 OS=Homo sapiens GN= | 73,901 | 15,71 | 31192 | 3 | 22,45 | 22,51 | 23,24 | 22,78 | 22,54 | 23,06 | 1,042 | 8,49E-01 | 0 | 0,0600001 | 0,07120293 |
| RET4_HUMA | RBP4 | sp\|P02753\|RET4_HUMAN Retinol-binding protein 4 OS=Homo sapiens GN=RBP4 PE=1 S' | 39,108 | 9,95 | 23010 | 2 | 22,25 | 20,33 | 22,73 | 23,14 | 20,12 | 22,23 | 1,042 | 9,61E-01 | 0 | 0,0600001 | 0,01722543 |
| EF1A1_HUM | EEF1A1 | sp\|P68104\|EF1A1_HUMAN Elongation factor 1-alpha 1 OS=Homo sapiens GN=EEF1A1 P | 2560,38 | 77,06 | 50141 | 56 | 30,93 | 30,53 | 32,02 | 30,82 | 31,11 | 31,73 | 1,042 | 9,14E-01 | 0 | 0,05999947 | 0,03920573 |
| UB2V2_HUM | UBE2V2 | sp\|Q15819\|UB2V2_HUMAN Ubiquitin-conjugating enzyme E2 variant 2 OS=Homo sapie | 332,483 | 55,17 | 16363 | 8 | 24,52 | 25,00 | 22,13 | 24,58 | 24,58 | 22,66 | 1,040 | 9,61E-01 | 0 | 0,05666669 | 0,01719452 |
| SERC_HUMA | PSAT1 | sp\|Q9Y617\|SERC_HUMAN Phosphoserine aminotransferase OS=Homo sapiens GN=PSA1 | 193,825 | 18,38 | 40423 | 7 | 26,19 | 26,28 | 24,18 | 26,04 | 26,46 | 24,32 | 1,040 | 9,55E-01 | 0 | 0,05666606 | 0,01990706 |
| ARI1A_HUM | ARID1A | sp\|O14497\|ARI1A_HUMAN AT-rich interactive domain-containing protein 1A OS=Homo | 14,227 | 0,31 | 242045 | 1 | 18,35 | 18,52 | 22,12 | 19,28 | 18,43 | 21,46 | 1,039 | 9,73E-01 | 0 | 0,05570539 | 0,01205927 |
| RL36A_HUM | RPL36A | sp\|P83881\|RL36A_HUMAN 60S ribosomal protein L36a OS=Homo sapiens GN=RPL36A P | 57,99 | 24,53 | 12441 | 3 | 25,16 | 24,76 | 24,16 | 25,20 | 24,70 | 24,34 | 1,038 | 8,96E-01 | 0 | 0,05333392 | 0,04771306 |
| ALDOC_HUN | ALDOC | sp\|P09972\|ALDOC_HUMAN Fructose-bisphosphate aldolase C OS=Homo sapiens GN=Al | 305,212 | 16,76 | 39456 | 5 | 22,64 | 22,93 | 24,14 | 22,52 | 23,21 | 24,14 | 1,038 | 9,39E-01 | 0 | 0,05333328 | 0,02725259 |
| IMDH2_HUN | IMPDH2 | sp\|P12268\|IMDH2_HUMAN Inosine-5'-monophosphate dehydrogenase 2 OS=Homo sap | 656,52 | 48,25 | 55805 | 18 | 25,84 | 26,04 | 27,19 | 26,09 | 26,28 | 26,86 | 1,038 | 9,17E-01 | 0 | 0,05333328 | 0,03767221 |
| VP26B_HUM | VPS26B | sp\|Q4G0F5\|VP26B_HUMAN Vacuolar protein sorting-associated protein 26B OS=Homo | 85,555 | 12,2 | 39155 | 3 | 21,95 | 22,38 | 22,90 | 21,96 | 22,12 | 23,31 | 1,038 | 9,21E-01 | 0 | 0,05333328 | 0,03562506 |
| MP2K1_HUN | MAP2K1 | sp\|Q02750\|MP2K1_HUMAN Dual specificity mitogen-activated protein kinase kinase 1 ( | 678,023 | 61,32 | 43439 | 18 | 26,33 | 26,62 | 26,08 | 26,43 | 26,48 | 26,28 | 1,038 | 7,66E-01 | 0 | 0,05333328 | 0,11593901 |
| IPO13_HUM | IPO13 | sp\|O94829\|IPO13_HUMAN Importin-13 OS=Homo sapiens GN=IPO13 PE=1 SV=3 | 78,428 | 3,12 | 108195 | 2 | 18,11 | 18,53 | 21,03 | 18,37 | 18,79 | 20,67 | 1,038 | 9,65E-01 | 0 | 0,05333328 | 0,01531929 |
| ARK74_HUM | AKR7L | sp\|Q8NHP1\|ARK74_HUMAN Aflatoxin B1 aldehyde reductase member 4 OS=Homo sapi | 46,437 | 8,46 | 36964 | 2 | 23,10 | 22,85 | 23,19 | 22,88 | 23,06 | 23,36 | 1,038 | 7,73E-01 | 0 | 0,05333265 | 0,11164509 |
| HS12A_HUM | HSPA12A | sp\|O43301\|HS12A_HUMAN Heat shock 70 kDa protein 12A OS=Homo sapiens GN=HSPA | 282,038 | 16,74 | 74978 | 9 | 22,52 | 22,60 | 24,16 | 22,49 | 22,71 | 24,24 | 1,038 | 9,48E-01 | 0 | 0,05333265 | 0,02324539 |
| PEDF_HUMA | SERPINF1 | sp\|P36955\|PEDF_HUMAN Pigment epithelium-derived factor OS=Homo sapiens GN=SEF | 207,965 | 9,33 | 46312 | 3 | 24,58 | 22,84 | 26,54 | 24,65 | 23,73 | 25,74 | 1,038 | 9,67E-01 | 0 | 0,05333265 | 0,01451516 |
| NAGK_HUM/ | NAGK | sp\|Q9UJ70\|NAGK_HUMAN N-acetyl-D-glucosamine kinase OS=Homo sapiens GN=NAGK | 151,62 | 19,19 | 37376 | 6 | 23,52 | 23,45 | 22,25 | 23,59 | 23,65 | 22,14 | 1,038 | 9,38E-01 | 0 | 0,05333265 | 0,02784837 |
| OSBL9_HUM | OSBPL9 | sp\|Q96SU4\|OSBL9_HUMAN Oxysterol-binding protein-related protein 9 OS=Homo sapi | 16,953 | 2,17 | 83185 | 1 | 18,11 | 18,73 | 19,04 | 18,25 | 18,62 | 19,16 | 1,036 | 9,00E-01 | 0 | 0,05100822 | 0,04594692 |
| WWP1_HUN | WWP1 | sp\|Q9H0M0\|WWP1_HUMAN NEDD4-like E3 ubiquitin-protein ligase WWP1 OS=Homo s | 264,561 | 10,3 | 105202 | 9 | 23,11 | 21,96 | 20,59 | 22,18 | 21,99 | 21,64 | 1,035 | 9,50E-01 | 0 | 0,05001005 | 0,02240162 |
| SHC1_HUMA | SHC1 | sp\|P29353\|SHC1_HUMAN SHC-transforming protein 1 OS=Homo sapiens GN=SHC1 PE=: | 170,541 | 11,66 | 62822 | 5 | 22,97 | 23,38 | 21,77 | 22,95 | 22,63 | 22,69 | 1,035 | 9,24E-01 | 0 | 0,05000051 | 0,03428691 |
| SERC1_HUM | SERINC1 | sp\|Q9NRX5\|SERC1_HUMAN Serine incorporator 1 OS=Homo sapiens GN=SERINC1 PE=1 | 153,869 | 6,62 | 50495 | 2 | 21,41 | 21,16 | 21,16 | 21,35 | 21,60 | 20,93 | 1,035 | 8,26E-01 | 0 | 0,05000051 | 0,08326574 |
| PA1B3_HUM | PAFAH1B3 | sp\|Q15102\|PA1B3_HUMAN Platelet-activating factor acetylhydrolase IB subunit gamma | 197,227 | 44,59 | 25734 | 6 | 24,74 | 24,96 | 24,28 | 24,74 | 24,62 | 24,77 | 1,035 | 8,20E-01 | 0 | 0,05000051 | 0,08632461 |
| KIF2C_HUM | KIF2C | sp\|Q99661\|KIF2C_HUMAN Kinesin-like protein KIF2C OS=Homo sapiens GN=KIF2C PE=1 | 19,984 | 1,52 | 81313 | 1 | 17,48 | 19,59 | 21,84 | 18,42 | 18,45 | 22,18 | 1,034 | 9,80E-01 | 0 | 0,04820824 | 0,0089495 |
| LRP8_HUMA | LRP8 | sp\|Q14114\|LRP8_HUMAN Low-density lipoprotein receptor-related protein 8 OS=Homc | 10,231 | 0,93 | 105634 | 1 | 21,50 | 21,30 | 19,99 | 21,08 | 21,53 | 20,33 | 1,034 | 9,39E-01 | 0 | 0,04805756 | 0,02743202 |
| AACS_HUMA | AACS | sp\|Q86V21\|AACS_HUMAN Acetoacetyl-CoA synthetase OS=Homo sapiens GN=AACS PE: | 10,836 | 1,19 | 75144 | 1 | 21,44 | 21,22 | 21,14 | 21,34 | 21,32 | 21,28 | 1,033 | 6,37E-01 | 0 | 0,0466671 | 0,19615792 |
| VKORL_HUM | VKORC1L1 | sp\|Q8N0U8\|VKORL_HUMAN Vitamin K epoxide reductase complex subunit 1-like protei | 45,504 | 6,25 | 19836 | 1 | 20,97 | 20,92 | 22,00 | 20,62 | 21,50 | 21,91 | 1,033 | 9,33E-01 | 0 | 0,0466671 | 0,03030857 |
| DUS3_HUM/ | DUSP3 | sp\|P51452\|DUS3_HUMAN Dual specificity protein phosphatase 3 OS=Homo sapiens GN: | 68,913 | 13,51 | 20478 | 2 | 21,44 | 22,43 | 20,55 | 21,69 | 22,04 | 20,83 | 1,033 | 9,46E-01 | 0 | 0,0466671 | 0,02396705 |
| PGAM1_HUN | PGAM1 | sp\|P18669\|PGAM1_HUMAN Phosphoglycerate mutase 1 OS=Homo sapiens GN=PGAM1 | 1199,398 | 80,71 | 28804 | 22 | 28,47 | 28,84 | 27,49 | 28,52 | 28,55 | 27,87 | 1,033 | 9,24E-01 | 0 | 0,0466671 | 0,0343077 |
| IF4E_HUMAI | EIF4E | sp\|P06730\|IF4E_HUMAN Eukaryotic translation initiation factor 4E OS=Homo sapiens GI | 134,418 | 14,75 | 25097 | 3 | 24,42 | 24,79 | 23,95 | 24,27 | 24,59 | 24,44 | 1,033 | 8,66E-01 | 0 | 0,04666646 | 0,06233029 |
| SNP23_HUM | SNAP23 | sp\|O00161\|SNP23_HUMAN Synaptosomal-associated protein 23 OS=Homo sapiens GN= | 149,854 | 21,8 | 23354 | 4 | 25,04 | 25,28 | 24,62 | 25,54 | 25,43 | 24,11 | 1,033 | 9,30E-01 | 0 | 0,04666646 | 0,03157029 |
| UB2V1_HUM | UBE2V1 | sp\|Q13404\|UB2V1_HUMAN Ubiquitin-conjugating enzyme E2 variant 1 OS=Homo sapie | 335,082 | 57,82 | 16495 | 8 | 24,57 | 24,67 | 22,53 | 24,57 | 24,70 | 22,64 | 1,033 | 9,64E-01 | 0 | 0,04666646 | 0,01604874 |
| HUWE1_HUN | HUWE1 | sp\|Q7Z6Z7\|HUWE1_HUMAN E3 ubiquitin-protein ligase HUWE1 OS=Homo sapiens GN= | 711,438 | 8,94 | 481891 | 25 | 24,49 | 24,95 | 25,07 | 24,41 | 24,93 | 25,31 | 1,033 | 8,89E-01 | 0 | 0,04666646 | 0,05088834 |
| PFKAL_HUM | PFKL | sp\|P17858\|PFKAL_HUMAN ATP-dependent 6-phosphofructokinase, liver type OS=Homc | 419,353 | 12,44 | 85018 | 8 | 24,69 | 24,90 | 25,22 | 24,69 | 25,05 | 25,21 | 1,033 | 8,41E-01 | 0 | 0,04666646 | 0,07533722 |
| AT1A1_HUM | ATP1A1 | sp\|P05023\|AT1A1_HUMAN Sodium/potassium-transporting ATPase subunit alpha-1 OS: | 2862,22 | 44,87 | 112896 | 59 | 30,12 | 30,38 | 28,98 | 30,22 | 30,30 | 29,10 | 1,033 | 9,40E-01 | 0 | 0,04666646 | 0,02705643 |
| PCY2_HUMA | PCYT2 | sp\|Q99447\|PCY2_HUMAN Ethanolamine-phosphate cytidylyltransferase OS=Homo sapi | 39,525 | 5,14 | 43835 | 2 | 22,67 | 23,18 | 20,44 | 22,66 | 22,39 | 21,38 | 1,033 | 9,62E-01 | 0 | 0,04666583 | 0,01670334 |
| PI42B_HUM | PIP4K2B | sp\|P78356\|PI42B_HUMAN Phosphatidylinositol 5-phosphate 4-kinase type-2 beta OS=H | 146,494 | 9,62 | 47378 | 3 | 20,57 | 20,93 | 20,18 | 20,57 | 20,50 | 20,75 | 1,032 | 8,51E-01 | 0 | 0,04581388 | 0,07020091 |
| GPTC8_HUM | GPATCH8 | sp\|Q9UKJ3\|GPTC8_HUMAN G patch domain-containing protein 8 OS=Homo sapiens GN | 9,236 | 0,67 | 164197 | 1 | 18,89 | 19,33 | 21,41 | 19,64 | 18,65 | 21,47 | 1,031 | 9,71E-01 | 0 | 0,04383787 | 0,01277568 |
| RAB7A_HUM | RAB7A | sp\|P51149\|RAB7A_HUMAN Ras-related protein Rab-7a OS=Homo sapiens GN=RAB7A PE | 779,851 | 72,46 | 23490 | 18 | 28,56 | 28,44 | 28,30 | 28,52 | 28,62 | 28,29 | 1,030 | 7,43E-01 | 0 | 0,04333433 | 0,12908902 |

| | | | | | | | | | | | | | | | | |
|---|---|---|---|---|---|---|---|---|---|---|---|---|---|---|---|---|
| LRC8A_HUM | LRRC8A | sp\|Q8IWT6\|LRC8A_HUMAN Volume-regulated anion channel subunit LRRC8A OS=Homo | 262,582 | 16,54 | 94199 | 11 | 24,73 | 24,98 | 23,80 | 24,80 | 25,08 | 23,76 | **1,030** | 9,40E-01 | **0** | 0,04333369 | 0,02699115 |
| SEP10_HUM | SEPT10 | sp\|Q9P0V9\|SEP10_HUMAN Septin-10 OS=Homo sapiens GN=SEPT10 PE=1 SV=2 | 39,298 | 6,17 | 52593 | 2 | 23,24 | 22,80 | 22,09 | 22,83 | 23,25 | 22,18 | **1,030** | 9,29E-01 | **0** | 0,04333369 | 0,03195593 |
| CLIC1_HUM | CLIC1 | sp\|O00299\|CLIC1_HUMAN Chloride intracellular channel protein 1 OS=Homo sapiens GI | 551,466 | 67,63 | 26923 | 13 | 27,87 | 27,67 | 28,01 | 27,79 | 27,74 | 28,15 | **1,030** | 8,03E-01 | **0** | 0,04333305 | 0,09532372 |
| FYN_HUMAN | FYN | sp\|P06241\|FYN_HUMAN Tyrosine-protein kinase Fyn OS=Homo sapiens GN=FYN PE=1 S | 160,901 | 14,34 | 60762 | 7 | 17,28 | 17,02 | 18,05 | 17,41 | 17,11 | 17,96 | **1,030** | 9,20E-01 | **0** | 0,04239019 | 0,03609794 |
| IMB1_HUMA | KPNB1 | sp\|Q14974\|IMB1_HUMAN Importin subunit beta-1 OS=Homo sapiens GN=KPNB1 PE=1 ! | 2126,989 | 62,1 | 97170 | 46 | 28,96 | 29,05 | 28,88 | 28,99 | 29,04 | 28,98 | **1,028** | 4,89E-01 | **0** | 0,04000092 | 0,31113025 |
| CSN7B_HUM | COPS7B | sp\|Q9H9Q2\|CSN7B_HUMAN COP9 signalosome complex subunit 7b OS=Homo sapiens ( | 59,797 | 6,82 | 29622 | 1 | 18,65 | 19,24 | 20,23 | 18,71 | 18,79 | 20,74 | **1,028** | 9,63E-01 | **0** | 0,04000028 | 0,01642131 |
| FNBP1_HUM | FNBP1 | sp\|Q96RU3\|FNBP1_HUMAN Formin-binding protein 1 OS=Homo sapiens GN=FNBP1 PE= | 93,497 | 5,35 | 71307 | 3 | 22,13 | 22,55 | 21,20 | 22,19 | 22,56 | 21,25 | **1,028** | 9,46E-01 | **0** | 0,04000028 | 0,0239832 |
| RAB5A_HUM | RAB5A | sp\|P20339\|RAB5A_HUMAN Ras-related protein Rab-5A OS=Homo sapiens GN=RAB5A PI | 368,326 | 53,49 | 23659 | 9 | 24,91 | 25,02 | 24,74 | 24,89 | 25,12 | 24,78 | **1,028** | 7,72E-01 | **0** | 0,04000028 | 0,1122965 |
| GDS1_HUMA | RAP1GDS1 | sp\|P52306\|GDS1_HUMAN Rap1 GTPase-GDP dissociation stimulator 1 OS=Homo sapiens | 211,806 | 12,36 | 66317 | 5 | 22,64 | 23,11 | 22,78 | 22,39 | 23,06 | 23,20 | **1,028** | 8,96E-01 | **0** | 0,03999964 | 0,04789075 |
| EIF3C_HUM | EIF3C | sp\|Q99613\|EIF3C_HUMAN Eukaryotic translation initiation factor 3 subunit C OS=Homo | 758,489 | 33,08 | 105344 | 25 | 26,28 | 26,62 | 26,61 | 26,44 | 26,45 | 26,74 | **1,028** | 8,01E-01 | **0** | 0,03999964 | 0,09613508 |
| DAAM1_HUM | DAAM1 | sp\|Q9Y4D1\|DAAM1_HUMAN Disheveled-associated activator of morphogenesis 1 OS=H | 54,035 | 3,34 | 123473 | 2 | 19,63 | 19,52 | 17,19 | 19,32 | 19,14 | 17,99 | **1,027** | 9,68E-01 | **0** | 0,03784307 | 0,01391887 |
| NEDD8_HUM | NEDD8 | sp\|Q15843\|NEDD8_HUMAN NEDD8 OS=Homo sapiens GN=NEDD8 PE=1 SV=1 | 124,049 | 40,74 | 9072 | 3 | 24,26 | 24,73 | 21,89 | 24,45 | 24,42 | 22,12 | **1,026** | 9,76E-01 | **0** | 0,03666751 | 0,01033136 |
| FSCN1_HUM | FSCN1 | sp\|Q16658\|FSCN1_HUMAN Fascin OS=Homo sapiens GN=FSCN1 PE=1 SV=3 | 724,705 | 41,38 | 54530 | 19 | 28,29 | 28,34 | 27,36 | 28,20 | 28,36 | 27,54 | **1,026** | 9,32E-01 | **0** | 0,03666687 | 0,03043675 |
| NHRF1_HUM | SLC9A3R1 | sp\|O14745\|NHRF1_HUMAN Na(+)/H(+) exchange regulatory cofactor NHE-RF1 OS=Hom | 204,781 | 17,6 | 38868 | 6 | 27,14 | 25,95 | 23,88 | 27,22 | 26,19 | 23,67 | **1,026** | 9,81E-01 | **0** | 0,03666687 | 0,00848643 |
| YLAT2_HUM | SLC7A6 | sp\|Q92536\|YLAT2_HUMAN Y+L amino acid transporter 2 OS=Homo sapiens GN=SLC7A6 | 57,584 | 4,27 | 56828 | 2 | 21,52 | 21,81 | 21,42 | 21,25 | 21,72 | 21,89 | **1,026** | 8,78E-01 | **0** | 0,03666623 | 0,05646781 |
| CNN3_HUM | CNN3 | sp\|Q15417\|CNN3_HUMAN Calponin-3 OS=Homo sapiens GN=CNN3 PE=1 SV=1 | 301,154 | 41,95 | 36414 | 10 | 24,92 | 25,02 | 24,89 | 24,92 | 25,14 | 24,88 | **1,026** | 7,04E-01 | **0** | 0,03666623 | 0,15230783 |
| DYHC1_HUM | DYNC1H1 | sp\|Q14204\|DYHC1_HUMAN Cytoplasmic dynein 1 heavy chain 1 OS=Homo sapiens GN= | 5957,578 | 45,14 | 532408 | 179 | 30,04 | 30,20 | 29,35 | 29,97 | 30,19 | 29,54 | **1,026** | 9,15E-01 | **0** | 0,03666623 | 0,03851428 |
| CUL3_HUMA | CUL3 | sp\|Q13618\|CUL3_HUMAN Cullin-3 OS=Homo sapiens GN=CUL3 PE=1 SV=2 | 170,476 | 15,23 | 88930 | 10 | 24,10 | 24,03 | 23,70 | 23,93 | 24,23 | 23,78 | **1,026** | 8,49E-01 | **0** | 0,03666623 | 0,07096954 |
| SRRT_HUMA | SRRT | sp\|Q9BXP5\|SRRT_HUMAN Serrate RNA effector molecule homolog OS=Homo sapiens G | 187,645 | 9,25 | 100666 | 8 | 23,06 | 23,23 | 24,77 | 23,01 | 23,16 | 24,99 | **1,023** | 9,70E-01 | **0** | 0,03333346 | 0,0131618 |
| S12A2_HUM | SLC12A2 | sp\|P55011\|S12A2_HUMAN Solute carrier family 12 member 2 OS=Homo sapiens GN=SL | 233,761 | 7,67 | 131447 | 8 | 25,05 | 25,12 | 22,70 | 24,88 | 25,03 | 23,06 | **1,023** | 9,75E-01 | **0** | 0,03333282 | 0,01081184 |
| GANAB_HUM | GANAB | sp\|Q14697\|GANAB_HUMAN Neutral alpha-glucosidase AB OS=Homo sapiens GN=GANA | 356,042 | 16,63 | 106874 | 13 | 25,36 | 25,29 | 26,33 | 25,47 | 25,65 | 25,96 | **1,023** | 9,32E-01 | **0** | 0,03333219 | 0,03077111 |
| RS16_HUMA | RPS16 | sp\|P62249\|RS16_HUMAN 40S ribosomal protein S16 OS=Homo sapiens GN=RPS16 PE=1 | 400,095 | 58,9 | 16445 | 12 | 28,25 | 27,89 | 27,95 | 28,28 | 27,78 | 28,12 | **1,021** | 8,79E-01 | **0** | 0,03000069 | 0,05607198 |
| IF4H_HUMA | EIF4H | sp\|Q15056\|IF4H_HUMAN Eukaryotic translation initiation factor 4H OS=Homo sapiens G | 89,452 | 16,94 | 27385 | 2 | 21,98 | 21,86 | 22,47 | 21,33 | 22,78 | 22,29 | **1,021** | 9,52E-01 | **0** | 0,03000069 | 0,02152459 |
| EIF2D_HUM | EIF2D | sp\|P41214\|EIF2D_HUMAN Eukaryotic translation initiation factor 2D OS=Homo sapiens ( | 23,637 | 2,4 | 64706 | 1 | 19,63 | 20,32 | 20,59 | 20,00 | 20,26 | 20,37 | **1,021** | 9,27E-01 | **0** | 0,03000069 | 0,03308248 |
| UBXN1_HUM | UBXN1 | sp\|Q04323\|UBXN1_HUMAN UBX domain-containing protein 1 OS=Homo sapiens GN=UI | 121,414 | 7,74 | 33325 | 3 | 20,07 | 20,48 | 20,85 | 20,03 | 20,24 | 21,22 | **1,021** | 9,48E-01 | **0** | 0,03000005 | 0,02329581 |
| PRDX2_HUM | PRDX2 | sp\|P32119\|PRDX2_HUMAN Peroxiredoxin-2 OS=Homo sapiens GN=PRDX2 PE=1 SV=5 | 158,534 | 18,69 | 21892 | 4 | 23,85 | 24,07 | 23,50 | 23,76 | 24,29 | 23,46 | **1,021** | 9,24E-01 | **0** | 0,03000005 | 0,03450251 |
| TM119_HUM | TMEM119 | sp\|Q4V9L6\|TM119_HUMAN Transmembrane protein 119 OS=Homo sapiens GN=TMEM | 49,456 | 3,89 | 29203 | 1 | 22,76 | 23,08 | 21,95 | 22,86 | 23,13 | 21,89 | **1,021** | 9,55E-01 | **0** | 0,02999942 | 0,01978683 |
| PFD1_HUMA | PFDN1 | sp\|O60925\|PFD1_HUMAN Prefoldin subunit 1 OS=Homo sapiens GN=PFDN1 PE=1 SV=2 | 41,141 | 9,02 | 14210 | 1 | 20,69 | 20,79 | 21,11 | 20,83 | 20,72 | 21,13 | **1,021** | 8,73E-01 | **0** | 0,02999878 | 0,05893948 |
| TT30A_HUM | TTC30A | sp\|Q86WT1\|TT30A_HUMAN Tetratricopeptide repeat protein 30A OS=Homo sapiens GI | 177,136 | 13,38 | 76136 | 6 | 18,75 | 18,27 | 22,60 | 19,07 | 18,44 | 22,18 | **1,020** | 9,88E-01 | **0** | 0,02838516 | 0,00518854 |
| SMAP_HUM | SMAP | sp\|O00193\|SMAP_HUMAN Small acidic protein OS=Homo sapiens GN=SMAP PE=1 SV=1 | 69,144 | 12,57 | 20333 | 1 | 19,95 | 20,06 | 18,98 | 20,18 | 19,81 | 19,08 | **1,019** | 9,56E-01 | **0** | 0,02774747 | 0,01961671 |
| RBM3_HUM | RBM3 | sp\|P98179\|RBM3_HUMAN RNA-binding protein 3 OS=Homo sapiens GN=RBM3 PE=1 SV | 111,057 | 20,38 | 17170 | 3 | 24,74 | 24,19 | 24,40 | 25,02 | 24,49 | 23,90 | **1,019** | 9,45E-01 | **0** | 0,02666664 | 0,02472484 |
| ESYT1_HUM | ESYT1 | sp\|Q9BSJ8\|ESYT1_HUMAN Extended synaptotagmin-1 OS=Homo sapiens GN=ESYT1 PE= | 310,582 | 10,33 | 122856 | 9 | 25,56 | 25,90 | 25,85 | 25,82 | 26,02 | 25,55 | **1,019** | 8,85E-01 | **0** | 0,02666664 | 0,0532171 |
| PP2BA_HUM | PPP3CA | sp\|Q08209\|PP2BA_HUMAN Serine/threonine-protein phosphatase 2B catalytic subunit | 66,328 | 6,14 | 58688 | 3 | 23,54 | 23,52 | 21,63 | 23,00 | 23,54 | 22,23 | **1,019** | 9,73E-01 | **0** | 0,02666664 | 0,01191777 |
| PPIA_HUMA | PPIA | sp\|P62937\|PPIA_HUMAN Peptidyl-prolyl cis-trans isomerase A OS=Homo sapiens GN=PI | 1170,011 | 72,12 | 18012 | 17 | 31,28 | 31,20 | 31,14 | 31,40 | 31,38 | 30,92 | **1,019** | 8,77E-01 | **0** | 0,02666601 | 0,05690745 |
| HSP76_HUM | HSPA6 | sp\|P17066\|HSP76_HUMAN Heat shock 70 kDa protein 6 OS=Homo sapiens GN=HSPA6 P | 562,022 | 17,88 | 71028 | 12 | 28,92 | 28,41 | 29,05 | 28,80 | 28,65 | 29,00 | **1,016** | 9,21E-01 | **0** | 0,02333323 | 0,03590126 |
| TXD12_HUM | TXNDC12 | sp\|O95881\|TXD12_HUMAN Thioredoxin domain-containing protein 12 OS=Homo sapier | 11,976 | 5,23 | 19206 | 1 | 21,18 | 21,38 | 22,70 | 21,83 | 21,85 | 21,65 | **1,016** | 9,64E-01 | **0** | 0,02333323 | 0,01608506 |
| BIN1_HUMA | BIN1 | sp\|O00499\|BIN1_HUMAN Myc box-dependent-interacting protein 1 OS=Homo sapiens | 605,27 | 32,88 | 64699 | 14 | 25,81 | 25,59 | 25,37 | 25,96 | 25,53 | 25,35 | **1,016** | 9,21E-01 | **0** | 0,02333323 | 0,03572737 |
| TIAR_HUMA | TIAL1 | sp\|Q01085\|TIAR_HUMAN Nucleolysin TIAR OS=Homo sapiens GN=TIAL1 PE=1 SV=1 | 139,073 | 15,73 | 41591 | 5 | 23,55 | 23,54 | 23,34 | 23,63 | 23,40 | 23,47 | **1,016** | 8,21E-01 | **0** | 0,0233326 | 0,08575051 |
| TRI56_HUM | TRIM56 | sp\|Q9BRZ2\|TRI56_HUMAN E3 ubiquitin-protein ligase TRIM56 OS=Homo sapiens GN=TI | 15,453 | 2,52 | 81488 | 1 | 18,48 | 17,45 | 20,08 | 18,03 | 17,87 | 20,17 | **1,016** | 9,84E-01 | **0** | 0,02258873 | 0,00695636 |
| COMD9_HUI | COMMD9 | sp\|Q9P000\|COMD9_HUMAN COMM domain-containing protein 9 OS=Homo sapiens GN | 39,8 | 8,08 | 21819 | 1 | 17,92 | 20,37 | 21,79 | 18,63 | 19,59 | 21,92 | **1,015** | 9,89E-01 | **0** | 0,02175458 | 0,00477079 |
| PGTB2_HUM | RABGGTB | sp\|P53611\|PGTB2_HUMAN Geranylgeranyl transferase type-2 subunit beta OS=Homo s | 49,822 | 3,32 | 36924 | 1 | 21,35 | 21,87 | 20,90 | 21,58 | 21,44 | 21,16 | **1,014** | 9,51E-01 | **0** | 0,01999982 | 0,02179011 |
| ECM29_HUN | ECM29 | sp\|Q5VYK3\|ECM29_HUMAN Proteasome-associated protein ECM29 homolog OS=Homo | 362,351 | 9,81 | 204291 | 11 | 22,44 | 22,53 | 24,76 | 22,69 | 22,41 | 24,69 | **1,014** | 9,86E-01 | **0** | 0,01999982 | 0,00628114 |
| WDR61_HUN | WDR61 | sp\|Q9GZS3\|WDR61_HUMAN WD repeat-containing protein 61 OS=Homo sapiens GN=W | 117,206 | 21,64 | 33581 | 3 | 19,28 | 20,05 | 22,62 | 18,80 | 20,15 | 23,06 | **1,014** | 9,91E-01 | **0** | 0,01999919 | 0,00405905 |
| SAMH1_HUN | SAMHD1 | sp\|Q9Y3Z3\|SAMH1_HUMAN Deoxynucleoside triphosphate triphosphohydrolase SAMH | 9,716 | 2,24 | 72201 | 1 | 19,51 | 19,77 | 20,78 | 20,00 | 19,55 | 20,57 | **1,014** | 9,69E-01 | **0** | 0,01999919 | 0,01358259 |
| F168A_HUM | FAM168A | sp\|Q92567\|F168A_HUMAN Protein FAM168A OS=Homo sapiens GN=FAM168A PE=1 SV | 12,057 | 4,1 | 26184 | 1 | 22,55 | 21,91 | 17,87 | 21,71 | 21,48 | 19,19 | **1,012** | 9,92E-01 | **0** | 0,01670647 | 0,00326997 |
| USP9X_HUM | USP9X | sp\|Q93008\|USP9X_HUMAN Probable ubiquitin carboxyl-terminal hydrolase FAF-X OS=H | 691,407 | 14,82 | 292280 | 30 | 25,49 | 25,67 | 25,51 | 25,34 | 25,77 | 25,61 | **1,012** | 9,10E-01 | **0** | 0,01666705 | 0,04116753 |
| GT251_HUM | COLGALT1 | sp\|Q8NBJ5\|GT251_HUMAN Procollagen galactosyltransferase 1 OS=Homo sapiens GN=( | 14,375 | 1,29 | 71636 | 1 | 21,16 | 20,92 | 20,88 | 21,44 | 20,85 | 20,72 | **1,012** | 9,48E-01 | **0** | 0,01666705 | 0,02338981 |
| RHG17_HUN | ARHGAP17 | sp\|Q68EM7\|RHG17_HUMAN Rho GTPase-activating protein 17 OS=Homo sapiens GN=A | 155,39 | 3,86 | 95437 | 3 | 22,19 | 22,09 | 22,78 | 22,28 | 22,56 | 22,27 | **1,012** | 9,47E-01 | **0** | 0,01666641 | 0,02367838 |
| OXSR1_HUM | OXSR1 | sp\|O95747\|OXSR1_HUMAN Serine/threonine-protein kinase OSR1 OS=Homo sapiens G | 201,817 | 13,28 | 58022 | 5 | 23,37 | 23,17 | 23,92 | 22,68 | 23,66 | 24,17 | **1,012** | 9,75E-01 | **0** | 0,01666641 | 0,01118864 |
| RAB8B_HUM | RAB8B | sp\|Q92930\|RAB8B_HUMAN Ras-related protein Rab-8B OS=Homo sapiens GN=RAB8B P | 320,503 | 47,34 | 23584 | 11 | 25,35 | 25,58 | 25,18 | 25,75 | 25,48 | 24,93 | **1,012** | 9,53E-01 | **0** | 0,01666641 | 0,02075302 |
| CSK21_HUM | CSNK2A1 | sp\|P68400\|CSK21_HUMAN Casein kinase II subunit alpha OS=Homo sapiens GN=CSNK2/ | 545,39 | 59,85 | 45144 | 20 | 25,42 | 25,82 | 27,49 | 25,60 | 25,87 | 27,30 | **1,009** | 9,88E-01 | **0** | 0,01333364 | 0,00529827 |
| SNAA_HUM | NAPA | sp\|P54920\|SNAA_HUMAN Alpha-soluble NSF attachment protein OS=Homo sapiens GN | 482,297 | 45,42 | 33233 | 10 | 25,73 | 25,68 | 25,10 | 25,64 | 25,83 | 25,08 | **1,009** | 9,67E-01 | **0** | 0,013333 | 0,01458915 |
| ARMC8_HUN | ARMC8 | sp\|Q8IUR7\|ARMC8_HUMAN Armadillo repeat-containing protein 8 OS=Homo sapiens G | 81,107 | 11,14 | 75509 | 4 | 18,44 | 19,62 | 21,04 | 18,11 | 19,10 | 21,93 | **1,009** | 9,93E-01 | **0** | 0,013333 | 0,00318312 |
| ICMT_HUMA | ICMT | sp\|O60725\|ICMT_HUMAN Protein-S-isoprenylcysteine O-methyltransferase OS=Homo s | 48,425 | 7,04 | 31938 | 1 | 18,39 | 18,84 | 19,09 | 18,97 | 18,39 | 19,00 | **1,009** | 9,65E-01 | **0** | 0,013333 | 0,01549191 |
| TCTP_HUMA | TPT1 | sp\|P13693\|TCTP_HUMAN Translationally-controlled tumor protein OS=Homo sapiens G | 440,402 | 68,6 | 19595 | 11 | 27,91 | 28,11 | 27,44 | 27,75 | 28,00 | 27,75 | **1,009** | 9,54E-01 | **0** | 0,013333 | 0,02063229 |
| HS71L_HUM | HSPA1L | sp\|P34931\|HS71L_HUMAN Heat shock 70 kDa protein 1-like OS=Homo sapiens GN=HSP/ | 420,629 | 20,59 | 70375 | 9 | 21,95 | 20,82 | 19,20 | 21,55 | 21,31 | 19,15 | **1,009** | 9,91E-01 | **0** | 0,01333237 | 0,00395059 |
| PIN4_HUMA | PIN4 | sp\|Q9Y237\|PIN4_HUMAN Peptidyl-prolyl cis-trans isomerase NIMA-interacting 4 OS=Ho | 58,306 | 16,03 | 13810 | 2 | 22,80 | 22,66 | 19,77 | 22,81 | 22,47 | 19,99 | **1,009** | 9,93E-01 | **0** | 0,0130043 | 0,00319923 |
| VAT1L_HUM | VAT1L | sp\|Q9HCJ6\|VAT1L_HUMAN Synaptic vesicle membrane protein VAT-1 homolog-like OS= | 26,366 | 3,1 | 45899 | 1 | 18,78 | 18,30 | 17,12 | 17,40 | 18,69 | 18,15 | **1,008** | 9,85E-01 | **0** | 0,01200549 | 0,00636112 |
| RBM4_HUM | RBM4 | sp\|Q9BWF3\|RBM4_HUMAN RNA-binding protein 4 OS=Homo sapiens GN=RBM4 PE=1 S | 222,189 | 28,02 | 40314 | 9 | 21,14 | 21,89 | 22,42 | 21,69 | 21,55 | 22,24 | **1,007** | 9,82E-01 | **0** | 0,01000023 | 0,00769747 |
| MK03_HUM | MAPK3 | sp\|P27361\|MK03_HUMAN Mitogen-activated protein kinase 3 OS=Homo sapiens GN=M | 189,674 | 25,33 | 43136 | 8 | 22,85 | 22,72 | 22,05 | 22,73 | 22,83 | 22,09 | **1,007** | 9,78E-01 | **0** | 0,01000023 | 0,00970413 |
| GDIR1_HUM | ARHGDIA | sp\|P52565\|GDIR1_HUMAN Rho GDP-dissociation inhibitor 1 OS=Homo sapiens GN=ARH | 419,34 | 48,04 | 23207 | 11 | 26,94 | 27,08 | 26,48 | 26,99 | 27,03 | 26,51 | **1,007** | 9,70E-01 | **0** | 0,01000023 | 0,01341541 |
| TBA4B_HUM | TUBA4B | sp\|Q9H853\|TBA4B_HUMAN Putative tubulin-like protein alpha-4B OS=Homo sapiens GN | 136,399 | 10,79 | 27551 | 2 | 25,97 | 26,81 | 27,34 | 26,15 | 26,65 | 27,35 | **1,007** | 9,86E-01 | **0** | 0,01000023 | 0,00619716 |
| CND3_HUM | NCAPG | sp\|Q9BPX3\|CND3_HUMAN Condensin complex subunit 3 OS=Homo sapiens GN=NCAPG | 57,274 | 2,46 | 114334 | 2 | 19,83 | 20,28 | 22,23 | 19,71 | 20,87 | 21,79 | **1,007** | 9,92E-01 | **0** | 0,01000023 | 0,00343817 |
| P4K2A_HUM | PI4K2A | sp\|Q9BTU6\|P4K2A_HUMAN Phosphatidylinositol 4-kinase type 2-alpha OS=Homo sapiei | 130,59 | 16,49 | 54022 | 7 | 23,79 | 23,66 | 22,68 | 23,64 | 24,10 | 22,42 | **1,007** | 9,88E-01 | **0** | 0,00999959 | 0,00535859 |
| RAB10_HUM | RAB10 | sp\|P61026\|RAB10_HUMAN Ras-related protein Rab-10 OS=Homo sapiens GN=RAB10 PE | 609,791 | 78,5 | 22541 | 17 | 26,84 | 26,41 | 27,38 | 26,87 | 26,74 | 27,04 | **1,005** | 9,83E-01 | **0** | 0,00666746 | 0,00745597 |
| ARL6_HUMA | ARL6 | sp\|Q9H0F7\|ARL6_HUMAN ADP-ribosylation factor-like protein 6 OS=Homo sapiens GN= | 10,719 | 4,84 | 21097 | 1 | 21,29 | 20,97 | 21,13 | 20,94 | 21,35 | 21,12 | **1,005** | 9,67E-01 | **0** | 0,00666746 | 0,01468204 |
| CDK1_HUMA | CDK1 | sp\|P06493\|CDK1_HUMAN Cyclin-dependent kinase 1 OS=Homo sapiens GN=CDK1 PE=1 | 650,356 | 68,35 | 34095 | 18 | 26,63 | 26,80 | 26,97 | 26,55 | 26,69 | 27,18 | **1,005** | 9,77E-01 | **0** | 0,00666746 | 0,01023106 |
| PSMD9_HUN | PSMD9 | sp\|O00233\|PSMD9_HUMAN 26S proteasome non-ATPase regulatory subunit 9 OS=Hom | 10,416 | 4,48 | 24682 | 1 | 22,22 | 21,46 | 21,31 | 22,42 | 21,42 | 21,17 | **1,005** | 9,89E-01 | **0** | 0,00666746 | 0,00460069 |
| TMOD3_HUI | TMOD3 | sp\|Q9NYL9\|TMOD3_HUMAN Tropomodulin-3 OS=Homo sapiens GN=TMOD3 PE=1 SV=: | 75,353 | 6,82 | 39595 | 2 | 22,07 | 22,51 | 22,31 | 22,62 | 22,55 | 21,74 | **1,005** | 9,84E-01 | **0** | 0,00666682 | 0,00706792 |
| SRGP2_HUM | SRGAP2 | sp\|O75044\|SRGP2_HUMAN SLIT-ROBO Rho GTPase-activating protein 2 OS=Homo sapie | 385,495 | 17,09 | 120881 | 14 | 25,15 | 24,90 | 22,05 | 24,91 | 24,75 | 22,46 | **1,005** | 9,96E-01 | **0** | 0,00666682 | 0,00171219 |
| NPTN_HUM | NPTN | sp\|Q9Y639\|NPTN_HUMAN Neuroplastin OS=Homo sapiens GN=NPTN PE=1 SV=2 | 110,434 | 6,78 | 44387 | 3 | 25,20 | 25,47 | 24,55 | 25,31 | 25,36 | 24,57 | **1,005** | 9,87E-01 | **0** | 0,00666682 | 0,00584705 |
| DJC13_HUM | DNAJC13 | sp\|O75165\|DJC13_HUMAN DnaJ homolog subfamily C member 13 OS=Homo sapiens GI | 283,518 | 6,55 | 254415 | 12 | 23,86 | 24,41 | 23,71 | 23,99 | 24,49 | 23,52 | **1,005** | 9,86E-01 | **0** | 0,00666682 | 0,00621774 |
| EXOC3_HUM | EXOC3 | sp\|O60645\|EXOC3_HUMAN Exocyst complex component 3 OS=Homo sapiens GN=EXO( | 30,51 | 3,7 | 86845 | 2 | 20,93 | 22,25 | 21,46 | 21,01 | 22,46 | 21,19 | **1,005** | 9,92E-01 | **0** | 0,00666682 | 0,00365839 |
| ERP29_HUM | ERP29 | sp\|P30040\|ERP29_HUMAN Endoplasmic reticulum resident protein 29 OS=Homo sapien | 90,322 | 17,24 | 28993 | 5 | 25,17 | 25,19 | 25,57 | 25,58 | 25,72 | 24,65 | **1,005** | 9,86E-01 | **0** | 0,00666618 | 0,00607143 |
| GNL1_HUM | GNL1 | sp\|P36915\|GNL1_HUMAN Guanine nucleotide-binding protein-like 1 OS=Homo sapiens | 48,275 | 4,61 | 68661 | 2 | 22,01 | 22,00 | 22,98 | 22,13 | 22,08 | 22,80 | **1,005** | 9,87E-01 | **0** | 0,00666618 | 0,0054706 |
| RS29_HUMA | RPS29 | sp\|P62273\|RS29_HUMAN 40S ribosomal protein S29 OS=Homo sapiens GN=RPS29 PE=1 | 51,883 | 46,43 | 6677 | 3 | 24,98 | 25,18 | 21,95 | 25,13 | 24,95 | 22,05 | **1,005** | 9,97E-01 | **0** | 0,00666618 | 0,00150528 |
| OBSCN_HUN | OBSCN | sp\|Q5VST9\|OBSCN_HUMAN Obscurin OS=Homo sapiens GN=OBSCN PE=1 SV=3 | 9,546 | 0,09 | 868484 | 1 | 21,11 | 19,62 | 21,94 | 21,13 | 19,52 | 22,04 | **1,005** | 9,95E-01 | **0** | 0,00666618 | 0,00217299 |
| VTNC_HUM | VTN | sp\|P04004\|VTNC_HUMAN Vitronectin OS=Homo sapiens GN=VTN PE=1 SV=1 | 98,274 | 10,67 | 54306 | 3 | 24,75 | 23,88 | 27,60 | 24,68 | 24,59 | 26,97 | **1,002** | 9,98E-01 | **0** | 0,00333341 | 0,00079502 |
| ILF2_HUMAN | ILF2 | sp\|Q12905\|ILF2_HUMAN Interleukin enhancer-binding factor 2 OS=Homo sapiens GN=I | 852,997 | 71,54 | 43062 | 18 | 27,32 | 27,30 | 27,53 | 27,23 | 27,28 | 27,65 | **1,002** | 9,84E-01 | **0** | 0,00333341 | 0,00722527 |
| STAT3_HUM | STAT3 | sp\|P40763\|STAT3_HUMAN Signal transducer and activator of transcription 3 OS=Homo | 267,266 | 23,9 | 88068 | 10 | 24,11 | 24,05 | 23,98 | 23,65 | 24,03 | 24,47 | **1,002** | 9,90E-01 | **0** | 0,00333341 | 0,00454976 |
| H31_HUMAN | HIST1H3A | sp\|P68431\|H31_HUMAN Histone H3.1 OS=Homo sapiens GN=HIST1H3A PE=1 SV=2 | 246,62 | 43,38 | 15404 | 6 | 22,02 | 22,14 | 22,81 | 22,35 | 22,40 | 22,23 | **1,002** | 9,90E-01 | **0** | 0,00333341 | 0,00434885 |
| ACTB_HUMA | ACTB | sp\|P60709\|ACTB_HUMAN Actin, cytoplasmic 1 OS=Homo sapiens GN=ACTB PE=1 SV=1 | 4172,782 | 97,6 | 41737 | 78 | 27,96 | 27,97 | 27,28 | 27,96 | 28,30 | 26,96 | **1,002** | 9,95E-01 | **0** | 0,00333277 | 0,00235364 |
| DNJB1_HUM | DNAJB1 | sp\|P25685\|DNJB1_HUMAN DnaJ homolog subfamily B member 1 OS=Homo sapiens GN: | 284,745 | 21,18 | 38044 | 8 | 25,08 | 25,18 | 24,46 | 24,96 | 25,06 | 24,71 | **1,002** | 9,90E-01 | **0** | 0,00333277 | 0,00439782 |
| MASP1_HUN | MASP1 | sp\|P48740\|MASP1_HUMAN Mannan-binding lectin serine protease 1 OS=Homo sapiens | 30,19 | 1,57 | 79247 | 1 | 18,41 | 19,78 | 20,99 | 20,11 | 18,94 | 20,14 | **1,002** | 9,97E-01 | **0** | 0,0031573 | 0,00122016 |
| HYAS1_HUM | HAS1 | sp\|Q92839\|HYAS1_HUMAN Hyaluronan synthase 1 OS=Homo sapiens GN=HAS1 PE=2 S\ | 36,827 | 1,38 | 64832 | 1 | 25,07 | 25,58 | 24,43 | 24,98 | 25,36 | 24,74 | **1,000** | 1,00E+00 | **0** | 3,5238E-15 | 0 |
| RRAS_HUMA | RRAS | sp\|P10301\|RRAS_HUMAN Ras-related protein R-Ras OS=Homo sapiens GN=RRAS PE=1 S | 618,746 | 70,18 | 23480 | 13 | 26,18 | 26,23 | 26,08 | 26,06 | 26,70 | 25,73 | **1,000** | 1,00E+00 | **0** | 0 | 0 |
| ARH_HUMAN | LDLRAP1 | sp\|Q5SW96\|ARH_HUMAN Low density lipoprotein receptor adapter protein 1 OS=Hom | 43,946 | 3,9 | 33885 | 1 | 20,48 | 20,56 | 21,28 | 20,37 | 20,52 | 21,42 | **0,998** | 9,94E-01 | **0** | -0,0033328 | 0,00262377 |
| DHX30_HUM | DHX30 | sp\|Q7L2E3\|DHX30_HUMAN Putative ATP-dependent RNA helicase DHX30 OS=Homo sap | 146,792 | 5,36 | 133938 | 5 | 21,40 | 21,26 | 22,59 | 21,27 | 20,79 | 23,18 | **0,998** | 9,97E-01 | **0** | -0,0033328 | 0,00128946 |
| GSTP1_HUM | GSTP1 | sp\|P09211\|GSTP1_HUMAN Glutathione S-transferase P OS=Homo sapiens GN=GSTP1 PI | 68,343 | 8,1 | 23356 | 2 | 29,04 | 28,92 | 26,78 | 28,84 | 28,86 | 27,03 | **0,998** | 9,97E-01 | **0** | -0,0033334 | 0,00114153 |
| M3K20_HUN | MAP3K20 | sp\|Q9NYL2\|M3K20_HUMAN Mitogen-activated protein kinase kinase kinase 20 OS=Hon | 31,915 | 2,75 | 91155 | 2 | 21,48 | 22,07 | 22,19 | 21,40 | 22,47 | 21,86 | **0,998** | 9,93E-01 | **0** | -0,0033334 | 0,00286888 |
| STML3_HUM | STOML3 | sp\|Q8TAV4\|STML3_HUMAN Stomatin-like protein 3 OS=Homo sapiens GN=STOML3 PE= | 17,798 | 2,41 | 32135 | 1 | 25,61 | 24,31 | 23,35 | 25,61 | 24,34 | 23,31 | **0,998** | 9,97E-01 | **0** | -0,0033334 | 0,00116476 |
| IPO7_HUMA | IPO7 | sp\|O95373\|IPO7_HUMAN Importin-7 OS=Homo sapiens GN=IPO7 PE=1 SV=1 | 898,784 | 29 | 119517 | 20 | 25,66 | 26,13 | 26,75 | 25,47 | 26,25 | 26,81 | **0,998** | 9,95E-01 | **0** | -0,0033334 | 0,00217429 |
| ACTG_HUM | ACTG1 | sp\|P63261\|ACTG_HUMAN Actin, cytoplasmic 2 OS=Homo sapiens GN=ACTG1 PE=1 SV=1 | 4186,057 | 97,6 | 41793 | 78 | 27,19 | 27,50 | 26,60 | 27,18 | 27,51 | 26,59 | **0,998** | 9,93E-01 | **0** | -0,0033334 | 0,00288996 |
| SRSF9_HUM | SRSF9 | sp\|Q13242\|SRSF9_HUMAN Serine/arginine-rich splicing factor 9 OS=Homo sapiens GN= | 38,327 | 9,5 | 25542 | 2 | 22,33 | 22,79 | 23,21 | 22,78 | 22,48 | 23,06 | **0,998** | 9,92E-01 | **0** | -0,0033334 | 0,00358225 |
| ACTN1_HUM | ACTN1 | sp\|P12814\|ACTN1_HUMAN Alpha-actinin-1 OS=Homo sapiens GN=ACTN1 PE=1 SV=2 | 3339,105 | 70,29 | 103058 | 71 | 28,94 | 29,18 | 28,47 | 28,90 | 29,21 | 28,47 | **0,998** | 9,92E-01 | **0** | -0,003334 | 0,00364492 |
| RAB2A_HUM | RAB2A | sp\|P61019\|RAB2A_HUMAN Ras-related protein Rab-2A OS=Homo sapiens GN=RAB2A PI | 635,434 | 76,42 | 23546 | 16 | 27,41 | 27,36 | 27,53 | 27,21 | 27,56 | 27,52 | **0,998** | 9,79E-01 | **0** | -0,003334 | 0,00902502 |
| UGDH_HUM | UGDH | sp\|O60701\|UGDH_HUMAN UDP-glucose 6-dehydrogenase OS=Homo sapiens GN=UGDI | 667,107 | 38,87 | 55024 | 13 | 26,20 | 26,45 | 26,08 | 26,06 | 26,58 | 26,08 | **0,998** | 9,88E-01 | **0** | -0,003334 | 0,00540884 |
| RL4_HUMAN | RPL4 | sp\|P36578\|RL4_HUMAN 60S ribosomal protein L4 OS=Homo sapiens GN=RPL4 PE=1 SV= | 588,527 | 37,7 | 47697 | 18 | 27,04 | 27,62 | 27,04 | 27,07 | 26,80 | 27,82 | **0,998** | 9,93E-01 | **0** | -0,0033347 | 0,00301739 |
| YIPF5_HUM | YIPF5 | sp\|Q969M3\|YIPF5_HUMAN Protein YIPF5 OS=Homo sapiens GN=YIPF5 PE=1 SV=1 | 15,491 | 3,5 | 27989 | 1 | 21,52 | 21,28 | 19,27 | 21,73 | 21,23 | 19,09 | **0,996** | 9,96E-01 | **0** | -0,0063063 | 0,00190462 |
| MARK3_HUN | MARK3 | sp\|P27448\|MARK3_HUMAN MAP/microtubule affinity-regulating kinase 3 OS=Homo sap | 148,244 | 9,96 | 84489 | 6 | 21,97 | 22,21 | 20,73 | 22,04 | 21,48 | 21,37 | **0,995** | 9,90E-01 | **0** | -0,0066655 | 0,00433486 |
| MARE1_HUN | MAPRE1 | sp\|Q15691\|MARE1_HUMAN Microtubule-associated protein RP/EB family member 1 OS | 362,552 | 57,09 | 29999 | 11 | 25,21 | 25,12 | 24,73 | 25,18 | 25,17 | 24,69 | **0,995** | 9,77E-01 | **0** | -0,0066662 | 0,01003999 |
| SMC2_HUM | SMC2 | sp\|O95347\|SMC2_HUMAN Structural maintenance of chromosomes protein 2 OS=Hom | 484,927 | 18,88 | 135656 | 19 | 24,52 | 24,79 | 25,89 | 24,52 | 24,70 | 25,96 | **0,995** | 9,92E-01 | **0** | -0,0066668 | 0,00353348 |
| STXB3_HUM | STXBP3 | sp\|O00186\|STXB3_HUMAN Syntaxin-binding protein 3 OS=Homo sapiens GN=STXBP3 P | 820,171 | 31,25 | 67764 | 21 | 28,18 | 27,92 | 26,72 | 28,06 | 27,90 | 26,84 | **0,995** | 9,92E-01 | **0** | -0,0066668 | 0,00369288 |
| NTM1A_HUN | NTMT1 | sp\|Q9BV86\|NTM1A_HUMAN N-terminal Xaa-Pro-Lys N-methyltransferase 1 OS=Homo s | 60,09 | 7,17 | 25387 | 2 | 22,66 | 22,64 | 23,54 | 22,55 | 22,77 | 23,50 | **0,995** | 9,88E-01 | **0** | -0,0066668 | 0,00529114 |
| IMA8_HUMA | KPNA7 | sp\|A9QM74\|IMA8_HUMAN Importin subunit alpha-8 OS=Homo sapiens GN=KPNA7 PE= | 37,132 | 2,13 | 56938 | 1 | 23,09 | 23,53 | 24,10 | 23,28 | 23,61 | 23,81 | **0,995** | 9,85E-01 | **0** | -0,0066668 | 0,00661624 |
| TBG1_HUMA | TUBG1 | sp\|P23258\|TBG1_HUMAN Tubulin gamma-1 chain OS=Homo sapiens GN=TUBG1 PE=1 S | 183,682 | 14,63 | 51170 | 5 | 22,92 | 22,68 | 23,62 | 22,57 | 22,79 | 23,84 | **0,995** | 9,90E-01 | **0** | -0,0066668 | 0,00452143 |
| VPS25_HUM | VPS25 | sp\|Q9BRG1\|VPS25_HUMAN Vacuolar protein-sorting-associated protein 25 OS=Homo s | 214,962 | 52,84 | 20748 | 7 | 25,77 | 25,45 | 24,00 | 25,49 | 25,42 | 24,29 | **0,995** | 9,93E-01 | **0** | -0,0066668 | 0,00325734 |
| SEPT4_HUM | SEPT4 | sp\|O43236\|SEPT4_HUMAN Septin-4 OS=Homo sapiens GN=SEPT4 PE=1 SV=1 | 11,63 | 1,88 | 55098 | 1 | 20,61 | 20,36 | 21,81 | 20,35 | 20,14 | 22,27 | **0,995** | 9,94E-01 | **0** | -0,0066668 | 0,00268204 |
| SEPT5_HUM | SEPT5 | sp\|Q99719\|SEPT5_HUMAN Septin-5 OS=Homo sapiens GN=SEPT5 PE=1 SV=1 | 11,63 | 2,44 | 42777 | 1 | 20,61 | 20,36 | 21,81 | 20,35 | 20,14 | 22,27 | **0,995** | 9,94E-01 | **0** | -0,0066668 | 0,00268204 |
| AL3A2_HUM | ALDH3A2 | sp\|P51648\|AL3A2_HUMAN Fatty aldehyde dehydrogenase OS=Homo sapiens GN=ALDH | 66,66 | 4,95 | 54848 | 2 | 22,11 | 22,18 | 22,79 | 22,13 | 22,30 | 22,63 | **0,995** | 9,81E-01 | **0** | -0,0066681 | 0,00839766 |
| PUS7_HUMA | PUS7 | sp\|Q96PZ0\|PUS7_HUMAN Pseudouridylate synthase 7 homolog OS=Homo sapiens GN= | 22,512 | 1,66 | 75035 | 1 | 18,27 | 19,93 | 17,65 | 17,37 | 18,25 | 20,20 | **0,995** | 9,95E-01 | **0** | -0,006752 | 0,00204647 |
| LRCH3_HUM | LRCH3 | sp\|Q96II8\|LRCH3_HUMAN Leucine-rich repeat and calponin homology domain-containi | 13,825 | 1,54 | 86083 | 1 | 17,85 | 18,13 | 19,78 | 18,47 | 17,60 | 19,67 | **0,995** | 9,93E-01 | **0** | -0,0074088 | 0,00284876 |
| SGPL1_HUM | SGPL1 | sp\|O95470\|SGPL1_HUMAN Sphingosine-1-phosphate lyase 1 OS=Homo sapiens GN=SG | 87,784 | 6,69 | 63524 | 2 | 18,40 | 17,56 | 20,90 | 17,39 | 18,14 | 21,31 | **0,994** | 9,96E-01 | **0** | -0,0092456 | 0,00192712 |
| RAB14_HUM | RAB14 | sp\|P61106\|RAB14_HUMAN Ras-related protein Rab-14 OS=Homo sapiens GN=RAB14 PE | 886,212 | 85,58 | 23897 | 21 | 26,98 | 26,93 | 27,67 | 26,94 | 27,10 | 27,51 | **0,993** | 9,74E-01 | **0** | -0,0099996 | 0,01126006 |
| S10A8_HUM | S100A8 | sp\|P05109\|S10A8_HUMAN Protein S100-A8 OS=Homo sapiens GN=S100A8 PE=1 SV=1 | 29,678 | 20,43 | 10835 | 2 | 22,08 | 21,20 | 22,85 | 22,03 | 21,70 | 22,37 | **0,993** | 9,85E-01 | **0** | -0,0099996 | 0,00637772 |
| RBM4B_HUN | RBM4B | sp\|Q9BQ04\|RBM4B_HUMAN RNA-binding protein 4B OS=Homo sapiens GN=RBM4B PE: | 187,885 | 25,07 | 40150 | 9 | 21,71 | 21,36 | 22,18 | 21,76 | 21,61 | 21,85 | **0,993** | 9,70E-01 | **0** | -0,0099996 | 0,01335028 |
| TMEM2_HUI | TMEM2 | sp\|Q9UHN6\|TMEM2_HUMAN Cell surface hyaluronidase OS=Homo sapiens GN=TMEM: | 40,497 | 1,59 | 154374 | 2 | 22,38 | 22,52 | 20,08 | 22,28 | 22,32 | 20,35 | **0,993** | 9,93E-01 | **0** | -0,0099996 | 0,0031927 |
| SNX5_HUMA | SNX5 | sp\|Q9Y5X3\|SNX5_HUMAN Sorting nexin-5 OS=Homo sapiens GN=SNX5 PE=1 SV=1 | 134,91 | 15,1 | 46816 | 5 | 24,92 | 24,94 | 24,01 | 24,80 | 24,95 | 24,09 | **0,993** | 9,82E-01 | **0** | -0,0100002 | 0,00810714 |
| RS30_HUMA | FAU | sp\|P62861\|RS30_HUMAN 40S ribosomal protein S30 OS=Homo sapiens GN=FAU PE=1 S' | 53,306 | 20,34 | 6648 | 3 | 24,89 | 24,87 | 21,95 | 24,97 | 24,85 | 21,86 | **0,993** | 9,95E-01 | **0** | -0,0100002 | 0,00231591 |
| GNA12_HUN | GNA12 | sp\|Q03113\|GNA12_HUMAN Guanine nucleotide-binding protein subunit alpha-12 OS=H | 85,815 | 7,09 | 44279 | 3 | 21,22 | 20,94 | 21,07 | 21,31 | 21,30 | 20,59 | **0,993** | 9,70E-01 | **0** | -0,0100002 | 0,01313322 |
| RB11B_HUM | RAB11B | sp\|Q15907\|RB11B_HUMAN Ras-related protein Rab-11B OS=Homo sapiens GN=RAB11B | 561,009 | 59,17 | 24489 | 14 | 23,91 | 24,20 | 22,97 | 23,88 | 24,31 | 22,86 | **0,993** | 9,87E-01 | **0** | -0,0100002 | 0,00577189 |
| TLN2_HUMA | TLN2 | sp\|Q9Y4G6\|TLN2_HUMAN Talin-2 OS=Homo sapiens GN=TLN2 PE=1 SV=4 | 544,03 | 6,22 | 271613 | 14 | 22,00 | 22,34 | 18,97 | 22,06 | 22,24 | 18,98 | **0,993** | 9,95E-01 | **0** | -0,0100002 | 0,00216892 |
| FLNB_HUMA | FLNB | sp\|O75369\|FLNB_HUMAN Filamin-B OS=Homo sapiens GN=FLNB PE=1 SV=2 | 2841,446 | 41,08 | 278164 | 77 | 28,32 | 28,60 | 27,93 | 28,29 | 28,55 | 27,98 | **0,993** | 9,71E-01 | **0** | -0,0100002 | 0,01297485 |
| ASPH_HUMA | ASPH | sp\|Q12797\|ASPH_HUMAN Aspartyl/asparaginyl beta-hydroxylase OS=Homo sapiens GN | 20,619 | 1,85 | 85863 | 1 | 18,44 | 18,38 | 19,78 | 18,10 | 18,60 | 19,87 | **0,993** | 9,88E-01 | **0** | -0,0107212 | 0,00503385 |
| HNRPQ_HUN | SYNCRIP | sp\|O60506\|HNRPQ_HUMAN Heterogeneous nuclear ribonucleoprotein Q OS=Homo sap | 689,755 | 33,55 | 69603 | 18 | 25,71 | 25,91 | 26,65 | 25,68 | 26,11 | 26,44 | **0,991** | 9,72E-01 | **0** | -0,0133324 | 0,01220405 |
| RFTN1_HUM | RFTN1 | sp\|Q14699\|RFTN1_HUMAN Raftlin OS=Homo sapiens GN=RFTN1 PE=1 SV=4 | 51,7 | 5,54 | 63146 | 4 | 25,35 | 25,05 | 21,88 | 25,05 | 24,84 | 22,35 | **0,991** | 9,93E-01 | **0** | -0,013333 | 0,00309406 |
| CYFP2_HUM | CYFIP2 | sp\|Q96F07\|CYFP2_HUMAN Cytoplasmic FMR1-interacting protein 2 OS=Homo sapiens ( | 728,218 | 17,45 | 148399 | 20 | 23,26 | 22,54 | 22,38 | 22,79 | 22,91 | 22,44 | **0,991** | 9,67E-01 | **0** | -0,013333 | 0,01446392 |
| NSF_HUMAN | NSF | sp\|P46459\|NSF_HUMAN Vesicle-fusing ATPase OS=Homo sapiens GN=NSF PE=1 SV=3 | 536,992 | 31,72 | 82594 | 19 | 25,68 | 25,41 | 25,71 | 25,62 | 25,57 | 25,57 | **0,991** | 8,97E-01 | **0** | -0,013333 | 0,04713781 |
| CYTB_HUMA | CSTB | sp\|P04080\|CYTB_HUMAN Cystatin-B OS=Homo sapiens GN=CSTB PE=1 SV=2 | 46,457 | 12,24 | 11140 | 1 | 22,66 | 22,72 | 21,95 | 22,27 | 22,76 | 22,26 | **0,991** | 9,66E-01 | **0** | -0,013333 | 0,01485341 |
| BUD31_HUN | BUD31 | sp\|P41223\|BUD31_HUMAN Protein BUD31 homolog OS=Homo sapiens GN=BUD31 PE=: | 33,967 | 13,19 | 17000 | 2 | 20,96 | 20,96 | 22,08 | 20,91 | 21,25 | 21,80 | **0,991** | 9,78E-01 | **0** | -0,013333 | 0,00965916 |
| THOP1_HUN | THOP1 | sp\|P52888\|THOP1_HUMAN Thimet oligopeptidase OS=Homo sapiens GN=THOP1 PE=1 S | 185,681 | 9,87 | 78840 | 6 | 23,57 | 23,90 | 22,04 | 23,99 | 23,77 | 21,71 | **0,991** | 9,89E-01 | **0** | -0,0133336 | 0,00472077 |
| CC116_HUM | CCDC116 | sp\|Q8IYX3\|CC116_HUMAN Coiled-coil domain-containing protein 116 OS=Homo sapien: | 12,309 | 1,36 | 56940 | 1 | 22,76 | 22,86 | 22,98 | 22,73 | 22,65 | 23,18 | **0,991** | 9,44E-01 | **0** | -0,0133336 | 0,0252571 |
| PCNP_HUMA | PCNP | sp\|Q8WW12\|PCNP_HUMAN PEST proteolytic signal-containing nuclear protein OS=Hom | 29,44 | 8,99 | 18925 | 1 | 20,42 | 20,70 | 19,38 | 20,85 | 20,21 | 19,40 | **0,991** | 9,83E-01 | **0** | -0,0133336 | 0,0075428 |
| EIF3E_HUM | EIF3E | sp\|P60228\|EIF3E_HUMAN Eukaryotic translation initiation factor 3 subunit E OS=Homo : | 654,222 | 53,03 | 52221 | 19 | 26,61 | 26,86 | 26,87 | 26,45 | 26,81 | 27,04 | **0,991** | 9,48E-01 | **0** | -0,0133336 | 0,02325593 |
| VP13C_HUM | VPS13C | sp\|Q709C8\|VP13C_HUMAN Vacuolar protein sorting-associated protein 13C OS=Homo : | 14,309 | 0,32 | 422390 | 1 | 20,12 | 20,68 | 19,52 | 20,22 | 20,35 | 19,71 | **0,991** | 9,74E-01 | **0** | -0,0133343 | 0,01134637 |
| ARP2_HUMA | ACTR2 | sp\|P61160\|ARP2_HUMAN Actin-related protein 2 OS=Homo sapiens GN=ACTR2 PE=1 SV | 772,031 | 56,35 | 44761 | 19 | 27,43 | 27,35 | 26,76 | 27,38 | 27,39 | 26,73 | **0,991** | 9,67E-01 | **0** | -0,0133343 | 0,01452976 |
| INCE_HUMA | INCENP | sp\|Q9NQS7\|INCE_HUMAN Inner centromere protein OS=Homo sapiens GN=INCENP PE: | 10,016 | 0,98 | 105429 | 1 | 17,53 | 18,27 | 21,73 | 18,99 | 17,80 | 20,69 | **0,989** | 9,92E-01 | **0** | -0,0163612 | 0,00346884 |
| KTHY_HUMA | DTYMK | sp\|P23919\|KTHY_HUMAN Thymidylate kinase OS=Homo sapiens GN=DTYMK PE=1 SV=4 | 35,741 | 5,19 | 23819 | 1 | 25,09 | 24,64 | 23,94 | 24,93 | 24,85 | 23,84 | **0,989** | 9,74E-01 | **0** | -0,0166664 | 0,01134296 |
| S10AA_HUM | S100A10 | sp\|P60903\|S10AA_HUMAN Protein S100-A10 OS=Homo sapiens GN=S100A10 PE=1 SV=: | 403,954 | 85,57 | 11203 | 11 | 27,64 | 27,99 | 27,25 | 27,74 | 27,95 | 27,14 | **0,989** | 9,61E-01 | **0** | -0,0166664 | 0,01711018 |
| MEMO1_HU | MEMO1 | sp\|Q9Y316\|MEMO1_HUMAN Protein MEMO1 OS=Homo sapiens GN=MEMO1 PE=1 SV= | 197,202 | 29,29 | 33733 | 7 | 24,99 | 24,55 | 23,18 | 24,64 | 24,52 | 23,51 | **0,989** | 9,81E-01 | **0** | -0,0166664 | 0,00840237 |
| MP2K3_HUN | MAP2K3 | sp\|P46734\|MP2K3_HUMAN Dual specificity mitogen-activated protein kinase kinase 3 O | 81,717 | 19,02 | 39318 | 5 | 22,71 | 22,95 | 23,10 | 23,07 | 22,85 | 22,79 | **0,989** | 9,12E-01 | **0** | -0,0166664 | 0,03991821 |
| VPS11_HUM | VPS11 | sp\|Q9H270\|VPS11_HUMAN Vacuolar protein sorting-associated protein 11 homolog OS | 45,032 | 2,55 | 107837 | 2 | 21,52 | 21,26 | 20,51 | 21,06 | 20,95 | 21,23 | **0,989** | 9,60E-01 | **0** | -0,016667 | 0,01765821 |
| GNA13_HUN | GNA13 | sp\|Q14344\|GNA13_HUMAN Guanine nucleotide-binding protein subunit alpha-13 OS=H | 474,23 | 37,93 | 44050 | 13 | 26,08 | 25,83 | 25,45 | 26,15 | 25,96 | 25,20 | **0,989** | 9,64E-01 | **0** | -0,016667 | 0,01610601 |
| HERC4_HUM | HERC4 | sp\|Q5GLZ8\|HERC4_HUMAN Probable E3 ubiquitin-protein ligase HERC4 OS=Homo sapie | 232,352 | 11,26 | 118563 | 8 | 23,86 | 23,80 | 23,33 | 23,39 | 24,15 | 23,40 | **0,989** | 9,59E-01 | **0** | -0,016667 | 0,01832456 |
| RN114_HUM | RNF114 | sp\|Q9Y508\|RN114_HUMAN E3 ubiquitin-protein ligase RNF114 OS=Homo sapiens GN=R | 66,244 | 14,47 | 25694 | 3 | 21,81 | 22,20 | 20,45 | 21,96 | 21,76 | 20,69 | **0,989** | 9,81E-01 | **0** | -0,016667 | 0,00829194 |
| GPC5C_HUM | GPRC5C | sp\|Q9NQ84\|GPC5C_HUMAN G-protein coupled receptor family C group 5 member C OS | 20,819 | 2,04 | 48193 | 1 | 24,56 | 23,87 | 19,94 | 24,20 | 23,55 | 20,57 | **0,988** | 9,93E-01 | **0** | -0,0168597 | 0,00302496 |
| WIPI2_HUM | WIPI2 | sp\|Q9Y4P8\|WIPI2_HUMAN WD repeat domain phosphoinositide-interacting protein 2 C | 26,515 | 3,52 | 49408 | 1 | 19,14 | 18,75 | 20,96 | 19,34 | 18,61 | 20,84 | **0,988** | 9,86E-01 | **0** | -0,0177574 | 0,00615576 |
| NUP88_HUN | NUP88 | sp\|Q99567\|NUP88_HUMAN Nuclear pore complex protein Nup88 OS=Homo sapiens GN | 56,244 | 1,75 | 83542 | 1 | 19,86 | 19,54 | 20,58 | 19,77 | 19,50 | 20,65 | **0,986** | 9,68E-01 | **0** | -0,0200005 | 0,01427222 |
| SPT4H_HUM | SUPT4H1 | sp\|P63272\|SPT4H_HUMAN Transcription elongation factor SPT4 OS=Homo sapiens GN= | 26,243 | 13,68 | 13193 | 1 | 21,39 | 21,82 | 18,04 | 21,60 | 21,98 | 17,61 | **0,986** | 9,92E-01 | **0** | -0,0203546 | 0,00361864 |
| ADH1A_HUN | ADH1A | sp\|P07327\|ADH1A_HUMAN Alcohol dehydrogenase 1A OS=Homo sapiens GN=ADH1A P | 20,911 | 2,67 | 39859 | 2 | 24,13 | 24,30 | 23,87 | 24,09 | 24,15 | 23,99 | **0,984** | 8,70E-01 | **0** | -0,0233332 | 0,06062984 |
| PURA_HUM | PURA | sp\|Q00577\|PURA_HUMAN Transcriptional activator protein Pur-alpha OS=Homo sapien | 259,516 | 23,29 | 34911 | 5 | 25,42 | 24,88 | 25,51 | 25,32 | 25,12 | 25,30 | **0,984** | 9,16E-01 | **0** | -0,0233332 | 0,03830375 |
| MYOF_HUM | MYOF | sp\|Q9NZM1\|MYOF_HUMAN Myoferlin OS=Homo sapiens GN=MYOF PE=1 SV=1 | 1438,361 | 25,42 | 234709 | 43 | 27,76 | 27,95 | 27,64 | 27,89 | 27,87 | 27,52 | **0,984** | 8,84E-01 | **0** | -0,0233332 | 0,05349052 |
| RHOB_HUM | RHOB | sp\|P62745\|RHOB_HUMAN Rho-related GTP-binding protein RhoB OS=Homo sapiens GN | 262,742 | 37,24 | 22123 | 7 | 24,77 | 23,92 | 23,77 | 24,78 | 24,23 | 23,38 | **0,984** | 9,66E-01 | **0** | -0,0233339 | 0,01507884 |
| XPO7_HUM | XPO7 | sp\|Q9UIA9\|XPO7_HUMAN Exportin-7 OS=Homo sapiens GN=XPO7 PE=1 SV=3 | 70,252 | 3,5 | 123907 | 3 | 20,97 | 20,61 | 20,70 | 19,88 | 21,23 | 21,10 | **0,984** | 9,61E-01 | **0** | -0,0233339 | 0,01747933 |
| ADAS_HUM | AGPS | sp\|O00116\|ADAS_HUMAN Alkyldihydroxyacetonephosphate synthase, peroxisomal OS= | 18,331 | 1,98 | 72912 | 1 | 18,14 | 17,69 | 21,61 | 17,59 | 18,37 | 21,41 | **0,983** | 9,89E-01 | **0** | -0,0246614 | 0,00474965 |
| PROF1_HUM | PFN1 | sp\|P07737\|PROF1_HUMAN Profilin-1 OS=Homo sapiens GN=PFN1 PE=1 SV=2 | 665,222 | 70 | 15054 | 17 | 29,89 | 29,87 | 29,06 | 29,84 | 29,95 | 28,95 | **0,982** | 9,52E-01 | **0** | -0,026666 | 0,02125839 |
| PGTB1_HUM | PGGT1B | sp\|P53609\|PGTB1_HUMAN Geranylgeranyl transferase type-1 subunit beta OS=Homo s | 12,14 | 2,65 | 42368 | 1 | 19,97 | 20,38 | 20,48 | 20,13 | 19,70 | 20,92 | **0,982** | 9,49E-01 | **0** | -0,026666 | 0,02284888 |
| TNPO3_HUN | TNPO3 | sp\|Q9Y5L0\|TNPO3_HUMAN Transportin-3 OS=Homo sapiens GN=TNPO3 PE=1 SV=3 | 662,987 | 29,79 | 104203 | 17 | 24,85 | 24,98 | 25,74 | 24,84 | 24,80 | 25,85 | **0,982** | 9,55E-01 | **0** | -0,0266666 | 0,02011102 |
| NUCB2_HUN | NUCB2 | sp\|P80303\|NUCB2_HUMAN Nucleobindin-2 OS=Homo sapiens GN=NUCB2 PE=1 SV=2 | 65,19 | 4,52 | 50196 | 2 | 22,12 | 21,65 | 22,71 | 22,24 | 22,09 | 22,07 | **0,982** | 9,36E-01 | **0** | -0,0266666 | 0,02879261 |
| MOB2_HUM | MOB2 | sp\|Q70IA6\|MOB2_HUMAN MOB kinase activator 2 OS=Homo sapiens GN=MOB2 PE=1 S | 35,787 | 4,64 | 26927 | 1 | 21,20 | 20,87 | 21,46 | 20,95 | 20,94 | 21,56 | **0,982** | 9,25E-01 | **0** | -0,0266666 | 0,03376876 |
| PAXI_HUMA | PXN | sp\|P49023\|PAXI_HUMAN Paxillin OS=Homo sapiens GN=PXN PE=1 SV=3 | 204,701 | 15,4 | 64505 | 7 | 25,08 | 24,93 | 24,45 | 25,04 | 25,03 | 24,31 | **0,982** | 9,35E-01 | **0** | -0,0266666 | 0,02916716 |
| EGFR_HUMA | EGFR | sp\|P00533\|EGFR_HUMAN Epidermal growth factor receptor OS=Homo sapiens GN=EGF | 217,988 | 6,94 | 134277 | 8 | 23,26 | 23,37 | 22,53 | 23,60 | 22,98 | 22,50 | **0,982** | 9,52E-01 | **0** | -0,0266673 | 0,02152033 |
| ARL3_HUMA | ARL3 | sp\|P36405\|ARL3_HUMAN ADP-ribosylation factor-like protein 3 OS=Homo sapiens GN=/ | 490,195 | 63,74 | 20456 | 9 | 25,31 | 25,45 | 26,54 | 25,22 | 25,44 | 26,56 | **0,982** | 9,65E-01 | **0** | -0,0266673 | 0,01554485 |
| SNAG_HUM | NAPG | sp\|Q99747\|SNAG_HUMAN Gamma-soluble NSF attachment protein OS=Homo sapiens ( | 342,842 | 42,63 | 34746 | 12 | 26,38 | 26,45 | 24,31 | 26,36 | 26,16 | 24,53 | **0,979** | 9,75E-01 | **0** | -0,0299994 | 0,01086686 |
| VP26A_HUM | VPS26A | sp\|O75436\|VP26A_HUMAN Vacuolar protein sorting-associated protein 26A OS=Homo : | 196,822 | 22,32 | 38170 | 5 | 23,30 | 22,69 | 23,46 | 22,51 | 23,43 | 23,42 | **0,979** | 9,42E-01 | **0** | -0,0299994 | 0,02613253 |
| MRP1_HUM | ABCC1 | sp\|P33527\|MRP1_HUMAN Multidrug resistance-associated protein 1 OS=Homo sapiens | 962,587 | 19,4 | 171591 | 25 | 27,29 | 27,61 | 25,34 | 27,18 | 27,36 | 25,61 | **0,979** | 9,75E-01 | **0** | -0,0300001 | 0,0109785 |
| ACTN4_HUM | ACTN4 | sp\|O43707\|ACTN4_HUMAN Alpha-actinin-4 OS=Homo sapiens GN=ACTN4 PE=1 SV=2 | 2763,969 | 61,25 | 104854 | 59 | 28,58 | 28,88 | 28,04 | 28,52 | 28,76 | 28,13 | **0,979** | 9,27E-01 | **0** | -0,0300001 | 0,03301505 |
| AMPB_HUM | RNPEP | sp\|Q9H4A4\|AMPB_HUMAN Aminopeptidase B OS=Homo sapiens GN=RNPEP PE=1 SV=2 | 107,22 | 7,23 | 72596 | 4 | 22,64 | 23,04 | 23,90 | 22,34 | 23,35 | 23,80 | **0,979** | 9,61E-01 | **0** | -0,0300001 | 0,01749084 |
| RAB1A_HUM | RAB1A | sp\|P62820\|RAB1A_HUMAN Ras-related protein Rab-1A OS=Homo sapiens GN=RAB1A PI | 641,55 | 78,05 | 22678 | 16 | 26,01 | 25,91 | 26,88 | 25,80 | 26,17 | 26,74 | **0,979** | 9,45E-01 | **0** | -0,0300001 | 0,02437111 |
| MRP_HUMA | MARCKSL1 | sp\|P49006\|MRP_HUMAN MARCKS-related protein OS=Homo sapiens GN=MARCKSL1 PE | 233,402 | 34,36 | 19529 | 5 | 26,49 | 26,52 | 23,32 | 26,48 | 26,17 | 23,59 | **0,979** | 9,84E-01 | **0** | -0,0300001 | 0,00702429 |
| STX18_HUM | STX18 | sp\|Q9P2W9\|STX18_HUMAN Syntaxin-18 OS=Homo sapiens GN=STX18 PE=1 SV=1 | 19,231 | 2,69 | 38674 | 1 | 19,04 | 17,24 | 18,43 | 17,58 | 18,16 | 18,87 | **0,977** | 9,62E-01 | **0** | -0,033069 | 0,01698936 |
| WDR1_HUM | WDR1 | sp\|O75083\|WDR1_HUMAN WD repeat-containing protein 1 OS=Homo sapiens GN=WDI | 1339,642 | 50,5 | 66194 | 24 | 30,04 | 29,31 | 28,81 | 29,33 | 29,92 | 28,81 | **0,977** | 9,48E-01 | **0** | -0,0333335 | 0,02320826 |
| ARPC4_HUM | ARPC4 | sp\|P59998\|ARPC4_HUMAN Actin-related protein 2/3 complex subunit 4 OS=Homo sapie | 298,907 | 71,43 | 19667 | 9 | 27,00 | 27,16 | 26,35 | 26,81 | 27,16 | 26,44 | **0,977** | 9,23E-01 | **0** | -0,0333335 | 0,03486659 |
| NIBL1_HUM | FAM129B | sp\|Q96TA1\|NIBL1_HUMAN Niban-like protein 1 OS=Homo sapiens GN=FAM129B PE=1 S | 611,781 | 19,44 | 84138 | 14 | 26,51 | 26,23 | 26,38 | 26,14 | 26,56 | 26,32 | **0,977** | 8,31E-01 | **0** | -0,0333335 | 0,08054976 |
| NDRG3_HUN | NDRG3 | sp\|Q9UGV2\|NDRG3_HUMAN Protein NDRG3 OS=Homo sapiens GN=NDRG3 PE=1 SV=2 | 143,559 | 13,07 | 41409 | 3 | 23,25 | 22,83 | 21,79 | 22,85 | 22,46 | 22,46 | **0,977** | 9,45E-01 | **0** | -0,0333341 | 0,02462632 |
| S23A2_HUM | SLC23A2 | sp\|Q9UGH3\|S23A2_HUMAN Solute carrier family 23 member 2 OS=Homo sapiens GN=S | 109,953 | 9,23 | 70337 | 4 | 23,51 | 22,69 | 25,83 | 23,50 | 23,13 | 25,30 | **0,977** | 9,78E-01 | **0** | -0,0333341 | 0,00950455 |
| LANC1_HUM | LANCL1 | sp\|O43813\|LANC1_HUMAN LanC-like protein 1 OS=Homo sapiens GN=LANCL1 PE=1 SV= | 24,978 | 2,76 | 45283 | 1 | 21,30 | 21,18 | 21,60 | 21,44 | 21,18 | 21,35 | **0,975** | 8,14E-01 | **0** | -0,0366662 | 0,08911765 |
| HEM3_HUM | HMBS | sp\|P08397\|HEM3_HUMAN Porphobilinogen deaminase OS=Homo sapiens GN=HMBS PE | 10,991 | 3,05 | 39330 | 1 | 21,18 | 21,59 | 20,13 | 21,38 | 21,12 | 20,29 | **0,975** | 9,50E-01 | **0** | -0,0366662 | 0,02246339 |
| INO1_HUMA | ISYNA1 | sp\|Q9NPH2\|INO1_HUMAN Inositol-3-phosphate synthase 1 OS=Homo sapiens GN=ISYN | 83,096 | 5,56 | 61068 | 4 | 23,77 | 24,02 | 23,66 | 23,71 | 23,97 | 23,66 | **0,975** | 8,11E-01 | **0** | -0,0366675 | 0,09105614 |
| TSN4_HUMA | TSPAN4 | sp\|O14817\|TSN4_HUMAN Tetraspanin-4 OS=Homo sapiens GN=TSPAN4 PE=1 SV=1 | 96,59 | 13,03 | 26118 | 2 | 28,15 | 27,96 | 28,15 | 28,08 | 28,28 | 27,78 | **0,973** | 8,13E-01 | **0** | -0,039999 | 0,08980257 |
| KIF2A_HUM | KIF2A | sp\|O00139\|KIF2A_HUMAN Kinesin-like protein KIF2A OS=Homo sapiens GN=KIF2A PE=1 | 40,673 | 2,69 | 79955 | 2 | 21,23 | 21,25 | 22,82 | 21,25 | 21,49 | 22,44 | **0,973** | 9,53E-01 | **0** | -0,0399996 | 0,02083721 |
| LTOR4_HUM | LAMTOR4 | sp\|Q0VGL1\|LTOR4_HUMAN Ragulator complex protein LAMTOR4 OS=Homo sapiens GN | 76,14 | 53,54 | 10741 | 2 | 22,56 | 21,37 | 20,64 | 22,11 | 21,04 | 21,30 | **0,973** | 9,54E-01 | **0** | -0,0399996 | 0,02064555 |
| GLCNE_HUM | GNE | sp\|Q9Y223\|GLCNE_HUMAN Bifunctional UDP-N-acetylglucosamine 2-epimerase/N-acet | 150,489 | 11,08 | 79275 | 7 | 22,76 | 23,69 | 24,51 | 22,44 | 23,33 | 25,07 | **0,973** | 9,68E-01 | **0** | -0,0400003 | 0,01434384 |
| GOLP3_HUM | GOLPH3 | sp\|Q9H4A6\|GOLP3_HUMAN Golgi phosphoprotein 3 OS=Homo sapiens GN=GOLPH3 PE | 181,147 | 16,44 | 33811 | 4 | 21,75 | 21,69 | 21,42 | 21,31 | 21,87 | 21,56 | **0,973** | 8,44E-01 | **0** | -0,0400003 | 0,07342117 |
| RS26_HUMA | RPS26 | sp\|P62854\|RS26_HUMAN 40S ribosomal protein S26 OS=Homo sapiens GN=RPS26 PE=1 | 131,137 | 31,3 | 13015 | 3 | 26,07 | 26,14 | 26,20 | 26,02 | 25,88 | 26,39 | **0,973** | 8,11E-01 | **0** | -0,0400003 | 0,0909214 |
| CSK22_HUM | CSNK2A2 | sp\|P19784\|CSK22_HUMAN Casein kinase II subunit alpha' OS=Homo sapiens GN=CSNK2 | 184,061 | 23,43 | 41213 | 8 | 22,75 | 23,24 | 25,23 | 22,79 | 23,28 | 25,02 | **0,970** | 9,68E-01 | **0** | -0,0433324 | 0,01410952 |
| PP1A_HUMA | PPP1CA | sp\|P62136\|PP1A_HUMAN Serine/threonine-protein phosphatase PP1-alpha catalytic sul | 851,993 | 60,91 | 37512 | 21 | 25,92 | 25,68 | 25,04 | 25,68 | 25,54 | 25,29 | **0,970** | 8,87E-01 | **0** | -0,0433331 | 0,05205859 |
| KAP2_HUMA | PRKAR2A | sp\|P13861\|KAP2_HUMAN cAMP-dependent protein kinase type II-alpha regulatory subu | 299,386 | 23,51 | 45518 | 8 | 24,48 | 24,36 | 23,88 | 24,38 | 24,45 | 23,76 | **0,970** | 8,87E-01 | **0** | -0,0433331 | 0,05216276 |
| APMAP_HUN | APMAP | sp\|Q9HDC9\|APMAP_HUMAN Adipocyte plasma membrane-associated protein OS=Hom | 171,607 | 11,06 | 46480 | 5 | 25,74 | 25,26 | 25,71 | 25,67 | 25,55 | 25,36 | **0,970** | 8,21E-01 | **0** | -0,0433331 | 0,0855653 |
| TX1B3_HUM | TAX1BP3 | sp\|O14907\|TX1B3_HUMAN Tax1-binding protein 3 OS=Homo sapiens GN=TAX1BP3 PE= | 156,559 | 28,23 | 13735 | 2 | 23,58 | 23,67 | 23,38 | 23,55 | 23,76 | 23,19 | **0,970** | 8,28E-01 | **0** | -0,0433331 | 0,08181097 |
| KIF3B_HUM | KIF3B | sp\|O15066\|KIF3B_HUMAN Kinesin-like protein KIF3B OS=Homo sapiens GN=KIF3B PE=1 | 204,511 | 14,73 | 85125 | 9 | 22,48 | 22,01 | 25,18 | 22,22 | 23,33 | 23,99 | **0,970** | 9,71E-01 | **0** | -0,0433337 | 0,01284811 |
| FA49B_HUM | FAM49B | sp\|Q9NUQ9\|FA49B_HUMAN Protein FAM49B OS=Homo sapiens GN=FAM49B PE=1 SV= | 340,508 | 29,32 | 36748 | 7 | 24,97 | 24,78 | 23,47 | 24,72 | 24,71 | 23,66 | **0,970** | 9,45E-01 | **0** | -0,0433337 | 0,02465458 |
| DNJC5_HUM | DNAJC5 | sp\|Q9H3Z4\|DNJC5_HUMAN DnaJ homolog subfamily C member 5 OS=Homo sapiens GN | 260,12 | 24,75 | 22149 | 6 | 25,31 | 25,16 | 24,95 | 25,25 | 25,33 | 24,74 | **0,970** | 8,54E-01 | **0** | -0,0433337 | 0,06851049 |
| KAPCA_HUM | PRKACA | sp\|P17612\|KAPCA_HUMAN cAMP-dependent protein kinase catalytic subunit alpha OS= | 237,082 | 33,9 | 40590 | 13 | 22,52 | 23,00 | 23,61 | 22,38 | 23,33 | 23,29 | **0,970** | 9,27E-01 | **0** | -0,0433337 | 0,03306939 |
| CLCA_HUMA | CLTA | sp\|P09496\|CLCA_HUMAN Clathrin light chain A OS=Homo sapiens GN=CLTA PE=1 SV=1 | 153,227 | 17,74 | 27077 | 7 | 27,16 | 26,30 | 26,72 | 27,33 | 26,69 | 26,02 | **0,968** | 9,23E-01 | **0** | -0,0466658 | 0,03488795 |
| SC23B_HUM | SEC23B | sp\|Q15437\|SC23B_HUMAN Protein transport protein Sec23B OS=Homo sapiens GN=SE( | 360,014 | 18,77 | 86479 | 10 | 23,47 | 23,60 | 24,57 | 23,43 | 23,62 | 24,45 | **0,968** | 9,25E-01 | **0** | -0,0466658 | 0,03372572 |
| PDLI1_HUM | PDLIM1 | sp\|O00151\|PDLI1_HUMAN PDZ and LIM domain protein 1 OS=Homo sapiens GN=PDLIM | 114,525 | 13,07 | 36072 | 3 | 23,96 | 24,31 | 22,60 | 24,27 | 24,35 | 22,11 | **0,968** | 9,61E-01 | **0** | -0,0466658 | 0,01721277 |
| CA2D1_HUM | CACNA2D1 | sp\|P54289\|CA2D1_HUMAN Voltage-dependent calcium channel subunit alpha-2/delta-: | 303,224 | 14,96 | 124568 | 13 | 25,22 | 24,97 | 24,99 | 24,86 | 24,84 | 25,34 | **0,968** | 8,10E-01 | **0** | -0,0466658 | 0,09133172 |
| DNJC7_HUM | DNAJC7 | sp\|Q99615\|DNJC7_HUMAN DnaJ homolog subfamily C member 7 OS=Homo sapiens GN | 708,053 | 36,64 | 56441 | 22 | 26,61 | 26,32 | 26,47 | 26,59 | 26,11 | 26,56 | **0,968** | 8,04E-01 | **0** | -0,0466665 | 0,09452202 |
| U2AF1_HUM | U2AF1 | sp\|Q01081\|U2AF1_HUMAN Splicing factor U2AF 35 kDa subunit OS=Homo sapiens GN= | 320,124 | 45,83 | 27872 | 9 | 24,38 | 25,23 | 25,68 | 24,58 | 24,56 | 26,01 | **0,968** | 9,43E-01 | **0** | -0,0466665 | 0,02550282 |
| UFD1_HUM | UFD1 | sp\|Q92890\|UFD1_HUMAN Ubiquitin recognition factor in ER-associated degradation pr | 137,068 | 20,2 | 34500 | 6 | 23,90 | 24,32 | 22,34 | 23,83 | 24,09 | 22,50 | **0,968** | 9,55E-01 | **0** | -0,0466665 | 0,01997548 |
| SAR1A_HUM | SAR1A | sp\|Q9NR31\|SAR1A_HUMAN GTP-binding protein SAR1a OS=Homo sapiens GN=SAR1A P | 570,86 | 75,25 | 22367 | 16 | 25,49 | 25,89 | 26,44 | 25,29 | 25,99 | 26,40 | **0,968** | 9,18E-01 | **0** | -0,0466665 | 0,03720312 |
| PLPP1_HUM | PLPP1 | sp\|O14494\|PLPP1_HUMAN Phospholipid phosphatase 1 OS=Homo sapiens GN=PLPP1 PI | 94,95 | 10,21 | 32156 | 2 | 24,40 | 23,58 | 23,67 | 23,87 | 24,24 | 23,40 | **0,968** | 9,02E-01 | **0** | -0,0466665 | 0,04482176 |
| RAB8A_HUM | RAB8A | sp\|P61006\|RAB8A_HUMAN Ras-related protein Rab-8A OS=Homo sapiens GN=RAB8A PI | 416,781 | 55,07 | 23668 | 11 | 26,34 | 26,36 | 25,97 | 26,56 | 26,18 | 25,79 | **0,968** | 8,64E-01 | **0** | -0,0466665 | 0,06339986 |
| RIPR1_HUM | RIPOR1 | sp\|Q6ZS17\|RIPR1_HUMAN Rho family-interacting cell polarization regulator 1 OS=Homo | 143,797 | 4,17 | 132308 | 4 | 21,72 | 22,27 | 21,04 | 21,29 | 22,00 | 21,60 | **0,968** | 9,15E-01 | **0** | -0,0466665 | 0,03856263 |
| M4K5_HUM | MAP4K5 | sp\|Q9Y4K4\|M4K5_HUMAN Mitogen-activated protein kinase kinase kinase kinase 5 OS= | 86,806 | 5,79 | 95040 | 4 | 22,18 | 21,34 | 23,05 | 22,22 | 21,92 | 22,29 | **0,968** | 9,31E-01 | **0** | -0,0466665 | 0,03103852 |
| EVA1A_HUM | EVA1A | sp\|Q9H8M9\|EVA1A_HUMAN Protein eva-1 homolog A OS=Homo sapiens GN=EVA1A PE | 33,606 | 7,24 | 17470 | 1 | 23,17 | 23,33 | 22,44 | 23,50 | 23,24 | 22,06 | **0,968** | 9,33E-01 | **0** | -0,0466671 | 0,03015511 |
| SLK_HUMAN | SLK | sp\|Q9H2G2\|SLK_HUMAN STE20-like serine/threonine-protein kinase OS=Homo sapiens | 203,995 | 9,23 | 142695 | 9 | 23,55 | 23,48 | 22,36 | 23,22 | 23,40 | 22,63 | **0,968** | 9,22E-01 | **0** | -0,0466671 | 0,03506401 |
| PLEK_HUMA | PLEK | sp\|P08567\|PLEK_HUMAN Pleckstrin OS=Homo sapiens GN=PLEK PE=1 SV=3 | 115,35 | 11,43 | 40125 | 3 | 23,27 | 23,09 | 22,92 | 23,84 | 23,00 | 22,30 | **0,968** | 9,23E-01 | **0** | -0,0466671 | 0,03456551 |
| PCID2_HUM | PCID2 | sp\|Q5JVF3\|PCID2_HUMAN PCI domain-containing protein 2 OS=Homo sapiens GN=PCID | 63,619 | 7,77 | 46030 | 3 | 20,54 | 20,47 | 21,73 | 19,89 | 20,48 | 22,23 | **0,968** | 9,57E-01 | **0** | -0,0466671 | 0,0190999 |
| RAP1A_HUM | RAP1A | sp\|P62834\|RAP1A_HUMAN Ras-related protein Rap-1A OS=Homo sapiens GN=RAP1A PI | 1051,188 | 83,15 | 20987 | 19 | 26,35 | 26,30 | 24,93 | 26,35 | 26,41 | 24,67 | **0,966** | 9,49E-01 | **0** | -0,0499999 | 0,0226903 |
| ARPC3_HUM | ARPC3 | sp\|O15145\|ARPC3_HUMAN Actin-related protein 2/3 complex subunit 3 OS=Homo sapi | 216,203 | 48,31 | 20547 | 7 | 26,27 | 26,25 | 25,25 | 26,12 | 26,10 | 25,40 | **0,966** | 9,09E-01 | **0** | -0,0499999 | 0,04135325 |
| HIP1_HUMA | HIP1 | sp\|O00291\|HIP1_HUMAN Huntingtin-interacting protein 1 OS=Homo sapiens GN=HIP1 F | 134,64 | 7,43 | 116221 | 5 | 21,99 | 22,26 | 22,96 | 22,28 | 21,88 | 22,90 | **0,966** | 9,10E-01 | **0** | -0,0499999 | 0,04108025 |
| PDIA4_HUM | PDIA4 | sp\|P13667\|PDIA4_HUMAN Protein disulfide-isomerase A4 OS=Homo sapiens GN=PDIA4 | 422,044 | 17,98 | 72932 | 14 | 25,76 | 26,10 | 27,08 | 26,29 | 26,08 | 26,42 | **0,966** | 9,08E-01 | **0** | -0,0499999 | 0,0417447 |
| BZW2_HUM | BZW2 | sp\|Q9Y6E2\|BZW2_HUMAN Basic leucine zipper and W2 domain-containing protein 2 OS | 193,53 | 15,75 | 48162 | 5 | 23,16 | 23,42 | 23,48 | 22,92 | 23,48 | 23,51 | **0,966** | 8,28E-01 | **0** | -0,0499999 | 0,08200188 |

| | | | | | | | | | | | | | | | | |
|---|---|---|---|---|---|---|---|---|---|---|---|---|---|---|---|---|
| UBP4_HUMA | USP4 | sp\|Q13107\|UBP4_HUMAN Ubiquitin carboxyl-terminal hydrolase 4 OS=Homo sapiens G | 81,9 | 4,78 | 108565 | 3 | 20,62 | 19,83 | 22,35 | 19,48 | 20,81 | 22,36 | 0,966 | 9,66E-01 | 0 | -0,0500005 | 0,01483284 |
| RU1C_HUMA | SNRPC | sp\|P09234\|RU1C_HUMAN U1 small nuclear ribonucleoprotein C OS=Homo sapiens GN= | 102,629 | 25,79 | 17394 | 3 | 22,30 | 22,61 | 21,28 | 22,41 | 22,49 | 21,14 | 0,966 | 9,37E-01 | 0 | -0,0500005 | 0,02828729 |
| ECE1_HUMA | ECE1 | sp\|P42892\|ECE1_HUMAN Endothelin-converting enzyme 1 OS=Homo sapiens GN=ECE1 | 57,975 | 4,16 | 87164 | 2 | 21,95 | 22,27 | 20,33 | 22,70 | 21,65 | 20,05 | 0,966 | 9,62E-01 | 0 | -0,0500005 | 0,01699158 |
| MIRO1_HUN | RHOT1 | sp\|Q8IXI2\|MIRO1_HUMAN Mitochondrial Rho GTPase 1 OS=Homo sapiens GN=RHOT1 | 105,497 | 6,15 | 70784 | 3 | 20,99 | 19,46 | 21,75 | 20,34 | 20,31 | 21,40 | 0,966 | 9,51E-01 | 0 | -0,0502078 | 0,02196933 |
| GPAT4_HUN | GPAT4 | sp\|Q86UL3\|GPAT4_HUMAN Glycerol-3-phosphate acyltransferase 4 OS=Homo sapiens | 29,693 | 2,63 | 52071 | 1 | 18,71 | 19,44 | 20,58 | 19,10 | 18,81 | 20,66 | 0,966 | 9,52E-01 | 0 | -0,0502415 | 0,02117449 |
| HGNAT_HUN | HGSNAT | sp\|Q68CP4\|HGNAT_HUMAN Heparan-alpha-glucosaminide N-acetyltransferase OS=Hor | 19,124 | 1,66 | 73293 | 1 | 22,10 | 22,20 | 17,86 | 21,73 | 21,27 | 19,01 | 0,965 | 9,77E-01 | 0 | -0,0515366 | 0,01024122 |
| AATC_HUMA | GOT1 | sp\|P17174\|AATC_HUMAN Aspartate aminotransferase, cytoplasmic OS=Homo sapiens | 88,038 | 12,11 | 46248 | 4 | 23,92 | 23,88 | 22,40 | 23,85 | 24,20 | 21,99 | 0,964 | 9,53E-01 | 0 | -0,0533326 | 0,02094606 |
| ARL1_HUMA | ARL1 | sp\|P40616\|ARL1_HUMAN ADP-ribosylation factor-like protein 1 OS=Homo sapiens GN= | 144,383 | 21,55 | 20418 | 3 | 23,33 | 23,20 | 23,88 | 23,04 | 23,45 | 23,76 | 0,964 | 8,65E-01 | 0 | -0,0533326 | 0,06286505 |
| BABA2_HUM | BABAM2 | sp\|Q9NXR7\|BABA2_HUMAN BRISC and BRCA1-A complex member 2 OS=Homo sapiens | 19,915 | 3,66 | 43552 | 1 | 19,44 | 19,32 | 20,34 | 19,45 | 19,31 | 20,18 | 0,964 | 9,05E-01 | 0 | -0,0533333 | 0,04331987 |
| IF4A3_HUM | EIF4A3 | sp\|P38919\|IF4A3_HUMAN Eukaryotic initiation factor 4A-III OS=Homo sapiens GN=EIF4 | 1178,976 | 68,37 | 46871 | 29 | 28,04 | 27,67 | 28,46 | 27,75 | 27,74 | 28,52 | 0,964 | 8,85E-01 | 0 | -0,0533333 | 0,05328533 |
| STRBP_HUM | STRBP | sp\|Q96SI9\|STRBP_HUMAN Spermatid perinuclear RNA-binding protein OS=Homo sapier | 196,961 | 6,7 | 73653 | 6 | 22,98 | 22,77 | 23,25 | 22,40 | 23,44 | 23,00 | 0,964 | 8,80E-01 | 0 | -0,0533333 | 0,05545869 |
| CHMP3_HUN | CHMP3 | sp\|Q9Y3E7\|CHMP3_HUMAN Charged multivesicular body protein 3 OS=Homo sapiens G | 233,026 | 27,93 | 25073 | 9 | 27,74 | 26,64 | 25,75 | 27,69 | 27,06 | 25,22 | 0,964 | 9,57E-01 | 0 | -0,0533333 | 0,01890834 |
| KC1G3_HUM | CSNK1G3 | sp\|Q9Y6M4\|KC1G3_HUMAN Casein kinase I isoform gamma-3 OS=Homo sapiens GN=CS | 278,623 | 31,54 | 51389 | 12 | 23,69 | 23,35 | 21,25 | 23,88 | 23,56 | 20,69 | 0,964 | 9,68E-01 | 0 | -0,0533339 | 0,01390272 |
| KPCD1_HUM | PRKD1 | sp\|Q15139\|KPCD1_HUMAN Serine/threonine-protein kinase D1 OS=Homo sapiens GN= | 49,817 | 4,39 | 101704 | 3 | 18,63 | 19,69 | 21,13 | 18,15 | 20,03 | 21,11 | 0,963 | 9,64E-01 | 0 | -0,0544281 | 0,01598649 |
| FBLN1_HUM | FBLN1 | sp\|P23142\|FBLN1_HUMAN Fibulin-1 OS=Homo sapiens GN=FBLN1 PE=1 SV=4 | 293,219 | 13,23 | 77214 | 9 | 25,22 | 23,64 | 25,98 | 25,60 | 23,76 | 25,31 | 0,961 | 9,53E-01 | 0 | -0,0566661 | 0,02110784 |
| SEPT8_HUM | SEPT8 | sp\|Q92599\|SEPT8_HUMAN Septin-8 OS=Homo sapiens GN=SEPT8 PE=1 SV=4 | 202,344 | 23,19 | 55756 | 8 | 23,32 | 23,30 | 23,12 | 23,27 | 23,51 | 22,79 | 0,961 | 8,10E-01 | 0 | -0,0566661 | 0,09135956 |
| CDK4_HUMA | CDK4 | sp\|P11802\|CDK4_HUMAN Cyclin-dependent kinase 4 OS=Homo sapiens GN=CDK4 PE=1 | 205,74 | 33 | 33730 | 10 | 25,24 | 25,23 | 25,38 | 25,04 | 25,16 | 25,48 | 0,961 | 7,06E-01 | 0 | -0,0566661 | 0,15101851 |
| CAZA2_HUM | CAPZA2 | sp\|P47755\|CAZA2_HUMAN F-actin-capping protein subunit alpha-2 OS=Homo sapiens G | 308,516 | 40,21 | 32949 | 8 | 25,00 | 24,80 | 24,29 | 24,85 | 24,71 | 24,36 | 0,961 | 8,36E-01 | 0 | -0,0566667 | 0,07773339 |
| LMAN2_HUN | LMAN2 | sp\|Q12907\|LMAN2_HUMAN Vesicular integral-membrane protein VIP36 OS=Homo sapi | 200,466 | 31,46 | 40229 | 8 | 24,94 | 24,77 | 24,85 | 25,16 | 24,94 | 24,29 | 0,961 | 8,42E-01 | 0 | -0,0566667 | 0,07490917 |
| PIGU_HUMA | PIGU | sp\|Q9H490\|PIGU_HUMAN Phosphatidylinositol glycan anchor biosynthesis class U prote | 56,881 | 3,91 | 50052 | 1 | 19,07 | 19,17 | 20,51 | 19,30 | 19,06 | 20,22 | 0,961 | 9,27E-01 | 0 | -0,0566673 | 0,03277629 |
| RGS19_HUM | RGS19 | sp\|P49795\|RGS19_HUMAN Regulator of G-protein signaling 19 OS=Homo sapiens GN=R | 57,097 | 5,99 | 24636 | 1 | 20,32 | 20,44 | 22,18 | 21,06 | 20,89 | 20,82 | 0,961 | 9,30E-01 | 0 | -0,0566673 | 0,03156213 |
| TLN1_HUMA | TLN1 | sp\|Q9Y490\|TLN1_HUMAN Talin-1 OS=Homo sapiens GN=TLN1 PE=1 SV=3 | 4734,389 | 51,52 | 269767 | 99 | 30,22 | 29,87 | 30,20 | 29,97 | 29,94 | 30,21 | 0,961 | 7,10E-01 | 0 | -0,0566673 | 0,14854315 |
| MACF1_HUN | MACF1 | sp\|Q9UPN3\|MACF1_HUMAN Microtubule-actin cross-linking factor 1, isoforms 1/2/3/5 | 256,231 | 1,73 | 838308 | 11 | 24,68 | 24,60 | 23,99 | 24,56 | 24,56 | 23,98 | 0,961 | 8,55E-01 | 0 | -0,0566673 | 0,06791225 |
| CENPV_HUM | CENPV | sp\|Q7Z7K6\|CENPV_HUMAN Centromere protein V OS=Homo sapiens GN=CENPV PE=1 S | 42,677 | 11,64 | 29946 | 2 | 18,74 | 19,99 | 23,45 | 17,76 | 21,31 | 22,94 | 0,960 | 9,79E-01 | 0 | -0,0581512 | 0,00920726 |
| HSP7C_HUM | HSPA8 | sp\|P11142\|HSP7C_HUMAN Heat shock cognate 71 kDa protein OS=Homo sapiens GN=H | 3783,389 | 87,62 | 70898 | 73 | 32,78 | 32,46 | 32,21 | 32,83 | 32,36 | 32,08 | 0,959 | 8,37E-01 | 0 | -0,0599976 | 0,07706348 |
| M3K3_HUM | MAP3K3 | sp\|Q99759\|M3K3_HUMAN Mitogen-activated protein kinase kinase kinase 3 OS=Homo | 10,282 | 1,12 | 70898 | 1 | 22,15 | 22,29 | 22,05 | 22,12 | 22,10 | 22,09 | 0,959 | 4,41E-01 | 0 | -0,0599995 | 0,35591162 |
| SAE2_HUMA | UBA2 | sp\|Q9UBT2\|SAE2_HUMAN SUMO-activating enzyme subunit 2 OS=Homo sapiens GN=U | 183,246 | 13,12 | 71224 | 8 | 24,50 | 23,80 | 24,30 | 24,56 | 23,79 | 24,07 | 0,959 | 8,54E-01 | 0 | -0,0599995 | 0,06835839 |
| SEP11_HUM | SEPT11 | sp\|Q9NVA2\|SEP11_HUMAN Septin-11 OS=Homo sapiens GN=SEPT11 PE=1 SV=3 | 484,676 | 40,79 | 49398 | 12 | 25,54 | 25,75 | 25,41 | 25,44 | 25,61 | 25,47 | 0,959 | 6,21E-01 | 0 | -0,0600001 | 0,20711019 |
| GRP78_HUM | HSPA5 | sp\|P11021\|GRP78_HUMAN 78 kDa glucose-regulated protein OS=Homo sapiens GN=HS | 2261,014 | 64,83 | 72333 | 46 | 29,20 | 28,71 | 30,12 | 29,38 | 28,95 | 29,52 | 0,959 | 9,00E-01 | 0 | -0,0600001 | 0,04585032 |
| TNR6_HUMA | FAS | sp\|P25445\|TNR6_HUMAN Tumor necrosis factor receptor superfamily member 6 OS=H | 32,583 | 2,69 | 37732 | 2 | 14,48 | 17,83 | 21,10 | 15,67 | 17,46 | 20,10 | 0,959 | 9,80E-01 | 0 | -0,0600001 | 0,00856563 |
| RAB18_HUM | RAB18 | sp\|Q9NP72\|RAB18_HUMAN Ras-related protein Rab-18 OS=Homo sapiens GN=RAB18 P | 424,743 | 60,68 | 22977 | 11 | 26,94 | 27,07 | 26,73 | 26,85 | 27,07 | 26,64 | 0,959 | 7,25E-01 | 0 | -0,0600001 | 0,13979811 |
| RHOA_HUM | RHOA | sp\|P61586\|RHOA_HUMAN Transforming protein RhoA OS=Homo sapiens GN=RHOA PE= | 855,124 | 80,31 | 21768 | 18 | 27,41 | 27,33 | 26,91 | 27,54 | 27,22 | 26,71 | 0,959 | 8,45E-01 | 0 | -0,0600001 | 0,07329898 |
| KINH_HUMA | KIF5B | sp\|P33176\|KINH_HUMAN Kinesin-1 heavy chain OS=Homo sapiens GN=KIF5B PE=1 SV= | 1640,37 | 52,34 | 109685 | 41 | 26,86 | 27,12 | 27,90 | 26,96 | 27,03 | 27,71 | 0,959 | 8,86E-01 | 0 | -0,0600007 | 0,05247043 |
| SF3A1_HUM | SF3A1 | sp\|Q15459\|SF3A1_HUMAN Splicing factor 3A subunit 1 OS=Homo sapiens GN=SF3A1 PE | 131,29 | 9,71 | 88886 | 6 | 22,37 | 22,10 | 24,29 | 22,44 | 22,00 | 24,14 | 0,959 | 9,53E-01 | 0 | -0,0600007 | 0,02107489 |
| LOXL4_HUM | LOXL4 | sp\|Q96JB6\|LOXL4_HUMAN Lysyl oxidase homolog 4 OS=Homo sapiens GN=LOXL4 PE=1 | 15,71 | 1,06 | 84483 | 1 | 20,24 | 18,14 | 20,82 | 18,22 | 19,73 | 21,06 | 0,959 | 9,61E-01 | 0 | -0,0609652 | 0,01749698 |
| LRC47_HUM | LRRC47 | sp\|Q8N1G4\|LRC47_HUMAN Leucine-rich repeat-containing protein 47 OS=Homo sapier | 34,981 | 1,54 | 63473 | 1 | 19,91 | 19,72 | 20,93 | 20,54 | 18,77 | 21,07 | 0,958 | 9,42E-01 | 0 | -0,0614414 | 0,02606939 |
| GAN_HUMA | GAN | sp\|Q9H2C0\|GAN_HUMAN Gigaxonin OS=Homo sapiens GN=GAN PE=1 SV=1 | 31,962 | 2,35 | 67638 | 1 | 19,74 | 19,14 | 19,52 | 19,12 | 19,06 | 20,03 | 0,958 | 8,70E-01 | 0 | -0,0625852 | 0,06041616 |
| CP062_HUM | C16orf62 | sp\|Q7Z3J2\|CP062_HUMAN UPF0505 protein C16orf62 OS=Homo sapiens GN=C16orf62 | 138,417 | 5,4 | 109563 | 4 | 20,30 | 20,12 | 22,24 | 19,52 | 20,65 | 22,30 | 0,957 | 9,55E-01 | 0 | -0,0633335 | 0,0200004 |
| GYS1_HUMA | GYS1 | sp\|P13807\|GYS1_HUMAN Glycogen [starch] synthase, muscle OS=Homo sapiens GN=G | 40,126 | 2,85 | 83786 | 2 | 21,73 | 22,37 | 21,90 | 21,56 | 22,30 | 21,95 | 0,957 | 8,36E-01 | 0 | -0,0633335 | 0,07774391 |
| GYS2_HUMA | GYS2 | sp\|P54840\|GYS2_HUMAN Glycogen [starch] synthase, liver OS=Homo sapiens GN=GYS2 | 40,126 | 2,99 | 80989 | 2 | 21,73 | 22,37 | 21,90 | 21,56 | 22,30 | 21,95 | 0,957 | 8,36E-01 | 0 | -0,0633335 | 0,07774391 |
| 1433Z_HUM | YWHAZ | sp\|P63104\|1433Z_HUMAN 14-3-3 protein zeta/delta OS=Homo sapiens GN=YWHAZ PE= | 1423,788 | 69,39 | 27745 | 28 | 29,87 | 29,83 | 28,91 | 29,73 | 29,71 | 28,98 | 0,957 | 8,82E-01 | 0 | -0,0633341 | 0,05474456 |
| SL9A1_HUM | SLC9A1 | sp\|P19634\|SL9A1_HUMAN Sodium/hydrogen exchanger 1 OS=Homo sapiens GN=SLC9A | 18,172 | 1,72 | 90763 | 1 | 21,58 | 21,84 | 18,48 | 21,70 | 21,45 | 18,56 | 0,957 | 9,68E-01 | 0 | -0,0633469 | 0,0141957 |
| KIF4B_HUM | KIF4B | sp\|Q2VIQ3\|KIF4B_HUMAN Chromosome-associated kinesin KIF4B OS=Homo sapiens GN | 379,78 | 11,67 | 140035 | 13 | 18,56 | 19,59 | 22,15 | 19,97 | 18,59 | 21,55 | 0,957 | 9,65E-01 | 0 | -0,0634867 | 0,0153816 |
| RAD50_HUM | RAD50 | sp\|Q92878\|RAD50_HUMAN DNA repair protein RAD50 OS=Homo sapiens GN=RAD50 PE | 47,922 | 1,83 | 153892 | 2 | 19,24 | 18,47 | 21,40 | 19,49 | 18,07 | 21,37 | 0,956 | 9,63E-01 | 0 | -0,0642808 | 0,01643775 |
| BCS1_HUMA | BCS1L | sp\|Q9Y276\|BCS1_HUMAN Mitochondrial chaperone BCS1 OS=Homo sapiens GN=BCS1L | 25,979 | 4,53 | 47534 | 1 | 16,33 | 15,61 | 19,13 | 17,41 | 16,43 | 17,03 | 0,955 | 9,56E-01 | 0 | -0,0658077 | 0,01971902 |
| UBE3A_HUM | UBE3A | sp\|Q05086\|UBE3A_HUMAN Ubiquitin-protein ligase E3A OS=Homo sapiens GN=UBE3A | 78,404 | 5,03 | 100688 | 4 | 21,11 | 22,63 | 23,35 | 21,67 | 21,95 | 23,27 | 0,955 | 9,39E-01 | 0 | -0,0666663 | 0,02714147 |
| RAB35_HUM | RAB35 | sp\|Q15286\|RAB35_HUMAN Ras-related protein Rab-35 OS=Homo sapiens GN=RAB35 P | 787,09 | 74,13 | 23025 | 16 | 28,69 | 28,19 | 28,32 | 28,45 | 28,48 | 28,07 | 0,955 | 7,55E-01 | 0 | -0,0666669 | 0,12195688 |
| ATP7A_HUM | ATP7A | sp\|Q04656\|ATP7A_HUMAN Copper-transporting ATPase 1 OS=Homo sapiens GN=ATP7 | 57,134 | 1,13 | 163374 | 1 | 18,63 | 19,70 | 19,19 | 18,61 | 19,71 | 19,00 | 0,955 | 8,88E-01 | 0 | -0,0666669 | 0,05135368 |
| ACTZ_HUMA | ACTR1A | sp\|P61163\|ACTZ_HUMAN Alpha-centractin OS=Homo sapiens GN=ACTR1A PE=1 SV=1 | 654,953 | 65,16 | 42614 | 17 | 25,34 | 25,51 | 26,09 | 24,63 | 25,66 | 26,45 | 0,955 | 9,13E-01 | 0 | -0,0666669 | 0,0394833 |
| FLOT2_HUM | FLOT2 | sp\|Q14254\|FLOT2_HUMAN Flotillin-2 OS=Homo sapiens GN=FLOT2 PE=1 SV=2 | 467,281 | 42,06 | 47064 | 14 | 26,46 | 26,24 | 26,54 | 26,56 | 26,31 | 26,17 | 0,955 | 6,70E-01 | 0 | -0,0666669 | 0,17406795 |
| LEG3_HUMA | LGALS3 | sp\|P17931\|LEG3_HUMAN Galectin-3 OS=Homo sapiens GN=LGALS3 PE=1 SV=5 | 133,36 | 14,4 | 26152 | 3 | 25,11 | 25,69 | 22,73 | 24,87 | 25,14 | 23,32 | 0,955 | 9,53E-01 | 0 | -0,0666669 | 0,02079643 |
| PYGL_HUMA | PYGL | sp\|P06737\|PYGL_HUMAN Glycogen phosphorylase, liver form OS=Homo sapiens GN=P | 297,145 | 9,56 | 97149 | 6 | 16,57 | 18,72 | 20,02 | 17,87 | 17,69 | 19,55 | 0,955 | 9,57E-01 | 0 | -0,0670439 | 0,01910739 |
| FIBG_HUMA | FGG | sp\|P02679\|FIBG_HUMAN Fibrinogen gamma chain OS=Homo sapiens GN=FGG PE=1 SV= | 15,791 | 3,97 | 51512 | 1 | 20,82 | 18,86 | 20,88 | 21,44 | 18,90 | 20,02 | 0,954 | 9,48E-01 | 0 | -0,0681483 | 0,02300884 |
| CYGB_HUMA | CYGB | sp\|Q8WWM9\|CYGB_HUMAN Cytoglobin OS=Homo sapiens GN=CYGB PE=1 SV=1 | 10,972 | 3,68 | 21405 | 1 | 18,29 | 19,94 | 25,47 | 18,63 | 20,43 | 24,44 | 0,954 | 9,81E-01 | 0 | -0,0683301 | 0,00811307 |
| LYPA2_HUM | LYPLA2 | sp\|O95372\|LYPA2_HUMAN Acyl-protein thioesterase 2 OS=Homo sapiens GN=LYPLA2 P | 65,897 | 25,54 | 24737 | 3 | 22,64 | 22,71 | 23,21 | 22,41 | 22,59 | 23,35 | 0,953 | 8,47E-01 | 0 | -0,0699991 | 0,07228545 |
| GEMI5_HUM | GEMIN5 | sp\|Q8TEQ6\|GEMI5_HUMAN Gem-associated protein 5 OS=Homo sapiens GN=GEMIN5 | 91,619 | 2,72 | 168589 | 4 | 20,22 | 19,64 | 22,41 | 19,99 | 19,02 | 23,05 | 0,953 | 9,65E-01 | 0 | -0,0699997 | 0,01569357 |
| AADAT_HUN | AADAT | sp\|Q8N5Z0\|AADAT_HUMAN Kynurenine/alpha-aminoadipate aminotransferase, mitoch | 55,409 | 4,24 | 47352 | 1 | 21,06 | 21,09 | 21,56 | 21,26 | 21,27 | 20,97 | 0,953 | 7,30E-01 | 0 | -0,0699997 | 0,13638924 |
| PCBP1_HUM | PCBP1 | sp\|Q15365\|PCBP1_HUMAN Poly(rC)-binding protein 1 OS=Homo sapiens GN=PCBP1 PE= | 827,918 | 69,38 | 37498 | 18 | 27,93 | 27,63 | 27,84 | 27,85 | 27,57 | 27,77 | 0,953 | 5,96E-01 | 0 | -0,0699997 | 0,22457666 |
| SEPT7_HUM | SEPT7 | sp\|Q16181\|SEPT7_HUMAN Septin-7 OS=Homo sapiens GN=SEPT7 PE=1 SV=2 | 815,702 | 53,09 | 50680 | 20 | 27,17 | 27,22 | 26,82 | 27,05 | 27,16 | 26,79 | 0,953 | 6,97E-01 | 0 | -0,0699997 | 0,15706852 |
| MPRD_HUM | M6PR | sp\|P20645\|MPRD_HUMAN Cation-dependent mannose-6-phosphate receptor OS=Hom | 253,082 | 24,91 | 30993 | 5 | 25,26 | 25,57 | 25,44 | 25,35 | 25,58 | 25,13 | 0,953 | 6,81E-01 | 0 | -0,0700003 | 0,16711256 |
| UBAC2_HUN | UBAC2 | sp\|Q8NBM4\|UBAC2_HUMAN Ubiquitin-associated domain-containing protein 2 OS=Hor | 9,608 | 2,33 | 38964 | 1 | 25,42 | 23,18 | 24,50 | 25,64 | 23,57 | 23,68 | 0,953 | 9,44E-01 | 0 | -0,0700003 | 0,02505958 |
| FA10_HUMA | F10 | sp\|P00742\|FA10_HUMAN Coagulation factor X OS=Homo sapiens GN=F10 PE=1 SV=2 | 126,677 | 9,84 | 54732 | 4 | 27,09 | 25,60 | 25,11 | 27,12 | 25,72 | 24,75 | 0,953 | 9,42E-01 | 0 | -0,0700003 | 0,02578124 |
| UTS2_HUMA | UTS2 | sp\|O95399\|UTS2_HUMAN Urotensin-2 OS=Homo sapiens GN=UTS2 PE=1 SV=1 | 12,621 | 6,45 | 14296 | 1 | 25,68 | 25,27 | 26,20 | 25,94 | 25,39 | 25,61 | 0,953 | 8,34E-01 | 0 | -0,0700003 | 0,07884349 |
| XP32_HUMA | XP32 | sp\|Q5T750\|XP32_HUMAN Skin-specific protein 32 OS=Homo sapiens GN=XP32 PE=1 SV | 12,795 | 3,2 | 26238 | 1 | 24,23 | 22,26 | 18,47 | 24,29 | 23,38 | 17,08 | 0,952 | 9,81E-01 | 0 | -0,070254 | 0,00816458 |
| SLIK6_HUMA | SLITRK6 | sp\|Q9H5Y7\|SLIK6_HUMAN SLIT and NTRK-like protein 6 OS=Homo sapiens GN=SLITRK6 | 34,849 | 3,33 | 95110 | 2 | 19,77 | 18,46 | 22,60 | 19,79 | 19,90 | 20,93 | 0,952 | 9,58E-01 | 0 | -0,0705827 | 0,01842419 |
| RBM14_HUN | RBM14 | sp\|Q96PK6\|RBM14_HUMAN RNA-binding protein 14 OS=Homo sapiens GN=RBM14 PE= | 90,512 | 7,32 | 69492 | 2 | 17,71 | 17,43 | 21,89 | 18,44 | 16,41 | 21,96 | 0,951 | 9,75E-01 | 0 | -0,0723139 | 0,01098952 |
| DD19A_HUN | DDX19A | sp\|Q9NUU7\|DD19A_HUMAN ATP-dependent RNA helicase DDX19A OS=Homo sapiens | 308,184 | 21,13 | 53975 | 7 | 24,35 | 24,64 | 25,17 | 24,27 | 24,34 | 25,33 | 0,950 | 8,69E-01 | 0 | -0,0733331 | 0,0608412 |
| MAP11_HUN | METAP1 | sp\|P53582\|MAP11_HUMAN Methionine aminopeptidase 1 OS=Homo sapiens GN=MET | 79,19 | 10,62 | 43215 | 3 | 22,69 | 22,88 | 22,91 | 22,71 | 22,65 | 22,90 | 0,950 | 5,12E-01 | 0 | -0,0733337 | 0,29049651 |
| ITIH3_HUMA | ITIH3 | sp\|Q06033\|ITIH3_HUMAN Inter-alpha-trypsin inhibitor heavy chain H3 OS=Homo sapier | 200,833 | 4,16 | 99849 | 4 | 25,29 | 22,97 | 26,36 | 25,40 | 23,30 | 25,70 | 0,950 | 9,56E-01 | 0 | -0,0733337 | 0,01947337 |
| M4K4_HUM | MAP4K4 | sp\|O95819\|M4K4_HUMAN Mitogen-activated protein kinase kinase kinase kinase 4 OS= | 543,471 | 20,1 | 142101 | 20 | 24,24 | 24,00 | 22,59 | 24,31 | 23,64 | 22,66 | 0,950 | 9,22E-01 | 0 | -0,0733337 | 0,0352885 |
| ERP44_HUM | ERP44 | sp\|Q9BS26\|ERP44_HUMAN Endoplasmic reticulum resident protein 44 OS=Homo sapier | 65,314 | 6,4 | 46971 | 3 | 22,45 | 22,19 | 24,27 | 22,79 | 22,30 | 23,60 | 0,950 | 9,27E-01 | 0 | -0,0733337 | 0,03272919 |
| CCD25_HUM | CCDC25 | sp\|Q86WR0\|CCD25_HUMAN Coiled-coil domain-containing protein 25 OS=Homo sapier | 38,071 | 11,06 | 24479 | 2 | 23,45 | 23,09 | 21,88 | 23,39 | 22,92 | 21,89 | 0,950 | 9,16E-01 | 0 | -0,0733337 | 0,03832878 |
| RFA2_HUMA | RPA2 | sp\|P15927\|RFA2_HUMAN Replication protein A 32 kDa subunit OS=Homo sapiens GN=R | 11,263 | 2,96 | 29247 | 1 | 18,39 | 18,55 | 21,94 | 17,56 | 18,79 | 22,30 | 0,949 | 9,69E-01 | 0 | -0,0758591 | 0,01369592 |
| WDR20_HUN | WDR20 | sp\|Q8TBZ3\|WDR20_HUMAN WD repeat-containing protein 20 OS=Homo sapiens GN=W | 93,451 | 12,13 | 62893 | 4 | 20,94 | 21,00 | 19,80 | 21,09 | 21,06 | 19,36 | 0,948 | 9,17E-01 | 0 | -0,0766665 | 0,03755723 |
| CO6A3_HUN | COL6A3 | sp\|P12111\|CO6A3_HUMAN Collagen alpha-3(VI) chain OS=Homo sapiens GN=COL6A3 P | 152,78 | 2,23 | 343669 | 6 | 23,95 | 22,43 | 23,49 | 23,36 | 22,83 | 23,45 | 0,948 | 8,83E-01 | 0 | -0,0766665 | 0,05393576 |
| TFR1_HUMA | TFRC | sp\|P02786\|TFR1_HUMAN Transferrin receptor protein 1 OS=Homo sapiens GN=TFRC PE | 158,125 | 7,24 | 84871 | 5 | 25,57 | 24,97 | 25,47 | 25,41 | 25,22 | 25,15 | 0,948 | 7,23E-01 | 0 | -0,0766665 | 0,14114339 |
| GBRAP_HUN | GABARAP | sp\|O95166\|GBRAP_HUMAN Gamma-aminobutyric acid receptor-associated protein OS= | 29,394 | 29,91 | 13918 | 3 | 22,37 | 22,65 | 21,42 | 22,86 | 21,93 | 21,42 | 0,948 | 8,98E-01 | 0 | -0,0766665 | 0,04665313 |
| TOM34_HUN | TOMM34 | sp\|Q15785\|TOM34_HUMAN Mitochondrial import receptor subunit TOM34 OS=Homo | 12,866 | 3,56 | 34559 | 1 | 21,46 | 21,52 | 20,76 | 21,65 | 21,31 | 20,55 | 0,948 | 8,60E-01 | 0 | -0,0766671 | 0,06571007 |
| INSR_HUMA | INSR | sp\|P06213\|INSR_HUMAN Insulin receptor OS=Homo sapiens GN=INSR PE=1 SV=4 | 482,243 | 14,04 | 156333 | 13 | 23,95 | 24,60 | 21,97 | 23,99 | 24,23 | 22,07 | 0,948 | 9,45E-01 | 0 | -0,0766671 | 0,02454273 |
| WDR36_HUN | WDR36 | sp\|Q8NI36\|WDR36_HUMAN WD repeat-containing protein 36 OS=Homo sapiens GN=W | 28,259 | 3,15 | 105322 | 1 | 19,27 | 18,49 | 18,73 | 18,67 | 17,18 | 20,40 | 0,948 | 9,40E-01 | 0 | -0,0769393 | 0,02693521 |
| UBL3_HUMA | UBL3 | sp\|O95164\|UBL3_HUMAN Ubiquitin-like protein 3 OS=Homo sapiens GN=UBL3 PE=1 SV | 19,766 | 10,26 | 13157 | 1 | 20,42 | 20,35 | 18,36 | 20,46 | 19,73 | 18,71 | 0,948 | 9,31E-01 | 0 | -0,0773214 | 0,0308459 |
| DBNL_HUMA | DBNL | sp\|Q9UJU6\|DBNL_HUMAN Drebrin-like protein OS=Homo sapiens GN=DBNL PE=1 SV=1 | 122,987 | 10 | 48207 | 4 | 24,62 | 24,67 | 23,63 | 24,57 | 24,61 | 23,50 | 0,946 | 8,80E-01 | 0 | -0,0799999 | 0,05557746 |
| AAKG1_HUN | PRKAG1 | sp\|P54619\|AAKG1_HUMAN 5'-AMP-activated protein kinase subunit gamma-1 OS=Hom | 135,949 | 18,43 | 37579 | 6 | 23,31 | 23,28 | 23,10 | 22,87 | 23,46 | 23,12 | 0,946 | 6,85E-01 | 0 | -0,0799999 | 0,16446521 |
| CTR3_HUMA | SLC7A3 | sp\|Q8WY07\|CTR3_HUMAN Cationic amino acid transporter 3 OS=Homo sapiens GN=SL | 79,278 | 2,1 | 67169 | 1 | 23,06 | 23,44 | 23,90 | 22,84 | 23,57 | 23,75 | 0,946 | 8,39E-01 | 0 | -0,0799999 | 0,07618455 |
| ARAF_HUMA | ARAF | sp\|P10398\|ARAF_HUMAN Serine/threonine-protein kinase A-Raf OS=Homo sapiens GN | 97,897 | 8,58 | 67585 | 4 | 22,84 | 22,82 | 23,00 | 22,53 | 23,23 | 22,66 | 0,946 | 7,37E-01 | 0 | -0,0799999 | 0,13239468 |
| SEPT9_HUM | SEPT9 | sp\|Q9UHD8\|SEPT9_HUMAN Septin-9 OS=Homo sapiens GN=SEPT9 PE=1 SV=2 | 323,965 | 19,28 | 65401 | 9 | 25,93 | 25,96 | 24,53 | 25,77 | 25,91 | 24,50 | 0,946 | 9,08E-01 | 0 | -0,0799999 | 0,04186111 |
| SF3B1_HUM | SF3B1 | sp\|O75533\|SF3B1_HUMAN Splicing factor 3B subunit 1 OS=Homo sapiens GN=SF3B1 PE | 908,248 | 28,91 | 145830 | 28 | 26,29 | 26,25 | 26,23 | 26,06 | 26,24 | 26,23 | 0,946 | 2,60E-01 | 0 | -0,0800006 | 0,58507145 |
| GCN1_HUM | GCN1 | sp\|Q92616\|GCN1_HUMAN eIF-2-alpha kinase activator GCN1 OS=Homo sapiens GN=GC | 1391,759 | 19,99 | 292758 | 41 | 32,62 | 32,34 | 32,85 | 32,69 | 32,37 | 32,51 | 0,946 | 6,70E-01 | 0 | -0,0800006 | 0,1740619 |
| ENPL_HUMA | HSP90B1 | sp\|P14625\|ENPL_HUMAN Endoplasmin OS=Homo sapiens GN=HSP90B1 PE=1 SV=1 | 1547,701 | 50,06 | 92469 | 39 | 28,29 | 27,68 | 29,04 | 28,30 | 28,09 | 28,38 | 0,946 | 8,52E-01 | 0 | -0,0800012 | 0,06945298 |
| ARF5_HUMA | ARF5 | sp\|P84085\|ARF5_HUMAN ADP-ribosylation factor 5 OS=Homo sapiens GN=ARF5 PE=1 S | 487,208 | 47,78 | 20530 | 9 | 22,66 | 22,42 | 24,56 | 22,03 | 23,18 | 24,18 | 0,944 | 9,32E-01 | 0 | -0,0833327 | 0,03054401 |
| UTRO_HUM | UTRN | sp\|P46939\|UTRO_HUMAN Utrophin OS=Homo sapiens GN=UTRN PE=1 SV=2 | 438,294 | 5,18 | 394466 | 14 | 24,94 | 25,01 | 21,22 | 24,34 | 24,15 | 22,43 | 0,944 | 9,55E-01 | 0 | -0,0833333 | 0,01994383 |
| FAM3C_HUN | FAM3C | sp\|Q92520\|FAM3C_HUMAN Protein FAM3C OS=Homo sapiens GN=FAM3C PE=1 SV=1 | 150,481 | 18,94 | 24680 | 5 | 24,98 | 24,66 | 24,60 | 25,23 | 24,02 | 24,74 | 0,944 | 8,33E-01 | 0 | -0,0833333 | 0,07928508 |
| RLA0_HUMA | RPLP0 | sp\|P05388\|RLA0_HUMAN 60S acidic ribosomal protein P0 OS=Homo sapiens GN=RPLP0 | 498,768 | 44,79 | 34274 | 14 | 28,21 | 28,18 | 28,26 | 27,94 | 28,13 | 28,33 | 0,944 | 5,09E-01 | 0 | -0,0833333 | 0,29349639 |
| RS27L_HUM | RPS27L | sp\|Q71UM5\|RS27L_HUMAN 40S ribosomal protein S27-like OS=Homo sapiens GN=RPS2 | 186,454 | 38,1 | 9477 | 4 | 22,52 | 22,87 | 21,40 | 22,78 | 22,65 | 21,11 | 0,944 | 9,10E-01 | 0 | -0,0833333 | 0,04074559 |
| SCFD1_HUM | SCFD1 | sp\|Q8WVM8\|SCFD1_HUMAN Sec1 family domain-containing protein 1 OS=Homo sapier | 164,797 | 14,8 | 72380 | 6 | 23,68 | 23,70 | 23,19 | 23,37 | 23,10 | 23,85 | 0,944 | 7,77E-01 | 0 | -0,0833333 | 0,10936778 |
| NEK7_HUMA | NEK7 | sp\|Q8TDX7\|NEK7_HUMAN Serine/threonine-protein kinase Nek7 OS=Homo sapiens GN | 37,004 | 8,61 | 34551 | 2 | 22,47 | 22,02 | 22,83 | 22,31 | 22,14 | 22,62 | 0,944 | 7,76E-01 | 0 | -0,0833333 | 0,11038032 |
| TPPC3_HUM | TRAPPC3 | sp\|O43617\|TPPC3_HUMAN Trafficking protein particle complex subunit 3 OS=Homo sap | 26,965 | 6,11 | 20274 | 1 | 22,26 | 22,44 | 22,18 | 22,50 | 22,42 | 21,71 | 0,944 | 7,67E-01 | 0 | -0,083334 | 0,11529773 |
| F10A1_HUM | ST13 | sp\|P50502\|F10A1_HUMAN Hsc70-interacting protein OS=Homo sapiens GN=ST13 PE=1 | 348,805 | 20,33 | 41332 | 10 | 28,07 | 28,07 | 26,62 | 28,08 | 27,88 | 26,55 | 0,944 | 9,09E-01 | 0 | -0,083334 | 0,04165292 |
| PRDX4_HUM | PRDX4 | sp\|Q13162\|PRDX4_HUMAN Peroxiredoxin-4 OS=Homo sapiens GN=PRDX4 PE=1 SV=1 | 412,898 | 49,82 | 30540 | 12 | 25,36 | 25,80 | 26,68 | 25,73 | 25,72 | 26,14 | 0,944 | 8,50E-01 | 0 | -0,083334 | 0,07080105 |
| SNR40_HUM | SNRNP40 | sp\|Q96DI7\|SNR40_HUMAN U5 small nuclear ribonucleoprotein 40 kDa protein OS=Hom | 54,415 | 24,37 | 39311 | 4 | 22,10 | 22,08 | 23,07 | 21,51 | 22,12 | 23,36 | 0,942 | 8,98E-01 | 0 | -0,0866661 | 0,04672235 |
| RAB34_HUM | RAB34 | sp\|Q9BZG1\|RAB34_HUMAN Ras-related protein Rab-34 OS=Homo sapiens GN=RAB34 P | 249,604 | 32,05 | 29044 | 7 | 24,92 | 24,92 | 24,42 | 24,64 | 24,87 | 24,49 | 0,942 | 6,87E-01 | 0 | -0,0866667 | 0,16299446 |
| PPIB_HUMA | PPIB | sp\|P23284\|PPIB_HUMAN Peptidyl-prolyl cis-trans isomerase B OS=Homo sapiens GN=P | 473,967 | 47,69 | 23743 | 12 | 26,76 | 26,76 | 27,65 | 26,77 | 27,06 | 27,08 | 0,942 | 7,96E-01 | 0 | -0,0866667 | 0,09927264 |
| KIF1B_HUM | KIF1B | sp\|O60333\|KIF1B_HUMAN Kinesin-like protein KIF1B OS=Homo sapiens GN=KIF1B PE=1 | 189,766 | 3,58 | 204476 | 5 | 21,69 | 21,60 | 21,42 | 21,77 | 21,58 | 21,10 | 0,942 | 7,07E-01 | 0 | -0,0866667 | 0,15060866 |
| DYN2_HUM | DNM2 | sp\|P50570\|DYN2_HUMAN Dynamin-2 OS=Homo sapiens GN=DNM2 PE=1 SV=2 | 571,241 | 32,99 | 98064 | 25 | 25,32 | 25,26 | 24,72 | 25,13 | 25,23 | 24,68 | 0,942 | 7,51E-01 | 0 | -0,0866667 | 0,12434492 |
| SQSTM_HUN | SQSTM1 | sp\|Q13501\|SQSTM_HUMAN Sequestosome-1 OS=Homo sapiens GN=SQSTM1 PE=1 SV= | 297,424 | 20 | 47687 | 7 | 23,81 | 23,45 | 24,31 | 23,74 | 23,35 | 24,22 | 0,942 | 8,19E-01 | 0 | -0,0866667 | 0,08684594 |
| MTMR2_HUI | MTMR2 | sp\|Q13614\|MTMR2_HUMAN Myotubularin-related protein 2 OS=Homo sapiens GN=M | 24,596 | 2,95 | 73381 | 2 | 19,87 | 20,75 | 19,22 | 20,01 | 20,38 | 19,19 | 0,942 | 8,86E-01 | 0 | -0,0866667 | 0,05272075 |
| DDAH1_HUN | DDAH1 | sp\|O94760\|DDAH1_HUMAN N(G),N(G)-dimethylarginine dimethylaminohydrolase 1 OS= | 222,947 | 44,21 | 31122 | 9 | 25,44 | 25,28 | 23,90 | 25,40 | 24,99 | 23,97 | 0,942 | 9,00E-01 | 0 | -0,0866674 | 0,04573762 |
| YJ017_HUMAN | | sp\|Q8NDZ9\|YJ017_HUMAN Putative uncharacterized protein LOC439951 OS=Homo sap | 16,815 | 5,12 | 22305 | 1 | 22,59 | 22,30 | 17,86 | 22,32 | 22,28 | 17,89 | 0,941 | 9,69E-01 | 0 | -0,0872733 | 0,013591 |
| SF3B5_HUM | SF3B5 | sp\|Q9BWJ5\|SF3B5_HUMAN Splicing factor 3B subunit 5 OS=Homo sapiens GN=SF3B5 PE | 14,6 | 17,44 | 10135 | 1 | 19,41 | 19,89 | 20,61 | 19,41 | 19,55 | 20,68 | 0,940 | 8,76E-01 | 0 | -0,088892 | 0,05770095 |
| AT2B1_HUM | ATP2B1 | sp\|P20020\|AT2B1_HUMAN Plasma membrane calcium-transporting ATPase 1 OS=Homo | 1799,04 | 35,69 | 138755 | 41 | 26,89 | 27,17 | 25,91 | 26,76 | 27,02 | 25,92 | 0,940 | 8,67E-01 | 0 | -0,0899995 | 0,06173703 |
| NRP1_HUMA | NRP1 | sp\|O14786\|NRP1_HUMAN Neuropilin-1 OS=Homo sapiens GN=NRP1 PE=1 SV=3 | 243,916 | 10,18 | 103134 | 6 | 24,99 | 24,60 | 24,89 | 24,99 | 24,66 | 24,56 | 0,940 | 6,34E-01 | 0 | -0,0900002 | 0,19805639 |
| DLGP4_HUM | DLGAP4 | sp\|Q9Y2H0\|DLGP4_HUMAN Disks large-associated protein 4 OS=Homo sapiens GN=DLG | 28,541 | 1,11 | 108012 | 1 | 21,60 | 21,29 | 20,25 | 21,37 | 21,10 | 20,40 | 0,940 | 8,66E-01 | 0 | -0,0900002 | 0,06250107 |
| STX4_HUMA | STX4 | sp\|Q12846\|STX4_HUMAN Syntaxin-4 OS=Homo sapiens GN=STX4 PE=1 SV=2 | 460,364 | 34,01 | 34180 | 11 | 27,42 | 27,47 | 26,15 | 27,13 | 27,00 | 26,64 | 0,940 | 8,53E-01 | 0 | -0,0900002 | 0,06895295 |
| FPRP_HUMA | PTGFRN | sp\|Q9P2B2\|FPRP_HUMAN Prostaglandin F2 receptor negative regulator OS=Homo sapie | 121,988 | 7,74 | 98556 | 6 | 24,27 | 23,58 | 22,35 | 24,34 | 23,27 | 22,32 | 0,940 | 9,17E-01 | 0 | -0,0900002 | 0,03769837 |
| K1C18_HUM | KRT18 | sp\|P05783\|K1C18_HUMAN Keratin, type I cytoskeletal 18 OS=Homo sapiens GN=KRT18 | 97,652 | 7,91 | 48058 | 3 | 22,94 | 22,63 | 21,00 | 22,84 | 23,07 | 20,39 | 0,940 | 9,36E-01 | 0 | -0,0900002 | 0,02887924 |
| KAD2_HUMA | AK2 | sp\|P54819\|KAD2_HUMAN Adenylate kinase 2, mitochondrial OS=Homo sapiens GN=AK | 179,112 | 31,38 | 26478 | 5 | 23,72 | 24,18 | 22,38 | 23,55 | 23,81 | 22,65 | 0,940 | 8,96E-01 | 0 | -0,0900002 | 0,04786351 |
| SC24A_HUM | SEC24A | sp\|O95486\|SC24A_HUMAN Protein transport protein Sec24A OS=Homo sapiens GN=SE | 166,671 | 5,49 | 119749 | 4 | 21,34 | 21,63 | 22,88 | 20,97 | 21,56 | 23,05 | 0,940 | 9,14E-01 | 0 | -0,0900002 | 0,03926782 |
| AHNK_HUM | AHNAK | sp\|Q09666\|AHNK_HUMAN Neuroblast differentiation-associated protein AHNAK OS=H | 1527,806 | 17,76 | 629101 | 43 | 28,98 | 29,02 | 27,07 | 29,15 | 28,92 | 26,73 | 0,940 | 9,33E-01 | 0 | -0,0900002 | 0,03016227 |
| CHM1A_HUN | CHMP1A | sp\|Q9HD42\|CHM1A_HUMAN Charged multivesicular body protein 1a OS=Homo sapiens | 243,639 | 22,45 | 21703 | 10 | 28,54 | 27,80 | 26,38 | 28,45 | 27,84 | 26,15 | 0,937 | 9,25E-01 | 0 | -0,0933329 | 0,03370954 |
| AP2A2_HUM | AP2A2 | sp\|O94973\|AP2A2_HUMAN AP-2 complex subunit alpha-2 OS=Homo sapiens GN=AP2A | 1223,995 | 39,3 | 103960 | 28 | 27,06 | 26,80 | 27,49 | 26,94 | 27,13 | 27,00 | 0,937 | 6,78E-01 | 0 | -0,0933329 | 0,16870808 |
| CIP2A_HUM | KIAA1524 | sp\|Q8TCG1\|CIP2A_HUMAN Protein CIP2A OS=Homo sapiens GN=KIAA1524 PE=1 SV=2 | 40,316 | 2,98 | 102185 | 1 | 18,14 | 19,08 | 19,87 | 18,47 | 18,48 | 19,86 | 0,937 | 8,98E-01 | 0 | -0,0933336 | 0,04694321 |
| ARL2_HUMA | ARL2 | sp\|P36404\|ARL2_HUMAN ADP-ribosylation factor-like protein 2 OS=Homo sapiens GN= | 141,317 | 23,37 | 20878 | 4 | 23,89 | 23,98 | 23,72 | 23,30 | 23,55 | 24,46 | 0,937 | 8,09E-01 | 0 | -0,0933336 | 0,09229326 |
| ROCK1_HUN | ROCK1 | sp\|Q13464\|ROCK1_HUMAN Rho-associated protein kinase 1 OS=Homo sapiens GN=RO | 33,944 | 1,7 | 158175 | 2 | 21,18 | 21,01 | 20,30 | 20,47 | 20,92 | 20,82 | 0,937 | 7,73E-01 | 0 | -0,0933336 | 0,11195561 |
| S10AB_HUM | S100A11 | sp\|P31949\|S10AB_HUMAN Protein S100-A11 OS=Homo sapiens GN=S100A11 PE=1 SV= | 271,708 | 35,24 | 11740 | 5 | 27,20 | 27,48 | 27,22 | 26,75 | 27,98 | 26,89 | 0,937 | 8,27E-01 | 0 | -0,0933336 | 0,08271856 |
| STK3_HUMA | STK3 | sp\|Q13188\|STK3_HUMAN Serine/threonine-protein kinase 3 OS=Homo sapiens GN=STK | 44,398 | 4,48 | 56301 | 2 | 21,86 | 21,55 | 20,21 | 20,98 | 21,40 | 20,96 | 0,937 | 8,68E-01 | 0 | -0,09342 | 0,06162912 |
| PTH2_HUMA | PTRH2 | sp\|Q9Y3E5\|PTH2_HUMAN Peptidyl-tRNA hydrolase 2, mitochondrial OS=Homo sapiens | 37,244 | 8,94 | 19194 | 2 | 17,79 | 18,63 | 20,71 | 17,78 | 19,81 | 19,26 | 0,936 | 9,33E-01 | 0 | -0,0948995 | 0,03019342 |
| HS71A_HUM | HSPA1A | sp\|P0DMV8\|HS71A_HUMAN Heat shock 70 kDa protein 1A OS=Homo sapiens GN=HSPA | 720,516 | 27,61 | 70052 | 18 | 26,57 | 25,89 | 26,21 | 26,40 | 26,00 | 25,98 | 0,935 | 7,07E-01 | 0 | -0,0966663 | 0,15059943 |
| RHG01_HUN | ARHGAP1 | sp\|Q07960\|RHG01_HUMAN Rho GTPase-activating protein 1 OS=Homo sapiens GN=AR | 474,69 | 47,38 | 50436 | 18 | 26,57 | 26,60 | 26,01 | 26,29 | 26,66 | 25,94 | 0,935 | 7,50E-01 | 0 | -0,0966663 | 0,12507791 |
| STOM_HUM | STOM | sp\|P27105\|STOM_HUMAN Erythrocyte band 7 integral membrane protein OS=Homo sa | 233,061 | 24,31 | 31731 | 5 | 26,31 | 25,09 | 24,51 | 26,31 | 25,19 | 24,12 | 0,935 | 9,12E-01 | 0 | -0,0966663 | 0,03981295 |
| CTL2_HUMA | SLC44A2 | sp\|Q8IWA5\|CTL2_HUMAN Choline transporter-like protein 2 OS=Homo sapiens GN=SL | 127,347 | 9,63 | 80124 | 5 | 24,21 | 23,77 | 22,80 | 23,65 | 24,05 | 22,79 | 0,935 | 8,71E-01 | 0 | -0,0966663 | 0,0600129 |
| CTR1_HUMA | SLC7A1 | sp\|P30825\|CTR1_HUMAN High affinity cationic amino acid transporter 1 OS=Homo sapi | 136,175 | 7,15 | 67638 | 3 | 20,65 | 21,46 | 24,15 | 20,98 | 21,05 | 23,94 | 0,935 | 9,50E-01 | 0 | -0,0966663 | 0,02243319 |
| DC1L2_HUM | DYNC1LI2 | sp\|O43237\|DC1L2_HUMAN Cytoplasmic dynein 1 light intermediate chain 2 OS=Homo s | 206,421 | 24,19 | 54099 | 7 | 24,01 | 23,78 | 24,54 | 23,62 | 23,95 | 24,47 | 0,935 | 7,87E-01 | 0 | -0,096667 | 0,10407724 |
| CAVN1_HUN | CAVIN1 | sp\|Q6NZI2\|CAVN1_HUMAN Caveolae-associated protein 1 OS=Homo sapiens GN=CAVI | 546,579 | 20 | 43476 | 11 | 26,72 | 26,94 | 27,28 | 26,71 | 26,95 | 26,99 | 0,935 | 6,29E-01 | 0 | -0,096667 | 0,20156962 |
| 4F2_HUMAN | SLC3A2 | sp\|P08195\|4F2_HUMAN 4F2 cell-surface antigen heavy chain OS=Homo sapiens GN=SL | 280,958 | 16,83 | 67994 | 10 | 28,58 | 28,70 | 26,93 | 28,56 | 28,37 | 26,99 | 0,935 | 9,04E-01 | 0 | -0,096667 | 0,04365481 |
| GMFB_HUM | GMFB | sp\|P60983\|GMFB_HUMAN Glia maturation factor beta OS=Homo sapiens GN=GMFB PE | 78,432 | 24,65 | 16713 | 3 | 25,02 | 25,04 | 23,01 | 25,05 | 24,73 | 23,00 | 0,935 | 9,22E-01 | 0 | -0,0966676 | 0,03529512 |
| MED1_HUM | MED1 | sp\|Q15648\|MED1_HUMAN Mediator of RNA polymerase II transcription subunit 1 OS=H | 29,872 | 0,57 | 168478 | 1 | 18,90 | 18,64 | 21,62 | 17,88 | 19,06 | 21,93 | 0,935 | 9,53E-01 | 0 | -0,097023 | 0,02108892 |
| ES1_HUMAN | C21orf33 | sp\|P30042\|ES1_HUMAN ES1 protein homolog, mitochondrial OS=Homo sapiens GN=C2 | 31,269 | 3,73 | 28170 | 1 | 17,78 | 18,74 | 22,84 | 19,18 | 17,87 | 22,02 | 0,935 | 9,63E-01 | 0 | -0,0975761 | 0,01637427 |
| DOCK5_HUN | DOCK5 | sp\|Q9H7D0\|DOCK5_HUMAN Dedicator of cytokinesis protein 5 OS=Homo sapiens GN=D | 18,613 | 0,64 | 215309 | 1 | 20,43 | 19,54 | 20,05 | 20,84 | 20,33 | 18,56 | 0,934 | 9,01E-01 | 0 | -0,0981007 | 0,04544872 |
| DDX20_HUM | DDX20 | sp\|Q9UHI6\|DDX20_HUMAN Probable ATP-dependent RNA helicase DDX20 OS=Homo sa | 11,741 | 3,16 | 92241 | 1 | 19,66 | 18,58 | 21,28 | 18,24 | 19,80 | 21,19 | 0,933 | 9,36E-01 | 0 | -0,0995496 | 0,02890816 |
| GALK1_HUM | GALK1 | sp\|P51570\|GALK1_HUMAN Galactokinase OS=Homo sapiens GN=GALK1 PE=1 SV=1 | 74,889 | 10,2 | 42272 | 4 | 23,90 | 24,26 | 24,51 | 23,86 | 24,11 | 24,40 | 0,933 | 6,94E-01 | 0 | -0,0999997 | 0,15892934 |
| PFKAM_HUN | PFKM | sp\|P08237\|PFKAM_HUMAN ATP-dependent 6-phosphofructokinase, muscle type OS=H | 536,308 | 26,15 | 85183 | 14 | 23,95 | 24,57 | 25,68 | 23,94 | 24,37 | 25,59 | 0,933 | 8,94E-01 | 0 | -0,0999997 | 0,04846622 |
| CHMP5_HUN | CHMP5 | sp\|Q9NZZ3\|CHMP5_HUMAN Charged multivesicular body protein 5 OS=Homo sapiens | 355,571 | 52,51 | 24571 | 7 | 26,73 | 26,50 | 24,50 | 26,88 | 26,33 | 24,22 | 0,933 | 9,30E-01 | 0 | -0,1000004 | 0,03130746 |
| STXB1_HUM | STXBP1 | sp\|P61764\|STXB1_HUMAN Syntaxin-binding protein 1 OS=Homo sapiens GN=STXBP1 PE | 631,877 | 39,39 | 67569 | 20 | 26,58 | 26,39 | 23,96 | 26,10 | 26,15 | 24,37 | 0,931 | 9,25E-01 | 0 | -0,1033325 | 0,03401542 |
| CARM1_HUN | CARM1 | sp\|Q86X55\|CARM1_HUMAN Histone-arginine methyltransferase CARM1 OS=Homo sapi | 225,596 | 12,5 | 65854 | 6 | 24,16 | 24,34 | 25,22 | 24,23 | 24,16 | 25,02 | 0,931 | 8,21E-01 | 0 | -0,1033332 | 0,0855849 |
| PLOD3_HUM | PLOD3 | sp\|O60568\|PLOD3_HUMAN Procollagen-lysine,2-oxoglutarate 5-dioxygenase 3 OS=Hon | 66,671 | 2,98 | 84785 | 2 | 21,41 | 20,42 | 22,15 | 21,47 | 20,19 | 22,01 | 0,931 | 8,95E-01 | 0 | -0,1033332 | 0,04808095 |
| MLP3B_HUN | MAP1LC3B | sp\|Q9GZQ8\|MLP3B_HUMAN Microtubule-associated proteins 1A/1B light chain 3B OS= | 45,707 | 22,4 | 14688 | 3 | 24,20 | 24,51 | 23,04 | 24,36 | 24,52 | 22,56 | 0,931 | 9,00E-01 | 0 | -0,1033338 | 0,04580529 |
| TPM2_HUM | TPM2 | sp\|P07951\|TPM2_HUMAN Tropomyosin beta chain OS=Homo sapiens GN=TPM2 PE=1 S | 831,95 | 46,13 | 32851 | 21 | 25,18 | 25,62 | 23,27 | 25,18 | 25,22 | 23,36 | 0,931 | 9,18E-01 | 0 | -0,1033338 | 0,03699053 |
| ELAV1_HUM | ELAVL1 | sp\|Q15717\|ELAV1_HUMAN ELAV-like protein 1 OS=Homo sapiens GN=ELAVL1 PE=1 SV= | 202,57 | 28,22 | 36092 | 7 | 24,69 | 24,59 | 25,70 | 24,58 | 24,82 | 25,27 | 0,931 | 8,13E-01 | 0 | -0,1033338 | 0,09011942 |
| LPPRC_HUM | LRPPRC | sp\|P42704\|LPPRC_HUMAN Leucine-rich PPR motif-containing protein, mitochondrial OS | 49,113 | 1,87 | 157905 | 3 | 23,12 | 23,10 | 23,21 | 23,21 | 23,43 | 22,48 | 0,931 | 7,39E-01 | 0 | -0,1033338 | 0,13145678 |
| PLMN_HUM | PLG | sp\|P00747\|PLMN_HUMAN Plasminogen OS=Homo sapiens GN=PLG PE=1 SV=2 | 250,83 | 7,65 | 90569 | 7 | 24,95 | 23,20 | 25,50 | 25,34 | 23,39 | 24,61 | 0,931 | 9,14E-01 | 0 | -0,1033338 | 0,03913637 |
| SRP14_HUM | SRP14 | sp\|P37108\|SRP14_HUMAN Signal recognition particle 14 kDa protein OS=Homo sapiens | 249,839 | 27,21 | 14570 | 6 | 26,33 | 26,36 | 26,25 | 26,38 | 26,58 | 25,67 | 0,931 | 7,29E-01 | 0 | -0,1033338 | 0,13726631 |
| GDIB_HUMA | GDI2 | sp\|P50395\|GDIB_HUMAN Rab GDP dissociation inhibitor beta OS=Homo sapiens GN=GD | 1377,202 | 58,43 | 50663 | 32 | 28,29 | 28,22 | 27,29 | 27,97 | 28,22 | 27,30 | 0,931 | 8,19E-01 | 0 | -0,1033344 | 0,08660271 |
| VPS35_HUM | VPS35 | sp\|Q96QK1\|VPS35_HUMAN Vacuolar protein sorting-associated protein 35 OS=Homo s | 834,279 | 30,65 | 91707 | 19 | 26,27 | 26,36 | 26,37 | 25,97 | 26,24 | 26,48 | 0,931 | 5,31E-01 | 0 | -0,1033344 | 0,27524377 |
| ZC3HF_HUM | ZC3H15 | sp\|Q8WU90\|ZC3HF_HUMAN Zinc finger CCCH domain-containing protein 15 OS=Homo | 23,502 | 2,11 | 48602 | 1 | 14,33 | 19,32 | 20,54 | 16,97 | 16,12 | 20,78 | 0,930 | 9,67E-01 | 0 | -0,1050905 | 0,01462634 |
| MORC4_HUN | MORC4 | sp\|Q8TE76\|MORC4_HUMAN MORC family CW-type zinc finger protein 4 OS=Homo sapi | 13,882 | 0,75 | 106348 | 1 | 18,92 | 19,33 | 22,89 | 18,62 | 19,18 | 23,02 | 0,929 | 9,57E-01 | 0 | -0,1064587 | 0,01891309 |
| CLIC2_HUM | CLIC2 | sp\|O15247\|CLIC2_HUMAN Chloride intracellular channel protein 2 OS=Homo sapiens G | 37,79 | 7,29 | 28356 | 2 | 21,62 | 21,71 | 21,89 | 21,43 | 21,86 | 21,61 | 0,929 | 5,10E-01 | 0 | -0,1066659 | 0,29207817 |
| ARPC2_HUM | ARPC2 | sp\|O15144\|ARPC2_HUMAN Actin-related protein 2/3 complex subunit 2 OS=Homo sapi | 573,131 | 57 | 34333 | 17 | 27,29 | 27,08 | 26,40 | 26,93 | 27,25 | 26,27 | 0,929 | 8,00E-01 | 0 | -0,1066666 | 0,0968601 |
| PDIA3_HUM | PDIA3 | sp\|P30101\|PDIA3_HUMAN Protein disulfide-isomerase A3 OS=Homo sapiens GN=PDIA3 | 974,704 | 43,37 | 56782 | 26 | 27,93 | 27,96 | 29,32 | 28,25 | 28,09 | 28,55 | 0,929 | 8,34E-01 | 0 | -0,1066666 | 0,07868165 |
| PCYOX_HUM | PCYOX1 | sp\|Q9UHG3\|PCYOX_HUMAN Prenylcysteine oxidase 1 OS=Homo sapiens GN=PCYOX1 P | 50,862 | 2,57 | 56640 | 1 | 22,40 | 22,24 | 23,10 | 22,32 | 22,32 | 22,78 | 0,929 | 7,44E-01 | 0 | -0,1066666 | 0,12815417 |
| DYL2_HUMA | DYNLL2 | sp\|Q96FJ2\|DYL2_HUMAN Dynein light chain 2, cytoplasmic OS=Homo sapiens GN=DYN | 165,994 | 58,43 | 10350 | 5 | 21,81 | 22,80 | 21,84 | 22,30 | 22,32 | 21,51 | 0,929 | 8,12E-01 | 0 | -0,1066666 | 0,09029911 |
| RAB1B_HUM | RAB1B | sp\|Q9H0U4\|RAB1B_HUMAN Ras-related protein Rab-1B OS=Homo sapiens GN=RAB1B | 640,495 | 75,12 | 22171 | 14 | 26,09 | 26,14 | 26,29 | 26,00 | 25,96 | 26,24 | 0,929 | 3,72E-01 | 0 | -0,1066672 | 0,4299459 |
| RTN4_HUMA | RTN4 | sp\|Q9NQC3\|RTN4_HUMAN Reticulon-4 OS=Homo sapiens GN=RTN4 PE=1 SV=2 | 200,299 | 4,03 | 129931 | 4 | 26,87 | 26,76 | 26,27 | 26,70 | 26,98 | 25,90 | 0,929 | 7,89E-01 | 0 | -0,1066672 | 0,10303285 |
| BRE1A_HUM | RNF20 | sp\|Q5VTR2\|BRE1A_HUMAN E3 ubiquitin-protein ligase BRE1A OS=Homo sapiens GN=R | 27,063 | 3,18 | 113662 | 2 | 20,10 | 18,32 | 19,32 | 20,58 | 16,48 | 20,36 | 0,928 | 9,43E-01 | 0 | -0,1076946 | 0,02526255 |
| EXC6B_HUM | EXOC6B | sp\|Q9Y2D4\|EXC6B_HUMAN Exocyst complex component 6B OS=Homo sapiens GN=EXC | 48,678 | 2,34 | 94201 | 1 | 18,54 | 19,73 | 19,05 | 18,54 | 19,18 | 19,27 | 0,927 | 8,04E-01 | 0 | -0,1099993 | 0,0948904 |
| RB39A_HUM | RAB39A | sp\|Q14964\|RB39A_HUMAN Ras-related protein Rab-39A OS=Homo sapiens GN=RAB39 | 87,435 | 5,99 | 25007 | 2 | 21,92 | 21,85 | 22,55 | 22,18 | 21,81 | 22,00 | 0,927 | 6,79E-01 | 0 | -0,11 | 0,16814259 |
| AK1C4_HUM | AKR1C4 | sp\|P17516\|AK1C4_HUMAN Aldo-keto reductase family 1 member C4 OS=Homo sapiens | 71,423 | 8,36 | 37067 | 2 | 23,40 | 23,14 | 21,18 | 23,15 | 22,48 | 21,76 | 0,927 | 8,98E-01 | 0 | -0,11 | 0,04661447 |
| CCDC6_HUM | CCDC6 | sp\|Q16204\|CCDC6_HUMAN Coiled-coil domain-containing protein 6 OS=Homo sapiens | 125,441 | 16,67 | 53291 | 7 | 23,20 | 23,32 | 23,51 | 23,29 | 23,03 | 23,38 | 0,927 | 4,71E-01 | 0 | -0,11 | 0,32676223 |
| ITB5_HUMA | ITGB5 | sp\|P18084\|ITB5_HUMAN Integrin beta-5 OS=Homo sapiens GN=ITGB5 PE=1 SV=1 | 169,007 | 8,89 | 88054 | 6 | 23,91 | 23,43 | 21,17 | 23,72 | 23,42 | 21,04 | 0,927 | 9,31E-01 | 0 | -0,11 | 0,03096271 |
| PI51C_HUM | PIP5K1C | sp\|O60331\|PI51C_HUMAN Phosphatidylinositol 4-phosphate 5-kinase type-1 gamma O | 72,89 | 6,29 | 73260 | 3 | 21,15 | 22,37 | 21,25 | 21,79 | 21,64 | 21,01 | 0,927 | 8,22E-01 | 0 | -0,11 | 0,08506845 |
| NECT2_HUM | NECTIN2 | sp\|Q92692\|NECT2_HUMAN Nectin-2 OS=Homo sapiens GN=NECTIN2 PE=1 SV=1 | 95,163 | 4,83 | 57742 | 2 | 23,85 | 23,87 | 23,04 | 23,91 | 24,00 | 22,52 | 0,927 | 8,52E-01 | 0 | -0,1100006 | 0,06973838 |
| HMGA1_HUI | HMGA1 | sp\|P17096\|HMGA1_HUMAN High mobility group protein HMG-I/HMG-Y OS=Homo sapi | 46,4 | 17,76 | 11676 | 1 | 21,07 | 21,69 | 19,55 | 21,74 | 21,42 | 18,82 | 0,926 | 9,26E-01 | 0 | -0,1102962 | 0,03316099 |
| TBC15_HUM | TBC1D15 | sp\|Q8TC07\|TBC15_HUMAN TBC1 domain family member 15 OS=Homo sapiens GN=TBC | 41,922 | 3,33 | 79491 | 2 | 20,31 | 19,85 | 21,85 | 19,08 | 20,54 | 22,05 | 0,925 | 9,20E-01 | 0 | -0,1119976 | 0,03618337 |
| EIF1A_HUM | EIF1AD | sp\|Q8N9N8\|EIF1A_HUMAN Probable RNA-binding protein EIF1AD OS=Homo sapiens G | 23,75 | 9,7 | 19053 | 1 | 20,72 | 20,88 | 19,15 | 20,68 | 20,49 | 19,24 | 0,924 | 8,81E-01 | 0 | -0,1133327 | 0,05480667 |
| IMA7_HUMA | KPNA6 | sp\|O60684\|IMA7_HUMAN Importin subunit alpha-7 OS=Homo sapiens GN=KPNA6 PE=1 | 562,202 | 40,3 | 60030 | 15 | 24,68 | 25,02 | 24,91 | 24,71 | 24,86 | 24,70 | 0,924 | 3,72E-01 | 0 | -0,1133334 | 0,42985279 |
| CAPZB_HUM | CAPZB | sp\|P47756\|CAPZB_HUMAN F-actin-capping protein subunit beta OS=Homo sapiens GN= | 1090,995 | 66,43 | 31350 | 23 | 28,02 | 28,16 | 27,21 | 27,92 | 27,88 | 27,25 | 0,924 | 7,73E-01 | 0 | -0,1133334 | 0,11185677 |
| FABP5_HUM | FABP5 | sp\|Q01469\|FABP5_HUMAN Fatty acid-binding protein, epidermal OS=Homo sapiens GN | 78,208 | 14,07 | 15164 | 3 | 24,77 | 25,06 | 24,19 | 24,88 | 25,05 | 23,75 | 0,924 | 8,25E-01 | 0 | -0,113334 | 0,08329539 |
| UTP15_HUM | UTP15 | sp\|Q8TED0\|UTP15_HUMAN U3 small nucleolar RNA-associated protein 15 homolog OS= | 26,319 | 1,93 | 58415 | 1 | 20,33 | 18,72 | 19,14 | 18,89 | 17,71 | 21,24 | 0,924 | 9,25E-01 | 0 | -0,1144218 | 0,0337869 |
| SAP18_HUM | SAP18 | sp\|O00422\|SAP18_HUMAN Histone deacetylase complex subunit SAP18 OS=Homo sapie | 20,607 | 20,26 | 17561 | 3 | 18,43 | 17,44 | 23,11 | 17,70 | 17,61 | 23,32 | 0,923 | 9,66E-01 | 0 | -0,1162917 | 0,01496454 |
| A2MG_HUM | A2M | sp\|P01023\|A2MG_HUMAN Alpha-2-macroglobulin OS=Homo sapiens GN=A2M PE=1 SV | 260,931 | 3,93 | 163291 | 7 | 28,15 | 26,48 | 29,31 | 28,26 | 26,65 | 28,68 | 0,922 | 9,15E-01 | 0 | -0,1166662 | 0,03851484 |
| PRRC1_HUM | PRRC1 | sp\|Q96M27\|PRRC1_HUMAN Protein PRRC1 OS=Homo sapiens GN=PRRC1 PE=1 SV=1 | 165,451 | 13,48 | 46701 | 3 | 24,39 | 24,20 | 23,44 | 24,10 | 24,04 | 23,54 | 0,922 | 7,49E-01 | 0 | -0,1166662 | 0,12556399 |
| CAZA1_HUM | CAPZA1 | sp\|P52907\|CAZA1_HUMAN F-actin-capping protein subunit alpha-1 OS=Homo sapiens G | 649,477 | 59,09 | 32923 | 12 | 26,57 | 26,52 | 26,41 | 26,31 | 26,66 | 26,18 | 0,922 | 4,83E-01 | 0 | -0,1166668 | 0,316367 |
| MAP2_HUM | METAP2 | sp\|P50579\|MAP2_HUMAN Methionine aminopeptidase 2 OS=Homo sapiens GN=META | 11,509 | 1,67 | 52892 | 1 | 20,97 | 20,64 | 20,79 | 20,80 | 20,46 | 20,79 | 0,922 | 4,72E-01 | 0 | -0,1166668 | 0,32649672 |
| ITA3_HUMA | ITGA3 | sp\|P26006\|ITA3_HUMAN Integrin alpha-3 OS=Homo sapiens GN=ITGA3 PE=1 SV=5 | 407,681 | 7,52 | 116612 | 12 | 27,89 | 27,50 | 26,62 | 27,66 | 27,58 | 26,42 | 0,922 | 8,42E-01 | 0 | -0,1166668 | 0,07460799 |
| SEPT2_HUM | SEPT2 | sp\|Q15019\|SEPT2_HUMAN Septin-2 OS=Homo sapiens GN=SEPT2 PE=1 SV=1 | 631,453 | 49,86 | 41487 | 14 | 26,85 | 26,85 | 26,74 | 26,62 | 26,80 | 26,67 | 0,922 | 1,47E-01 | 0 | -0,1166668 | 0,83264961 |
| VPP3_HUMA | TCIRG1 | sp\|Q13488\|VPP3_HUMAN V-type proton ATPase 116 kDa subunit a isoform 3 OS=Homo | 131,256 | 3,86 | 92968 | 2 | 22,43 | 22,35 | 20,21 | 21,73 | 22,01 | 20,90 | 0,922 | 8,91E-01 | 0 | -0,1168594 | 0,05018281 |
| ILF3_HUMA | ILF3 | sp\|Q12906\|ILF3_HUMAN Interleukin enhancer-binding factor 3 OS=Homo sapiens GN= | 789,013 | 28,41 | 95338 | 23 | 27,64 | 27,45 | 27,30 | 27,57 | 27,37 | 27,09 | 0,920 | 5,20E-01 | 0 | -0,1199996 | 0,28377698 |
| FLNA_HUMA | FLNA | sp\|P21333\|FLNA_HUMAN Filamin-A OS=Homo sapiens GN=FLNA PE=1 SV=4 | 5626,116 | 56,4 | 280739 | 130 | 30,71 | 30,55 | 30,25 | 30,68 | 30,47 | 30,00 | 0,920 | 6,46E-01 | 0 | -0,1199996 | 0,18972342 |
| ANFY1_HUM | ANKFY1 | sp\|Q9P2R3\|ANFY1_HUMAN Rabankyrin-5 OS=Homo sapiens GN=ANKFY1 PE=1 SV=2 | 139,621 | 7,7 | 128399 | 8 | 24,39 | 23,94 | 23,50 | 23,76 | 23,96 | 23,75 | 0,920 | 6,75E-01 | 0 | -0,1200002 | 0,17061237 |
| CNPY2_HUM | CNPY2 | sp\|Q9Y2B0\|CNPY2_HUMAN Protein canopy homolog 2 OS=Homo sapiens GN=CNPY2 PE | 157,984 | 33,52 | 20652 | 5 | 23,42 | 23,56 | 23,28 | 23,74 | 23,63 | 22,53 | 0,920 | 7,76E-01 | 0 | -0,1200002 | 0,10999821 |
| TSP1_HUMA | THBS1 | sp\|P07996\|TSP1_HUMAN Thrombospondin-1 OS=Homo sapiens GN=THBS1 PE=1 SV=2 | 715,983 | 21,79 | 129383 | 21 | 25,78 | 24,71 | 27,18 | 26,13 | 24,84 | 26,34 | 0,920 | 8,95E-01 | 0 | -0,1200002 | 0,04808557 |
| SATT_HUMA | SLC1A4 | sp\|P43007\|SATT_HUMAN Neutral amino acid transporter A OS=Homo sapiens GN=SLC1 | 146,88 | 6,39 | 55723 | 2 | 26,79 | 26,95 | 25,85 | 26,54 | 27,01 | 25,68 | 0,920 | 8,28E-01 | 0 | -0,1200002 | 0,08170972 |
| SBDS_HUMA | SBDS | sp\|Q9Y3A5\|SBDS_HUMAN Ribosome maturation protein SBDS OS=Homo sapiens GN=S | 47,854 | 10,4 | 28764 | 3 | 24,35 | 24,30 | 23,25 | 24,06 | 23,99 | 23,48 | 0,918 | 7,75E-01 | 0 | -0,1233336 | 0,1109316 |
| 1433E_HUM | YWHAE | sp\|P62258\|1433E_HUMAN 14-3-3 protein epsilon OS=Homo sapiens GN=YWHAE PE=1 | 1529,164 | 77,65 | 29174 | 36 | 29,69 | 29,66 | 29,08 | 29,46 | 29,51 | 29,09 | 0,918 | 6,33E-01 | 0 | -0,1233336 | 0,1988964 |
| CHM2A_HUN | CHMP2A | sp\|O43633\|CHM2A_HUMAN Charged multivesicular body protein 2a OS=Homo sapiens | 208,046 | 22,07 | 25104 | 12 | 28,76 | 28,09 | 26,74 | 28,66 | 28,09 | 26,47 | 0,918 | 8,96E-01 | 0 | -0,1233336 | 0,0477416 |
| DYN3_HUM | DNM3 | sp\|Q9UQ16\|DYN3_HUMAN Dynamin-3 OS=Homo sapiens GN=DNM3 PE=1 SV=4 | 138,965 | 10,82 | 97746 | 9 | 23,00 | 22,77 | 22,57 | 22,77 | 22,64 | 22,56 | 0,918 | 4,23E-01 | 0 | -0,1233336 | 0,37317514 |
| MPRI_HUMA | IGF2R | sp\|P11717\|MPRI_HUMAN Cation-independent mannose-6-phosphate receptor OS=Hor | 10,7 | 0,36 | 274375 | 1 | 21,93 | 21,30 | 21,24 | 22,05 | 21,48 | 20,57 | 0,918 | 8,11E-01 | 0 | -0,1233336 | 0,09071161 |
| RM16_HUM | MRPL16 | sp\|Q9NX20\|RM16_HUMAN 39S ribosomal protein L16, mitochondrial OS=Homo sapiens | 11,703 | 3,98 | 28449 | 1 | 18,94 | 18,16 | 21,07 | 19,30 | 18,29 | 20,20 | 0,917 | 9,09E-01 | 0 | -0,1253217 | 0,04138374 |
| RLA1_HUMA | RPLP1 | sp\|P05386\|RLA1_HUMAN 60S acidic ribosomal protein P1 OS=Homo sapiens GN=RPLP1 | 249,247 | 57,02 | 11514 | 4 | 25,39 | 26,02 | 26,17 | 25,19 | 25,72 | 26,29 | 0,916 | 7,66E-01 | 0 | -0,1266664 | 0,11582148 |
| AGM1_HUM | PGM3 | sp\|O95394\|AGM1_HUMAN Phosphoacetylglucosamine mutase OS=Homo sapiens GN= | 86,134 | 11,07 | 59852 | 5 | 23,47 | 23,91 | 21,90 | 23,32 | 23,60 | 21,98 | 0,916 | 8,80E-01 | 0 | -0,1266664 | 0,05542231 |
| IF2B2_HUM | IGF2BP2 | sp\|Q9Y6M1\|IF2B2_HUMAN Insulin-like growth factor 2 mRNA-binding protein 2 OS=Ho | 120,267 | 8,85 | 66121 | 4 | 21,10 | 21,68 | 22,78 | 21,08 | 21,18 | 22,92 | 0,916 | 8,78E-01 | 0 | -0,126667 | 0,05651093 |
| FAF1_HUMA | FAF1 | sp\|Q9UNN5\|FAF1_HUMAN FAS-associated factor 1 OS=Homo sapiens GN=FAF1 PE=1 S | 105,489 | 5,38 | 73954 | 3 | 22,59 | 22,41 | 21,93 | 22,50 | 22,32 | 21,73 | 0,916 | 6,99E-01 | 0 | -0,126667 | 0,15551694 |
| CPNS1_HUM | CAPNS1 | sp\|P04632\|CPNS1_HUMAN Calpain small subunit 1 OS=Homo sapiens GN=CAPNS1 PE=1 | 613,072 | 40,67 | 28316 | 13 | 26,01 | 26,31 | 26,15 | 25,96 | 26,33 | 25,79 | 0,914 | 5,13E-01 | 0 | -0,1299998 | 0,28962378 |
| GRK6_HUMA | GRK6 | sp\|P43250\|GRK6_HUMAN G protein-coupled receptor kinase 6 OS=Homo sapiens GN= | 152,11 | 10,07 | 65991 | 4 | 23,15 | 23,08 | 21,97 | 23,13 | 22,78 | 21,90 | 0,914 | 8,18E-01 | 0 | -0,1299998 | 0,08724173 |
| TSN9_HUMA | TSPAN9 | sp\|O75954\|TSN9_HUMAN Tetraspanin-9 OS=Homo sapiens GN=TSPAN9 PE=1 SV=1 | 156,765 | 13,81 | 26779 | 3 | 25,93 | 25,49 | 24,24 | 25,82 | 25,67 | 23,78 | 0,914 | 8,83E-01 | 0 | -0,1299998 | 0,05405003 |
| SPAG1_HUM | SPAG1 | sp\|Q07617\|SPAG1_HUMAN Sperm-associated antigen 1 OS=Homo sapiens GN=SPAG1 | 66,755 | 3,02 | 103639 | 3 | 25,23 | 24,25 | 24,01 | 24,14 | 25,16 | 23,80 | 0,914 | 8,26E-01 | 0 | -0,1300004 | 0,08312287 |
| KPRA_HUMA | PRPSAP1 | sp\|Q14558\|KPRA_HUMAN Phosphoribosyl pyrophosphate synthase-associated protein | 138,754 | 17,13 | 39394 | 4 | 22,01 | 22,05 | 23,65 | 21,70 | 21,98 | 23,63 | 0,912 | 8,77E-01 | 0 | -0,1333332 | 0,05696755 |
| FA9_HUMAN | F9 | sp\|P00740\|FA9_HUMAN Coagulation factor IX OS=Homo sapiens GN=F9 PE=1 SV=2 | 15,828 | 2,6 | 51778 | 1 | 21,48 | 20,79 | 21,77 | 21,67 | 20,61 | 21,36 | 0,912 | 7,71E-01 | 0 | -0,1333332 | 0,11286407 |
| MYH14_HUN | MYH14 | sp\|Q7Z406\|MYH14_HUMAN Myosin-14 OS=Homo sapiens GN=MYH14 PE=1 SV=2 | 420,682 | 5,06 | 227871 | 11 | 24,97 | 24,96 | 25,67 | 24,88 | 25,10 | 25,22 | 0,912 | 6,29E-01 | 0 | -0,1333332 | 0,2013551 |
| GAPR1_HUM | GLIPR2 | sp\|Q9H4G4\|GAPR1_HUMAN Golgi-associated plant pathogenesis-related protein 1 OS= | 418,602 | 45,45 | 17218 | 6 | 28,45 | 28,01 | 26,56 | 28,32 | 28,04 | 26,26 | 0,912 | 8,84E-01 | 0 | -0,1333332 | 0,053304 |
| KI13B_HUM | KIF13B | sp\|Q9NQT8\|KI13B_HUMAN Kinesin-like protein KIF13B OS=Homo sapiens GN=KIF13B P | 50,946 | 1,53 | 202789 | 2 | 17,50 | 18,66 | 19,09 | 17,96 | 18,55 | 18,34 | 0,912 | 8,05E-01 | 0 | -0,1333338 | 0,09425715 |
| AT1B1_HUM | ATP1B1 | sp\|P05026\|AT1B1_HUMAN Sodium/potassium-transporting ATPase subunit beta-1 OS= | 170,072 | 11,55 | 35061 | 6 | 27,43 | 27,67 | 25,10 | 27,35 | 27,40 | 25,05 | 0,912 | 9,12E-01 | 0 | -0,1333338 | 0,04019045 |
| RRAS2_HUM | RRAS2 | sp\|P62070\|RRAS2_HUMAN Ras-related protein R-Ras2 OS=Homo sapiens GN=RRAS2 PE | 959,111 | 75 | 23400 | 19 | 28,70 | 28,43 | 27,42 | 28,57 | 28,42 | 27,16 | 0,912 | 8,33E-01 | 0 | -0,1333338 | 0,07929393 |
| LMO7_HUM | LMO7 | sp\|Q8WWI1\|LMO7_HUMAN LIM domain only protein 7 OS=Homo sapiens GN=LMO7 P | 180,047 | 2,97 | 192696 | 3 | 21,68 | 21,61 | 22,18 | 21,06 | 21,59 | 22,42 | 0,912 | 7,74E-01 | 0 | -0,1333338 | 0,11109802 |
| STAU1_HUM | STAU1 | sp\|O95793\|STAU1_HUMAN Double-stranded RNA-binding protein Staufen homolog 1 C | 29,312 | 3,99 | 63182 | 2 | 21,50 | 21,15 | 21,83 | 21,52 | 22,12 | 20,43 | 0,910 | 8,10E-01 | 0 | -0,136666 | 0,09151477 |
| ATX2L_HUM | ATXN2L | sp\|Q8WWM7\|ATX2L_HUMAN Ataxin-2-like protein OS=Homo sapiens GN=ATXN2L PE= | 388,053 | 15,26 | 113374 | 13 | 24,88 | 24,79 | 24,78 | 24,93 | 24,82 | 24,29 | 0,910 | 5,32E-01 | 0 | -0,1366666 | 0,27397422 |

| | | | | | | | | | | | | | | | | | |
|---|---|---|---|---|---|---|---|---|---|---|---|---|---|---|---|---|---|
| CRLF3_HUM | CRLF3 | sp\|Q8IUI8\|CRLF3_HUMAN Cytokine receptor-like factor 3 OS=Homo sapiens GN=CRLF3 | 55,604 | 7,69 | 49766 | 2 | 21,08 | 21,51 | 21,10 | 20,81 | 21,16 | 21,31 | **0,910** | 5,39E-01 | **0** | -0,1366673 | 0,26808093 |
| PHP14_HUM | PHPT1 | sp\|Q9NRX4\|PHP14_HUMAN 14 kDa phosphohistidine phosphatase OS=Homo sapiens G | 52,25 | 8,8 | 13833 | 1 | 20,76 | 20,52 | 18,70 | 20,62 | 20,75 | 18,19 | **0,908** | 9,01E-01 | **0** | -0,1397324 | 0,04519392 |
| NXN_HUMA | NXN | sp\|Q6DKJ4\|NXN_HUMAN Nucleoredoxin OS=Homo sapiens GN=NXN PE=1 SV=2 | 337,75 | 30,11 | 48392 | 10 | 25,07 | 25,08 | 25,56 | 24,81 | 24,86 | 25,62 | **0,908** | 6,73E-01 | **0** | -0,1399994 | 0,17201966 |
| CNNM4_HU | CNNM4 | sp\|Q6P4Q7\|CNNM4_HUMAN Metal transporter CNNM4 OS=Homo sapiens GN=CNNM4 | 27,002 | 1,42 | 86607 | 1 | 21,74 | 22,23 | 20,66 | 21,69 | 22,03 | 20,49 | **0,908** | 8,42E-01 | **0** | -0,1399994 | 0,07470557 |
| ANO6_HUM | ANO6 | sp\|Q4KMQ2\|ANO6_HUMAN Anoctamin-6 OS=Homo sapiens GN=ANO6 PE=1 SV=2 | 83,251 | 5,82 | 106165 | 6 | 25,47 | 25,56 | 24,73 | 25,39 | 25,58 | 24,37 | **0,908** | 7,75E-01 | **0** | -0,1399994 | 0,11049133 |
| VPS4B_HUM | VPS4B | sp\|O75351\|VPS4B_HUMAN Vacuolar protein sorting-associated protein 4B OS=Homo sa | 582,222 | 36,04 | 49302 | 15 | 27,30 | 27,11 | 25,73 | 27,29 | 27,01 | 25,42 | **0,908** | 8,64E-01 | **0** | -0,1399994 | 0,06372073 |
| SAP_HUMAN | PSAP | sp\|P07602\|SAP_HUMAN Prosaposin OS=Homo sapiens GN=PSAP PE=1 SV=2 | 41,619 | 2,86 | 58113 | 1 | 17,49 | 18,44 | 19,87 | 17,26 | 18,08 | 20,04 | **0,908** | 9,03E-01 | **0** | -0,14 | 0,04441231 |
| MOT2_HUM | SLC16A7 | sp\|O60669\|MOT2_HUMAN Monocarboxylate transporter 2 OS=Homo sapiens GN=SLC1 | 26,919 | 1,67 | 52200 | 1 | 21,80 | 21,24 | 21,84 | 21,23 | 21,76 | 21,47 | **0,908** | 6,01E-01 | **0** | -0,14 | 0,22106492 |
| STK10_HUM | STK10 | sp\|O94804\|STK10_HUMAN Serine/threonine-protein kinase 10 OS=Homo sapiens GN=S | 193,391 | 9,3 | 112135 | 7 | 23,34 | 22,46 | 22,03 | 23,39 | 21,86 | 22,16 | **0,908** | 8,29E-01 | **0** | -0,14 | 0,0815786 |
| RAB13_HUM | RAB13 | sp\|P51153\|RAB13_HUMAN Ras-related protein Rab-13 OS=Homo sapiens GN=RAB13 PE | 209,386 | 31,53 | 22774 | 6 | 24,92 | 24,45 | 24,73 | 24,74 | 24,47 | 24,47 | **0,908** | 4,40E-01 | **0** | -0,1400007 | 0,3564288 |
| NECP1_HUM | NECAP1 | sp\|Q8NC96\|NECP1_HUMAN Adaptin ear-binding coat-associated protein 1 OS=Homo sa | 21,585 | 3,27 | 29737 | 1 | 21,28 | 21,70 | 21,66 | 21,51 | 21,49 | 21,22 | **0,908** | 4,40E-01 | **0** | -0,1400007 | 0,35703503 |
| GSK3A_HUM | GSK3A | sp\|P49840\|GSK3A_HUMAN Glycogen synthase kinase-3 alpha OS=Homo sapiens GN=GS | 50,791 | 4,76 | 50981 | 2 | 20,55 | 20,61 | 21,36 | 20,39 | 20,66 | 21,05 | **0,908** | 6,87E-01 | **0** | -0,1400007 | 0,16277596 |
| SF3A3_HUM | SF3A3 | sp\|Q12874\|SF3A3_HUMAN Splicing factor 3A subunit 3 OS=Homo sapiens GN=SF3A3 PE | 278,003 | 26,35 | 58849 | 9 | 24,39 | 24,38 | 24,72 | 24,42 | 24,24 | 24,40 | **0,905** | 3,17E-01 | **0** | -0,1433328 | 0,49926855 |
| DP13A_HUM | APPL1 | sp\|Q9UKG1\|DP13A_HUMAN DCC-interacting protein 13-alpha OS=Homo sapiens GN=AI | 202,558 | 12,83 | 79663 | 7 | 24,99 | 24,90 | 22,14 | 24,24 | 24,46 | 22,90 | **0,905** | 8,98E-01 | **0** | -0,1433334 | 0,04649526 |
| FUMH_HUM | FH | sp\|P07954\|FUMH_HUMAN Fumarate hydratase, mitochondrial OS=Homo sapiens GN=F | 546,202 | 39,02 | 54637 | 12 | 25,53 | 25,80 | 26,61 | 25,82 | 25,75 | 25,94 | **0,905** | 6,86E-01 | **0** | -0,1433334 | 0,16383821 |
| PELP1_HUM | PELP1 | sp\|Q8IZL8\|PELP1_HUMAN Proline-, glutamic acid- and leucine-rich protein 1 OS=Homo | 56,337 | 1,24 | 119700 | 1 | 17,32 | 19,16 | 19,68 | 17,06 | 19,05 | 19,62 | **0,905** | 8,99E-01 | **0** | -0,1433334 | 0,04644863 |
| CALU_HUMA | CALU | sp\|O43852\|CALU_HUMAN Calumenin OS=Homo sapiens GN=CALU PE=1 SV=2 | 348,438 | 38,73 | 37107 | 11 | 23,91 | 24,25 | 26,02 | 24,53 | 24,21 | 25,01 | **0,905** | 8,46E-01 | **0** | -0,1433334 | 0,07237485 |
| RL36_HUMA | RPL36 | sp\|Q9Y3U8\|RL36_HUMAN 60S ribosomal protein L36 OS=Homo sapiens GN=RPL36 PE= | 19,138 | 9,52 | 12254 | 1 | 22,65 | 23,14 | 21,80 | 22,83 | 22,11 | 22,21 | **0,903** | 7,62E-01 | **0** | -0,1466662 | 0,11825498 |
| ENOB_HUM | ENO3 | sp\|P13929\|ENOB_HUMAN Beta-enolase OS=Homo sapiens GN=ENO3 PE=1 SV=5 | 477,425 | 19,35 | 46987 | 6 | 27,76 | 27,97 | 26,21 | 27,61 | 27,71 | 26,18 | **0,903** | 8,53E-01 | **0** | -0,1466662 | 0,0689673 |
| HEBP1_HUM | HEBP1 | sp\|Q9NRV9\|HEBP1_HUMAN Heme-binding protein 1 OS=Homo sapiens GN=HEBP1 PE= | 209,068 | 22,75 | 21097 | 3 | 25,69 | 25,78 | 24,96 | 25,52 | 25,63 | 24,84 | **0,903** | 7,03E-01 | **0** | -0,1466668 | 0,15284014 |
| CD276_HUM | CD276 | sp\|Q5ZPR3\|CD276_HUMAN CD276 antigen OS=Homo sapiens GN=CD276 PE=1 SV=1 | 134,003 | 8,24 | 57235 | 2 | 24,89 | 25,05 | 24,70 | 24,80 | 24,80 | 24,60 | **0,903** | 2,93E-01 | **0** | -0,1466668 | 0,53360088 |
| DNJC8_HUM | DNAJC8 | sp\|O75937\|DNJC8_HUMAN DnaJ homolog subfamily C member 8 OS=Homo sapiens GN | 21,022 | 3,16 | 29842 | 1 | 20,94 | 21,14 | 19,26 | 21,19 | 20,95 | 18,75 | **0,902** | 8,87E-01 | **0** | -0,1482557 | 0,05206879 |
| TLK1_HUMA | TLK1 | sp\|Q9UKI8\|TLK1_HUMAN Serine/threonine-protein kinase tousled-like 1 OS=Homo sap | 38,589 | 2,22 | 86700 | 2 | 18,48 | 18,97 | 17,12 | 18,95 | 18,86 | 16,31 | **0,902** | 8,92E-01 | **0** | -0,1485062 | 0,04957526 |
| L2GL1_HUM | LLGL1 | sp\|Q15334\|L2GL1_HUMAN Lethal(2) giant larvae protein homolog 1 OS=Homo sapiens | 439,082 | 13,82 | 115418 | 11 | 25,97 | 25,92 | 24,45 | 25,88 | 25,65 | 24,36 | **0,901** | 8,38E-01 | **0** | -0,1500003 | 0,07680559 |
| RAB5B_HUM | RAB5B | sp\|P61020\|RAB5B_HUMAN Ras-related protein Rab-5B OS=Homo sapiens GN=RAB5B PE | 457,987 | 53,49 | 23707 | 10 | 25,37 | 25,19 | 25,59 | 25,25 | 25,11 | 25,34 | **0,901** | 3,24E-01 | **0** | -0,1500003 | 0,48884959 |
| MYH9_HUM | MYH9 | sp\|P35579\|MYH9_HUMAN Myosin-9 OS=Homo sapiens GN=MYH9 PE=1 SV=4 | 6029,003 | 51,38 | 226532 | 132 | 29,94 | 30,28 | 30,65 | 30,13 | 30,23 | 30,06 | **0,901** | 5,16E-01 | **0** | -0,1500009 | 0,28721776 |
| DNJA2_HUM | DNAJA2 | sp\|O60884\|DNJA2_HUMAN DnaJ homolog subfamily A member 2 OS=Homo sapiens GN | 1101,692 | 63,11 | 45746 | 27 | 29,24 | 28,80 | 28,25 | 29,20 | 28,59 | 28,04 | **0,899** | 7,45E-01 | **0** | -0,1533324 | 0,12758209 |
| IMDH1_HUM | IMPDH1 | sp\|P20839\|IMDH1_HUMAN Inosine-5'-monophosphate dehydrogenase 1 OS=Homo sap | 44,003 | 6,61 | 55406 | 3 | 21,64 | 21,64 | 24,39 | 21,27 | 21,72 | 24,22 | **0,899** | 9,12E-01 | **0** | -0,153333 | 0,04019969 |
| MARE2_HUM | MAPRE2 | sp\|Q15555\|MARE2_HUMAN Microtubule-associated protein RP/EB family member 2 OS | 91,35 | 14,68 | 37031 | 3 | 21,85 | 21,60 | 23,06 | 21,35 | 21,78 | 22,92 | **0,899** | 8,25E-01 | **0** | -0,153333 | 0,08347868 |
| F120A_HUM | FAM120A | sp\|Q9NZB2\|F120A_HUMAN Constitutive coactivator of PPAR-gamma-like protein 1 OS=I | 135,443 | 5,37 | 121888 | 5 | 23,09 | 23,25 | 23,54 | 22,85 | 22,96 | 23,61 | **0,899** | 6,02E-01 | **0** | -0,1533337 | 0,2203438 |
| 1433B_HUM | YWHAB | sp\|P31946\|1433B_HUMAN 14-3-3 protein beta/alpha OS=Homo sapiens GN=YWHAB PE | 1217,941 | 75,61 | 28082 | 29 | 27,27 | 27,26 | 26,45 | 27,25 | 26,84 | 26,43 | **0,899** | 6,92E-01 | **0** | -0,1533337 | 0,15967493 |
| AIDA_HUMA | AIDA | sp\|Q96BJ3\|AIDA_HUMAN Axin interactor, dorsalization-associated protein OS=Homo sa | 161,663 | 26,47 | 35023 | 6 | 24,52 | 24,34 | 23,54 | 24,49 | 24,29 | 23,16 | **0,899** | 7,79E-01 | **0** | -0,1533337 | 0,10818908 |
| FAT4_HUMA | FAT4 | sp\|Q6V0I7\|FAT4_HUMAN Protocadherin Fat 4 OS=Homo sapiens GN=FAT4 PE=1 SV=2 | 83,831 | 0,88 | 542687 | 3 | 22,70 | 22,11 | 21,48 | 22,26 | 20,81 | 22,76 | **0,899** | 8,33E-01 | **0** | -0,1533337 | 0,07920483 |
| CHM4A_HU | CHMP4A | sp\|Q9BY43\|CHM4A_HUMAN Charged multivesicular body protein 4a OS=Homo sapiens | 172,953 | 18,47 | 25098 | 5 | 27,38 | 26,57 | 26,22 | 27,47 | 26,45 | 25,78 | **0,897** | 8,07E-01 | **0** | -0,1566658 | 0,09326131 |
| SAHH3_HUM | AHCYL2 | sp\|Q96HN2\|SAHH3_HUMAN Adenosylhomocysteinase 3 OS=Homo sapiens GN=AHCYL2 | 286,078 | 21,28 | 66721 | 14 | 22,75 | 21,96 | 22,09 | 22,36 | 22,21 | 21,76 | **0,897** | 6,33E-01 | **0** | -0,1566664 | 0,19839422 |
| HBA_HUMA | HBA1 | sp\|P69905\|HBA_HUMAN Hemoglobin subunit alpha OS=Homo sapiens GN=HBA1 PE=1 S | 247,546 | 34,51 | 15258 | 6 | 30,64 | 29,74 | 31,43 | 30,60 | 29,56 | 31,18 | **0,897** | 8,29E-01 | **0** | -0,1566664 | 0,0813485 |
| VAT1_HUMA | VAT1 | sp\|Q99536\|VAT1_HUMAN Synaptic vesicle membrane protein VAT-1 homolog OS=Hom | 601,585 | 30,03 | 41920 | 11 | 27,69 | 27,89 | 26,82 | 27,56 | 27,65 | 26,72 | **0,897** | 7,41E-01 | **0** | -0,1566671 | 0,13016257 |
| TWF1_HUM | TWF1 | sp\|Q12792\|TWF1_HUMAN Twinfilin-1 OS=Homo sapiens GN=TWF1 PE=1 SV=3 | 244,306 | 27,14 | 40283 | 7 | 25,18 | 25,24 | 23,78 | 24,93 | 24,80 | 24,00 | **0,897** | 7,93E-01 | **0** | -0,1566671 | 0,10070689 |
| IFRD2_HUM | IFRD2 | sp\|Q12894\|IFRD2_HUMAN Interferon-related developmental regulator 2 OS=Homo sap | 13,128 | 1,78 | 54814 | 1 | 20,51 | 20,51 | 20,83 | 20,43 | 19,32 | 21,62 | **0,895** | 8,24E-01 | **0** | -0,1599998 | 0,08425186 |
| PTPRA_HUM | PTPRA | sp\|P18433\|PTPRA_HUMAN Receptor-type tyrosine-protein phosphatase alpha OS=Hom | 390,625 | 24,94 | 90600 | 14 | 25,55 | 25,35 | 26,08 | 25,56 | 25,53 | 25,41 | **0,895** | 5,12E-01 | **0** | -0,1599998 | 0,29076568 |
| ITB6_HUMA | ITGB6 | sp\|P18564\|ITB6_HUMAN Integrin beta-6 OS=Homo sapiens GN=ITGB6 PE=1 SV=2 | 48,319 | 1,14 | 85936 | 1 | 27,83 | 27,60 | 26,73 | 27,87 | 27,47 | 26,34 | **0,895** | 7,92E-01 | **0** | -0,1599998 | 0,10128201 |
| GBP7_HUMA | GBP7 | sp\|Q8N8V2\|GBP7_HUMAN Guanylate-binding protein 7 OS=Homo sapiens GN=GBP7 PE | 18,556 | 1,57 | 72513 | 1 | 21,69 | 21,66 | 21,11 | 20,87 | 21,00 | 22,11 | **0,895** | 7,32E-01 | **0** | -0,1599998 | 0,13525045 |
| MCES_HUM | RNMT | sp\|O43148\|MCES_HUMAN mRNA cap guanine-N7 methyltransferase OS=Homo sapiens | 42,435 | 4,2 | 54844 | 2 | 22,55 | 22,22 | 21,93 | 22,00 | 20,56 | 23,66 | **0,895** | 8,69E-01 | **0** | -0,1599998 | 0,06075409 |
| LRC49_HUM | LRRC49 | sp\|Q8IUZ0\|LRC49_HUMAN Leucine-rich repeat-containing protein 49 OS=Homo sapiens | 15,521 | 1,31 | 78894 | 1 | 23,43 | 23,70 | 22,54 | 23,61 | 23,49 | 22,09 | **0,895** | 8,03E-01 | **0** | -0,1600005 | 0,09521138 |
| 1433F_HUM | YWHAH | sp\|Q04917\|1433F_HUMAN 14-3-3 protein eta OS=Homo sapiens GN=YWHAH PE=1 SV= | 654,697 | 53,25 | 28219 | 18 | 28,07 | 27,99 | 27,04 | 27,97 | 27,71 | 26,94 | **0,895** | 7,42E-01 | **0** | -0,1600005 | 0,1297836 |
| CAVN2_HUM | CAVIN2 | sp\|O95810\|CAVN2_HUMAN Caveolae-associated protein 2 OS=Homo sapiens GN=CAVI | 96,471 | 6,59 | 47173 | 3 | 24,12 | 24,07 | 24,51 | 24,20 | 23,80 | 24,22 | **0,895** | 4,58E-01 | **0** | -0,1600005 | 0,33897132 |
| ARF4_HUMA | ARF4 | sp\|P18085\|ARF4_HUMAN ADP-ribosylation factor 4 OS=Homo sapiens GN=ARF4 PE=1 S | 693,818 | 68,33 | 20511 | 15 | 26,51 | 26,53 | 26,49 | 26,16 | 26,55 | 26,34 | **0,895** | 2,31E-01 | **0** | -0,1600005 | 0,63695489 |
| NCDN_HUM | NCDN | sp\|Q9UBB6\|NCDN_HUMAN Neurochondrin OS=Homo sapiens GN=NCDN PE=1 SV=1 | 139,991 | 8,78 | 78864 | 4 | 21,96 | 21,93 | 21,72 | 21,77 | 21,58 | 21,77 | **0,893** | 1,73E-01 | **0** | -0,1633326 | 0,76252932 |
| TRIPC_HUM | TRIP12 | sp\|Q14669\|TRIPC_HUMAN E3 ubiquitin-protein ligase TRIP12 OS=Homo sapiens GN=TR | 47,291 | 0,6 | 220434 | 1 | 19,96 | 19,96 | 20,73 | 19,47 | 19,82 | 20,87 | **0,893** | 7,57E-01 | **0** | -0,1633326 | 0,1209402 |
| GNAQ_HUM | GNAQ | sp\|P50148\|GNAQ_HUMAN Guanine nucleotide-binding protein G(q) subunit alpha OS=I | 747,73 | 58,77 | 42142 | 19 | 26,16 | 26,29 | 26,10 | 26,23 | 26,15 | 25,68 | **0,893** | 4,17E-01 | **0** | -0,1633339 | 0,38019933 |
| DHB4_HUM | HSD17B4 | sp\|P51659\|DHB4_HUMAN Peroxisomal multifunctional enzyme type 2 OS=Homo sapien | 22,953 | 1,49 | 79686 | 1 | 20,86 | 20,37 | 21,57 | 20,76 | 20,31 | 21,24 | **0,893** | 7,29E-01 | **0** | -0,1633339 | 0,13715284 |
| CASP7_HUM | CASP7 | sp\|P55210\|CASP7_HUMAN Caspase-7 OS=Homo sapiens GN=CASP7 PE=1 SV=1 | 40,119 | 3,96 | 34277 | 1 | 21,28 | 21,22 | 18,81 | 21,73 | 20,63 | 18,45 | **0,893** | 9,03E-01 | **0** | -0,1640015 | 0,04441607 |
| IFT56_HUM | TTC26 | sp\|A0AVF1\|IFT56_HUMAN Intraflagellar transport protein 56 OS=Homo sapiens GN=TT( | 219,502 | 18,59 | 64178 | 9 | 17,90 | 20,35 | 25,58 | 21,73 | 17,35 | 24,26 | **0,891** | 9,59E-01 | **0** | -0,1662731 | 0,01821646 |
| ACSL4_HUM | ACSL4 | sp\|O60488\|ACSL4_HUMAN Long-chain-fatty-acid--CoA ligase 4 OS=Homo sapiens GN=A | 957,88 | 42,62 | 79188 | 22 | 27,22 | 27,21 | 25,81 | 26,93 | 27,00 | 25,81 | **0,891** | 7,97E-01 | **0** | -0,166666 | 0,09847854 |
| MTOR_HUM | MTOR | sp\|P42345\|MTOR_HUMAN Serine/threonine-protein kinase mTOR OS=Homo sapiens G | 93,122 | 2 | 288892 | 3 | 20,30 | 20,12 | 21,23 | 19,94 | 20,41 | 20,80 | **0,891** | 7,15E-01 | **0** | -0,1666667 | 0,14596223 |
| SRP09_HUM | SRP9 | sp\|P49458\|SRP09_HUMAN Signal recognition particle 9 kDa protein OS=Homo sapiens G | 93,097 | 30,23 | 10112 | 3 | 26,60 | 26,74 | 24,61 | 26,66 | 26,59 | 24,20 | **0,891** | 8,83E-01 | **0** | -0,1666667 | 0,05411215 |
| F234A_HUM | FAM234A | sp\|Q9H0X4\|F234A_HUMAN Protein FAM234A OS=Homo sapiens GN=FAM234A PE=1 SV | 25,006 | 2,17 | 59660 | 1 | 22,83 | 22,78 | 20,93 | 22,54 | 22,53 | 20,97 | **0,891** | 8,48E-01 | **0** | -0,1666667 | 0,07170296 |
| RAB21_HUM | RAB21 | sp\|Q9UL25\|RAB21_HUMAN Ras-related protein Rab-21 OS=Homo sapiens GN=RAB21 PI | 406,889 | 44,44 | 24348 | 9 | 26,11 | 26,04 | 25,00 | 25,56 | 25,99 | 25,10 | **0,891** | 7,25E-01 | **0** | -0,1666673 | 0,13970728 |
| RL18_HUMA | RPL18 | sp\|Q07020\|RL18_HUMAN 60S ribosomal protein L18 OS=Homo sapiens GN=RPL18 PE=1 | 177,783 | 26,06 | 21634 | 5 | 25,20 | 25,66 | 24,86 | 24,91 | 24,49 | 25,82 | **0,891** | 7,33E-01 | **0** | -0,1666673 | 0,13479468 |
| S10A4_HUM | S100A4 | sp\|P26447\|S10A4_HUMAN Protein S100-A4 OS=Homo sapiens GN=S100A4 PE=1 SV=1 | 25,879 | 16,83 | 11729 | 2 | 24,13 | 24,23 | 20,02 | 24,12 | 23,91 | 19,84 | **0,889** | 9,36E-01 | **0** | -0,1689809 | 0,02888663 |
| RRAGA_HUM | RRAGA | sp\|Q7L523\|RRAGA_HUMAN Ras-related GTP-binding protein A OS=Homo sapiens GN=R | 128,903 | 12,14 | 36566 | 3 | 23,21 | 22,79 | 22,57 | 22,59 | 22,61 | 22,86 | **0,889** | 4,57E-01 | **0** | -0,1699994 | 0,33976473 |
| CNBP_HUMA | CNBP | sp\|P62633\|CNBP_HUMAN Cellular nucleic acid-binding protein OS=Homo sapiens GN=C | 228,594 | 30,51 | 19463 | 4 | 24,59 | 24,72 | 24,01 | 24,52 | 24,19 | 24,10 | **0,889** | 5,38E-01 | **0** | -0,1699994 | 0,26904718 |
| TM14C_HUM | TMEM14C | sp\|Q9P0S9\|TM14C_HUMAN Transmembrane protein 14C OS=Homo sapiens GN=TMEM | 120,831 | 41,07 | 11565 | 2 | 20,79 | 21,12 | 19,88 | 20,34 | 21,12 | 19,82 | **0,889** | 7,64E-01 | **0** | -0,1700001 | 0,1168108 |
| DNJA1_HUM | DNAJA1 | sp\|P31689\|DNJA1_HUMAN DnaJ homolog subfamily A member 1 OS=Homo sapiens GN | 1742,566 | 74,81 | 44868 | 32 | 29,23 | 29,13 | 28,29 | 29,15 | 28,82 | 28,17 | **0,889** | 7,03E-01 | **0** | -0,1700001 | 0,15325918 |
| CASP3_HUM | CASP3 | sp\|P42574\|CASP3_HUMAN Caspase-3 OS=Homo sapiens GN=CASP3 PE=1 SV=2 | 27,263 | 3,97 | 31608 | 1 | 21,41 | 20,50 | 20,71 | 21,48 | 19,82 | 20,81 | **0,889** | 7,75E-01 | **0** | -0,1700001 | 0,11086721 |
| STRN3_HUM | STRN3 | sp\|Q13033\|STRN3_HUMAN Striatin-3 OS=Homo sapiens GN=STRN3 PE=1 SV=3 | 84,21 | 3,89 | 87209 | 3 | 22,08 | 22,54 | 21,93 | 21,72 | 21,93 | 22,39 | **0,889** | 5,63E-01 | **0** | -0,1700007 | 0,24956151 |
| LITAF_HUM | LITAF | sp\|Q99732\|LITAF_HUMAN Lipopolysaccharide-induced tumor necrosis factor-alpha fact | 18,422 | 4,35 | 17107 | 1 | 25,45 | 25,16 | 19,16 | 25,37 | 24,65 | 19,23 | **0,887** | 9,54E-01 | **0** | -0,1725629 | 0,02037003 |
| AGO2_HUM | AGO2 | sp\|Q9UKV8\|AGO2_HUMAN Protein argonaute-2 OS=Homo sapiens GN=AGO2 PE=1 SV= | 20,056 | 1,98 | 97208 | 1 | 20,74 | 19,08 | 18,75 | 19,05 | 20,24 | 18,76 | **0,887** | 8,32E-01 | **0** | -0,172718 | 0,07966034 |
| RADI_HUMA | RDX | sp\|P35241\|RADI_HUMAN Radixin OS=Homo sapiens GN=RDX PE=1 SV=1 | 1644,023 | 61,06 | 68564 | 47 | 27,90 | 27,65 | 27,81 | 27,85 | 27,37 | 27,62 | **0,887** | 3,31E-01 | **0** | -0,1733322 | 0,4805419 |
| MOES_HUM | MSN | sp\|P26038\|MOES_HUMAN Moesin OS=Homo sapiens GN=MSN PE=1 SV=3 | 2790,976 | 71,06 | 67820 | 65 | 30,81 | 30,46 | 29,04 | 30,65 | 30,28 | 28,86 | **0,887** | 8,33E-01 | **0** | -0,1733329 | 0,07956486 |
| GNA11_HUM | GNA11 | sp\|P29992\|GNA11_HUMAN Guanine nucleotide-binding protein subunit alpha-11 OS=H | 525,37 | 46,8 | 42123 | 16 | 24,36 | 24,19 | 24,64 | 24,45 | 24,12 | 24,10 | **0,887** | 3,74E-01 | **0** | -0,1733329 | 0,42687428 |
| RAP1B_HUM | RAP1B | sp\|P61224\|RAP1B_HUMAN Ras-related protein Rap-1b OS=Homo sapiens GN=RAP1B PE | 1286,798 | 83,15 | 20825 | 22 | 27,57 | 27,41 | 26,49 | 27,54 | 27,34 | 26,07 | **0,887** | 7,76E-01 | **0** | -0,1733329 | 0,10997384 |
| VPS4A_HUM | VPS4A | sp\|Q9UN37\|VPS4A_HUMAN Vacuolar protein sorting-associated protein 4A OS=Homo s | 694,246 | 52,63 | 48898 | 22 | 28,07 | 27,63 | 26,77 | 28,02 | 27,64 | 26,29 | **0,887** | 8,03E-01 | **0** | -0,1733329 | 0,09548368 |
| DCTN1_HUM | DCTN1 | sp\|Q14203\|DCTN1_HUMAN Dynactin subunit 1 OS=Homo sapiens GN=DCTN1 PE=1 SV= | 255,713 | 11,11 | 141695 | 10 | 24,33 | 24,29 | 23,60 | 24,36 | 23,86 | 23,48 | **0,887** | 6,45E-01 | **0** | -0,1733335 | 0,19075653 |
| MYO1D_HUM | MYO1D | sp\|O94832\|MYO1D_HUMAN Unconventional myosin-Id OS=Homo sapiens GN=MYO1D | 478,33 | 20,08 | 116202 | 16 | 26,34 | 25,83 | 24,92 | 25,96 | 25,59 | 25,02 | **0,887** | 7,45E-01 | **0** | -0,1733335 | 0,12786808 |
| PIPNB_HUM | PITPNB | sp\|P48739\|PIPNB_HUMAN Phosphatidylinositol transfer protein beta isoform OS=Homo | 222,379 | 42,8 | 31540 | 10 | 25,74 | 25,04 | 24,42 | 25,03 | 25,47 | 24,18 | **0,887** | 7,63E-01 | **0** | -0,1733335 | 0,11737997 |
| GSK3B_HUM | GSK3B | sp\|P49841\|GSK3B_HUMAN Glycogen synthase kinase-3 beta OS=Homo sapiens GN=GSI | 77,353 | 14,76 | 46744 | 3 | 21,88 | 20,57 | 22,28 | 21,13 | 20,96 | 22,12 | **0,887** | 7,97E-01 | **0** | -0,1733335 | 0,09854849 |
| FLOT1_HUM | FLOT1 | sp\|O75955\|FLOT1_HUMAN Flotillin-1 OS=Homo sapiens GN=FLOT1 PE=1 SV=3 | 547,608 | 44,03 | 47355 | 15 | 26,06 | 26,08 | 26,29 | 26,01 | 26,00 | 25,90 | **0,887** | 1,01E-01 | **0** | -0,1733335 | 0,99735292 |
| FARP2_HUM | FARP2 | sp\|O94887\|FARP2_HUMAN FERM, ARHGEF and pleckstrin domain-containing protein 2 | 12,468 | 0,76 | 119888 | 1 | 20,62 | 21,10 | 18,51 | 20,20 | 20,11 | 19,40 | **0,886** | 8,45E-01 | **0** | -0,1742617 | 0,07330853 |
| TIMP3_HUM | TIMP3 | sp\|P35625\|TIMP3_HUMAN Metalloproteinase inhibitor 3 OS=Homo sapiens GN=TIMP3 | 92,551 | 17,06 | 24145 | 3 | 24,06 | 22,83 | 23,33 | 23,74 | 23,35 | 22,60 | **0,885** | 7,36E-01 | **0** | -0,1766663 | 0,13292604 |
| PA24A_HUM | PLA2G4A | sp\|P47712\|PA24A_HUMAN Cytosolic phospholipase A2 OS=Homo sapiens GN=PLA2G4A | 222,142 | 13,35 | 85239 | 8 | 24,26 | 24,77 | 24,04 | 24,24 | 24,68 | 23,62 | **0,885** | 6,63E-01 | **0** | -0,1766669 | 0,17860466 |
| S4A7_HUMA | SLC4A7 | sp\|Q9Y6M7\|S4A7_HUMAN Sodium bicarbonate cotransporter 3 OS=Homo sapiens GN= | 477,181 | 12,52 | 136044 | 13 | 24,70 | 25,18 | 23,83 | 24,94 | 24,81 | 23,43 | **0,885** | 7,91E-01 | **0** | -0,1766669 | 0,10173132 |
| COF1_HUMA | CFL1 | sp\|P23528\|COF1_HUMAN Cofilin-1 OS=Homo sapiens GN=CFL1 PE=1 SV=3 | 852,227 | 77,71 | 18502 | 15 | 30,47 | 30,48 | 29,30 | 30,31 | 30,29 | 29,11 | **0,883** | 7,63E-01 | **0** | -0,179999 | 0,11749744 |
| CALM1_HUM | CALM1 | sp\|P0DP23\|CALM1_HUMAN Calmodulin-1 OS=Homo sapiens GN=CALM1 PE=1 SV=1 | 652,395 | 85,91 | 16838 | 14 | 26,56 | 26,36 | 27,57 | 26,35 | 26,36 | 27,24 | **0,883** | 7,25E-01 | **0** | -0,1799997 | 0,13970649 |
| RAB3A_HUM | RAB3A | sp\|P20336\|RAB3A_HUMAN Ras-related protein Rab-3A OS=Homo sapiens GN=RAB3A PI | 227,243 | 30,45 | 24984 | 6 | 23,67 | 23,93 | 23,34 | 23,36 | 23,62 | 23,42 | **0,883** | 3,92E-01 | **0** | -0,1799997 | 0,40623801 |
| SNG2_HUMA | SYNGR2 | sp\|O43760\|SNG2_HUMAN Synaptogyrin-2 OS=Homo sapiens GN=SYNGR2 PE=1 SV=1 | 29,216 | 4,46 | 24810 | 1 | 22,69 | 22,93 | 22,21 | 22,61 | 22,81 | 21,87 | **0,883** | 6,39E-01 | **0** | -0,1799997 | 0,19418958 |
| EIF1_HUMA | EIF1 | sp\|P41567\|EIF1_HUMAN Eukaryotic translation initiation factor 1 OS=Homo sapiens GN | 237,417 | 61,06 | 12732 | 5 | 24,16 | 24,62 | 22,08 | 24,10 | 24,43 | 21,79 | **0,883** | 8,82E-01 | **0** | -0,1799997 | 0,05442378 |
| CB39L_HUM | CAB39L | sp\|Q9H9S4\|CB39L_HUMAN Calcium-binding protein 39-like OS=Homo sapiens GN=CAB3 | 43,664 | 11,28 | 39088 | 4 | 23,96 | 23,61 | 23,15 | 23,51 | 23,94 | 22,73 | **0,883** | 6,94E-01 | **0** | -0,1799997 | 0,15893418 |
| ALBU_HUMA | ALB | sp\|P02768\|ALBU_HUMAN Serum albumin OS=Homo sapiens GN=ALB PE=1 SV=2 | 267,946 | 9,85 | 69367 | 7 | 30,89 | 29,84 | 31,85 | 31,02 | 29,97 | 31,05 | **0,883** | 8,04E-01 | **0** | -0,1800003 | 0,09450673 |
| ARC1A_HUM | ARPC1A | sp\|Q92747\|ARC1A_HUMAN Actin-related protein 2/3 complex subunit 1A OS=Homo sap | 330,95 | 31,62 | 41569 | 9 | 25,36 | 25,30 | 24,67 | 24,74 | 25,25 | 24,80 | **0,883** | 5,46E-01 | **0** | -0,1800003 | 0,26289898 |
| SMRC1_HUM | SMARCC1 | sp\|Q92922\|SMRC1_HUMAN SWI/SNF complex subunit SMARCC1 OS=Homo sapiens GN | 165,111 | 6,61 | 122867 | 6 | 18,00 | 19,06 | 18,47 | 18,34 | 18,22 | 18,42 | **0,881** | 5,89E-01 | **0** | -0,1828899 | 0,22959766 |
| CND1_HUM | NCAPD2 | sp\|Q15021\|CND1_HUMAN Condensin complex subunit 1 OS=Homo sapiens GN=NCAPD | 44,392 | 1,86 | 157182 | 2 | 20,42 | 21,41 | 21,18 | 19,85 | 20,96 | 21,65 | **0,881** | 7,76E-01 | **0** | -0,1833337 | 0,10986549 |
| ABR_HUMA | ABR | sp\|Q12979\|ABR_HUMAN Active breakpoint cluster region-related protein OS=Homo sap | 129,456 | 5,59 | 97598 | 5 | 22,59 | 22,81 | 20,79 | 22,00 | 22,50 | 21,14 | **0,881** | 8,20E-01 | **0** | -0,1833337 | 0,08638055 |
| CDC42_HUM | CDC42 | sp\|P60953\|CDC42_HUMAN Cell division control protein 42 homolog OS=Homo sapiens | 587,557 | 64,4 | 21259 | 11 | 29,15 | 29,04 | 28,37 | 29,07 | 28,71 | 28,23 | **0,881** | 6,23E-01 | **0** | -0,1833344 | 0,20572617 |
| MKRN2_HUM | MKRN2 | sp\|Q9H000\|MKRN2_HUMAN Probable E3 ubiquitin-protein ligase makorin-2 OS=Homo | 75,622 | 8,89 | 46940 | 3 | 21,93 | 20,92 | 22,62 | 21,05 | 21,55 | 22,32 | **0,881** | 7,81E-01 | **0** | -0,1833344 | 0,10735321 |
| S12A4_HUM | SLC12A4 | sp\|Q9UP95\|S12A4_HUMAN Solute carrier family 12 member 4 OS=Homo sapiens GN=SI | 111,775 | 6,08 | 120650 | 6 | 24,75 | 25,13 | 23,73 | 24,87 | 24,83 | 23,35 | **0,879** | 7,89E-01 | **0** | -0,1866659 | 0,10303082 |
| 2A5D_HUMA | PPP2R5D | sp\|Q14738\|2A5D_HUMAN Serine/threonine-protein phosphatase 2A 56 kDa regulatory | 119,682 | 9,3 | 69992 | 4 | 22,07 | 21,18 | 23,01 | 20,19 | 22,61 | 22,90 | **0,879** | 8,62E-01 | **0** | -0,1866665 | 0,06440754 |
| CLIC4_HUM | CLIC4 | sp\|Q9Y696\|CLIC4_HUMAN Chloride intracellular channel protein 4 OS=Homo sapiens GI | 1453,286 | 81,82 | 28772 | 29 | 29,43 | 29,21 | 27,52 | 29,11 | 28,97 | 27,52 | **0,879** | 8,25E-01 | **0** | -0,1866665 | 0,08376732 |
| DCAF7_HUM | DCAF7 | sp\|P61962\|DCAF7_HUMAN DDB1- and CUL4-associated factor 7 OS=Homo sapiens GN= | 71,799 | 15,2 | 38926 | 4 | 23,17 | 23,35 | 23,41 | 23,38 | 23,14 | 22,85 | **0,879** | 3,32E-01 | **0** | -0,1866671 | 0,47860353 |
| RPAC1_HUM | POLR1C | sp\|O15160\|RPAC1_HUMAN DNA-directed RNA polymerases I and III subunit RPAC1 OS= | 76,619 | 10,69 | 39250 | 2 | 20,77 | 21,42 | 21,73 | 19,56 | 21,80 | 22,00 | **0,878** | 8,32E-01 | **0** | -0,1883316 | 0,07973609 |
| H2A2A_HUM | HIST2H2AA3 | sp\|Q6FI13\|H2A2A_HUMAN Histone H2A type 2-A OS=Homo sapiens GN=HIST2H2AA3 PI | 297,633 | 57,69 | 14095 | 6 | 17,64 | 19,71 | 18,73 | 18,19 | 18,27 | 19,05 | **0,877** | 7,87E-01 | **0** | -0,1899993 | 0,10400132 |
| CPSF6_HUM | CPSF6 | sp\|Q16630\|CPSF6_HUMAN Cleavage and polyadenylation specificity factor subunit 6 OS | 346,52 | 17,97 | 59210 | 8 | 25,57 | 25,53 | 25,71 | 25,62 | 25,44 | 25,18 | **0,877** | 2,43E-01 | **0** | -0,1899993 | 0,6141905 |
| STK38_HUM | STK38 | sp\|Q15208\|STK38_HUMAN Serine/threonine-protein kinase 38 OS=Homo sapiens GN=S | 173,777 | 21,94 | 54190 | 6 | 23,47 | 23,81 | 22,44 | 23,43 | 23,06 | 22,66 | **0,877** | 7,06E-01 | **0** | -0,1899999 | 0,15146491 |
| SCRIB_HUM | SCRIB | sp\|Q14160\|SCRIB_HUMAN Protein scribble homolog OS=Homo sapiens GN=SCRIB PE=1 | 115,35 | 3,07 | 174885 | 4 | 23,09 | 23,00 | 22,22 | 22,90 | 22,50 | 22,34 | **0,877** | 5,87E-01 | **0** | -0,1899999 | 0,23101089 |
| DHX36_HUM | DHX36 | sp\|Q9H2U1\|DHX36_HUMAN ATP-dependent RNA helicase DHX36 OS=Homo sapiens GN | 25,597 | 1,29 | 114760 | 1 | 17,95 | 17,74 | 18,77 | 17,85 | 17,34 | 18,70 | **0,877** | 7,26E-01 | **0** | -0,1899999 | 0,13883391 |
| CTGF_HUMA | CTGF | sp\|P29279\|CTGF_HUMAN Connective tissue growth factor OS=Homo sapiens GN=CTGF | 86,337 | 16,62 | 38091 | 4 | 21,19 | 20,64 | 24,23 | 21,47 | 20,47 | 23,55 | **0,877** | 9,01E-01 | **0** | -0,1900005 | 0,04513359 |
| FIBB_HUMA | FGB | sp\|P02675\|FIBB_HUMAN Fibrinogen beta chain OS=Homo sapiens GN=FGB PE=1 SV=2 | 75,558 | 5,3 | 55928 | 3 | 23,29 | 21,37 | 25,15 | 23,61 | 21,16 | 24,47 | **0,877** | 9,04E-01 | **0** | -0,1900005 | 0,04398193 |
| RAB43_HUM | RAB43 | sp\|Q86YS6\|RAB43_HUMAN Ras-related protein Rab-43 OS=Homo sapiens GN=RAB43 PI | 79,779 | 13,21 | 23339 | 4 | 21,25 | 21,25 | 21,58 | 21,96 | 21,79 | 19,76 | **0,876** | 8,03E-01 | **0** | -0,1909968 | 0,0953396 |
| VPP1_HUMA | ATP6V0A1 | sp\|Q93050\|VPP1_HUMAN V-type proton ATPase 116 kDa subunit a isoform 1 OS=Homo | 234,35 | 13,26 | 96413 | 8 | 24,41 | 24,51 | 24,57 | 24,44 | 24,38 | 24,09 | **0,875** | 1,76E-01 | **0** | -0,1933333 | 0,75488408 |
| PDLI5_HUM | PDLIM5 | sp\|Q96HC4\|PDLI5_HUMAN PDZ and LIM domain protein 5 OS=Homo sapiens GN=PDLIM | 307,512 | 17,45 | 63945 | 12 | 25,55 | 25,81 | 26,10 | 25,64 | 25,51 | 25,73 | **0,875** | 3,22E-01 | **0** | -0,1933333 | 0,49224206 |
| SDCB1_HUM | SDCBP | sp\|O00560\|SDCB1_HUMAN Syntenin-1 OS=Homo sapiens GN=SDCBP PE=1 SV=1 | 1281,036 | 65,77 | 32444 | 33 | 31,95 | 31,47 | 31,08 | 31,76 | 31,56 | 30,60 | **0,875** | 6,81E-01 | **0** | -0,1933333 | 0,16656395 |
| S12A9_HUM | SLC12A9 | sp\|Q9BXP2\|S12A9_HUMAN Solute carrier family 12 member 9 OS=Homo sapiens GN=SL | 9,983 | 0,77 | 96110 | 1 | 21,59 | 21,67 | 21,54 | 21,00 | 21,64 | 21,58 | **0,875** | 4,04E-01 | **0** | -0,1933339 | 0,39326777 |
| CEP55_HUM | CEP55 | sp\|Q53EZ4\|CEP55_HUMAN Centrosomal protein of 55 kDa OS=Homo sapiens GN=CEP5 | 842,233 | 27,16 | 54178 | 15 | 28,36 | 27,67 | 28,16 | 28,38 | 27,84 | 27,39 | **0,875** | 6,12E-01 | **0** | -0,1933339 | 0,21319963 |
| TFG_HUMA | TFG | sp\|Q92734\|TFG_HUMAN Protein TFG OS=Homo sapiens GN=TFG PE=1 SV=2 | 106,263 | 13,75 | 43448 | 5 | 25,82 | 25,49 | 24,86 | 25,71 | 25,73 | 24,15 | **0,875** | 7,61E-01 | **0** | -0,1933339 | 0,11847004 |
| NEXN_HUM | NEXN | sp\|Q0ZGT2\|NEXN_HUMAN Nexilin OS=Homo sapiens GN=NEXN PE=1 SV=1 | 43,662 | 3,85 | 80658 | 2 | 22,53 | 22,91 | 21,81 | 22,80 | 22,24 | 21,63 | **0,875** | 7,00E-01 | **0** | -0,1933339 | 0,15479017 |
| EMP3_HUM | EMP3 | sp\|P54852\|EMP3_HUMAN Epithelial membrane protein 3 OS=Homo sapiens GN=EMP3 | 17,669 | 4,91 | 18429 | 1 | 23,12 | 21,86 | 23,71 | 21,66 | 22,82 | 23,63 | **0,875** | 8,19E-01 | **0** | -0,1933339 | 0,08684083 |
| AGAP1_HUM | AGAP1 | sp\|Q9UPQ3\|AGAP1_HUMAN Arf-GAP with GTPase, ANK repeat and PH domain-containi | 42,265 | 1,87 | 94470 | 1 | 19,55 | 19,67 | 18,77 | 20,08 | 18,47 | 18,86 | **0,873** | 7,46E-01 | **0** | -0,1953894 | 0,1274226 |
| ARC1B_HUM | ARPC1B | sp\|O15143\|ARC1B_HUMAN Actin-related protein 2/3 complex subunit 1B OS=Homo sap | 462,746 | 43,82 | 40950 | 13 | 26,65 | 26,58 | 25,47 | 26,36 | 26,49 | 25,26 | **0,873** | 7,37E-01 | **0** | -0,1966661 | 0,13253952 |
| IMA3_HUMA | KPNA4 | sp\|O00629\|IMA3_HUMAN Importin subunit alpha-3 OS=Homo sapiens GN=KPNA4 PE=1 | 189,269 | 20,15 | 57887 | 6 | 22,52 | 22,35 | 23,31 | 22,32 | 22,70 | 22,57 | **0,873** | 5,68E-01 | **0** | -0,1966667 | 0,24601411 |
| B4GA1_HUM | B4GAT1 | sp\|O43505\|B4GA1_HUMAN Beta-1,4-glucuronyltransferase 1 OS=Homo sapiens GN=B4 | 80,037 | 3,86 | 47119 | 1 | 20,58 | 20,14 | 22,02 | 20,43 | 20,32 | 21,40 | **0,873** | 7,82E-01 | **0** | -0,1966667 | 0,10699927 |
| GNAS1_HUM | GNAS | sp\|Q5JWF2\|GNAS1_HUMAN Guanine nucleotide-binding protein G(s) subunit alpha isof | 760,599 | 20,54 | 111025 | 18 | 27,87 | 27,85 | 27,27 | 27,74 | 27,78 | 26,88 | **0,873** | 6,08E-01 | **0** | -0,1966674 | 0,21643338 |
| UBA6_HUM | UBA6 | sp\|A0AVT1\|UBA6_HUMAN Ubiquitin-like modifier-activating enzyme 6 OS=Homo sapier | 26,755 | 0,95 | 117970 | 1 | 19,71 | 20,51 | 21,51 | 17,64 | 21,46 | 22,03 | **0,871** | 8,99E-01 | **0** | -0,1991603 | 0,04626625 |
| CBR3_HUMA | CBR3 | sp\|O75828\|CBR3_HUMAN Carbonyl reductase [NADPH] 3 OS=Homo sapiens GN=CBR3 I | 28,449 | 2,89 | 30850 | 1 | 24,04 | 23,90 | 23,14 | 23,71 | 23,62 | 23,15 | **0,871** | 5,76E-01 | **0** | -0,2000001 | 0,23941688 |
| APOE_HUM | APOE | sp\|P02649\|APOE_HUMAN Apolipoprotein E OS=Homo sapiens GN=APOE PE=1 SV=1 | 119,127 | 9,15 | 36154 | 4 | 27,06 | 26,67 | 29,53 | 27,09 | 26,74 | 28,83 | **0,871** | 8,65E-01 | **0** | -0,2000001 | 0,06293964 |
| RFC5_HUMA | RFC5 | sp\|P40937\|RFC5_HUMAN Replication factor C subunit 5 OS=Homo sapiens GN=RFC5 PE | 85,087 | 12,35 | 38497 | 3 | 20,74 | 21,51 | 23,10 | 19,84 | 21,60 | 23,31 | **0,871** | 8,78E-01 | **0** | -0,2000001 | 0,05668022 |
| TAB1_HUMA | TAB1 | sp\|Q15750\|TAB1_HUMAN TGF-beta-activated kinase 1 and MAP3K7-binding protein 1 C | 71,214 | 7,14 | 54644 | 2 | 21,75 | 21,69 | 21,87 | 21,45 | 21,75 | 21,51 | **0,871** | 1,32E-01 | **0** | -0,2000001 | 0,88016002 |
| IF172_HUM | IFT172 | sp\|Q9UG01\|IF172_HUMAN Intraflagellar transport protein 172 homolog OS=Homo sapi | 602,058 | 13,84 | 197576 | 20 | 24,59 | 21,47 | 27,03 | 23,39 | 23,04 | 26,06 | **0,871** | 9,20E-01 | **0** | -0,2000001 | 0,03621448 |
| AP2B1_HUM | AP2B1 | sp\|P63010\|AP2B1_HUMAN AP-2 complex subunit beta OS=Homo sapiens GN=AP2B1 PE | 1932,283 | 65,85 | 104553 | 51 | 27,69 | 27,61 | 28,24 | 27,54 | 27,75 | 27,65 | **0,871** | 3,89E-01 | **0** | -0,2000001 | 0,41020189 |
| TNIK_HUMA | TNIK | sp\|Q9UKE5\|TNIK_HUMAN TRAF2 and NCK-interacting protein kinase OS=Homo sapiens | 461,314 | 13,68 | 154943 | 15 | 22,58 | 22,47 | 18,77 | 22,86 | 21,73 | 18,62 | **0,869** | 9,15E-01 | **0** | -0,2021929 | 0,03850546 |
| IFT74_HUM | IFT74 | sp\|Q96LB3\|IFT74_HUMAN Intraflagellar transport protein 74 homolog OS=Homo sapien | 48,834 | 4,5 | 69239 | 3 | 20,50 | 18,22 | 24,29 | 19,86 | 19,61 | 22,93 | **0,869** | 9,27E-01 | **0** | -0,202261 | 0,03301957 |
| PROF2_HUM | PFN2 | sp\|P35080\|PROF2_HUMAN Profilin-2 OS=Homo sapiens GN=PFN2 PE=1 SV=3 | 189,481 | 56,43 | 15046 | 5 | 24,95 | 25,05 | 21,17 | 24,95 | 24,77 | 20,84 | **0,869** | 9,18E-01 | **0** | -0,2033329 | 0,03722722 |
| CSRP1_HUM | CSRP1 | sp\|P21291\|CSRP1_HUMAN Cysteine and glycine-rich protein 1 OS=Homo sapiens GN=C | 529,651 | 50,78 | 20567 | 10 | 27,01 | 26,83 | 26,59 | 26,52 | 26,61 | 26,69 | **0,869** | 1,96E-01 | **0** | -0,2033329 | 0,70752677 |
| ITB1_HUMA | ITGB1 | sp\|P05556\|ITB1_HUMAN Integrin beta-1 OS=Homo sapiens GN=ITGB1 PE=1 SV=2 | 1664,134 | 34,34 | 88415 | 36 | 30,89 | 30,79 | 29,82 | 30,94 | 30,54 | 29,41 | **0,869** | 7,40E-01 | **0** | -0,2033329 | 0,13085273 |
| OTUD4_HUM | OTUD4 | sp\|Q01804\|OTUD4_HUMAN OTU domain-containing protein 4 OS=Homo sapiens GN=O | 51,19 | 1,08 | 124045 | 1 | 19,94 | 20,39 | 21,10 | 20,33 | 19,66 | 20,83 | **0,869** | 6,93E-01 | **0** | -0,2033335 | 0,15945663 |
| AMPD2_HUM | AMPD2 | sp\|Q01433\|AMPD2_HUMAN AMP deaminase 2 OS=Homo sapiens GN=AMPD2 PE=1 SV= | 62,095 | 2,16 | 100688 | 2 | 22,54 | 22,67 | 21,24 | 22,27 | 22,21 | 21,36 | **0,869** | 7,27E-01 | **0** | -0,2033335 | 0,13844479 |
| CBR1_HUMA | CBR1 | sp\|P16152\|CBR1_HUMAN Carbonyl reductase [NADPH] 1 OS=Homo sapiens GN=CBR1 P | 182,051 | 17,33 | 30375 | 4 | 24,49 | 24,75 | 24,93 | 24,33 | 24,32 | 24,91 | **0,869** | 4,32E-01 | **0** | -0,2033335 | 0,36420369 |
| NCS1_HUMA | NCS1 | sp\|P62166\|NCS1_HUMAN Neuronal calcium sensor 1 OS=Homo sapiens GN=NCS1 PE=1 | 151,586 | 31,58 | 21879 | 5 | 23,03 | 23,46 | 21,76 | 22,65 | 22,60 | 22,39 | **0,869** | 7,14E-01 | **0** | -0,2033335 | 0,14635859 |
| GSHR_HUM | GSR | sp\|P00390\|GSHR_HUMAN Glutathione reductase, mitochondrial OS=Homo sapiens GN= | 100,317 | 6,9 | 56257 | 3 | 24,08 | 24,02 | 22,53 | 24,35 | 24,23 | 21,44 | **0,869** | 8,59E-01 | **0** | -0,2033335 | 0,06575857 |
| SH3K1_HUM | SH3KBP1 | sp\|Q96B97\|SH3K1_HUMAN SH3 domain-containing kinase-binding protein 1 OS=Homo | 313,356 | 26,17 | 73126 | 12 | 24,64 | 25,08 | 21,60 | 24,94 | 24,81 | 20,95 | **0,867** | 9,09E-01 | **0** | -0,2066663 | 0,04123732 |
| XPOT_HUM | XPOT | sp\|O43592\|XPOT_HUMAN Exportin-T OS=Homo sapiens GN=XPOT PE=1 SV=2 | 138,725 | 4,78 | 109964 | 4 | 22,27 | 22,56 | 22,75 | 21,57 | 22,86 | 22,53 | **0,867** | 6,42E-01 | **0** | -0,2066663 | 0,19263524 |
| RAB5C_HUM | RAB5C | sp\|P51148\|RAB5C_HUMAN Ras-related protein Rab-5C OS=Homo sapiens GN=RAB5C PE | 573,419 | 66,67 | 23483 | 13 | 26,77 | 26,90 | 26,72 | 26,59 | 26,70 | 26,48 | **0,867** | 6,78E-02 | **0** | -0,2066663 | 1,16891853 |
| HEP2_HUMA | SERPIND1 | sp\|P05546\|HEP2_HUMAN Heparin cofactor 2 OS=Homo sapiens GN=SERPIND1 PE=1 SV | 68,918 | 5,01 | 57071 | 3 | 23,95 | 23,28 | 25,06 | 24,05 | 23,04 | 24,58 | **0,867** | 7,79E-01 | **0** | -0,2066669 | 0,10853579 |
| P4HA2_HUM | P4HA2 | sp\|O15460\|P4HA2_HUMAN Prolyl 4-hydroxylase subunit alpha-2 OS=Homo sapiens GN | 127,444 | 8,41 | 60902 | 3 | 20,58 | 20,19 | 23,46 | 21,17 | 19,87 | 22,57 | **0,866** | 8,80E-01 | **0** | -0,2072589 | 0,05533648 |
| 3BP1_HUMA | SH3BP1 | sp\|Q9Y3L3\|3BP1_HUMAN SH3 domain-binding protein 1 OS=Homo sapiens GN=SH3BP1 | 54,987 | 2 | 75713 | 1 | 20,80 | 21,21 | 19,58 | 20,51 | 20,31 | 20,14 | **0,865** | 6,97E-01 | **0** | -0,2099902 | 0,15696413 |
| PDE8A_HUM | PDE8A | sp\|O60658\|PDE8A_HUMAN High affinity cAMP-specific and IBMX-insensitive 3',5'-cyclic | 37,428 | 3,26 | 93304 | 1 | 18,80 | 19,94 | 18,55 | 19,05 | 19,12 | 18,49 | **0,865** | 6,79E-01 | **0** | -0,2099997 | 0,16788892 |
| MYO1C_HUM | MYO1C | sp\|O00159\|MYO1C_HUMAN Unconventional myosin-Ic OS=Homo sapiens GN=MYO1C I | 2898,992 | 53,06 | 121682 | 66 | 31,61 | 31,47 | 30,18 | 31,42 | 31,33 | 29,88 | **0,865** | 7,71E-01 | **0** | -0,2100004 | 0,11273007 |
| STX7_HUMA | STX7 | sp\|O15400\|STX7_HUMAN Syntaxin-7 OS=Homo sapiens GN=STX7 PE=1 SV=4 | 72,738 | 7,28 | 29816 | 2 | 22,43 | 21,97 | 21,62 | 22,47 | 21,67 | 21,25 | **0,865** | 6,49E-01 | **0** | -0,2100004 | 0,18758283 |
| PTGR2_HUM | PTGR2 | sp\|Q8N8N7\|PTGR2_HUMAN Prostaglandin reductase 2 OS=Homo sapiens GN=PTGR2 PI | 13,386 | 1,99 | 38499 | 1 | 21,28 | 20,75 | 19,46 | 20,81 | 21,23 | 18,82 | **0,864** | 8,30E-01 | **0** | -0,210804 | 0,08114166 |
| COX6C_HUM | COX6C | sp\|P09669\|COX6C_HUMAN Cytochrome c oxidase subunit 6C OS=Homo sapiens GN=CC | 14,431 | 9,33 | 8781 | 1 | 20,47 | 21,33 | 18,67 | 19,65 | 20,33 | 19,85 | **0,863** | 8,06E-01 | **0** | -0,2129205 | 0,0938034 |
| TM222_HUM | TMEM222 | sp\|Q9H0R3\|TM222_HUMAN Transmembrane protein 222 OS=Homo sapiens GN=TMEM | 67,325 | 6,73 | 23230 | 1 | 20,23 | 20,22 | 22,64 | 20,18 | 20,06 | 22,21 | **0,863** | 8,51E-01 | **0** | -0,2133331 | 0,07005465 |
| PDCD6_HUM | PDCD6 | sp\|O75340\|PDCD6_HUMAN Programmed cell death protein 6 OS=Homo sapiens GN=PD | 531,261 | 62,83 | 21868 | 13 | 27,69 | 27,98 | 26,64 | 27,65 | 27,93 | 26,09 | **0,863** | 7,76E-01 | **0** | -0,2133331 | 0,10987279 |
| PI42C_HUM | PIP4K2C | sp\|Q8TBX8\|PI42C_HUMAN Phosphatidylinositol 5-phosphate 4-kinase type-2 gamma O | 54,152 | 12,35 | 47300 | 3 | 22,31 | 22,49 | 21,38 | 22,08 | 22,22 | 21,24 | **0,863** | 6,67E-01 | **0** | -0,2133331 | 0,17575965 |
| MYL6_HUM | MYL6 | sp\|P60660\|MYL6_HUMAN Myosin light polypeptide 6 OS=Homo sapiens GN=MYL6 PE= | 769,065 | 66,23 | 16930 | 15 | 28,48 | 28,66 | 27,97 | 28,56 | 28,65 | 27,26 | **0,863** | 6,88E-01 | **0** | -0,2133331 | 0,16221408 |
| RLA2_HUMA | RPLP2 | sp\|P05387\|RLA2_HUMAN 60S acidic ribosomal protein P2 OS=Homo sapiens GN=RPLP2 | 459,273 | 63,48 | 11665 | 7 | 27,21 | 27,45 | 25,63 | 27,23 | 27,32 | 25,10 | **0,863** | 8,29E-01 | **0** | -0,2133331 | 0,08166412 |
| CPSF5_HUM | NUDT21 | sp\|O43809\|CPSF5_HUMAN Cleavage and polyadenylation specificity factor subunit 5 OS | 478,899 | 64,32 | 26227 | 14 | 26,54 | 26,60 | 26,80 | 26,33 | 26,22 | 26,75 | **0,863** | 3,01E-01 | **0** | -0,2133338 | 0,52199893 |
| FOLH1_HUM | FOLH1 | sp\|Q04609\|FOLH1_HUMAN Glutamate carboxypeptidase 2 OS=Homo sapiens GN=FOLH | 55,988 | 2,4 | 84331 | 2 | 21,50 | 21,40 | 21,27 | 21,48 | 21,21 | 20,84 | **0,863** | 3,40E-01 | **0** | -0,2133338 | 0,46857599 |
| VTA1_HUMA | VTA1 | sp\|Q9NP79\|VTA1_HUMAN Vacuolar protein sorting-associated protein VTA1 homolog C | 215,625 | 28,34 | 33879 | 9 | 26,12 | 26,10 | 24,91 | 26,17 | 25,92 | 24,40 | **0,863** | 7,70E-01 | **0** | -0,2133338 | 0,1133712 |
| APOA1_HUM | APOA1 | sp\|P02647\|APOA1_HUMAN Apolipoprotein A-I OS=Homo sapiens GN=APOA1 PE=1 SV= | 59,49 | 5,99 | 30778 | 1 | 23,73 | 23,60 | 27,11 | 23,07 | 23,84 | 26,89 | **0,863** | 9,03E-01 | **0** | -0,2133338 | 0,04449615 |
| ASSY_HUMA | ASS1 | sp\|P00966\|ASSY_HUMAN Argininosuccinate synthase OS=Homo sapiens GN=ASS1 PE=1 | 106,768 | 17,96 | 46530 | 8 | 25,48 | 25,14 | 24,68 | 25,06 | 25,07 | 24,52 | **0,861** | 5,03E-01 | **0** | -0,2166665 | 0,29864758 |
| CTNB1_HUM | CTNNB1 | sp\|P35222\|CTNB1_HUMAN Catenin beta-1 OS=Homo sapiens GN=CTNNB1 PE=1 SV=1 | 1303,667 | 49,94 | 85497 | 30 | 27,55 | 27,49 | 25,54 | 27,09 | 27,04 | 25,80 | **0,861** | 7,96E-01 | **0** | -0,2166665 | 0,09917262 |
| MET_HUMA | MET | sp\|P08581\|MET_HUMAN Hepatocyte growth factor receptor OS=Homo sapiens GN=ME | 236,316 | 9,28 | 155541 | 10 | 26,72 | 26,31 | 28,35 | 26,89 | 26,03 | 27,81 | **0,861** | 8,02E-01 | **0** | -0,2166665 | 0,09595735 |
| SC6A9_HUM | SLC6A9 | sp\|P48067\|SC6A9_HUMAN Sodium- and chloride-dependent glycine transporter 1 OS=I | 118,213 | 5,38 | 78260 | 4 | 24,24 | 24,04 | 20,91 | 23,69 | 24,03 | 20,82 | **0,861** | 8,91E-01 | **0** | -0,2166665 | 0,05016963 |
| CTDS1_HUM | CTDSP1 | sp\|Q9GZU7\|CTDS1_HUMAN Carboxy-terminal domain RNA polymerase II polypeptide A | 60,719 | 9,58 | 29203 | 2 | 22,17 | 21,94 | 21,64 | 21,97 | 21,83 | 21,30 | **0,861** | 4,44E-01 | **0** | -0,2166672 | 0,35274956 |
| EHD1_HUM | EHD1 | sp\|Q9H4M9\|EHD1_HUMAN EH domain-containing protein 1 OS=Homo sapiens GN=EHD | 2360,298 | 80,15 | 60627 | 55 | 29,91 | 29,54 | 28,02 | 29,72 | 29,28 | 27,82 | **0,861** | 8,03E-01 | **0** | -0,2166672 | 0,09502288 |
| DREB_HUMA | DBN1 | sp\|Q16643\|DREB_HUMAN Drebrin OS=Homo sapiens GN=DBN1 PE=1 SV=4 | 309,363 | 19,11 | 71429 | 8 | 25,52 | 25,20 | 25,62 | 25,11 | 25,03 | 25,55 | **0,861** | 3,51E-01 | **0** | -0,2166672 | 0,45477604 |
| EI2BB_HUM | EIF2B2 | sp\|P49770\|EI2BB_HUMAN Translation initiation factor eIF-2B subunit beta OS=Homo sa | 23,862 | 9,4 | 38990 | 1 | 18,09 | 17,72 | 20,21 | 17,89 | 18,83 | 18,66 | **0,859** | 8,04E-01 | **0** | -0,2194888 | 0,09477687 |
| 1433G_HUM | YWHAG | sp\|P61981\|1433G_HUMAN 14-3-3 protein gamma OS=Homo sapiens GN=YWHAG PE=1 | 690,172 | 68,02 | 28303 | 20 | 27,63 | 27,38 | 26,80 | 27,38 | 27,12 | 26,65 | **0,859** | 5,36E-01 | **0** | -0,2199993 | 0,27051941 |
| AP2S1_HUM | AP2S1 | sp\|P53680\|AP2S1_HUMAN AP-2 complex subunit sigma OS=Homo sapiens GN=AP2S1 PI | 84,86 | 33,8 | 17018 | 6 | 25,80 | 26,04 | 24,59 | 25,78 | 25,90 | 24,09 | **0,859** | 7,80E-01 | **0** | -0,2199999 | 0,10784252 |
| EXOC1_HUM | EXOC1 | sp\|Q9NV70\|EXOC1_HUMAN Exocyst complex component 1 OS=Homo sapiens GN=EXO | 63,843 | 3,47 | 101982 | 3 | 23,06 | 22,62 | 22,38 | 22,70 | 23,05 | 21,65 | **0,859** | 6,61E-01 | **0** | -0,2199999 | 0,17976455 |
| SH3L1_HUM | SH3BGRL | sp\|O75368\|SH3L1_HUMAN SH3 domain-binding glutamic acid-rich-like protein OS=Hom | 171,694 | 47,37 | 12774 | 5 | 25,00 | 25,15 | 22,96 | 24,65 | 24,98 | 22,87 | **0,859** | 8,33E-01 | **0** | -0,2199999 | 0,07961367 |
| PP1B_HUMA | PPP1CB | sp\|P62140\|PP1B_HUMAN Serine/threonine-protein phosphatase PP1-beta catalytic sub | 817,248 | 59,94 | 37187 | 18 | 25,11 | 25,41 | 25,34 | 24,85 | 25,38 | 24,97 | **0,859** | 2,98E-01 | **0** | -0,2200006 | 0,52508725 |
| RPEL1_HUM | RPEL1 | sp\|Q2QD12\|RPEL1_HUMAN Ribulose-phosphate 3-epimerase-like protein 1 OS=Homo s | 18,979 | 3,07 | 25023 | 1 | 21,98 | 22,34 | 19,31 | 22,14 | 22,04 | 18,79 | **0,858** | 8,87E-01 | **0** | -0,2202911 | 0,05203394 |
| GRK4_HUMA | GRK4 | sp\|P32298\|GRK4_HUMAN G protein-coupled receptor kinase 4 OS=Homo sapiens GN=C | 56,618 | 2,42 | 66583 | 1 | 20,63 | 20,42 | 21,27 | 20,34 | 20,27 | 21,04 | **0,857** | 5,63E-01 | **0** | -0,2233327 | 0,24944987 |
| H12_HUMAN | HIST1H1C | sp\|P16403\|H12_HUMAN Histone H1.2 OS=Homo sapiens GN=HIST1H1C PE=1 SV=2 | 111,436 | 15,49 | 21365 | 4 | 24,09 | 24,73 | 25,37 | 24,74 | 24,34 | 24,44 | **0,857** | 5,96E-01 | **0** | -0,2233334 | 0,22458948 |
| AKIB1_HUM | ANKIB1 | sp\|Q9P2G1\|AKIB1_HUMAN Ankyrin repeat and IBR domain-containing protein 1 OS=Ho | 95,291 | 8,54 | 122002 | 4 | 21,53 | 21,93 | 19,44 | 21,20 | 21,06 | 19,97 | **0,857** | 8,09E-01 | **0** | -0,223334 | 0,09212237 |
| EXOC2_HUM | EXOC2 | sp\|Q96KP1\|EXOC2_HUMAN Exocyst complex component 2 OS=Homo sapiens GN=EXO( | 30,966 | 2,6 | 104066 | 2 | 20,79 | 20,75 | 21,94 | 20,23 | 20,70 | 21,88 | **0,857** | 7,40E-01 | **0** | -0,223334 | 0,13095483 |
| MMS19_HUM | MMS19 | sp\|Q96T76\|MMS19_HUMAN MMS19 nucleotide excision repair protein homolog OS=Ho | 248,699 | 8,93 | 113290 | 7 | 22,20 | 22,75 | 23,61 | 21,72 | 22,49 | 23,68 | **0,857** | 7,66E-01 | **0** | -0,223334 | 0,1155293 |
| SC6A6_HUM | SLC6A6 | sp\|P31641\|SC6A6_HUMAN Sodium- and chloride-dependent taurine transporter OS=Ho | 10,7 | 2,74 | 69830 | 1 | 20,95 | 20,43 | 18,35 | 20,47 | 19,26 | 19,32 | **0,856** | 8,13E-01 | **0** | -0,2246278 | 0,09004474 |
| RL29_HUMA | RPL29 | sp\|P47914\|RL29_HUMAN 60S ribosomal protein L29 OS=Homo sapiens GN=RPL29 PE=1 | 30,762 | 5,03 | 17752 | 1 | 23,63 | 23,39 | 22,32 | 23,79 | 22,37 | 22,50 | **0,855** | 7,27E-01 | **0** | -0,2266655 | 0,13818358 |
| UBQL4_HUM | UBQLN4 | sp\|Q9NRR5\|UBQL4_HUMAN Ubiquilin-4 OS=Homo sapiens GN=UBQLN4 PE=1 SV=2 | 89,859 | 9,98 | 63853 | 5 | 22,07 | 22,67 | 21,71 | 21,99 | 21,24 | 22,54 | **0,855** | 6,54E-01 | **0** | -0,2266661 | 0,18414191 |
| CRUM2_HUM | CRB2 | sp\|Q5IJ48\|CRUM2_HUMAN Protein crumbs homolog 2 OS=Homo sapiens GN=CRB2 PE= | 210,776 | 4,36 | 134265 | 4 | 24,55 | 24,32 | 22,26 | 24,21 | 23,97 | 22,27 | **0,855** | 8,23E-01 | **0** | -0,2266668 | 0,08450535 |
| ITA5_HUMA | ITGA5 | sp\|P08648\|ITA5_HUMAN Integrin alpha-5 OS=Homo sapiens GN=ITGA5 PE=1 SV=2 | 357,726 | 19,26 | 114536 | 14 | 27,26 | 27,41 | 25,60 | 27,06 | 27,24 | 25,29 | **0,855** | 8,03E-01 | **0** | -0,2266668 | 0,09526635 |
| MYO1E_HUM | MYO1E | sp\|Q12965\|MYO1E_HUMAN Unconventional myosin-Ie OS=Homo sapiens GN=MYO1E I | 612,683 | 26,35 | 127062 | 22 | 26,09 | 25,72 | 25,85 | 25,98 | 26,08 | 24,92 | **0,855** | 5,89E-01 | **0** | -0,2266668 | 0,22975176 |
| TSN14_HUM | TSPAN14 | sp\|Q8NG11\|TSN14_HUMAN Tetraspanin-14 OS=Homo sapiens GN=TSPAN14 PE=1 SV=1 | 274,259 | 26,3 | 30691 | 7 | 27,36 | 26,91 | 25,58 | 27,05 | 26,82 | 25,30 | **0,855** | 7,82E-01 | **0** | -0,2266674 | 0,10675739 |
| TM245_HUM | TMEM245 | sp\|Q9H330\|TM245_HUMAN Transmembrane protein 245 OS=Homo sapiens GN=TMEM | 72,296 | 2,31 | 100945 | 2 | 18,15 | 19,97 | 21,91 | 18,25 | 19,49 | 21,61 | **0,855** | 8,84E-01 | **0** | -0,2267367 | 0,05334385 |
| STX1B_HUM | STX1B | sp\|P61266\|STX1B_HUMAN Syntaxin-1B OS=Homo sapiens GN=STX1B PE=1 SV=1 | 24,003 | 4,86 | 33245 | 1 | 20,27 | 20,37 | 18,66 | 19,56 | 20,28 | 18,78 | **0,854** | 7,62E-01 | **0** | -0,2284082 | 0,11790577 |
| LIMA1_HUM | LIMA1 | sp\|Q9UHB6\|LIMA1_HUMAN LIM domain and actin-binding protein 1 OS=Homo sapiens | 133,037 | 6,59 | 85226 | 5 | 24,32 | 25,14 | 23,43 | 24,27 | 24,34 | 23,59 | **0,853** | 6,97E-01 | **0** | -0,2299995 | 0,15701724 |
| LIS1_HUMA | PAFAH1B1 | sp\|P43034\|LIS1_HUMAN Platelet-activating factor acetylhydrolase IB subunit alpha OS=I | 601,914 | 58,78 | 46638 | 19 | 26,33 | 26,39 | 25,84 | 25,66 | 26,19 | 26,02 | **0,853** | 3,81E-01 | **0** | -0,2299995 | 0,41873109 |
| NDRG2_HUM | NDRG2 | sp\|Q9UN36\|NDRG2_HUMAN Protein NDRG2 OS=Homo sapiens GN=NDRG2 PE=1 SV=2 | 29,413 | 4,04 | 40798 | 1 | 20,76 | 20,76 | 20,63 | 20,73 | 20,29 | 20,44 | **0,853** | 1,67E-01 | **0** | -0,2299995 | 0,77834219 |
| RHOC_HUM | RHOC | sp\|P08134\|RHOC_HUMAN Rho-related GTP-binding protein RhoC OS=Homo sapiens GN | 696,341 | 71,5 | 22006 | 14 | 25,72 | 25,70 | 25,55 | 25,35 | 25,79 | 25,14 | **0,853** | 3,12E-01 | **0** | -0,2299995 | 0,50603206 |
| DDX3X_HUM | DDX3X | sp\|O00571\|DDX3X_HUMAN ATP-dependent RNA helicase DDX3X OS=Homo sapiens GN | 1307,572 | 54,98 | 73243 | 36 | 28,64 | 28,42 | 29,04 | 28,35 | 28,21 | 28,85 | **0,853** | 4,36E-01 | **0** | -0,2300002 | 0,36078217 |
| FABP4_HUM | FABP4 | sp\|P15090\|FABP4_HUMAN Fatty acid-binding protein, adipocyte OS=Homo sapiens GN= | 53,11 | 15,91 | 14719 | 2 | 21,17 | 21,79 | 24,66 | 21,17 | 21,73 | 24,03 | **0,853** | 8,76E-01 | **0** | -0,2300002 | 0,05736615 |
| HBB_HUMA | HBB | sp\|P68871\|HBB_HUMAN Hemoglobin subunit beta OS=Homo sapiens GN=HBB PE=1 SV= | 69,996 | 12,93 | 15998 | 2 | 28,58 | 27,94 | 29,70 | 28,59 | 27,78 | 29,16 | **0,853** | 7,42E-01 | **0** | -0,2300002 | 0,12959998 |
| GNPI2_HUM | GNPDA2 | sp\|Q8TDQ7\|GNPI2_HUMAN Glucosamine-6-phosphate isomerase 2 OS=Homo sapiens ( | 183,213 | 39,13 | 31085 | 8 | 23,58 | 24,22 | 23,52 | 23,71 | 24,12 | 22,80 | **0,853** | 6,36E-01 | **0** | -0,2300002 | 0,19650995 |
| CN37_HUMA | CNP | sp\|P09543\|CN37_HUMAN 2',3'-cyclic-nucleotide 3'-phosphodiesterase OS=Homo sapie | 126,855 | 11,4 | 47579 | 5 | 25,89 | 24,96 | 23,88 | 25,72 | 24,92 | 23,39 | **0,851** | 8,08E-01 | **0** | -0,233333 | 0,0928056 |
| TPPC5_HUM | TRAPPC5 | sp\|Q8IUR0\|TPPC5_HUMAN Trafficking protein particle complex subunit 5 OS=Homo sap | 75,025 | 19,15 | 20783 | 2 | 20,83 | 21,07 | 21,74 | 20,67 | 21,17 | 21,10 | **0,851** | 4,99E-01 | **0** | -0,233333 | 0,30219734 |
| BAG5_HUMA | BAG5 | sp\|Q9UL15\|BAG5_HUMAN BAG family molecular chaperone regulator 5 OS=Homo sapie | 183,116 | 10,96 | 51200 | 4 | 23,98 | 24,12 | 23,80 | 23,80 | 23,64 | 23,76 | **0,851** | 8,90E-02 | **0** | -0,2333336 | 1,05056126 |
| PLTP_HUMA | PLTP | sp\|P55058\|PLTP_HUMAN Phospholipid transfer protein OS=Homo sapiens GN=PLTP PE= | 34,719 | 2,23 | 54739 | 1 | 21,10 | 20,24 | 23,15 | 20,88 | 19,98 | 22,93 | **0,851** | 8,59E-01 | **0** | -0,2333336 | 0,06625666 |
| RBP56_HUM | TAF15 | sp\|Q92804\|RBP56_HUMAN TATA-binding protein-associated factor 2N OS=Homo sapie | 71,987 | 5,41 | 61830 | 2 | 20,64 | 21,09 | 18,98 | 20,62 | 21,02 | 18,36 | **0,849** | 8,33E-01 | **0** | -0,2356669 | 0,0793779 |
| TB10A_HUM | TBC1D10A | sp\|Q9BXI6\|TB10A_HUMAN TBC1 domain family member 10A OS=Homo sapiens GN=TB( | 129,426 | 11,81 | 57118 | 5 | 24,92 | 24,46 | 22,93 | 24,77 | 24,34 | 22,49 | **0,849** | 8,10E-01 | **0** | -0,2366664 | 0,09141655 |
| TB10B_HUM | TBC1D10B | sp\|Q4KMP7\|TB10B_HUMAN TBC1 domain family member 10B OS=Homo sapiens GN=TI | 19,203 | 1,86 | 87199 | 1 | 21,67 | 21,80 | 21,18 | 21,53 | 21,66 | 20,75 | **0,849** | 5,26E-01 | **0** | -0,2366664 | 0,27899666 |
| PZP_HUMAN | PZP | sp\|P20742\|PZP_HUMAN Pregnancy zone protein OS=Homo sapiens GN=PZP PE=1 SV=4 | 148,274 | 2,29 | 163832 | 4 | 24,30 | 23,06 | 26,51 | 24,39 | 23,32 | 25,45 | **0,849** | 8,51E-01 | **0** | -0,2366664 | 0,0700613 |
| ITIH2_HUM | ITIH2 | sp\|P19823\|ITIH2_HUMAN Inter-alpha-trypsin inhibitor heavy chain H2 OS=Homo sapien | 327,358 | 10,25 | 106463 | 9 | 26,00 | 24,84 | 27,72 | 26,01 | 24,75 | 27,09 | **0,849** | 8,37E-01 | **0** | -0,2366664 | 0,07746493 |
| LRP12_HUM | LRP12 | sp\|Q9Y561\|LRP12_HUMAN Low-density lipoprotein receptor-related protein 12 OS=Ho | 14,235 | 0,81 | 94984 | 1 | 18,71 | 18,54 | 22,73 | 18,10 | 18,05 | 23,12 | **0,848** | 9,18E-01 | **0** | -0,2375673 | 0,03714106 |
| PEA15_HUM | PEA15 | sp\|Q15121\|PEA15_HUMAN Astrocytic phosphoprotein PEA-15 OS=Homo sapiens GN=P | 37,484 | 9,23 | 15040 | 1 | 20,91 | 21,14 | 18,15 | 20,46 | 20,90 | 18,13 | **0,848** | 8,63E-01 | **0** | -0,237819 | 0,06420965 |
| LXN_HUMAN | LXN | sp\|Q9BS40\|LXN_HUMAN Latexin OS=Homo sapiens GN=LXN PE=1 SV=2 | 24,687 | 4,5 | 25750 | 1 | 21,75 | 21,63 | 20,97 | 21,42 | 21,37 | 20,84 | **0,847** | 4,76E-01 | **0** | -0,2399991 | 0,32256215 |
| PEF1_HUMA | PEF1 | sp\|Q9UBV8\|PEF1_HUMAN Peflin OS=Homo sapiens GN=PEF1 PE=1 SV=1 | 124,712 | 12,32 | 30381 | 3 | 22,58 | 22,22 | 24,06 | 21,91 | 22,87 | 23,36 | **0,847** | 7,51E-01 | **0** | -0,2399991 | 0,12437518 |
| SRBS2_HUM | SORBS2 | sp\|O94875\|SRBS2_HUMAN Sorbin and SH3 domain-containing protein 2 OS=Homo sapi | 27,912 | 1,91 | 124108 | 1 | 20,84 | 20,60 | 20,97 | 20,58 | 20,52 | 20,59 | **0,847** | 9,57E-02 | **0** | -0,2399998 | 1,01901904 |
| 1433T_HUM | YWHAQ | sp\|P27348\|1433T_HUMAN 14-3-3 protein theta OS=Homo sapiens GN=YWHAQ PE=1 SV | 1032,333 | 68,57 | 27764 | 23 | 27,18 | 27,42 | 26,58 | 26,87 | 26,93 | 26,66 | **0,847** | 4,13E-01 | **0** | -0,2399998 | 0,38416221 |
| PICAL_HUM | PICALM | sp\|Q13492\|PICAL_HUMAN Phosphatidylinositol-binding clathrin assembly protein OS=H | 678,073 | 29,6 | 70755 | 18 | 26,02 | 25,48 | 26,99 | 25,64 | 25,53 | 26,60 | **0,847** | 6,89E-01 | **0** | -0,2399998 | 0,16182302 |
| EHD3_HUM | EHD3 | sp\|Q9NZN3\|EHD3_HUMAN EH domain-containing protein 3 OS=Homo sapiens GN=EHD | 1128,738 | 52,52 | 60887 | 31 | 25,52 | 24,91 | 24,43 | 25,40 | 24,84 | 23,90 | **0,847** | 6,79E-01 | **0** | -0,2400004 | 0,16787573 |
| SNTB2_HUM | SNTB2 | sp\|Q13425\|SNTB2_HUMAN Beta-2-syntrophin OS=Homo sapiens GN=SNTB2 PE=1 SV=1 | 383,134 | 22,59 | 57950 | 12 | 26,49 | 26,58 | 25,03 | 26,17 | 26,21 | 25,00 | **0,847** | 7,27E-01 | **0** | -0,2400004 | 0,13860539 |
| GNAI2_HUM | GNAI2 | sp\|P04899\|GNAI2_HUMAN Guanine nucleotide-binding protein G(i) subunit alpha-2 OS | 1299,339 | 62,82 | 40451 | 29 | 28,04 | 27,93 | 27,88 | 27,75 | 27,94 | 27,43 | **0,845** | 1,94E-01 | **0** | -0,2433332 | 0,71192916 |
| TMA7_HUM | TMA7 | sp\|Q9Y2S6\|TMA7_HUMAN Translation machinery-associated protein 7 OS=Homo sapie | 24,575 | 29,69 | 7066 | 2 | 24,62 | 23,47 | 18,85 | 23,72 | 23,62 | 18,87 | **0,845** | 9,24E-01 | **0** | -0,2433332 | 0,03455295 |
| CNN1_HUM | CNN1 | sp\|P51911\|CNN1_HUMAN Calponin-1 OS=Homo sapiens GN=CNN1 PE=1 SV=2 | 90,961 | 21,89 | 33170 | 4 | 23,86 | 24,11 | 22,77 | 24,25 | 24,33 | 21,43 | **0,845** | 8,26E-01 | **0** | -0,2433338 | 0,08288482 |

| | | | | | | | | | | | | | | | | |
|---|---|---|---|---|---|---|---|---|---|---|---|---|---|---|---|---|
| RASA3_HUM | RASA3 | sp\|Q14644\|RASA3_HUMAN Ras GTPase-activating protein 3 OS=Homo sapiens GN=RAS/ | 68,584 | 3,48 | 95699 | 2 | 21,04 | 20,01 | 19,45 | 21,55 | 19,77 | 18,44 | **0,844** | 8,21E-01 | **0** | -0,2450752 | 0,08580103 |
| NU107_HUN | NUP107 | sp\|P57740\|NU107_HUMAN Nuclear pore complex protein Nup107 OS=Homo sapiens G | 98,93 | 2,59 | 106374 | 2 | 19,14 | 21,16 | 22,16 | 17,47 | 21,13 | 23,13 | **0,843** | 9,02E-01 | **0** | -0,2462076 | 0,04471774 |
| MARCS_HUN | MARCKS | sp\|P29966\|MARCS_HUMAN Myristoylated alanine-rich C-kinase substrate OS=Homo sa | 209,651 | 15,66 | 31555 | 5 | 26,94 | 26,45 | 24,57 | 26,59 | 26,45 | 24,18 | **0,843** | 8,28E-01 | **0** | -0,2466666 | 0,08195707 |
| TOLIP_HUM/ | TOLLIP | sp\|Q9H0E2\|TOLIP_HUMAN Toll-interacting protein OS=Homo sapiens GN=TOLLIP PE=1 | 92,048 | 13,5 | 30282 | 3 | 24,95 | 25,04 | 24,83 | 24,85 | 24,90 | 24,33 | **0,843** | 2,68E-01 | **0** | -0,2466672 | 0,57105613 |
| MOT4_HUM. | SLC16A3 | sp\|O15427\|MOT4_HUMAN Monocarboxylate transporter 4 OS=Homo sapiens GN=SLC1 | 51,864 | 3,23 | 49469 | 2 | 23,59 | 22,92 | 21,94 | 22,80 | 23,32 | 21,59 | **0,843** | 7,43E-01 | **0** | -0,2466672 | 0,12908296 |
| MALD1_HUN | MARVELD1 | sp\|Q9BSK0\|MALD1_HUMAN MARVEL domain-containing protein 1 OS=Homo sapiens G | 32,356 | 4,05 | 18914 | 1 | 23,79 | 23,99 | 19,20 | 23,61 | 23,58 | 19,04 | **0,842** | 9,15E-01 | **0** | -0,2481346 | 0,03864591 |
| LYN_HUMAN | LYN | sp\|P07948\|LYN_HUMAN Tyrosine-protein kinase Lyn OS=Homo sapiens GN=LYN PE=1 S | 254,716 | 17,38 | 58574 | 8 | 24,14 | 23,51 | 22,04 | 24,12 | 23,50 | 21,32 | **0,841** | 8,24E-01 | **0** | -0,25 | 0,08410564 |
| KIF23_HUM/ | KIF23 | sp\|Q02241\|KIF23_HUMAN Kinesin-like protein KIF23 OS=Homo sapiens GN=KIF23 PE=1 | 1880,483 | 33,75 | 110059 | 43 | 28,00 | 27,75 | 30,69 | 28,93 | 27,51 | 29,25 | **0,841** | 8,29E-01 | **0** | -0,25 | 0,08161712 |
| ADA10_HUM | ADAM10 | sp\|O14672\|ADA10_HUMAN Disintegrin and metalloproteinase domain-containing prote | 891,373 | 37,3 | 84142 | 24 | 28,78 | 28,38 | 27,52 | 28,64 | 28,18 | 27,11 | **0,841** | 6,92E-01 | **0** | -0,25 | 0,16008594 |
| DCNL1_HUM | DCUN1D1 | sp\|Q96GG9\|DCNL1_HUMAN DCN1-like protein 1 OS=Homo sapiens GN=DCUN1D1 PE=1 | 37,522 | 10,42 | 30124 | 3 | 22,17 | 22,01 | 21,48 | 21,98 | 21,46 | 21,47 | **0,841** | 4,07E-01 | **0** | -0,2500006 | 0,39034863 |
| TARB1_HUM | TARBP1 | sp\|Q13395\|TARB1_HUMAN Probable methyltransferase TARBP1 OS=Homo sapiens GN= | 57,7 | 0,86 | 181675 | 1 | 18,97 | 18,68 | 18,67 | 17,85 | 18,74 | 18,98 | **0,840** | 5,20E-01 | **0** | -0,2509441 | 0,28365063 |
| RL26L_HUM/ | RPL26L1 | sp\|Q9UNX3\|RL26L_HUMAN 60S ribosomal protein L26-like 1 OS=Homo sapiens GN=RPL | 232,853 | 45,52 | 17256 | 10 | 22,26 | 21,71 | 22,89 | 22,28 | 20,65 | 23,17 | **0,839** | 7,71E-01 | **0** | -0,2533328 | 0,11302988 |
| TBCE_HUMA | TBCE | sp\|Q15813\|TBCE_HUMAN Tubulin-specific chaperone E OS=Homo sapiens GN=TBCE PE | 25,578 | 2,28 | 59346 | 1 | 19,97 | 20,82 | 19,79 | 19,62 | 19,71 | 20,49 | **0,839** | 5,80E-01 | **0** | -0,2533334 | 0,23677914 |
| MDR1_HUM. | ABCB1 | sp\|P08183\|MDR1_HUMAN Multidrug resistance protein 1 OS=Homo sapiens GN=ABCB1 | 417,677 | 12,5 | 141479 | 13 | 26,07 | 26,18 | 24,80 | 25,67 | 25,88 | 24,74 | **0,839** | 6,77E-01 | **0** | -0,2533334 | 0,16948336 |
| CHM4B_HUN | CHMP4B | sp\|Q9H444\|CHM4B_HUMAN Charged multivesicular body protein 4b OS=Homo sapiens | 414,724 | 41,52 | 24950 | 13 | 29,93 | 29,64 | 27,45 | 30,02 | 29,23 | 27,01 | **0,839** | 8,42E-01 | **0** | -0,2533334 | 0,0745379 |
| MVP_HUMA | MVP | sp\|Q14764\|MVP_HUMAN Major vault protein OS=Homo sapiens GN=MVP PE=1 SV=4 | 873,215 | 28,67 | 99327 | 19 | 27,13 | 27,47 | 26,74 | 26,81 | 27,22 | 26,55 | **0,839** | 4,28E-01 | **0** | -0,2533334 | 0,36893946 |
| PHS_HUMAN | PCBD1 | sp\|P61457\|PHS_HUMAN Pterin-4-alpha-carbinolamine dehydratase OS=Homo sapiens ( | 59,969 | 15,38 | 12000 | 2 | 19,22 | 19,26 | 19,39 | 19,18 | 19,61 | 18,31 | **0,837** | 5,42E-01 | **0** | -0,2564322 | 0,26584423 |
| MINY3_HUM | MINDY3 | sp\|Q9H8M7\|MINY3_HUMAN Ubiquitin carboxyl-terminal hydrolase MINDY-3 OS=Homo | 185,87 | 17,75 | 49725 | 6 | 23,81 | 23,93 | 22,84 | 23,86 | 23,34 | 22,61 | **0,837** | 6,35E-01 | **0** | -0,2566662 | 0,19715389 |
| CO3_HUMAN | C3 | sp\|P01024\|CO3_HUMAN Complement C3 OS=Homo sapiens GN=C3 PE=1 SV=2 | 589,788 | 9,98 | 187148 | 16 | 28,13 | 26,24 | 28,85 | 27,87 | 26,40 | 28,18 | **0,837** | 8,01E-01 | **0** | -0,2566662 | 0,09642966 |
| LIMS1_HUM. | LIMS1 | sp\|P48059\|LIMS1_HUMAN LIM and senescent cell antigen-like-containing domain prote | 246,443 | 35,08 | 37251 | 9 | 25,99 | 25,41 | 24,83 | 25,77 | 25,16 | 24,53 | **0,837** | 6,28E-01 | **0** | -0,2566662 | 0,20189078 |
| DHYS_HUMA | DHPS | sp\|P49366\|DHYS_HUMAN Deoxyhypusine synthase OS=Homo sapiens GN=DHPS PE=1 S | 51,519 | 5,42 | 40971 | 1 | 18,04 | 18,72 | 18,31 | 18,00 | 17,99 | 18,31 | **0,837** | 3,15E-01 | **0** | -0,2566668 | 0,50102721 |
| STABP_HUM | STAMBP | sp\|O95630\|STABP_HUMAN STAM-binding protein OS=Homo sapiens GN=STAMBP PE=1 | 329,172 | 25,71 | 48077 | 9 | 25,94 | 25,29 | 24,11 | 25,69 | 25,18 | 23,70 | **0,837** | 7,65E-01 | **0** | -0,2566668 | 0,11637794 |
| RAB15_HUM | RAB15 | sp\|P59190\|RAB15_HUMAN Ras-related protein Rab-15 OS=Homo sapiens GN=RAB15 PE | 95,32 | 10,38 | 24391 | 2 | 27,51 | 27,25 | 27,21 | 26,75 | 27,40 | 27,05 | **0,837** | 2,89E-01 | **0** | -0,2566668 | 0,53936822 |
| RAB32_HUM | RAB32 | sp\|Q13637\|RAB32_HUMAN Ras-related protein Rab-32 OS=Homo sapiens GN=RAB32 PI | 55,678 | 14,22 | 24997 | 3 | 22,27 | 21,61 | 21,80 | 21,82 | 21,07 | 22,02 | **0,837** | 5,03E-01 | **0** | -0,2566668 | 0,29813651 |
| RB33B_HUM | RAB33B | sp\|Q9H082\|RB33B_HUMAN Ras-related protein Rab-33B OS=Homo sapiens GN=RAB33E | 87,435 | 5,68 | 25718 | 2 | 27,54 | 27,27 | 27,27 | 26,81 | 27,41 | 27,09 | **0,837** | 2,59E-01 | **0** | -0,2566675 | 0,58656516 |
| SENP3_HUM | SENP3 | sp\|Q9H4L4\|SENP3_HUMAN Sentrin-specific protease 3 OS=Homo sapiens GN=SENP3 PE | 14,876 | 1,57 | 65010 | 1 | 17,37 | 17,95 | 19,52 | 17,47 | 16,99 | 19,60 | **0,835** | 8,13E-01 | **0** | -0,2598565 | 0,09001014 |
| BROX_HUM/ | BROX | sp\|Q5VW32\|BROX_HUMAN BRO1 domain-containing protein BROX OS=Homo sapiens ( | 383,33 | 32,36 | 46476 | 11 | 26,98 | 26,48 | 26,76 | 26,81 | 26,43 | 26,20 | **0,835** | 3,20E-01 | **0** | -0,2599996 | 0,49465553 |
| MILK1_HUM | MICALL1 | sp\|Q8N3F8\|MILK1_HUMAN MICAL-like protein 1 OS=Homo sapiens GN=MICALL1 PE=1 | 111,643 | 2,78 | 93441 | 2 | 22,58 | 22,38 | 20,22 | 22,56 | 22,23 | 19,61 | **0,835** | 8,39E-01 | **0** | -0,2599996 | 0,07615293 |
| OPA1_HUM/ | OPA1 | sp\|O60313\|OPA1_HUMAN Dynamin-like 120 kDa protein, mitochondrial OS=Homo sapi | 126,056 | 7,81 | 111631 | 5 | 22,00 | 22,89 | 22,58 | 22,64 | 22,26 | 21,79 | **0,835** | 5,08E-01 | **0** | -0,2599996 | 0,29381899 |
| MMP14_HUI | MMP14 | sp\|P50281\|MMP14_HUMAN Matrix metalloproteinase-14 OS=Homo sapiens GN=MMP1 | 126,251 | 11 | 65894 | 5 | 25,16 | 24,70 | 24,51 | 24,55 | 24,75 | 24,29 | **0,835** | 3,30E-01 | **0** | -0,2600002 | 0,48198547 |
| NECP2_HUM | NECAP2 | sp\|Q9NVZ3\|NECP2_HUMAN Adaptin ear-binding coat-associated protein 2 OS=Homo sa | 16,962 | 5,32 | 28339 | 1 | 18,72 | 21,26 | 21,73 | 18,24 | 21,00 | 21,69 | **0,835** | 8,62E-01 | **0** | -0,2601693 | 0,06429171 |
| RB11A_HUM | RAB11A | sp\|P62491\|RB11A_HUMAN Ras-related protein Rab-11A OS=Homo sapiens GN=RAB11A | 464,309 | 59,72 | 24394 | 12 | 23,90 | 23,75 | 17,77 | 23,73 | 23,19 | 17,72 | **0,835** | 9,30E-01 | **0** | -0,2606958 | 0,03154974 |
| AP3B1_HUM | AP3B1 | sp\|O00203\|AP3B1_HUMAN AP-3 complex subunit beta-1 OS=Homo sapiens GN=AP3B1 | 15,781 | 1,19 | 121320 | 1 | 19,50 | 18,12 | 20,09 | 17,71 | 19,06 | 20,15 | **0,834** | 7,89E-01 | **0** | -0,2612998 | 0,10270169 |
| EHD4_HUM/ | EHD4 | sp\|Q9H223\|EHD4_HUMAN EH domain-containing protein 4 OS=Homo sapiens GN=EHD | 1076,375 | 53,23 | 61175 | 31 | 28,16 | 27,96 | 26,97 | 28,03 | 27,78 | 26,49 | **0,833** | 6,85E-01 | **0** | -0,2633324 | 0,16454399 |
| EDF1_HUMA | EDF1 | sp\|O60869\|EDF1_HUMAN Endothelial differentiation-related factor 1 OS=Homo sapiens | 68,05 | 16,89 | 16369 | 2 | 23,79 | 24,17 | 22,35 | 23,91 | 23,77 | 21,84 | **0,833** | 7,77E-01 | **0** | -0,2633336 | 0,1097407 |
| L2GL2_HUM. | LLGL2 | sp\|Q6P1M3\|L2GL2_HUMAN Lethal(2) giant larvae protein homolog 2 OS=Homo sapiens | 40,957 | 3,14 | 113448 | 2 | 21,89 | 22,02 | 20,45 | 21,46 | 21,95 | 20,16 | **0,833** | 7,38E-01 | **0** | -0,2633336 | 0,13205959 |
| AP2M1_HUN | AP2M1 | sp\|Q96CW1\|AP2M1_HUMAN AP-2 complex subunit mu OS=Homo sapiens GN=AP2M1 F | 697,281 | 54,71 | 49655 | 22 | 27,44 | 27,25 | 27,79 | 27,21 | 27,26 | 27,22 | **0,833** | 1,73E-01 | **0** | -0,2633343 | 0,76250209 |
| HTRA1_HUM | HTRA1 | sp\|Q92743\|HTRA1_HUMAN Serine protease HTRA1 OS=Homo sapiens GN=HTRA1 PE=1 | 431,486 | 32,08 | 51287 | 11 | 26,56 | 25,66 | 26,05 | 26,08 | 25,40 | 25,99 | **0,831** | 4,73E-01 | **0** | -0,2666664 | 0,32541327 |
| GBB1_HUM/ | GNB1 | sp\|P62873\|GBB1_HUMAN Guanine nucleotide-binding protein G(I)/G(S)/G(T) subunit be | 1317,351 | 68,53 | 37377 | 22 | 29,10 | 29,21 | 28,28 | 29,02 | 28,94 | 27,83 | **0,831** | 6,10E-01 | **0** | -0,2666664 | 0,2143365 |
| MYH10_HUN | MYH10 | sp\|P35580\|MYH10_HUMAN Myosin-10 OS=Homo sapiens GN=MYH10 PE=1 SV=3 | 4002,118 | 51,27 | 228999 | 96 | 28,09 | 28,51 | 28,33 | 28,19 | 28,32 | 27,62 | **0,831** | 3,41E-01 | **0** | -0,2666664 | 0,46715663 |
| PACN3_HUM | PACSIN3 | sp\|Q9UKS6\|PACN3_HUMAN Protein kinase C and casein kinase substrate in neurons pro | 749,44 | 44,1 | 48487 | 17 | 28,84 | 28,25 | 27,39 | 28,67 | 28,14 | 26,87 | **0,831** | 7,15E-01 | **0** | -0,2666664 | 0,14572232 |
| VINC_HUMA | VCL | sp\|P18206\|VINC_HUMAN Vinculin OS=Homo sapiens GN=VCL PE=1 SV=4 | 3238,763 | 59,26 | 123799 | 82 | 30,27 | 30,30 | 29,67 | 30,15 | 30,05 | 29,24 | **0,831** | 4,93E-01 | **0** | -0,266667 | 0,30728488 |
| STEA1_HUM. | STEAP1 | sp\|Q9UHE8\|STEA1_HUMAN Metalloreductase STEAP1 OS=Homo sapiens GN=STEAP1 PI | 23,962 | 4,72 | 39851 | 2 | 24,69 | 23,24 | 22,45 | 23,97 | 23,70 | 21,91 | **0,831** | 7,87E-01 | **0** | -0,266667 | 0,10428382 |
| GBB2_HUM/ | GNB2 | sp\|P62879\|GBB2_HUMAN Guanine nucleotide-binding protein G(I)/G(S)/G(T) subunit be | 1260,408 | 68,82 | 37331 | 21 | 27,85 | 27,99 | 26,95 | 27,59 | 27,85 | 26,55 | **0,831** | 6,31E-01 | **0** | -0,266667 | 0,19990368 |
| ARPC5_HUM | ARPC5 | sp\|O15511\|ARPC5_HUMAN Actin-related protein 2/3 complex subunit 5 OS=Homo sapi | 205,021 | 39,74 | 16320 | 4 | 24,16 | 26,03 | 24,49 | 24,42 | 25,41 | 24,05 | **0,831** | 7,24E-01 | **0** | -0,266667 | 0,14000583 |
| RD23B_HUM | RAD23B | sp\|P54727\|RD23B_HUMAN UV excision repair protein RAD23 homolog B OS=Homo sapi | 444,204 | 37,16 | 43171 | 13 | 27,09 | 27,28 | 25,36 | 26,88 | 26,81 | 25,24 | **0,831** | 7,59E-01 | **0** | -0,2666677 | 0,11967826 |
| TRUA_HUM/ | PUS1 | sp\|Q9Y606\|TRUA_HUMAN tRNA pseudouridine synthase A OS=Homo sapiens GN=PUS1 | 42,971 | 2,81 | 47470 | 1 | 18,24 | 19,71 | 21,23 | 17,06 | 19,56 | 21,76 | **0,831** | 8,76E-01 | **0** | -0,2672494 | 0,05740209 |
| ARP5L_HUM. | ARPC5L | sp\|Q9BPX5\|ARP5L_HUMAN Actin-related protein 2/3 complex subunit 5-like protein OS | 32,197 | 8,5 | 16941 | 1 | 21,19 | 21,44 | 19,21 | 21,07 | 21,57 | 18,40 | **0,830** | 8,36E-01 | **0** | -0,2680753 | 0,07804495 |
| PLXA1_HUM. | PLXNA1 | sp\|Q9UIW2\|PLXA1_HUMAN Plexin-A1 OS=Homo sapiens GN=PLXNA1 PE=1 SV=3 | 540,089 | 13,29 | 211067 | 18 | 25,40 | 25,37 | 23,54 | 25,06 | 25,26 | 23,18 | **0,829** | 7,80E-01 | **0** | -0,2700005 | 0,1078691 |
| MSPD2_HUN | MOSPD2 | sp\|Q8NHP6\|MSPD2_HUMAN Motile sperm domain-containing protein 2 OS=Homo sapi | 34,709 | 2,9 | 59746 | 1 | 20,18 | 19,32 | 19,92 | 20,95 | 18,55 | 19,11 | **0,829** | 7,42E-01 | **0** | -0,2705498 | 0,12932594 |
| CAD11_HUM | CDH11 | sp\|P55287\|CAD11_HUMAN Cadherin-11 OS=Homo sapiens GN=CDH11 PE=2 SV=2 | 149,996 | 8,04 | 87965 | 5 | 24,80 | 24,48 | 21,30 | 24,21 | 23,88 | 21,67 | **0,827** | 8,52E-01 | **0** | -0,2733332 | 0,06963034 |
| RASH_HUMA | HRAS | sp\|P01112\|RASH_HUMAN GTPase HRas OS=Homo sapiens GN=HRAS PE=1 SV=1 | 543,447 | 64,55 | 21298 | 12 | 24,10 | 24,30 | 23,11 | 23,91 | 23,93 | 22,85 | **0,827** | 6,22E-01 | **0** | -0,2733332 | 0,20619854 |
| CADM4_HUN | CADM4 | sp\|Q8NFZ8\|CADM4_HUMAN Cell adhesion molecule 4 OS=Homo sapiens GN=CADM4 PI | 57,866 | 4,64 | 42785 | 1 | 22,01 | 22,16 | 21,50 | 21,59 | 22,09 | 21,17 | **0,827** | 4,57E-01 | **0** | -0,2733332 | 0,33977443 |
| NED4L_HUM | NEDD4L | sp\|Q96PU5\|NED4L_HUMAN E3 ubiquitin-protein ligase NEDD4-like OS=Homo sapiens G | 522,099 | 19,9 | 111932 | 15 | 25,87 | 25,43 | 25,08 | 25,71 | 25,30 | 24,55 | **0,827** | 5,41E-01 | **0** | -0,2733345 | 0,26690692 |
| 2A5G_HUM/ | PPP2R5C | sp\|Q13362\|2A5G_HUMAN Serine/threonine-protein phosphatase 2A 56 kDa regulatory | 74,109 | 9,35 | 61061 | 4 | 20,32 | 17,49 | 22,87 | 17,37 | 20,27 | 22,22 | **0,827** | 9,02E-01 | **0** | -0,2745209 | 0,04467664 |
| CLH1_HUMA | CLTC | sp\|Q00610\|CLH1_HUMAN Clathrin heavy chain 1 OS=Homo sapiens GN=CLTC PE=1 SV= | 7353,286 | 73,13 | 191615 | 140 | 31,14 | 31,16 | 31,98 | 31,02 | 31,06 | 31,37 | **0,825** | 4,06E-01 | **0** | -0,276666 | 0,39172682 |
| FADS2_HUM | FADS2 | sp\|O95864\|FADS2_HUMAN Fatty acid desaturase 2 OS=Homo sapiens GN=FADS2 PE=1 | 23,272 | 3,38 | 52259 | 1 | 21,51 | 20,96 | 20,85 | 21,06 | 20,82 | 20,61 | **0,825** | 3,17E-01 | **0** | -0,2766666 | 0,49927121 |
| RTN1_HUMA | RTN1 | sp\|Q16799\|RTN1_HUMAN Reticulon-1 OS=Homo sapiens GN=RTN1 PE=1 SV=1 | 10,614 | 1,03 | 83618 | 1 | 26,54 | 25,59 | 25,09 | 25,85 | 26,03 | 24,51 | **0,825** | 6,88E-01 | **0** | -0,2766666 | 0,16227419 |
| LAMB2_HUN | LAMB2 | sp\|P55268\|LAMB2_HUMAN Laminin subunit beta-2 OS=Homo sapiens GN=LAMB2 PE=1 | 40,709 | 0,83 | 195981 | 1 | 20,47 | 18,93 | 20,69 | 20,36 | 18,46 | 20,43 | **0,825** | 7,60E-01 | **0** | -0,2782288 | 0,11931 |
| CAVN3_HUN | CAVIN3 | sp\|Q969G5\|CAVN3_HUMAN Caveolae-associated protein 3 OS=Homo sapiens GN=CAVI | 38,959 | 8,05 | 27701 | 2 | 22,71 | 23,77 | 23,47 | 23,06 | 23,22 | 22,83 | **0,824** | 4,50E-01 | **0** | -0,2800001 | 0,34634608 |
| CHM1B_HUN | CHMP1B | sp\|Q7LBR1\|CHM1B_HUMAN Charged multivesicular body protein 1b OS=Homo sapiens | 55,502 | 8,54 | 22109 | 3 | 25,92 | 25,69 | 22,66 | 25,80 | 25,35 | 22,28 | **0,824** | 8,63E-01 | **0** | -0,2800001 | 0,06384702 |
| DEST_HUMA | DSTN | sp\|P60981\|DEST_HUMAN Destrin OS=Homo sapiens GN=DSTN PE=1 SV=3 | 908,408 | 80,61 | 18506 | 18 | 28,83 | 28,74 | 27,87 | 28,48 | 28,54 | 27,58 | **0,824** | 5,56E-01 | **0** | -0,2800001 | 0,25515799 |
| TYDP2_HUM | TDP2 | sp\|O95551\|TYDP2_HUMAN Tyrosyl-DNA phosphodiesterase 2 OS=Homo sapiens GN=TI | 28,231 | 3,04 | 40930 | 1 | 19,69 | 19,74 | 18,45 | 20,01 | 17,91 | 19,11 | **0,822** | 7,22E-01 | **0** | -0,2828979 | 0,14153923 |
| CRIP1_HUM/ | CRIP1 | sp\|P50238\|CRIP1_HUMAN Cysteine-rich protein 1 OS=Homo sapiens GN=CRIP1 PE=1 SV | 38,485 | 54,55 | 8533 | 3 | 25,35 | 25,39 | 21,14 | 24,93 | 24,67 | 21,43 | **0,822** | 8,83E-01 | **0** | -0,2833328 | 0,05412216 |
| PFD2_HUMA | PFDN2 | sp\|Q9UHV9\|PFD2_HUMAN Prefoldin subunit 2 OS=Homo sapiens GN=PFDN2 PE=1 SV=1 | 101,378 | 14,94 | 16648 | 2 | 22,86 | 23,23 | 22,89 | 22,76 | 22,67 | 22,70 | **0,822** | 8,02E-02 | **0** | -0,2833328 | 1,09589342 |
| FINC_HUMAI | FN1 | sp\|P02751\|FINC_HUMAN Fibronectin OS=Homo sapiens GN=FN1 PE=1 SV=4 | 1897,314 | 25,82 | 262607 | 49 | 28,03 | 27,20 | 29,27 | 27,92 | 26,95 | 28,78 | **0,822** | 7,41E-01 | **0** | -0,2833335 | 0,12999617 |
| ABI1_HUMAI | ABI1 | sp\|Q8IZP0\|ABI1_HUMAN Abl interactor 1 OS=Homo sapiens GN=ABI1 PE=1 SV=4 | 491,956 | 32,09 | 55081 | 13 | 26,76 | 26,65 | 25,65 | 26,28 | 26,13 | 25,80 | **0,822** | 4,98E-01 | **0** | -0,2833335 | 0,30289391 |
| PCLO_HUMA | PCLO | sp\|Q9Y6V0\|PCLO_HUMAN Protein piccolo OS=Homo sapiens GN=PCLO PE=1 SV=4 | 15,818 | 0,39 | 553277 | 2 | 27,64 | 25,83 | 29,86 | 26,23 | 26,96 | 29,29 | **0,822** | 8,58E-01 | **0** | -0,2833335 | 0,06645583 |
| 3HIDH_HUM | HIBADH | sp\|P31937\|3HIDH_HUMAN 3-hydroxyisobutyrate dehydrogenase, mitochondrial OS=Hc | 39,088 | 3,87 | 35329 | 1 | 17,84 | 20,11 | 21,14 | 20,46 | 17,72 | 20,06 | **0,821** | 8,36E-01 | **0** | -0,2854188 | 0,0775448 |
| LFG3_HUMA | TMBIM1 | sp\|Q969X1\|LFG3_HUMAN Protein lifeguard 3 OS=Homo sapiens GN=TMBIM1 PE=1 SV= | 132,741 | 8,68 | 34607 | 3 | 25,25 | 25,14 | 24,51 | 24,95 | 25,03 | 24,06 | **0,820** | 5,00E-01 | **0** | -0,2866662 | 0,30103962 |
| C2D1B_HUM | CC2D1B | sp\|Q5T0F9\|C2D1B_HUMAN Coiled-coil and C2 domain-containing protein 1B OS=Homo | 263,145 | 14,57 | 94224 | 9 | 25,12 | 24,72 | 23,31 | 24,78 | 24,68 | 22,83 | **0,820** | 7,50E-01 | **0** | -0,2866662 | 0,12512222 |
| ILK_HUMAN | ILK | sp\|Q13418\|ILK_HUMAN Integrin-linked protein kinase OS=Homo sapiens GN=ILK PE=1 | 446,978 | 36,28 | 51419 | 15 | 26,95 | 26,60 | 26,92 | 26,78 | 26,14 | 26,69 | **0,820** | 2,79E-01 | **0** | -0,2866669 | 0,55394369 |
| ITSN1_HUM/ | ITSN1 | sp\|Q15811\|ITSN1_HUMAN Intersectin-1 OS=Homo sapiens GN=ITSN1 PE=1 SV=3 | 13,99 | 0,7 | 195422 | 1 | 20,45 | 20,69 | 19,23 | 20,60 | 19,72 | 19,19 | **0,820** | 6,63E-01 | **0** | -0,2866669 | 0,17822296 |
| TPM1_HUM/ | TPM1 | sp\|P09493\|TPM1_HUMAN Tropomyosin alpha-1 chain OS=Homo sapiens GN=TPM1 PE= | 634,371 | 40,14 | 32709 | 18 | 24,77 | 25,01 | 22,43 | 24,37 | 24,93 | 22,05 | **0,820** | 8,24E-01 | **0** | -0,2866669 | 0,08418192 |
| ITA6_HUMAI | ITGA6 | sp\|P23229\|ITA6_HUMAN Integrin alpha-6 OS=Homo sapiens GN=ITGA6 PE=1 SV=5 | 718,268 | 18,85 | 126606 | 19 | 27,78 | 27,43 | 27,18 | 27,66 | 27,20 | 26,67 | **0,820** | 4,40E-01 | **0** | -0,2866669 | 0,35642151 |
| PPM1A_HUN | PPM1A | sp\|P35813\|PPM1A_HUMAN Protein phosphatase 1A OS=Homo sapiens GN=PPM1A PE= | 178,415 | 18,59 | 42448 | 6 | 23,45 | 23,84 | 23,54 | 23,46 | 23,39 | 23,12 | **0,820** | 1,42E-01 | **0** | -0,2866675 | 0,84810536 |
| NUDT4_HUN | NUDT4 | sp\|Q9NZJ9\|NUDT4_HUMAN Diphosphoinositol polyphosphate phosphohydrolase 2 OS= | 55,175 | 7,78 | 20306 | 1 | 18,90 | 20,00 | 20,67 | 19,04 | 18,95 | 20,72 | **0,818** | 7,27E-01 | **0** | -0,2891229 | 0,13825275 |
| TFIP8_HUM/ | TNFAIP8 | sp\|O95379\|TFIP8_HUMAN Tumor necrosis factor alpha-induced protein 8 OS=Homo sa | 75,959 | 10,1 | 23003 | 1 | 20,17 | 19,39 | 20,31 | 19,86 | 19,75 | 19,39 | **0,818** | 4,15E-01 | **0** | -0,2899996 | 0,38159537 |
| SH3B4_HUM | SH3BP4 | sp\|Q9P0V3\|SH3B4_HUMAN SH3 domain-binding protein 4 OS=Homo sapiens GN=SH3BI | 91,121 | 5,19 | 107496 | 4 | 23,30 | 23,47 | 21,10 | 23,19 | 22,43 | 21,38 | **0,818** | 7,70E-01 | **0** | -0,2899996 | 0,11359321 |
| RAB23_HUM | RAB23 | sp\|Q9ULC3\|RAB23_HUMAN Ras-related protein Rab-23 OS=Homo sapiens GN=RAB23 P | 371,648 | 40,51 | 26659 | 9 | 26,80 | 26,71 | 26,17 | 26,56 | 26,49 | 25,76 | **0,818** | 4,20E-01 | **0** | -0,2899996 | 0,37713116 |
| THBG_HUM/ | SERPINA7 | sp\|P05543\|THBG_HUMAN Thyroxine-binding globulin OS=Homo sapiens GN=SERPINA7 | 65,915 | 6,27 | 46325 | 3 | 23,26 | 21,35 | 24,75 | 23,20 | 21,60 | 23,69 | **0,818** | 8,16E-01 | **0** | -0,2899996 | 0,08815812 |
| CASC3_HUM | CASC3 | sp\|O15234\|CASC3_HUMAN Protein CASC3 OS=Homo sapiens GN=CASC3 PE=1 SV=2 | 137,253 | 9,67 | 76278 | 6 | 24,86 | 24,25 | 24,25 | 24,85 | 23,87 | 23,77 | **0,818** | 5,09E-01 | **0** | -0,2899996 | 0,29357089 |
| ZO2_HUMAN | TJP2 | sp\|Q9UDY2\|ZO2_HUMAN Tight junction protein ZO-2 OS=Homo sapiens GN=TJP2 PE=1 | 302,017 | 10,42 | 133958 | 11 | 24,76 | 24,30 | 24,53 | 24,13 | 24,29 | 24,30 | **0,818** | 1,14E-01 | **0** | -0,2900003 | 0,94365728 |
| KAPCG_HUN | PRKACG | sp\|P22612\|KAPCG_HUMAN cAMP-dependent protein kinase catalytic subunit gamma O | 65,984 | 9,4 | 40434 | 2 | 22,28 | 22,62 | 22,95 | 21,74 | 22,81 | 22,43 | **0,818** | 4,75E-01 | **0** | -0,2900009 | 0,32344909 |
| MYL9_HUM/ | MYL9 | sp\|P24844\|MYL9_HUMAN Myosin regulatory light polypeptide 9 OS=Homo sapiens GN= | 430,279 | 55,81 | 19827 | 9 | 24,99 | 25,23 | 24,39 | 24,77 | 25,03 | 23,93 | **0,816** | 5,19E-01 | **0** | -0,2933324 | 0,28476565 |
| CHMP6_HUN | CHMP6 | sp\|Q96FZ7\|CHMP6_HUMAN Charged multivesicular body protein 6 OS=Homo sapiens G | 119,646 | 11,94 | 23485 | 3 | 23,16 | 23,28 | 21,80 | 22,86 | 22,81 | 21,69 | **0,816** | 6,55E-01 | **0** | -0,2933331 | 0,18355079 |
| RHOQ_HUM. | RHOQ | sp\|P17081\|RHOQ_HUMAN Rho-related GTP-binding protein RhoQ OS=Homo sapiens GI | 104,419 | 12,2 | 22659 | 2 | 18,67 | 18,66 | 19,05 | 18,31 | 18,44 | 18,75 | **0,816** | 1,84E-01 | **0** | -0,2933331 | 0,73440256 |
| MOB1A_HUN | MOB1A | sp\|Q9H8S9\|MOB1A_HUMAN MOB kinase activator 1A OS=Homo sapiens GN=MOB1A PI | 155,257 | 39,35 | 25080 | 7 | 25,72 | 25,38 | 24,03 | 25,30 | 25,18 | 23,77 | **0,816** | 7,02E-01 | **0** | -0,2933331 | 0,15387083 |
| PLAK_HUMA | JUP | sp\|P14923\|PLAK_HUMAN Junction plakoglobin OS=Homo sapiens GN=JUP PE=1 SV=3 | 1602,76 | 67,52 | 81745 | 36 | 28,77 | 28,73 | 26,11 | 28,49 | 28,31 | 25,93 | **0,816** | 8,20E-01 | **0** | -0,2933337 | 0,08627445 |
| DIAP1_HUM. | DIAPH1 | sp\|O60610\|DIAP1_HUMAN Protein diaphanous homolog 1 OS=Homo sapiens GN=DIAPH | 417,189 | 15,57 | 141347 | 14 | 24,87 | 26,18 | 23,00 | 24,72 | 24,91 | 23,54 | **0,816** | 7,87E-01 | **0** | -0,2933337 | 0,10379281 |
| MINK1_HUM | MINK1 | sp\|Q8N4C8\|MINK1_HUMAN Misshapen-like kinase 1 OS=Homo sapiens GN=MINK1 PE= | 407,988 | 12,24 | 149822 | 15 | 22,20 | 21,74 | 23,18 | 22,75 | 21,03 | 22,46 | **0,816** | 6,89E-01 | **0** | -0,2933337 | 0,16203376 |
| CDV3_HUM/ | CDV3 | sp\|Q9UKY7\|CDV3_HUMAN Protein CDV3 homolog OS=Homo sapiens GN=CDV3 PE=1 SV | 20 | 3,1 | 27335 | 1 | 21,92 | 22,13 | 20,34 | 22,33 | 22,30 | 18,88 | **0,816** | 8,30E-01 | **0** | -0,2934424 | 0,0811757 |
| RABP2_HUM | CRABP2 | sp\|P29373\|RABP2_HUMAN Cellular retinoic acid-binding protein 2 OS=Homo sapiens GI | 150,904 | 26,81 | 15693 | 3 | 26,07 | 26,05 | 17,65 | 25,76 | 26,04 | 17,09 | **0,815** | 9,45E-01 | **0** | -0,2956346 | 0,02435511 |
| ABCD3_HUM | ABCD3 | sp\|P28288\|ABCD3_HUMAN ATP-binding cassette sub-family D member 3 OS=Homo sap | 32,985 | 2,88 | 75476 | 1 | 18,20 | 18,56 | 20,78 | 18,12 | 18,83 | 19,70 | **0,815** | 7,66E-01 | **0** | -0,2959474 | 0,1160127 |
| AP2A1_HUM | AP2A1 | sp\|O95782\|AP2A1_HUMAN AP-2 complex subunit alpha-1 OS=Homo sapiens GN=AP2A1 | 676,283 | 27,23 | 107546 | 20 | 25,78 | 25,13 | 25,89 | 25,27 | 25,44 | 25,20 | **0,814** | 2,97E-01 | **0** | -0,2966658 | 0,52723124 |
| PTPRF_HUM. | PTPRF | sp\|P10586\|PTPRF_HUMAN Receptor-type tyrosine-protein phosphatase F OS=Homo sa | 120,48 | 4,93 | 212879 | 6 | 23,67 | 23,15 | 22,32 | 23,12 | 23,25 | 21,88 | **0,814** | 6,40E-01 | **0** | -0,2966665 | 0,19372136 |
| RTN3_HUMA | RTN3 | sp\|O95197\|RTN3_HUMAN Reticulon-3 OS=Homo sapiens GN=RTN3 PE=1 SV=2 | 114,355 | 2,03 | 112611 | 3 | 27,48 | 27,02 | 26,44 | 26,98 | 27,29 | 25,78 | **0,814** | 6,18E-01 | **0** | -0,2966665 | 0,20882657 |
| HGS_HUMAN | HGS | sp\|O14964\|HGS_HUMAN Hepatocyte growth factor-regulated tyrosine kinase substrate | 444,583 | 18,28 | 86192 | 14 | 27,04 | 26,78 | 25,59 | 26,75 | 26,40 | 25,37 | **0,814** | 6,52E-01 | **0** | -0,2966671 | 0,18601431 |
| RBGP1_HUM | RABGAP1 | sp\|Q9Y3P9\|RBGP1_HUMAN Rab GTPase-activating protein 1 OS=Homo sapiens GN=RAE | 29,778 | 1,12 | 121737 | 1 | 18,83 | 17,33 | 20,28 | 17,77 | 17,80 | 19,97 | **0,812** | 8,02E-01 | **0** | -0,2999776 | 0,09581307 |
| NCK1_HUM/ | NCK1 | sp\|P16333\|NCK1_HUMAN Cytoplasmic protein NCK1 OS=Homo sapiens GN=NCK1 PE=1 | 66,759 | 8,49 | 42864 | 3 | 22,16 | 21,99 | 21,88 | 22,51 | 21,61 | 21,01 | **0,812** | 5,36E-01 | **0** | -0,2999992 | 0,27099543 |
| SCAM2_HUN | SCAMP2 | sp\|O15127\|SCAM2_HUMAN Secretory carrier-associated membrane protein 2 OS=Hom | 22,353 | 3,65 | 36649 | 1 | 23,37 | 23,12 | 23,48 | 23,12 | 23,34 | 22,61 | **0,812** | 2,81E-01 | **0** | -0,2999999 | 0,55101839 |
| LSM12_HUM | LSM12 | sp\|Q3MHD2\|LSM12_HUMAN Protein LSM12 homolog OS=Homo sapiens GN=LSM12 PE | 153,268 | 12,31 | 21701 | 3 | 22,17 | 22,03 | 23,38 | 21,98 | 21,95 | 22,75 | **0,812** | 5,82E-01 | **0** | -0,2999999 | 0,23474569 |
| ULA1_HUMA | NAE1 | sp\|Q13564\|ULA1_HUMAN NEDD8-activating enzyme E1 regulatory subunit OS=Homo s | 46,11 | 3,56 | 60246 | 2 | 23,41 | 23,00 | 22,59 | 22,77 | 22,33 | 23,00 | **0,812** | 3,85E-01 | **0** | -0,2999999 | 0,41482031 |
| RENT1_HUM | UPF1 | sp\|Q92900\|RENT1_HUMAN Regulator of nonsense transcripts 1 OS=Homo sapiens GN= | 1560,61 | 41,45 | 124345 | 41 | 28,21 | 28,24 | 27,53 | 28,01 | 27,82 | 27,25 | **0,812** | 4,09E-01 | **0** | -0,2999999 | 0,38855196 |
| LIPL_HUMAN | LPL | sp\|P06858\|LIPL_HUMAN Lipoprotein lipase OS=Homo sapiens GN=LPL PE=1 SV=1 | 123,676 | 11,16 | 53162 | 4 | 20,20 | 21,40 | 24,53 | 20,61 | 20,65 | 23,97 | **0,812** | 8,69E-01 | **0** | -0,3000005 | 0,0610573 |
| VIME_HUMA | VIM | sp\|P08670\|VIME_HUMAN Vimentin OS=Homo sapiens GN=VIM PE=1 SV=4 | 1780,092 | 54,72 | 53652 | 42 | 28,99 | 29,60 | 30,33 | 28,88 | 29,18 | 29,96 | **0,812** | 5,83E-01 | **0** | -0,3000005 | 0,23399472 |
| TOM70_HUN | TOMM70 | sp\|O94826\|TOM70_HUMAN Mitochondrial import receptor subunit TOM70 OS=Homo | 301,282 | 16,28 | 67455 | 8 | 23,93 | 24,28 | 24,65 | 24,09 | 23,82 | 24,05 | **0,812** | 2,52E-01 | **0** | -0,3000005 | 0,59866813 |
| VPS45_HUM. | VPS45 | sp\|Q9NRW7\|VPS45_HUMAN Vacuolar protein sorting-associated protein 45 OS=Homo | 66,514 | 5,96 | 65077 | 3 | 22,16 | 23,89 | 21,63 | 21,66 | 21,98 | 23,13 | **0,810** | 7,29E-01 | **0** | -0,3033333 | 0,13742042 |
| CHP1_HUMA | CHP1 | sp\|Q99653\|CHP1_HUMAN Calcineurin B homologous protein 1 OS=Homo sapiens GN=C | 145,374 | 25,13 | 22456 | 4 | 21,66 | 23,06 | 23,03 | 21,49 | 22,19 | 23,16 | **0,810** | 6,74E-01 | **0** | -0,3033333 | 0,17147673 |
| GP1BB_HUM | GP1BB | sp\|P13224\|GP1BB_HUMAN Platelet glycoprotein Ib beta chain OS=Homo sapiens GN=GI | 39,954 | 9,71 | 21718 | 2 | 25,62 | 24,66 | 24,49 | 25,52 | 24,55 | 23,79 | **0,810** | 6,46E-01 | **0** | -0,3033333 | 0,18971978 |
| GPC4_HUMA | GPC4 | sp\|O75487\|GPC4_HUMAN Glypican-4 OS=Homo sapiens GN=GPC4 PE=1 SV=4 | 182,158 | 7,19 | 62412 | 4 | 21,99 | 21,69 | 21,91 | 22,08 | 21,38 | 21,22 | **0,810** | 3,38E-01 | **0** | -0,3033339 | 0,47121817 |
| ZFN2B_HUM | ZFAND2B | sp\|Q8WV99\|ZFN2B_HUMAN AN1-type zinc finger protein 2B OS=Homo sapiens GN=ZFA | 73,73 | 12,84 | 28023 | 3 | 23,61 | 23,42 | 22,87 | 23,34 | 23,13 | 22,52 | **0,810** | 4,12E-01 | **0** | -0,3033339 | 0,38553296 |
| IMA4_HUMA | KPNA3 | sp\|O00505\|IMA4_HUMAN Importin subunit alpha-4 OS=Homo sapiens GN=KPNA3 PE=1 | 343,612 | 30,9 | 57811 | 9 | 23,82 | 24,10 | 24,70 | 23,27 | 24,06 | 24,38 | **0,810** | 5,10E-01 | **0** | -0,3033339 | 0,29252855 |
| FKB1A_HUM | FKBP1A | sp\|P62942\|FKB1A_HUMAN Peptidyl-prolyl cis-trans isomerase FKBP1A OS=Homo sapier | 505,479 | 73,15 | 11951 | 11 | 28,09 | 28,12 | 26,11 | 27,96 | 27,81 | 25,64 | **0,810** | 7,77E-01 | **0** | -0,3033346 | 0,10946448 |
| DIP2B_HUM. | DIP2B | sp\|Q9P265\|DIP2B_HUMAN Disco-interacting protein 2 homolog B OS=Homo sapiens GN | 1040,926 | 28,62 | 171492 | 36 | 27,64 | 27,07 | 25,40 | 27,32 | 26,87 | 25,00 | **0,809** | 7,70E-01 | **0** | -0,3066661 | 0,11379127 |
| SNF8_HUMA | SNF8 | sp\|Q96H20\|SNF8_HUMAN Vacuolar-sorting protein SNF8 OS=Homo sapiens GN=SNF8 P | 212,071 | 27,52 | 28864 | 5 | 24,71 | 24,46 | 22,25 | 24,53 | 24,34 | 21,63 | **0,809** | 8,14E-01 | **0** | -0,3066661 | 0,08942496 |
| LARP1_HUM. | LARP1 | sp\|Q6PKG0\|LARP1_HUMAN La-related protein 1 OS=Homo sapiens GN=LARP1 PE=1 SV= | 228,338 | 7,3 | 123510 | 7 | 23,69 | 23,26 | 24,14 | 23,22 | 23,09 | 23,86 | **0,809** | 4,28E-01 | **0** | -0,3066667 | 0,36846236 |
| ATLA2_HUM | ATL2 | sp\|Q8NHH9\|ATLA2_HUMAN Atlastin-2 OS=Homo sapiens GN=ATL2 PE=1 SV=2 | 102,722 | 10,29 | 66229 | 5 | 22,89 | 23,35 | 22,75 | 22,62 | 23,22 | 22,23 | **0,809** | 4,18E-01 | **0** | -0,3066667 | 0,37846566 |
| YBOX1_HUM | YBX1 | sp\|P67809\|YBOX1_HUMAN Nuclease-sensitive element-binding protein 1 OS=Homo sa | 816,604 | 51,23 | 35924 | 15 | 26,15 | 26,29 | 26,24 | 25,98 | 25,77 | 26,01 | **0,809** | 2,34E-02 | **0** | -0,3066667 | 1,63134804 |
| TADBP_HUM | TARDBP | sp\|Q13148\|TADBP_HUMAN TAR DNA-binding protein 43 OS=Homo sapiens GN=TARDBI | 269,522 | 26,33 | 44740 | 7 | 24,66 | 24,68 | 25,84 | 24,37 | 24,23 | 25,66 | **0,809** | 6,36E-01 | **0** | -0,3066667 | 0,19663622 |
| HD_HUMAN | HTT | sp\|P42858\|HD_HUMAN Huntingtin OS=Homo sapiens GN=HTT PE=1 SV=2 | 19,335 | 0,48 | 347603 | 1 | 17,81 | 18,23 | 18,91 | 17,91 | 18,02 | 18,10 | **0,808** | 3,97E-01 | **0** | -0,3068129 | 0,4007362 |
| KHNYN_HUN | KHNYN | sp\|O15037\|KHNYN_HUMAN Protein KHNYN OS=Homo sapiens GN=KHNYN PE=1 SV=3 | 53,703 | 2,36 | 74534 | 1 | 21,24 | 20,62 | 19,12 | 20,92 | 20,41 | 18,73 | **0,807** | 7,52E-01 | **0** | -0,3085918 | 0,12368282 |
| DPOLA_HUN | POLA1 | sp\|P09884\|DPOLA_HUMAN DNA polymerase alpha catalytic subunit OS=Homo sapiens ( | 26,929 | 0,62 | 165913 | 1 | 19,21 | 19,00 | 21,59 | 20,03 | 17,56 | 21,28 | **0,807** | 8,33E-01 | **0** | -0,3089269 | 0,07938175 |
| C2C2L_HUM. | C2CD2L | sp\|O14523\|C2C2L_HUMAN Phospholipid transfer protein C2CD2L OS=Homo sapiens GN | 31,435 | 2,83 | 76181 | 2 | 21,67 | 21,41 | 27,30 | 21,53 | 21,43 | 26,49 | **0,807** | 9,09E-01 | **0** | -0,3099995 | 0,04145573 |
| QORL1_HUN | CRYZL1 | sp\|O95825\|QORL1_HUMAN Quinone oxidoreductase-like protein 1 OS=Homo sapiens G | 14,235 | 3,44 | 38697 | 1 | 18,55 | 18,75 | 19,10 | 18,68 | 18,10 | 18,69 | **0,807** | 2,87E-01 | **0** | -0,3099995 | 0,54178888 |
| FMNL3_HUN | FMNL3 | sp\|Q8IVF7\|FMNL3_HUMAN Formin-like protein 3 OS=Homo sapiens GN=FMNL3 PE=1 S' | 326,981 | 12,94 | 117213 | 11 | 24,26 | 24,12 | 22,22 | 24,25 | 23,61 | 21,81 | **0,807** | 7,68E-01 | **0** | -0,3100001 | 0,11449022 |
| LYPA1_HUM. | LYPLA1 | sp\|O75608\|LYPA1_HUMAN Acyl-protein thioesterase 1 OS=Homo sapiens GN=LYPLA1 P | 66,419 | 19,57 | 24670 | 3 | 24,95 | 24,86 | 24,19 | 24,78 | 24,66 | 23,63 | **0,807** | 5,17E-01 | **0** | -0,3100007 | 0,28654526 |
| PIN1_HUMAI | PIN1 | sp\|Q13526\|PIN1_HUMAN Peptidyl-prolyl cis-trans isomerase NIMA-interacting 1 OS=Ho | 342,103 | 52,76 | 18243 | 6 | 25,56 | 25,78 | 25,20 | 25,30 | 25,43 | 24,88 | **0,807** | 2,61E-01 | **0** | -0,3100007 | 0,58368722 |
| CTR2_HUMA | SLC7A2 | sp\|P52569\|CTR2_HUMAN Cationic amino acid transporter 2 OS=Homo sapiens GN=SLC7 | 79,844 | 7,14 | 71673 | 4 | 23,90 | 24,05 | 19,07 | 23,37 | 23,42 | 19,30 | **0,806** | 8,91E-01 | **0** | -0,3113155 | 0,05015483 |
| ZW10_HUM/ | ZW10 | sp\|O43264\|ZW10_HUMAN Centromere/kinetochore protein zw10 homolog OS=Homo | 35,15 | 1,8 | 88829 | 1 | 17,55 | 18,92 | 20,28 | 17,22 | 18,09 | 20,51 | **0,806** | 8,17E-01 | **0** | -0,3120035 | 0,08787666 |
| SYHM_HUM/ | HARS2 | sp\|P49590\|SYHM_HUMAN Probable histidine--tRNA ligase, mitochondrial OS=Homo sap | 86,435 | 7,51 | 56888 | 4 | 22,63 | 22,96 | 19,90 | 22,68 | 22,47 | 19,40 | **0,805** | 8,38E-01 | **0** | -0,3127511 | 0,07667244 |
| NCKP1_HUM | NCKAP1 | sp\|Q9Y2A7\|NCKP1_HUMAN Nck-associated protein 1 OS=Homo sapiens GN=NCKAP1 PE | 1378,343 | 40,78 | 128790 | 40 | 28,05 | 27,60 | 27,05 | 27,38 | 27,45 | 26,93 | **0,805** | 3,99E-01 | **0** | -0,3133329 | 0,39947281 |
| VA0D1_HUM | ATP6V0D1 | sp\|P61421\|VA0D1_HUMAN V-type proton ATPase subunit d 1 OS=Homo sapiens GN=AT | 197,2 | 36,18 | 40329 | 7 | 25,28 | 25,28 | 25,72 | 24,94 | 25,25 | 25,15 | **0,805** | 1,44E-01 | **0** | -0,3133335 | 0,8417106 |
| PLXB2_HUM. | PLXNB2 | sp\|O15031\|PLXB2_HUMAN Plexin-B2 OS=Homo sapiens GN=PLXNB2 PE=1 SV=3 | 933,028 | 16,05 | 205127 | 25 | 28,24 | 28,25 | 26,40 | 27,88 | 27,92 | 26,15 | **0,805** | 7,30E-01 | **0** | -0,3133335 | 0,13643594 |
| FA5_HUMAN | F5 | sp\|P12259\|FA5_HUMAN Coagulation factor V OS=Homo sapiens GN=F5 PE=1 SV=4 | 477,52 | 5,44 | 251703 | 13 | 27,18 | 26,94 | 27,37 | 27,07 | 26,77 | 26,71 | **0,805** | 1,34E-01 | **0** | -0,3133341 | 0,87353895 |
| DNJB6_HUM | DNAJB6 | sp\|O75190\|DNJB6_HUMAN DnaJ homolog subfamily B member 6 OS=Homo sapiens GN | 62,2 | 6,44 | 36087 | 2 | 22,05 | 21,81 | 18,89 | 22,00 | 21,54 | 18,26 | **0,803** | 8,49E-01 | **0** | -0,3160013 | 0,07110141 |
| LDOC1_HUM | LDOC1 | sp\|O95751\|LDOC1_HUMAN Protein LDOC1 OS=Homo sapiens GN=LDOC1 PE=1 SV=1 | 108,409 | 23,97 | 16968 | 2 | 23,74 | 23,38 | 23,01 | 23,37 | 23,54 | 22,27 | **0,803** | 5,21E-01 | **0** | -0,3166656 | 0,28336355 |
| PDC6I_HUM/ | PDCD6IP | sp\|Q8WUM4\|PDC6I_HUMAN Programmed cell death 6-interacting protein OS=Homo sa | 3111,818 | 57,14 | 96023 | 59 | 32,90 | 32,54 | 31,91 | 32,72 | 32,33 | 31,35 | **0,803** | 5,61E-01 | **0** | -0,3166663 | 0,25122766 |
| CAV1_HUMA | CAV1 | sp\|Q03135\|CAV1_HUMAN Caveolin-1 OS=Homo sapiens GN=CAV1 PE=1 SV=4 | 289,788 | 41,57 | 20472 | 8 | 27,73 | 28,06 | 28,98 | 27,56 | 27,54 | 28,72 | **0,803** | 5,89E-01 | **0** | -0,3166663 | 0,22963198 |
| MLF2_HUM/ | MLF2 | sp\|Q15773\|MLF2_HUMAN Myeloid leukemia factor 2 OS=Homo sapiens GN=MLF2 PE=1 | 74,34 | 14,92 | 28147 | 3 | 22,49 | 23,39 | 21,99 | 23,30 | 21,93 | 21,69 | **0,803** | 6,50E-01 | **0** | -0,3166663 | 0,18681729 |
| ZO1_HUMAN | TJP1 | sp\|Q07157\|ZO1_HUMAN Tight junction protein ZO-1 OS=Homo sapiens GN=TJP1 PE=1 | 587,327 | 11,67 | 195459 | 16 | 25,69 | 25,79 | 25,42 | 25,49 | 25,45 | 25,01 | **0,803** | 1,70E-01 | **0** | -0,3166669 | 0,77012752 |
| DPOD2_HUN | POLD2 | sp\|P49005\|DPOD2_HUMAN DNA polymerase delta subunit 2 OS=Homo sapiens GN=POI | 12,406 | 3,2 | 51289 | 1 | 20,99 | 21,16 | 20,54 | 20,46 | 21,19 | 20,09 | **0,803** | 4,43E-01 | **0** | -0,3166669 | 0,35361183 |
| UAP1_HUM/ | UAP1 | sp\|Q16222\|UAP1_HUMAN UDP-N-acetylhexosamine pyrophosphorylase OS=Homo sapi | 152,16 | 13,22 | 58769 | 6 | 23,21 | 23,45 | 23,92 | 23,46 | 23,85 | 22,32 | **0,803** | 5,64E-01 | **0** | -0,3166669 | 0,24871174 |
| B3A2_HUMA | SLC4A2 | sp\|P04920\|B3A2_HUMAN Anion exchange protein 2 OS=Homo sapiens GN=SLC4A2 PE= | 21,095 | 0,81 | 137009 | 1 | 20,50 | 20,61 | 17,72 | 20,38 | 20,31 | 17,19 | **0,803** | 8,34E-01 | **0** | -0,3166669 | 0,07893211 |
| PKHO2_HUN | PLEKHO2 | sp\|Q8TD55\|PKHO2_HUMAN Pleckstrin homology domain-containing family O member | 74,868 | 6,33 | 53350 | 3 | 23,04 | 23,11 | 21,29 | 23,48 | 22,77 | 20,24 | **0,803** | 7,97E-01 | **0** | -0,3166676 | 0,09876906 |
| PCDGB_HUN | PCDHGA11 | sp\|Q9Y5H2\|PCDGB_HUMAN Protocadherin gamma-A11 OS=Homo sapiens GN=PCDHGA | 117,919 | 4,71 | 101543 | 3 | 22,55 | 22,38 | 19,41 | 22,83 | 22,09 | 18,47 | **0,803** | 8,60E-01 | **0** | -0,3171298 | 0,06536436 |
| FMNL1_HUN | FMNL1 | sp\|O95466\|FMNL1_HUMAN Formin-like protein 1 OS=Homo sapiens GN=FMNL1 PE=1 S | 29,834 | 1,09 | 121854 | 1 | 21,23 | 20,63 | 20,33 | 21,05 | 21,33 | 18,85 | **0,802** | 7,20E-01 | **0** | -0,318353 | 0,14292926 |
| RECK_HUMA | RECK | sp\|O95980\|RECK_HUMAN Reversion-inducing cysteine-rich protein with Kazal motifs O | 220,702 | 6,69 | 106457 | 5 | 23,42 | 24,07 | 22,46 | 23,23 | 23,61 | 22,15 | **0,801** | 6,43E-01 | **0** | -0,3199997 | 0,19148039 |
| PCBP2_HUM | PCBP2 | sp\|Q15366\|PCBP2_HUMAN Poly(rC)-binding protein 2 OS=Homo sapiens GN=PCBP2 PE= | 1103,637 | 62,47 | 38580 | 22 | 28,88 | 28,84 | 29,40 | 28,57 | 28,56 | 29,03 | **0,801** | 2,50E-01 | **0** | -0,3199997 | 0,60260196 |
| IST1_HUMAN | IST1 | sp\|P53990\|IST1_HUMAN IST1 homolog OS=Homo sapiens GN=IST1 PE=1 SV=1 | 369,689 | 27,47 | 39751 | 11 | 28,75 | 28,44 | 26,95 | 28,53 | 28,12 | 26,53 | **0,801** | 7,18E-01 | **0** | -0,3199997 | 0,14395081 |
| HYOU1_HUN | HYOU1 | sp\|Q9Y4L1\|HYOU1_HUMAN Hypoxia up-regulated protein 1 OS=Homo sapiens GN=HYC | 359,941 | 13,61 | 111335 | 8 | 24,44 | 24,07 | 25,08 | 23,94 | 24,17 | 24,52 | **0,801** | 4,00E-01 | **0** | -0,3199997 | 0,39831256 |
| AT11B_HUM | ATP11B | sp\|Q9Y2G3\|AT11B_HUMAN Probable phospholipid-transporting ATPase IF OS=Homo sa | 14,669 | 0,59 | 134190 | 1 | 22,13 | 22,71 | 21,82 | 22,14 | 22,30 | 21,26 | **0,801** | 4,84E-01 | **0** | -0,3199997 | 0,31508031 |
| AT8B3_HUM | ATP8B3 | sp\|O60423\|AT8B3_HUMAN Phospholipid-transporting ATPase IK OS=Homo sapiens GN= | 14,669 | 0,54 | 146752 | 1 | 22,13 | 22,71 | 21,82 | 22,14 | 22,30 | 21,26 | **0,801** | 4,84E-01 | **0** | -0,3199997 | 0,31508031 |
| IGL1_HUMAN | | sp\|P0DOX8\|IGL1_HUMAN Immunoglobulin lambda-1 light chain OS=Homo sapiens PE=1 | 23,636 | 3,7 | 22830 | 1 | 24,27 | 22,46 | 23,81 | 24,39 | 22,02 | 23,17 | **0,801** | 7,33E-01 | **0** | -0,3199997 | 0,1350733 |
| PLS3_HUMAI | PLSCR3 | sp\|Q9NRY6\|PLS3_HUMAN Phospholipid scramblase 3 OS=Homo sapiens GN=PLSCR3 PE | 269,139 | 31,19 | 31648 | 7 | 27,93 | 27,44 | 27,00 | 27,58 | 27,36 | 26,47 | **0,801** | 5,01E-01 | **0** | -0,3200003 | 0,30042917 |
| RB27A_HUM | RAB27A | sp\|P51159\|RB27A_HUMAN Ras-related protein Rab-27A OS=Homo sapiens GN=RAB27A | 58,283 | 18,55 | 24868 | 4 | 23,85 | 22,70 | 22,40 | 23,56 | 22,10 | 22,33 | **0,801** | 6,40E-01 | **0** | -0,3200003 | 0,19397502 |
| CO4A_HUM/ | C4A | sp\|P0C0L4\|CO4A_HUMAN Complement C4-A OS=Homo sapiens GN=C4A PE=1 SV=2 | 281,265 | 5,68 | 192785 | 10 | 27,00 | 25,01 | 27,29 | 27,21 | 24,81 | 26,32 | **0,801** | 7,65E-01 | **0** | -0,320001 | 0,11608198 |
| ARF3_HUMA | ARF3 | sp\|P61204\|ARF3_HUMAN ADP-ribosylation factor 3 OS=Homo sapiens GN=ARF3 PE=1 S' | 925,55 | 80,11 | 20601 | 20 | 20,60 | 21,44 | 17,81 | 20,44 | 20,70 | 17,75 | **0,801** | 8,36E-01 | **0** | -0,320797 | 0,07803016 |
| UBQL2_HUN | UBQLN2 | sp\|Q9UHD9\|UBQL2_HUMAN Ubiquilin-2 OS=Homo sapiens GN=UBQLN2 PE=1 SV=2 | 246,164 | 16,03 | 65696 | 7 | 23,44 | 23,40 | 22,30 | 23,42 | 23,54 | 21,21 | **0,799** | 7,21E-01 | **0** | -0,3233331 | 0,14187997 |
| XPR1_HUMA | XPR1 | sp\|Q9UBH6\|XPR1_HUMAN Xenotropic and polytropic retrovirus receptor 1 OS=Homo s | 12,669 | 1,72 | 81535 | 1 | 18,93 | 19,21 | 18,87 | 19,15 | 18,33 | 18,56 | **0,799** | 2,91E-01 | **0** | -0,3233337 | 0,53675237 |
| DDR1_HUM/ | DDR1 | sp\|Q08345\|DDR1_HUMAN Epithelial discoidin domain-containing receptor 1 OS=Homo | 18,631 | 1,1 | 101128 | 1 | 20,73 | 20,57 | 18,92 | 20,53 | 20,02 | 18,70 | **0,799** | 7,05E-01 | **0** | -0,323892 | 0,15190318 |
| IFT20_HUM/ | IFT20 | sp\|Q8IY31\|IFT20_HUMAN Intraflagellar transport protein 20 homolog OS=Homo sapien | 145,781 | 33,33 | 15281 | 3 | 17,95 | 19,67 | 24,37 | 21,15 | 17,09 | 22,77 | **0,798** | 9,05E-01 | **0** | -0,3263486 | 0,04351982 |
| 6PGL_HUMA | PGLS | sp\|O95336\|6PGL_HUMAN 6-phosphogluconolactonase OS=Homo sapiens GN=PGLS PE= | 115,29 | 16,67 | 27547 | 3 | 22,48 | 22,97 | 23,42 | 21,93 | 22,35 | 23,61 | **0,797** | 5,99E-01 | **0** | -0,3266659 | 0,22243855 |
| ITIH1_HUMA | ITIH1 | sp\|P19827\|ITIH1_HUMAN Inter-alpha-trypsin inhibitor heavy chain H1 OS=Homo sapien | 63,108 | 2,41 | 101389 | 2 | 23,38 | 21,63 | 23,01 | 23,20 | 21,41 | 22,43 | **0,797** | 6,83E-01 | **0** | -0,3266659 | 0,1656056 |
| MK08_HUM/ | MAPK8 | sp\|P45983\|MK08_HUMAN Mitogen-activated protein kinase 8 OS=Homo sapiens GN=M | 60,476 | 6,79 | 48296 | 3 | 19,62 | 19,55 | 21,57 | 19,30 | 18,70 | 21,76 | **0,797** | 7,90E-01 | **0** | -0,3266665 | 0,10245177 |
| CAB39_HUM | CAB39 | sp\|Q9Y376\|CAB39_HUMAN Calcium-binding protein 39 OS=Homo sapiens GN=CAB39 PI | 254,754 | 33,72 | 39869 | 10 | 25,81 | 25,48 | 25,02 | 25,27 | 25,58 | 24,48 | **0,797** | 4,60E-01 | **0** | -0,3266665 | 0,3376316 |
| TBC8B_HUM | TBC1D8B | sp\|Q0IIM8\|TBC8B_HUMAN TBC1 domain family member 8B OS=Homo sapiens GN=TBC | 32,429 | 0,62 | 128709 | 1 | 24,84 | 24,80 | 24,22 | 24,71 | 24,26 | 23,91 | **0,797** | 3,46E-01 | **0** | -0,3266665 | 0,46077006 |
| TRFL_HUMA | LTF | sp\|P02788\|TRFL_HUMAN Lactotransferrin OS=Homo sapiens GN=LTF PE=1 SV=6 | 42,709 | 2,96 | 78182 | 2 | 24,99 | 24,72 | 25,30 | 25,07 | 24,34 | 24,61 | **0,796** | 2,90E-01 | **0** | -0,3299993 | 0,53702038 |
| CD9_HUMAN | CD9 | sp\|P21926\|CD9_HUMAN CD9 antigen OS=Homo sapiens GN=CD9 PE=1 SV=4 | 123,134 | 9,65 | 25416 | 3 | 28,88 | 28,12 | 27,73 | 27,66 | 28,54 | 27,54 | **0,796** | 5,14E-01 | **0** | -0,3299993 | 0,28864118 |
| NCAM1_HUN | NCAM1 | sp\|P13591\|NCAM1_HUMAN Neural cell adhesion molecule 1 OS=Homo sapiens GN=NC | 605,143 | 28,79 | 94574 | 18 | 26,99 | 27,10 | 24,47 | 26,74 | 26,69 | 24,14 | **0,796** | 7,99E-01 | **0** | -0,3299999 | 0,09730852 |
| 41_HUMAN | EPB41 | sp\|P11171\|41_HUMAN Protein 4.1 OS=Homo sapiens GN=EPB41 PE=1 SV=4 | 87,974 | 6,48 | 97017 | 5 | 24,62 | 23,58 | 23,06 | 24,32 | 23,65 | 22,30 | **0,796** | 6,83E-01 | **0** | -0,3300006 | 0,16567724 |
| HDAC6_HUN | HDAC6 | sp\|Q9UBN7\|HDAC6_HUMAN Histone deacetylase 6 OS=Homo sapiens GN=HDAC6 PE=1 | 42,987 | 1,98 | 131419 | 1 | 19,45 | 18,95 | 18,80 | 20,19 | 16,82 | 19,20 | **0,795** | 7,62E-01 | **0** | -0,3304119 | 0,11810966 |
| MAAI_HUM/ | GSTZ1 | sp\|O43708\|MAAI_HUMAN Maleylacetoacetate isomerase OS=Homo sapiens GN=GSTZ1 | 12,378 | 9,26 | 24212 | 1 | 19,43 | 19,72 | 17,69 | 19,53 | 19,46 | 16,86 | **0,795** | 7,75E-01 | **0** | -0,330609 | 0,11058311 |
| UBTD2_HUN | UBTD2 | sp\|Q8WUN7\|UBTD2_HUMAN Ubiquitin domain-containing protein 2 OS=Homo sapiens | 43,256 | 16,24 | 26190 | 3 | 21,57 | 21,92 | 19,52 | 21,41 | 21,12 | 19,48 | **0,794** | 7,46E-01 | **0** | -0,3328527 | 0,12733114 |
| RIPL1_HUM/ | RILPL1 | sp\|Q5EBL4\|RIPL1_HUMAN RILP-like protein 1 OS=Homo sapiens GN=RILPL1 PE=1 SV=1 | 36,602 | 2,73 | 47108 | 1 | 19,83 | 19,88 | 19,97 | 19,61 | 19,28 | 19,79 | **0,794** | 9,77E-02 | **0** | -0,3333321 | 1,01014448 |
| PI51A_HUM/ | PIP5K1A | sp\|Q99755\|PI51A_HUMAN Phosphatidylinositol 4-phosphate 5-kinase type-1 alpha OS= | 317,059 | 20,64 | 62633 | 8 | 23,72 | 23,52 | 22,20 | 23,29 | 23,45 | 21,70 | **0,794** | 6,73E-01 | **0** | -0,3333327 | 0,17168773 |
| PCDG5_HUN | PCDHGA5 | sp\|Q9Y5G8\|PCDG5_HUMAN Protocadherin gamma-A5 OS=Homo sapiens GN=PCDHGA5 | 137,703 | 7,09 | 100935 | 4 | 22,44 | 22,49 | 21,81 | 22,32 | 22,06 | 21,36 | **0,794** | 4,08E-01 | **0** | -0,3333333 | 0,38972223 |
| APOC3_HUN | APOC3 | sp\|P02656\|APOC3_HUMAN Apolipoprotein C-III OS=Homo sapiens GN=APOC3 PE=1 SV= | 93,085 | 16,16 | 10852 | 1 | 21,89 | 21,42 | 22,70 | 21,88 | 20,94 | 22,19 | **0,794** | 5,64E-01 | **0** | -0,3333333 | 0,24904798 |
| RAB3C_HUM | RAB3C | sp\|Q96E17\|RAB3C_HUMAN Ras-related protein Rab-3C OS=Homo sapiens GN=RAB3C PI | 124,342 | 14,1 | 25952 | 4 | 23,35 | 23,71 | 23,23 | 22,96 | 23,28 | 23,05 | **0,794** | 1,26E-01 | **0** | -0,3333333 | 0,89948306 |
| RAB3D_HUM | RAB3D | sp\|O95716\|RAB3D_HUMAN Ras-related protein Rab-3D OS=Homo sapiens GN=RAB3D F | 159,576 | 23,29 | 24267 | 5 | 23,35 | 23,71 | 23,23 | 22,96 | 23,28 | 23,05 | **0,794** | 1,26E-01 | **0** | -0,3333333 | 0,89948306 |
| COPT1_HUM | SLC31A1 | sp\|O15431\|COPT1_HUMAN High affinity copper uptake protein 1 OS=Homo sapiens GN | 77,612 | 15,79 | 21091 | 2 | 24,23 | 23,95 | 22,54 | 23,83 | 23,76 | 22,13 | **0,794** | 6,85E-01 | **0** | -0,333334 | 0,16450727 |
| BGAL_HUM/ | GLB1 | sp\|P16278\|BGAL_HUMAN Beta-galactosidase OS=Homo sapiens GN=GLB1 PE=1 SV=2 | 38,047 | 2,95 | 76075 | 1 | 19,76 | 19,60 | 20,87 | 19,20 | 19,98 | 20,05 | **0,794** | 5,28E-01 | **0** | -0,333526 | 0,27703522 |
| TRAD1_HUM | TRAFD1 | sp\|O14545\|TRAD1_HUMAN TRAF-type zinc finger domain-containing protein 1 OS=Hom | 30,059 | 1,89 | 64841 | 1 | 16,92 | 18,79 | 20,88 | 18,69 | 16,82 | 20,08 | **0,793** | 8,33E-01 | **0** | -0,3338534 | 0,07938073 |
| PMGT2_HUN | POMGNT2 | sp\|Q8NAT1\|PMGT2_HUMAN Protein O-linked-mannose beta-1,4-N-acetylglucosaminyl | 41,291 | 3,28 | 66615 | 1 | 18,92 | 16,77 | 19,57 | 18,21 | 17,60 | 18,43 | **0,793** | 7,24E-01 | **0** | -0,3340454 | 0,14023008 |
| CROCC_HUN | CROCC | sp\|Q5TZA2\|CROCC_HUMAN Rootletin OS=Homo sapiens GN=CROCC PE=1 SV=1 | 15,331 | 0,4 | 228523 | 1 | 24,74 | 24,52 | 20,46 | 24,61 | 24,13 | 19,98 | **0,793** | 8,77E-01 | **0** | -0,3342241 | 0,0570544 |
| S52A1_HUM | SLC52A1 | sp\|Q9NWF4\|S52A1_HUMAN Solute carrier family 52, riboflavin transporter, member 1 | 72,556 | 5,36 | 46317 | 1 | 17,77 | 19,90 | 18,01 | 18,62 | 17,99 | 18,06 | **0,793** | 6,59E-01 | **0** | -0,3352057 | 0,18140148 |
| IQGA2_HUM | IQGAP2 | sp\|Q13576\|IQGA2_HUMAN Ras GTPase-activating-like protein IQGAP2 OS=Homo sapier | 52,17 | 1,08 | 180578 | 2 | 24,39 | 24,55 | 19,12 | 24,41 | 24,07 | 18,58 | **0,792** | 9,03E-01 | **0** | -0,3361085 | 0,04415913 |
| RHOG_HUM. | RHOG | sp\|P84095\|RHOG_HUMAN Rho-related GTP-binding protein RhoG OS=Homo sapiens GI | 542,859 | 62,3 | 21309 | 11 | 27,48 | 27,47 | 27,14 | 27,25 | 27,12 | 26,71 | **0,792** | 1,63E-01 | **0** | -0,3366661 | 0,78714033 |
| COF2_HUMA | CFL2 | sp\|Q9Y281\|COF2_HUMAN Cofilin-2 OS=Homo sapiens GN=CFL2 PE=1 SV=1 | 747,442 | 85,54 | 18737 | 15 | 28,59 | 28,34 | 27,15 | 28,22 | 28,33 | 26,52 | **0,792** | 6,71E-01 | **0** | -0,3366667 | 0,17341065 |
| VAMP3_HUN | VAMP3 | sp\|Q15836\|VAMP3_HUMAN Vesicle-associated membrane protein 3 OS=Homo sapiens | 339,597 | 50 | 11309 | 7 | 25,33 | 25,37 | 25,74 | 25,26 | 25,26 | 24,91 | **0,792** | 1,27E-01 | **0** | -0,3366667 | 0,89686855 |
| BI2L1_HUM/ | BAIAP2L1 | sp\|Q9UHR4\|BI2L1_HUMAN Brain-specific angiogenesis inhibitor 1-associated protein 2- | 30,782 | 3,52 | 56883 | 2 | 23,47 | 22,36 | 19,69 | 22,53 | 22,54 | 19,43 | **0,790** | 8,35E-01 | **0** | -0,3399798 | 0,0785031 |
| APOB_HUM/ | APOB | sp\|P04114\|APOB_HUMAN Apolipoprotein B-100 OS=Homo sapiens GN=APOB PE=1 SV= | 186,936 | 1,88 | 515605 | 10 | 27,18 | 26,50 | 27,65 | 27,03 | 26,60 | 26,68 | **0,790** | 3,97E-01 | **0** | -0,3399995 | 0,40098539 |
| GIPC1_HUM. | GIPC1 | sp\|O14908\|GIPC1_HUMAN PDZ domain-containing protein GIPC1 OS=Homo sapiens GN | 230,485 | 19,52 | 36049 | 6 | 24,31 | 24,49 | 23,77 | 24,23 | 24,20 | 23,12 | **0,790** | 4,68E-01 | **0** | -0,3399995 | 0,32982401 |
| MYO10_HUN | MYO10 | sp\|Q9HD67\|MYO10_HUMAN Unconventional myosin-X OS=Homo sapiens GN=MYO10 | 173,695 | 4,37 | 237347 | 7 | 24,36 | 23,77 | 23,61 | 24,03 | 23,64 | 23,05 | **0,790** | 4,04E-01 | **0** | -0,3400008 | 0,39339269 |
| CA021_HUM | C1orf21 | sp\|Q9H246\|CA021_HUMAN Uncharacterized protein C1orf21 OS=Homo sapiens GN=C1 | 15,491 | 8,26 | 13865 | 1 | 22,07 | 21,35 | 19,41 | 21,58 | 20,69 | 19,54 | **0,790** | 7,48E-01 | **0** | -0,3406143 | 0,12612136 |
| MPP5_HUM/ | MPP5 | sp\|Q8N3R9\|MPP5_HUMAN MAGUK p55 subfamily member 5 OS=Homo sapiens GN=MI | 180,167 | 12,59 | 77294 | 7 | 24,63 | 24,39 | 22,42 | 24,27 | 23,97 | 22,17 | **0,788** | 7,38E-01 | **0** | -0,3433329 | 0,13165507 |
| TT30B_HUM. | TTC30B | sp\|Q8N4P2\|TT30B_HUMAN Tetratricopeptide repeat protein 30B OS=Homo sapiens GN | 160,607 | 15,94 | 76099 | 7 | 26,68 | 26,57 | 22,67 | 26,17 | 26,79 | 21,93 | **0,788** | 8,73E-01 | **0** | -0,3433329 | 0,05891271 |
| GNAI1_HUM | GNAI1 | sp\|P63096\|GNAI1_HUMAN Guanine nucleotide-binding protein G(i) subunit alpha-1 OS | 993,602 | 66,67 | 40361 | 26 | 24,65 | 24,49 | 23,83 | 24,35 | 24,19 | 23,40 | **0,788** | 4,24E-01 | **0** | -0,3433329 | 0,37230618 |
| GIT2_HUMA | GIT2 | sp\|Q14161\|GIT2_HUMAN ARF GTPase-activating protein GIT2 OS=Homo sapiens GN=GI | 307,853 | 17,92 | 84543 | 10 | 23,51 | 23,92 | 21,22 | 23,46 | 23,45 | 20,71 | **0,788** | 7,96E-01 | **0** | -0,3433336 | 0,09912454 |
| P4HA1_HUM | P4HA1 | sp\|P13674\|P4HA1_HUMAN Prolyl 4-hydroxylase subunit alpha-1 OS=Homo sapiens GN= | 409,844 | 33,52 | 61049 | 15 | 25,36 | 25,17 | 25,59 | 25,66 | 25,10 | 24,33 | **0,788** | 4,43E-01 | **0** | -0,3433336 | 0,35310691 |
| CXG1_HUMA | GJC1 | sp\|P36383\|CXG1_HUMAN Gap junction gamma-1 protein OS=Homo sapiens GN=GJC1 P | 14,133 | 2,27 | 45470 | 1 | 21,51 | 20,37 | 21,60 | 20,72 | 20,64 | 21,09 | **0,788** | 4,59E-01 | **0** | -0,3433342 | 0,33820636 |
| MYO1B_HUN | MYO1B | sp\|O43795\|MYO1B_HUMAN Unconventional myosin-Ib OS=Homo sapiens GN=MYO1B | 1357,964 | 39,61 | 131985 | 37 | 28,79 | 28,53 | 26,77 | 28,25 | 28,26 | 26,55 | **0,788** | 7,08E-01 | **0** | -0,3433342 | 0,1502683 |
| ZWILC_HUM | ZWILCH | sp\|Q9H900\|ZWILC_HUMAN Protein zwilch homolog OS=Homo sapiens GN=ZWILCH PE= | 37,291 | 1,69 | 67214 | 1 | 20,16 | 19,08 | 21,25 | 20,07 | 18,23 | 21,15 | **0,787** | 7,61E-01 | **0** | -0,3448912 | 0,11867478 |
| VP37D_HUM | VPS37D | sp\|Q86XT2\|VP37D_HUMAN Vacuolar protein sorting-associated protein 37D OS=Homo | 14,728 | 4,78 | 27730 | 1 | 20,29 | 19,72 | 18,61 | 20,28 | 19,00 | 18,31 | **0,787** | 6,73E-01 | **0** | -0,3452435 | 0,17212235 |
| GET4_HUMA | GET4 | sp\|Q7L5D6\|GET4_HUMAN Golgi to ER traffic protein 4 homolog OS=Homo sapiens GN= | 21,209 | 3,36 | 36504 | 1 | 18,16 | 20,01 | 20,37 | 18,58 | 20,41 | 18,51 | **0,787** | 7,27E-01 | **0** | -0,3459644 | 0,13855509 |
| STX12_HUM. | STX12 | sp\|Q86Y82\|STX12_HUMAN Syntaxin-12 OS=Homo sapiens GN=STX12 PE=1 SV=1 | 131,408 | 14,49 | 31642 | 3 | 24,25 | 24,40 | 23,17 | 24,17 | 24,04 | 22,57 | **0,786** | 6,18E-01 | **0** | -0,3466663 | 0,20877114 |
| CBPN_HUMA | CPN1 | sp\|P15169\|CBPN_HUMAN Carboxypeptidase N catalytic chain OS=Homo sapiens GN=CP | 42,906 | 5,46 | 52286 | 2 | 21,96 | 20,93 | 23,25 | 21,64 | 20,76 | 22,70 | **0,786** | 7,12E-01 | **0** | -0,3466663 | 0,14747721 |
| PALLD_HUM. | PALLD | sp\|Q8WX93\|PALLD_HUMAN Palladin OS=Homo sapiens GN=PALLD PE=1 SV=3 | 281,948 | 7,74 | 150564 | 10 | 24,28 | 24,11 | 25,25 | 23,69 | 23,96 | 24,95 | **0,786** | 5,43E-01 | **0** | -0,346667 | 0,26508061 |
| IFIT5_HUMA | IFIT5 | sp\|Q13325\|IFIT5_HUMAN Interferon-induced protein with tetratricopeptide repeats 5 ( | 96,307 | 6,02 | 55847 | 3 | 24,32 | 24,29 | 24,51 | 24,10 | 23,97 | 24,01 | **0,786** | 1,17E-02 | **0** | -0,346667 | 1,93038355 |
| BMR1A_HUN | BMPR1A | sp\|P36894\|BMR1A_HUMAN Bone morphogenetic protein receptor type-1A OS=Homo s | 65,084 | 2,82 | 60198 | 1 | 21,26 | 21,03 | 18,36 | 20,75 | 20,48 | 18,38 | **0,786** | 7,86E-01 | **0** | -0,3472131 | 0,10452829 |
| CALD1_HUM | CALD1 | sp\|Q05682\|CALD1_HUMAN Caldesmon OS=Homo sapiens GN=CALD1 PE=1 SV=3 | 219,927 | 11,6 | 93231 | 10 | 25,65 | 25,52 | 24,34 | 25,28 | 25,01 | 24,17 | **0,785** | 5,48E-01 | **0** | -0,3499997 | 0,26114713 |
| OGT1_HUM/ | OGT | sp\|O15294\|OGT1_HUMAN UDP-N-acetylglucosamine--peptide N-acetylglucosaminyltra | 77,426 | 4,21 | 116925 | 3 | 18,83 | 19,10 | 20,60 | 18,31 | 17,64 | 21,53 | **0,783** | 8,03E-01 | **0** | -0,3526611 | 0,09543831 |
| AKT1_HUMA | AKT1 | sp\|P31749\|AKT1_HUMAN RAC-alpha serine/threonine-protein kinase OS=Homo sapien | 39,5 | 3,54 | 55686 | 1 | 20,78 | 20,61 | 21,33 | 20,13 | 20,56 | 20,97 | **0,783** | 3,39E-01 | **0** | -0,3533344 | 0,46994988 |
| CAN5_HUMA | CAPN5 | sp\|O15484\|CAN5_HUMAN Calpain-5 OS=Homo sapiens GN=CAPN5 PE=1 SV=2 | 523,33 | 22,34 | 73169 | 14 | 27,84 | 27,17 | 26,72 | 27,34 | 26,84 | 26,48 | **0,781** | 4,33E-01 | **0** | -0,3566666 | 0,36313253 |
| NXF1_HUMA | NXF1 | sp\|Q9UBU9\|NXF1_HUMAN Nuclear RNA export factor 1 OS=Homo sapiens GN=NXF1 PE | 138,868 | 11,47 | 70182 | 6 | 23,82 | 23,54 | 21,32 | 23,17 | 22,77 | 21,67 | **0,781** | 7,15E-01 | **0** | -0,3566666 | 0,14579917 |

| | | | | | | | | | | | | | | | | | |
|---|---|---|---|---|---|---|---|---|---|---|---|---|---|---|---|---|---|
| ITB3_HUMA | ITGB3 | sp\|P05106\|ITB3_HUMAN Integrin beta-3 OS=Homo sapiens GN=ITGB3 PE=1 SV=2 | 1239,885 | 38,96 | 87058 | 27 | 29,55 | 28,54 | 27,96 | 29,48 | 28,49 | 27,01 | **0,781** | 6,98E-01 | **0** | -0,3566666 | 0,15617208 |
| CDC16_HUM | CDC16 | sp\|Q13042\|CDC16_HUMAN Cell division cycle protein 16 homolog OS=Homo sapiens G | 22,943 | 2,58 | 71656 | 2 | 22,59 | 21,35 | 22,98 | 20,98 | 21,90 | 22,97 | **0,781** | 6,62E-01 | **0** | -0,3566672 | 0,17925903 |
| TYRP1_HUM | TYRP1 | sp\|P17643\|TYRP1_HUMAN 5,6-dihydroxyindole-2-carboxylic acid oxidase OS=Homo sap | 15,269 | 1,3 | 60724 | 1 | 26,92 | 25,77 | 25,62 | 26,94 | 25,66 | 24,64 | **0,781** | 6,72E-01 | **0** | -0,3566672 | 0,17267194 |
| TRIP6_HUM | TRIP6 | sp\|Q15654\|TRIP6_HUMAN Thyroid receptor-interacting protein 6 OS=Homo sapiens G | 17,112 | 3,99 | 50288 | 1 | 20,17 | 20,09 | 18,53 | 18,60 | 19,59 | 19,52 | **0,779** | 5,95E-01 | **0** | -0,3595568 | 0,22565573 |
| THRB_HUMA | F2 | sp\|P00734\|THRB_HUMAN Prothrombin OS=Homo sapiens GN=F2 PE=1 SV=2 | 586,384 | 10,93 | 70037 | 11 | 27,10 | 26,22 | 27,39 | 26,75 | 26,19 | 26,69 | **0,779** | 4,13E-01 | **0** | -0,3599993 | 0,38447898 |
| SEPR_HUMA | FAP | sp\|Q12884\|SEPR_HUMAN Prolyl endopeptidase FAP OS=Homo sapiens GN=FAP PE=1 S | 650,842 | 27,63 | 87713 | 19 | 26,15 | 26,23 | 26,82 | 25,57 | 26,28 | 26,27 | **0,779** | 3,18E-01 | **0** | -0,3599993 | 0,49727318 |
| SNP25_HUM | SNAP25 | sp\|P60880\|SNP25_HUMAN Synaptosomal-associated protein 25 OS=Homo sapiens GN= | 42,434 | 4,85 | 23315 | 1 | 25,45 | 25,71 | 23,50 | 25,15 | 25,18 | 23,25 | **0,779** | 7,23E-01 | **0** | -0,36 | 0,14102172 |
| UBTD1_HUM | UBTD1 | sp\|Q9HAC8\|UBTD1_HUMAN Ubiquitin domain-containing protein 1 OS=Homo sapiens | 71,278 | 18,06 | 25938 | 3 | 22,23 | 22,31 | 21,01 | 22,31 | 21,69 | 20,46 | **0,777** | 6,25E-01 | **0** | -0,3633334 | 0,2040762 |
| EFR3A_HUM | EFR3A | sp\|Q14156\|EFR3A_HUMAN Protein EFR3 homolog A OS=Homo sapiens GN=EFR3A PE=1 | 564,399 | 22,41 | 92924 | 14 | 25,36 | 25,48 | 23,37 | 24,96 | 25,29 | 22,87 | **0,777** | 7,40E-01 | **0** | -0,3633334 | 0,13083063 |
| ANGE1_HUM | ANGEL1 | sp\|Q9UNK9\|ANGE1_HUMAN Protein angel homolog 1 OS=Homo sapiens GN=ANGEL1 P | 10,975 | 1,79 | 75276 | 1 | 19,79 | 19,50 | 19,36 | 19,61 | 18,91 | 19,04 | **0,777** | 2,19E-01 | **0** | -0,3633334 | 0,65951199 |
| APOH_HUM | APOH | sp\|P02749\|APOH_HUMAN Beta-2-glycoprotein 1 OS=Homo sapiens GN=APOH PE=1 SV= | 65,044 | 3,48 | 38298 | 2 | 25,45 | 25,34 | 24,92 | 25,38 | 24,70 | 24,54 | **0,777** | 2,98E-01 | **0** | -0,3633334 | 0,52582525 |
| TGFI1_HUM | TGFB1I1 | sp\|O43294\|TGFI1_HUMAN Transforming growth factor beta-1-induced transcript 1 pro | 280,092 | 27,33 | 49814 | 9 | 24,13 | 24,45 | 25,09 | 24,32 | 23,85 | 24,41 | **0,777** | 3,34E-01 | **0** | -0,3633334 | 0,47570143 |
| PI4KA_HUM | PI4KA | sp\|P42356\|PI4KA_HUMAN Phosphatidylinositol 4-kinase alpha OS=Homo sapiens GN=P | 627,251 | 12,57 | 231319 | 18 | 25,31 | 25,29 | 23,41 | 24,88 | 25,16 | 22,88 | **0,777** | 7,23E-01 | **0** | -0,363334 | 0,14086774 |
| GBB4_HUMA | GNB4 | sp\|Q9HAV0\|GBB4_HUMAN Guanine nucleotide-binding protein subunit beta-4 OS=Hom | 929,969 | 67,35 | 37567 | 17 | 27,20 | 26,61 | 25,87 | 26,51 | 26,62 | 25,46 | **0,777** | 5,33E-01 | **0** | -0,363334 | 0,27303454 |
| CSKP_HUMA | CASK | sp\|O14936\|CSKP_HUMAN Peripheral plasma membrane protein CASK OS=Homo sapien | 905,786 | 38,23 | 105123 | 29 | 27,52 | 27,53 | 25,72 | 27,14 | 27,13 | 25,41 | **0,777** | 6,85E-01 | **0** | -0,363334 | 0,16433678 |
| PCCB_HUMA | PCCB | sp\|P05166\|PCCB_HUMAN Propionyl-CoA carboxylase beta chain, mitochondrial OS=Hor | 17,208 | 2,04 | 58216 | 1 | 18,06 | 18,32 | 18,47 | 19,01 | 18,26 | 16,50 | **0,777** | 6,54E-01 | **0** | -0,3638961 | 0,18458935 |
| SC5A3_HUM | SLC5A3 | sp\|P53794\|SC5A3_HUMAN Sodium/myo-inositol cotransporter OS=Homo sapiens GN= | 125,814 | 7,1 | 79694 | 5 | 24,64 | 24,13 | 22,55 | 23,53 | 24,20 | 22,49 | **0,776** | 6,71E-01 | **0** | -0,3666655 | 0,17309364 |
| CHM2B_HUM | CHMP2B | sp\|Q9UQN3\|CHM2B_HUMAN Charged multivesicular body protein 2b OS=Homo sapien | 240,887 | 18,78 | 23907 | 7 | 26,80 | 26,65 | 24,93 | 26,71 | 25,97 | 24,60 | **0,776** | 6,92E-01 | **0** | -0,3666668 | 0,15974504 |
| ACTH_HUMA | ACTG2 | sp\|P63267\|ACTH_HUMAN Actin, gamma-enteric smooth muscle OS=Homo sapiens GN= | 2823,464 | 84,57 | 41877 | 56 | 22,52 | 21,98 | 22,83 | 22,51 | 22,17 | 21,55 | **0,776** | 3,84E-01 | **0** | -0,3666668 | 0,41612743 |
| ITA2B_HUM | ITGA2B | sp\|P08514\|ITA2B_HUMAN Integrin alpha-IIb OS=Homo sapiens GN=ITGA2B PE=1 SV=3 | 102,969 | 2,41 | 113377 | 3 | 26,75 | 25,71 | 24,58 | 26,12 | 25,72 | 24,09 | **0,774** | 6,96E-01 | **0** | -0,3699996 | 0,15712467 |
| CAD13_HUM | CDH13 | sp\|P55290\|CAD13_HUMAN Cadherin-13 OS=Homo sapiens GN=CDH13 PE=1 SV=1 | 166,794 | 7,43 | 78287 | 4 | 24,74 | 25,07 | 23,51 | 24,71 | 24,53 | 22,97 | **0,774** | 6,38E-01 | **0** | -0,3700002 | 0,1950505 |
| CYFP1_HUM | CYFIP1 | sp\|Q7L576\|CYFP1_HUMAN Cytoplasmic FMR1-interacting protein 1 OS=Homo sapiens | 1641,779 | 44,05 | 145182 | 45 | 27,10 | 26,67 | 26,35 | 26,53 | 26,67 | 25,81 | **0,774** | 3,42E-01 | **0** | -0,3700002 | 0,46546733 |
| MK15_HUM | MAPK15 | sp\|Q8TD08\|MK15_HUMAN Mitogen-activated protein kinase 15 OS=Homo sapiens GN= | 15,881 | 1,47 | 59832 | 1 | 24,20 | 24,43 | 23,69 | 23,87 | 23,94 | 23,40 | **0,774** | 2,52E-01 | **0** | -0,3700002 | 0,5983949 |
| TOM20_HUM | TOMM20 | sp\|Q15388\|TOM20_HUMAN Mitochondrial import receptor subunit TOM20 homolog O | 35,784 | 13,79 | 16298 | 1 | 19,44 | 18,53 | 19,50 | 18,14 | 19,19 | 19,03 | **0,774** | 4,60E-01 | **0** | -0,3700002 | 0,33738062 |
| RO60_HUMA | TROVE2 | sp\|P10155\|RO60_HUMAN 60 kDa SS-A/Ro ribonucleoprotein OS=Homo sapiens GN=TR | 439,378 | 25,65 | 60671 | 12 | 26,45 | 26,29 | 26,61 | 26,25 | 25,86 | 26,13 | **0,774** | 6,65E-02 | **0** | -0,3700008 | 1,17732136 |
| TRI23_HUM | TRIM23 | sp\|P36406\|TRI23_HUMAN E3 ubiquitin-protein ligase TRIM23 OS=Homo sapiens GN=TR | 31,745 | 1,74 | 64067 | 1 | 25,42 | 25,27 | 26,10 | 25,05 | 25,30 | 25,33 | **0,774** | 2,43E-01 | **0** | -0,3700008 | 0,6144792 |
| RAC1_HUMA | RAC1 | sp\|P63000\|RAC1_HUMAN Ras-related C3 botulinum toxin substrate 1 OS=Homo sapiens | 705,164 | 88,02 | 21450 | 16 | 28,98 | 28,79 | 28,44 | 28,78 | 28,37 | 27,94 | **0,772** | 2,67E-01 | **0** | -0,373333 | 0,57391447 |
| ZC3HA_HUM | ZC3H10 | sp\|Q96K80\|ZC3HA_HUMAN Zinc finger CCCH domain-containing protein 10 OS=Homo s | 143,191 | 13,59 | 46052 | 5 | 24,67 | 24,42 | 24,21 | 24,54 | 23,84 | 23,80 | **0,772** | 2,46E-01 | **0** | -0,373333 | 0,60980642 |
| MAOM_HUM | ME2 | sp\|P23368\|MAOM_HUMAN NAD-dependent malic enzyme, mitochondrial OS=Homo sa | 173,222 | 8,39 | 65444 | 3 | 22,07 | 21,06 | 23,27 | 21,37 | 21,91 | 22,00 | **0,772** | 6,06E-01 | **0** | -0,373333 | 0,21735506 |
| CALR_HUMA | CALR | sp\|P27797\|CALR_HUMAN Calreticulin OS=Homo sapiens GN=CALR PE=1 SV=1 | 484,08 | 41,73 | 48142 | 12 | 28,95 | 28,66 | 29,00 | 29,27 | 28,15 | 28,07 | **0,772** | 4,05E-01 | **0** | -0,3733336 | 0,39234623 |
| MGRN1_HUM | MGRN1 | sp\|O60291\|MGRN1_HUMAN E3 ubiquitin-protein ligase MGRN1 OS=Homo sapiens GN= | 38,566 | 4,71 | 60753 | 2 | 22,72 | 22,52 | 19,54 | 22,17 | 22,13 | 19,36 | **0,772** | 8,01E-01 | **0** | -0,3733336 | 0,09634184 |
| SERPH_HUM | SERPINH1 | sp\|P50454\|SERPH_HUMAN Serpin H1 OS=Homo sapiens GN=SERPINH1 PE=1 SV=2 | 695,274 | 41,15 | 46441 | 17 | 27,36 | 27,40 | 29,26 | 27,30 | 27,16 | 28,44 | **0,772** | 6,43E-01 | **0** | -0,3733336 | 0,19163573 |
| NAGAB_HUM | NAGA | sp\|P17050\|NAGAB_HUMAN Alpha-N-acetylgalactosaminidase OS=Homo sapiens GN=N | 35,515 | 2,68 | 46565 | 1 | 22,46 | 22,16 | 21,63 | 22,07 | 21,80 | 21,25 | **0,770** | 3,33E-01 | **0** | -0,3766664 | 0,47783885 |
| ANT3_HUMA | SERPINC1 | sp\|P01008\|ANT3_HUMAN Antithrombin-III OS=Homo sapiens GN=SERPINC1 PE=1 SV=1 | 150,987 | 12,07 | 52602 | 6 | 26,32 | 25,40 | 26,96 | 26,13 | 25,39 | 26,03 | **0,770** | 5,00E-01 | **0** | -0,3766664 | 0,30095648 |
| IKKA_HUMA | CHUK | sp\|O15111\|IKKA_HUMAN Inhibitor of nuclear factor kappa-B kinase subunit alpha OS= | 11,9 | 1,48 | 84640 | 1 | 20,76 | 20,03 | 20,76 | 20,00 | 19,87 | 20,55 | **0,770** | 3,05E-01 | **0** | -0,376667 | 0,51581343 |
| RLGPB_HUM | RALGAPB | sp\|Q86X10\|RLGPB_HUMAN Ral GTPase-activating protein subunit beta OS=Homo sapie | 61,456 | 1,2 | 166799 | 1 | 18,11 | 18,03 | 18,95 | 17,35 | 17,59 | 19,02 | **0,770** | 5,63E-01 | **0** | -0,376667 | 0,24923184 |
| HPBP1_HUM | HSPBP1 | sp\|Q9NZL4\|HPBP1_HUMAN Hsp70-binding protein 1 OS=Homo sapiens GN=HSPBP1 PE= | 110,154 | 10,22 | 39474 | 3 | 21,95 | 22,86 | 22,77 | 21,51 | 22,72 | 22,22 | **0,770** | 4,54E-01 | **0** | -0,3766677 | 0,34264622 |
| CUL4B_HUM | CUL4B | sp\|Q13620\|CUL4B_HUMAN Cullin-4B OS=Homo sapiens GN=CUL4B PE=1 SV=4 | 144,797 | 7,01 | 103982 | 6 | 18,77 | 19,56 | 17,79 | 18,67 | 17,89 | 18,43 | **0,769** | 5,37E-01 | **0** | -0,3782177 | 0,26973801 |
| AASD1_HUM | AARSD1 | sp\|Q9BTE6\|AASD1_HUMAN Alanyl-tRNA editing protein Aarsd1 OS=Homo sapiens GN= | 15,657 | 2,67 | 45480 | 1 | 21,02 | 20,87 | 18,09 | 20,76 | 18,95 | 19,13 | **0,768** | 7,50E-01 | **0** | -0,3799845 | 0,12491194 |
| PACN2_HUM | PACSIN2 | sp\|Q9UNF0\|PACN2_HUMAN Protein kinase C and casein kinase substrate in neurons pr | 665,89 | 31,07 | 55739 | 19 | 29,35 | 29,26 | 27,46 | 29,08 | 28,94 | 26,91 | **0,768** | 7,05E-01 | **0** | -0,3799998 | 0,15202188 |
| C2D1A_HUM | CC2D1A | sp\|Q6P1N0\|C2D1A_HUMAN Coiled-coil and C2 domain-containing protein 1A OS=Homo | 331,064 | 13,35 | 104062 | 12 | 26,33 | 26,06 | 23,41 | 26,19 | 25,57 | 22,90 | **0,768** | 7,96E-01 | **0** | -0,3799998 | 0,09922276 |
| LIN7C_HUM | LIN7C | sp\|Q9NUP9\|LIN7C_HUMAN Protein lin-7 homolog C OS=Homo sapiens GN=LIN7C PE=1 | 411,293 | 57,36 | 21834 | 11 | 26,20 | 25,98 | 24,35 | 25,96 | 25,71 | 23,72 | **0,768** | 7,00E-01 | **0** | -0,3800011 | 0,15481624 |
| LRC40_HUM | LRRC40 | sp\|Q9H9A6\|LRC40_HUMAN Leucine-rich repeat-containing protein 40 OS=Homo sapien | 26,301 | 3,32 | 68250 | 2 | 21,93 | 21,97 | 21,59 | 20,97 | 22,02 | 21,35 | **0,767** | 3,10E-01 | **0** | -0,3833332 | 0,50908378 |
| PTPA_HUMA | PTPA | sp\|Q15257\|PTPA_HUMAN Serine/threonine-protein phosphatase 2A activator OS=Hom | 84,554 | 10,34 | 40668 | 3 | 23,51 | 23,57 | 21,92 | 23,07 | 23,07 | 21,71 | **0,767** | 6,16E-01 | **0** | -0,3833338 | 0,21067161 |
| CPTP_HUMA | CPTP | sp\|Q5TA50\|CPTP_HUMAN Ceramide-1-phosphate transfer protein OS=Homo sapiens G | 18,047 | 3,27 | 24365 | 1 | 21,57 | 20,88 | 19,91 | 20,86 | 21,02 | 19,33 | **0,766** | 6,22E-01 | **0** | -0,3849525 | 0,20608727 |
| SRGP1_HUM | SRGAP1 | sp\|Q7Z6B7\|SRGP1_HUMAN SLIT-ROBO Rho GTPase-activating protein 1 OS=Homo sapie | 185,277 | 6,36 | 124264 | 5 | 21,16 | 21,24 | 17,07 | 19,79 | 20,89 | 17,63 | **0,765** | 8,29E-01 | **0** | -0,3866666 | 0,08146179 |
| TGM2_HUM | TGM2 | sp\|P21980\|TGM2_HUMAN Protein-glutamine gamma-glutamyltransferase 2 OS=Homo s | 303,477 | 13,97 | 77329 | 9 | 27,94 | 28,13 | 26,42 | 27,65 | 27,96 | 25,72 | **0,765** | 6,85E-01 | **0** | -0,3866673 | 0,16440985 |
| UBB_HUMA | UBB | sp\|P0CG47\|UBB_HUMAN Polyubiquitin-B OS=Homo sapiens GN=UBB PE=1 SV=1 | 896,325 | 98,69 | 25762 | 18 | 25,27 | 24,76 | 22,91 | 24,55 | 24,32 | 22,91 | **0,765** | 6,84E-01 | **0** | -0,3866673 | 0,16524396 |
| S10A1_HUM | S100A1 | sp\|P23297\|S10A1_HUMAN Protein S100-A1 OS=Homo sapiens GN=S100A1 PE=1 SV=2 | 51,481 | 22,34 | 10546 | 1 | 20,22 | 19,38 | 19,40 | 20,19 | 18,95 | 18,69 | **0,763** | 5,10E-01 | **0** | -0,3899988 | 0,29281605 |
| PTN23_HUM | PTPN23 | sp\|Q9H3S7\|PTN23_HUMAN Tyrosine-protein phosphatase non-receptor type 23 OS=H | 932,934 | 19,19 | 178974 | 23 | 26,83 | 26,43 | 27,46 | 26,60 | 26,20 | 26,75 | **0,763** | 3,18E-01 | **0** | -0,3899994 | 0,49821918 |
| TS101_HUM | TSG101 | sp\|Q99816\|TS101_HUMAN Tumor susceptibility gene 101 protein OS=Homo sapiens G | 1291,649 | 59,49 | 43944 | 29 | 30,04 | 29,71 | 28,75 | 29,67 | 29,33 | 28,33 | **0,763** | 5,23E-01 | **0** | -0,39 | 0,28137516 |
| PSD10_HUM | PSMD10 | sp\|O75832\|PSD10_HUMAN 26S proteasome non-ATPase regulatory subunit 10 OS=Hom | 173,606 | 42,92 | 24428 | 6 | 22,60 | 23,31 | 23,19 | 21,90 | 23,13 | 22,90 | **0,763** | 4,22E-01 | **0** | -0,3900007 | 0,37440258 |
| PRDX5_HUM | PRDX5 | sp\|P30044\|PRDX5_HUMAN Peroxiredoxin-5, mitochondrial OS=Homo sapiens GN=PRD | 324,161 | 33,18 | 22086 | 8 | 25,88 | 26,36 | 26,51 | 25,81 | 26,38 | 25,39 | **0,763** | 3,20E-01 | **0** | -0,3900007 | 0,49435299 |
| WIPF2_HUM | WIPF2 | sp\|Q8TF74\|WIPF2_HUMAN WAS/WASL-interacting protein family member 2 OS=Homo | 67,809 | 8,41 | 46289 | 3 | 23,72 | 23,68 | 21,52 | 23,64 | 22,98 | 21,13 | **0,763** | 7,28E-01 | **0** | -0,3900007 | 0,1378749 |
| PRAG1_HUM | PRAG1 | sp\|Q86YV5\|PRAG1_HUMAN Tyrosine-protein kinase PRAG1 OS=Homo sapiens GN=PRA | 45,852 | 1,42 | 149624 | 2 | 22,34 | 22,36 | 20,50 | 21,99 | 21,97 | 20,07 | **0,763** | 6,82E-01 | **0** | -0,3910796 | 0,16613834 |
| CD2AP_HUM | CD2AP | sp\|Q9Y5K6\|CD2AP_HUMAN CD2-associated protein OS=Homo sapiens GN=CD2AP PE=1 | 585,927 | 27,7 | 71451 | 16 | 27,56 | 27,49 | 26,05 | 27,29 | 27,10 | 25,53 | **0,761** | 6,25E-01 | **0** | -0,3933322 | 0,20419004 |
| MKRN1_HUM | MKRN1 | sp\|Q9UHC7\|MKRN1_HUMAN E3 ubiquitin-protein ligase makorin-1 OS=Homo sapiens G | 334,782 | 20,12 | 53349 | 10 | 26,26 | 26,15 | 25,66 | 26,13 | 25,75 | 25,01 | **0,761** | 3,56E-01 | **0** | -0,3933334 | 0,44888671 |
| CLAP1_HUM | CLASP1 | sp\|Q7Z460\|CLAP1_HUMAN CLIP-associating protein 1 OS=Homo sapiens GN=CLASP1 PE | 59,075 | 0,98 | 169451 | 1 | 17,97 | 17,78 | 19,69 | 17,26 | 17,31 | 19,69 | **0,761** | 7,16E-01 | **0** | -0,3933334 | 0,14525556 |
| FBLI1_HUMA | FBLIM1 | sp\|Q8WUP2\|FBLI1_HUMAN Filamin-binding LIM protein 1 OS=Homo sapiens GN=FBLIM | 51,803 | 6,7 | 40670 | 2 | 21,56 | 21,63 | 19,40 | 20,69 | 21,77 | 18,95 | **0,761** | 7,38E-01 | **0** | -0,3948816 | 0,13190747 |
| TPC6B_HUM | TRAPPC6B | sp\|Q86SZ2\|TPC6B_HUMAN Trafficking protein particle complex subunit 6B OS=Homo sa | 46,607 | 12,03 | 17983 | 1 | 19,35 | 18,22 | 19,67 | 17,80 | 18,80 | 19,45 | **0,760** | 5,75E-01 | **0** | -0,3965556 | 0,24002316 |
| UB2D3_HUM | UBE2D3 | sp\|P61077\|UB2D3_HUMAN Ubiquitin-conjugating enzyme E2 D3 OS=Homo sapiens GN= | 182,679 | 42,86 | 16687 | 4 | 21,58 | 21,63 | 21,25 | 21,52 | 20,63 | 21,12 | **0,760** | 2,35E-01 | **0** | -0,3966662 | 0,62985089 |
| MPP6_HUM | MPP6 | sp\|Q9NZW5\|MPP6_HUMAN MAGUK p55 subfamily member 6 OS=Homo sapiens GN=M | 826,651 | 56,48 | 61117 | 28 | 27,57 | 27,78 | 25,77 | 27,25 | 27,23 | 25,45 | **0,760** | 6,73E-01 | **0** | -0,3966668 | 0,17180036 |
| PABP1_HUM | PABPC1 | sp\|P11940\|PABP1_HUMAN Polyadenylate-binding protein 1 OS=Homo sapiens GN=PAB | 2879,836 | 65,88 | 70671 | 59 | 31,97 | 31,88 | 31,62 | 31,81 | 31,43 | 31,03 | **0,758** | 1,83E-01 | **0** | -0,3999996 | 0,7383236 |
| ACAD9_HUM | ACAD9 | sp\|Q9H845\|ACAD9_HUMAN Acyl-CoA dehydrogenase family member 9, mitochondrial | 72,091 | 6,12 | 68760 | 2 | 19,95 | 19,88 | 22,79 | 19,66 | 20,06 | 21,69 | **0,756** | 7,42E-01 | **0** | -0,4031467 | 0,12951267 |
| TIA1_HUMA | TIA1 | sp\|P31483\|TIA1_HUMAN Nucleolysin TIA-1 isoform p40 OS=Homo sapiens GN=TIA1 PE= | 101,954 | 12,44 | 42963 | 4 | 21,18 | 21,05 | 17,85 | 20,62 | 20,72 | 17,53 | **0,756** | 8,03E-01 | **0** | -0,403333 | 0,09545696 |
| SMUF1_HUM | SMURF1 | sp\|Q9HCE7\|SMUF1_HUMAN E3 ubiquitin-protein ligase SMURF1 OS=Homo sapiens GN= | 75,515 | 6,74 | 86114 | 5 | 24,86 | 24,38 | 23,65 | 24,21 | 24,36 | 23,11 | **0,756** | 4,88E-01 | **0** | -0,403333 | 0,31186513 |
| PAIP1_HUM | PAIP1 | sp\|Q9H074\|PAIP1_HUMAN Polyadenylate-binding protein-interacting protein 1 OS=Hor | 194,588 | 12,53 | 53525 | 6 | 25,66 | 25,47 | 25,35 | 25,44 | 25,17 | 24,66 | **0,756** | 1,76E-01 | **0** | -0,403333 | 0,75390075 |
| PDC10_HUM | PDCD10 | sp\|Q9BUL8\|PDC10_HUMAN Programmed cell death protein 10 OS=Homo sapiens GN=P | 75,808 | 24,06 | 24702 | 3 | 24,44 | 24,26 | 23,14 | 23,76 | 24,06 | 22,81 | **0,756** | 5,07E-01 | **0** | -0,4033337 | 0,29483385 |
| LIN7A_HUM | LIN7A | sp\|O14910\|LIN7A_HUMAN Protein lin-7 homolog A OS=Homo sapiens GN=LIN7A PE=1 | 248,29 | 30,04 | 25997 | 7 | 25,62 | 25,52 | 23,82 | 25,39 | 25,01 | 23,35 | **0,756** | 6,62E-01 | **0** | -0,4033337 | 0,17900917 |
| EMC8_HUM | EMC8 | sp\|O43402\|EMC8_HUMAN ER membrane protein complex subunit 8 OS=Homo sapiens | 11,091 | 4,29 | 23773 | 1 | 22,31 | 20,25 | 21,96 | 19,43 | 21,89 | 21,99 | **0,755** | 7,20E-01 | **0** | -0,4047038 | 0,14254034 |
| SCPDL_HUM | SCCPDH | sp\|Q8NBX0\|SCPDL_HUMAN Saccharopine dehydrogenase-like oxidoreductase OS=Hom | 65,211 | 4,9 | 47151 | 2 | 22,07 | 22,48 | 22,58 | 21,68 | 22,40 | 21,83 | **0,754** | 2,05E-01 | **0** | -0,4066664 | 0,68752975 |
| ITIH4_HUMA | ITIH4 | sp\|Q14624\|ITIH4_HUMAN Inter-alpha-trypsin inhibitor heavy chain H4 OS=Homo sapie | 38,818 | 1,94 | 103357 | 2 | 25,49 | 23,40 | 26,03 | 25,41 | 22,86 | 25,43 | **0,754** | 7,46E-01 | **0** | -0,4066664 | 0,12732363 |
| S22A8_HUM | SLC22A8 | sp\|Q8TCC7\|S22A8_HUMAN Solute carrier family 22 member 8 OS=Homo sapiens GN=SL | 10,43 | 1,29 | 59856 | 1 | 24,08 | 23,92 | 23,84 | 23,64 | 23,20 | 23,78 | **0,754** | 9,71E-02 | **0** | -0,4066664 | 1,01276408 |
| PP1R7_HUM | PPP1R7 | sp\|Q15435\|PP1R7_HUMAN Protein phosphatase 1 regulatory subunit 7 OS=Homo sapie | 113,435 | 7,78 | 41564 | 2 | 21,13 | 21,11 | 22,67 | 19,54 | 20,89 | 23,26 | **0,754** | 7,52E-01 | **0** | -0,4066664 | 0,12350511 |
| ALDH2_HUM | ALDH2 | sp\|P05091\|ALDH2_HUMAN Aldehyde dehydrogenase, mitochondrial OS=Homo sapiens | 87,274 | 11,61 | 56381 | 4 | 24,00 | 22,48 | 23,92 | 22,82 | 23,31 | 23,05 | **0,754** | 4,73E-01 | **0** | -0,4066671 | 0,32521973 |
| MBNL1_HUM | MBNL1 | sp\|Q9NR56\|MBNL1_HUMAN Muscleblind-like protein 1 OS=Homo sapiens GN=MBNL1 | 97,907 | 13,92 | 41817 | 4 | 24,31 | 24,05 | 23,97 | 24,03 | 23,47 | 23,60 | **0,753** | 1,07E-01 | **0** | -0,4099992 | 0,97051267 |
| GAPD1_HUM | GAPVD1 | sp\|Q14C86\|GAPD1_HUMAN GTPase-activating protein and VPS9 domain-containing pro | 94,7 | 3,18 | 164980 | 4 | 22,93 | 23,07 | 22,54 | 22,16 | 22,70 | 22,45 | **0,753** | 1,39E-01 | **0** | -0,4099998 | 0,85664012 |
| AT11C_HUM | ATP11C | sp\|Q8NB49\|AT11C_HUMAN Phospholipid-transporting ATPase IG OS=Homo sapiens GN | 277,476 | 9,63 | 129477 | 10 | 25,04 | 25,22 | 22,63 | 24,83 | 24,84 | 21,99 | **0,753** | 7,62E-01 | **0** | -0,4099998 | 0,11814313 |
| LRC8C_HUM | LRRC8C | sp\|Q8TDW0\|LRC8C_HUMAN Volume-regulated anion channel subunit LRRC8C OS=Hom | 61,762 | 7,47 | 92450 | 6 | 23,76 | 23,90 | 21,07 | 23,17 | 23,68 | 20,65 | **0,753** | 7,71E-01 | **0** | -0,4099998 | 0,11321065 |
| MPZL1_HUM | MPZL1 | sp\|O95297\|MPZL1_HUMAN Myelin protein zero-like protein 1 OS=Homo sapiens GN=M | 144,741 | 16,36 | 29082 | 5 | 26,06 | 25,99 | 24,10 | 25,73 | 25,41 | 23,78 | **0,753** | 6,66E-01 | **0** | -0,4099998 | 0,17654456 |
| FHL1_HUMA | FHL1 | sp\|Q13642\|FHL1_HUMAN Four and a half LIM domains protein 1 OS=Homo sapiens GN= | 268,168 | 27,55 | 36263 | 6 | 25,78 | 25,67 | 26,64 | 25,18 | 25,38 | 26,30 | **0,753** | 4,25E-01 | **0** | -0,4100005 | 0,37212021 |
| WASF2_HUM | WASF2 | sp\|Q9Y6W5\|WASF2_HUMAN Wiskott-Aldrich syndrome protein family member 2 OS=H | 244,193 | 12,85 | 54284 | 6 | 25,11 | 24,85 | 24,68 | 24,95 | 24,50 | 23,96 | **0,753** | 2,59E-01 | **0** | -0,4100005 | 0,58587128 |
| ARHG7_HUM | ARHGEF7 | sp\|Q14155\|ARHG7_HUMAN Rho guanine nucleotide exchange factor 7 OS=Homo sapie | 116,486 | 6,23 | 90012 | 4 | 23,31 | 23,45 | 21,55 | 23,05 | 22,88 | 21,15 | **0,753** | 6,59E-01 | **0** | -0,4100005 | 0,18114146 |
| LMF2_HUM | LMF2 | sp\|Q9BU23\|LMF2_HUMAN Lipase maturation factor 2 OS=Homo sapiens GN=LMF2 PE= | 33,135 | 2,55 | 79698 | 1 | 21,53 | 21,08 | 21,57 | 21,82 | 21,99 | 19,14 | **0,752** | 6,84E-01 | **0** | -0,4103018 | 0,16498718 |
| UB2G1_HUM | UBE2G1 | sp\|P62253\|UB2G1_HUMAN Ubiquitin-conjugating enzyme E2 G1 OS=Homo sapiens GN= | 36,041 | 7,06 | 19509 | 1 | 21,33 | 21,55 | 19,87 | 20,97 | 21,25 | 19,29 | **0,752** | 6,36E-01 | **0** | -0,4120413 | 0,19623742 |
| BAIP2_HUM | BAIAP2 | sp\|Q9UQB8\|BAIP2_HUMAN Brain-specific angiogenesis inhibitor 1-associated protein 2 | 467,837 | 27,36 | 60868 | 14 | 27,46 | 26,92 | 25,05 | 27,11 | 26,88 | 24,20 | **0,751** | 7,45E-01 | **0** | -0,4133326 | 0,12788008 |
| IPO4_HUMA | IPO4 | sp\|Q8TEX9\|IPO4_HUMAN Importin-4 OS=Homo sapiens GN=IPO4 PE=1 SV=2 | 70,563 | 2,31 | 118715 | 2 | 21,75 | 22,39 | 21,87 | 20,61 | 22,11 | 22,05 | **0,751** | 4,78E-01 | **0** | -0,4133333 | 0,32091565 |
| CELF2_HUM | CELF2 | sp\|O95319\|CELF2_HUMAN CUGBP Elav-like family member 2 OS=Homo sapiens GN=CE | 175,959 | 15,75 | 54285 | 6 | 23,81 | 23,78 | 24,07 | 23,57 | 23,40 | 23,45 | **0,751** | 1,70E-02 | **0** | -0,4133333 | 1,76948805 |
| DAG1_HUM | DAG1 | sp\|Q14118\|DAG1_HUMAN Dystroglycan OS=Homo sapiens GN=DAG1 PE=1 SV=2 | 203,886 | 7,15 | 97441 | 7 | 26,08 | 25,89 | 24,63 | 25,46 | 25,55 | 24,35 | **0,751** | 5,27E-01 | **0** | -0,4133333 | 0,27857114 |
| DLG1_HUMA | DLG1 | sp\|Q12959\|DLG1_HUMAN Disks large homolog 1 OS=Homo sapiens GN=DLG1 PE=1 SV= | 585,09 | 22,23 | 100455 | 13 | 26,45 | 26,65 | 24,28 | 25,97 | 26,16 | 24,01 | **0,751** | 7,07E-01 | **0** | -0,4133339 | 0,15054419 |
| PLXB1_HUM | PLXNB1 | sp\|O43157\|PLXB1_HUMAN Plexin-B1 OS=Homo sapiens GN=PLXNB1 PE=1 SV=3 | 17,169 | 1,17 | 232298 | 2 | 20,97 | 20,72 | 18,47 | 20,47 | 20,43 | 18,02 | **0,751** | 7,34E-01 | **0** | -0,4136054 | 0,13429859 |
| UN45A_HUM | UNC45A | sp\|Q9H3U1\|UN45A_HUMAN Protein unc-45 homolog A OS=Homo sapiens GN=UNC45A | 192,914 | 10,38 | 103077 | 8 | 24,13 | 24,62 | 23,06 | 23,37 | 23,85 | 23,34 | **0,749** | 4,42E-01 | **0** | -0,416666 | 0,35411964 |
| MYL1_HUM | MYL1 | sp\|P05976\|MYL1_HUMAN Myosin light chain 1/3, skeletal muscle isoform OS=Homo sa | 169,09 | 8,25 | 21145 | 4 | 25,80 | 26,15 | 23,75 | 25,68 | 25,79 | 22,98 | **0,749** | 7,43E-01 | **0** | -0,416666 | 0,12906904 |
| STX2_HUMA | STX2 | sp\|P32856\|STX2_HUMAN Syntaxin-2 OS=Homo sapiens GN=STX2 PE=1 SV=3 | 58,669 | 9,38 | 33341 | 3 | 24,00 | 23,90 | 21,30 | 23,53 | 23,23 | 21,19 | **0,749** | 7,35E-01 | **0** | -0,416666 | 0,13350479 |
| ITB2_HUMA | ITGB2 | sp\|P05107\|ITB2_HUMAN Integrin beta-2 OS=Homo sapiens GN=ITGB2 PE=1 SV=2 | 14,623 | 1,17 | 84782 | 1 | 21,19 | 20,65 | 20,42 | 20,86 | 20,77 | 19,38 | **0,749** | 4,76E-01 | **0** | -0,4166667 | 0,32220109 |
| CCM2_HUM | CCM2 | sp\|Q9BSQ5\|CCM2_HUMAN Cerebral cavernous malformations 2 protein OS=Homo sapi | 53,306 | 7,66 | 48837 | 3 | 23,32 | 22,77 | 19,04 | 22,85 | 22,18 | 18,85 | **0,749** | 8,31E-01 | **0** | -0,4166667 | 0,08052041 |
| GPX4_HUMA | GPX4 | sp\|P36969\|GPX4_HUMAN Phospholipid hydroperoxide glutathione peroxidase, mitoch | 38,791 | 12,69 | 22175 | 2 | 22,68 | 22,74 | 21,42 | 22,13 | 22,37 | 21,09 | **0,749** | 5,14E-01 | **0** | -0,4166667 | 0,28897662 |
| ENOPH_HUM | ENOPH1 | sp\|Q9UHY7\|ENOPH_HUMAN Enolase-phosphatase E1 OS=Homo sapiens GN=ENOPH1 | 48,259 | 9,2 | 28933 | 2 | 21,14 | 18,04 | 17,85 | 17,33 | 20,23 | 18,22 | **0,749** | 7,76E-01 | **0** | -0,4166673 | 0,11016085 |
| DCTP1_HUM | DCTPP1 | sp\|Q9H773\|DCTP1_HUMAN dCTP pyrophosphatase 1 OS=Homo sapiens GN=DCTPP1 PE | 43,793 | 11,76 | 18681 | 1 | 18,69 | 17,36 | 19,10 | 18,15 | 16,08 | 19,66 | **0,748** | 7,37E-01 | **0** | -0,4192956 | 0,13252902 |
| STK26_HUM | STK26 | sp\|Q9P289\|STK26_HUMAN Serine/threonine-protein kinase 26 OS=Homo sapiens GN= | 407,931 | 35,58 | 46529 | 10 | 25,31 | 25,48 | 25,30 | 24,85 | 25,12 | 24,86 | **0,747** | 1,66E-02 | **0** | -0,4199988 | 1,77956634 |
| SRC_HUMAN | SRC | sp\|P12931\|SRC_HUMAN Proto-oncogene tyrosine-protein kinase Src OS=Homo sapiens | 506,559 | 41,42 | 59835 | 17 | 26,60 | 26,24 | 25,23 | 26,00 | 25,85 | 24,96 | **0,747** | 4,67E-01 | **0** | -0,4200001 | 0,33076137 |
| DHRS4_HUM | DHRS4 | sp\|Q9BTZ2\|DHRS4_HUMAN Dehydrogenase/reductase SDR family member 4 OS=Homo | 14,581 | 3,6 | 29537 | 1 | 21,52 | 21,13 | 22,09 | 20,50 | 20,90 | 22,08 | **0,747** | 4,88E-01 | **0** | -0,4200001 | 0,31182043 |
| SMAD2_HUM | SMAD2 | sp\|Q15796\|SMAD2_HUMAN Mothers against decapentaplegic homolog 2 OS=Homo sa | 65,563 | 11,99 | 52306 | 4 | 22,71 | 22,59 | 22,33 | 22,46 | 22,13 | 21,78 | **0,747** | 1,37E-01 | **0** | -0,4200001 | 0,86396287 |
| ITAV_HUMA | ITGAV | sp\|P06756\|ITAV_HUMAN Integrin alpha-V OS=Homo sapiens GN=ITGAV PE=1 SV=2 | 1229,178 | 32,44 | 116038 | 30 | 28,52 | 28,13 | 27,68 | 28,13 | 27,91 | 27,03 | **0,747** | 3,68E-01 | **0** | -0,4200001 | 0,43383319 |
| RHOF_HUM | RHOF | sp\|Q9HBH0\|RHOF_HUMAN Rho-related GTP-binding protein RhoF OS=Homo sapiens G | 153,491 | 27,96 | 23625 | 7 | 26,23 | 25,79 | 23,90 | 25,71 | 25,40 | 23,55 | **0,747** | 6,91E-01 | **0** | -0,4200007 | 0,16048191 |
| NAA50_HUM | NAA50 | sp\|Q9GZZ1\|NAA50_HUMAN N-alpha-acetyltransferase 50 OS=Homo sapiens GN=NAA5 | 12,875 | 8,88 | 19398 | 1 | 20,06 | 21,20 | 21,78 | 23,19 | 19,49 | 19,10 | **0,747** | 7,78E-01 | **0** | -0,4213225 | 0,10900543 |
| PLCA_HUMA | AGPAT1 | sp\|Q99943\|PLCA_HUMAN 1-acyl-sn-glycerol-3-phosphate acyltransferase alpha OS=Ho | 34,663 | 7,42 | 31717 | 1 | 18,68 | 19,17 | 20,76 | 18,70 | 19,10 | 19,54 | **0,746** | 5,64E-01 | **0** | -0,423048 | 0,24887939 |
| GRB2_HUMA | GRB2 | sp\|P62993\|GRB2_HUMAN Growth factor receptor-bound protein 2 OS=Homo sapiens G | 546,492 | 75,12 | 25206 | 16 | 27,94 | 27,82 | 26,63 | 27,51 | 27,28 | 26,33 | **0,746** | 4,86E-01 | **0** | -0,4233329 | 0,31313252 |
| TSN5_HUMA | TSPAN5 | sp\|P62079\|TSN5_HUMAN Tetraspanin-5 OS=Homo sapiens GN=TSPAN5 PE=1 SV=1 | 102,587 | 11,19 | 30337 | 2 | 22,48 | 22,14 | 19,93 | 22,13 | 21,51 | 19,64 | **0,746** | 7,19E-01 | **0** | -0,4233335 | 0,14342778 |
| TAOK1_HUM | TAOK1 | sp\|Q7L7X3\|TAOK1_HUMAN Serine/threonine-protein kinase TAO1 OS=Homo sapiens G | 500,19 | 24,98 | 116070 | 21 | 25,68 | 25,80 | 24,56 | 25,88 | 25,30 | 23,59 | **0,746** | 6,22E-01 | **0** | -0,4233335 | 0,20647713 |
| YKT6_HUMA | YKT6 | sp\|O15498\|YKT6_HUMAN Synaptobrevin homolog YKT6 OS=Homo sapiens GN=YKT6 PE | 121,965 | 23,74 | 22418 | 5 | 25,69 | 25,79 | 24,58 | 25,29 | 25,29 | 24,21 | **0,746** | 4,68E-01 | **0** | -0,4233335 | 0,32931492 |
| MPRIP_HUM | MPRIP | sp\|Q6WCQ1\|MPRIP_HUMAN Myosin phosphatase Rho-interacting protein OS=Homo sa | 45,135 | 1,95 | 116533 | 1 | 19,93 | 20,52 | 20,22 | 20,01 | 19,75 | 19,64 | **0,746** | 1,05E-01 | **0** | -0,4237073 | 0,97945285 |
| ODB2_HUM | DBT | sp\|P11182\|ODB2_HUMAN Lipoamide acyltransferase component of branched-chain alp | 20,735 | 1,87 | 53487 | 1 | 19,22 | 18,56 | 20,90 | 18,70 | 18,59 | 20,12 | **0,745** | 6,45E-01 | **0** | -0,424708 | 0,19051315 |
| KC1G1_HUM | CSNK1G1 | sp\|Q9HCP0\|KC1G1_HUMAN Casein kinase I isoform gamma-1 OS=Homo sapiens GN=CS | 197,273 | 22,27 | 48511 | 8 | 20,94 | 22,00 | 19,53 | 21,44 | 21,27 | 18,48 | **0,744** | 7,39E-01 | **0** | -0,4266173 | 0,13120249 |
| GRSF1_HUM | GRSF1 | sp\|Q12849\|GRSF1_HUMAN G-rich sequence factor 1 OS=Homo sapiens GN=GRSF1 PE= | 35,469 | 4,58 | 53126 | 1 | 17,64 | 17,57 | 20,60 | 17,86 | 17,24 | 19,43 | **0,744** | 7,39E-01 | **0** | -0,4266663 | 0,13161409 |
| GID8_HUMA | GID8 | sp\|Q9NWU2\|GID8_HUMAN Glucose-induced degradation protein 8 homolog OS=Homo | 53,959 | 20,18 | 26749 | 2 | 19,42 | 20,43 | 22,05 | 19,24 | 18,74 | 22,64 | **0,744** | 7,82E-01 | **0** | -0,4266669 | 0,10653204 |
| TOE1_HUMA | TOE1 | sp\|Q96GM8\|TOE1_HUMAN Target of EGR1 protein 1 OS=Homo sapiens GN=TOE1 PE=1 | 22,025 | 2,16 | 56548 | 1 | 17,60 | 19,47 | 20,40 | 18,50 | 17,54 | 20,15 | **0,744** | 7,23E-01 | **0** | -0,427379 | 0,14092763 |
| MENT_HUM | MENT | sp\|Q9BUN1\|MENT_HUMAN Protein MENT OS=Homo sapiens GN=MENT PE=2 SV=1 | 10,541 | 2,05 | 36769 | 1 | 20,75 | 20,45 | 19,08 | 20,62 | 19,98 | 18,40 | **0,743** | 6,35E-01 | **0** | -0,4282786 | 0,1971011 |
| RHOD_HUM | RHOD | sp\|O00212\|RHOD_HUMAN Rho-related GTP-binding protein RhoD OS=Homo sapiens G | 40,844 | 9,05 | 23413 | 2 | 22,72 | 22,50 | 20,28 | 22,11 | 21,93 | 20,17 | **0,742** | 6,88E-01 | **0** | -0,4299997 | 0,16245528 |
| ITA1_HUMA | ITGA1 | sp\|P56199\|ITA1_HUMAN Integrin alpha-1 OS=Homo sapiens GN=ITGA1 PE=1 SV=2 | 86,269 | 2,37 | 130848 | 3 | 22,06 | 22,60 | 22,77 | 21,77 | 22,09 | 22,28 | **0,742** | 1,74E-01 | **0** | -0,4299997 | 0,75848335 |
| TXTP_HUMA | SLC25A1 | sp\|P53007\|TXTP_HUMAN Tricarboxylate transport protein, mitochondrial OS=Homo sa | 49 | 15,11 | 34013 | 2 | 20,52 | 21,56 | 22,11 | 20,07 | 21,05 | 21,77 | **0,741** | 5,58E-01 | **0** | -0,4326477 | 0,25336825 |
| TRI26_HUM | TRIM26 | sp\|Q12899\|TRI26_HUMAN Tripartite motif-containing protein 26 OS=Homo sapiens GN= | 283,393 | 25,23 | 62166 | 12 | 26,47 | 26,62 | 25,66 | 26,08 | 26,34 | 25,03 | **0,741** | 4,34E-01 | **0** | -0,4333331 | 0,36213182 |
| SCAM3_HUM | SCAMP3 | sp\|O14828\|SCAM3_HUMAN Secretory carrier-associated membrane protein 3 OS=Hom | 168,384 | 15,27 | 38287 | 3 | 23,91 | 24,17 | 24,83 | 23,85 | 23,60 | 24,16 | **0,741** | 2,45E-01 | **0** | -0,4333331 | 0,61114091 |
| FRIH_HUMA | FTH1 | sp\|P02794\|FRIH_HUMAN Ferritin heavy chain OS=Homo sapiens GN=FTH1 PE=1 SV=2 | 139,049 | 37,7 | 21226 | 6 | 23,97 | 24,28 | 23,93 | 23,77 | 23,57 | 23,54 | **0,741** | 3,05E-02 | **0** | -0,4333331 | 1,51605571 |
| RASK_HUMA | KRAS | sp\|P01116\|RASK_HUMAN GTPase KRas OS=Homo sapiens GN=KRAS PE=1 SV=1 | 497,898 | 63,49 | 21656 | 11 | 25,74 | 25,58 | 24,30 | 25,39 | 24,99 | 23,94 | **0,741** | 5,28E-01 | **0** | -0,4333331 | 0,27717709 |
| SEM3C_HUM | SEMA3C | sp\|Q99985\|SEM3C_HUMAN Semaphorin-3C OS=Homo sapiens GN=SEMA3C PE=2 SV=2 | 45,859 | 2,93 | 85207 | 2 | 23,42 | 22,95 | 21,16 | 23,22 | 22,70 | 20,31 | **0,741** | 7,21E-01 | **0** | -0,4333337 | 0,14214441 |
| SNX3_HUMA | SNX3 | sp\|O60493\|SNX3_HUMAN Sorting nexin-3 OS=Homo sapiens GN=SNX3 PE=1 SV=3 | 75,987 | 27,16 | 18762 | 5 | 24,37 | 24,31 | 23,83 | 24,13 | 24,08 | 23,00 | **0,741** | 3,46E-01 | **0** | -0,4333337 | 0,46058011 |
| PHLB1_HUM | PHLDB1 | sp\|Q86UU1\|PHLB1_HUMAN Pleckstrin homology-like domain family B member 1 OS=H | 64,958 | 2,47 | 151162 | 3 | 22,63 | 22,36 | 22,83 | 22,32 | 21,88 | 22,32 | **0,741** | 9,63E-02 | **0** | -0,4333337 | 1,01623825 |
| A2AP_HUMA | SERPINF2 | sp\|P08697\|A2AP_HUMAN Alpha-2-antiplasmin OS=Homo sapiens GN=SERPINF2 PE=1 S | 43,862 | 6,11 | 54566 | 2 | 23,29 | 22,36 | 24,97 | 23,36 | 22,12 | 23,83 | **0,739** | 6,59E-01 | **0** | -0,4366665 | 0,18094724 |
| TCF25_HUM | TCF25 | sp\|Q9BQ70\|TCF25_HUMAN Transcription factor 25 OS=Homo sapiens GN=TCF25 PE=1 | 85,276 | 5,33 | 76667 | 4 | 22,82 | 22,66 | 23,24 | 22,66 | 21,97 | 22,78 | **0,739** | 2,27E-01 | **0** | -0,4366665 | 0,64456515 |
| SF3A2_HUM | SF3A2 | sp\|Q15428\|SF3A2_HUMAN Splicing factor 3A subunit 2 OS=Homo sapiens GN=SF3A2 PE | 95,484 | 9,91 | 49256 | 3 | 22,34 | 21,88 | 23,08 | 21,65 | 21,47 | 22,87 | **0,739** | 4,80E-01 | **0** | -0,4366665 | 0,31842611 |
| E41L2_HUM | EPB41L2 | sp\|O43491\|E41L2_HUMAN Band 4.1-like protein 2 OS=Homo sapiens GN=EPB41L2 PE=1 | 890,319 | 26,57 | 112588 | 26 | 27,62 | 27,58 | 26,69 | 27,25 | 27,21 | 26,12 | **0,739** | 4,13E-01 | **0** | -0,4366671 | 0,38374191 |
| RBM15_HUM | RBM15 | sp\|Q96T37\|RBM15_HUMAN Putative RNA-binding protein 15 OS=Homo sapiens GN=RB | 41,741 | 1,74 | 107189 | 1 | 20,19 | 16,58 | 20,66 | 18,29 | 17,10 | 20,73 | **0,738** | 8,07E-01 | **0** | -0,4378332 | 0,09335339 |
| TAGL_HUMA | TAGLN | sp\|Q01995\|TAGL_HUMAN Transgelin OS=Homo sapiens GN=TAGLN PE=1 SV=4 | 983,472 | 78,61 | 22611 | 25 | 29,55 | 29,62 | 29,67 | 29,12 | 29,19 | 29,21 | **0,737** | 5,73E-04 | **0** | -0,4399999 | 3,2418141 |
| TETN_HUMA | CLEC3B | sp\|P05452\|TETN_HUMAN Tetranectin OS=Homo sapiens GN=CLEC3B PE=1 SV=3 | 30,519 | 5,45 | 22537 | 1 | 21,51 | 18,94 | 22,59 | 21,16 | 18,12 | 22,44 | **0,737** | 8,06E-01 | **0** | -0,4409536 | 0,09393117 |
| IFT80_HUMA | IFT80 | sp\|Q9P2H3\|IFT80_HUMAN Intraflagellar transport protein 80 homolog OS=Homo sapie | 341,596 | 14,54 | 88035 | 10 | 19,80 | 20,57 | 25,56 | 19,65 | 20,77 | 24,18 | **0,735** | 8,54E-01 | **0** | -0,4433327 | 0,06846531 |
| GNS_HUMA | GNS | sp\|P15586\|GNS_HUMAN N-acetylglucosamine-6-sulfatase OS=Homo sapiens GN=GNS | 121,258 | 8,15 | 62082 | 4 | 23,24 | 22,46 | 21,80 | 23,09 | 22,58 | 20,50 | **0,735** | 6,46E-01 | **0** | -0,4433327 | 0,18958886 |
| PTK7_HUMA | PTK7 | sp\|Q13308\|PTK7_HUMAN Inactive tyrosine-protein kinase 7 OS=Homo sapiens GN=PTK | 429,235 | 15,33 | 118392 | 14 | 26,75 | 26,80 | 25,67 | 26,31 | 26,29 | 25,29 | **0,735** | 4,25E-01 | **0** | -0,4433327 | 0,3718988 |
| LTOR1_HUM | LAMTOR1 | sp\|Q6IAA8\|LTOR1_HUMAN Ragulator complex protein LAMTOR1 OS=Homo sapiens GN | 37,215 | 14,29 | 17745 | 2 | 23,21 | 23,07 | 21,60 | 22,73 | 22,74 | 21,08 | **0,735** | 5,88E-01 | **0** | -0,4433333 | 0,23029191 |
| CPNE2_HUM | CPNE2 | sp\|Q96FN4\|CPNE2_HUMAN Copine-2 OS=Homo sapiens GN=CPNE2 PE=1 SV=3 | 480,796 | 28,47 | 61190 | 12 | 25,74 | 25,76 | 25,94 | 25,35 | 25,93 | 24,83 | **0,735** | 2,43E-01 | **0** | -0,4433333 | 0,61432484 |
| S20A1_HUM | SLC20A1 | sp\|Q8WUM9\|S20A1_HUMAN Sodium-dependent phosphate transporter 1 OS=Homo sa | 82,865 | 6,77 | 73700 | 3 | 22,26 | 22,05 | 18,96 | 21,98 | 21,89 | 18,07 | **0,735** | 8,04E-01 | **0** | -0,4433333 | 0,09469457 |
| LRP1_HUMA | LRP1 | sp\|Q07954\|LRP1_HUMAN Prolow-density lipoprotein receptor-related protein 1 OS=Ho | 1244,753 | 12,59 | 504606 | 41 | 26,75 | 26,75 | 25,57 | 26,44 | 26,48 | 24,82 | **0,735** | 5,46E-01 | **0** | -0,4433333 | 0,26251265 |
| SR140_HUM | U2SURP | sp\|O15042\|SR140_HUMAN U2 snRNP-associated SURP motif-containing protein OS=Ho | 78,464 | 3,6 | 118292 | 3 | 23,15 | 23,15 | 23,51 | 22,75 | 22,00 | 23,73 | **0,735** | 4,38E-01 | **0** | -0,4433333 | 0,35861614 |
| RND3_HUM | RND3 | sp\|P61587\|RND3_HUMAN Rho-related GTP-binding protein RhoE OS=Homo sapiens GN | 72,903 | 9,84 | 27368 | 2 | 23,38 | 23,59 | 21,72 | 23,14 | 22,72 | 21,50 | **0,735** | 5,95E-01 | **0** | -0,4433333 | 0,225287 |
| U3IP2_HUM | RRP9 | sp\|O43818\|U3IP2_HUMAN U3 small nucleolar RNA-interacting protein 2 OS=Homo sapi | 23,544 | 2,53 | 51841 | 1 | 16,82 | 19,03 | 18,25 | 16,94 | 16,68 | 19,15 | **0,735** | 6,84E-01 | **0** | -0,4451084 | 0,16495756 |
| PTTG_HUMA | PTTG1IP | sp\|P53801\|PTTG_HUMAN Pituitary tumor-transforming gene 1 protein-interacting prot | 36,593 | 6,67 | 20324 | 1 | 23,48 | 23,07 | 22,08 | 23,29 | 22,45 | 21,55 | **0,734** | 5,31E-01 | **0** | -0,4466661 | 0,27496714 |
| RL26_HUMA | RPL26 | sp\|P61254\|RL26_HUMAN 60S ribosomal protein L26 OS=Homo sapiens GN=RPL26 PE=1 | 262,1 | 46,9 | 17258 | 10 | 23,83 | 24,54 | 24,96 | 23,70 | 22,96 | 25,33 | **0,734** | 5,95E-01 | **0** | -0,4466667 | 0,22568185 |
| RASN_HUMA | NRAS | sp\|P01111\|RASN_HUMAN GTPase NRas OS=Homo sapiens GN=NRAS PE=1 SV=1 | 476,677 | 59,79 | 21229 | 10 | 24,02 | 24,88 | 23,28 | 24,24 | 23,76 | 22,84 | **0,734** | 5,10E-01 | **0** | -0,4466667 | 0,29234703 |
| RB27B_HUM | RAB27B | sp\|O00194\|RB27B_HUMAN Ras-related protein Rab-27B OS=Homo sapiens GN=RAB27B | 127,243 | 19,72 | 24608 | 4 | 24,67 | 23,38 | 22,68 | 24,49 | 23,55 | 21,35 | **0,734** | 7,05E-01 | **0** | -0,4466667 | 0,15183622 |
| PABP4_HUM | PABPC4 | sp\|Q13310\|PABP4_HUMAN Polyadenylate-binding protein 4 OS=Homo sapiens GN=PAB | 2177,941 | 57,92 | 70783 | 49 | 29,77 | 29,62 | 29,09 | 29,42 | 29,15 | 28,57 | **0,734** | 2,41E-01 | **0** | -0,4466674 | 0,61811324 |
| BAX_HUMA | BAX | sp\|Q07812\|BAX_HUMAN Apoptosis regulator BAX OS=Homo sapiens GN=BAX PE=1 SV= | 49,467 | 6,77 | 21184 | 1 | 17,44 | 20,13 | 20,21 | 16,78 | 19,22 | 20,43 | **0,733** | 7,66E-01 | **0** | -0,4484247 | 0,11583673 |
| CYTM1_HUM | CYSTM1 | sp\|Q9H1C7\|CYTM1_HUMAN Cysteine-rich and transmembrane domain-containing prot | 44,945 | 11,34 | 10631 | 2 | 24,96 | 24,80 | 22,34 | 24,11 | 24,25 | 22,39 | **0,732** | 6,87E-01 | **0** | -0,4499995 | 0,16312021 |
| ARRC_HUMA | ARR3 | sp\|P36575\|ARRC_HUMAN Arrestin-C OS=Homo sapiens GN=ARR3 PE=1 SV=2 | 14,478 | 1,8 | 42778 | 1 | 21,72 | 21,80 | 25,79 | 21,56 | 21,31 | 25,09 | **0,732** | 8,16E-01 | **0** | -0,4500001 | 0,0880949 |
| ADCY6_HUM | ADCY6 | sp\|O43306\|ADCY6_HUMAN Adenylate cyclase type 6 OS=Homo sapiens GN=ADCY6 PE= | 31,514 | 1,63 | 130615 | 2 | 20,02 | 18,78 | 19,92 | 19,86 | 19,48 | 18,03 | **0,732** | 5,47E-01 | **0** | -0,4500001 | 0,26198878 |
| ARF6_HUMA | ARF6 | sp\|P62330\|ARF6_HUMAN ADP-ribosylation factor 6 OS=Homo sapiens GN=ARF6 PE=1 S | 602,824 | 70,86 | 20082 | 13 | 28,35 | 27,96 | 28,41 | 27,75 | 27,99 | 27,63 | **0,732** | 6,32E-02 | **0** | -0,4500001 | 1,19934742 |
| LUM_HUMA | LUM | sp\|P51884\|LUM_HUMAN Lumican OS=Homo sapiens GN=LUM PE=1 SV=2 | 94,326 | 13,91 | 38429 | 4 | 25,17 | 23,87 | 25,74 | 25,10 | 23,80 | 24,53 | **0,732** | 5,38E-01 | **0** | -0,4500001 | 0,26912467 |
| IMPA1_HUM | IMPA1 | sp\|P29218\|IMPA1_HUMAN Inositol monophosphatase 1 OS=Homo sapiens GN=IMPA1 | 29,485 | 3,97 | 30189 | 1 | 22,04 | 22,16 | 19,93 | 22,20 | 21,76 | 18,81 | **0,731** | 7,43E-01 | **0** | -0,4514192 | 0,12880477 |
| KC1E_HUMA | CSNK1E | sp\|P49674\|KC1E_HUMAN Casein kinase I isoform epsilon OS=Homo sapiens GN=CSNK1 | 109,903 | 13,46 | 47315 | 5 | 19,36 | 19,85 | 19,48 | 19,82 | 18,34 | 19,18 | **0,731** | 3,75E-01 | **0** | -0,4524307 | 0,42652538 |
| INT9_HUMA | INTS9 | sp\|Q9NV88\|INT9_HUMAN Integrator complex subunit 9 OS=Homo sapiens GN=INTS9 P | 56,089 | 1,82 | 73814 | 1 | 19,91 | 18,51 | 19,85 | 17,58 | 18,90 | 20,44 | **0,731** | 6,57E-01 | **0** | -0,4525655 | 0,18227971 |
| CAN7_HUMA | CAPN7 | sp\|Q9Y6W3\|CAN7_HUMAN Calpain-7 OS=Homo sapiens GN=CAPN7 PE=1 SV=1 | 35,53 | 4,92 | 92652 | 3 | 22,32 | 22,13 | 21,07 | 22,42 | 21,49 | 20,25 | **0,730** | 5,73E-01 | **0** | -0,4533329 | 0,24198497 |
| EPS15_HUM | EPS15 | sp\|P42566\|EPS15_HUMAN Epidermal growth factor receptor substrate 15 OS=Homo sa | 130,325 | 6,25 | 98656 | 5 | 23,93 | 23,59 | 24,06 | 23,54 | 23,58 | 23,10 | **0,730** | 9,48E-02 | **0** | -0,4533329 | 1,02302813 |
| VP37A_HUM | VPS37A | sp\|Q8NEZ2\|VP37A_HUMAN Vacuolar protein sorting-associated protein 37A OS=Homo | 67,56 | 4,03 | 44314 | 1 | 22,38 | 22,41 | 21,29 | 22,31 | 21,99 | 20,42 | **0,730** | 5,47E-01 | **0** | -0,4533335 | 0,26173737 |
| KIF7_HUMA | KIF7 | sp\|Q2M1P5\|KIF7_HUMAN Kinesin-like protein KIF7 OS=Homo sapiens GN=KIF7 PE=1 SV | 119,372 | 3,72 | 150587 | 3 | 17,84 | 20,04 | 21,98 | 19,16 | 18,29 | 21,05 | **0,730** | 7,69E-01 | **0** | -0,454944 | 0,11405304 |
| VP37C_HUM | VPS37C | sp\|A5D8V6\|VP37C_HUMAN Vacuolar protein sorting-associated protein 37C OS=Homo | 228,83 | 16,06 | 38659 | 6 | 23,67 | 23,61 | 25,67 | 23,57 | 22,95 | 25,06 | **0,729** | 6,46E-01 | **0** | -0,4566669 | 0,18950739 |
| APAF_HUMA | APAF1 | sp\|O14727\|APAF_HUMAN Apoptotic protease-activating factor 1 OS=Homo sapiens GN | 65,872 | 1,12 | 141840 | 1 | 18,84 | 19,41 | 19,94 | 18,42 | 18,35 | 20,05 | **0,729** | 5,15E-01 | **0** | -0,4566669 | 0,28836812 |
| ITA2_HUMA | ITGA2 | sp\|P17301\|ITA2_HUMAN Integrin alpha-2 OS=Homo sapiens GN=ITGA2 PE=1 SV=1 | 32,516 | 0,93 | 129295 | 1 | 21,03 | 20,26 | 19,83 | 21,21 | 20,62 | 17,92 | **0,728** | 6,92E-01 | **0** | -0,4572964 | 0,16004482 |
| 1C02_HUMA | HLA-C | sp\|P30501\|1C02_HUMAN HLA class I histocompatibility antigen, Cw-2 alpha chain OS=H | 124,615 | 16,12 | 41095 | 4 | 21,79 | 20,29 | 20,70 | 21,96 | 21,06 | 18,38 | **0,727** | 7,14E-01 | **0** | -0,4591783 | 0,14645061 |
| IGSF8_HUM | IGSF8 | sp\|Q969P0\|IGSF8_HUMAN Immunoglobulin superfamily member 8 OS=Homo sapiens G | 689,805 | 26,26 | 65034 | 13 | 28,00 | 27,91 | 26,65 | 27,78 | 27,40 | 26,00 | **0,727** | 5,44E-01 | **0** | -0,4599997 | 0,26427083 |
| MCTS1_HUM | MCTS1 | sp\|Q9ULC4\|MCTS1_HUMAN Malignant T-cell-amplified sequence 1 OS=Homo sapiens G | 37,522 | 9,39 | 20555 | 1 | 18,62 | 19,35 | 19,08 | 17,94 | 19,16 | 18,57 | **0,727** | 3,26E-01 | **0** | -0,4600004 | 0,48621573 |
| ARFP1_HUM | ARFIP1 | sp\|P53367\|ARFP1_HUMAN Arfaptin-1 OS=Homo sapiens GN=ARFIP1 PE=1 SV=2 | 32,36 | 2,95 | 41738 | 1 | 19,49 | 20,81 | 21,53 | 18,17 | 20,89 | 21,38 | **0,726** | 7,12E-01 | **0** | -0,4613489 | 0,14755952 |
| NOTC2_HUM | NOTCH2 | sp\|Q04721\|NOTC2_HUMAN Neurogenic locus notch homolog protein 2 OS=Homo sapie | 95,619 | 2,35 | 265405 | 4 | 22,86 | 23,00 | 21,07 | 22,70 | 22,70 | 20,14 | **0,725** | 6,83E-01 | **0** | -0,4633331 | 0,16534367 |
| GTPB1_HUM | GTPBP1 | sp\|O00178\|GTPB1_HUMAN GTP-binding protein 1 OS=Homo sapiens GN=GTPBP1 PE=1 | 243,875 | 17,64 | 72454 | 7 | 23,99 | 24,09 | 24,57 | 22,67 | 24,27 | 24,32 | **0,725** | 4,62E-01 | **0** | -0,4633331 | 0,33499223 |
| CSPG4_HUM | CSPG4 | sp\|Q6UVK1\|CSPG4_HUMAN Chondroitin sulfate proteoglycan 4 OS=Homo sapiens GN= | 410,599 | 2,45 | 250537 | 11 | 28,06 | 27,66 | 25,85 | 27,64 | 27,35 | 25,19 | **0,725** | 6,76E-01 | **0** | -0,4633331 | 0,17009078 |
| AGFG1_HUM | AGFG1 | sp\|P52594\|AGFG1_HUMAN Arf-GAP domain and FG repeat-containing protein 1 OS=Ho | 69,272 | 8,19 | 58260 | 3 | 20,86 | 21,16 | 22,32 | 20,90 | 20,25 | 21,80 | **0,725** | 5,04E-01 | **0** | -0,4633338 | 0,29715414 |
| UFO_HUMA | AXL | sp\|P30530\|UFO_HUMAN Tyrosine-protein kinase receptor UFO OS=Homo sapiens GN= | 306,221 | 12,64 | 98336 | 9 | 25,43 | 25,45 | 23,95 | 25,28 | 25,19 | 22,97 | **0,725** | 6,35E-01 | **0** | -0,4633338 | 0,19700847 |
| SPCS1_HUM | SPCS1 | sp\|Q9Y6A9\|SPCS1_HUMAN Signal peptidase complex subunit 1 OS=Homo sapiens GN=S | 18,969 | 14,71 | 11805 | 1 | 21,84 | 22,12 | 19,23 | 22,04 | 21,83 | 17,92 | **0,724** | 7,89E-01 | **0** | -0,4655253 | 0,10295236 |
| ACE_HUMAN | ACE | sp\|P12821\|ACE_HUMAN Angiotensin-converting enzyme OS=Homo sapiens GN=ACE PE | 31,222 | 1,07 | 149715 | 1 | 17,13 | 18,74 | 19,75 | 17,53 | 17,54 | 19,15 | **0,724** | 6,44E-01 | **0** | -0,4656671 | 0,1910338 |
| DCTN2_HUM | DCTN2 | sp\|Q13561\|DCTN2_HUMAN Dynactin subunit 2 OS=Homo sapiens GN=DCTN2 PE=1 SV= | 146,402 | 17,96 | 44231 | 6 | 24,63 | 24,52 | 23,52 | 23,54 | 24,12 | 23,61 | **0,724** | 3,06E-01 | **0** | -0,4666659 | 0,51480263 |
| BAF_HUMAN | BANF1 | sp\|O75531\|BAF_HUMAN Barrier-to-autointegration factor OS=Homo sapiens GN=BANF | 62,969 | 40,45 | 10059 | 2 | 22,24 | 22,71 | 22,45 | 21,61 | 22,53 | 21,86 | **0,724** | 2,02E-01 | **0** | -0,4666659 | 0,69364212 |
| NUP43_HUM | NUP43 | sp\|Q8NFH3\|NUP43_HUMAN Nucleoporin Nup43 OS=Homo sapiens GN=NUP43 PE=1 SV | 29,037 | 4,47 | 42151 | 1 | 23,43 | 21,35 | 22,50 | 22,34 | 21,27 | 22,27 | **0,724** | 5,38E-01 | **0** | -0,4666665 | 0,26919751 |
| SH3G1_HUM | SH3GL1 | sp\|Q99961\|SH3G1_HUMAN Endophilin-A2 OS=Homo sapiens GN=SH3GL1 PE=1 SV=1 | 465,74 | 19,57 | 41490 | 13 | 29,27 | 29,10 | 28,65 | 29,01 | 28,52 | 28,09 | **0,724** | 2,23E-01 | **0** | -0,4666665 | 0,65176841 |
| EXOC4_HUM | EXOC4 | sp\|Q96A65\|EXOC4_HUMAN Exocyst complex component 4 OS=Homo sapiens GN=EXO | 154,833 | 7,6 | 110498 | 5 | 21,58 | 22,15 | 23,97 | 21,97 | 22,00 | 22,33 | **0,724** | 5,57E-01 | **0** | -0,4666665 | 0,25386859 |
| DNJB4_HUM | DNAJB4 | sp\|Q9UDY4\|DNJB4_HUMAN DnaJ homolog subfamily B member 4 OS=Homo sapiens GN | 361,958 | 27 | 37807 | 10 | 26,37 | 26,12 | 25,92 | 25,86 | 25,72 | 25,43 | **0,724** | 6,20E-02 | **0** | -0,4666672 | 1,20765657 |
| PGFRB_HUM | PDGFRB | sp\|P09619\|PGFRB_HUMAN Platelet-derived growth factor receptor beta OS=Homo sapi | 412,287 | 14,65 | 123968 | 13 | 26,29 | 25,99 | 26,20 | 25,73 | 25,73 | 25,62 | **0,724** | 8,32E-03 | **0** | -0,4666672 | 2,08000353 |
| SLAI2_HUMA | SLAIN2 | sp\|Q9P270\|SLAI2_HUMAN SLAIN motif-containing protein 2 OS=Homo sapiens GN=SLA | 11,76 | 2,07 | 62543 | 1 | 20,44 | 20,60 | 19,85 | 19,89 | 20,24 | 19,35 | **0,722** | 2,45E-01 | **0** | -0,4696433 | 0,61066611 |
| FZD7_HUMA | FZD7 | sp\|O75084\|FZD7_HUMAN Frizzled-7 OS=Homo sapiens GN=FZD7 PE=1 SV=2 | 59,215 | 4,18 | 63620 | 3 | 22,13 | 22,55 | 21,57 | 21,75 | 22,01 | 21,08 | **0,722** | 3,02E-01 | **0** | -0,4699993 | 0,52053145 |
| RELL1_HUM | RELL1 | sp\|Q8IUW5\|RELL1_HUMAN RELT-like protein 1 OS=Homo sapiens GN=RELL1 PE=1 SV=1 | 53,168 | 13,65 | 29340 | 3 | 22,46 | 22,99 | 22,20 | 22,94 | 22,24 | 21,06 | **0,722** | 4,74E-01 | **0** | -0,4699999 | 0,32393214 |
| TP4A2_HUM | PTP4A2 | sp\|Q12974\|TP4A2_HUMAN Protein tyrosine phosphatase type IVA 2 OS=Homo sapiens | 257,763 | 73,05 | 19127 | 9 | 25,04 | 25,27 | 24,17 | 24,71 | 24,58 | 23,78 | **0,722** | 3,49E-01 | **0** | -0,4700006 | 0,45707989 |
| LZTR1_HUM | LZTR1 | sp\|Q8N653\|LZTR1_HUMAN Leucine-zipper-like transcriptional regulator 1 OS=Homo sa | 31,787 | 2,5 | 94719 | 1 | 17,07 | 18,59 | 18,37 | 18,01 | 16,82 | 17,78 | **0,721** | 4,74E-01 | **0** | -0,4722678 | 0,32448009 |
| MB12B_HUM | MVB12B | sp\|Q9H7P6\|MB12B_HUMAN Multivesicular body subunit 12B OS=Homo sapiens GN=M | 548,971 | 59,25 | 35620 | 12 | 26,65 | 26,47 | 25,40 | 26,37 | 26,18 | 24,55 | **0,720** | 5,34E-01 | **0** | -0,4733327 | 0,27217265 |
| FETA_HUMA | AFP | sp\|P02771\|FETA_HUMAN Alpha-fetoprotein OS=Homo sapiens GN=AFP PE=1 SV=1 | 63,409 | 6,73 | 68678 | 4 | 24,89 | 23,49 | 25,84 | 24,84 | 22,92 | 25,04 | **0,720** | 6,48E-01 | **0** | -0,4733327 | 0,18843921 |
| PGS1_HUMA | BGN | sp\|P21810\|PGS1_HUMAN Biglycan OS=Homo sapiens GN=BGN PE=1 SV=2 | 138,484 | 16,3 | 41654 | 6 | 25,41 | 24,80 | 25,37 | 25,33 | 24,26 | 24,57 | **0,720** | 2,74E-01 | **0** | -0,4733334 | 0,56176022 |
| VPS16_HUM | VPS16 | sp\|Q9H269\|VPS16_HUMAN Vacuolar protein sorting-associated protein 16 homolog OS | 46,269 | 3,69 | 94694 | 2 | 23,28 | 23,34 | 23,51 | 21,95 | 22,94 | 23,81 | **0,719** | 4,29E-01 | **0** | -0,4766668 | 0,36796448 |
| RICTR_HUM | RICTOR | sp\|Q6R327\|RICTR_HUMAN Rapamycin-insensitive companion of mTOR OS=Homo sapie | 28,803 | 1 | 192218 | 1 | 19,18 | 19,37 | 18,27 | 18,27 | 18,95 | 18,17 | **0,719** | 3,18E-01 | **0** | -0,4766668 | 0,49695434 |
| YIPF6_HUMA | YIPF6 | sp\|Q96EC8\|YIPF6_HUMAN Protein YIPF6 OS=Homo sapiens GN=YIPF6 PE=1 SV=2 | 18,19 | 3,39 | 26256 | 1 | 21,26 | 20,61 | 18,78 | 21,42 | 20,97 | 16,83 | **0,719** | 7,86E-01 | **0** | -0,4769052 | 0,10482398 |
| CRIM1_HUM | CRIM1 | sp\|Q9NZV1\|CRIM1_HUMAN Cysteine-rich motor neuron 1 protein OS=Homo sapiens G | 55,591 | 2,9 | 113738 | 2 | 21,72 | 21,30 | 21,87 | 21,38 | 21,38 | 20,69 | **0,717** | 1,69E-01 | **0** | -0,4800002 | 0,77211323 |
| ANKH_HUM | ANKH | sp\|Q9HCJ1\|ANKH_HUMAN Progressive ankylosis protein homolog OS=Homo sapiens G | 23,59 | 4,27 | 54241 | 1 | 18,77 | 18,92 | 19,07 | 18,59 | 17,92 | 18,81 | **0,717** | 1,63E-01 | **0** | -0,4800002 | 0,78743606 |
| MYH11_HUM | MYH11 | sp\|P35749\|MYH11_HUMAN Myosin-11 OS=Homo sapiens GN=MYH11 PE=1 SV=3 | 723,698 | 8,06 | 227339 | 17 | 23,94 | 24,35 | 22,84 | 23,85 | 23,81 | 22,03 | **0,717** | 5,57E-01 | **0** | -0,4800002 | 0,25392596 |
| ANTR1_HUM | ANTXR1 | sp\|Q9H6X2\|ANTR1_HUMAN Anthrax toxin receptor 1 OS=Homo sapiens GN=ANTXR1 P | 107,728 | 7,45 | 62789 | 3 | 21,79 | 22,12 | 21,72 | 21,24 | 21,21 | 21,74 | **0,717** | 8,58E-02 | **0** | -0,4800008 | 1,06635781 |
| RAB12_HUM | RAB12 | sp\|Q6IQ22\|RAB12_HUMAN Ras-related protein Rab-12 OS=Homo sapiens GN=RAB12 P | 205,81 | 27,46 | 27248 | 8 | 25,05 | 25,02 | 23,96 | 24,37 | 24,69 | 23,52 | **0,715** | 3,89E-01 | **0** | -0,4833323 | 0,41032153 |
| MYD88_HUM | MYD88 | sp\|Q99836\|MYD88_HUMAN Myeloid differentiation primary response protein MyD88 | 40,103 | 6,76 | 33233 | 2 | 19,25 | 20,25 | 20,85 | 19,04 | 18,28 | 21,58 | **0,715** | 6,83E-01 | **0** | -0,484258 | 0,16572732 |
| FEN1_HUMA | FEN1 | sp\|P39748\|FEN1_HUMAN Flap endonuclease 1 OS=Homo sapiens GN=FEN1 PE=1 SV=1 | 63,208 | 7,89 | 42593 | 4 | 26,12 | 24,66 | 25,13 | 25,10 | 24,93 | 24,42 | **0,714** | 3,65E-01 | **0** | -0,4866664 | 0,43808279 |
| VPS36_HUM | VPS36 | sp\|Q86VN1\|VPS36_HUMAN Vacuolar protein-sorting-associated protein 36 OS=Homo s | 385,948 | 25,65 | 43817 | 10 | 26,49 | 26,21 | 24,56 | 25,95 | 25,63 | 24,22 | **0,714** | 5,77E-01 | **0** | -0,4866664 | 0,23865402 |
| RAB4A_HUM | RAB4A | sp\|P20338\|RAB4A_HUMAN Ras-related protein Rab-4A OS=Homo sapiens GN=RAB4A P | 151,436 | 20,64 | 24390 | 4 | 23,44 | 22,62 | 22,43 | 23,17 | 22,60 | 21,26 | **0,714** | 4,93E-01 | **0** | -0,486667 | 0,30736475 |
| EPHB4_HUM | EPHB4 | sp\|P54760\|EPHB4_HUMAN Ephrin type-B receptor 4 OS=Homo sapiens GN=EPHB4 PE= | 503,768 | 15,81 | 108270 | 13 | 25,78 | 25,59 | 24,66 | 25,14 | 25,13 | 24,30 | **0,714** | 3,35E-01 | **0** | -0,4866676 | 0,47535528 |
| DDX18_HUM | DDX18 | sp\|Q9NVP1\|DDX18_HUMAN ATP-dependent RNA helicase DDX18 OS=Homo sapiens GN | 25,653 | 2,09 | 75407 | 1 | 18,04 | 18,46 | 20,52 | 17,31 | 17,79 | 20,46 | **0,713** | 7,15E-01 | **0** | -0,4879328 | 0,14569213 |
| GNAI3_HUM | GNAI3 | sp\|P08754\|GNAI3_HUMAN Guanine nucleotide-binding protein G(k) subunit alpha OS= | 967,459 | 55,93 | 40532 | 20 | 27,73 | 27,76 | 26,84 | 27,35 | 27,44 | 26,07 | **0,712** | 4,12E-01 | **0** | -0,4899998 | 0,38511159 |
| NB5R1_HUM | CYB5R1 | sp\|Q9UHQ9\|NB5R1_HUMAN NADH-cytochrome b5 reductase 1 OS=Homo sapiens GN= | 52,404 | 7,54 | 34095 | 2 | 21,92 | 22,34 | 22,96 | 21,72 | 21,76 | 22,27 | **0,712** | 2,34E-01 | **0** | -0,4899998 | 0,63033673 |
| GGA3_HUM | GGA3 | sp\|Q9NZ52\|GGA3_HUMAN ADP-ribosylation factor-binding protein GGA3 OS=Homo sa | 16,456 | 3,46 | 78315 | 1 | 18,44 | 18,56 | 19,37 | 17,95 | 17,92 | 19,03 | **0,712** | 3,54E-01 | **0** | -0,4899998 | 0,45127232 |
| MB12A_HUM | MVB12A | sp\|Q96EY5\|MB12A_HUMAN Multivesicular body subunit 12A OS=Homo sapiens GN=M | 61,841 | 6,59 | 28783 | 1 | 25,57 | 25,60 | 24,70 | 25,15 | 25,21 | 24,04 | **0,712** | 3,66E-01 | **0** | -0,4900004 | 0,43610524 |
| DAB2_HUMA | DAB2 | sp\|P98082\|DAB2_HUMAN Disabled homolog 2 OS=Homo sapiens GN=DAB2 PE=1 SV=3 | 279,707 | 9,74 | 82448 | 7 | 25,32 | 24,82 | 25,28 | 24,82 | 24,66 | 24,47 | **0,712** | 6,11E-02 | **0** | -0,4900004 | 1,21405157 |
| MOT1_HUM | SLC16A1 | sp\|P53985\|MOT1_HUMAN Monocarboxylate transporter 1 OS=Homo sapiens GN=SLC1 | 147,663 | 11,6 | 53944 | 6 | 23,67 | 24,36 | 21,91 | 23,08 | 23,61 | 21,77 | **0,710** | 6,17E-01 | **0** | -0,4933332 | 0,20962643 |
| CPSF7_HUM | CPSF7 | sp\|Q8N684\|CPSF7_HUMAN Cleavage and polyadenylation specificity factor subunit 7 O | 144,286 | 9,98 | 52050 | 4 | 23,80 | 23,14 | 24,56 | 23,30 | 22,99 | 23,73 | **0,710** | 3,47E-01 | **0** | -0,4933332 | 0,46009345 |
| MRCKA_HUM | CDC42BPA | sp\|Q5VT25\|MRCKA_HUMAN Serine/threonine-protein kinase MRCK alpha OS=Homo sa | 53,128 | 2,94 | 197307 | 5 | 21,72 | 22,08 | 18,18 | 21,23 | 21,25 | 18,02 | **0,710** | 7,79E-01 | **0** | -0,4933332 | 0,10847652 |
| MYO3B_HUM | MYO3B | sp\|Q8WXR4\|MYO3B_HUMAN Myosin-IIIb OS=Homo sapiens GN=MYO3B PE=2 SV=4 | 23,214 | 0,6 | 151829 | 1 | 24,23 | 24,08 | 22,81 | 23,64 | 23,79 | 22,21 | **0,710** | 5,06E-01 | **0** | -0,4933332 | 0,29608997 |

| | | | | | | | | | | | | | | | | | |
|---|---|---|---|---|---|---|---|---|---|---|---|---|---|---|---|---|---|
| PP1G_HUMA | PPP1CC | sp\|P36873\|PP1G_HUMAN Serine/threonine-protein phosphatase PP1-gamma catalytic s | 800,33 | 54,8 | 36984 | 18 | 23,66 | 23,97 | 23,50 | 23,17 | 23,49 | 22,98 | **0,709** | 7,06E-02 | **0** | -0,4966666 | 1,15100793 |
| KPCB_HUMA | PRKCB | sp\|P05771\|KPCB_HUMAN Protein kinase C beta type OS=Homo sapiens GN=PRKCB PE=1 | 122,049 | 7,15 | 76869 | 4 | 23,57 | 22,92 | 22,20 | 22,81 | 22,99 | 21,40 | **0,709** | 4,81E-01 | **0** | -0,4966672 | 0,31796767 |
| UEVLD_HUM | UEVLD | sp\|Q8IX04\|UEVLD_HUMAN Ubiquitin-conjugating enzyme E2 variant 3 OS=Homo sapier | 58,653 | 2,55 | 52264 | 1 | 22,20 | 21,57 | 21,68 | 21,50 | 21,50 | 20,96 | **0,709** | 1,34E-01 | **0** | -0,4966672 | 0,87286611 |
| EFL1_HUMA | EFL1 | sp\|Q7Z2Z2\|EFL1_HUMAN Elongation factor-like GTPase 1 OS=Homo sapiens GN=EFL1 P | 62,459 | 1,43 | 125430 | 1 | 17,72 | 19,13 | 21,69 | 18,04 | 17,46 | 21,55 | **0,708** | 7,87E-01 | **0** | -0,4981467 | 0,10391568 |
| EEA1_HUMA | EEA1 | sp\|Q15075\|EEA1_HUMAN Early endosome antigen 1 OS=Homo sapiens GN=EEA1 PE=1 | 28,709 | 1,2 | 162466 | 1 | 19,90 | 19,74 | 19,64 | 20,03 | 19,99 | 17,76 | **0,708** | 5,45E-01 | **0** | -0,4982751 | 0,26383268 |
| PKP3_HUMA | PKP3 | sp\|Q9Y446\|PKP3_HUMAN Plakophilin-3 OS=Homo sapiens GN=PKP3 PE=1 SV=1 | 23,093 | 1,38 | 87082 | 1 | 21,87 | 21,80 | 18,20 | 21,71 | 20,71 | 17,95 | **0,708** | 7,78E-01 | **0** | -0,4984678 | 0,10892151 |
| S10AD_HUM | S100A13 | sp\|Q99584\|S10AD_HUMAN Protein S100-A13 OS=Homo sapiens GN=S100A13 PE=1 SV= | 21,302 | 11,22 | 11471 | 1 | 21,49 | 22,07 | 18,90 | 21,15 | 20,63 | 19,18 | **0,708** | 6,84E-01 | **0** | -0,498923 | 0,16518514 |
| CTNA2_HUM | CTNNA2 | sp\|P26232\|CTNA2_HUMAN Catenin alpha-2 OS=Homo sapiens GN=CTNNA2 PE=1 SV=5 | 397,661 | 11,65 | 105313 | 10 | 23,38 | 23,64 | 22,67 | 23,02 | 23,26 | 21,91 | **0,707** | 3,80E-01 | **0** | -0,4999994 | 0,42047539 |
| WWP2_HUM | WWP2 | sp\|O00308\|WWP2_HUMAN NEDD4-like E3 ubiquitin-protein ligase WWP2 OS=Homo sa | 421,65 | 16,9 | 98912 | 13 | 25,01 | 24,76 | 25,07 | 24,32 | 24,53 | 24,49 | **0,707** | 1,21E-02 | **0** | -0,5 | 1,91833268 |
| KIRR1_HUM | KIRREL1 | sp\|Q96J84\|KIRR1_HUMAN Kin of IRRE-like protein 1 OS=Homo sapiens GN=KIRREL1 PE= | 677,537 | 28,8 | 83536 | 16 | 26,47 | 26,28 | 24,97 | 26,07 | 25,82 | 24,33 | **0,707** | 5,25E-01 | **0** | -0,5 | 0,27964804 |
| MOB3C_HUI | MOB3C | sp\|Q70IA8\|MOB3C_HUMAN MOB kinase activator 3C OS=Homo sapiens GN=MOB3C PE | 26,019 | 9,72 | 25623 | 1 | 18,20 | 18,10 | 20,73 | 18,60 | 18,60 | 18,33 | **0,707** | 5,94E-01 | **0** | -0,5 | 0,22599794 |
| BLK_HUMAN | BLK | sp\|P51451\|BLK_HUMAN Tyrosine-protein kinase Blk OS=Homo sapiens GN=BLK PE=1 SV | 10,183 | 2,97 | 57706 | 2 | 21,67 | 21,54 | 20,18 | 21,36 | 21,12 | 19,40 | **0,706** | 5,54E-01 | **0** | -0,5032495 | 0,25665251 |
| MARK2_HUN | MARK2 | sp\|Q7KZI7\|MARK2_HUMAN Serine/threonine-protein kinase MARK2 OS=Homo sapiens | 244,626 | 14,97 | 87911 | 9 | 24,57 | 24,37 | 22,30 | 24,02 | 23,87 | 21,84 | **0,705** | 6,44E-01 | **0** | -0,5033328 | 0,19077858 |
| ZYX_HUMAN | ZYX | sp\|Q15942\|ZYX_HUMAN Zyxin OS=Homo sapiens GN=ZYX PE=1 SV=1 | 126,207 | 8,57 | 61277 | 3 | 20,39 | 21,52 | 20,46 | 20,92 | 20,50 | 19,44 | **0,705** | 4,29E-01 | **0** | -0,5033328 | 0,36773477 |
| EVA1B_HUM | EVA1B | sp\|Q9NVM1\|EVA1B_HUMAN Protein eva-1 homolog B OS=Homo sapiens GN=EVA1B PE | 134,791 | 10,3 | 18374 | 2 | 23,98 | 23,47 | 21,04 | 23,97 | 23,81 | 19,20 | **0,705** | 7,95E-01 | **0** | -0,5033334 | 0,09990111 |
| EPHB2_HUM | EPHB2 | sp\|P29323\|EPHB2_HUMAN Ephrin type-B receptor 2 OS=Homo sapiens GN=EPHB2 PE=1 | 626,37 | 22,18 | 117493 | 15 | 24,53 | 25,12 | 22,58 | 24,01 | 24,17 | 22,54 | **0,705** | 6,16E-01 | **0** | -0,5033334 | 0,21059971 |
| HEG1_HUM | HEG1 | sp\|Q9ULI3\|HEG1_HUMAN Protein HEG homolog 1 OS=Homo sapiens GN=HEG1 PE=1 SV | 205,192 | 5,72 | 147461 | 7 | 25,78 | 24,86 | 23,00 | 25,31 | 24,77 | 22,05 | **0,705** | 7,18E-01 | **0** | -0,503334 | 0,14384869 |
| DCBD2_HUN | DCBLD2 | sp\|Q96PD2\|DCBD2_HUMAN Discoidin, CUB and LCCL domain-containing protein 2 OS=H | 56,263 | 4,26 | 85035 | 2 | 22,53 | 21,86 | 21,36 | 21,97 | 21,60 | 20,67 | **0,705** | 3,83E-01 | **0** | -0,503334 | 0,4167036 |
| SYTL4_HUM | SYTL4 | sp\|Q96C24\|SYTL4_HUMAN Synaptotagmin-like protein 4 OS=Homo sapiens GN=SYTL4 F | 16,625 | 1,64 | 76024 | 1 | 21,22 | 20,71 | 19,73 | 21,06 | 20,27 | 18,81 | **0,704** | 5,57E-01 | **0** | -0,5056248 | 0,25432979 |
| FHL2_HUMA | FHL2 | sp\|Q14192\|FHL2_HUMAN Four and a half LIM domains protein 2 OS=Homo sapiens GN= | 180,481 | 24,73 | 32193 | 6 | 23,68 | 24,00 | 24,73 | 23,46 | 23,28 | 24,15 | **0,704** | 2,83E-01 | **0** | -0,5066668 | 0,54881389 |
| HECD1_HUM | HECTD1 | sp\|Q9ULT8\|HECD1_HUMAN E3 ubiquitin-protein ligase HECTD1 OS=Homo sapiens GN=I | 260,549 | 5,21 | 289384 | 9 | 22,80 | 23,36 | 24,38 | 22,46 | 22,98 | 23,58 | **0,704** | 4,20E-01 | **0** | -0,5066668 | 0,37660551 |
| CLCN3_HUM | CLCN3 | sp\|P51790\|CLCN3_HUMAN H(+)/Cl(-) exchange transporter 3 OS=Homo sapiens GN=CL | 13,847 | 0,86 | 90966 | 1 | 21,69 | 21,54 | 21,17 | 21,12 | 21,41 | 20,35 | **0,703** | 2,24E-01 | **0** | -0,5076059 | 0,65046554 |
| IGSF3_HUM | IGSF3 | sp\|O75054\|IGSF3_HUMAN Immunoglobulin superfamily member 3 OS=Homo sapiens G | 169,569 | 6,87 | 135196 | 7 | 24,40 | 24,07 | 23,67 | 24,11 | 23,82 | 22,68 | **0,702** | 3,52E-01 | **0** | -0,5099996 | 0,45326911 |
| PLCD1_HUM | PLCD1 | sp\|P51178\|PLCD1_HUMAN 1-phosphatidylinositol 4,5-bisphosphate phosphodiesterase | 138,605 | 7,94 | 85665 | 7 | 27,28 | 27,42 | 25,19 | 26,88 | 26,86 | 24,62 | **0,702** | 6,50E-01 | **0** | -0,5100002 | 0,18729617 |
| EPN1_HUMA | EPN1 | sp\|Q9Y6I3\|EPN1_HUMAN Epsin-1 OS=Homo sapiens GN=EPN1 PE=1 SV=2 | 153,178 | 9,72 | 60293 | 5 | 24,56 | 24,43 | 24,35 | 24,13 | 24,03 | 23,65 | **0,702** | 3,24E-02 | **0** | -0,5100002 | 1,49009834 |
| LPP_HUMAN | LPP | sp\|Q93052\|LPP_HUMAN Lipoma-preferred partner OS=Homo sapiens GN=LPP PE=1 SV= | 78,89 | 9,8 | 65746 | 4 | 22,97 | 23,43 | 22,85 | 22,75 | 22,49 | 22,48 | **0,702** | 6,13E-02 | **0** | -0,5100002 | 1,21264204 |
| SPAST_HUM | SPAST | sp\|Q9UBP0\|SPAST_HUMAN Spastin OS=Homo sapiens GN=SPAST PE=1 SV=1 | 66,79 | 3,08 | 67197 | 1 | 18,95 | 18,92 | 19,91 | 18,51 | 18,13 | 19,61 | **0,702** | 4,06E-01 | **0** | -0,5100358 | 0,39110992 |
| RIT1_HUMA | RIT1 | sp\|Q92963\|RIT1_HUMAN GTP-binding protein Rit1 OS=Homo sapiens GN=RIT1 PE=1 SV | 26,816 | 5,48 | 25145 | 1 | 20,25 | 20,16 | 19,87 | 19,86 | 20,02 | 18,86 | **0,701** | 2,49E-01 | **0** | -0,5130564 | 0,60434297 |
| CNTFR_HUM | CNTFR | sp\|P26992\|CNTFR_HUMAN Ciliary neurotrophic factor receptor subunit alpha OS=Homo | 13,419 | 3,49 | 40633 | 1 | 20,05 | 20,03 | 20,40 | 19,87 | 19,11 | 19,96 | **0,701** | 1,57E-01 | **0** | -0,513333 | 0,80416957 |
| ASPC1_HUM | ASPSCR1 | sp\|Q9BZE9\|ASPC1_HUMAN Tether containing UBX domain for GLUT4 OS=Homo sapiens | 17,506 | 2,89 | 60183 | 1 | 19,15 | 19,60 | 19,89 | 17,74 | 19,49 | 19,87 | **0,701** | 4,98E-01 | **0** | -0,513333 | 0,30240427 |
| WDFY1_HUN | WDFY1 | sp\|Q8IWB7\|WDFY1_HUMAN WD repeat and FYVE domain-containing protein 1 OS=Hor | 52,378 | 10 | 46324 | 3 | 21,42 | 21,66 | 21,47 | 20,97 | 20,54 | 21,50 | **0,701** | 1,48E-01 | **0** | -0,513333 | 0,82889216 |
| NEUL_HUMA | NLN | sp\|Q9BYT8\|NEUL_HUMAN Neurolysin, mitochondrial OS=Homo sapiens GN=NLN PE=1 S | 111,545 | 5,54 | 80652 | 4 | 22,55 | 23,07 | 21,86 | 22,68 | 22,75 | 20,50 | **0,699** | 5,62E-01 | **0** | -0,5166664 | 0,25048289 |
| SPNS1_HUM | SPNS1 | sp\|Q9H2V7\|SPNS1_HUMAN Protein spinster homolog 1 OS=Homo sapiens GN=SPNS1 P | 144,878 | 10,61 | 56630 | 2 | 20,70 | 21,15 | 18,79 | 19,70 | 20,21 | 19,18 | **0,699** | 5,45E-01 | **0** | -0,516667 | 0,26361256 |
| TFCP2_HUM | TFCP2 | sp\|Q12800\|TFCP2_HUMAN Alpha-globin transcription factor CP2 OS=Homo sapiens GN= | 28,766 | 3,39 | 57256 | 1 | 18,31 | 19,53 | 19,53 | 17,40 | 19,37 | 19,05 | **0,699** | 5,20E-01 | **0** | -0,516667 | 0,28399343 |
| VLDLR_HUM | VLDLR | sp\|P98155\|VLDLR_HUMAN Very low-density lipoprotein receptor OS=Homo sapiens GN | 10,119 | 1,15 | 96098 | 1 | 20,04 | 18,25 | 19,45 | 19,01 | 18,59 | 18,59 | **0,699** | 3,96E-01 | **0** | -0,5170193 | 0,40217621 |
| PFD5_HUMA | PFDN5 | sp\|Q99471\|PFD5_HUMAN Prefoldin subunit 5 OS=Homo sapiens GN=PFDN5 PE=1 SV=2 | 33,659 | 9,74 | 17328 | 1 | 18,53 | 18,78 | 20,59 | 18,05 | 17,56 | 20,73 | **0,698** | 6,83E-01 | **0** | -0,5180728 | 0,1656589 |
| IF122_HUMA | IFT122 | sp\|Q9HBG6\|IF122_HUMAN Intraflagellar transport protein 122 homolog OS=Homo sapi | 11,881 | 1,05 | 141825 | 1 | 17,53 | 18,41 | 18,70 | 17,41 | 17,28 | 18,40 | **0,698** | 3,57E-01 | **0** | -0,5180982 | 0,44694389 |
| KCC2G_HUM | CAMK2G | sp\|Q13555\|KCC2G_HUMAN Calcium/calmodulin-dependent protein kinase type II subu | 82,515 | 10,04 | 62609 | 5 | 21,49 | 21,93 | 22,83 | 20,70 | 21,37 | 22,62 | **0,697** | 4,91E-01 | **0** | -0,5199992 | 0,3086745 |
| CTNA1_HUM | CTNNA1 | sp\|P35221\|CTNA1_HUMAN Catenin alpha-1 OS=Homo sapiens GN=CTNNA1 PE=1 SV=1 | 1766,971 | 55,96 | 100071 | 42 | 28,38 | 28,33 | 27,12 | 27,88 | 27,95 | 26,44 | **0,697** | 4,63E-01 | **0** | -0,5199998 | 0,33421471 |
| IGF1R_HUM | IGF1R | sp\|P08069\|IGF1R_HUMAN Insulin-like growth factor 1 receptor OS=Homo sapiens GN=I | 1131,189 | 28,24 | 154793 | 29 | 27,27 | 27,13 | 25,60 | 26,98 | 26,70 | 24,76 | **0,697** | 5,86E-01 | **0** | -0,5199998 | 0,23202482 |
| PAK4_HUMA | PAK4 | sp\|O96013\|PAK4_HUMAN Serine/threonine-protein kinase PAK 4 OS=Homo sapiens GN | 434,897 | 24,03 | 64072 | 11 | 25,94 | 25,78 | 24,41 | 25,50 | 25,02 | 24,05 | **0,697** | 4,66E-01 | **0** | -0,5200005 | 0,33150124 |
| IMA5_HUMA | KPNA1 | sp\|P52294\|IMA5_HUMAN Importin subunit alpha-5 OS=Homo sapiens GN=KPNA1 PE=1 | 194,521 | 13,01 | 60222 | 5 | 22,30 | 22,08 | 22,34 | 22,14 | 21,31 | 21,70 | **0,696** | 1,07E-01 | **0** | -0,5233332 | 0,96893373 |
| SHLB1_HUM | SH3GLB1 | sp\|Q9Y371\|SHLB1_HUMAN Endophilin-B1 OS=Homo sapiens GN=SH3GLB1 PE=1 SV=1 | 105,551 | 8,22 | 40796 | 2 | 22,36 | 22,23 | 21,72 | 21,52 | 21,89 | 21,33 | **0,696** | 1,10E-01 | **0** | -0,5233332 | 0,95977242 |
| ADT3_HUMA | SLC25A6 | sp\|P12236\|ADT3_HUMAN ADP/ATP translocase 3 OS=Homo sapiens GN=SLC25A6 PE=1 | 534,867 | 39,93 | 32866 | 13 | 23,86 | 24,06 | 25,84 | 23,44 | 23,86 | 24,89 | **0,696** | 5,30E-01 | **0** | -0,5233332 | 0,27549701 |
| ELN_HUMAN | ELN | sp\|P15502\|ELN_HUMAN Elastin OS=Homo sapiens GN=ELN PE=1 SV=3 | 37,231 | 2,42 | 68469 | 1 | 23,79 | 21,85 | 26,07 | 23,06 | 21,49 | 25,59 | **0,696** | 7,74E-01 | **0** | -0,5233339 | 0,1110055 |
| DCTN3_HUN | DCTN3 | sp\|O75935\|DCTN3_HUMAN Dynactin subunit 3 OS=Homo sapiens GN=DCTN3 PE=1 SV= | 22,015 | 4,3 | 21119 | 1 | 19,36 | 20,53 | 21,48 | 18,05 | 19,82 | 21,93 | **0,696** | 7,03E-01 | **0** | -0,5233339 | 0,15293037 |
| STT3B_HUM | STT3B | sp\|Q8TCJ2\|STT3B_HUMAN Dolichyl-diphosphooligosaccharide--protein glycosyltransfe | 43,177 | 3,15 | 93674 | 3 | 21,47 | 20,56 | 22,64 | 17,72 | 22,06 | 23,32 | **0,695** | 7,85E-01 | **0** | -0,5240364 | 0,10486825 |
| ITM2B_HUM | ITM2B | sp\|Q9Y287\|ITM2B_HUMAN Integral membrane protein 2B OS=Homo sapiens GN=ITM2 | 34,887 | 3,76 | 30338 | 1 | 21,97 | 21,74 | 19,20 | 22,00 | 21,41 | 17,93 | **0,695** | 7,51E-01 | **0** | -0,5254695 | 0,12408783 |
| SRC8_HUMA | CTTN | sp\|Q14247\|SRC8_HUMAN Src substrate cortactin OS=Homo sapiens GN=CTTN PE=1 SV= | 283,904 | 17,27 | 61586 | 10 | 24,22 | 24,58 | 24,05 | 24,17 | 23,59 | 23,51 | **0,694** | 1,13E-01 | **0** | -0,526666 | 0,94739007 |
| TRI18_HUM | MID1 | sp\|O15344\|TRI18_HUMAN E3 ubiquitin-protein ligase Midline-1 OS=Homo sapiens GN= | 21,344 | 1,8 | 75251 | 1 | 19,85 | 19,43 | 19,57 | 19,28 | 18,74 | 19,25 | **0,694** | 6,99E-02 | **0** | -0,5266666 | 1,1554559 |
| CO6A1_HUN | COL6A1 | sp\|P12109\|CO6A1_HUMAN Collagen alpha-1(VI) chain OS=Homo sapiens GN=COL6A1 P | 234,014 | 10,12 | 108529 | 8 | 27,20 | 26,63 | 26,92 | 26,85 | 25,40 | 26,92 | **0,694** | 3,70E-01 | **0** | -0,5266666 | 0,43167904 |
| VPS28_HUM | VPS28 | sp\|Q9UK41\|VPS28_HUMAN Vacuolar protein sorting-associated protein 28 homolog OS | 728,786 | 47,06 | 25425 | 16 | 28,27 | 28,17 | 27,16 | 27,68 | 27,71 | 26,63 | **0,694** | 3,53E-01 | **0** | -0,5266673 | 0,4520687 |
| PLCD3_HUM | PLCD3 | sp\|Q8N3E9\|PLCD3_HUMAN 1-phosphatidylinositol 4,5-bisphosphate phosphodiesteras | 121,981 | 6,84 | 89258 | 5 | 23,61 | 23,75 | 20,91 | 23,46 | 23,21 | 20,02 | **0,694** | 7,33E-01 | **0** | -0,5266673 | 0,13460386 |
| ATG2A_HUN | ATG2A | sp\|Q2TAZ0\|ATG2A_HUMAN Autophagy-related protein 2 homolog A OS=Homo sapiens | 10,557 | 0,36 | 212860 | 1 | 24,95 | 23,35 | 23,74 | 24,07 | 23,72 | 22,67 | **0,694** | 4,56E-01 | **0** | -0,5266673 | 0,34060145 |
| IBP2_HUMA | IGFBP2 | sp\|P18065\|IBP2_HUMAN Insulin-like growth factor-binding protein 2 OS=Homo sapiens | 31,756 | 3,69 | 34814 | 1 | 18,72 | 19,75 | 19,35 | 17,79 | 20,37 | 18,08 | **0,694** | 5,77E-01 | **0** | -0,52742 | 0,23873074 |
| ES8L2_HUM | EPS8L2 | sp\|Q9H6S3\|ES8L2_HUMAN Epidermal growth factor receptor kinase substrate 8-like pr | 70,075 | 4,76 | 80621 | 3 | 21,34 | 22,35 | 18,72 | 20,19 | 21,99 | 18,64 | **0,693** | 7,34E-01 | **0** | -0,5293204 | 0,13435514 |
| CELF1_HUM | CELF1 | sp\|Q92879\|CELF1_HUMAN CUGBP Elav-like family member 1 OS=Homo sapiens GN=CE | 551,655 | 34,77 | 52063 | 15 | 25,74 | 25,81 | 25,71 | 25,53 | 24,87 | 25,27 | **0,693** | 5,25E-02 | **0** | -0,5299988 | 1,27978841 |
| TM1L2_HUN | TOM1L2 | sp\|Q6ZVM7\|TM1L2_HUMAN TOM1-like protein 2 OS=Homo sapiens GN=TOM1L2 PE=1 | 280,639 | 16,77 | 55556 | 6 | 24,72 | 24,56 | 23,62 | 24,20 | 24,18 | 22,93 | **0,693** | 3,84E-01 | **0** | -0,5299994 | 0,41588819 |
| RB22A_HUM | RAB22A | sp\|Q9UL26\|RB22A_HUMAN Ras-related protein Rab-22A OS=Homo sapiens GN=RAB22 | 280,519 | 42,78 | 21855 | 6 | 23,35 | 23,22 | 21,27 | 23,24 | 22,90 | 20,11 | **0,693** | 6,81E-01 | **0** | -0,5300001 | 0,16677924 |
| KC1A_HUMA | CSNK1A1 | sp\|P48729\|KC1A_HUMAN Casein kinase I isoform alpha OS=Homo sapiens GN=CSNK1A | 285,471 | 29,08 | 38915 | 10 | 26,01 | 25,69 | 26,15 | 25,26 | 25,35 | 25,65 | **0,693** | 4,23E-02 | **0** | -0,5300001 | 1,37401547 |
| ITCH_HUMA | ITCH | sp\|Q96J02\|ITCH_HUMAN E3 ubiquitin-protein ligase Itchy homolog OS=Homo sapiens G | 1717,414 | 48,62 | 102803 | 39 | 28,60 | 28,22 | 27,84 | 28,07 | 27,79 | 27,21 | **0,693** | 1,89E-01 | **0** | -0,5300001 | 0,72386568 |
| KDIS_HUMA | KIDINS220 | sp\|Q9ULH0\|KDIS_HUMAN Kinase D-interacting substrate of 220 kDa OS=Homo sapiens | 32,662 | 1,19 | 196542 | 2 | 21,90 | 21,10 | 18,94 | 20,62 | 20,96 | 18,76 | **0,692** | 6,59E-01 | **0** | -0,5321547 | 0,18117838 |
| K2C74_HUM | KRT74 | sp\|Q7RTS7\|K2C74_HUMAN Keratin, type II cytoskeletal 74 OS=Homo sapiens GN=KRT7 | 177,196 | 9,26 | 57865 | 7 | 21,74 | 17,90 | 23,16 | 21,92 | 16,82 | 22,46 | **0,691** | 8,34E-01 | **0** | -0,5331039 | 0,07865543 |
| DYL1_HUMA | DYNLL1 | sp\|P63167\|DYL1_HUMAN Dynein light chain 1, cytoplasmic OS=Homo sapiens GN=DYNL | 236,104 | 65,17 | 10366 | 6 | 23,33 | 23,13 | 23,57 | 23,00 | 22,70 | 22,73 | **0,691** | 2,84E-02 | **0** | -0,5333328 | 1,54592528 |
| MOV10_HUN | MOV10 | sp\|Q9HCE1\|MOV10_HUMAN Putative helicase MOV-10 OS=Homo sapiens GN=MOV10 | 666,371 | 22,03 | 113671 | 19 | 27,70 | 27,35 | 26,27 | 26,97 | 26,71 | 26,04 | **0,691** | 3,56E-01 | **0** | -0,5333341 | 0,44827198 |
| QCR9_HUM | UQCR10 | sp\|Q9UDW1\|QCR9_HUMAN Cytochrome b-c1 complex subunit 9 OS=Homo sapiens GN | 18,489 | 11,11 | 7308 | 1 | 21,17 | 20,99 | 17,53 | 18,13 | 20,84 | 19,12 | **0,691** | 7,27E-01 | **0** | -0,5342236 | 0,13868938 |
| RAI14_HUM | RAI14 | sp\|Q9P0K7\|RAI14_HUMAN Ankycorbin OS=Homo sapiens GN=RAI14 PE=1 SV=2 | 139,742 | 5 | 110041 | 4 | 22,76 | 23,03 | 19,92 | 22,87 | 22,57 | 18,67 | **0,690** | 7,66E-01 | **0** | -0,5347411 | 0,11564676 |
| CO1A1_HUN | COL1A1 | sp\|P02452\|CO1A1_HUMAN Collagen alpha-1(I) chain OS=Homo sapiens GN=COL1A1 PE | 389,211 | 9,08 | 138941 | 12 | 25,57 | 24,61 | 26,95 | 25,29 | 23,98 | 26,25 | **0,689** | 6,01E-01 | **0** | -0,5366669 | 0,22137459 |
| OGDHL_HUN | OGDHL | sp\|Q9ULD0\|OGDHL_HUMAN 2-oxoglutarate dehydrogenase-like, mitochondrial OS=Ho | 77,161 | 3,47 | 114481 | 4 | 21,96 | 22,10 | 19,33 | 21,02 | 21,01 | 19,74 | **0,689** | 6,18E-01 | **0** | -0,5381165 | 0,20894586 |
| S10A6_HUM | S100A6 | sp\|P06703\|S10A6_HUMAN Protein S100-A6 OS=Homo sapiens GN=S100A6 PE=1 SV=1 | 16,981 | 18,89 | 10180 | 1 | 18,03 | 17,88 | 20,35 | 17,63 | 17,15 | 19,87 | **0,688** | 6,66E-01 | **0** | -0,5385558 | 0,17643648 |
| MIRO2_HUN | RHOT2 | sp\|Q8IXI1\|MIRO2_HUMAN Mitochondrial Rho GTPase 2 OS=Homo sapiens GN=RHOT2 | 30,9 | 1,46 | 68118 | 1 | 24,29 | 24,28 | 22,84 | 23,85 | 23,76 | 22,18 | **0,688** | 4,98E-01 | **0** | -0,5400003 | 0,30282718 |
| AT2B3_HUM | ATP2B3 | sp\|Q16720\|AT2B3_HUMAN Plasma membrane calcium-transporting ATPase 3 OS=Hom | 773,611 | 12,7 | 134197 | 18 | 24,43 | 24,33 | 22,74 | 23,20 | 23,96 | 22,72 | **0,688** | 4,56E-01 | **0** | -0,5400003 | 0,34057134 |
| NLTP_HUMA | SCP2 | sp\|P22307\|NLTP_HUMAN Non-specific lipid-transfer protein OS=Homo sapiens GN=SCP | 41,322 | 3,66 | 58994 | 2 | 20,28 | 20,58 | 22,76 | 20,70 | 20,05 | 21,24 | **0,686** | 5,59E-01 | **0** | -0,5433337 | 0,25247316 |
| NEDD4_HUN | NEDD4 | sp\|P46934\|NEDD4_HUMAN E3 ubiquitin-protein ligase NEDD4 OS=Homo sapiens GN=N | 371,704 | 12,51 | 149114 | 13 | 25,33 | 25,05 | 24,40 | 24,59 | 24,53 | 24,02 | **0,685** | 1,72E-01 | **0** | -0,5466658 | 0,76333526 |
| CTDS2_HUM | CTDSP2 | sp\|O14595\|CTDS2_HUMAN Carboxy-terminal domain RNA polymerase II polypeptide A | 92,262 | 18,45 | 30664 | 4 | 22,64 | 22,25 | 22,75 | 22,37 | 21,98 | 21,65 | **0,685** | 1,01E-01 | **0** | -0,5466665 | 0,99562665 |
| STEA2_HUM | STEAP2 | sp\|Q8NFT2\|STEA2_HUMAN Metalloreductase STEAP2 OS=Homo sapiens GN=STEAP2 PE | 19,41 | 1,63 | 56056 | 1 | 23,73 | 23,70 | 22,05 | 23,42 | 22,48 | 21,94 | **0,685** | 4,81E-01 | **0** | -0,5466665 | 0,31823324 |
| COMP_HUM | COMP | sp\|P49747\|COMP_HUMAN Cartilage oligomeric matrix protein OS=Homo sapiens GN=C | 56,25 | 6,34 | 82860 | 3 | 21,27 | 18,52 | 21,82 | 21,10 | 17,93 | 20,94 | **0,685** | 7,25E-01 | **0** | -0,5466665 | 0,13938286 |
| EHD2_HUMA | EHD2 | sp\|Q9NZN4\|EHD2_HUMAN EH domain-containing protein 2 OS=Homo sapiens GN=EHD | 1878,257 | 54,33 | 61161 | 42 | 30,79 | 30,42 | 30,65 | 30,38 | 30,03 | 29,81 | **0,685** | 5,08E-02 | **0** | -0,5466671 | 1,29454492 |
| CTND1_HUN | CTNND1 | sp\|O60716\|CTND1_HUMAN Catenin delta-1 OS=Homo sapiens GN=CTNND1 PE=1 SV=1 | 881,032 | 30,68 | 108170 | 26 | 27,79 | 27,78 | 26,51 | 27,27 | 27,10 | 26,07 | **0,685** | 3,89E-01 | **0** | -0,5466671 | 0,40964127 |
| SCAR3_HUM | SCARA3 | sp\|Q6AZY7\|SCAR3_HUMAN Scavenger receptor class A member 3 OS=Homo sapiens GN | 15,866 | 1,16 | 65137 | 1 | 21,00 | 20,65 | 18,67 | 20,28 | 20,16 | 18,23 | **0,683** | 6,07E-01 | **0** | -0,5490233 | 0,21683673 |
| STAM1_HUN | STAM | sp\|Q92783\|STAM1_HUMAN Signal transducing adapter molecule 1 OS=Homo sapiens G | 556,062 | 28,52 | 59180 | 13 | 26,55 | 26,57 | 25,80 | 26,23 | 26,16 | 24,88 | **0,683** | 3,39E-01 | **0** | -0,5499999 | 0,47009459 |
| CTL1_HUMA | SLC44A1 | sp\|Q8WWI5\|CTL1_HUMAN Choline transporter-like protein 1 OS=Homo sapiens GN=SL | 500,75 | 18,87 | 73302 | 11 | 26,66 | 26,59 | 25,67 | 26,19 | 26,23 | 24,85 | **0,683** | 3,77E-01 | **0** | -0,5499999 | 0,42326431 |
| CATD_HUMA | CTSD | sp\|P07339\|CATD_HUMAN Cathepsin D OS=Homo sapiens GN=CTSD PE=1 SV=1 | 171,155 | 13,59 | 44552 | 5 | 26,77 | 26,83 | 25,50 | 26,07 | 26,29 | 25,09 | **0,683** | 3,89E-01 | **0** | -0,5499999 | 0,41036684 |
| BCAM_HUM | BCAM | sp\|P50895\|BCAM_HUMAN Basal cell adhesion molecule OS=Homo sapiens GN=BCAM P | 26,534 | 1,91 | 67405 | 1 | 21,28 | 21,12 | 20,79 | 20,60 | 20,89 | 20,05 | **0,683** | 1,26E-01 | **0** | -0,5500011 | 0,89861808 |
| CPNS2_HUM | CAPNS2 | sp\|Q96L46\|CPNS2_HUMAN Calpain small subunit 2 OS=Homo sapiens GN=CAPNS2 PE=2 | 39,228 | 6,05 | 27660 | 3 | 20,85 | 21,32 | 20,06 | 20,63 | 20,63 | 19,32 | **0,683** | 3,90E-01 | **0** | -0,5506134 | 0,40904407 |
| FZD1_HUMA | FZD1 | sp\|Q9UP38\|FZD1_HUMAN Frizzled-1 OS=Homo sapiens GN=FZD1 PE=1 SV=2 | 12,679 | 3,4 | 71158 | 1 | 17,83 | 19,97 | 20,29 | 18,62 | 18,92 | 18,89 | **0,682** | 5,17E-01 | **0** | -0,5513859 | 0,2863625 |
| GTR8_HUMA | SLC2A8 | sp\|Q9NY64\|GTR8_HUMAN Solute carrier family 2, facilitated glucose transporter memb | 30,922 | 2,52 | 50819 | 1 | 20,98 | 21,10 | 19,25 | 20,89 | 20,65 | 18,14 | **0,682** | 6,31E-01 | **0** | -0,5516904 | 0,19975137 |
| CCD80_HUM | CCDC80 | sp\|Q76M96\|CCD80_HUMAN Coiled-coil domain-containing protein 80 OS=Homo sapien | 55,497 | 2,11 | 108174 | 2 | 22,87 | 22,13 | 18,02 | 21,76 | 21,30 | 18,30 | **0,682** | 7,81E-01 | **0** | -0,5531788 | 0,10740628 |
| SNX27_HUM | SNX27 | sp\|Q96L92\|SNX27_HUMAN Sorting nexin-27 OS=Homo sapiens GN=SNX27 PE=1 SV=2 | 45,094 | 4,07 | 61265 | 3 | 22,50 | 22,85 | 21,55 | 22,15 | 22,23 | 20,86 | **0,681** | 4,01E-01 | **0** | -0,5533333 | 0,39652918 |
| NDKM_HUM | NME4 | sp\|O00746\|NDKM_HUMAN Nucleoside diphosphate kinase, mitochondrial OS=Homo sa | 29,589 | 7,49 | 20659 | 1 | 21,40 | 21,52 | 20,58 | 21,16 | 20,53 | 20,15 | **0,681** | 2,55E-01 | **0** | -0,5533333 | 0,59291031 |
| MYLK_HUM | MYLK | sp\|Q15746\|MYLK_HUMAN Myosin light chain kinase, smooth muscle OS=Homo sapiens | 202,812 | 5,64 | 210715 | 9 | 24,31 | 23,63 | 24,64 | 23,44 | 23,57 | 23,90 | **0,680** | 1,64E-01 | **0** | -0,5566661 | 0,7844932 |
| F13A_HUMA | F13A1 | sp\|P00488\|F13A_HUMAN Coagulation factor XIII A chain OS=Homo sapiens GN=F13A1 F | 26,249 | 1,37 | 83267 | 1 | 22,65 | 21,46 | 22,72 | 22,33 | 20,76 | 22,07 | **0,680** | 4,30E-01 | **0** | -0,5566661 | 0,36637924 |
| PARVB_HUM | PARVB | sp\|Q9HBI1\|PARVB_HUMAN Beta-parvin OS=Homo sapiens GN=PARVB PE=1 SV=1 | 202,893 | 13,19 | 41714 | 4 | 22,83 | 21,64 | 21,60 | 22,25 | 21,38 | 20,77 | **0,680** | 3,98E-01 | **0** | -0,5566667 | 0,39978315 |
| RHOJ_HUMA | RHOJ | sp\|Q9H4E5\|RHOJ_HUMAN Rho-related GTP-binding protein RhoJ OS=Homo sapiens GN | 214,854 | 24,3 | 23821 | 4 | 24,92 | 24,59 | 24,57 | 24,49 | 24,44 | 23,48 | **0,680** | 1,85E-01 | **0** | -0,5566667 | 0,73371385 |
| RFA1_HUMA | RPA1 | sp\|P27694\|RFA1_HUMAN Replication protein A 70 kDa DNA-binding subunit OS=Homo | 87,498 | 6,33 | 68138 | 3 | 22,67 | 22,79 | 21,17 | 22,54 | 22,67 | 19,75 | **0,680** | 6,35E-01 | **0** | -0,5566667 | 0,19712415 |
| TBC24_HUM | TBC1D24 | sp\|Q9ULP9\|TBC24_HUMAN TBC1 domain family member 24 OS=Homo sapiens GN=TBC | 57,041 | 2,68 | 62919 | 1 | 20,55 | 20,42 | 18,61 | 20,16 | 20,07 | 17,67 | **0,678** | 6,15E-01 | **0** | -0,559768 | 0,2112985 |
| DMD_HUMA | DMD | sp\|P11532\|DMD_HUMAN Dystrophin OS=Homo sapiens GN=DMD PE=1 SV=3 | 498,696 | 4,26 | 426750 | 12 | 26,14 | 25,58 | 24,29 | 25,62 | 25,10 | 23,61 | **0,678** | 5,29E-01 | **0** | -0,5599995 | 0,27624676 |
| AF1L2_HUM | AFAP1L2 | sp\|Q8N4X5\|AF1L2_HUMAN Actin filament-associated protein 1-like 2 OS=Homo sapiens | 53,928 | 1,71 | 91300 | 1 | 18,63 | 18,94 | 19,50 | 18,54 | 17,92 | 18,93 | **0,678** | 2,23E-01 | **0** | -0,5599995 | 0,65100227 |
| NEGR1_HUN | NEGR1 | sp\|Q7Z3B1\|NEGR1_HUMAN Neuronal growth regulator 1 OS=Homo sapiens GN=NEGR1 | 347,354 | 29,66 | 38719 | 9 | 24,80 | 24,93 | 25,59 | 24,28 | 24,48 | 24,88 | **0,678** | 1,37E-01 | **0** | -0,5600001 | 0,86376447 |
| MA1A1_HUN | MAN1A1 | sp\|P33908\|MA1A1_HUMAN Mannosyl-oligosaccharide 1,2-alpha-mannosidase IA OS=H | 77,913 | 4,44 | 72969 | 2 | 22,06 | 20,90 | 22,44 | 21,31 | 20,68 | 21,73 | **0,678** | 3,70E-01 | **0** | -0,5600001 | 0,43205962 |
| EP15R_HUM | EPS15L1 | sp\|Q9UBC2\|EP15R_HUMAN Epidermal growth factor receptor substrate 15-like 1 OS=H | 296,997 | 15,62 | 94255 | 10 | 25,18 | 24,89 | 24,76 | 24,71 | 24,48 | 23,95 | **0,677** | 9,35E-02 | **0** | -0,5633335 | 1,02928512 |
| CD166_HUM | ALCAM | sp\|Q13740\|CD166_HUMAN CD166 antigen OS=Homo sapiens GN=ALCAM PE=1 SV=2 | 320,53 | 13,04 | 65102 | 11 | 26,74 | 26,86 | 25,23 | 26,27 | 26,22 | 24,65 | **0,677** | 4,93E-01 | **0** | -0,5633335 | 0,30740347 |
| RFC3_HUMA | RFC3 | sp\|P40938\|RFC3_HUMAN Replication factor C subunit 3 OS=Homo sapiens GN=RFC3 PE | 135,725 | 16,57 | 40556 | 4 | 21,37 | 21,87 | 22,90 | 20,10 | 21,15 | 23,20 | **0,677** | 6,09E-01 | **0** | -0,5633335 | 0,21561591 |
| PARVA_HUM | PARVA | sp\|Q9NVD7\|PARVA_HUMAN Alpha-parvin OS=Homo sapiens GN=PARVA PE=1 SV=1 | 302,546 | 19,89 | 42244 | 6 | 23,38 | 23,66 | 24,52 | 22,76 | 23,36 | 23,74 | **0,675** | 2,73E-01 | **0** | -0,5666663 | 0,56404857 |
| OTU7B_HUN | OTUD7B | sp\|Q6GQQ9\|OTU7B_HUMAN OTU domain-containing protein 7B OS=Homo sapiens GN | 130,649 | 9,02 | 92526 | 5 | 24,03 | 23,33 | 22,53 | 23,25 | 23,23 | 21,71 | **0,675** | 4,45E-01 | **0** | -0,5666676 | 0,3517928 |
| TM87A_HUN | TMEM87A | sp\|Q8NBN3\|TM87A_HUMAN Transmembrane protein 87A OS=Homo sapiens GN=TMEI | 30,791 | 3,6 | 63430 | 1 | 18,43 | 20,07 | 19,04 | 18,57 | 18,19 | 19,08 | **0,674** | 3,55E-01 | **0** | -0,568484 | 0,44960993 |
| NAA35_HUN | NAA35 | sp\|Q5VZE5\|NAA35_HUMAN N-alpha-acetyltransferase 35, NatC auxiliary subunit OS=Ho | 11,459 | 2,34 | 83639 | 1 | 18,98 | 18,78 | 20,16 | 18,71 | 19,16 | 18,34 | **0,674** | 3,10E-01 | **0** | -0,5687993 | 0,5079878 |
| CRIP2_HUM | CRIP2 | sp\|P52943\|CRIP2_HUMAN Cysteine-rich protein 2 OS=Homo sapiens GN=CRIP2 PE=1 SV | 91,294 | 35,1 | 22493 | 4 | 25,31 | 25,18 | 21,97 | 25,20 | 24,86 | 20,69 | **0,674** | 7,69E-01 | **0** | -0,5699991 | 0,11394505 |
| PRKX_HUMA | PRKX | sp\|P51817\|PRKX_HUMAN cAMP-dependent protein kinase catalytic subunit PRKX OS=H | 18,016 | 3,07 | 40896 | 1 | 21,16 | 20,84 | 22,71 | 20,81 | 20,67 | 21,52 | **0,674** | 4,20E-01 | **0** | -0,5699997 | 0,37692723 |
| DDX6_HUMA | DDX6 | sp\|P26196\|DDX6_HUMAN Probable ATP-dependent RNA helicase DDX6 OS=Homo sapie | 311,773 | 30,23 | 54417 | 10 | 24,86 | 25,25 | 24,63 | 24,34 | 24,47 | 24,22 | **0,674** | 4,30E-02 | **0** | -0,5700003 | 1,36657343 |
| HSP72_HUM | HSPA2 | sp\|P54652\|HSP72_HUMAN Heat shock-related 70 kDa protein 2 OS=Homo sapiens GN=I | 1868,459 | 63,38 | 70021 | 39 | 29,29 | 29,27 | 28,45 | 28,80 | 28,54 | 27,96 | **0,674** | 2,00E-01 | **0** | -0,570001 | 0,69894458 |
| SMCE1_HUN | SMARCE1 | sp\|Q969G3\|SMCE1_HUMAN SWI/SNF-related matrix-associated actin-dependent regula | 47,14 | 3,16 | 46649 | 1 | 18,19 | 18,60 | 21,09 | 18,30 | 16,67 | 21,18 | **0,672** | 7,38E-01 | **0** | -0,573089 | 0,13185722 |
| PAN3_HUMA | PAN3 | sp\|Q58A45\|PAN3_HUMAN PAN2-PAN3 deadenylation complex subunit PAN3 OS=Homo | 256,478 | 11,16 | 95613 | 8 | 24,50 | 24,04 | 25,13 | 24,09 | 23,42 | 24,44 | **0,672** | 2,58E-01 | **0** | -0,5733331 | 0,58829443 |
| ACTA_HUMA | ACTA2 | sp\|P62736\|ACTA_HUMAN Actin, aortic smooth muscle OS=Homo sapiens GN=ACTA2 PE | 2941,583 | 88,33 | 42009 | 57 | 19,90 | 18,62 | 18,52 | 18,04 | 18,29 | 18,98 | **0,671** | 3,35E-01 | **0** | -0,575964 | 0,47479503 |
| RIR2_HUMA | RRM2 | sp\|P31350\|RIR2_HUMAN Ribonucleoside-diphosphate reductase subunit M2 OS=Homo | 61,278 | 2,83 | 44878 | 1 | 18,91 | 21,08 | 20,74 | 18,45 | 20,78 | 19,76 | **0,670** | 5,77E-01 | **0** | -0,5785942 | 0,23896749 |
| CAH6_HUMA | CA6 | sp\|P23280\|CAH6_HUMAN Carbonic anhydrase 6 OS=Homo sapiens GN=CA6 PE=1 SV=3 | 19,567 | 2,6 | 35367 | 1 | 19,89 | 20,69 | 19,46 | 22,38 | 17,91 | 18,01 | **0,670** | 7,22E-01 | **0** | -0,5787366 | 0,1413069 |
| FRG1_HUMA | FRG1 | sp\|Q14331\|FRG1_HUMAN Protein FRG1 OS=Homo sapiens GN=FRG1 PE=1 SV=1 | 13,756 | 6,2 | 29172 | 1 | 19,22 | 18,91 | 20,28 | 18,49 | 17,37 | 20,81 | **0,669** | 6,24E-01 | **0** | -0,5794385 | 0,2046691 |
| GCP3_HUMA | TUBGCP3 | sp\|Q96CW5\|GCP3_HUMAN Gamma-tubulin complex component 3 OS=Homo sapiens G | 109,879 | 3,2 | 103571 | 2 | 18,87 | 18,84 | 21,24 | 17,90 | 18,23 | 21,07 | **0,669** | 6,74E-01 | **0** | -0,5809161 | 0,17138442 |
| DNJA4_HUM | DNAJA4 | sp\|Q8WW22\|DNJA4_HUMAN DnaJ homolog subfamily A member 4 OS=Homo sapiens G | 25,396 | 2,27 | 44798 | 1 | 24,29 | 23,79 | 19,05 | 24,07 | 22,63 | 18,68 | **0,668** | 8,14E-01 | **0** | -0,5824083 | 0,0892941 |
| LAMP2_HUN | LAMP2 | sp\|P13473\|LAMP2_HUMAN Lysosome-associated membrane glycoprotein 2 OS=Homo | 35,214 | 2,2 | 44961 | 1 | 22,85 | 22,06 | 22,63 | 21,27 | 22,08 | 22,44 | **0,667** | 2,36E-01 | **0** | -0,5833327 | 0,62751134 |
| VPS18_HUM | VPS18 | sp\|Q9P253\|VPS18_HUMAN Vacuolar protein sorting-associated protein 18 homolog OS | 88,659 | 4,11 | 110186 | 3 | 22,59 | 22,22 | 21,99 | 21,94 | 21,69 | 21,42 | **0,667** | 6,45E-02 | **0** | -0,5833327 | 1,19012345 |
| NALD2_HUN | NAALAD2 | sp\|Q9Y3Q0\|NALD2_HUMAN N-acetylated-alpha-linked acidic dipeptidase 2 OS=Homo s | 358,773 | 17,84 | 83592 | 10 | 26,51 | 26,87 | 27,45 | 26,14 | 26,17 | 26,77 | **0,667** | 1,63E-01 | **0** | -0,583334 | 0,78667229 |
| PCDGK_HUN | PCDHGC3 | sp\|Q9UN70\|PCDGK_HUMAN Protocadherin gamma-C3 OS=Homo sapiens GN=PCDHGC | 272,537 | 11,03 | 101077 | 8 | 26,71 | 26,46 | 25,07 | 26,37 | 25,66 | 24,45 | **0,666** | 4,82E-01 | **0** | -0,5866655 | 0,3168763 |
| CSRP2_HUM | CSRP2 | sp\|Q16527\|CSRP2_HUMAN Cysteine and glycine-rich protein 2 OS=Homo sapiens GN=C | 216,769 | 47,15 | 20954 | 6 | 24,45 | 24,22 | 23,37 | 24,27 | 23,78 | 22,23 | **0,666** | 4,47E-01 | **0** | -0,5866667 | 0,34933471 |
| RHDF1_HUM | RHBDF1 | sp\|Q96CC6\|RHDF1_HUMAN Inactive rhomboid protein 1 OS=Homo sapiens GN=RHBDF | 63,916 | 3,51 | 97401 | 2 | 20,78 | 21,72 | 17,90 | 20,30 | 19,70 | 18,63 | **0,665** | 6,62E-01 | **0** | -0,5886841 | 0,17930037 |
| UBP10_HUM | USP10 | sp\|Q14694\|UBP10_HUMAN Ubiquitin carboxyl-terminal hydrolase 10 OS=Homo sapiens | 26,07 | 1,25 | 87134 | 1 | 21,24 | 17,52 | 21,92 | 18,81 | 18,00 | 22,11 | **0,665** | 7,67E-01 | **0** | -0,589325 | 0,11528085 |
| CRYAB_HUM | CRYAB | sp\|P02511\|CRYAB_HUMAN Alpha-crystallin B chain OS=Homo sapiens GN=CRYAB PE=1 | 24,447 | 11,43 | 20159 | 2 | 20,57 | 21,79 | 22,65 | 19,51 | 21,19 | 22,54 | **0,664** | 6,09E-01 | **0** | -0,5899995 | 0,21553091 |
| FZD2_HUMA | FZD2 | sp\|Q14332\|FZD2_HUMAN Frizzled-2 OS=Homo sapiens GN=FZD2 PE=1 SV=1 | 105,548 | 12,57 | 63554 | 6 | 24,37 | 23,48 | 22,57 | 23,18 | 23,38 | 22,09 | **0,664** | 4,19E-01 | **0** | -0,5900002 | 0,37730071 |
| SNX18_HUM | SNX18 | sp\|Q96RF0\|SNX18_HUMAN Sorting nexin-18 OS=Homo sapiens GN=SNX18 PE=1 SV=2 | 557,149 | 27,07 | 68894 | 13 | 27,35 | 27,41 | 26,36 | 26,97 | 26,78 | 25,60 | **0,664** | 3,42E-01 | **0** | -0,5900002 | 0,46636242 |
| LARP4_HUM | LARP4 | sp\|Q71RC2\|LARP4_HUMAN La-related protein 4 OS=Homo sapiens GN=LARP4 PE=1 SV= | 59,047 | 1,8 | 80596 | 1 | 20,45 | 20,12 | 22,27 | 20,73 | 18,25 | 22,08 | **0,663** | 6,73E-01 | **0** | -0,5927118 | 0,17180023 |
| NPC1_HUMA | NPC1 | sp\|O15118\|NPC1_HUMAN Niemann-Pick C1 protein OS=Homo sapiens GN=NPC1 PE=1 S | 85,557 | 2,82 | 142167 | 2 | 23,55 | 23,35 | 22,23 | 22,77 | 23,01 | 21,57 | **0,663** | 3,83E-01 | **0** | -0,5933329 | 0,41691891 |
| BACD3_HUM | KCTD10 | sp\|Q9H3F6\|BACD3_HUMAN BTB/POZ domain-containing adapter for CUL3-mediated Rh | 35,903 | 3,51 | 35432 | 1 | 21,95 | 21,91 | 21,93 | 21,60 | 21,12 | 21,28 | **0,661** | 1,35E-02 | **0** | -0,5966663 | 1,86843396 |
| GOGA7_HUN | GOLGA7 | sp\|Q7Z5G4\|GOGA7_HUMAN Golgin subfamily A member 7 OS=Homo sapiens GN=GOL | 175,056 | 32,85 | 15824 | 6 | 24,93 | 25,90 | 23,28 | 24,69 | 24,73 | 22,90 | **0,661** | 5,73E-01 | **0** | -0,596667 | 0,24159828 |
| PGAM5_HUN | PGAM5 | sp\|Q96HS1\|PGAM5_HUMAN Serine/threonine-protein phosphatase PGAM5, mitochon | 52,777 | 4,84 | 32004 | 2 | 21,30 | 20,99 | 21,91 | 20,64 | 20,13 | 21,64 | **0,661** | 3,15E-01 | **0** | -0,596667 | 0,50220589 |
| TOM7_HUM | TOMM7 | sp\|Q9P0U1\|TOM7_HUMAN Mitochondrial import receptor subunit TOM7 homolog OS= | 17,919 | 30,91 | 6248 | 1 | 18,78 | 19,33 | 19,53 | 18,28 | 18,47 | 19,09 | **0,660** | 1,45E-01 | **0** | -0,5987848 | 0,83935935 |
| KPCD_HUMA | PRKCD | sp\|Q05655\|KPCD_HUMAN Protein kinase C delta type OS=Homo sapiens GN=PRKCD PE= | 20,197 | 1,48 | 77505 | 1 | 20,97 | 21,39 | 19,87 | 21,59 | 21,08 | 17,76 | **0,660** | 6,65E-01 | **0** | -0,5989958 | 0,17733458 |
| LYOX_HUMA | LOX | sp\|P28300\|LYOX_HUMAN Protein-lysine 6-oxidase OS=Homo sapiens GN=LOX PE=1 SV= | 278,87 | 22,06 | 46944 | 7 | 25,49 | 24,29 | 26,37 | 24,93 | 23,83 | 25,59 | **0,660** | 4,91E-01 | **0** | -0,6000004 | 0,30912974 |
| RPC2_HUMA | POLR3B | sp\|Q9NW08\|RPC2_HUMAN DNA-directed RNA polymerase III subunit RPC2 OS=Homo s | 9,752 | 0,62 | 127785 | 1 | 23,18 | 22,25 | 25,24 | 22,80 | 22,33 | 23,74 | **0,660** | 5,72E-01 | **0** | -0,6000004 | 0,24267232 |
| AGRL2_HUM | ADGRL2 | sp\|O95490\|AGRL2_HUMAN Adhesion G protein-coupled receptor L2 OS=Homo sapiens | 358,437 | 10,01 | 163349 | 10 | 25,56 | 25,60 | 24,65 | 24,89 | 25,05 | 24,07 | **0,660** | 2,39E-01 | **0** | -0,6000004 | 0,62158768 |
| DESI1_HUM | DESI1 | sp\|Q6ICB0\|DESI1_HUMAN Desumoylating isopeptidase 1 OS=Homo sapiens GN=DESI1 F | 26,534 | 9,52 | 18263 | 2 | 23,04 | 23,73 | 21,88 | 22,50 | 22,50 | 21,84 | **0,658** | 3,59E-01 | **0** | -0,6033332 | 0,44478915 |
| CYH1_HUMA | CYTH1 | sp\|Q15438\|CYH1_HUMAN Cytohesin-1 OS=Homo sapiens GN=CYTH1 PE=1 SV=1 | 23,337 | 2,76 | 46413 | 1 | 21,87 | 21,72 | 20,93 | 21,50 | 21,35 | 19,86 | **0,658** | 3,71E-01 | **0** | -0,6033332 | 0,43069789 |
| NACAD_HUN | NACAD | sp\|O15069\|NACAD_HUMAN NAC-alpha domain-containing protein 1 OS=Homo sapiens | 30,335 | 0,83 | 161101 | 1 | 21,47 | 21,75 | 20,44 | 21,38 | 21,68 | 18,79 | **0,658** | 5,79E-01 | **0** | -0,6034126 | 0,23742418 |
| MYO9B_HUN | MYO9B | sp\|Q13459\|MYO9B_HUMAN Unconventional myosin-IXb OS=Homo sapiens GN=MYO9 | 109,641 | 1,9 | 243401 | 3 | 20,62 | 19,49 | 20,79 | 17,65 | 20,15 | 21,28 | **0,657** | 6,25E-01 | **0** | -0,6066672 | 0,20413059 |
| DAB2P_HUM | DAB2IP | sp\|Q5VWQ8\|DAB2P_HUMAN Disabled homolog 2-interacting protein OS=Homo sapiens | 64,24 | 1,26 | 131625 | 1 | 20,15 | 20,17 | 19,54 | 19,99 | 19,85 | 18,19 | **0,656** | 3,78E-01 | **0** | -0,6088117 | 0,42305671 |
| TARA_HUMA | TRIOBP | sp\|Q9H2D6\|TARA_HUMAN TRIO and F-actin-binding protein OS=Homo sapiens GN=TRI | 13,082 | 0,51 | 261376 | 1 | 16,12 | 19,50 | 17,75 | 17,52 | 16,86 | 17,16 | **0,656** | 5,73E-01 | **0** | -0,6089141 | 0,24179597 |
| P5CR1_HUM | PYCR1 | sp\|P32322\|P5CR1_HUMAN Pyrroline-5-carboxylate reductase 1, mitochondrial OS=Hom | 28,992 | 3,45 | 33361 | 1 | 21,65 | 21,39 | 22,72 | 21,15 | 21,00 | 21,78 | **0,655** | 2,66E-01 | **0** | -0,6099993 | 0,57546688 |
| PAN2_HUMA | PAN2 | sp\|Q504Q3\|PAN2_HUMAN PAN2-PAN3 deadenylation complex catalytic subunit PAN2 C | 313,325 | 11,15 | 135368 | 10 | 24,42 | 24,13 | 24,70 | 23,69 | 23,52 | 24,21 | **0,655** | 8,27E-02 | **0** | -0,61 | 1,0826865 |
| GRPE1_HUM | GRPEL1 | sp\|Q9HAV7\|GRPE1_HUMAN GrpE protein homolog 1, mitochondrial OS=Homo sapiens | 22,758 | 5,07 | 24279 | 1 | 20,96 | 21,34 | 22,65 | 20,57 | 20,97 | 21,58 | **0,655** | 3,60E-01 | **0** | -0,61 | 0,44406458 |
| ARL8B_HUM | ARL8B | sp\|Q9NVJ2\|ARL8B_HUMAN ADP-ribosylation factor-like protein 8B OS=Homo sapiens G | 423,201 | 67,74 | 21539 | 13 | 26,54 | 26,79 | 25,62 | 25,81 | 26,19 | 25,12 | **0,655** | 2,67E-01 | **0** | -0,6100006 | 0,57267969 |
| SEP15_HUM | SELENOF | sp\|O60613\|SEP15_HUMAN Selenoprotein F OS=Homo sapiens GN=SELENOF PE=1 SV=3 | 57,397 | 7,41 | 17790 | 1 | 18,99 | 18,70 | 19,48 | 17,33 | 19,16 | 18,84 | **0,655** | 3,72E-01 | **0** | -0,6110153 | 0,42950112 |
| RABP1_HUM | CRABP1 | sp\|P29762\|RABP1_HUMAN Cellular retinoic acid-binding protein 1 OS=Homo sapiens GN | 262,561 | 62,04 | 15566 | 9 | 24,64 | 24,89 | 24,64 | 23,79 | 24,33 | 24,21 | **0,654** | 2,89E-02 | **0** | -0,6133327 | 1,53960027 |
| NRDC_HUM | NRDC | sp\|O43847\|NRDC_HUMAN Nardilysin OS=Homo sapiens GN=NRDC PE=1 SV=2 | 34,595 | 2,52 | 131572 | 3 | 21,28 | 20,54 | 22,64 | 20,27 | 20,33 | 22,02 | **0,654** | 5,06E-01 | **0** | -0,6133334 | 0,29567234 |
| FERM2_HUN | FERMT2 | sp\|Q96AC1\|FERM2_HUMAN Fermitin family homolog 2 OS=Homo sapiens GN=FERMT2 | 1299,133 | 57,5 | 77861 | 35 | 28,26 | 27,98 | 27,83 | 27,47 | 27,59 | 27,17 | **0,654** | 2,59E-02 | **0** | -0,6133334 | 1,58687743 |
| TP4A1_HUM | PTP4A1 | sp\|Q93096\|TP4A1_HUMAN Protein tyrosine phosphatase type IVA 1 OS=Homo sapiens | 241,91 | 41,04 | 19815 | 6 | 24,38 | 24,60 | 23,29 | 23,95 | 23,79 | 22,68 | **0,652** | 3,39E-01 | **0** | -0,6166662 | 0,46946875 |
| WDR48_HUN | WDR48 | sp\|Q8TAF3\|WDR48_HUMAN WD repeat-containing protein 48 OS=Homo sapiens GN=W | 82,471 | 6,94 | 76210 | 3 | 21,40 | 20,71 | 19,82 | 20,23 | 20,86 | 18,99 | **0,652** | 4,37E-01 | **0** | -0,6166662 | 0,35958799 |
| SNX9_HUMA | SNX9 | sp\|Q9Y5X1\|SNX9_HUMAN Sorting nexin-9 OS=Homo sapiens GN=SNX9 PE=1 SV=1 | 499,271 | 25,71 | 66592 | 14 | 27,38 | 27,18 | 26,16 | 26,90 | 26,81 | 25,16 | **0,652** | 4,16E-01 | **0** | -0,6166668 | 0,38104549 |
| BDH_HUMAI | BDH1 | sp\|Q02338\|BDH_HUMAN D-beta-hydroxybutyrate dehydrogenase, mitochondrial OS=H | 70,329 | 8,75 | 38157 | 3 | 23,58 | 23,46 | 24,20 | 23,23 | 23,28 | 22,88 | **0,652** | 7,79E-02 | **0** | -0,6166668 | 1,10869073 |
| F1712_HUM | FAM171A2 | sp\|A8MVW0\|F1712_HUMAN Protein FAM171A2 OS=Homo sapiens GN=FAM171A2 PE= | 178,256 | 8,23 | 87435 | 6 | 23,97 | 23,67 | 20,49 | 23,20 | 22,62 | 20,44 | **0,649** | 6,78E-01 | **0** | -0,6233323 | 0,16871766 |
| MOB3A_HUI | MOB3A | sp\|Q96BX8\|MOB3A_HUMAN MOB kinase activator 3A OS=Homo sapiens GN=MOB3A P | 17,881 | 5,53 | 25464 | 1 | 21,51 | 20,91 | 20,66 | 21,22 | 20,22 | 19,77 | **0,649** | 2,78E-01 | **0** | -0,6233336 | 0,55557875 |
| FBW1B_HUN | FBXW11 | sp\|Q9UKB1\|FBW1B_HUMAN F-box/WD repeat-containing protein 11 OS=Homo sapiens | 46,927 | 6,27 | 62091 | 2 | 19,74 | 21,38 | 18,93 | 20,57 | 19,47 | 18,14 | **0,649** | 5,69E-01 | **0** | -0,6235148 | 0,24483179 |
| CD123_HUM | CDC123 | sp\|O75794\|CD123_HUMAN Cell division cycle protein 123 homolog OS=Homo sapiens G | 41,677 | 2,98 | 39135 | 1 | 19,54 | 20,76 | 21,15 | 17,33 | 20,70 | 21,55 | **0,649** | 6,74E-01 | **0** | -0,6235269 | 0,17124315 |
| EDIL3_HUM | EDIL3 | sp\|O43854\|EDIL3_HUMAN EGF-like repeat and discoidin I-like domain-containing protei | 10,212 | 2,08 | 53765 | 1 | 17,76 | 19,18 | 21,49 | 17,46 | 17,42 | 21,68 | **0,649** | 7,44E-01 | **0** | -0,6241767 | 0,12845361 |
| KI20A_HUM | KIF20A | sp\|O95235\|KI20A_HUMAN Kinesin-like protein KIF20A OS=Homo sapiens GN=KIF20A PE | 185,6 | 7,64 | 100278 | 5 | 20,68 | 21,15 | 24,78 | 20,59 | 20,04 | 24,10 | **0,648** | 7,47E-01 | **0** | -0,6266664 | 0,1264936 |
| T132A_HUM | TMEM132A | sp\|Q24JP5\|T132A_HUMAN Transmembrane protein 132A OS=Homo sapiens GN=TMEM | 14,338 | 0,98 | 110110 | 1 | 17,77 | 16,32 | 18,94 | 17,03 | 16,94 | 17,17 | **0,647** | 4,54E-01 | **0** | -0,6287645 | 0,34257658 |
| CARL1_HUM | CARMIL1 | sp\|Q5VZK9\|CARL1_HUMAN F-actin-uncapping protein LRRC16A OS=Homo sapiens GN= | 169,108 | 4,67 | 151557 | 4 | 22,52 | 22,67 | 22,39 | 21,66 | 22,37 | 21,66 | **0,646** | 6,54E-02 | **0** | -0,6299998 | 1,18418935 |
| XPO5_HUMA | XPO5 | sp\|Q9HAV4\|XPO5_HUMAN Exportin-5 OS=Homo sapiens GN=XPO5 PE=1 SV=1 | 105,511 | 3,65 | 136311 | 3 | 20,89 | 20,51 | 21,80 | 19,04 | 20,83 | 21,43 | **0,645** | 4,80E-01 | **0** | -0,6333326 | 0,31898135 |
| RAP2C_HUM | RAP2C | sp\|Q9Y3L5\|RAP2C_HUMAN Ras-related protein Rap-2c OS=Homo sapiens GN=RAP2C PE | 602,567 | 63,93 | 20745 | 11 | 25,96 | 25,79 | 24,25 | 25,50 | 25,45 | 23,15 | **0,645** | 5,40E-01 | **0** | -0,6333332 | 0,26743064 |
| TM1L1_HUN | TOM1L1 | sp\|O75674\|TM1L1_HUMAN TOM1-like protein 1 OS=Homo sapiens GN=TOM1L1 PE=1 S | 144,888 | 7,35 | 52989 | 4 | 25,36 | 25,37 | 24,71 | 24,89 | 24,61 | 24,04 | **0,645** | 1,29E-01 | **0** | -0,6333332 | 0,8890879 |
| VASP_HUMA | VASP | sp\|P50552\|VASP_HUMAN Vasodilator-stimulated phosphoprotein OS=Homo sapiens GN | 170,153 | 13,95 | 39830 | 5 | 24,62 | 24,93 | 22,99 | 24,79 | 24,55 | 21,30 | **0,645** | 6,46E-01 | **0** | -0,6333338 | 0,18994083 |
| CPNE8_HUM | CPNE8 | sp\|Q86YQ8\|CPNE8_HUMAN Copine-8 OS=Homo sapiens GN=CPNE8 PE=1 SV=2 | 773,32 | 46,81 | 63108 | 21 | 26,61 | 26,06 | 25,01 | 26,00 | 25,74 | 24,03 | **0,643** | 4,58E-01 | **0** | -0,6366666 | 0,33912613 |
| LETM1_HUN | LETM1 | sp\|O95202\|LETM1_HUMAN Mitochondrial proton/calcium exchanger protein OS=Homo | 68,964 | 6,63 | 83354 | 4 | 22,30 | 22,01 | 23,62 | 21,82 | 21,96 | 22,23 | **0,642** | 2,78E-01 | **0** | -0,6400007 | 0,55639636 |
| APH1A_HUM | APH1A | sp\|Q96BI3\|APH1A_HUMAN Gamma-secretase subunit APH-1A OS=Homo sapiens GN=A | 32,197 | 4,91 | 28996 | 1 | 21,12 | 20,02 | 19,16 | 18,92 | 20,23 | 19,23 | **0,641** | 4,06E-01 | **0** | -0,6413708 | 0,39138687 |
| CIP4_HUMA | TRIP10 | sp\|Q15642\|CIP4_HUMAN Cdc42-interacting protein 4 OS=Homo sapiens GN=TRIP10 PE= | 268,498 | 14,14 | 68352 | 6 | 23,14 | 23,89 | 23,27 | 22,55 | 23,25 | 22,57 | **0,640** | 1,20E-01 | **0** | -0,6433334 | 0,92097395 |
| CY1_HUMAN | CYC1 | sp\|P08574\|CY1_HUMAN Cytochrome c1, heme protein, mitochondrial OS=Homo sapien | 117,702 | 16,62 | 35422 | 4 | 22,16 | 23,25 | 24,12 | 22,15 | 22,31 | 23,14 | **0,640** | 3,75E-01 | **0** | -0,6433341 | 0,42619637 |
| STX11_HUM | STX11 | sp\|O75558\|STX11_HUMAN Syntaxin-11 OS=Homo sapiens GN=STX11 PE=1 SV=1 | 15,96 | 3,14 | 33196 | 1 | 23,88 | 23,65 | 21,99 | 23,55 | 22,79 | 21,24 | **0,639** | 5,14E-01 | **0** | -0,6466662 | 0,28924191 |
| RING1_HUM | RING1 | sp\|Q06587\|RING1_HUMAN E3 ubiquitin-protein ligase RING1 OS=Homo sapiens GN=RI | 26,253 | 2,96 | 42429 | 1 | 17,60 | 19,26 | 19,77 | 17,43 | 17,79 | 19,46 | **0,637** | 5,12E-01 | **0** | -0,6499685 | 0,29044529 |
| QCR2_HUMA | UQCRC2 | sp\|P22695\|QCR2_HUMAN Cytochrome b-c1 complex subunit 2, mitochondrial OS=Hom | 75,053 | 7,73 | 48443 | 2 | 21,52 | 21,54 | 23,54 | 21,57 | 20,73 | 22,35 | **0,637** | 4,71E-01 | **0** | -0,6500009 | 0,32708334 |
| FARP1_HUM | FARP1 | sp\|Q9Y4F1\|FARP1_HUMAN FERM, ARHGEF and pleckstrin domain-containing protein 1 | 1356,87 | 36,46 | 118633 | 34 | 28,28 | 28,27 | 28,62 | 27,89 | 27,62 | 27,71 | **0,637** | 9,66E-03 | **0** | -0,6500009 | 2,01521188 |
| RDH14_HUM | RDH14 | sp\|Q9HBH5\|RDH14_HUMAN Retinol dehydrogenase 14 OS=Homo sapiens GN=RDH14 P | 29,244 | 2,38 | 36865 | 1 | 23,00 | 22,94 | 20,29 | 22,94 | 23,17 | 18,17 | **0,637** | 7,44E-01 | **0** | -0,6515865 | 0,12856487 |
| K2C73_HUM | KRT73 | sp\|Q86Y46\|K2C73_HUMAN Keratin, type II cytoskeletal 73 OS=Homo sapiens GN=KRT7 | 234,339 | 11,3 | 58923 | 8 | 21,74 | 19,08 | 23,16 | 21,92 | 17,65 | 22,46 | **0,636** | 7,53E-01 | **0** | -0,6519356 | 0,12312393 |
| INT3_HUMA | INTS3 | sp\|Q68E01\|INT3_HUMAN Integrator complex subunit 3 OS=Homo sapiens GN=INTS3 PE | 26,45 | 2,49 | 118070 | 1 | 20,36 | 19,06 | 20,46 | 18,67 | 18,40 | 20,85 | **0,636** | 5,07E-01 | **0** | -0,6525777 | 0,29514451 |
| COBL1_HUM | COBLL1 | sp\|Q53SF7\|COBL1_HUMAN Cordon-bleu protein-like 1 OS=Homo sapiens GN=COBLL1 F | 33,534 | 1,5 | 131787 | 2 | 21,83 | 21,49 | 21,67 | 21,57 | 20,80 | 21,08 | **0,636** | 1,08E-02 | **0** | -0,6533337 | 1,9657037 |
| ULK3_HUMA | ULK3 | sp\|Q6PHR2\|ULK3_HUMAN Serine/threonine-protein kinase ULK3 OS=Homo sapiens GN | 70,07 | 4,87 | 53444 | 2 | 23,72 | 23,53 | 22,03 | 23,35 | 22,97 | 21,00 | **0,636** | 5,10E-01 | **0** | -0,6533337 | 0,2927857 |
| CCAR2_HUM | CCAR2 | sp\|Q8N163\|CCAR2_HUMAN Cell cycle and apoptosis regulator protein 2 OS=Homo sapi | 27,547 | 1,95 | 102902 | 2 | 19,77 | 20,21 | 22,76 | 18,22 | 20,06 | 22,49 | **0,635** | 6,94E-01 | **0** | -0,6550763 | 0,15862903 |
| CETN2_HUM | CETN2 | sp\|P41208\|CETN2_HUMAN Centrin-2 OS=Homo sapiens GN=CETN2 PE=1 SV=1 | 31,876 | 6,98 | 19738 | 1 | 21,76 | 21,54 | 21,42 | 21,63 | 21,19 | 19,93 | **0,635** | 2,74E-01 | **0** | -0,6555837 | 0,56165526 |
| PM2PB_HUN | PMS2P11 | sp\|Q13670\|PM2PB_HUMAN Putative postmeiotic segregation increased 2-like protein 1 | 28,597 | 5,19 | 28555 | 1 | 20,44 | 20,36 | 19,64 | 20,39 | 19,66 | 18,42 | **0,635** | 3,57E-01 | **0** | -0,6557312 | 0,44777237 |
| CCNB2_HUM | CCNB2 | sp\|O95067\|CCNB2_HUMAN G2/mitotic-specific cyclin-B2 OS=Homo sapiens GN=CCNB2 | 16,447 | 2,76 | 45282 | 1 | 19,04 | 19,10 | 20,52 | 18,94 | 16,58 | 21,17 | **0,634** | 6,66E-01 | **0** | -0,6566671 | 0,17666046 |
| FBP1L_HUM | FNBP1L | sp\|Q5T0N5\|FBP1L_HUMAN Formin-binding protein 1-like OS=Homo sapiens GN=FNBP1 | 128,691 | 11,4 | 69977 | 6 | 24,52 | 24,37 | 23,14 | 24,07 | 23,69 | 22,30 | **0,634** | 3,97E-01 | **0** | -0,6566671 | 0,40107726 |
| MIC60_HUM | IMMT | sp\|Q16891\|MIC60_HUMAN MICOS complex subunit MIC60 OS=Homo sapiens GN=IMM | 341,018 | 18,21 | 83678 | 13 | 25,26 | 25,47 | 25,97 | 25,06 | 24,87 | 24,79 | **0,633** | 4,28E-02 | **0** | -0,6599992 | 1,36822526 |
| IQGA3_HUM | IQGAP3 | sp\|Q86VI3\|IQGA3_HUMAN Ras GTPase-activating-like protein IQGAP3 OS=Homo sapien | 488,125 | 14,96 | 184699 | 17 | 25,43 | 25,26 | 25,03 | 24,83 | 24,59 | 24,32 | **0,633** | 2,44E-02 | **0** | -0,6600005 | 1,61217121 |
| K2C8_HUMA | KRT8 | sp\|P05787\|K2C8_HUMAN Keratin, type II cytoskeletal 8 OS=Homo sapiens GN=KRT8 PE= | 532,469 | 36,02 | 53704 | 20 | 25,72 | 25,35 | 22,85 | 24,77 | 24,79 | 22,37 | **0,631** | 6,12E-01 | **0** | -0,6633326 | 0,21327136 |
| RET1_HUMA | RBP1 | sp\|P09455\|RET1_HUMAN Retinol-binding protein 1 OS=Homo sapiens GN=RBP1 PE=1 S | 178,297 | 41,48 | 15850 | 7 | 25,15 | 25,21 | 24,22 | 24,37 | 24,64 | 23,58 | **0,631** | 2,16E-01 | **0** | -0,6633326 | 0,66612999 |
| SNX33_HUM | SNX33 | sp\|Q8WV41\|SNX33_HUMAN Sorting nexin-33 OS=Homo sapiens GN=SNX33 PE=1 SV=1 | 459,292 | 20,21 | 65265 | 13 | 27,39 | 26,99 | 26,04 | 26,60 | 26,38 | 25,44 | **0,630** | 2,81E-01 | **0** | -0,6666667 | 0,55107139 |
| UBAP1_HUM | UBAP1 | sp\|Q9NZ09\|UBAP1_HUMAN Ubiquitin-associated protein 1 OS=Homo sapiens GN=UBA | 28,675 | 4,18 | 55084 | 2 | 22,52 | 22,40 | 20,02 | 22,38 | 21,64 | 18,92 | **0,630** | 6,43E-01 | **0** | -0,6666673 | 0,19209062 |
| NTNG1_HUN | NTNG1 | sp\|Q9Y2I2\|NTNG1_HUMAN Netrin-G1 OS=Homo sapiens GN=NTNG1 PE=1 SV=3 | 49,466 | 3,34 | 60541 | 1 | 20,58 | 18,85 | 21,58 | 20,93 | 16,99 | 21,08 | **0,629** | 6,90E-01 | **0** | -0,6692816 | 0,16139772 |
| TSN6_HUMA | TSPAN6 | sp\|O43657\|TSN6_HUMAN Tetraspanin-6 OS=Homo sapiens GN=TSPAN6 PE=1 SV=1 | 147,891 | 13,06 | 27563 | 3 | 27,27 | 26,37 | 25,79 | 26,27 | 26,39 | 24,76 | **0,629** | 3,79E-01 | **0** | -0,6700007 | 0,42095297 |
| PP4P1_HUM | PIP4P1 | sp\|Q86T03\|PP4P1_HUMAN Type 1 phosphatidylinositol 4,5-bisphosphate 4-phosphatas | 55,731 | 7,58 | 29470 | 2 | 22,16 | 22,14 | 20,06 | 21,73 | 21,62 | 18,99 | **0,627** | 5,85E-01 | **0** | -0,6724173 | 0,23278314 |
| NPS3A_HUM | NIPSNAP3A | sp\|Q9UFN0\|NPS3A_HUMAN Protein NipSnap homolog 3A OS=Homo sapiens GN=NIPSN | 11,618 | 5,67 | 28467 | 1 | 21,02 | 20,74 | 22,44 | 20,06 | 20,92 | 21,20 | **0,627** | 3,44E-01 | **0** | -0,6733335 | 0,46328812 |
| AGRG6_HUN | ADGRG6 | sp\|Q86SQ4\|AGRG6_HUMAN Adhesion G-protein coupled receptor G6 OS=Homo sapien | 126,175 | 2,29 | 136695 | 2 | 23,06 | 23,15 | 22,11 | 22,55 | 22,77 | 20,98 | **0,627** | 3,62E-01 | **0** | -0,6733335 | 0,44165826 |
| REEP5_HUM | REEP5 | sp\|Q00765\|REEP5_HUMAN Receptor expression-enhancing protein 5 OS=Homo sapiens | 16,317 | 4,23 | 21493 | 1 | 19,27 | 23,67 | 18,34 | 17,00 | 19,25 | 23,00 | **0,626** | 7,92E-01 | **0** | -0,6761665 | 0,10119212 |
| RD23A_HUM | RAD23A | sp\|P54725\|RD23A_HUMAN UV excision repair protein RAD23 homolog A OS=Homo sapi | 214,486 | 23,69 | 39609 | 5 | 23,59 | 23,90 | 22,14 | 22,90 | 23,35 | 21,35 | **0,626** | 4,52E-01 | **0** | -0,6766663 | 0,34475912 |
| E41L1_HUM | EPB41L1 | sp\|Q9H4G0\|E41L1_HUMAN Band 4.1-like protein 1 OS=Homo sapiens GN=EPB41L1 PE= | 120,055 | 7,95 | 98503 | 6 | 23,07 | 23,42 | 20,19 | 23,10 | 22,82 | 18,73 | **0,626** | 7,18E-01 | **0** | -0,6766669 | 0,14399927 |
| CH60_HUMA | HSPD1 | sp\|P10809\|CH60_HUMAN 60 kDa heat shock protein, mitochondrial OS=Homo sapiens | 1724,916 | 55,85 | 61055 | 59 | 27,84 | 27,82 | 29,35 | 27,33 | 27,30 | 28,35 | **0,626** | 3,32E-01 | **0** | -0,6766669 | 0,47934099 |
| RFIP5_HUM | RAB11FIP5 | sp\|Q9BXF6\|RFIP5_HUMAN Rab11 family-interacting protein 5 OS=Homo sapiens GN=RA | 12,115 | 1,84 | 70415 | 1 | 19,70 | 20,47 | 20,60 | 19,21 | 20,61 | 18,92 | **0,625** | 3,18E-01 | **0** | -0,6774216 | 0,49821491 |
| RSPRY_HUM | RSPRY1 | sp\|Q96DX4\|RSPRY_HUMAN RING finger and SPRY domain-containing protein 1 OS=Hom | 235,106 | 17,36 | 64180 | 6 | 24,54 | 24,27 | 22,56 | 23,86 | 23,87 | 21,60 | **0,624** | 5,25E-01 | **0** | -0,6799997 | 0,2800836 |
| ADT2_HUMA | SLC25A5 | sp\|P05141\|ADT2_HUMAN ADP/ATP translocase 2 OS=Homo sapiens GN=SLC25A5 PE=1 | 729,811 | 52,35 | 32852 | 19 | 26,08 | 26,07 | 26,99 | 25,40 | 25,69 | 26,01 | **0,624** | 1,26E-01 | **0** | -0,6799997 | 0,90054092 |
| ERBB2_HUM | ERBB2 | sp\|P04626\|ERBB2_HUMAN Receptor tyrosine-protein kinase erbB-2 OS=Homo sapiens | 113,604 | 2,79 | 137910 | 3 | 22,44 | 21,95 | 21,94 | 21,97 | 21,75 | 20,57 | **0,624** | 2,17E-01 | **0** | -0,6800009 | 0,66272195 |
| NIPS2_HUM | NIPSNAP2 | sp\|O75323\|NIPS2_HUMAN Protein NipSnap homolog 2 OS=Homo sapiens GN=NIPSNAP | 38,999 | 10,49 | 33743 | 2 | 18,42 | 16,45 | 20,68 | 18,00 | 16,04 | 19,47 | **0,624** | 6,88E-01 | **0** | -0,6809451 | 0,16256022 |
| ACBD5_HUM | ACBD5 | sp\|Q5T8D3\|ACBD5_HUMAN Acyl-CoA-binding domain-containing protein 5 OS=Homo s | 20,253 | 2,06 | 60092 | 1 | 18,60 | 19,51 | 20,37 | 17,01 | 19,29 | 20,14 | **0,624** | 5,58E-01 | **0** | -0,6809947 | 0,25339793 |
| PRC1_HUMA | PRC1 | sp\|O43663\|PRC1_HUMAN Protein regulator of cytokinesis 1 OS=Homo sapiens GN=PRC | 299,758 | 13,71 | 71607 | 10 | 23,22 | 23,90 | 28,26 | 23,25 | 23,15 | 26,93 | **0,623** | 7,51E-01 | **0** | -0,6833331 | 0,12437643 |
| MICU1_HUM | MICU1 | sp\|Q9BPX6\|MICU1_HUMAN Calcium uptake protein 1, mitochondrial OS=Homo sapiens | 47,787 | 3,15 | 54351 | 1 | 20,64 | 20,90 | 20,62 | 19,94 | 19,79 | 20,38 | **0,623** | 2,63E-02 | **0** | -0,6833331 | 1,5798293 |
| ENAH_HUM | ENAH | sp\|Q8N8S7\|ENAH_HUMAN Protein enabled homolog OS=Homo sapiens GN=ENAH PE=1 | 460,286 | 27,75 | 66510 | 14 | 25,73 | 25,77 | 24,52 | 25,20 | 25,23 | 23,54 | **0,623** | 3,80E-01 | **0** | -0,6833331 | 0,42039929 |
| COCA1_HUN | COL12A1 | sp\|Q99715\|COCA1_HUMAN Collagen alpha-1(XII) chain OS=Homo sapiens GN=COL12A1 | 854,139 | 10,71 | 333147 | 30 | 26,48 | 25,10 | 27,29 | 26,13 | 24,70 | 25,99 | **0,623** | 4,33E-01 | **0** | -0,6833337 | 0,36348588 |
| MIC19_HUM | CHCHD3 | sp\|Q9NX63\|MIC19_HUMAN MICOS complex subunit MIC19 OS=Homo sapiens GN=CHC | 57,106 | 4,85 | 26152 | 1 | 21,98 | 22,32 | 22,02 | 21,66 | 21,70 | 20,90 | **0,621** | 7,13E-02 | **0** | -0,6866665 | 1,14713011 |
| URP2_HUMA | FERMT3 | sp\|Q86UX7\|URP2_HUMAN Fermitin family homolog 3 OS=Homo sapiens GN=FERMT3 P | 151,727 | 10,49 | 75953 | 6 | 25,07 | 23,68 | 22,81 | 24,62 | 23,80 | 21,08 | **0,621** | 6,14E-01 | **0** | -0,6866665 | 0,21206793 |

| | | | | | | | | | | | | | | | | |
|---|---|---|---|---|---|---|---|---|---|---|---|---|---|---|---|---|
| TOM1_HUM | TOM1 | sp\|O60784\|TOM1_HUMAN Target of Myb protein 1 OS=Homo sapiens GN=TOM1 PE=1 | 164,733 | 11,59 | 53818 | 4 | 23,88 | 23,70 | 23,30 | 22,89 | 23,05 | 22,88 | 0,621 | 1,89E-02 | 0 | -0,6866671 | 1,72416865 |
| TEX10_HUM | TEX10 | sp\|Q9NXF1\|TEX10_HUMAN Testis-expressed protein 10 OS=Homo sapiens GN=TEX10 P | 68,913 | 2,58 | 105674 | 2 | 20,76 | 20,00 | 21,46 | 18,39 | 20,45 | 21,31 | 0,620 | 5,14E-01 | 0 | -0,6891352 | 0,28904755 |
| ECHA_HUMA | HADHA | sp\|P40939\|ECHA_HUMAN Trifunctional enzyme subunit alpha, mitochondrial OS=Homo | 70,037 | 3,54 | 83000 | 2 | 22,20 | 22,27 | 22,93 | 21,99 | 21,69 | 21,65 | 0,620 | 5,44E-02 | 0 | -0,6900005 | 1,26425079 |
| C1QT3_HUM | C1QTNF3 | sp\|Q9BXJ4\|C1QT3_HUMAN Complement C1q tumor necrosis factor-related protein 3 O | 98,438 | 11,79 | 26994 | 3 | 23,68 | 20,77 | 24,45 | 23,38 | 21,13 | 22,32 | 0,620 | 6,22E-01 | 0 | -0,6900012 | 0,20586724 |
| ABCB7_HUM | ABCB7 | sp\|O75027\|ABCB7_HUMAN ATP-binding cassette sub-family B member 7, mitochondria | 25,381 | 1,86 | 82641 | 1 | 19,08 | 18,56 | 19,72 | 18,87 | 18,00 | 18,41 | 0,619 | 1,75E-01 | 0 | -0,6928565 | 0,75654979 |
| P3C2B_HUM | PIK3C2B | sp\|O00750\|P3C2B_HUMAN Phosphatidylinositol 4-phosphate 3-kinase C2 domain-conta | 10,278 | 0,73 | 184768 | 1 | 21,15 | 19,68 | 21,11 | 20,20 | 19,72 | 19,94 | 0,618 | 2,40E-01 | 0 | -0,6933333 | 0,61953473 |
| CTRO_HUMA | CIT | sp\|O14578\|CTRO_HUMAN Citron Rho-interacting kinase OS=Homo sapiens GN=CIT PE= | 959,239 | 18,2 | 231431 | 29 | 24,63 | 23,98 | 27,15 | 24,34 | 23,75 | 25,58 | 0,617 | 5,63E-01 | 0 | -0,6966661 | 0,24928009 |
| IDH3A_HUM | IDH3A | sp\|P50213\|IDH3A_HUMAN Isocitrate dehydrogenase [NAD] subunit alpha, mitochondri | 310,029 | 19,95 | 39592 | 8 | 22,65 | 22,18 | 25,26 | 21,85 | 22,02 | 24,13 | 0,617 | 5,95E-01 | 0 | -0,6966667 | 0,22579278 |
| KRIT1_HUMA | KRIT1 | sp\|O00522\|KRIT1_HUMAN Krev interaction trapped protein 1 OS=Homo sapiens GN=KF | 86,48 | 7,88 | 84348 | 4 | 22,77 | 21,98 | 21,22 | 22,16 | 22,05 | 19,67 | 0,617 | 4,94E-01 | 0 | -0,6966667 | 0,30600622 |
| CLUS_HUMA | CLU | sp\|P10909\|CLUS_HUMAN Clusterin OS=Homo sapiens GN=CLU PE=1 SV=1 | 35,712 | 1,78 | 52495 | 1 | 23,90 | 22,77 | 21,23 | 23,25 | 22,04 | 20,51 | 0,616 | 5,62E-01 | 0 | -0,6999995 | 0,25043608 |
| DIP2C_HUM | DIP2C | sp\|Q9Y2E4\|DIP2C_HUMAN Disco-interacting protein 2 homolog C OS=Homo sapiens GN | 160,307 | 4,5 | 170767 | 6 | 22,78 | 22,40 | 22,35 | 22,96 | 22,20 | 20,27 | 0,616 | 4,37E-01 | 0 | -0,7000001 | 0,35921607 |
| SMUF2_HUN | SMURF2 | sp\|Q9HAU4\|SMUF2_HUMAN E3 ubiquitin-protein ligase SMURF2 OS=Homo sapiens GN | 310,865 | 22,59 | 86196 | 15 | 25,61 | 25,04 | 23,02 | 25,24 | 24,61 | 21,72 | 0,616 | 6,29E-01 | 0 | -0,7000008 | 0,20159391 |
| FAT1_HUMA | FAT1 | sp\|Q14517\|FAT1_HUMAN Protocadherin Fat 1 OS=Homo sapiens GN=FAT1 PE=1 SV=2 | 354,438 | 3,31 | 506273 | 11 | 25,19 | 24,97 | 22,36 | 24,57 | 23,88 | 21,97 | 0,616 | 5,90E-01 | 0 | -0,7000008 | 0,22927629 |
| NPRL2_HUM | NPRL2 | sp\|Q8WTW4\|NPRL2_HUMAN GATOR complex protein NPRL2 OS=Homo sapiens GN=NP | 32,844 | 4,47 | 43658 | 1 | 17,96 | 18,29 | 18,50 | 17,66 | 17,37 | 17,61 | 0,614 | 1,77E-02 | 0 | -0,7033329 | 1,7518066 |
| ADDA_HUM | ADD1 | sp\|P35611\|ADDA_HUMAN Alpha-adducin OS=Homo sapiens GN=ADD1 PE=1 SV=2 | 37,77 | 2,44 | 80955 | 1 | 20,63 | 20,82 | 21,67 | 19,85 | 20,34 | 20,82 | 0,614 | 1,73E-01 | 0 | -0,7033329 | 0,76118856 |
| AKA12_HUM | AKAP12 | sp\|Q02952\|AKA12_HUMAN A-kinase anchor protein 12 OS=Homo sapiens GN=AKAP12 | 80,851 | 1,4 | 191482 | 2 | 23,02 | 22,99 | 21,41 | 22,26 | 22,39 | 20,66 | 0,614 | 4,12E-01 | 0 | -0,7033335 | 0,38464155 |
| PCKGM_HUM | PCK2 | sp\|Q16822\|PCKGM_HUMAN Phosphoenolpyruvate carboxykinase [GTP], mitochondrial | 382,826 | 21,41 | 70730 | 11 | 24,93 | 25,59 | 26,16 | 24,50 | 24,71 | 25,35 | 0,613 | 1,82E-01 | 0 | -0,7066669 | 0,74046256 |
| CD151_HUM | CD151 | sp\|P48509\|CD151_HUMAN CD151 antigen OS=Homo sapiens GN=CD151 PE=1 SV=3 | 158,383 | 10,28 | 28295 | 3 | 26,77 | 26,61 | 25,10 | 25,64 | 26,30 | 24,42 | 0,613 | 4,08E-01 | 0 | -0,7066676 | 0,38904059 |
| JAK1_HUMA | JAK1 | sp\|P23458\|JAK1_HUMAN Tyrosine-protein kinase JAK1 OS=Homo sapiens GN=JAK1 PE= | 281,149 | 12,22 | 133277 | 10 | 24,40 | 24,06 | 23,62 | 23,79 | 23,39 | 22,77 | 0,611 | 1,30E-01 | 0 | -0,7099997 | 0,88737111 |
| GRP75_HUM | HSPA9 | sp\|P38646\|GRP75_HUMAN Stress-70 protein, mitochondrial OS=Homo sapiens GN=HSP | 1371,794 | 45,51 | 73680 | 29 | 26,75 | 27,04 | 28,49 | 26,13 | 26,50 | 27,52 | 0,611 | 3,55E-01 | 0 | -0,7100004 | 0,44930027 |
| SH3G3_HUM | SH3GL3 | sp\|Q99963\|SH3G3_HUMAN Endophilin-A3 OS=Homo sapiens GN=SH3GL3 PE=1 SV=1 | 28,489 | 2,88 | 39285 | 1 | 25,16 | 24,99 | 23,79 | 24,75 | 23,74 | 23,32 | 0,611 | 3,06E-01 | 0 | -0,7100004 | 0,51478738 |
| ATPG_HUMA | ATP5C1 | sp\|P36542\|ATPG_HUMAN ATP synthase subunit gamma, mitochondrial OS=Homo sapie | 61,738 | 6,71 | 32996 | 2 | 21,97 | 22,70 | 24,35 | 21,53 | 22,19 | 23,16 | 0,610 | 4,48E-01 | 0 | -0,7133331 | 0,34897561 |
| UB2D2_HUN | UBE2D2 | sp\|P62837\|UB2D2_HUMAN Ubiquitin-conjugating enzyme E2 D2 OS=Homo sapiens GN= | 181,991 | 42,86 | 16735 | 4 | 20,91 | 20,97 | 20,54 | 20,79 | 19,39 | 20,10 | 0,610 | 1,69E-01 | 0 | -0,7133331 | 0,77135067 |
| SESD1_HUM | SESTD1 | sp\|Q86VW0\|SESD1_HUMAN SEC14 domain and spectrin repeat-containing protein 1 OS | 151,489 | 11,93 | 79348 | 6 | 23,10 | 22,50 | 21,81 | 22,17 | 21,91 | 21,18 | 0,609 | 2,07E-01 | 0 | -0,7166665 | 0,68458062 |
| MDHM_HUN | MDH2 | sp\|P40926\|MDHM_HUMAN Malate dehydrogenase, mitochondrial OS=Homo sapiens G | 466,748 | 34,02 | 35503 | 10 | 27,07 | 27,10 | 28,57 | 26,53 | 26,67 | 27,39 | 0,609 | 2,71E-01 | 0 | -0,7166665 | 0,56636339 |
| T106B_HUM | TMEM106B | sp\|Q9NUM4\|T106B_HUMAN Transmembrane protein 106B OS=Homo sapiens GN=TME | 30,653 | 4,01 | 31127 | 2 | 21,86 | 21,53 | 21,88 | 21,21 | 21,10 | 20,81 | 0,609 | 1,21E-02 | 0 | -0,7166672 | 1,91604161 |
| MYADM_HU | MYADM | sp\|Q96S97\|MYADM_HUMAN Myeloid-associated differentiation marker OS=Homo sapi | 70,479 | 5,59 | 35274 | 2 | 24,93 | 24,36 | 25,39 | 23,76 | 24,36 | 24,40 | 0,607 | 1,18E-01 | 0 | -0,7199999 | 0,92757715 |
| TYB10_HUM | TMSB10 | sp\|P63313\|TYB10_HUMAN Thymosin beta-10 OS=Homo sapiens GN=TMSB10 PE=1 SV= | 52,991 | 31,82 | 5026 | 1 | 21,81 | 21,70 | 20,78 | 22,11 | 21,58 | 18,44 | 0,607 | 5,78E-01 | 0 | -0,7201258 | 0,23803499 |
| RNF31_HUM | RNF31 | sp\|Q96EP0\|RNF31_HUMAN E3 ubiquitin-protein ligase RNF31 OS=Homo sapiens GN=RN | 46,008 | 2,33 | 119652 | 2 | 21,01 | 20,94 | 19,31 | 19,97 | 20,63 | 18,49 | 0,606 | 4,40E-01 | 0 | -0,722215 | 0,35684792 |
| LMOD1_HUN | LMOD1 | sp\|P29536\|LMOD1_HUMAN Leiomodin-1 OS=Homo sapiens GN=LMOD1 PE=1 SV=3 | 47,725 | 3 | 67030 | 2 | 22,59 | 21,80 | 23,00 | 21,18 | 21,69 | 22,35 | 0,606 | 2,13E-01 | 0 | -0,7233327 | 0,671917 |
| ADAM9_HUN | ADAM9 | sp\|Q13443\|ADAM9_HUMAN Disintegrin and metalloproteinase domain-containing prot | 335,923 | 11,97 | 90556 | 12 | 25,78 | 25,76 | 24,67 | 25,03 | 25,16 | 23,84 | 0,604 | 2,62E-01 | 0 | -0,7266668 | 0,58097172 |
| RHG05_HUN | ARHGAP5 | sp\|Q13017\|RHG05_HUMAN Rho GTPase-activating protein 5 OS=Homo sapiens GN=ARH | 9,997 | 0,73 | 172460 | 1 | 20,51 | 20,50 | 20,82 | 19,57 | 19,57 | 20,51 | 0,604 | 9,28E-02 | 0 | -0,7266668 | 1,03260021 |
| ADCY9_HUN | ADCY9 | sp\|O60503\|ADCY9_HUMAN Adenylate cyclase type 9 OS=Homo sapiens GN=ADCY9 PE= | 57,444 | 3,84 | 150701 | 4 | 22,13 | 22,68 | 19,69 | 21,49 | 22,25 | 18,57 | 0,603 | 6,41E-01 | 0 | -0,7290332 | 0,19289496 |
| DDR2_HUMA | DDR2 | sp\|Q16832\|DDR2_HUMAN Discoidin domain-containing receptor 2 OS=Homo sapiens G | 151,706 | 7,84 | 96736 | 6 | 23,91 | 24,13 | 23,48 | 22,81 | 23,79 | 22,73 | 0,603 | 1,35E-01 | 0 | -0,7299995 | 0,86967087 |
| EPHB3_HUM | EPHB3 | sp\|P54753\|EPHB3_HUMAN Ephrin type-B receptor 3 OS=Homo sapiens GN=EPHB3 PE= | 478,377 | 19,64 | 110330 | 15 | 24,12 | 23,95 | 23,22 | 23,40 | 23,33 | 22,36 | 0,602 | 1,67E-01 | 0 | -0,7333336 | 0,77786416 |
| KIF14_HUMA | KIF14 | sp\|Q15058\|KIF14_HUMAN Kinesin-like protein KIF14 OS=Homo sapiens GN=KIF14 PE=1 | 184,844 | 4,79 | 186492 | 8 | 22,76 | 21,41 | 25,66 | 21,94 | 21,50 | 24,18 | 0,600 | 6,50E-01 | 0 | -0,7366664 | 0,18721185 |
| AKT2_HUMA | AKT2 | sp\|P31751\|AKT2_HUMAN RAC-beta serine/threonine-protein kinase OS=Homo sapiens | 31,419 | 2,29 | 55769 | 1 | 18,39 | 19,43 | 19,00 | 17,46 | 18,07 | 19,07 | 0,599 | 2,56E-01 | 0 | -0,7398307 | 0,59236327 |
| EPN2_HUMA | EPN2 | sp\|O95208\|EPN2_HUMAN Epsin-2 OS=Homo sapiens GN=EPN2 PE=1 SV=3 | 155,271 | 7,64 | 68482 | 4 | 24,32 | 23,95 | 24,68 | 23,78 | 23,56 | 23,38 | 0,597 | 3,65E-02 | 0 | -0,7433338 | 1,43773334 |
| ABRX2_HUM | ABRAXAS2 | sp\|Q15018\|ABRX2_HUMAN BRISC complex subunit Abraxas 2 OS=Homo sapiens GN=AB | 29,701 | 5,78 | 46901 | 2 | 18,49 | 18,81 | 20,77 | 18,67 | 17,94 | 19,23 | 0,597 | 4,08E-01 | 0 | -0,7437509 | 0,38945862 |
| IMPA3_HUM | IMPAD1 | sp\|Q9NX62\|IMPA3_HUMAN Inositol monophosphatase 3 OS=Homo sapiens GN=IMPAD | 10,166 | 2,79 | 38681 | 1 | 18,96 | 20,94 | 19,86 | 16,99 | 19,98 | 20,56 | 0,597 | 5,82E-01 | 0 | -0,7448279 | 0,2347997 |
| S10A7_HUM | S100A7 | sp\|P31151\|S10A7_HUMAN Protein S100-A7 OS=Homo sapiens GN=S100A7 PE=1 SV=4 | 11,184 | 10,89 | 11471 | 1 | 21,14 | 20,36 | 21,68 | 21,14 | 20,55 | 19,25 | 0,596 | 3,32E-01 | 0 | -0,7461948 | 0,47877696 |
| PTBP3_HUM | PTBP3 | sp\|O95758\|PTBP3_HUMAN Polypyrimidine tract-binding protein 3 OS=Homo sapiens GN | 125,456 | 13,59 | 59690 | 6 | 21,88 | 22,18 | 22,71 | 21,53 | 21,64 | 21,36 | 0,596 | 4,34E-02 | 0 | -0,746666 | 1,36287263 |
| GPM6A_HUN | GPM6A | sp\|P51674\|GPM6A_HUMAN Neuronal membrane glycoprotein M6-a OS=Homo sapiens | 230,284 | 15,83 | 31210 | 7 | 26,91 | 26,07 | 24,75 | 25,88 | 25,58 | 24,03 | 0,596 | 4,30E-01 | 0 | -0,7466666 | 0,36684034 |
| PR38A_HUM | PRPF38A | sp\|Q8NAV1\|PR38A_HUMAN Pre-mRNA-splicing factor 38A OS=Homo sapiens GN=PRPF3 | 59,275 | 8,97 | 37477 | 2 | 18,26 | 20,65 | 22,87 | 18,69 | 18,14 | 22,70 | 0,596 | 7,22E-01 | 0 | -0,7475135 | 0,14133635 |
| SHSA4_HUM | SHISA4 | sp\|Q96DD7\|SHSA4_HUMAN Protein shisa-4 OS=Homo sapiens GN=SHISA4 PE=2 SV=3 | 61,342 | 23,86 | 21522 | 3 | 25,80 | 25,86 | 24,59 | 25,72 | 25,16 | 23,11 | 0,593 | 4,47E-01 | 0 | -0,7533334 | 0,34951017 |
| SMAD5_HUN | SMAD5 | sp\|Q99717\|SMAD5_HUMAN Mothers against decapentaplegic homolog 5 OS=Homo sap | 104,235 | 13,55 | 52258 | 4 | 22,28 | 22,34 | 20,55 | 21,50 | 20,97 | 20,44 | 0,593 | 3,19E-01 | 0 | -0,7533334 | 0,49674653 |
| PGAM4_HUN | PGAM4 | sp\|Q8N0Y7\|PGAM4_HUMAN Probable phosphoglycerate mutase 4 OS=Homo sapiens G | 383,006 | 31,5 | 28777 | 8 | 20,80 | 20,00 | 19,83 | 20,50 | 18,28 | 19,59 | 0,593 | 3,49E-01 | 0 | -0,7535191 | 0,45702883 |
| S35U4_HUM | SLC35A4 | sp\|L0R6Q1\|S35U4_HUMAN SLC35A4 upstream open reading frame protein OS=Homo s | 16,798 | 9,71 | 11133 | 1 | 21,15 | 21,12 | 19,18 | 20,21 | 20,72 | 18,26 | 0,593 | 4,90E-01 | 0 | -0,7541784 | 0,31009236 |
| C42S1_HUM | CDC42SE1 | sp\|Q9NRR8\|C42S1_HUMAN CDC42 small effector protein 1 OS=Homo sapiens GN=CDC4 | 52,869 | 15,19 | 8925 | 1 | 21,73 | 21,84 | 18,95 | 21,45 | 20,52 | 18,28 | 0,591 | 6,00E-01 | 0 | -0,7581825 | 0,22155293 |
| ECHB_HUMA | HADHB | sp\|P55084\|ECHB_HUMAN Trifunctional enzyme subunit beta, mitochondrial OS=Homo | 38,242 | 5,91 | 51294 | 3 | 20,82 | 21,37 | 23,60 | 21,20 | 19,97 | 22,34 | 0,590 | 5,24E-01 | 0 | -0,7600002 | 0,28026897 |
| PHB2_HUMA | PHB2 | sp\|Q99623\|PHB2_HUMAN Prohibitin-2 OS=Homo sapiens GN=PHB2 PE=1 SV=2 | 573,609 | 58,19 | 33296 | 17 | 27,19 | 26,97 | 28,58 | 26,47 | 26,49 | 27,50 | 0,590 | 2,79E-01 | 0 | -0,7600002 | 0,55370974 |
| KIF3A_HUM | KIF3A | sp\|Q9Y496\|KIF3A_HUMAN Kinesin-like protein KIF3A OS=Homo sapiens GN=KIF3A PE=1 | 290,38 | 13,73 | 80041 | 7 | 20,61 | 19,46 | 23,84 | 18,68 | 19,88 | 23,07 | 0,590 | 7,03E-01 | 0 | -0,7600002 | 0,15318849 |
| ACOT9_HUN | ACOT9 | sp\|Q9Y305\|ACOT9_HUMAN Acyl-coenzyme A thioesterase 9, mitochondrial OS=Homo s | 193,13 | 18 | 49902 | 7 | 22,93 | 22,67 | 25,54 | 22,39 | 22,13 | 24,34 | 0,590 | 5,45E-01 | 0 | -0,7600009 | 0,26332829 |
| APC4_HUMA | ANAPC4 | sp\|Q9UJX5\|APC4_HUMAN Anaphase-promoting complex subunit 4 OS=Homo sapiens G | 69,784 | 3,22 | 92116 | 2 | 20,07 | 21,14 | 20,48 | 19,34 | 20,09 | 19,96 | 0,588 | 1,19E-01 | 0 | -0,7666664 | 0,92268983 |
| VP37B_HUM | VPS37B | sp\|Q9H9H4\|VP37B_HUMAN Vacuolar protein sorting-associated protein 37B OS=Homo | 190,925 | 28,42 | 31307 | 6 | 25,81 | 25,23 | 24,34 | 24,42 | 24,72 | 23,94 | 0,588 | 1,88E-01 | 0 | -0,7666664 | 0,72485309 |
| PCDH7_HUM | PCDH7 | sp\|O60245\|PCDH7_HUMAN Protocadherin-7 OS=Homo sapiens GN=PCDH7 PE=1 SV=2 | 474,94 | 14,78 | 116071 | 13 | 26,84 | 26,61 | 25,88 | 26,05 | 25,69 | 25,29 | 0,588 | 1,02E-01 | 0 | -0,7666664 | 0,98970914 |
| PLXA2_HUM | PLXNA2 | sp\|O75051\|PLXA2_HUMAN Plexin-A2 OS=Homo sapiens GN=PLXNA2 PE=1 SV=4 | 126,123 | 3,17 | 211104 | 6 | 22,44 | 22,54 | 21,92 | 21,81 | 21,74 | 21,05 | 0,588 | 6,84E-02 | 0 | -0,7666677 | 1,16507741 |
| NRBP_HUMA | NRBP1 | sp\|Q9UHY1\|NRBP_HUMAN Nuclear receptor-binding protein OS=Homo sapiens GN=NR | 33,022 | 2,8 | 59845 | 1 | 20,79 | 21,16 | 21,39 | 17,85 | 21,66 | 21,52 | 0,587 | 5,74E-01 | 0 | -0,7687689 | 0,24090903 |
| MICA2_HUM | MICAL2 | sp\|O94851\|MICA2_HUMAN [F-actin]-monooxygenase MICAL2 OS=Homo sapiens GN=M | 13,972 | 1,07 | 126689 | 1 | 18,10 | 19,27 | 20,85 | 18,22 | 16,80 | 20,89 | 0,586 | 6,21E-01 | 0 | -0,771143 | 0,20703932 |
| MRC2_HUM | MRC2 | sp\|Q9UBG0\|MRC2_HUMAN C-type mannose receptor 2 OS=Homo sapiens GN=MRC2 P | 15,565 | 1,08 | 166674 | 1 | 20,51 | 19,91 | 20,44 | 20,07 | 18,68 | 19,79 | 0,585 | 1,71E-01 | 0 | -0,7727928 | 0,76609746 |
| LIX1L_HUMA | LIX1L | sp\|Q8IVB5\|LIX1L_HUMAN LIX1-like protein OS=Homo sapiens GN=LIX1L PE=2 SV=1 | 98,115 | 11,28 | 36563 | 3 | 22,01 | 21,65 | 20,05 | 21,50 | 20,81 | 19,08 | 0,585 | 4,56E-01 | 0 | -0,7733332 | 0,34078404 |
| ASH2L_HUM | ASH2L | sp\|Q9UBL3\|ASH2L_HUMAN Set1/Ash2 histone methyltransferase complex subunit ASH | 30,988 | 2,39 | 68723 | 1 | 19,04 | 20,17 | 20,74 | 17,45 | 19,65 | 20,51 | 0,582 | 4,94E-01 | 0 | -0,7799842 | 0,30615479 |
| CCD50_HUM | CCDC50 | sp\|Q8IVM0\|CCD50_HUMAN Coiled-coil domain-containing protein 50 OS=Homo sapien | 144,562 | 19,28 | 35822 | 3 | 24,68 | 24,48 | 21,84 | 23,62 | 23,55 | 21,49 | 0,582 | 5,35E-01 | 0 | -0,7800001 | 0,27142819 |
| MPDZ_HUM | MPDZ | sp\|O75970\|MPDZ_HUMAN Multiple PDZ domain protein OS=Homo sapiens GN=MPDZ P | 51,221 | 1,5 | 221618 | 3 | 23,28 | 23,28 | 24,45 | 22,18 | 22,41 | 24,08 | 0,582 | 3,36E-01 | 0 | -0,7800007 | 0,47323071 |
| RCC2_HUMA | RCC2 | sp\|Q9P258\|RCC2_HUMAN Protein RCC2 OS=Homo sapiens GN=RCC2 PE=1 SV=2 | 198,277 | 15,52 | 56085 | 7 | 26,20 | 26,33 | 24,65 | 24,57 | 25,81 | 24,46 | 0,582 | 3,23E-01 | 0 | -0,7800007 | 0,49137178 |
| UB2D1_HUN | UBE2D1 | sp\|P51668\|UB2D1_HUMAN Ubiquitin-conjugating enzyme E2 D1 OS=Homo sapiens GN= | 52,244 | 30,61 | 16602 | 2 | 19,19 | 20,23 | 20,36 | 19,18 | 19,68 | 18,58 | 0,582 | 1,85E-01 | 0 | -0,7812144 | 0,73235694 |
| PRUN1_HUN | PRUNE1 | sp\|Q86TP1\|PRUN1_HUMAN Exopolyphosphatase PRUNE1 OS=Homo sapiens GN=PRUN | 12,631 | 2,43 | 50200 | 1 | 21,53 | 21,77 | 21,32 | 18,40 | 22,00 | 21,87 | 0,581 | 5,45E-01 | 0 | -0,7824987 | 0,26343894 |
| ADT1_HUMA | SLC25A4 | sp\|P12235\|ADT1_HUMAN ADP/ATP translocase 1 OS=Homo sapiens GN=SLC25A4 PE=1 | 688,69 | 50,34 | 33064 | 15 | 26,33 | 26,55 | 27,12 | 25,47 | 25,93 | 26,25 | 0,581 | 7,45E-02 | 0 | -0,7833335 | 1,12801363 |
| NHRF2_HUM | SLC9A3R2 | sp\|Q15599\|NHRF2_HUMAN Na(+)/H(+) exchange regulatory cofactor NHE-RF2 OS=Hom | 25,376 | 10,09 | 37414 | 2 | 22,62 | 22,83 | 19,76 | 22,72 | 22,02 | 18,11 | 0,580 | 6,76E-01 | 0 | -0,7858696 | 0,17032194 |
| PHB_HUMAN | PHB | sp\|P35232\|PHB_HUMAN Prohibitin OS=Homo sapiens GN=PHB PE=1 SV=1 | 581,249 | 56,25 | 29804 | 11 | 26,45 | 26,28 | 28,00 | 25,88 | 25,63 | 26,86 | 0,580 | 3,01E-01 | 0 | -0,7866675 | 0,52082193 |
| GLYM_HUM | SHMT2 | sp\|P34897\|GLYM_HUMAN Serine hydroxymethyltransferase, mitochondrial OS=Homo s | 468,389 | 36,9 | 55993 | 15 | 26,73 | 26,95 | 27,50 | 26,11 | 26,18 | 26,52 | 0,578 | 3,92E-02 | 0 | -0,7899996 | 1,40676613 |
| TYSY_HUMA | TYMS | sp\|P04818\|TYSY_HUMAN Thymidylate synthase OS=Homo sapiens GN=TYMS PE=1 SV= | 43,808 | 3,51 | 35716 | 1 | 18,16 | 19,23 | 19,11 | 18,20 | 18,01 | 17,92 | 0,578 | 8,59E-02 | 0 | -0,7905814 | 1,06609889 |
| TR112_HUM | TRMT112 | sp\|Q9UI30\|TR112_HUMAN Multifunctional methyltransferase subunit TRM112-like prot | 19,531 | 9,6 | 14199 | 1 | 22,42 | 22,55 | 19,69 | 22,44 | 22,27 | 17,56 | 0,576 | 6,89E-01 | 0 | -0,7958336 | 0,16165266 |
| CO4A1_HUN | COL4A1 | sp\|P02462\|CO4A1_HUMAN Collagen alpha-1(IV) chain OS=Homo sapiens GN=COL4A1 P | 27,622 | 0,84 | 160615 | 1 | 17,91 | 19,72 | 21,32 | 18,07 | 17,14 | 21,33 | 0,573 | 6,44E-01 | 0 | -0,8022162 | 0,1910662 |
| MTMR5_HUI | SBF1 | sp\|O95248\|MTMR5_HUMAN Myotubularin-related protein 5 OS=Homo sapiens GN=SBF | 67,415 | 1,66 | 208315 | 3 | 21,44 | 21,61 | 20,98 | 21,11 | 21,54 | 18,97 | 0,573 | 3,81E-01 | 0 | -0,8033333 | 0,4190748 |
| LIPB1_HUMA | PPFIBP1 | sp\|Q86W92\|LIPB1_HUMAN Liprin-beta-1 OS=Homo sapiens GN=PPFIBP1 PE=1 SV=2 | 89,013 | 3,56 | 114024 | 2 | 21,99 | 22,01 | 21,28 | 21,06 | 21,86 | 19,93 | 0,571 | 2,54E-01 | 0 | -0,8087578 | 0,59464865 |
| SDC1_HUMA | SDC1 | sp\|P18827\|SDC1_HUMAN Syndecan-1 OS=Homo sapiens GN=SDC1 PE=1 SV=3 | 180,626 | 4,19 | 32462 | 3 | 24,95 | 24,82 | 21,14 | 24,28 | 23,76 | 20,44 | 0,570 | 6,65E-01 | 0 | -0,8099995 | 0,17737631 |
| ATPA_HUMA | ATP5A1 | sp\|P25705\|ATPA_HUMAN ATP synthase subunit alpha, mitochondrial OS=Homo sapiens | 1189,516 | 49,91 | 59751 | 27 | 28,53 | 28,62 | 29,80 | 27,93 | 27,90 | 28,69 | 0,570 | 1,70E-01 | 0 | -0,8100001 | 0,7708092 |
| VANG1_HUN | VANGL1 | sp\|Q8TAA9\|VANG1_HUMAN Vang-like protein 1 OS=Homo sapiens GN=VANGL1 PE=1 S | 371,125 | 22,71 | 59975 | 9 | 26,58 | 24,57 | 24,47 | 24,58 | 24,64 | 23,97 | 0,570 | 3,23E-01 | 0 | -0,8100001 | 0,49022406 |
| ACADL_HUM | ACADL | sp\|P28330\|ACADL_HUMAN Long-chain specific acyl-CoA dehydrogenase, mitochondrial | 78,019 | 5,58 | 47656 | 3 | 22,05 | 20,30 | 24,57 | 21,01 | 20,14 | 23,33 | 0,569 | 6,30E-01 | 0 | -0,8133329 | 0,20049651 |
| TBB2B_HUM | TUBB2B | sp\|Q9BVA1\|TBB2B_HUMAN Tubulin beta-2B chain OS=Homo sapiens GN=TUBB2B PE=1 | 2598,45 | 84,72 | 49953 | 55 | 25,74 | 24,76 | 25,68 | 23,76 | 24,07 | 25,91 | 0,569 | 3,35E-01 | 0 | -0,8133335 | 0,47537184 |
| PAB4L_HUM | PABPC4L | sp\|P0CB38\|PAB4L_HUMAN Polyadenylate-binding protein 4-like OS=Homo sapiens GN= | 128,804 | 5,95 | 41854 | 3 | 21,61 | 21,37 | 19,45 | 21,03 | 20,71 | 18,25 | 0,568 | 5,04E-01 | 0 | -0,8154329 | 0,2972013 |
| NDUS1_HUN | NDUFS1 | sp\|P28331\|NDUS1_HUMAN NADH-ubiquinone oxidoreductase 75 kDa subunit, mitocho | 10,616 | 1,38 | 79468 | 1 | 19,68 | 18,14 | 20,95 | 18,75 | 18,38 | 19,19 | 0,567 | 3,88E-01 | 0 | -0,8189831 | 0,41162074 |
| ZDHC5_HUM | ZDHHC5 | sp\|Q9C0B5\|ZDHC5_HUMAN Palmitoyltransferase ZDHHC5 OS=Homo sapiens GN=ZDHH | 72,032 | 10,07 | 77545 | 4 | 23,23 | 23,03 | 20,71 | 23,21 | 22,62 | 18,68 | 0,566 | 6,43E-01 | 0 | -0,8199997 | 0,19212429 |
| PLK1_HUMA | PLK1 | sp\|P53350\|PLK1_HUMAN Serine/threonine-protein kinase PLK1 OS=Homo sapiens GN= | 216,918 | 6,47 | 68255 | 4 | 21,15 | 21,57 | 23,81 | 19,94 | 20,25 | 23,88 | 0,566 | 6,16E-01 | 0 | -0,8199997 | 0,21038835 |
| CXA1_HUMA | GJA1 | sp\|P17302\|CXA1_HUMAN Gap junction alpha-1 protein OS=Homo sapiens GN=GJA1 PE= | 650,104 | 47,38 | 43008 | 17 | 28,27 | 27,39 | 26,29 | 27,31 | 26,85 | 25,33 | 0,566 | 3,78E-01 | 0 | -0,8200003 | 0,42229554 |
| ATPB_HUMA | ATP5B | sp\|P06576\|ATPB_HUMAN ATP synthase subunit beta, mitochondrial OS=Homo sapiens | 1806,889 | 73,91 | 56560 | 36 | 28,35 | 28,58 | 29,86 | 27,72 | 27,92 | 28,68 | 0,565 | 2,11E-01 | 0 | -0,8233337 | 0,67579528 |
| MK_HUMAN | MDK | sp\|P21741\|MK_HUMAN Midkine OS=Homo sapiens GN=MDK PE=1 SV=1 | 34,406 | 6,99 | 15585 | 1 | 22,01 | 20,82 | 20,74 | 21,90 | 20,61 | 18,58 | 0,564 | 4,76E-01 | 0 | -0,8262609 | 0,32251941 |
| RGAP1_HUM | RACGAP1 | sp\|Q9H0H5\|RGAP1_HUMAN Rac GTPase-activating protein 1 OS=Homo sapiens GN=RAC | 1098,336 | 39,56 | 71027 | 26 | 27,54 | 27,40 | 30,64 | 26,95 | 26,94 | 29,20 | 0,563 | 5,57E-01 | 0 | -0,8299993 | 0,25403762 |
| MKLN1_HUN | MKLN1 | sp\|Q9UL63\|MKLN1_HUMAN Muskelin OS=Homo sapiens GN=MKLN1 PE=1 SV=2 | 19,4 | 1,77 | 84768 | 1 | 20,44 | 20,72 | 20,90 | 18,56 | 20,55 | 20,45 | 0,562 | 2,76E-01 | 0 | -0,8326314 | 0,5593167 |
| ERBIN_HUM | ERBIN | sp\|Q96RT1\|ERBIN_HUMAN Erbin OS=Homo sapiens GN=ERBIN PE=1 SV=2 | 174,284 | 6,02 | 158298 | 6 | 23,34 | 23,02 | 23,35 | 22,50 | 22,47 | 22,24 | 0,561 | 3,59E-03 | 0 | -0,833334 | 2,44470891 |
| ITBP1_HUM | ITGB1BP1 | sp\|O14713\|ITBP1_HUMAN Integrin beta-1-binding protein 1 OS=Homo sapiens GN=ITG | 28,878 | 7,5 | 21782 | 1 | 17,28 | 21,33 | 21,50 | 20,92 | 18,35 | 18,34 | 0,561 | 6,34E-01 | 0 | -0,8345 | 0,19768053 |
| ROMO1_HUI | ROMO1 | sp\|P60602\|ROMO1_HUMAN Reactive oxygen species modulator 1 OS=Homo sapiens GI | 48,2 | 21,52 | 8183 | 1 | 20,25 | 20,46 | 19,17 | 18,32 | 19,89 | 19,16 | 0,560 | 2,39E-01 | 0 | -0,835818 | 0,62222068 |
| S39AA_HUM | SLC39A10 | sp\|Q9ULF5\|S39AA_HUMAN Zinc transporter ZIP10 OS=Homo sapiens GN=SLC39A10 PE= | 31,092 | 2,17 | 94132 | 2 | 23,00 | 23,43 | 21,30 | 22,33 | 22,77 | 20,12 | 0,560 | 4,69E-01 | 0 | -0,8366661 | 0,32902948 |
| BGH3_HUM | TGFBI | sp\|Q15582\|BGH3_HUMAN Transforming growth factor-beta-induced protein ig-h3 OS= | 109,019 | 8,35 | 74681 | 4 | 21,29 | 19,52 | 23,24 | 20,44 | 18,27 | 22,83 | 0,560 | 6,48E-01 | 0 | -0,8366667 | 0,18822413 |
| WDR26_HUN | WDR26 | sp\|Q9H7D7\|WDR26_HUMAN WD repeat-containing protein 26 OS=Homo sapiens GN=W | 21,622 | 1,51 | 72124 | 1 | 17,75 | 19,89 | 20,85 | 16,91 | 18,56 | 20,51 | 0,559 | 5,78E-01 | 0 | -0,8382924 | 0,23821709 |
| MCU_HUMA | MCU | sp\|Q8NE86\|MCU_HUMAN Calcium uniporter protein, mitochondrial OS=Homo sapiens | 269,347 | 27,92 | 39867 | 7 | 23,74 | 24,15 | 25,61 | 23,10 | 23,43 | 24,45 | 0,559 | 2,95E-01 | 0 | -0,8399995 | 0,52998915 |
| HCD2_HUM | HSD17B10 | sp\|Q99714\|HCD2_HUMAN 3-hydroxyacyl-CoA dehydrogenase type-2 OS=Homo sapiens | 193,501 | 29,5 | 26923 | 4 | 23,95 | 24,06 | 25,65 | 23,04 | 23,72 | 24,38 | 0,559 | 2,79E-01 | 0 | -0,8400002 | 0,55389779 |
| NFKB2_HUM | NFKB2 | sp\|Q00653\|NFKB2_HUMAN Nuclear factor NF-kappa-B p100 subunit OS=Homo sapiens | 20,159 | 2 | 96749 | 1 | 20,15 | 20,45 | 20,96 | 18,58 | 19,70 | 20,76 | 0,558 | 2,80E-01 | 0 | -0,841156 | 0,55337432 |
| PTPRC_HUM | PTPRC | sp\|P08575\|PTPRC_HUMAN Receptor-type tyrosine-protein phosphatase C OS=Homo sa | 54,946 | 2,15 | 147254 | 2 | 22,04 | 20,93 | 20,87 | 21,13 | 21,11 | 19,07 | 0,557 | 3,42E-01 | 0 | -0,843249 | 0,46649504 |
| CRCM1_HUN | ORAI1 | sp\|Q96D31\|CRCM1_HUMAN Calcium release-activated calcium channel protein 1 OS=H | 20,187 | 3,99 | 32668 | 1 | 20,79 | 20,56 | 20,50 | 20,56 | 19,42 | 19,34 | 0,557 | 1,05E-01 | 0 | -0,8433336 | 0,97876862 |
| PAK6_HUMA | PAK6 | sp\|Q9NQU5\|PAK6_HUMAN Serine/threonine-protein kinase PAK 6 OS=Homo sapiens G | 280,315 | 15,71 | 74869 | 7 | 24,37 | 23,85 | 20,01 | 24,00 | 23,65 | 18,05 | 0,557 | 7,40E-01 | 0 | -0,8433342 | 0,13094489 |
| AL1L2_HUM | ALDH1L2 | sp\|Q3SY69\|AL1L2_HUMAN Mitochondrial 10-formyltetrahydrofolate dehydrogenase O | 31,375 | 2,93 | 101746 | 2 | 20,48 | 20,88 | 21,30 | 18,22 | 20,19 | 21,71 | 0,557 | 4,60E-01 | 0 | -0,8454793 | 0,3367795 |
| ARVC_HUMA | ARVCF | sp\|O00192\|ARVC_HUMAN Armadillo repeat protein deleted in velo-cardio-facial syndro | 153,265 | 6,44 | 104642 | 4 | 23,21 | 23,05 | 20,81 | 22,43 | 22,44 | 19,66 | 0,556 | 5,22E-01 | 0 | -0,8466657 | 0,28271362 |
| INT14_HUM | INTS14 | sp\|Q96SY0\|INT14_HUMAN Integrator complex subunit 14 OS=Homo sapiens GN=INTS1 | 42,003 | 2,32 | 57471 | 1 | 18,71 | 20,03 | 19,85 | 18,08 | 18,09 | 19,88 | 0,555 | 3,08E-01 | 0 | -0,8490213 | 0,51204802 |
| NOTC3_HUN | NOTCH3 | sp\|Q9UM47\|NOTC3_HUMAN Neurogenic locus notch homolog protein 3 OS=Homo sap | 34,459 | 0,95 | 243631 | 2 | 21,42 | 21,14 | 20,65 | 21,45 | 20,60 | 18,61 | 0,554 | 3,84E-01 | 0 | -0,8518143 | 0,41554819 |
| GRIN1_HUM | GPRIN1 | sp\|Q7Z2K8\|GRIN1_HUMAN G protein-regulated inducer of neurite outgrowth 1 OS=Hor | 16,199 | 0,79 | 102399 | 1 | 22,31 | 22,16 | 22,81 | 21,87 | 21,83 | 21,02 | 0,554 | 6,58E-02 | 0 | -0,8533325 | 1,18155489 |
| IDH3B_HUM | IDH3B | sp\|O43837\|IDH3B_HUMAN Isocitrate dehydrogenase [NAD] subunit beta, mitochondria | 101,975 | 12,99 | 42184 | 4 | 22,46 | 21,70 | 24,12 | 21,33 | 21,89 | 22,50 | 0,554 | 3,41E-01 | 0 | -0,8533338 | 0,46718992 |
| PSME1_HUN | PSME1 | sp\|Q06323\|PSME1_HUMAN Proteasome activator complex subunit 1 OS=Homo sapiens | 162,021 | 36,55 | 28723 | 7 | 24,28 | 27,45 | 22,82 | 24,16 | 24,26 | 23,57 | 0,554 | 5,71E-01 | 0 | -0,8533338 | 0,24356393 |
| BRCC3_HUM | BRCC3 | sp\|P46736\|BRCC3_HUMAN Lys-63-specific deubiquitinase BRCC36 OS=Homo sapiens GI | 10,081 | 3,48 | 36072 | 1 | 20,71 | 19,06 | 20,89 | 19,93 | 18,01 | 20,16 | 0,553 | 3,94E-01 | 0 | -0,8551782 | 0,40459647 |
| ATD3A_HUM | ATAD3A | sp\|Q9NVI7\|ATD3A_HUMAN ATPase family AAA domain-containing protein 3A OS=Homo | 71,819 | 7,41 | 71369 | 4 | 21,65 | 20,28 | 22,53 | 20,83 | 19,89 | 21,17 | 0,552 | 3,22E-01 | 0 | -0,8566672 | 0,49245134 |
| FMNL2_HUN | FMNL2 | sp\|Q96PY5\|FMNL2_HUMAN Formin-like protein 2 OS=Homo sapiens GN=FMNL2 PE=1 S | 196,425 | 7,46 | 123321 | 7 | 21,67 | 22,12 | 20,33 | 21,05 | 21,58 | 18,92 | 0,552 | 4,29E-01 | 0 | -0,8575191 | 0,36764279 |
| ACADM_HUI | ACADM | sp\|P11310\|ACADM_HUMAN Medium-chain specific acyl-CoA dehydrogenase, mitochon | 20,37 | 2,14 | 46588 | 1 | 18,53 | 19,77 | 22,09 | 18,86 | 17,87 | 21,09 | 0,552 | 5,77E-01 | 0 | -0,8580914 | 0,23919064 |
| PEBB_HUMA | CBFB | sp\|Q13951\|PEBB_HUMAN Core-binding factor subunit beta OS=Homo sapiens GN=CBFE | 74,435 | 14,29 | 21508 | 2 | 21,44 | 21,12 | 21,10 | 21,13 | 21,34 | 18,62 | 0,552 | 3,86E-01 | 0 | -0,85823 | 0,41370335 |
| CDK2_HUMA | CDK2 | sp\|P24941\|CDK2_HUMAN Cyclin-dependent kinase 2 OS=Homo sapiens GN=CDK2 PE=1 | 121,121 | 19,8 | 33930 | 5 | 20,99 | 20,84 | 22,49 | 18,80 | 20,58 | 22,36 | 0,551 | 4,98E-01 | 0 | -0,8594824 | 0,3028307 |
| THIL_HUMAI | ACAT1 | sp\|P24752\|THIL_HUMAN Acetyl-CoA acetyltransferase, mitochondrial OS=Homo sapien | 175,042 | 13,82 | 45200 | 4 | 23,99 | 24,16 | 24,75 | 23,24 | 23,37 | 23,71 | 0,551 | 3,32E-02 | 0 | -0,86 | 1,47877252 |
| GPC6_HUMA | GPC6 | sp\|Q9Y625\|GPC6_HUMAN Glypican-6 OS=Homo sapiens GN=GPC6 PE=1 SV=1 | 567,164 | 34,05 | 62736 | 16 | 25,92 | 25,71 | 26,96 | 24,98 | 25,07 | 25,95 | 0,550 | 1,56E-01 | 0 | -0,8633327 | 0,80651718 |
| TRI32_HUM | TRIM32 | sp\|Q13049\|TRI32_HUMAN E3 ubiquitin-protein ligase TRIM32 OS=Homo sapiens GN=TF | 45,553 | 4,13 | 71989 | 2 | 20,98 | 21,02 | 19,59 | 20,22 | 20,17 | 18,59 | 0,547 | 2,89E-01 | 0 | -0,8700002 | 0,53904483 |
| PHOCN_HUN | MOB4 | sp\|Q9Y3A3\|PHOCN_HUMAN MOB-like protein phocein OS=Homo sapiens GN=MOB4 PE | 24,893 | 6,22 | 26032 | 1 | 19,79 | 20,16 | 20,50 | 16,62 | 19,96 | 21,26 | 0,547 | 5,67E-01 | 0 | -0,871268 | 0,2464313 |
| VDAC1_HUN | VDAC1 | sp\|P21796\|VDAC1_HUMAN Voltage-dependent anion-selective channel protein 1 OS=H | 864,35 | 66,08 | 30773 | 15 | 27,44 | 27,53 | 28,16 | 26,63 | 26,80 | 27,08 | 0,546 | 2,89E-02 | 0 | -0,8733342 | 1,53868119 |
| DYM_HUMA | DYM | sp\|Q7RTS9\|DYM_HUMAN Dymeclin OS=Homo sapiens GN=DYM PE=1 SV=1 | 24,731 | 1,49 | 75935 | 1 | 19,82 | 20,11 | 19,32 | 18,19 | 18,19 | 20,24 | 0,546 | 2,91E-01 | 0 | -0,8741627 | 0,53562839 |
| ODO2_HUM | DLST | sp\|P36957\|ODO2_HUMAN Dihydrolipoyllysine-residue succinyltransferase component | 171,838 | 8,39 | 48755 | 4 | 24,65 | 24,32 | 25,24 | 24,15 | 23,86 | 23,57 | 0,545 | 5,05E-02 | 0 | -0,8766664 | 1,29643886 |
| PPIF_HUMAI | PPIF | sp\|P30405\|PPIF_HUMAN Peptidyl-prolyl cis-trans isomerase F, mitochondrial OS=Homo | 11,178 | 5,31 | 22040 | 1 | 21,71 | 21,83 | 21,78 | 21,29 | 21,19 | 20,20 | 0,543 | 6,56E-02 | 0 | -0,8799992 | 1,18338514 |
| P5CS_HUMA | ALDH18A1 | sp\|P54886\|P5CS_HUMAN Delta-1-pyrroline-5-carboxylate synthase OS=Homo sapiens G | 362,584 | 17,11 | 87302 | 10 | 23,62 | 23,52 | 25,27 | 23,03 | 22,78 | 23,96 | 0,543 | 2,60E-01 | 0 | -0,8800004 | 0,58474635 |
| CISY_HUMAI | CS | sp\|O75390\|CISY_HUMAN Citrate synthase, mitochondrial OS=Homo sapiens GN=CS PE= | 292,83 | 17,38 | 51712 | 7 | 25,27 | 25,31 | 26,68 | 24,65 | 24,60 | 25,36 | 0,542 | 1,67E-01 | 0 | -0,8833332 | 0,77623004 |
| GIT1_HUMAI | GIT1 | sp\|Q9Y2X7\|GIT1_HUMAN ARF GTPase-activating protein GIT1 OS=Homo sapiens GN=GI | 160,27 | 8,15 | 84341 | 5 | 20,27 | 20,91 | 20,24 | 20,04 | 20,19 | 18,54 | 0,542 | 1,97E-01 | 0 | -0,8843746 | 0,7066096 |
| ATPD_HUMA | ATP5D | sp\|P30049\|ATPD_HUMAN ATP synthase subunit delta, mitochondrial OS=Homo sapiens | 26,122 | 5,36 | 17490 | 1 | 23,03 | 23,68 | 23,17 | 22,57 | 22,53 | 22,12 | 0,541 | 2,22E-02 | 0 | -0,8866666 | 1,65413445 |
| AL1L1_HUM | ALDH1L1 | sp\|O75891\|AL1L1_HUMAN Cytosolic 10-formyltetrahydrofolate dehydrogenase OS=Hor | 17,45 | 1,44 | 98829 | 1 | 20,48 | 20,88 | 20,50 | 18,23 | 20,19 | 20,77 | 0,540 | 3,17E-01 | 0 | -0,8899422 | 0,49831807 |
| RAB7L_HUM | RAB29 | sp\|O14966\|RAB7L_HUMAN Ras-related protein Rab-7L1 OS=Homo sapiens GN=RAB29 F | 28,829 | 10,84 | 23155 | 2 | 21,58 | 22,17 | 19,33 | 21,65 | 21,24 | 17,52 | 0,539 | 6,01E-01 | 0 | -0,8917732 | 0,22101512 |
| HBS1L_HUM | HBS1L | sp\|Q9Y450\|HBS1L_HUMAN HBS1-like protein OS=Homo sapiens GN=HBS1L PE=1 SV=1 | 44,194 | 3,65 | 75473 | 2 | 22,24 | 21,03 | 22,23 | 21,11 | 20,79 | 20,92 | 0,538 | 9,62E-02 | 0 | -0,8933328 | 1,01702991 |
| SUN2_HUMA | SUN2 | sp\|Q9UH99\|SUN2_HUMAN SUN domain-containing protein 2 OS=Homo sapiens GN=SU | 84,635 | 4,74 | 80311 | 2 | 19,26 | 19,38 | 21,62 | 18,14 | 19,09 | 20,35 | 0,538 | 4,22E-01 | 0 | -0,8933334 | 0,37480825 |
| OR1M1_HUN | OR1M1 | sp\|Q8NGA1\|OR1M1_HUMAN Olfactory receptor 1M1 OS=Homo sapiens GN=OR1M1 PE | 15,448 | 2,24 | 34840 | 1 | 25,32 | 26,86 | 26,73 | 25,06 | 25,13 | 26,03 | 0,537 | 1,99E-01 | 0 | -0,8966668 | 0,7005124 |
| SSBP_HUMA | SSBP1 | sp\|Q04837\|SSBP_HUMAN Single-stranded DNA-binding protein, mitochondrial OS=Hom | 102,788 | 27,7 | 17260 | 3 | 20,00 | 22,64 | 23,59 | 19,33 | 21,77 | 22,42 | 0,535 | 5,61E-01 | 0 | -0,903333 | 0,25084029 |
| SUCA_HUMA | SUCLG1 | sp\|P53597\|SUCA_HUMAN Succinate--CoA ligase [ADP/GDP-forming] subunit alpha, mito | 148,106 | 12,43 | 36250 | 4 | 24,16 | 24,30 | 24,82 | 23,61 | 23,19 | 23,76 | 0,533 | 2,63E-02 | 0 | -0,9066658 | 1,58069636 |
| EFTU_HUMA | TUFM | sp\|P49411\|EFTU_HUMAN Elongation factor Tu, mitochondrial OS=Homo sapiens GN=TU | 497,429 | 29,2 | 49542 | 12 | 26,54 | 26,18 | 27,13 | 25,70 | 25,27 | 26,14 | 0,531 | 7,10E-02 | 0 | -0,9133333 | 1,1489307 |
| ARL8A_HUM | ARL8A | sp\|Q96BM9\|ARL8A_HUMAN ADP-ribosylation factor-like protein 8A OS=Homo sapiens | 367,897 | 70,43 | 21416 | 12 | 24,85 | 25,28 | 24,17 | 23,53 | 24,44 | 23,58 | 0,530 | 1,04E-01 | 0 | -0,9166667 | 0,98144234 |
| PCP_HUMAN | PRCP | sp\|P42785\|PCP_HUMAN Lysosomal Pro-X carboxypeptidase OS=Homo sapiens GN=PRC | 70,297 | 5,65 | 55800 | 2 | 22,73 | 22,44 | 21,99 | 21,60 | 22,47 | 20,34 | 0,530 | 2,34E-01 | 0 | -0,9166667 | 0,63065526 |
| WASF1_HUN | WASF1 | sp\|Q92558\|WASF1_HUMAN Wiskott-Aldrich syndrome protein family member 1 OS=Hc | 15,932 | 1,61 | 61652 | 1 | 19,24 | 19,49 | 18,25 | 19,14 | 17,90 | 17,18 | 0,529 | 2,51E-01 | 0 | -0,9200001 | 0,60008585 |
| TSN7_HUMA | TSPAN7 | sp\|P41732\|TSN7_HUMAN Tetraspanin-7 OS=Homo sapiens GN=TSPAN7 PE=1 SV=2 | 19,853 | 3,61 | 27574 | 1 | 23,52 | 23,47 | 24,04 | 22,56 | 22,33 | 23,36 | 0,526 | 6,24E-02 | 0 | -0,9266669 | 1,20489109 |
| TGON2_HUN | TGOLN2 | sp\|O43493\|TGON2_HUMAN Trans-Golgi network integral membrane protein 2 OS=Hom | 28,69 | 1,88 | 51113 | 1 | 20,61 | 19,33 | 20,76 | 17,76 | 18,72 | 21,43 | 0,525 | 4,78E-01 | 0 | -0,9282907 | 0,32028902 |
| CO5_HUMAI | C5 | sp\|P01031\|CO5_HUMAN Complement C5 OS=Homo sapiens GN=C5 PE=1 SV=4 | 76,303 | 1,91 | 188305 | 3 | 22,35 | 20,64 | 23,55 | 21,36 | 19,55 | 22,84 | 0,525 | 5,05E-01 | 0 | -0,9299997 | 0,2965225 |
| FACR1_HUM | FAR1 | sp\|Q8WVX9\|FACR1_HUMAN Fatty acyl-CoA reductase 1 OS=Homo sapiens GN=FAR1 PE | 17,459 | 2,33 | 59357 | 1 | 20,47 | 19,73 | 21,08 | 18,13 | 19,76 | 20,59 | 0,524 | 3,19E-01 | 0 | -0,9320443 | 0,49587228 |
| TOM22_HUN | TOMM22 | sp\|Q9NS69\|TOM22_HUMAN Mitochondrial import receptor subunit TOM22 homolog C | 109,515 | 16,2 | 15522 | 2 | 20,13 | 21,41 | 22,81 | 19,49 | 20,19 | 21,86 | 0,522 | 4,21E-01 | 0 | -0,9366659 | 0,3757572 |
| AN13A_HUN | ANKRD13A | sp\|Q8IZ07\|AN13A_HUMAN Ankyrin repeat domain-containing protein 13A OS=Homo sa | 173,986 | 12,54 | 67619 | 6 | 24,23 | 23,57 | 22,01 | 22,65 | 22,87 | 21,48 | 0,522 | 3,00E-01 | 0 | -0,9366665 | 0,52323225 |
| DGKA_HUM | DGKA | sp\|P23743\|DGKA_HUMAN Diacylglycerol kinase alpha OS=Homo sapiens GN=DGKA PE= | 66,506 | 5,71 | 82630 | 3 | 23,20 | 23,18 | 22,27 | 22,90 | 23,08 | 19,86 | 0,522 | 4,38E-01 | 0 | -0,9366671 | 0,35841104 |
| AIFM1_HUM | AIFM1 | sp\|O95831\|AIFM1_HUMAN Apoptosis-inducing factor 1, mitochondrial OS=Homo sapie | 35,553 | 5,55 | 66901 | 2 | 20,75 | 20,87 | 21,43 | 20,18 | 20,16 | 19,90 | 0,522 | 1,48E-02 | 0 | -0,9366671 | 1,83024453 |
| ADA17_HUM | ADAM17 | sp\|P78536\|ADA17_HUMAN Disintegrin and metalloproteinase domain-containing prote | 157,987 | 13,11 | 93021 | 6 | 23,86 | 24,03 | 21,72 | 23,48 | 23,40 | 19,92 | 0,522 | 5,37E-01 | 0 | -0,9366671 | 0,26994753 |
| TRIO_HUMA | TRIO | sp\|O75962\|TRIO_HUMAN Triple functional domain protein OS=Homo sapiens GN=TRIO | 23,769 | 0,32 | 346900 | 1 | 19,10 | 19,40 | 19,55 | 17,56 | 19,57 | 18,11 | 0,522 | 2,02E-01 | 0 | -0,9370238 | 0,69465813 |
| PAIP2_HUM | PAIP2 | sp\|Q9BPZ3\|PAIP2_HUMAN Polyadenylate-binding protein-interacting protein 2 OS=Hon | 101,36 | 34,65 | 14984 | 2 | 22,14 | 22,21 | 22,81 | 20,68 | 21,28 | 22,37 | 0,520 | 1,55E-01 | 0 | -0,943332 | 0,81066712 |
| SUCB1_HUM | SUCLA2 | sp\|Q9P2R7\|SUCB1_HUMAN Succinate--CoA ligase [ADP-forming] subunit beta, mitocho | 86,559 | 6,7 | 50317 | 3 | 22,61 | 22,69 | 23,79 | 21,74 | 21,60 | 22,92 | 0,520 | 1,71E-01 | 0 | -0,9433339 | 0,76747272 |
| UBS3B_HUM | UBASH3B | sp\|Q8TF42\|UBS3B_HUMAN Ubiquitin-associated and SH3 domain-containing protein B | 67,45 | 5,08 | 72696 | 2 | 22,85 | 22,58 | 18,03 | 21,23 | 21,52 | 17,87 | 0,519 | 6,54E-01 | 0 | -0,945357 | 0,18466144 |
| HXK1_HUMA | HK1 | sp\|P19367\|HXK1_HUMAN Hexokinase-1 OS=Homo sapiens GN=HK1 PE=1 SV=3 | 422,874 | 15,38 | 102486 | 14 | 24,07 | 25,06 | 26,11 | 23,53 | 24,08 | 24,79 | 0,519 | 2,44E-01 | 0 | -0,9466661 | 0,61341147 |
| CANB1_HUN | PPP3R1 | sp\|P63098\|CANB1_HUMAN Calcineurin subunit B type 1 OS=Homo sapiens GN=PPP3R1 | 23,038 | 7,06 | 19300 | 1 | 19,52 | 20,22 | 20,55 | 19,06 | 18,20 | 20,18 | 0,518 | 2,17E-01 | 0 | -0,9481564 | 0,66344513 |
| TGFR1_HUM | TGFBR1 | sp\|P36897\|TGFR1_HUMAN TGF-beta receptor type-1 OS=Homo sapiens GN=TGFBR1 PE | 25,216 | 4,97 | 55960 | 2 | 22,44 | 22,80 | 21,82 | 21,14 | 21,61 | 21,46 | 0,518 | 4,04E-02 | 0 | -0,9500001 | 1,39318916 |
| CADM3_HUN | CADM3 | sp\|Q8N126\|CADM3_HUMAN Cell adhesion molecule 3 OS=Homo sapiens GN=CADM3 P | 11,553 | 2,26 | 43300 | 1 | 19,58 | 19,08 | 20,37 | 18,87 | 17,87 | 19,44 | 0,517 | 1,84E-01 | 0 | -0,9517511 | 0,73437452 |
| ETFA_HUMA | ETFA | sp\|P13804\|ETFA_HUMAN Electron transfer flavoprotein subunit alpha, mitochondrial O | 358,888 | 39,04 | 35080 | 10 | 23,58 | 23,70 | 25,97 | 23,13 | 22,40 | 24,85 | 0,515 | 4,19E-01 | 0 | -0,9566669 | 0,37739938 |
| SDHA_HUM | SDHA | sp\|P31040\|SDHA_HUMAN Succinate dehydrogenase [ubiquinone] flavoprotein subunit, | 292,329 | 14,76 | 72692 | 6 | 23,27 | 23,44 | 25,44 | 22,34 | 22,80 | 24,14 | 0,515 | 3,39E-01 | 0 | -0,9566676 | 0,47004874 |
| LYSC_HUMA | LYZ | sp\|P61626\|LYSC_HUMAN Lysozyme C OS=Homo sapiens GN=LYZ PE=1 SV=1 | 28,015 | 8,11 | 16537 | 1 | 22,05 | 21,34 | 23,77 | 21,87 | 19,18 | 23,23 | 0,514 | 5,28E-01 | 0 | -0,9589996 | 0,27704471 |
| CHD3_HUM | CHD3 | sp\|Q12873\|CHD3_HUMAN Chromodomain-helicase-DNA-binding protein 3 OS=Homo s | 127,017 | 3,4 | 226592 | 5 | 19,02 | 18,38 | 18,93 | 18,62 | 17,89 | 16,93 | 0,513 | 1,43E-01 | 0 | -0,9620291 | 0,84431941 |
| NEO1_HUM | NEO1 | sp\|Q92859\|NEO1_HUMAN Neogenin OS=Homo sapiens GN=NEO1 PE=1 SV=2 | 21,537 | 1,03 | 160017 | 1 | 18,24 | 19,76 | 19,87 | 17,73 | 18,68 | 18,58 | 0,513 | 1,88E-01 | 0 | -0,9623534 | 0,72615383 |
| GLI3_HUMAI | GLI3 | sp\|P10071\|GLI3_HUMAN Transcriptional activator GLI3 OS=Homo sapiens GN=GLI3 PE= | 49,225 | 2,72 | 169863 | 3 | 19,54 | 17,94 | 22,77 | 18,61 | 18,06 | 20,69 | 0,513 | 5,86E-01 | 0 | -0,9633138 | 0,23176059 |
| SMCA4_HUN | SMARCA4 | sp\|P51532\|SMCA4_HUMAN Transcription activator BRG1 OS=Homo sapiens GN=SMAR( | 24,484 | 1,28 | 184646 | 2 | 20,56 | 20,62 | 21,07 | 18,65 | 19,23 | 21,48 | 0,512 | 3,34E-01 | 0 | -0,9644407 | 0,47625041 |
| PLCB3_HUM | PLCB3 | sp\|Q01970\|PLCB3_HUMAN 1-phosphatidylinositol 4,5-bisphosphate phosphodiesterase | 238,003 | 11,26 | 138799 | 10 | 25,24 | 25,14 | 25,95 | 25,18 | 24,68 | 23,57 | 0,512 | 1,48E-01 | 0 | -0,9666665 | 0,83032459 |
| LYAR_HUMA | LYAR | sp\|Q9NX58\|LYAR_HUMAN Cell growth-regulating nucleolar protein OS=Homo sapiens G | 14,031 | 3,43 | 43615 | 1 | 19,63 | 20,18 | 22,07 | 19,08 | 18,23 | 21,66 | 0,511 | 4,87E-01 | 0 | -0,9696674 | 0,31202663 |
| ETFB_HUMA | ETFB | sp\|P38117\|ETFB_HUMAN Electron transfer flavoprotein subunit beta OS=Homo sapiens | 161,621 | 21,96 | 27844 | 6 | 23,32 | 23,01 | 25,30 | 22,47 | 22,54 | 23,71 | 0,511 | 3,04E-01 | 0 | -0,9699999 | 0,51779535 |
| SUCB2_HUM | SUCLG2 | sp\|Q96I99\|SUCB2_HUMAN Succinate--CoA ligase [GDP-forming] subunit beta, mitochor | 112,216 | 8,56 | 46511 | 4 | 23,37 | 23,21 | 25,09 | 22,30 | 22,30 | 24,15 | 0,509 | 3,22E-01 | 0 | -0,973334 | 0,49243918 |
| RAC3_HUMA | RAC3 | sp\|P60763\|RAC3_HUMAN Ras-related C3 botulinum toxin substrate 3 OS=Homo sapiens | 438,099 | 87,5 | 21379 | 13 | 24,12 | 24,10 | 21,86 | 23,37 | 22,41 | 21,37 | 0,508 | 3,60E-01 | 0 | -0,9766668 | 0,44312091 |
| WASL_HUM | WASL | sp\|O00401\|WASL_HUMAN Neural Wiskott-Aldrich syndrome protein OS=Homo sapiens | 131,502 | 16,04 | 54827 | 6 | 24,24 | 24,92 | 23,96 | 23,31 | 24,11 | 22,76 | 0,507 | 1,13E-01 | 0 | -0,9799995 | 0,94610374 |
| ACON_HUM | ACO2 | sp\|Q99798\|ACON_HUMAN Aconitate hydratase, mitochondrial OS=Homo sapiens GN=A | 366,944 | 15,51 | 85425 | 11 | 24,18 | 24,76 | 25,44 | 23,88 | 23,93 | 23,63 | 0,507 | 5,95E-02 | 0 | -0,9800008 | 1,22520519 |
| PHLA3_HUM | PHLDA3 | sp\|Q9Y5J5\|PHLA3_HUMAN Pleckstrin homology-like domain family A member 3 OS=Hor | 34,961 | 13,39 | 13891 | 2 | 22,87 | 22,78 | 21,87 | 21,94 | 21,98 | 20,63 | 0,503 | 1,44E-01 | 0 | -0,990001 | 0,84072132 |
| ZZEF1_HUM | ZZEF1 | sp\|O43149\|ZZEF1_HUMAN Zinc finger ZZ-type and EF-hand domain-containing protein | 145,426 | 1,32 | 331075 | 3 | 21,12 | 21,91 | 22,42 | 21,34 | 21,68 | 19,44 | 0,501 | 2,77E-01 | 0 | -0,9966666 | 0,55725749 |
| VASN_HUM | VASN | sp\|Q6EMK4\|VASN_HUMAN Vasorin OS=Homo sapiens GN=VASN PE=1 SV=1 | 55,222 | 2,53 | 71713 | 1 | 19,03 | 18,07 | 21,74 | 18,94 | 16,10 | 20,81 | 0,501 | 6,00E-01 | 0 | -0,9984296 | 0,22183974 |
| VDAC2_HUN | VDAC2 | sp\|P45880\|VDAC2_HUMAN Voltage-dependent anion-selective channel protein 2 OS=H | 707,047 | 57,14 | 31567 | 13 | 27,51 | 27,63 | 28,94 | 26,76 | 26,60 | 27,72 | 0,500 | 1,58E-01 | 0 | -1 | 0,80220316 |
| VDAC3_HUN | VDAC3 | sp\|Q9Y277\|VDAC3_HUMAN Voltage-dependent anion-selective channel protein 3 OS=H | 513,659 | 44,52 | 30659 | 11 | 26,23 | 26,15 | 27,26 | 25,24 | 25,30 | 26,09 | 0,499 | 8,98E-02 | 0 | -1,0033334 | 1,04678186 |
| 2A5E_HUMA | PPP2R5E | sp\|Q16537\|2A5E_HUMAN Serine/threonine-protein phosphatase 2A 56 kDa regulatory | 149,285 | 18,84 | 54699 | 8 | 23,26 | 23,31 | 22,74 | 22,47 | 22,48 | 21,35 | 0,499 | 7,38E-02 | 0 | -1,0033334 | 1,13168836 |
| OAT_HUMAI | OAT | sp\|P04181\|OAT_HUMAN Ornithine aminotransferase, mitochondrial OS=Homo sapiens | 203,438 | 11,85 | 48535 | 5 | 24,46 | 23,91 | 25,29 | 23,75 | 23,09 | 23,80 | 0,498 | 9,47E-02 | 0 | -1,0066668 | 1,02347722 |
| QBK84356.1 | | nuclear egress membrane protein [Human betaherpesvirus 5] | 11,937 | 2,26 | 42988 | 1 | 17,42 | 18,60 | 19,04 | 16,93 | 15,90 | 19,21 | 0,498 | 4,08E-01 | 0 | -1,0067743 | 0,38934972 |
| ZPI_HUMAN | SERPINA10 | sp\|Q9UK55\|ZPI_HUMAN Protein Z-dependent protease inhibitor OS=Homo sapiens GN= | 20,13 | 2,25 | 50707 | 1 | 19,88 | 18,94 | 21,97 | 17,36 | 18,83 | 21,55 | 0,494 | 5,40E-01 | 0 | -1,0166664 | 0,26758582 |
| PXDC2_HUM | PLXDC2 | sp\|Q6UX71\|PXDC2_HUMAN Plexin domain-containing protein 2 OS=Homo sapiens GN= | 21,041 | 1,89 | 59583 | 1 | 20,14 | 19,67 | 21,00 | 19,48 | 18,27 | 19,98 | 0,491 | 1,84E-01 | 0 | -1,0266666 | 0,73560375 |
| EPS8_HUMA | EPS8 | sp\|Q12929\|EPS8_HUMAN Epidermal growth factor receptor kinase substrate 8 OS=Hon | 515,999 | 17,88 | 91882 | 11 | 25,80 | 25,47 | 24,07 | 25,02 | 24,70 | 22,52 | 0,489 | 3,37E-01 | 0 | -1,0333322 | 0,47257657 |
| STML2_HUM | STOML2 | sp\|Q9UJZ1\|STML2_HUMAN Stomatin-like protein 2, mitochondrial OS=Homo sapiens GI | 333,546 | 25 | 38534 | 5 | 22,74 | 22,98 | 25,13 | 21,69 | 22,10 | 23,95 | 0,487 | 3,71E-01 | 0 | -1,0366656 | 0,43054842 |
| KLC2_HUMA | KLC2 | sp\|Q9H0B6\|KLC2_HUMAN Kinesin light chain 2 OS=Homo sapiens GN=KLC2 PE=1 SV=1 | 248,122 | 17,68 | 68935 | 7 | 17,84 | 19,11 | 21,36 | 17,98 | 18,12 | 19,10 | 0,487 | 3,95E-01 | 0 | -1,0368633 | 0,40389502 |
| THIM_HUMA | ACAA2 | sp\|P42765\|THIM_HUMAN 3-ketoacyl-CoA thiolase, mitochondrial OS=Homo sapiens GN | 14,169 | 2,27 | 41924 | 1 | 18,29 | 20,75 | 22,81 | 17,61 | 19,29 | 21,84 | 0,487 | 5,94E-01 | 0 | -1,0371202 | 0,2260426 |
| GPC1_HUMA | GPC1 | sp\|P35052\|GPC1_HUMAN Glypican-1 OS=Homo sapiens GN=GPC1 PE=1 SV=2 | 20,224 | 4,48 | 61680 | 2 | 22,52 | 22,64 | 21,25 | 21,77 | 21,84 | 19,67 | 0,485 | 2,82E-01 | 0 | -1,0433331 | 0,55009621 |
| ENPLL_HUM | HSP90B2P | sp\|Q58FF3\|ENPLL_HUMAN Putative endoplasmin-like protein OS=Homo sapiens GN=HS | 131,877 | 13,03 | 45859 | 7 | 22,23 | 21,82 | 23,52 | 22,33 | 22,55 | 19,53 | 0,482 | 3,92E-01 | 0 | -1,0521679 | 0,4066105 |
| CU129_HUM | LINC00479 | sp\|Q96M42\|CU129_HUMAN Putative uncharacterized protein encoded by LINC00479 O | 17,263 | 4,93 | 15208 | 1 | 21,92 | 21,79 | 28,44 | 19,42 | 21,41 | 28,16 | 0,481 | 7,74E-01 | 0 | -1,0548878 | 0,11110062 |
| DCTN6_HUN | DCTN6 | sp\|O00399\|DCTN6_HUMAN Dynactin subunit 6 OS=Homo sapiens GN=DCTN6 PE=1 SV= | 47,459 | 7,37 | 20747 | 1 | 17,09 | 20,54 | 18,83 | 17,60 | 17,13 | 18,57 | 0,481 | 3,84E-01 | 0 | -1,0557613 | 0,41530272 |
| C99L2_HUM | CD99L2 | sp\|Q8TCZ2\|C99L2_HUMAN CD99 antigen-like protein 2 OS=Homo sapiens GN=CD99L2 F | 27,088 | 5,73 | 27986 | 1 | 17,89 | 20,95 | 20,97 | 19,07 | 16,84 | 20,73 | 0,481 | 5,26E-01 | 0 | -1,056956 | 0,279215 |
| RPP30_HUM | RPP30 | sp\|P78346\|RPP30_HUMAN Ribonuclease P protein subunit p30 OS=Homo sapiens GN=F | 60,022 | 3,36 | 29321 | 1 | 20,27 | 20,35 | 21,34 | 15,62 | 21,09 | 22,07 | 0,480 | 6,30E-01 | 0 | -1,0600004 | 0,20058006 |
| IASPP_HUM | PPP1R13L | sp\|Q8WUF5\|IASPP_HUMAN RelA-associated inhibitor OS=Homo sapiens GN=PPP1R13L | 57,322 | 2,9 | 89091 | 1 | 17,15 | 19,53 | 19,98 | 17,48 | 17,92 | 18,08 | 0,479 | 3,02E-01 | 0 | -1,06106 | 0,51942628 |
| PKP2_HUMA | PKP2 | sp\|Q99959\|PKP2_HUMAN Plakophilin-2 OS=Homo sapiens GN=PKP2 PE=1 SV=2 | 18,98 | 1,25 | 97415 | 1 | 21,43 | 21,42 | 19,02 | 21,28 | 20,95 | 16,45 | 0,479 | 5,77E-01 | 0 | -1,0614948 | 0,23872542 |
| IFT52_HUMA | IFT52 | sp\|Q9Y366\|IFT52_HUMAN Intraflagellar transport protein 52 homolog OS=Homo sapien | 149,08 | 14,19 | 49706 | 4 | 21,42 | 20,43 | 24,73 | 19,87 | 19,77 | 23,75 | 0,479 | 5,95E-01 | 0 | -1,0633329 | 0,22516456 |
| LONM_HUM | LONP1 | sp\|P36776\|LONM_HUMAN Lon protease homolog, mitochondrial OS=Homo sapiens GN | 135,889 | 9,7 | 106489 | 6 | 22,72 | 23,08 | 24,40 | 21,83 | 22,01 | 23,15 | 0,476 | 1,79E-01 | 0 | -1,0699997 | 0,74785949 |
| CH10_HUMA | HSPE1 | sp\|P61604\|CH10_HUMAN 10 kDa heat shock protein, mitochondrial OS=Homo sapiens | 147,418 | 47,06 | 10932 | 5 | 25,46 | 25,56 | 25,20 | 24,36 | 25,29 | 23,31 | 0,471 | 1,35E-01 | 0 | -1,0866661 | 0,86892075 |
| RBM12_HUN | RBM12 | sp\|Q9NTZ6\|RBM12_HUMAN RNA-binding protein 12 OS=Homo sapiens GN=RBM12 PE= | 25,544 | 1,82 | 97395 | 2 | 21,19 | 21,26 | 20,76 | 21,25 | 21,68 | 17,02 | 0,471 | 5,08E-01 | 0 | -1,087176 | 0,29436846 |
| M2OM_HUN | SLC25A11 | sp\|Q02978\|M2OM_HUMAN Mitochondrial 2-oxoglutarate/malate carrier protein OS=H | 137,737 | 11,78 | 34062 | 3 | 20,13 | 20,98 | 23,09 | 19,00 | 19,84 | 22,08 | 0,469 | 4,39E-01 | 0 | -1,0929076 | 0,35777481 |
| IDHP_HUMA | IDH2 | sp\|P48735\|IDHP_HUMAN Isocitrate dehydrogenase [NADP], mitochondrial OS=Homo sa | 583,076 | 38,94 | 50909 | 16 | 25,54 | 25,74 | 27,40 | 24,63 | 24,69 | 26,06 | 0,467 | 2,17E-01 | 0 | -1,1000004 | 0,66276927 |
| 1C06_HUMA | HLA-C | sp\|Q29963\|1C06_HUMAN HLA class I histocompatibility antigen, Cw-6 alpha chain OS=H | 94,653 | 16,67 | 40969 | 4 | 23,10 | 22,35 | 19,73 | 21,75 | 22,32 | 17,77 | 0,462 | 5,61E-01 | 0 | -1,1134618 | 0,25074677 |
| SPAT5_HUM | SPATA5 | sp\|Q8NB90\|SPAT5_HUMAN Spermatogenesis-associated protein 5 OS=Homo sapiens G | 103,302 | 3,47 | 97904 | 2 | 18,94 | 19,29 | 19,68 | 17,17 | 17,87 | 19,51 | 0,461 | 1,98E-01 | 0 | -1,1179841 | 0,7023873 |
| COX5A_HUN | COX5A | sp\|P20674\|COX5A_HUMAN Cytochrome c oxidase subunit 5A, mitochondrial OS=Homo | 25,077 | 24,67 | 16762 | 2 | 21,62 | 21,82 | 23,30 | 21,10 | 20,34 | 21,93 | 0,459 | 1,84E-01 | 0 | -1,123333 | 0,73434642 |
| DSCR3_HUM | DSCR3 | sp\|O14972\|DSCR3_HUMAN Down syndrome critical region protein 3 OS=Homo sapiens | 28,025 | 4,04 | 33010 | 1 | 19,88 | 19,63 | 19,93 | 17,00 | 19,06 | 20,00 | 0,458 | 2,75E-01 | 0 | -1,1267859 | 0,56128422 |
| NAV1_HUMA | NAV1 | sp\|Q8NEY1\|NAV1_HUMAN Neuron navigator 1 OS=Homo sapiens GN=NAV1 PE=1 SV=2 | 14,319 | 0,64 | 202472 | 1 | 19,36 | 19,97 | 21,22 | 17,42 | 18,35 | 21,38 | 0,456 | 4,38E-01 | 0 | -1,1331336 | 0,35885486 |
| DIC_HUMAN | SLC25A10 | sp\|Q9UBX3\|DIC_HUMAN Mitochondrial dicarboxylate carrier OS=Homo sapiens GN=SL( | 139,772 | 18,47 | 31282 | 5 | 23,79 | 23,68 | 25,46 | 21,99 | 22,75 | 24,78 | 0,455 | 3,24E-01 | 0 | -1,1366666 | 0,48890425 |
| ANLN_HUM | ANLN | sp\|Q9NQW6\|ANLN_HUMAN Anillin OS=Homo sapiens GN=ANLN PE=1 SV=2 | 132,621 | 4,18 | 124199 | 4 | 24,31 | 24,02 | 23,40 | 23,12 | 23,25 | 21,92 | 0,452 | 8,41E-02 | 0 | -1,1466662 | 1,07533362 |
| DSG2_HUM | DSG2 | sp\|Q14126\|DSG2_HUMAN Desmoglein-2 OS=Homo sapiens GN=DSG2 PE=1 SV=2 | 28,268 | 1,61 | 122294 | 2 | 22,59 | 23,44 | 21,26 | 20,99 | 21,49 | 21,36 | 0,451 | 1,52E-01 | 0 | -1,1500003 | 0,81687101 |
| PTPRD_HUM | PTPRD | sp\|P23468\|PTPRD_HUMAN Receptor-type tyrosine-protein phosphatase delta OS=Hom | 43,246 | 2,04 | 214760 | 3 | 21,23 | 20,74 | 18,52 | 20,80 | 19,34 | 16,88 | 0,448 | 4,59E-01 | 0 | -1,1592019 | 0,33835378 |
| CO6A2_HUN | COL6A2 | sp\|P12110\|CO6A2_HUMAN Collagen alpha-2(VI) chain OS=Homo sapiens GN=COL6A2 P | 141,85 | 7,07 | 108579 | 6 | 22,57 | 20,40 | 24,14 | 21,15 | 19,22 | 23,25 | 0,446 | 5,05E-01 | 0 | -1,1633333 | 0,29664532 |
| MPCP_HUM | SLC25A3 | sp\|Q00325\|MPCP_HUMAN Phosphate carrier protein, mitochondrial OS=Homo sapiens | 353,391 | 33,7 | 40095 | 10 | 25,95 | 26,56 | 27,56 | 24,97 | 25,42 | 26,19 | 0,446 | 1,20E-01 | 0 | -1,1633333 | 0,92247268 |
| LTOR5_HUM | LAMTOR5 | sp\|O43504\|LTOR5_HUMAN Ragulator complex protein LAMTOR5 OS=Homo sapiens GN | 30,238 | 21,98 | 9614 | 1 | 21,21 | 21,50 | 19,18 | 18,13 | 21,07 | 19,20 | 0,446 | 3,60E-01 | 0 | -1,1651262 | 0,44408022 |
| CADH2_HUN | CDH2 | sp\|P19022\|CADH2_HUMAN Cadherin-2 OS=Homo sapiens GN=CDH2 PE=1 SV=4 | 172,507 | 11,26 | 99809 | 6 | 25,17 | 24,97 | 22,05 | 24,09 | 24,20 | 20,40 | 0,445 | 5,08E-01 | 0 | -1,166666 | 0,29451986 |
| PRDX3_HUM | PRDX3 | sp\|P30048\|PRDX3_HUMAN Thioredoxin-dependent peroxide reductase, mitochondrial | 69,446 | 10,94 | 27693 | 3 | 23,96 | 23,93 | 25,41 | 22,31 | 23,28 | 24,21 | 0,445 | 1,87E-01 | 0 | -1,1666667 | 0,72728892 |
| HYAS2_HUM | HAS2 | sp\|Q92819\|HYAS2_HUMAN Hyaluronan synthase 2 OS=Homo sapiens GN=HAS2 PE=2 S | 87,818 | 16,12 | 63566 | 7 | 25,10 | 24,63 | 21,84 | 24,07 | 24,06 | 19,92 | 0,443 | 5,32E-01 | 0 | -1,1733335 | 0,27437566 |
| LTOR2_HUM | LAMTOR2 | sp\|Q9Y2Q5\|LTOR2_HUMAN Ragulator complex protein LAMTOR2 OS=Homo sapiens GN | 112,294 | 31,2 | 13508 | 3 | 22,48 | 21,93 | 19,90 | 20,10 | 21,40 | 19,29 | 0,443 | 3,04E-01 | 0 | -1,1734486 | 0,51782255 |
| FGRL1_HUM | FGFRL1 | sp\|Q8N441\|FGRL1_HUMAN Fibroblast growth factor receptor-like 1 OS=Homo sapiens | 28,884 | 4,37 | 54537 | 2 | 21,06 | 21,93 | 20,60 | 20,78 | 19,37 | 19,89 | 0,440 | 1,05E-01 | 0 | -1,1833331 | 0,97793607 |
| IFT57_HUMA | IFT57 | sp\|Q9NWB7\|IFT57_HUMAN Intraflagellar transport protein 57 homolog OS=Homo sapie | 49,837 | 6,76 | 49108 | 3 | 21,73 | 20,99 | 23,81 | 20,75 | 20,72 | 21,51 | 0,440 | 2,51E-01 | 0 | -1,1833331 | 0,60000228 |

| | | | | | | | | | | | | | | | | |
|---|---|---|---|---|---|---|---|---|---|---|---|---|---|---|---|---|
| NNTM_HUM | NNT | sp\|Q13423\|NNTM_HUMAN NAD(P) transhydrogenase, mitochondrial OS=Homo sapiens | 148,087 | 8,1 | 113896 | 6 | 21,73 | 22,77 | 23,68 | 22,39 | 21,62 | 20,62 | **0,440** | 1,95E-01 | **0** | -1,1833331 | 0,70959896 |
| WNT5A_HUI | WNT5A | sp\|P41221\|WNT5A_HUMAN Protein Wnt-5a OS=Homo sapiens GN=WNT5A PE=1 SV=2 | 113,025 | 11,05 | 42339 | 3 | 22,02 | 21,24 | 20,73 | 20,79 | 21,26 | 18,38 | **0,439** | 2,87E-01 | **0** | -1,1868521 | 0,54158133 |
| SCOT1_HUM | OXCT1 | sp\|P55809\|SCOT1_HUMAN Succinyl-CoA:3-ketoacid coenzyme A transferase 1, mitocho | 211,385 | 14,42 | 56158 | 8 | 23,69 | 23,55 | 25,58 | 22,60 | 22,66 | 23,99 | **0,438** | 2,09E-01 | **0** | -1,1899999 | 0,6789509 |
| ATAD1_HUM | ATAD1 | sp\|Q8NBU5\|ATAD1_HUMAN ATPase family AAA domain-containing protein 1 OS=Homo | 68,591 | 4,43 | 40744 | 1 | 19,55 | 18,73 | 21,14 | 17,60 | 17,60 | 20,65 | **0,437** | 3,90E-01 | **0** | -1,19266 | 0,40882587 |
| FR8P1_HUM. | FRMD8P1 | sp\|Q9BZ68\|FR8P1_HUMAN Putative FERM domain-containing protein FRMD8P1 OS=Ho | 12,64 | 2,17 | 41136 | 1 | 21,99 | 21,92 | 18,61 | 18,37 | 21,36 | 19,21 | **0,436** | 4,49E-01 | **0** | -1,1961295 | 0,34754093 |
| SFXN3_HUM | SFXN3 | sp\|Q9BWM7\|SFXN3_HUMAN Sideroflexin-3 OS=Homo sapiens GN=SFXN3 PE=1 SV=3 | 173,274 | 17,13 | 35503 | 4 | 22,81 | 23,17 | 24,26 | 21,67 | 21,66 | 23,32 | **0,436** | 1,64E-01 | **0** | -1,1966667 | 0,78522292 |
| DLDH_HUM/ | DLD | sp\|P09622\|DLDH_HUMAN Dihydrolipoyl dehydrogenase, mitochondrial OS=Homo sapie | 284,015 | 21,22 | 54177 | 9 | 23,03 | 23,21 | 26,15 | 21,64 | 22,30 | 24,85 | **0,435** | 4,42E-01 | **0** | -1,2000001 | 0,3546782 |
| DHE3_HUM/ | GLUD1 | sp\|P00367\|DHE3_HUMAN Glutamate dehydrogenase 1, mitochondrial OS=Homo sapier | 884,613 | 43,37 | 61398 | 27 | 26,38 | 26,46 | 28,39 | 25,18 | 25,52 | 26,92 | **0,434** | 2,28E-01 | **0** | -1,2033323 | 0,64232291 |
| C1QBP_HUM | C1QBP | sp\|Q07021\|C1QBP_HUMAN Complement component 1 Q subcomponent-binding prote | 69,181 | 10,64 | 31362 | 1 | 18,72 | 21,68 | 20,48 | 19,09 | 19,36 | 18,81 | **0,433** | 2,40E-01 | **0** | -1,2061761 | 0,62032165 |
| CMTD1_HUI | COMTD1 | sp\|Q86VU5\|CMTD1_HUMAN Catechol O-methyltransferase domain-containing protein | 52,897 | 9,92 | 28809 | 1 | 16,79 | 18,38 | 19,75 | 16,46 | 16,70 | 18,13 | **0,433** | 2,94E-01 | **0** | -1,207866 | 0,53146187 |
| COEA1_HUN | COL14A1 | sp\|Q05707\|COEA1_HUMAN Collagen alpha-1(XIV) chain OS=Homo sapiens GN=COL14A | 22,534 | 1,28 | 193515 | 2 | 21,79 | 21,84 | 20,75 | 21,35 | 20,93 | 18,47 | **0,432** | 2,79E-01 | **0** | -1,2104232 | 0,55504241 |
| PKN2_HUM/ | PKN2 | sp\|Q16513\|PKN2_HUMAN Serine/threonine-protein kinase N2 OS=Homo sapiens GN=P | 17,579 | 0,91 | 112035 | 1 | 20,28 | 20,48 | 20,10 | 19,75 | 20,26 | 17,22 | **0,432** | 2,70E-01 | **0** | -1,2120228 | 0,5685879 |
| PSMG3_HUI | PSMG3 | sp\|Q9BT73\|PSMG3_HUMAN Proteasome assembly chaperone 3 OS=Homo sapiens GN= | 19,222 | 10,66 | 13104 | 1 | 20,69 | 20,03 | 22,27 | 20,54 | 20,50 | 18,31 | **0,431** | 2,88E-01 | **0** | -1,2140293 | 0,53989108 |
| SC6A8_HUM | SLC6A8 | sp\|P48029\|SC6A8_HUMAN Sodium- and chloride-dependent creatine transporter 1 OS= | 34,263 | 4,41 | 70523 | 2 | 22,45 | 22,60 | 20,24 | 22,05 | 21,65 | 17,94 | **0,430** | 4,67E-01 | **0** | -1,2169272 | 0,33085953 |
| AATM_HUM. | GOT2 | sp\|P00505\|AATM_HUMAN Aspartate aminotransferase, mitochondrial OS=Homo sapier | 214,553 | 23,26 | 47518 | 8 | 23,92 | 23,96 | 26,22 | 22,68 | 23,01 | 24,75 | **0,429** | 2,87E-01 | **0** | -1,2199993 | 0,54151685 |
| IFT81_HUM/ | IFT81 | sp\|Q8WYA0\|IFT81_HUMAN Intraflagellar transport protein 81 homolog OS=Homo sapie | 284,638 | 13,91 | 79746 | 10 | 22,70 | 21,11 | 25,58 | 20,76 | 20,28 | 24,68 | **0,428** | 5,57E-01 | **0** | -1,2233334 | 0,25416469 |
| TENS3_HUM | TNS3 | sp\|Q68CZ2\|TENS3_HUMAN Tensin-3 OS=Homo sapiens GN=TNS3 PE=1 SV=2 | 107,181 | 1,38 | 155266 | 2 | 22,00 | 22,04 | 21,68 | 21,69 | 21,88 | 18,48 | **0,428** | 3,32E-01 | **0** | -1,2239908 | 0,4790332 |
| BMPR2_HUN | BMPR2 | sp\|Q13873\|BMPR2_HUMAN Bone morphogenetic protein receptor type-2 OS=Homo sa | 19,991 | 1,83 | 115201 | 1 | 20,96 | 20,79 | 23,02 | 20,60 | 20,21 | 20,28 | **0,427** | 1,67E-01 | **0** | -1,2266668 | 0,77814501 |
| NDE1_HUM/ | NDE1 | sp\|Q9NXR1\|NDE1_HUMAN Nuclear distribution protein nudE homolog 1 OS=Homo sapi | 50,079 | 3,47 | 38808 | 1 | 19,76 | 19,90 | 20,42 | 16,47 | 19,72 | 20,20 | **0,426** | 3,59E-01 | **0** | -1,2300002 | 0,44464022 |
| DIP2A_HUM/ | DIP2A | sp\|Q14689\|DIP2A_HUMAN Disco-interacting protein 2 homolog A OS=Homo sapiens GN | 93,858 | 4,07 | 170369 | 6 | 21,19 | 20,72 | 21,75 | 18,48 | 20,78 | 20,68 | **0,424** | 1,99E-01 | **0** | -1,2392133 | 0,70046595 |
| KIFA3_HUM/ | KIFAP3 | sp\|Q92845\|KIFA3_HUMAN Kinesin-associated protein 3 OS=Homo sapiens GN=KIFAP3 F | 193,827 | 9,72 | 91205 | 6 | 21,71 | 21,38 | 24,68 | 19,64 | 21,06 | 23,29 | **0,418** | 4,46E-01 | **0** | -1,2599996 | 0,35040679 |
| RCN1_HUM/ | RCN1 | sp\|Q15293\|RCN1_HUMAN Reticulocalbin-1 OS=Homo sapiens GN=RCN1 PE=1 SV=1 | 39,029 | 7,85 | 38890 | 2 | 19,17 | 18,53 | 22,25 | 19,23 | 15,63 | 21,26 | **0,413** | 5,60E-01 | **0** | -1,2754571 | 0,25201886 |
| RAPH1_HUM | RAPH1 | sp\|Q70E73\|RAPH1_HUMAN Ras-associated and pleckstrin homology domains-containin | 15,696 | 0,72 | 135256 | 1 | 21,62 | 21,04 | 19,88 | 18,67 | 20,82 | 19,21 | **0,412** | 1,95E-01 | **0** | -1,2805119 | 0,70946124 |
| MYRF_HUM/ | MYRF | sp\|Q9Y2G1\|MYRF_HUMAN Myelin regulatory factor OS=Homo sapiens GN=MYRF PE=1 | 172,409 | 4,34 | 124397 | 4 | 23,54 | 23,28 | 20,57 | 23,23 | 22,30 | 17,99 | **0,409** | 5,29E-01 | **0** | -1,2901796 | 0,27671875 |
| PLSI_HUMAN | PLS1 | sp\|Q14651\|PLSI_HUMAN Plastin-1 OS=Homo sapiens GN=PLS1 PE=1 SV=2 | 148,174 | 7,63 | 70253 | 4 | 20,90 | 20,31 | 20,23 | 18,41 | 20,16 | 18,99 | **0,407** | 8,06E-02 | **0** | -1,2953097 | 1,09360598 |
| RFFL_HUMAI | RFFL | sp\|Q8WZ73\|RFFL_HUMAN E3 ubiquitin-protein ligase rififylin OS=Homo sapiens GN=RFI | 99,197 | 14,05 | 40514 | 4 | 25,37 | 24,82 | 21,96 | 24,63 | 24,10 | 19,53 | **0,407** | 5,39E-01 | **0** | -1,2966665 | 0,26827273 |
| ABCBB_HUM | ABCB11 | sp\|O95342\|ABCBB_HUMAN Bile salt export pump OS=Homo sapiens GN=ABCB11 PE=1 S | 27,735 | 1,29 | 146407 | 1 | 21,08 | 21,71 | 21,61 | 18,12 | 20,82 | 21,57 | **0,407** | 2,91E-01 | **0** | -1,2973633 | 0,5366664 |
| ANR50_HUM | ANKRD50 | sp\|Q9ULJ7\|ANR50_HUMAN Ankyrin repeat domain-containing protein 50 OS=Homo sap | 459,215 | 13,37 | 155859 | 13 | 24,81 | 24,69 | 22,83 | 23,21 | 23,37 | 21,82 | **0,403** | 1,80E-01 | **0** | -1,3100001 | 0,74396028 |
| DCNL3_HUM | DCUN1D3 | sp\|Q8IWE4\|DCNL3_HUMAN DCN1-like protein 3 OS=Homo sapiens GN=DCUN1D3 PE=1 | 86,701 | 15,13 | 34291 | 3 | 22,41 | 22,27 | 20,93 | 21,65 | 22,30 | 17,71 | **0,401** | 4,32E-01 | **0** | -1,318107 | 0,36427128 |
| TNS2_HUMA | TNS2 | sp\|Q63HR2\|TNS2_HUMAN Tensin-2 OS=Homo sapiens GN=TNS2 PE=1 SV=2 | 244,933 | 6,17 | 152580 | 8 | 25,65 | 24,36 | 21,84 | 24,08 | 23,87 | 19,93 | **0,400** | 4,92E-01 | **0** | -1,3233331 | 0,30772934 |
| ANGL2_HUN | ANGPTL2 | sp\|Q9UKU9\|ANGL2_HUMAN Angiopoietin-related protein 2 OS=Homo sapiens GN=ANG | 49,325 | 3,45 | 57104 | 1 | 20,71 | 19,79 | 18,23 | 19,91 | 17,96 | 16,89 | **0,400** | 3,11E-01 | **0** | -1,3233827 | 0,50696903 |
| ZHANG_HUN | CREBZF | sp\|Q9NS37\|ZHANG_HUMAN CREB/ATF bZIP transcription factor OS=Homo sapiens GN= | 15,835 | 1,98 | 37134 | 1 | 22,41 | 22,20 | 22,70 | 22,12 | 22,52 | 18,68 | **0,398** | 3,39E-01 | **0** | -1,3285586 | 0,46920668 |
| RASEF_HUM. | RASEF | sp\|Q8IZ41\|RASEF_HUMAN Ras and EF-hand domain-containing protein OS=Homo sapie | 61,906 | 1,76 | 82879 | 1 | 21,62 | 21,55 | 22,02 | 21,75 | 21,83 | 17,61 | **0,397** | 3,95E-01 | **0** | -1,3316816 | 0,40315173 |
| BIG3_HUMAI | ARFGEF3 | sp\|Q5TH69\|BIG3_HUMAN Brefeldin A-inhibited guanine nucleotide-exchange protein 3 | 168,071 | 3,45 | 240652 | 5 | 22,54 | 22,29 | 17,69 | 19,52 | 20,05 | 18,94 | **0,396** | 4,53E-01 | **0** | -1,3360551 | 0,34393335 |
| ARG39_HUM | ARHGEF39 | sp\|Q8N4T4\|ARG39_HUMAN Rho guanine nucleotide exchange factor 39 OS=Homo sapi | 12,669 | 6,27 | 38295 | 1 | 22,33 | 21,95 | 20,98 | 22,05 | 21,55 | 17,56 | **0,387** | 4,07E-01 | **0** | -1,3682035 | 0,39044273 |
| TIM44_HUM | TIMM44 | sp\|O43615\|TIM44_HUMAN Mitochondrial import inner membrane translocase subunit | 45,978 | 3,32 | 51356 | 1 | 21,45 | 21,39 | 22,89 | 18,70 | 21,23 | 21,69 | **0,387** | 2,62E-01 | **0** | -1,3699557 | 0,58115645 |
| PREP_HUMA | PITRM1 | sp\|Q5JRX3\|PREP_HUMAN Presequence protease, mitochondrial OS=Homo sapiens GN= | 76,496 | 3,95 | 117413 | 4 | 22,56 | 21,95 | 23,36 | 20,75 | 21,29 | 21,72 | **0,387** | 5,06E-02 | **0** | -1,3700002 | 1,29613383 |
| FOXP1_HUM | FOXP1 | sp\|Q9H334\|FOXP1_HUMAN Forkhead box protein P1 OS=Homo sapiens GN=FOXP1 PE= | 11,018 | 1,62 | 75317 | 1 | 19,85 | 18,79 | 19,23 | 16,89 | 18,18 | 18,63 | **0,382** | 8,34E-02 | **0** | -1,3887272 | 1,07907983 |
| IFT46_HUM/ | IFT46 | sp\|Q9NQC8\|IFT46_HUMAN Intraflagellar transport protein 46 homolog OS=Homo sapie | 155,542 | 23,68 | 34286 | 4 | 19,46 | 19,27 | 25,32 | 17,95 | 17,65 | 24,28 | **0,382** | 6,61E-01 | **0** | -1,3899994 | 0,18011551 |
| DNJB2_HUM | DNAJB2 | sp\|P25686\|DNJB2_HUMAN DnaJ homolog subfamily B member 2 OS=Homo sapiens GN= | 69,634 | 6,79 | 35580 | 2 | 22,57 | 22,62 | 22,20 | 22,34 | 21,86 | 18,93 | **0,374** | 2,57E-01 | **0** | -1,418841 | 0,59087769 |
| GLSK_HUMA | GLS | sp\|O94925\|GLSK_HUMAN Glutaminase kidney isoform, mitochondrial OS=Homo sapien | 266,258 | 20,03 | 73461 | 8 | 23,37 | 22,67 | 25,05 | 22,10 | 21,11 | 23,61 | **0,373** | 2,33E-01 | **0** | -1,4233329 | 0,63288771 |
| ZCHC8_HUM | ZCCHC8 | sp\|Q6NZY4\|ZCHC8_HUMAN Zinc finger CCHC domain-containing protein 8 OS=Homo sa | 13,708 | 1,7 | 78577 | 1 | 18,33 | 17,94 | 18,90 | 16,27 | 15,70 | 18,92 | **0,372** | 2,39E-01 | **0** | -1,4258019 | 0,62203315 |
| ARLY_HUMA | ASL | sp\|P04424\|ARLY_HUMAN Argininosuccinate lyase OS=Homo sapiens GN=ASL PE=1 SV=4 | 35,057 | 2,37 | 51658 | 1 | 20,45 | 18,72 | 20,19 | 17,36 | 20,71 | 16,98 | **0,370** | 3,32E-01 | **0** | -1,4353301 | 0,47838662 |
| ACADS_HUM | ACADS | sp\|P16219\|ACADS_HUMAN Short-chain specific acyl-CoA dehydrogenase, mitochondria | 72,122 | 5,58 | 44297 | 2 | 20,31 | 21,00 | 23,06 | 17,05 | 20,76 | 22,17 | **0,362** | 4,47E-01 | **0** | -1,4643523 | 0,35003728 |
| PPAL_HUMA | ACP2 | sp\|P11117\|PPAL_HUMAN Lysosomal acid phosphatase OS=Homo sapiens GN=ACP2 PE= | 71,552 | 12,53 | 48344 | 4 | 24,79 | 24,46 | 22,34 | 23,83 | 24,34 | 19,02 | **0,362** | 4,75E-01 | **0** | -1,4660498 | 0,32363025 |
| QCR1_HUM/ | UQCRC1 | sp\|P31930\|QCR1_HUMAN Cytochrome b-c1 complex subunit 1, mitochondrial OS=Hom | 47,101 | 5,62 | 52646 | 2 | 21,25 | 21,28 | 23,85 | 20,42 | 18,50 | 23,02 | **0,358** | 3,99E-01 | **0** | -1,4808858 | 0,39954451 |
| TIM50_HUM | TIMM50 | sp\|Q3ZCQ8\|TIM50_HUMAN Mitochondrial import inner membrane translocase subunit | 85,4 | 7,93 | 39646 | 3 | 22,79 | 23,20 | 23,04 | 21,78 | 22,46 | 20,34 | **0,358** | 8,01E-02 | **0** | -1,4833342 | 1,09619986 |
| SNG1_HUM/ | SYNGR1 | sp\|O43759\|SNG1_HUMAN Synaptogyrin-1 OS=Homo sapiens GN=SYNGR1 PE=1 SV=2 | 35,75 | 5,15 | 25456 | 1 | 22,09 | 21,85 | 21,52 | 18,16 | 21,45 | 21,36 | **0,355** | 2,43E-01 | **0** | -1,4952475 | 0,61410485 |
| IF2B3_HUM/ | IGF2BP3 | sp\|O00425\|IF2B3_HUMAN Insulin-like growth factor 2 mRNA-binding protein 3 OS=Hon | 52,166 | 2,42 | 63705 | 1 | 18,67 | 18,50 | 22,11 | 16,63 | 16,08 | 22,05 | **0,353** | 5,39E-01 | **0** | -1,5025361 | 0,26856264 |
| CEP41_HUM. | CEP41 | sp\|Q9BYV8\|CEP41_HUMAN Centrosomal protein of 41 kDa OS=Homo sapiens GN=CEP4 | 55,241 | 3,22 | 41368 | 1 | 19,43 | 19,55 | 19,88 | 17,08 | 19,00 | 18,24 | **0,350** | 5,79E-02 | **0** | -1,5144857 | 1,23696835 |
| ODP2_HUM/ | DLAT | sp\|P10515\|ODP2_HUMAN Dihydrolipoyllysine-residue acetyltransferase component of | 94,76 | 6,34 | 68997 | 4 | 20,40 | 21,71 | 24,58 | 18,93 | 20,03 | 23,11 | **0,344** | 4,30E-01 | **0** | -1,539999 | 0,36617781 |
| TOM40_HUI | TOMM40 | sp\|O96008\|TOM40_HUMAN Mitochondrial import receptor subunit TOM40 homolog O | 170,184 | 8,03 | 37893 | 3 | 22,59 | 22,53 | 23,92 | 19,69 | 21,73 | 22,98 | **0,342** | 2,19E-01 | **0** | -1,5466671 | 0,66043537 |
| S29A2_HUM. | SLC29A2 | sp\|Q14542\|S29A2_HUMAN Equilibrative nucleoside transporter 2 OS=Homo sapiens GN | 43,888 | 3,29 | 50113 | 1 | 20,93 | 21,20 | 20,31 | 18,97 | 20,82 | 17,81 | **0,327** | 1,53E-01 | **0** | -1,613334 | 0,81609483 |
| MFN2_HUM. | MFN2 | sp\|O95140\|MFN2_HUMAN Mitofusin-2 OS=Homo sapiens GN=MFN2 PE=1 SV=3 | 20,138 | 2,25 | 86402 | 2 | 18,64 | 19,92 | 21,44 | 18,02 | 17,21 | 19,93 | **0,326** | 2,30E-01 | **0** | -1,6150087 | 0,63789345 |
| MERL_HUM/ | NF2 | sp\|P35240\|MERL_HUMAN Merlin OS=Homo sapiens GN=NF2 PE=1 SV=1 | 14,844 | 1,51 | 69690 | 1 | 20,63 | 20,66 | 19,06 | 16,85 | 19,80 | 18,75 | **0,318** | 1,78E-01 | **0** | -1,6517359 | 0,74873447 |
| PPM1B_HUI | PPM1B | sp\|O75688\|PPM1B_HUMAN Protein phosphatase 1B OS=Homo sapiens GN=PPM1B PE= | 80,437 | 7,31 | 52643 | 3 | 21,11 | 21,18 | 18,19 | 21,14 | 16,37 | 17,99 | **0,316** | 3,87E-01 | **0** | -1,6609713 | 0,41261689 |
| NUMBL_HUI | NUMBL | sp\|Q9Y6R0\|NUMBL_HUMAN Numb-like protein OS=Homo sapiens GN=NUMBL PE=1 SV | 15,387 | 1,81 | 64891 | 1 | 21,41 | 21,23 | 21,95 | 18,27 | 20,49 | 20,74 | **0,309** | 1,05E-01 | **0** | -1,6951974 | 0,97730833 |
| HCDH_HUM/ | HADH | sp\|Q16836\|HCDH_HUMAN Hydroxyacyl-coenzyme A dehydrogenase, mitochondrial OS | 92,622 | 10,51 | 34294 | 2 | 20,34 | 20,11 | 23,02 | 16,78 | 19,89 | 21,58 | **0,299** | 3,61E-01 | **0** | -1,7400004 | 0,44272311 |
| SODM_HUM | SOD2 | sp\|P04179\|SODM_HUMAN Superoxide dismutase [Mn], mitochondrial OS=Homo sapier | 12,225 | 4,05 | 24722 | 1 | 21,06 | 21,48 | 22,71 | 20,34 | 20,97 | 18,70 | **0,298** | 1,05E-01 | **0** | -1,7453486 | 0,97676722 |
| EPDR1_HUM | EPDR1 | sp\|Q9UM22\|EPDR1_HUMAN Mammalian ependymin-related protein 1 OS=Homo sapier | 33,847 | 10,27 | 25437 | 2 | 18,58 | 24,61 | 23,89 | 19,35 | 18,91 | 23,45 | **0,289** | 4,96E-01 | **0** | -1,7891668 | 0,30494826 |
| RRAGC_HUN | RRAGC | sp\|Q9HB90\|RRAGC_HUMAN Ras-related GTP-binding protein C OS=Homo sapiens GN=F | 90,07 | 19,55 | 44224 | 4 | 22,45 | 22,61 | 26,76 | 22,06 | 22,12 | 22,19 | **0,284** | 2,67E-01 | **0** | -1,8166669 | 0,57277904 |
| EF1D_HUMA | EEF1D | sp\|P29692\|EF1D_HUMAN Elongation factor 1-delta OS=Homo sapiens GN=EEF1D PE=1 S | 53,093 | 13,52 | 31122 | 4 | 27,91 | 28,07 | 27,94 | 25,57 | 27,80 | 24,95 | **0,274** | 9,76E-02 | **0** | -1,8666668 | 1,01060554 |
| EPN3_HUMA | EPN3 | sp\|Q9H201\|EPN3_HUMAN Epsin-3 OS=Homo sapiens GN=EPN3 PE=2 SV=1 | 105,2 | 3,48 | 68222 | 2 | 22,32 | 21,92 | 23,05 | 17,95 | 21,57 | 21,90 | **0,258** | 2,09E-01 | **0** | -1,9566663 | 0,67996847 |
| CLH2_HUMA | CLTCL1 | sp\|P53675\|CLH2_HUMAN Clathrin heavy chain 2 OS=Homo sapiens GN=CLTCL1 PE=1 SV | 1471,648 | 12,13 | 187030 | 30 | 21,84 | 21,43 | 23,91 | 17,87 | 20,54 | 22,76 | **0,249** | 2,81E-01 | **0** | -2,0036418 | 0,55138452 |
| SCMC1_HUN | SLC25A24 | sp\|Q6NUK1\|SCMC1_HUMAN Calcium-binding mitochondrial carrier protein SCaMC-1 O: | 58,022 | 9,22 | 53354 | 4 | 21,67 | 21,08 | 24,41 | 18,49 | 20,11 | 22,28 | **0,234** | 2,36E-01 | **0** | -2,0933329 | 0,6270626 |
| SFXN1_HUM | SFXN1 | sp\|Q9H9B4\|SFXN1_HUMAN Sideroflexin-1 OS=Homo sapiens GN=SFXN1 PE=1 SV=4 | 152,357 | 21,74 | 35619 | 4 | 22,06 | 22,55 | 23,48 | 21,12 | 18,08 | 22,44 | **0,225** | 1,88E-01 | **0** | -2,1506049 | 0,72546681 |
| IFT27_HUM/ | IFT27 | sp\|Q9BW83\|IFT27_HUMAN Intraflagellar transport protein 27 homolog OS=Homo sapie | 11,513 | 5,91 | 20480 | 1 | 21,26 | 20,98 | 22,54 | 17,31 | 19,28 | 21,63 | **0,219** | 1,77E-01 | **0** | -2,1881708 | 0,75103791 |
| ATP5H_HUM | ATP5H | sp\|O75947\|ATP5H_HUMAN ATP synthase subunit d, mitochondrial OS=Homo sapiens G | 16,034 | 5,59 | 18491 | 1 | 21,95 | 21,67 | 22,48 | 20,11 | 20,97 | 18,22 | **0,208** | 5,52E-02 | **0** | -2,2663377 | 1,2576992 |
| BRD9_HUM/ | BRD9 | sp\|Q9H8M2\|BRD9_HUMAN Bromodomain-containing protein 9 OS=Homo sapiens GN=I | 9,926 | 1,17 | 67000 | 1 | 19,22 | 23,56 | 25,33 | 18,77 | 17,59 | 24,66 | **0,195** | 4,53E-01 | **0** | -2,3608818 | 0,3440641 |
| DJC11_HUM. | DNAJC11 | sp\|Q9NVH1\|DJC11_HUMAN DnaJ homolog subfamily C member 11 OS=Homo sapiens G | 39,503 | 6,26 | 63278 | 2 | 20,10 | 21,15 | 22,47 | 17,97 | 20,42 | 18,02 | **0,185** | 8,30E-02 | **0** | -2,437912 | 1,0806697 |
| KI16B_HUM/ | KIF16B | sp\|Q96L93\|KI16B_HUMAN Kinesin-like protein KIF16B OS=Homo sapiens GN=KIF16B PE | 21,862 | 1,52 | 152011 | 2 | 18,92 | 22,29 | 23,09 | 17,34 | 17,60 | 21,97 | **0,182** | 2,80E-01 | **0** | -2,4611511 | 0,55305829 |
| DPYL4_HUM. | DPYSL4 | sp\|O14531\|DPYL4_HUMAN Dihydropyrimidinase-related protein 4 OS=Homo sapiens GI | 20,816 | 2,8 | 61878 | 1 | 22,75 | 23,11 | 19,58 | 19,11 | 18,88 | 19,58 | **0,162** | 8,25E-02 | **0** | -2,6257833 | 1,08373341 |
| VRK1_HUMA | VRK1 | sp\|Q99986\|VRK1_HUMAN Serine/threonine-protein kinase VRK1 OS=Homo sapiens GN | 23,272 | 3,54 | 45476 | 1 | 22,09 | 22,50 | 21,92 | 17,91 | 18,56 | 22,10 | **0,160** | 1,14E-01 | **0** | -2,6460705 | 0,94246986 |
| CO5A1_HUN | COL5A1 | sp\|P20908\|CO5A1_HUMAN Collagen alpha-1(V) chain OS=Homo sapiens GN=COL5A1 PE | 54,35 | 1,58 | 183560 | 1 | 21,94 | 20,36 | 20,13 | 15,34 | 19,26 | 19,13 | **0,134** | 1,08E-01 | **0** | -2,8993667 | 0,96557486 |
